# Combined analysis of somatic mutations and gene expression reveals nuclear speckles-associated enhanced stemness in gingivobuccal carcinoma under DNA damage response

**DOI:** 10.1101/2024.11.16.623931

**Authors:** Sachendra Kumar, Tamasa De, Janavi Subramani, Annapoorni Rangarajan, Debnath Pal

## Abstract

Smokeless tobacco chewing habits in India lead to a high prevalence of Gingivobuccal oral squamous cell carcinoma (OSCC-GB). Cancer stem cells (CSCs) are a sub-population of cancer cells within a tumor with stem-like properties and are believed to contribute to tumor initiation, progression, increased resistance to drug therapy, and promote post-therapeutic cancer relapse. An RNA-seq data-based integrative analysis of somatic mutation and gene expression was performed to explore the role of CSCs in disease progression using the novel Indian-origin OSCC-GB cell line ‘IIOC019’ from a patient with tobacco-chewing habit. The identified DNA damage-related known mutational signature (ID1, SBS5) indicates the impact of smokeless tobacco-related carcinogens in the IIOC019 cell line. The differentially expressed somatic variants, functional impact predictions, and survival analysis reveal the role of DNA damage response (DDR)-related genes in OSCC-GB, with the SON gene as a significant player. The study suggests that the loss of function in a somatic variant of the SON gene is associated with enhanced stemness and increased risk of disease progression in OSCC-GB under DDR conditions. The newly identified CSC-associated somatic variants appear to promote cancer spread, local recurrence, and resistance to chemotherapy or radiotherapy, contributing to the high mortality rates among Indian OSCC-GB patients.

## 1 INTRODUCTION

Oral Squamous Cell Carcinoma (OSCC) is a prominent and distinct subtype of Head and Neck Squamous Cell Carcinoma (HNSCC). It primarily affects the oral gingivobuccal region comprising of buccal mucosa, gingivobuccal sulcus, alveolus and retromolar triangle, and is known as Gingivobuccal Oral Squamous Cell Carcinoma (OSCC-GB). Due to its predominance in the Indian subcontinent, it is also aptly described as the ‘Indian Oral Cancer’ ^1, 2^. Smokeless tobacco chewing habit, such as khaini, gutka, and betel quid with slaked lime, is the major cause of this disease in India ^3, 4^. Additional risk factors include cigarette smoking, alcohol use, and human papillomavirus (HPV) infection ^5^. Patients with OSCC-GB often face high rates of locoregional recurrence and poor 5-year survival rate (20%). This is linked to late-stage (III and IV) detection and limited success with conventional treatments involving surgery, adjuvant radiation, and chemotherapy ^4, 6, 7^.

Cancer Stem Cells (CSCs) are characterized by their stemness imparting self-renewal property (unlimited division) to maintain their population in an undifferentiated stem-like state and their ability to generate diverse cell types within a cancer (multipotent nature) - properties similar to normal stem cells. CSCs may arise through the accumulation of DNA-damaging mutations in long-lived tissue resident adult stem cells (inherent stemness) or from non-stem cancer cells (induced stemness). It may involve a progressive loss of cell-of-origin memory and acquiring stem cell-like features by hijacking developmental and/or oncogenic pathways influenced by the tumor microenvironment (TME), including immune and stromal cells. CSCs exhibit dynamic plasticity, which leads to identity change between CSC and non-CSC states and vice-versa, leading to intra-tumour heterogeneity that predisposes patients to poor clinical outcomes ^8, 9^.

CSC phenotypes have been studied using anchorage-independent growth, sphere formation assay, cancer stemness-related marker identification, Aldehyde dehydrogenase (ALDH) activity assay, and side population assay ^10–14^. Several studies characterized CD44 glycoproteins as oral cancer stem cell markers showing CSC phenotypes such as cancer metastasis, locoregional relapse, and chemo and/or radio-resistance ^12, 15^. The orosphere assay showed a higher abundance of CSC markers (CD44- positive cells) compared to regular cell culture, reflecting that orospheres can maintain stem-like properties; anchorage-independent orospheres thus serve as a valuable model for investigating CSCs in HNSCC ^11^. A CD44-based differential gene expression analysis on Indian OSCC-GB primary tumors from patients with a smokeless tobacco chewing habit revealed decreased expression of genes involved in cell adhesion ^12^. Analyzing the same RNA-seq dataset we performed differential somatic gene mutation analysis, and our work revealed the presence of DNA damage response (DDR) related genes-mediated stemness phenotype. The possible DNA damage and repair response involves a higher prevalence of C>T mutations and 1 bp T/(A) nucleotide insertions, resulting in genomic instability and indicating the role of smokeless tobacco chewing-related carcinogens in OSCC-GB ^16^. The International Cancer Genome Consortium’s (ICGC) Oral Cancer India project has identified frequently mutated genes in various tumorigenic pathways (such as the TP53 signaling pathway) and specific mutational signatures (C>T, C>A, C>G) in patients with OSCC-GB exposed to tobacco smoking or chewing ^17^. This differential regulation is likely enhanced by the involvement of nuclear speckles that store RNA and proteins central to complex cellular processes, including transcription, splicing, and RNA modification, crucial for gene expression ^18, 19^. Nuclear speckles association leads to enhanced TP53-mediated target gene expression, which plays a central role in the DDR ^20^. Understanding such insights promises to improve the OSCC-GB diagnosis, prognosis, and targeted therapies involving CSCs.

Patient-derived cancer cell lines can be excellent models due to the presence of homogeneous cell populations, ease of culture, and genetic stability. The previous reports show that CSCs get enriched upon growing/culturing the cells in a three-dimensional (3D) environment (non-adherent condition), the orosphere assay ^11^. The early passage cell culture could mimic the patient tumor microenvironment, while late passage culture is used for high throughput studies due to its stability (phenotypic and genotypic) and associated reproducibility ^21, 22^. Thus far, very few studies have documented OSCC-GB cell lines particularly originating from Indian patients, mainly due to the challenges in culturing primary cells derived from these locally advanced tumor ^10, 23, 24^. Our lab has recently established a novel Indian OSCC-GB patient-derived cancer cell line, IIOC019 (**I**ndian **I**nstitute of Science **O**ral **C**ancer **019**) which harbors cancer stem cells ^25^. Previously, Indian-origin OSCC-GB cell lines have indicated the presence of cancer stem-like cells through orosphere assays. Nevertheless, a comprehensive genomic characterization needs to be improved ^10, 24^. Therefore, IIOC019 can be valuable for CSC-associated studies involving aetiologies that may be prevalent in the Indian population.

Our current study focuses on the integrative analysis of somatic mutations or variants and gene expression of orosphere (stem cell enriched) as compared to the adherent cells using the novel IIOC019 OSCC-GB cell line derived from an Indian patient with long-term smokeless tobacco chewing habit. We use our previous transcriptomic study ^16^ as a reference for somatic mutational analysis. Further, the DESeq2 pipeline analysis of gene expression ^26^, and Seurat pipeline ^27^ analysis of single-cell RNA sequencing (scRNA-seq) reads from public repositories are used to understand the role of CSCs in oral cancer. The analysis reveals differentially expressed somatic variants as candidate genes based on the mutational signature, functional impact, and survival analysis in the subgroups of OSCC-GB cells. The study suggests that the loss of function in the candidate gene somatic variants is associated with enhanced stemness and a higher likelihood of disease progression in OSCC-GB under DDR conditions.

## 2 MATERIALS AND METHODS

### 2.1 Data sources

The FASTQ files obtained from the RNA-sequencing of a newly established OSCC-GB female patient-derived cell line namely, IIOC019 ^25^ were used for integrative analysis of somatic mutations and gene expression. The patient had smokeless tobacco exposure for more than 30 years with no such records of cigarette smoking or alcohol consumption. The primary cell culture is described in detail in Section S1 ^25^. In brief, the primary oral cancer cells derived from the OSCC-GB patient were grown in regular cell culture followed by trypsinization to obtain a single-cell suspension. Further, these single cells were grown in ultra-low attachment conditions in serum-free media with specific growth factors for 7 days to obtain orospheres using the established method for orosphere assay ^11^. For the current study, IIOC019 cells were cultured in both adherent (Adh) and orosphere (Oro) conditions. Additionally, two different passages, early passage (P8) and late passage (P24) with three technical triplicates (N=3) for each condition (Adh and Oro) were used to generate the RNA-seq data (**Figure S1)**. Briefly, RNA isolation and quality checks were done for all the samples, and then they were sent to an outsourced company (Next Generation Sequencing facility, Centre for Cellular and Molecular Platforms) that utilized the standard manufacturer’s protocol for RNA sequencing based on paired-end chemistry (**Table S1**). Also, a commercially available oral cancer cell line, FaDu (hypopharyngeal origin), was grown in Adh and Oro conditions in three technical triplicates (N=3) and sent for RNA-sequencing. The somatic mutations and gene expression analysis were performed for Oro as compared to Adh by combining passages (P8+P24; N=6) and at individual passage levels (P8, P24; N=3 each) of IIOC019 and FaDu (N=3) cell line subgroups. To create the panel of normals (PoN), we downloaded Indian normal control OSCC-GB raw RNA sequencing BAM files (N=30) after controlled-data access agreements with ICGC. The European Genome-phenome Archive (EGA) dataset accession codes used in this study are EGAD00001004430 and EGAD00001003981. All studies cited the approval of the respective institutional ethics committee in their study.

### 2.2 RNA sequencing data analysis

We have used our established pipeline for RNA sequencing-based somatic mutation analysis. The brief details are mentioned in the supplementary section **(Section S1-S5)**, and a detailed version with commands for each step is mentioned in our previously published paper ^16^. Briefly, ICGC-acquired BAM files were converted into FASTQ files using the Genome Analysis Toolkit (GATK) v4.1.6.0 (van der Auwera et al. 2013). IIOC019, FaDu, and ICGC FASTQ files containing paired-end reads were rigorously checked for base quality, adapter contamination, and other quality parameters using FastQC v0.11.8 (http://www.bioinformatics.babraham.ac.uk/projects/fastqc). We employed Trimmomatic v0.39 to filter out low-quality reads and trim the adapter sequences **(Table S2)** ^28^. The quality-checked FASTQ files were aligned to the human genome, utilizing Gencode GRCh38.p12 and its corresponding gene annotation file ^29^. This genome alignment was done using STAR (Spliced Transcripts Alignment to a Reference) v2.7.1a aligner ^30^. Before somatic variant calling, the BAM files underwent preprocessing following the GATK best practices ^31^. The PoN was created using ICGC RNA-seq data from normal controls adjacent to the tumor of Indian OSCC-GB patient tissues using GATK preprocessed BAM files followed by variant calling using the Mutect2 pipeline ^31, 32^. This PoN filter accounts for variants originating from ‘normal contamination’ or RNA sequencing artefacts in the tumor **(Section S2-S3)**.

### 2.3 Somatic mutation analysis

Somatic variants were identified using GATK-preprocessed BAM files with the Mutect2 pipeline (GATK v4.1.6.0) in tumor-only mode, focusing on genome intervals (chr 1-22, XY) following GATK best practices (https://gatk.broadinstitute.org/hc/en-us/articles/360035531132). Selecting potential somatic variants required filtering steps, including checking against PoN variants, gnomAD germline variants, known RNA editing sites from the RADAR database ^33^, and other quality controls ^16^. Somatic variants were annotated using Ensembl Variant Effect Predictor (VEP v97) ^34^ with the human GRCh38.p12 gene assembly, Single Nucleotide Polymorphism Database (dbSNP) build 151 ^35^ and the Catalogue of Somatic Mutations in Cancer variants (COSMIC v88) ^36^. Predictions of missense mutation’s functional impact involving amino acid substitutions were generated through Polyphen2 v2.2.2 ^37^. Following annotation, the VCF files underwent a rigorous filtering procedure, excluding variants with minor allele frequencies, dbSNP variants using BCFtools v1.9, and filtering out non-coding variants. SIFT Indel algorithm (https://sift.bii.a-star.edu.sg/www/SIFT_indels2.html) was utilized to predict their functional impact, focusing on prioritizing frameshift variants ^38^. VCF v4.2 files containing exonic or coding sequence variants were converted to Mutation Annotation Format (MAF) using the vcf2maf v1.6.18 tool (https://github.com/mskcc/vcf2maf). The MAF files from IIOC019 Adh (N=6), IIOC019 Oro cohorts (N=6), FaDu Adh (N=3), and FaDu Oro (N=3) cohorts were used for identifying novel and COSMIC variants, and those present in at least three samples were further used for downstream analysis. This analysis was conducted using Bioconductor (https://www.bioconductor.org/) R programming v4.0.5 (https://www.r-project.org/about.html), leveraging the Maftools v2.6.05 R package ^39^. The differential mutational analysis compared IIOC019 Oro and IIOC019 Adh samples cohorts by combining early (P8) and late passage (P24) and individually applying Fisher’s exact test to all genes with Maftools-mafCompare **(Section S4-S6)**.

### 2.7 Mutational signature analysis

Mutational signatures were identified for genes mutated in three or more samples for each IIOC019 Oro (N=6, including P8 and P24) and IIOC019 Adh (N=6, including P8 and P24). This was done by using the SignatureAnalyzer v0.0.7 Python tool, which employs nonnegative matrix factorization (NMF) with default settings on MAF files. Assignments of mutational signatures to samples were based on known COSMIC signatures: cosmic3_exome for single base substitutions (SBS) and cosmic3_ID for small insertions and deletions (indels). This assignment was considered when the cosine similarity score was ≥0.75 ^40^.

### 2.8. Expression and survival analysis from TCGA data

We conducted a gene expression analysis using the t-test on our genes of interest. This analysis utilized batch-normalized (log2) RNA-Seq data processed with the Expectation Maximization (RSEM) method on Illumina Hiseq mRNA gene expression data obtained from the HNSCC TCGA Firehouse Legacy cohort, which was downloaded from cBioPortal (https://www.cbioportal.org; 06 Jan 2024). The boxplot was created using ggplot2 within the ggpubr v0.4.0 package in RStudio R v4.0.2. Additionally, Kaplan–Meier plots were generated to investigate the correlation between mRNA gene expression and survival outcomes in the HNSCC TCGA Firehose Legacy cohort for our genes of interest (data from cBioPortal; 06 Jan 2024) ^41^.

### 2.9 Differential gene expression analysis

The read counts were extracted from BAM files using the Subread (featureCounts) v2.0.1 package **(Table S3)** ^42^. The differential gene expression (DGE) analysis comparison of IIOC019 Oro (N=6, including P8 and P24) and IIOC019 Adh (N=6, including P8 and P24; reference) cohort was performed using DESeq2 v1.30.1 package in R v4.0.5 ^26^. The genes with at least a sum of 5 reads across all samples were used for downstream analysis. The statistical significance cutoff was set at false discovery rate (FDR)-adjusted p-value<0.05 and log2-fold change (log2FC) values of 0.06. The differentially expressed genes were annotated for gene symbol and Entrez ID using org.Hs.eg.db v3.12.0 and AnnotationDbi v1.52.0 Bioconductor package (doi:10.18129/B9.bioc.AnnotationDbi). The gene ontology was performed using GO.db v3.12.1 and GOstats v2.56.0 Bioconductor package (hyperGTest, p-value cutoff=0.01). KEGG Pathways were analyzed using GAGE v2.40.2 (p-value cutoff=0.05) ^43^ and visualized using Pathview package v1.30.1 ^44^. Similarly, the DGE analysis comparison was performed at each passage level for IIOC019 Oro (N=3, including P8 or P24) and IIOC019 Adh (N=3, including P8 or P24; reference) cohort.

### 2.10 Single cell RNA sequencing analysis

The scRNA-seq processed HNSCC primary tumor data were downloaded from the Gene Expression Omnibus database (https://www.ncbi.nlm.nih.gov/geo/) under accession codes GSE188737 (4 patient samples) ^45^ and GSE164690 (8 samples from 4 patients) ^46^. The GSE188737 samples Seurat object converts to raw counts using DropletUtils-demultiplex_convert_to_10x R package. Both the sample sources were combined and reanalyzed to understand CD44-mediated cancer stemness **(Table S4)**. Both HNSCC oral cavity scRNA-seq samples used for analysis were HPV-negative, higher grade (T3 or T4), chemo-naive, sequenced using droplet-based 10X genomics technology, and aligned to GRCh38 reference genome. The scRNA-seq data analysis was performed using the Seurat v4.3.0.1 R package ^27^. The samples were filtered for genes expressing in less than 3 cells and 200 genes. Cells with greater than 20% mitochondrial reads and feature counts over 6000 or less than 200 (nFeature_RNA) were excluded as quality control. The filtered data was further normalized (NormalizeData), followed by feature selection (FindVariableFeatures) using default parameters. Further, the scaling of the data was performed with all genes and variation due to ribosomal reads regress out (ScaleData) using default parameters. The principal Component Analysis (PCA) was used to reduce the dimensionality of the data, and the first 15 principal components were selected based on the Elbow plot. The batch correction was performed to integrate the sample from two studies using Harmony v0.1.1 (RunHarmony) ^47^. The neighbors were found with harmony-corrected cell embedding (dims = 1:15) using the shared nearest neighbor (SNN) method (FindNeighbors). The clustering was performed with a resolution of 0.07 using the Louvain algorithm (FindClusters), and this was visualized using Uniform Manifold Approximation and Projection (UMAP) plots. The cell classes were automatically annotated using singleR v1.4.1 R package ^48^ using HumanPrimaryCellAtlasData reference from the celldex v1.0.0 R package ^48^. The ‘Tissue_stem_cells’ and ‘Epithelial_cells’ were used to find the differential gene expression between CD44-positive (WhichCells, expression=CD44>0) and CD44-negative using FindMarkers with min.pct 0.1 and logfc.threshold=0.25 employing the Wilcoxon Rank Sum test (default).

## 3 RESULTS

### 3.1. Somatic mutational landscape of novel IIOC019 cell line grown as Orospheres (stem-like) and adherent (differentiated) conditions

A somatic mutational analysis based on RNA-sequencing data was performed by combining early passage (P8; N=3) and late passage (P24, N=3) for respective conditions, IIOC019 Oro (N=6) and IIOC019 Adh (N=6) to get a comprehensive understanding of gene mutations that are passed onto the subsequent generation of the cell line (**Figure S1)**. This composite data may capture heterogeneity and disease progression. The somatic mutational profile unveils a total of 1442 somatic variants, comprising 1109 frameshift insertions, 4 frameshift deletions, 239 missense mutations, and 38 nonsense mutations, excluding 52 silent mutations (**Figure 1a**). The IIOC019 Oro (N=6) somatic mutational profile unveils 643 somatic variants across 142 genes, comprising 471 frameshift insertions, 2 frameshift deletions, 123 missense mutations, and 14 nonsense mutations, excluding 33 silent mutations. On the contrary, the IIOC019 Adh mutation profile detected 799 somatic variants within 176 genes, encompassing 638 frameshift insertions, 2 frameshift deletions, 116 missense mutations, and 24 nonsense mutations, excluding 19 silent mutations. There were no inframe deletions in the IIOC019 cohorts. It was also noticed that 61 and 95 uniquely mutated genes were present in IIOC019 Oro and IIOC019 Adh cohorts, respectively; besides, 81 mutated genes were shared between both subgroups (**Figure 1a)**. The median variants per sample were 97.5 and 136 in IIOC019 Oro and IIOC019 Adh, respectively **(Figure 1b)**. Also, 471 Insertion (INS) variant types and 76 C>T Single Nucleotide Variant (SNV) were observed in IIOC019 Oro and 638 INS variant types and 74 C>T SNV in IIOC019 Adh as dominant variants **(Figure 1c-d)**.

**Figure 1.**
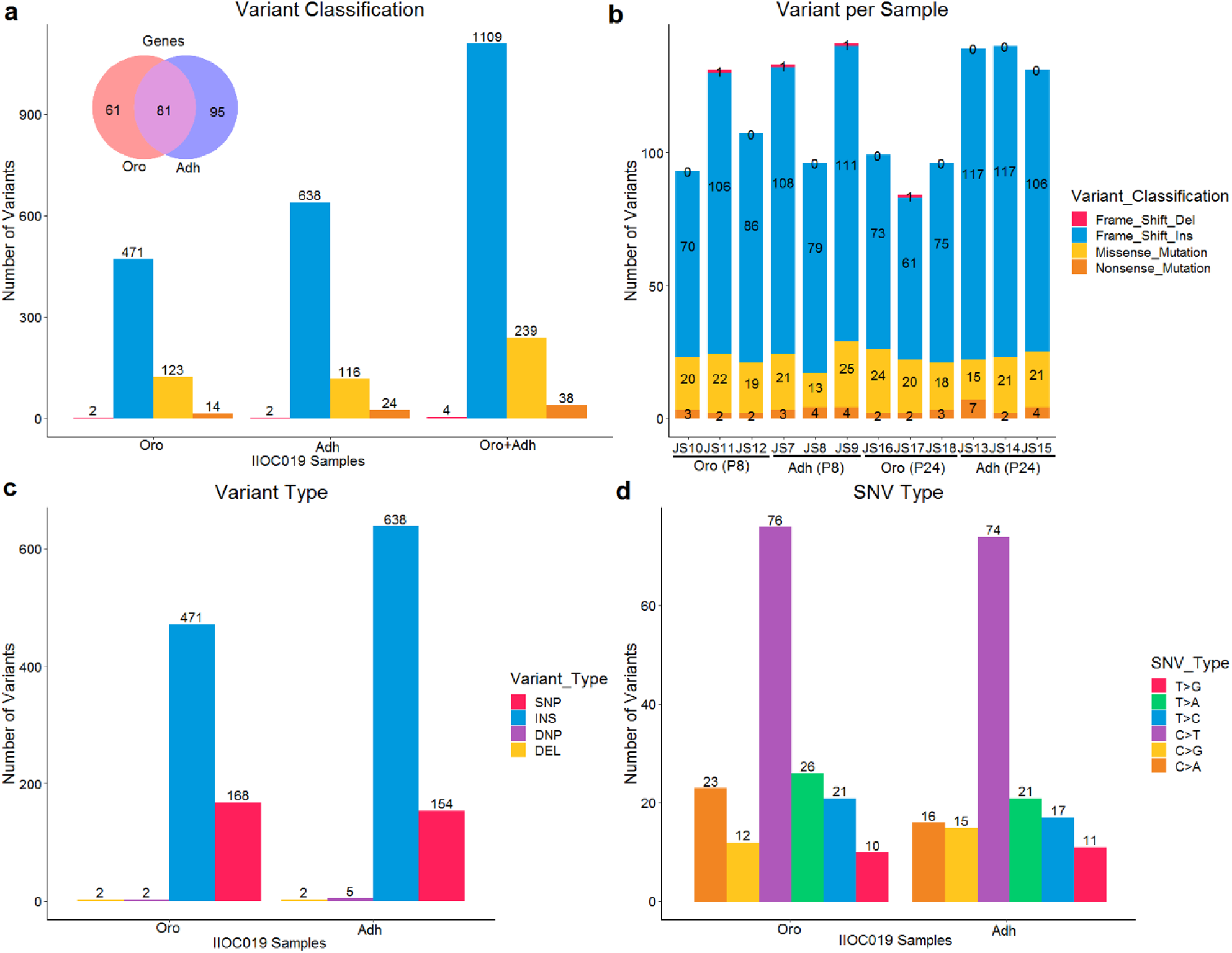
RNA-seq based somatic variant profile for Oro (N=6, including P8 and P24) and its matched Adh (N=6, including P8 and P24) of novel IIOC019 cell line using Maftools v2.6.05 R package. **a** Grouped bar plot for variant classification indicating higher frameshift insertions than missense mutations, frameshift deletions, and nonsense mutations annotated using VEP and Venn diagram showing common mutated genes. **b** Stacked bar plot showing accumulative variants frequency per sample (JS10, JS11, JS12) in IIOC019 Oro, its corresponding (JS7, JS8, JS9) in IIOC019 Adh for early passage (P8); (JS16, JS17, JS18) in IIOC019 Oro and its corresponding (JS13, JS14, JS15) in IIOC019 Adh for late passage (P24), here, the names of the samples within subgroups do not correspond to the order of the samples. **c** Grouped bar plot depicting higher insertions (INS) than single-nucleotide polymorphism (SNP), di-nucleotide polymorphism (DNP), and deletion (DEL) in IIOC019 subgroups. **d** Grouped bar plot for indicating a more prominent number of C>T SNV than other SNV types in IIOC019 subgroups.

### 3.2. Somatic mutational signature investigation indicates DDR-related etiology

Somatic mutational signatures analyzed in genes mutated in three or more samples for each IIOC019 Oro (N=6, including P8 and P24) and IIOC019 Adh (N=6, including P8 and P24) cohort suggest an etiology related to DNA damage. The most prominent COSMIC mutational signature (cosine similarity enrichment score≥0.75) identified in both IIOC019 Oro and IIOC019 Adh was DNA mismatch repair deficiency (ID1) **(Figure 2a-b)**, defective homologous recombination-based DNA damage repair (ID6), and DNA double-strand breaks-based DNA damage repair (ID8) (**Figure S2a-d)** due to small insertion and deletion (ID). Specifically, T/(A) nucleotide (1 base pair) insertion in the repetitive context. Also, the single base substitution (SBS), specifically C>T mutation, showed tobacco-associated carcinogens (SBS5) mutational signatures in both subgroups of the IIOC019 cell line **(Figure 2c-d, S2e-h)**.

**Figure 2.**
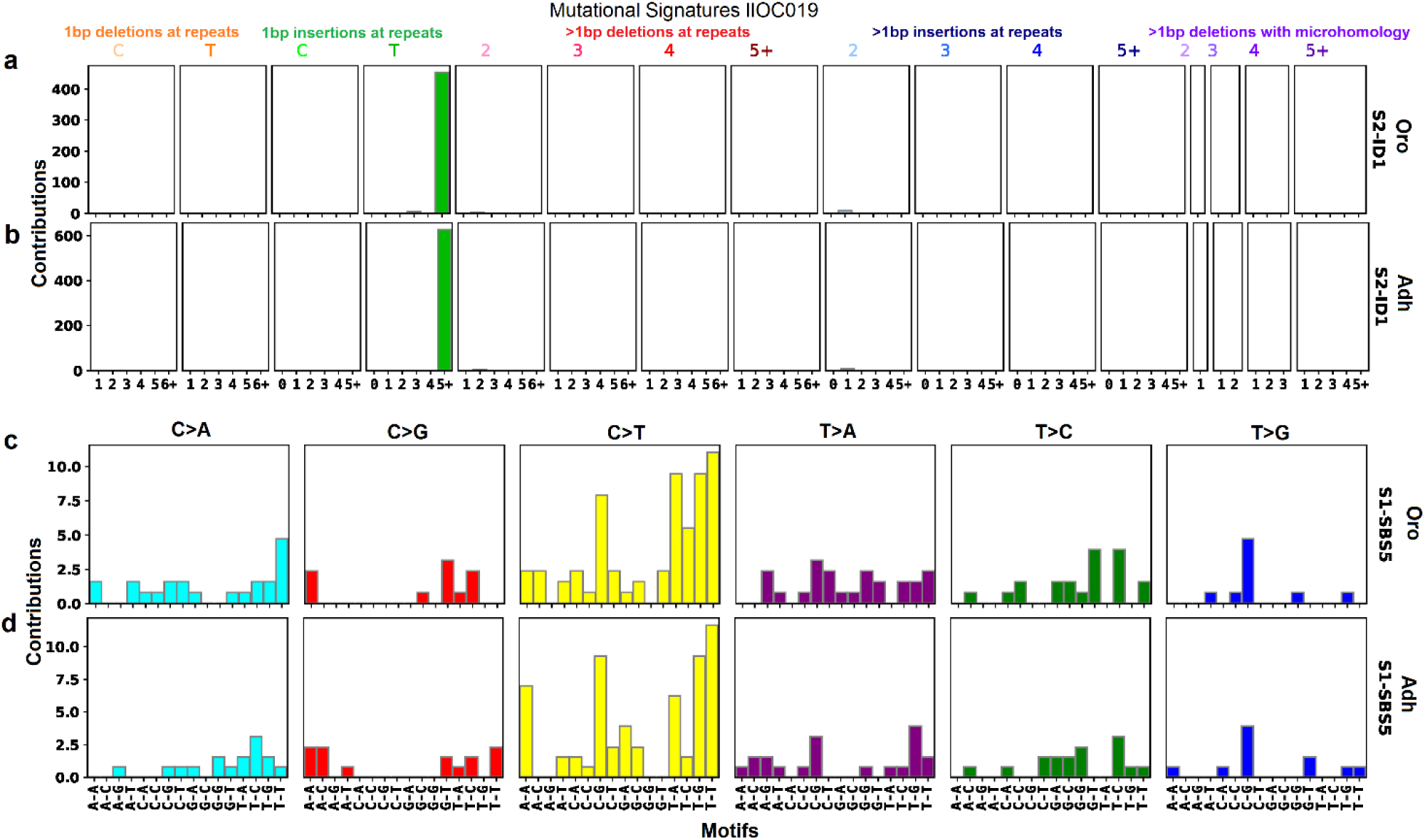
Mutational signature profile of IIOC019 Oro (N=6, including P8 and P24) and IIOC019 Adh cohort (N=6, including P8 and P24) using SignatureAnalyzer v0.0.7 python tool. **a-b** DNA mismatch repair deficiency (COSMIC ID1) characterized by prominent 1-base pair T/(A) nucleotide insertion. **c-d** Tobacco-associated carcinogens (SBS5) show the C>T single base substitution prevalence. Here, S1 exhibits the prevailing signature pattern among the corresponding COSMIC mutational signatures.

### 3.3 Distinctly mutated and functionally damaging DDR-related somatic variants in IIOC019 Oro and IIOC019 Adh cells

The comparative somatic mutational gene analysis revealed that 18 genes in IIOC019 Oro and 38 genes in IIOC019 Adh were distinctly present in at least 4/6 samples in respective subgroups (**Table S5**). Of these, 18 genes were distinctly mutated and statistically significant with a p-value<0.05, precisely 11 in IIOC019 Oro and 7 in IIOC019 Adh (**Table S5,** highlighted in bold). Further, the statistically significant mutated genes were selected in either five or six samples, with varying somatic variants for each gene. Subsequently, these somatic variants were prioritized, considering the functional impact prediction for missense mutations through PolyPhen2, and frameshift insertion mutations using SIFT Indel. SIFT Indel additionally supplied a report on nonsense-mediated decay (NMD) that leads to decreased gene expression, aiding in prioritizing the frameshift insertion somatic variants (**Table S6**).

In the IIOC019 Oro cohort, the identified NMD-associated frameshift insertion DDR somatic variants which were predicted to be damaging with a confidence score of 0.858 included SON DNA and RNA binding protein (SON, p.E2187Rfs*4), Heterogeneous nuclear ribonucleoprotein A3 (HNRNPA3, p.R167Efs*9), and Thyroid hormone receptor interactor 4 (TRIP4, p.G61Rfs*15) (**Figure 3a, S3a– b**). Similarly, IIOC019 Adh cohort, the identified NMD-associated frameshift insertion DNA damage-response somatic variants were predicted as damaging with a confidence score of 0.858 including Cell division cycle 23 (CDC23, p.L234Sfs*23) and Deltex E3 ubiquitin ligase 3L (DTX3L, p.S233Kfs*5) (**Figure S3c–d**).

**Figure. 3.**
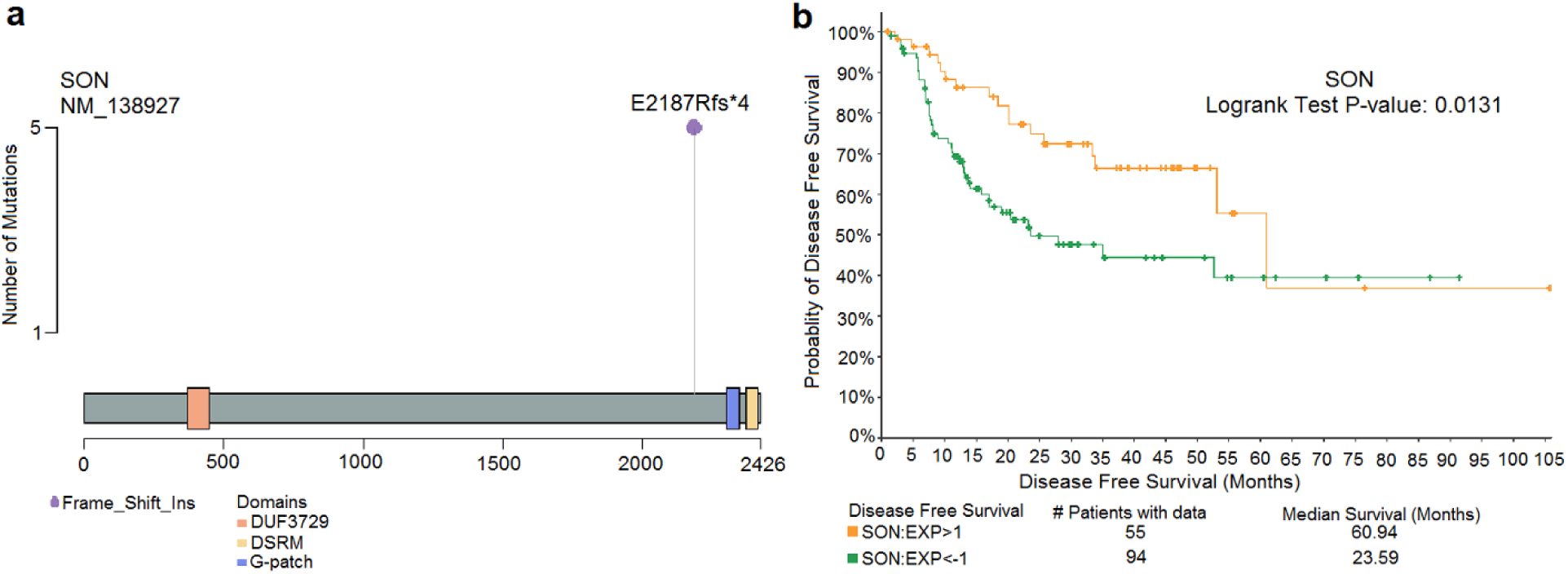
**a** LollipopPlot showing the position of SON protein frameshift insertion somatic variant (p.E2187Rfs*4/ p.Glu2187ArgfsTer4) before G-patch and Double-Stranded RNA-binding Motif domains (DSRM/DRBM) present in five out of the six IIOC019 Oro cohort (83.33% somatic mutation rate) using Maftools v2.6.05 R package. The prefix NM_ signifies the RefSeq transcript identifier and various domains on the corresponding protein structure are highlighted using distinct colors. **b** The Kaplan–Meier survival analysis correlated SON mRNA gene expression, revealing a reduced expression (EXP<-1, indicating the expression Z-score relative to diploid tumor samples, RNA Seq V2 RSEM) in the HNSCC TCGA Firehose Legacy cohort, which was linked to a poorer disease-free survival outcome with the statistically significant Logrank test p*-*value (0.0131) on 06 Jan 2024 calculated using the URL (https://www.cbioportal.org/).

Also, Kaplan–Meier survival analysis correlating mRNA gene expression was performed for SON, HNRNPA3 from IIOC019 Oro and CDC23, DTX3L from IIOC019 Adh using the HNSCC TCGA Firehose Legacy cohort in cBioPortal (**Figure 3b, S4a-c**). Decreased gene expression of SON linked to poor disease-free survival outcomes with a statistically significant Logrank test p-value (0.0131) (**Figure 3b**).

### 3.4. Comparative somatic mutational analysis of IIOC019 Oro and IIOC019 Adh cells based on early and late passages

A somatic analysis was performed for the different passages of IIOC019 subgroups to understand somatic mutational dynamics and associated heterogeneity due to the evolution of orosphere culture. The comparative somatic mutations of early passage (P8) IIOC019 Oro (N=3) and IIOC019 Adh (N=3) that are present in all 3 samples of the respective subgroups reveal a total of 58 mutated genes, precisely 29 in orosphere and 29 in adherent (**Table S7**). On the same line, there was a total of 63 mutated genes, precisely 19 in IIOC019 Oro (N=3) and 44 in IIOC019 Adh (N=3) in the late passage (P24) (**Table S8**). This data suggests that late passage (P24) may attain genomic homogeneity (stability), potentially resulting in decreased orosphere mutations than those from early passage (P8) orospheres. Also, we have compared our mutated gene results with previously established oral cancer cell line FaDu (hypopharyngeal origin). The comparative somatic mutational analysis of FaDu Oro (N=3) and FaDu Adh (N=3) that present in all 3 samples of the respective subgroups reveal a total of 66 mutated genes, precisely 34 in Oro and 32 in Adh (**Table S9**). The integrated results of early (P8) and late (P24) passage of the IIOC019 cell line reveal 14/96 commonly mutated unique genes combining all orosphere and adherent data concerning the FaDu cell line, indicating genomic similarities with the existing HNSCC cell line. Also, 12/96 commonly mutated unique genes combining CD44^+^Lin^−^ and CD44^−^Lin^−^ OSCC-GB primary tumor data concerning our previously published data indicated the relatedness of our novel IIOC019 cell line to OSCC-GB (**Figure S5**).

### 3.5. Differential gene expression analysis of IIOC019 Oro and IIOC019 Adh cells reveals DDR-related pathways

A DGE analysis was performed by combining early (P8) and late passage (P24) IIOC019 Oro (N=6) and IIOC019 Adh (N=6) cell line RNA-seq data to identify altered DDR-related pathways using the DESeq2 R package. The results revealed 4554 differentially and statistically significant genes (Log2fold 0.6 and padj<0.05), specifically 3134 up-regulated and 1420 down-regulated genes (**Figure S6a, Supplementary File2**). Similarly, the DGE of early passage (P8) IIOC019 Oro (N=3) and IIOC019 Adh (N=3) reveals 2438 differentially and statistically significant genes, specifically 1633 up-regulated and 805 down-regulated genes (**Supplementary File2**). Similarly, the DGE of late passage (P24) IIOC019 Oro (N=3) and IIOC019 Adh (N=3) reveals 5807 differentially and statistically significant genes, specifically 3513 up-regulated and 2294 down-regulated genes (**Supplementary File2**). The DGE were annotated by the KEGG pathway, showing 6 down-regulated statistically significant DDR pathways (p-value<0.05) in IIOC019 Oro (N=6) compared to IIOC019 Adh (N=6). Among the down-regulated DDR pathways, 6 and 1 overlapped with late passage (P24) and early passages (P8), respectively. The prominent down-regulated pathways were associated with DDR, such TP53 signaling pathway (**Figure S6b, S7**), Base excision repair, Nucleotide excision repair, Mismatch repair, Homologous recombination, and Cell cycle (**Figure S8-12, Table S10**). Also, the gene ontology analysis showed down-regulation of cellular activity of the nucleus in biological process (BP), cellular component (CC), and molecular function (MF) level among the top 10 identified differentially and statistically significant gene ontology (p-value<0.05) for IIOC019 Oro (N=6) versus IIOC019 Adh (N=6) (**Table S11-13**).

### 3.6. Single-cell transcriptional study to understand SON-mediated cancer stemness

As we have already seen in **Figure 3a**, the candidate SON gene somatic variant was distinctly mutated in IIOC019 Oro under DDR conditions. SON frameshift insertion somatic variants were predicted to be functionally damaging and NMD-associated, leading to decreased gene expression. A Kaplan– Meier survival analysis correlating CD44 (oral cancer stem cell marker) mRNA gene expression showed elevated expression associated with poor overall survival outcome with the statistically significant LogRank test p-value (5.701e-3) in the HNSCC TCGA Firehose Legacy cohort using cBioPortal (**Figure S13**). Single-cell sequencing data helps in identifying a rare subset of stem-like cells, specifically CD44-enriched cells, within primary tumor datasets. Therefore, a scRNA-seq analysis was performed to understand SON-mediated CD44-based cancer stemness regulations by re-analyzing the two published HNSCC (N=8) scRNA-seq data, having both immune (CD45-positive) and non-immune cells (CD45-negative) using Seurat R package. After the quality control analysis, 39313 cells were identified for further analysis (**Figure S14a-c**). The data were integrated using Harmony (**Figure 4a, S15a-b**), and cell types were annotated using the singleR package with HumanPrimaryCellAtlasData as a reference (**Figure S16-17**). The combined ‘Tissue stem cells’ and ‘Epithelial cells’ cell types were used to find the differential gene expression between CD44-positive and CD44-negative cells (**Figure 4b**). The results revealed 326 differentially and statistically significant genes (Log2fold 0.6 and padj<0.05), specifically 159 up-regulated and 167 down-regulated genes (**Supplementary File2**). The SON gene showed decreased expression in CD44- positive cells compared to CD44-negative cells (avg_log2FC change −0.447 and p_val_adj 4.336e-16 calculated using Seurat R package) (**Figure 4c**), indicating the SON NMD-associated frameshift insertion somatic variants impact cancer stemness.

**Figure 4.**
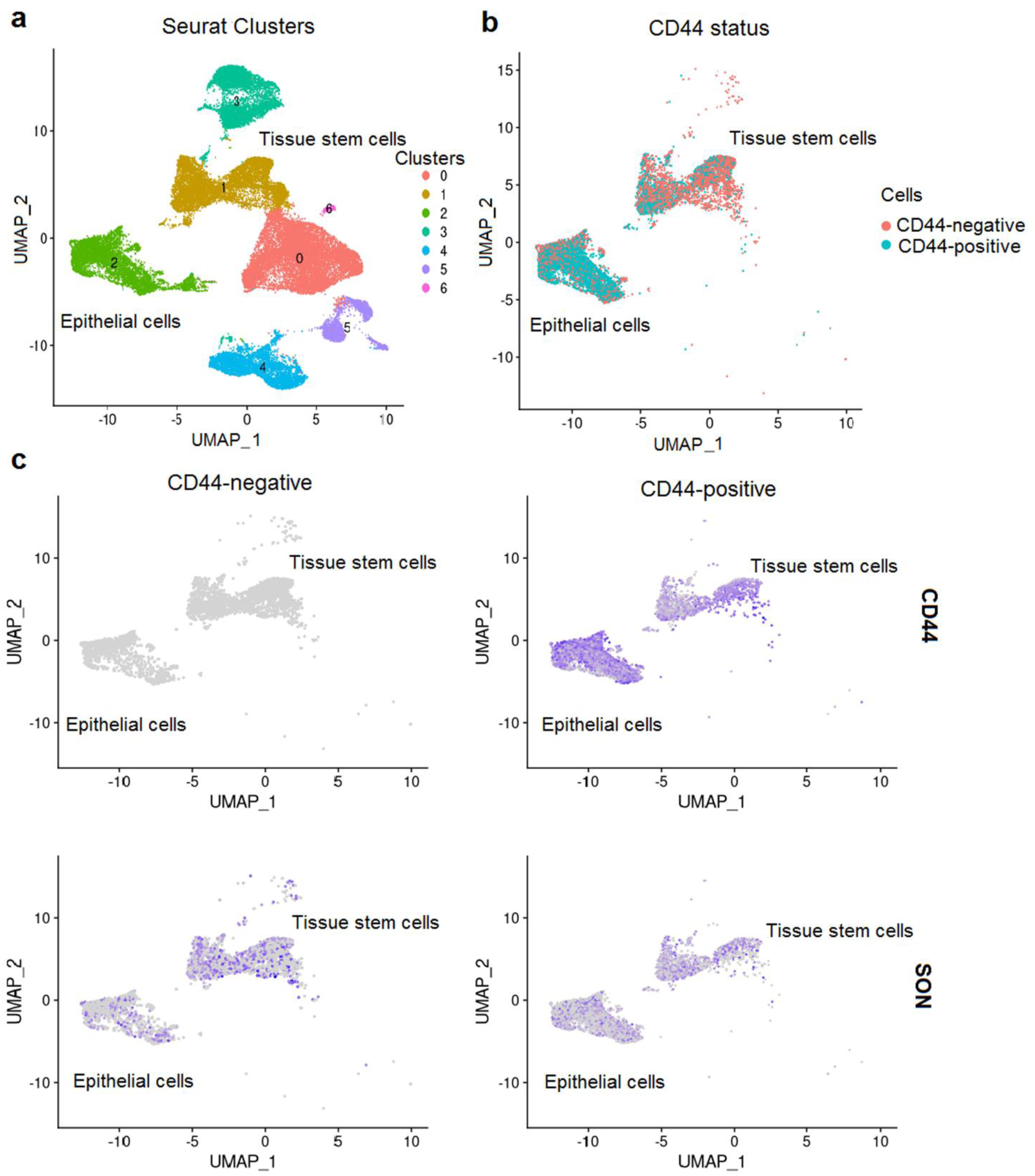
scRNA-seq analysis was performed to understand SON-mediated CD44-based cancer stemness regulations in HNSCC (N=8) using the Seurat v4.3.0.1 and Harmony v0.1.1 R package UMAP plot showing **a** Harmony corrected cell clusters (N=39313), highlighting Tissue stem cells (N=3714) and Epithelial cells (N=4874), **b** CD44-positive (WhichCells, expression=CD44>0) and CD44-negative cells in the Tissue stem cells and Epithelial cells, **c** FeaturePlot showing decreased SON expression in CD44-positive relative to CD44-negative cells in the selected cell types in UMAP dimension.

### 3.7. SON mutation impacts cancer stemness and patient survival via TP53

As we have previously seen in **Figure 3a**, SON (p.Glu2187ArgfsTer4) frameshift insertion somatic variant associated with NMD, caused by a 1-bp T nucleotide insertion, leads to a change in the affected amino acid from Glutamic acid to Arginine at position 2187. This alteration results in a new reading frame starting with asparagine at position 2187, causing a frameshift at 4 positions downstream until the end in 5 out of 6 samples IIOC019 Oro (N=6). This novel SON (p.Glu2187ArgfsTer4) frameshift insertion somatic variant is not reported in the COSMIC (v88; also the recent v99). SON (p.Glu2187ArgfsTer4) frameshift mutations lead to truncation of G-patch and Double-Stranded RNA-binding Motif (DSRM/DRBM) domains that are important for its mRNA-splicing function in nuclear speckles ^49^. SON frameshift insertion somatic variants were predicted to be NMD-associated, leading to decreased gene expression. As we have already seen in **Figure 3b**, a decreased SON gene expression showed poor disease-free survival outcomes in the HNSCC cohort. Previous studies have shown that the SON is essential for nuclear speckles and involved in regulating TP53-mediated enhanced expression of its downstream target (p21). The tumor suppressor TP53 gene plays a central role in the DDR ^50, 51^, and its PRD (proline-rich domain) is crucial in its association with nuclear speckles ^20^. We hypothesize that SON (p.Glu2187ArgfsTer4) frameshift mutation might disrupt the nuclear speckles and lead to loss of interaction with TP53. To test this hypothesis, we investigated TP53 gene expression levels in the IOC019 cells. As we have previously seen in **Figure S6b, S7**, DGE analysis showed down-regulation of TP53 gene expression in IIOC019 Oro (N=6) as compared to IIOC019 Adh (N=6) (Log2fold change −1.641 and padj. value 8.257e-08 calculated using DESeq2 R package) and overall, down-regulation of TP53 signaling KEGG pathway (p-value<0.05) for IIOC019 Oro using GAGE R package. In conclusion, NMD-associated SON (p.Glu2187ArgfsTer4) frameshift somatic mutation could potentially enhance stemness via TP53 in IIOC019 Oro in DDR conditions.

Additionally, this hypothesis was further supported by comparing the mean of RNA sequencing-based gene expression indicating lower TP53 expression in SON_Low tumors with a statistically significant p-value (0.00148) (**Figure 5a**) and CD44_High tumors with a statistically significant p-value (4.155236e-05) (**Figure 5b**) HNSCC TCGA Firehose Legacy cohort using t-test. As we have already seen in **Figure 4c**, SON-mediated cancer stemness was further validated using sc-RNAseq analysis. The analysis showed decreased expression of SON in CD44-positive cells as compared to CD44- negative cells. Also, TP53’s known downstream target CDKN1A/p21 showed lower expression in CD44-positive as compared to CD44-negative cells (avg_log2FC change −0.667 and p_val_adj 5.436e-11 calculated using Seurat R package) in HNSCC (**Figure S18**).

**Figure 5.**
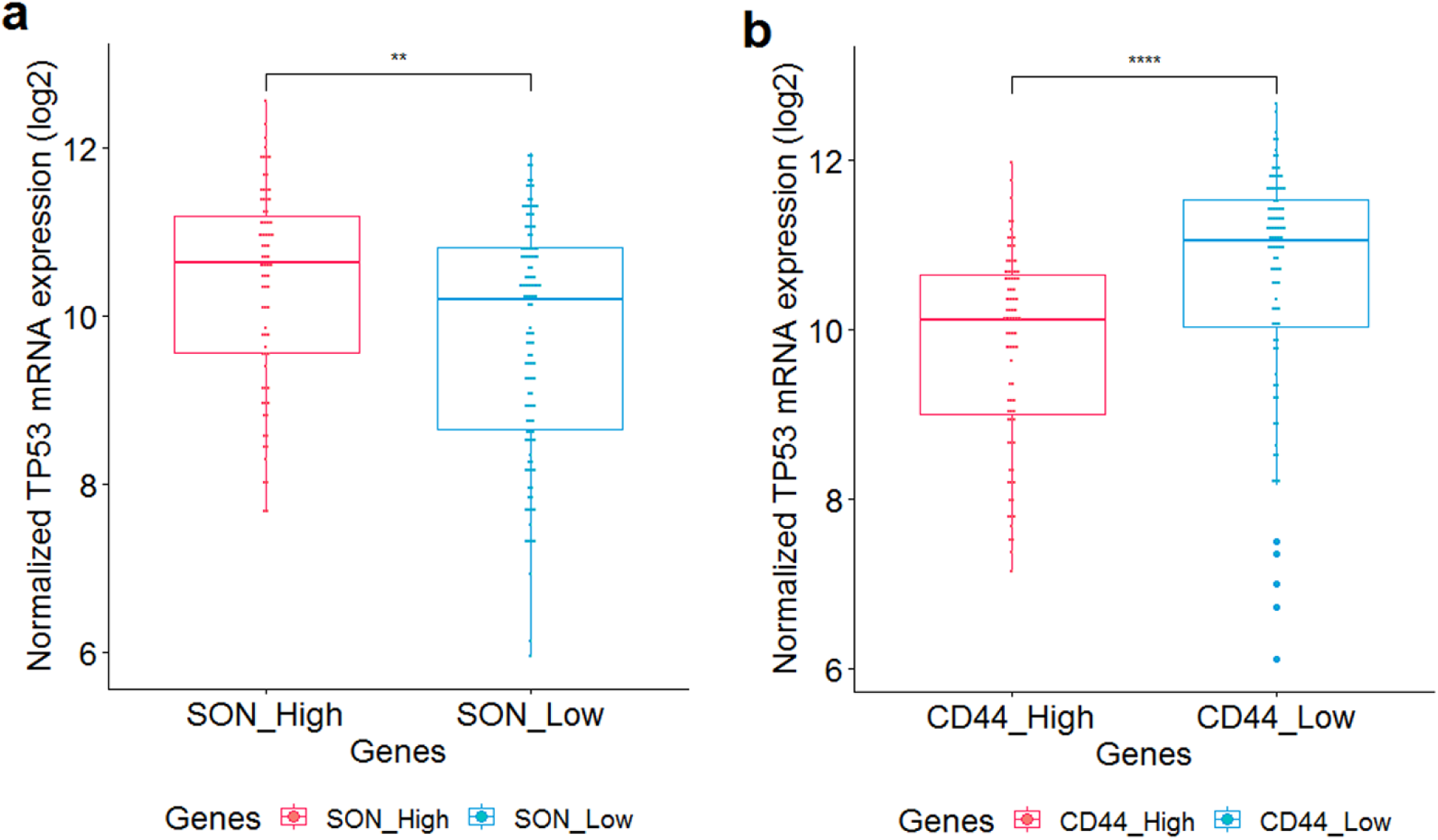
**a** Boxplot indicating the comparatively reduced expression of the TP53 gene within the SON_Low group (EXP<-1, N=118) in contrast to the SON_High group (EXP>1, N=77) with a statistically significant t-test p-value (0.00148) and **b** Boxplot showing relatively lower expression of TP53 gene in the CD44_High (EXP>1, N=86) compared to CD44_Low (EXP<-1, N=94) with a statistically significant t-test p-value (4.155236e-05) based on batch normalized (log2) RSEM Illumina Hiseq RNA sequencing mRNA gene expression in the HNSCC TCGA Firehose Legacy cohort via cBioPortal downloaded on 06 Jan 2024. The boxplot was plotted using ggplot2 based ggpubr v0.4.0 package in Rstudio R v4.0.2. The statistical significance is indicated as **p-value<0.01 and ****p-value<0.0001.

## DISCUSSION

An integrative analysis of somatic variants and gene expression was conducted to understand the role of orosphere-enriched CSCs in the novel IIOC019 OSCC-GB cell line. The capacity for self-renewal and multipotency exhibited by CSCs enables them to thrive and proliferate within the orosphere, in contrast to non-CSCs. DNA-damaging mutations accumulating within orospheres may induce the cancer stemness state, possibly through altering gene expression that creates a favorable TME. Comparing, a few identified somatically mutated genes in the novel IIOC019 cell line overlap with previously established the FaDu cell line (of hypopharyngeal origin) and our previously reported OSCC-GB primary tumor data, indicating genomic similarities of IIOC019 with HNSCC. The somatic mutational signature analysis identified our previously reported COSMIC signature T/(A) nucleotide (1 base pair) insertion (ID1) and related etiology (ID6, ID8) and C>T mutation (SBS5), indicating DDR due to tobacco carcinogens in both subgroups of the IIOC019 ^52^. Additionally, a COSMIC mutational signature for tobacco-associated carcinogens (ID3), defective DNA mismatch repair (ID7), DNA double-strand breaks by non-homologous DNA end-joining (ID8), and signatures with unknown importance (ID11, ID16) were identified in IIOC019 Adh, indicating intra-tumor heterogeneity within the subgroup of the IIOC019 cell line. Also, the intra-tumor heterogeneity analysis of distinctly mutated genes with significantly higher mutant-allele tumor heterogeneity (MATH) scores in IIOC019 Oro as compared to IIOC019 Adh using Maftools-inferHeterogeneity (data not shown), which is associated with poor survival outcomes ^53^. The somatic variant analysis revealed NMD-linked frameshift insertion somatic variants in the IIOC019 subgroups. NMD usually results in reduced gene expression, primarily due to loss-of-function (LoF), which is observed in various cancers ^54^ and is considered a crucial target for anticancer strategies ^55^.

The identified DDR-related mutated genes in IIOC019 Adh, such as CDC23, an E3 ubiquitin ligase play an important role in cell cycle regulation ^56^. CDC23 knockdown results in reduced invasion and proliferation of liver cancer ^57^. DTX3L E3 ligase knockout leads to p53 retention at poly ADP-ribose polymerase-associated DNA damage sites ^58^. DTX3L gene silencing results in reduced invasion and migration in glioma ^59^. The possible role of CDC23 (p.L234Sfs*23) and DTX3L (p.S233Kfs*5) NMD-associated LoF related to stemness requires more investigation in the context of DDR conditions in OSCC-GB.

The identified DDR-related mutated genes in IIOC019 Oro, such as TRIP4, a transcription coactivator, interact with EP300/CREBBP ^60^. TRIP4 (p.G61Rfs*15) LoF may increase CSC phenotype via our previously reported CREBBP-TP53-CD44 regulation in a DNA damage response context ^16^. HNRNPA3 plays an essential role in RNA transport and pre-mRNA splicing ^61^. HNRNPA3 (p.R167Efs*9) LoF might reduce its downstream target’s splicing, leading to an increased DDR. The possible role of TRIP4 (p.G61Rfs*15) and HNRNPA3 (p.R167Efs*9) NMD-associated LoF related to stemness requires more investigation in the context of DDR conditions in OSCC-GB. SON, a primary resident protein of nuclear speckles, acts as a regulatory hub for coordinating mRNA processing and transcription regulation that is essential for cell cycle progression ^19, 49, 62^. Our prior study showed increased CD44-enriched CSC phenotype in CD44^+^Lin^−^ as compared to CD44^-^Lin^-^ OSCC-GB primary tumors under DDR conditions, due to the loss of CREBBP acetyltransferase activity that reduced TP53 activity consequently, hindered its binding to the CD44 promoter ^16, 63, 64^. SON-associated nuclear speckles with TP53 could further enhance the regulation. This hypothesis is supported by experimental evidence showing SON knockdown leads to a reduced ability of TP53 to induce p21 (downstream target) at both RNA and protein levels in lung fibroblast IMR90 cells and requires TP53 proline-rich domain (PRD) ^20^. TP53-induced ZMAT3 splicing regulator was reported to inhibit CD44 variants in colorectal carcinoma, and ZMAT3 was observed to co-localize within nuclear speckles, suggesting the regulation of CD44 through nuclear speckles ^65^. SON (p.Glu2187ArgfsTer4) frameshift mutations lead to truncation of G-patch and DRBM domains that are important for their mRNA-splicing function in nuclear speckles. SON (p.Glu2187ArgfsTer4) frameshift mutation might disrupt the nuclear speckles, lead to loss of interaction with TP53 protein, and drive enhanced CD44-mediated CSC phenotype. Alternatively, the inefficient SON-induced TP53 mRNA splicing might decrease TP53 gene expressions, leading to lower protein expression and, consequently, weakening its interaction with nuclear speckles under DDR contextually. SON is known to enhance the mRNA splicing in the spliceosome complex involving nuclear speckles ^49, 62^. Our data also showed decreased TP53 gene expression and its overall down-regulation of the pathway in the IIOC019 Oro as compared to the IIOC019 Adh cell line. SON (nuclear speckles) and TP53 (DDR) gene expression were positively correlated, and SON exhibited decreased expression associated with poor disease-free survival outcomes, indicating disease recurrence after treatment HNSCC-TCGA data. Also, TP53 (DDR) and CD44 (stemness) gene expression were negatively correlated, and CD44 exhibited increased expression associated with poor overall survival outcomes, indicating disease progression in HNSCC TCGA data. Also, SON (nuclear speckles) and CD44 (stemness) gene expression were negatively correlated based on scRNA-seq analysis, indicating SON-mediated stemness in IIOC019 Oro through DDR. This regulation of nuclear speckles-associated enhanced stemness under DDR likely contributes to CSC dynamic plasticity within the TME, influencing locoregional recurrence and disease progression in OSCC-GB patients. In conclusion, NMD-associated SON (p.Glu2187ArgfsTer4) frameshift somatic mutation may contextually drive stemness and increase risk of poor disease-free survival outcomes in OSCC-GB patients via TP53 under DDR conditions, possibly induced by smokeless tobacco carcinogens (**Figure 6**).

**Figure. 6.**
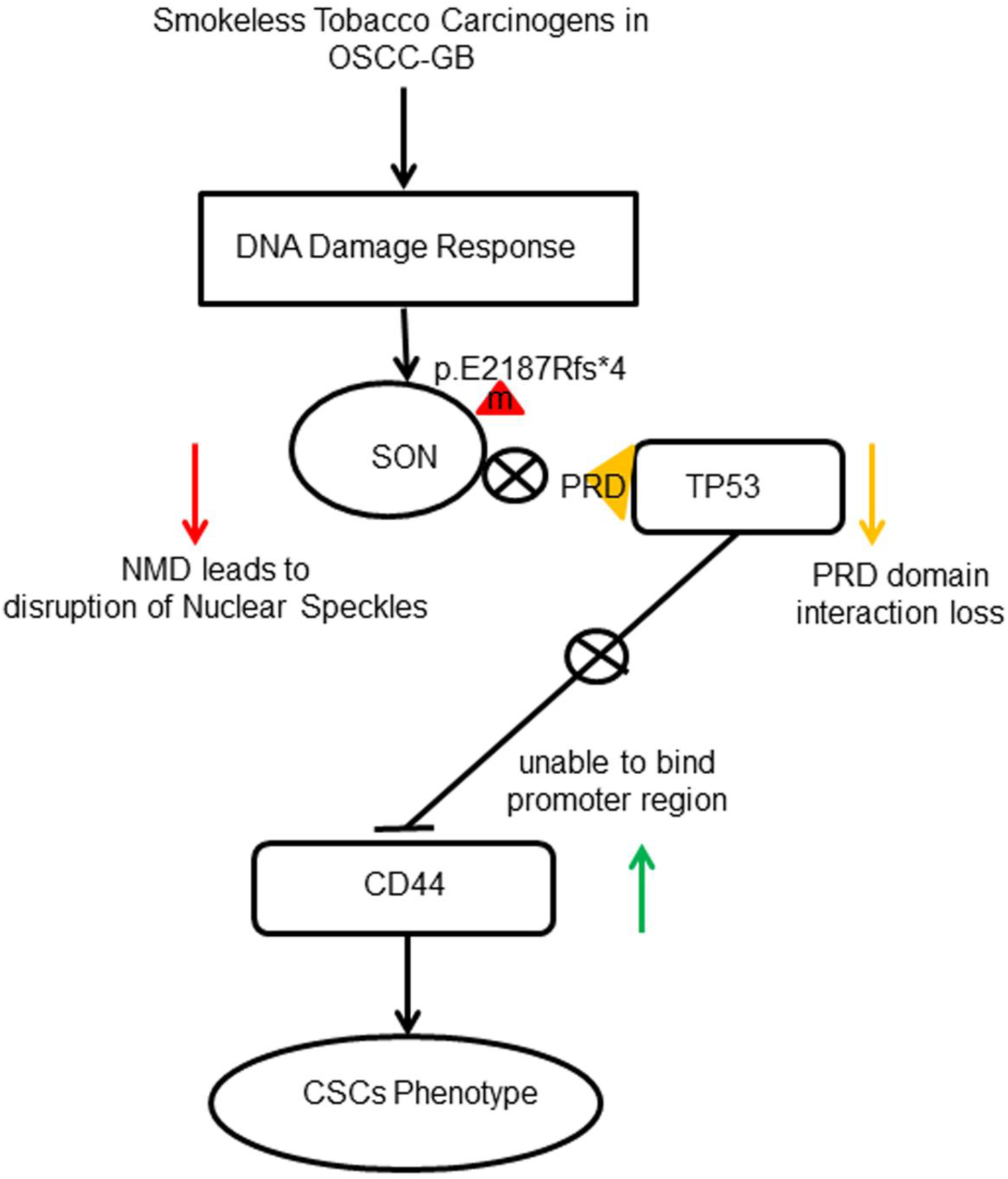
Schematic summary of smokeless tobacco carcinogen-induced CSC phenotype resulting from unrepaired DNA damage. Here, the loss of interactions between the PRD domain of TP53 with nuclear speckles SON (p.E2187Rfs*4) NMD-associated novel frameshift somatic variant drives reduced TP53 tumor suppressor expression. Since TP53 is known to bind to the promoter and inhibit CD44 expression, eventually, reduced TP53 could lead to enhanced CD44 gene expression, thus, promoting CSC phenotypes in IIOC019 Oro and other OSCC-GB patients under DDR context. The triangle with ‘m’ denoting mutation, ‘PRD’ represents the proline-rich domain, and the crossed circle denotes pathway repression.

## CONCLUSION

The integrative analysis of somatic variant and gene expression was performed for OSCC-GB patient-derived novel cell line ‘IIOC019’ of Indian origin. The study showed increased genome instability characterized by DDR-related mutational signature and TP53 signaling pathway down-regulation in IIOC019 Oro. The smokeless tobacco carcinogen-linked CD44-mediated CSC phenotypes were elucidated for SON, and its reduced expression correlated with an elevated risk of poor disease-free survival outcomes using the large HNSCC TCGA dataset. The identified SON (p.E2187Rfs*4) somatic variant possibly hinders interactions between the PRD domain of TP53 with SON nuclear speckles protein that may lead to enhanced expression of CD44-mediated CSCs phenotype in IIOC019 Oro under DDR conditions. The mechanism of structural association between SON and TP53 protein/gene requires further investigation. The anticancer therapeutic approach targeting the newly identified NMD-associated SON (p.E2187Rfs*4) frameshift insertion somatic variant could potentially enable the restoration of SON expression to reduce stemness and recurrence in OSCC-GB patients eventually.

## ACKNOWLEDGEMENTS

The authors would like to thank the Department of Biotechnology, New Delhi for providing funding support vide Sanction No: BT/PR17576/MED/30/1690/2016 under the project “Virtual National Oral Cancer Institute”. SK is thankful to the Department of Biotechnology, Government of India, for the Junior and Senior Research Fellowship award. The authors thank the ICGC for granting access to the ICGC-controlled data (ICGC Project #DACO-5868, “Studying genomic alterations of cancer-associated stem cells”). We thank the Next Generation Sequencing (NGS) facility at the Centre for Cellular and Molecular Platforms (CCAMP) for RNA-seq experiments.

## CONFLICT OF INTEREST

The authors confirm that there are no conflicts of interest.

## AUTHOR CONTRIBUTION

SK performed all computational data analysis and took the lead in manuscript writing. JS and TD contributed to cell line derivation and sample preparation for RNA sequencing and data collection. TD additionally contributed with scientific discussion and feedback for experiment design and data analyses, and critically reviewed the final manuscript. DP and AR conceived the original idea, supervised the project, designed experiments, reviewed the scientific content, and finalized the manuscript.

## AVAILABILITY OF DATA AND MATERIAL

The RNA-seq data utilized in this study will be available on request from the authors for research purposes. The processed data generated during this study are included in this article and its supplementary information.

## CODE AVAILABILITY

(software application or custom code) Not applicable.

## ETHICS APPROVAL

Ethical review/approval and written informed consent were not required for the study due to the computational nature of the work.

## CONSENT TO PARTICIPATE

Not applicable.

## CONSENT FOR PUBLICATION

Not applicable.

## Notes

### Competing Interest Statement

The authors have declared no competing interest.

