## Supplementary File 2 for "Combined analysis of somatic mutations and gene expression reveals nuclear speckles-associated enhanced stemness in gingivobuccal carcinoma under DNA damage response"

**The list of identified differentially and statistically significant genes (Log2fold 0.6 and padj<0.05; showing up- and down-regulated genes) for IIOC019 Oro as compared to IIOC019 Adh cohorts (N=6) RNA-seq using R package.**

| ensembl | baseMean | log2FoldChange | lfcSE | stat | pvalue | padj | ensembl | SYMBOL | ENTREZ |
|---|---|---|---|---|---|---|---|---|---|
| ENSG00000170373.8 | 3194.693 | 10.4498 | 0.4486037 | 21.95658 | 7.49E-107 | 3.27E-104 | ENSG00000170373 | CST1 | 1469 |
| ENSG00000151572.18 | 114.8281267 | 10.10899014 | 0.852263626 | 11.15733 | 6.59E-29 | 4.24E-27 | ENSG00000151572 | ANO4 | 121601 |
| ENSG00000167749.11 | 5443.794 | 10.10435 | 0.2141007 | 44.39195 | 0 | 0 | ENSG00000167749 | KLK4 | 9622 |
| ENSG00000242221.9 | 95.46373 | 10.08098 | 0.8506875 | 11.14508 | 7.57E-29 | 4.83E-27 | ENSG00000242221 | PSG2 | 5670 |
| ENSG00000180777.14 | 195.4123009 | 10.06650799 | 0.786178561 | 12.04117 | 2.16E-33 | 1.69E-31 | ENSG00000180777 | ANKRD30I | 374860 |
| ENSG00000101441.4 | 100.833 | 10.05214 | 0.8426028 | 11.21779 | 3.33E-29 | 2.20E-27 | ENSG00000101441 | CST4 | 1472 |
| ENSG00000221878.12 | 500.105 | 10.03228 | 0.5818742 | 16.21017 | 4.27E-59 | 7.96E-57 | ENSG00000221878 | PSG7 | 5676 |
| ENSG00000185966.3 | 15002.28922 | 9.820533499 | 0.381601133 | 24.16275 | 5.48E-129 | 3.17E-126 | ENSG00000185966 | LCE3E | 353145 |
| ENSG00000231672.7 | 119.968122 | 9.814735008 | 0.85686423 | 10.75402 | 5.67E-27 | 3.19E-25 | ENSG00000231672 | NA | NA |
| ENSG00000105205.7 | 111.259 | 9.742797 | 0.8558154 | 10.68314 | 1.22E-26 | 6.73E-25 | ENSG00000105205 | CLC | 1178 |
| ENSG00000250771.2 | 448.3958828 | 9.690469996 | 0.580535973 | 15.65875 | 2.90E-55 | 4.82E-53 | ENSG00000250771 | LOC10041 | 1E+08 |
| ENSG00000186844.5 | 2907.476579 | 9.495391383 | 0.249561625 | 35.64407 | 2.91E-278 | 5.54E-275 | ENSG00000186844 | LCE1A | 353131 |
| ENSG00000240386.3 | 1799.96331 | 9.49440693 | 0.363909051 | 24.44129 | 6.23E-132 | 3.68E-129 | ENSG00000240386 | LCE1F | 353137 |
| ENSG00000234756.1 | 65.98644504 | 9.419396939 | 0.888532712 | 9.925799 | 3.22E-23 | 1.47E-21 | ENSG00000234756 | LINC0262 | 1.05E+08 |
| ENSG00000164266.10 | 128.3428111 | 9.378624391 | 0.808327759 | 10.86023 | 1.78E-27 | 1.05E-25 | ENSG00000164266 | SPINK1 | 6690 |
| ENSG00000172155.9 | 273.9276838 | 9.363809856 | 0.602798623 | 14.53854 | 6.91E-48 | 9.15E-46 | ENSG00000172155 | LCE1D | 353134 |
| ENSG00000172551.11 | 1185.822127 | 9.208368404 | 0.315606153 | 27.27567 | 8.25E-164 | 6.27E-161 | ENSG00000172551 | MUCL1 | 118430 |
| ENSG00000285753.1 | 241.7869399 | 9.197222795 | 0.607435995 | 14.1533 | 1.78E-45 | 2.19E-43 | ENSG00000285753 | LCEP3 | 450211 |
| ENSG00000260454.2 | 315.4677296 | 9.175616849 | 0.572782074 | 14.97187 | 1.12E-50 | 1.61E-48 | ENSG00000260454 | NA | NA |
| ENSG00000108417.3 | 99.83081555 | 9.169173441 | 0.850611141 | 10.07414 | 7.19E-24 | 3.42E-22 | ENSG00000108417 | KRT37 | 8688 |
| ENSG00000172752.14 | 1478.954736 | 8.896570244 | 0.27239129 | 30.45828 | 9.31E-204 | 1.18E-200 | ENSG00000172752 | COL6A5 | 256076 |
| ENSG00000256955.2 | 483.1948926 | 8.762831596 | 0.393741532 | 20.73145 | 1.80E-95 | 7.16E-93 | ENSG00000256955 | NA | NA |
| ENSG00000149970.16 | 74.00014 | 8.67957 | 0.852613 | 9.476244 | 2.64E-21 | 1.08E-19 | ENSG00000149970 | CNKSR2 | 22866 |
| ENSG00000187180.3 | 488.2439754 | 8.654149381 | 0.418430116 | 19.24849 | 1.45E-82 | 4.55E-80 | ENSG00000187180 | LCE2C | 353140 |
| ENSG00000121898.13 | 76.60624274 | 8.621724637 | 0.817354702 | 9.814252 | 9.78E-23 | 4.36E-21 | ENSG00000121898 | CPXM2 | 119587 |
| ENSG00000186226.9 | 256.6002818 | 8.585193027 | 0.554678496 | 14.39607 | 5.48E-47 | 7.01E-45 | ENSG00000186226 | LCE1E | 353135 |
| ENSG00000102962.5 | 187.3140889 | 8.57292375 | 0.578109089 | 13.79138 | 2.87E-43 | 3.25E-41 | ENSG00000102962 | CCL22 | 6367 |
| ENSG00000167210.17 | 73.14736164 | 8.505509013 | 0.824823634 | 9.584484 | 9.29E-22 | 3.95E-20 | ENSG00000167210 | LOXHD1 | 125336 |
| ENSG00000244057.4 | 457.876204 | 8.498287729 | 0.388805105 | 20.31426 | 9.62E-92 | 3.46E-89 | ENSG00000244057 | LCE3C | 353144 |
| ENSG00000010438.16 | 69.8583853 | 8.468448003 | 0.805015656 | 9.774279 | 1.45E-22 | 6.37E-21 | ENSG00000010438 | PRSS3 | 5646 |
| ENSG00000187173.4 | 321.703507 | 8.459098094 | 0.464442822 | 16.92156 | 3.12E-64 | 6.65E-62 | ENSG00000187173 | LCE2A | 353139 |
| ENSG00000159455.8 | 384.4693635 | 8.431803874 | 0.400722349 | 19.54422 | 4.62E-85 | 1.50E-82 | ENSG00000159455 | LCE2B | 26239 |
| ENSG00000163202.5 | 45152.08673 | 8.430771882 | 0.164358689 | 47.6444 | 0 | 0 | ENSG00000163202 | LCE3D | 84648 |
| ENSG00000204363.4 | 108.8072 | 8.41495 | 0.7390582 | 10.5742 | 3.93E-26 | 2.11E-24 | ENSG00000204363 | SPANXN5 | 494197 |
| ENSG00000144285.20 | 106.73392 | 8.381599765 | 0.706666971 | 11.01169 | 3.36E-28 | 2.07E-26 | ENSG00000144285 | SCN1A | 6323 |

| | | | | | | | | | |
|---|---|---|---|---|---|---|---|---|---|
| ENSG00000287893.1 | 26.66728201 | 8.248750834 | 0.886657555 | 8.626499 | 6.33E-18 | 2.13E-16 | ENSG00000287893 | NA | NA |
| ENSG00000228013.1 | 1702.476747 | 8.245512115 | 0.268880218 | 28.43464 | 7.55E-178 | 6.93E-175 | ENSG00000228013 | IL6R-AS1 | 1.02E+08 |
| ENSG00000138670.17 | 2082.827943 | 8.241634906 | 0.220998079 | 34.57783 | 5.44E-262 | 9.05E-259 | ENSG00000138670 | RASGEF1I | 153020 |
| ENSG00000153012.12 | 302.7807596 | 8.241569393 | 0.460380729 | 16.59837 | 7.16E-62 | 1.46E-59 | ENSG00000153012 | LGI2 | 55203 |
| ENSG00000170369.4 | 94.19835 | 8.133236 | 0.7334268 | 10.27129 | 9.49E-25 | 4.72E-23 | ENSG00000170369 | CST2 | 1470 |
| ENSG00000187223.3 | 589.4266949 | 8.108786577 | 0.37574696 | 19.98363 | 7.65E-89 | 2.68E-86 | ENSG00000187223 | LCE2D | 353141 |
| ENSG00000084674.14 | 24.7273946 | 8.094502006 | 0.859792325 | 8.716642 | 2.87E-18 | 9.81E-17 | ENSG00000084674 | APOB | 338 |
| ENSG00000124089.4 | 23.53327 | 8.035892 | 0.8638337 | 8.608014 | 7.43E-18 | 2.49E-16 | ENSG00000124089 | MC3R | 4159 |
| ENSG00000278998.1 | 117.3274766 | 7.967081344 | 0.614635622 | 11.9861 | 4.20E-33 | 3.27E-31 | ENSG00000278998 | NA | NA |
| ENSG00000232679.2 | 171.9083709 | 7.95579186 | 0.530207112 | 13.87343 | 9.18E-44 | 1.06E-41 | ENSG00000232679 | NA | NA |
| ENSG00000182352.8 | 21.48748973 | 7.925266375 | 0.880511079 | 8.319335 | 8.85E-17 | 2.74E-15 | ENSG00000182352 | C17orf77 | 146723 |
| ENSG00000174038.13 | 82.51498604 | 7.894933193 | 0.667789963 | 10.92399 | 8.85E-28 | 5.26E-26 | ENSG00000174038 | C9orf131 | 138724 |
| ENSG00000110786.18 | 152.2051815 | 7.89252697 | 0.544598391 | 13.39065 | 6.86E-41 | 7.05E-39 | ENSG00000110786 | PTPN5 | 84867 |
| ENSG00000170848.15 | 137.154 | 7.742021 | 0.5545159 | 12.87974 | 5.85E-38 | 5.43E-36 | ENSG00000170848 | PSG6 | 5675 |
| ENSG00000087128.10 | 2062.058236 | 7.696432828 | 0.255092309 | 27.81908 | 2.55E-170 | 2.19E-167 | ENSG00000087128 | TMPRSS1 | 28983 |
| ENSG00000237250.3 | 45.75916511 | 7.693708732 | 0.755675183 | 9.387246 | 6.16E-21 | 2.51E-19 | ENSG00000237250 | LOC10013 | 1E+08 |
| ENSG00000235942.2 | 73.99665164 | 7.662017052 | 0.643901975 | 10.96753 | 5.47E-28 | 3.30E-26 | ENSG00000235942 | LCE6A | 448835 |
| ENSG00000169509.6 | 51673.74519 | 7.599505744 | 0.321051363 | 21.80183 | 2.23E-105 | 9.42E-103 | ENSG00000169509 | CRCT1 | 54544 |
| ENSG00000176136.6 | 47.59083321 | 7.565586784 | 0.723138676 | 9.632436 | 5.83E-22 | 2.50E-20 | ENSG00000176136 | MC5R | 4161 |
| ENSG00000144834.14 | 84.62033691 | 7.535748758 | 0.666748253 | 10.40235 | 2.42E-25 | 1.26E-23 | ENSG00000144834 | TAGLN3 | 29114 |
| ENSG00000286501.1 | 17.17041989 | 7.528972497 | 0.888213741 | 7.801019 | 6.14E-15 | 1.67E-13 | ENSG00000286501 | NA | NA |
| ENSG00000271573.1 | 141.5375721 | 7.523846046 | 0.529131525 | 13.0853 | 4.00E-39 | 3.85E-37 | ENSG00000271573 | NA | NA |
| ENSG00000183128.7 | 18.05338608 | 7.472663331 | 0.893438494 | 7.692374 | 1.44E-14 | 3.82E-13 | ENSG00000183128 | CALHM3 | 119395 |
| ENSG00000134853.12 | 458.1120314 | 7.407732429 | 0.388114044 | 17.54055 | 7.02E-69 | 1.68E-66 | ENSG00000134853 | PDGFRA | 5156 |
| ENSG00000110484.7 | 15.49548703 | 7.403162515 | 0.878281616 | 7.745992 | 9.48E-15 | 2.54E-13 | ENSG00000110484 | SCGB2A2 | 4250 |
| ENSG00000238042.5 | 21.1051673 | 7.384843931 | 0.871279179 | 7.787221 | 6.85E-15 | 1.86E-13 | ENSG00000238042 | LINC0225 | 1.05E+08 |
| ENSG00000243486.1 | 21.20450803 | 7.360818543 | 0.860482133 | 7.857012 | 3.93E-15 | 1.09E-13 | ENSG00000243486 | NA | NA |
| ENSG00000106565.18 | 41.62008029 | 7.345329837 | 0.764552301 | 8.822588 | 1.12E-18 | 3.94E-17 | ENSG00000106565 | TMEM176I | 28959 |
| ENSG00000183760.10 | 12686.67 | 7.325882 | 0.235865 | 28.51581 | 7.46E-179 | 7.09E-176 | ENSG00000183760 | ACP7 | 390928 |
| ENSG00000272021.1 | 30.49046799 | 7.315298566 | 0.859516061 | 7.812883 | 5.59E-15 | 1.53E-13 | ENSG00000272021 | NA | NA |
| ENSG00000253314.7 | 21.53538075 | 7.311440045 | 0.889522414 | 7.544993 | 4.52E-14 | 1.15E-12 | ENSG00000253314 | LINC0029 | 497634 |
| ENSG00000103316.11 | 14.01026779 | 7.30652491 | 0.879943313 | 7.621542 | 2.51E-14 | 6.51E-13 | ENSG00000103316 | CRYM | 1428 |
| ENSG00000150361.12 | 17.88419455 | 7.284706765 | 0.879251355 | 7.602726 | 2.90E-14 | 7.48E-13 | ENSG00000150361 | KLHL1 | 57626 |
| ENSG00000074317.11 | 90.65180722 | 7.271169737 | 0.543475728 | 12.27501 | 1.23E-34 | 9.95E-33 | ENSG00000074317 | SNCB | 6620 |
| ENSG00000249345.8 | 13.61428034 | 7.247573125 | 0.874903048 | 7.598068 | 3.01E-14 | 7.75E-13 | ENSG00000249345 | LINC0240 | 1.02E+08 |
| ENSG00000248444.1 | 14.42182121 | 7.239007277 | 0.875857763 | 7.580006 | 3.46E-14 | 8.87E-13 | ENSG00000248444 | NA | NA |
| ENSG00000159899.14 | 409.7605683 | 7.180065367 | 0.689665836 | 9.540947 | 1.42E-21 | 5.92E-20 | ENSG00000159899 | NPR2 | 4882 |
| ENSG00000203783.5 | 2100.383743 | 7.145552685 | 0.17576888 | 37.23954 | 1.56E-303 | 4.63E-300 | ENSG00000203783 | PRR9 | 574414 |
| ENSG00000241220.1 | 30.21992427 | 7.143397292 | 0.779170492 | 8.397902 | 4.55E-17 | 1.44E-15 | ENSG00000241220 | NA | NA |
| ENSG00000232624.8 | 12.59114683 | 7.134326603 | 0.890275373 | 7.339669 | 2.14E-13 | 5.21E-12 | ENSG00000232624 | C10orf126 | 283080 |
| ENSG00000128917.8 | 219.7443096 | 7.126483634 | 0.35827925 | 18.21619 | 3.84E-74 | 1.01E-71 | ENSG00000128917 | DLL4 | 54567 |
| ENSG00000203786.6 | 549.1051993 | 7.110486845 | 0.278682221 | 23.36169 | 1.05E-120 | 5.47E-118 | ENSG00000203786 | KPRP | 448834 |

| | | | | | | | | | |
|---|---|---|---|---|---|---|---|---|---|
| ENSG00000163395.16 | 2683.562346 | 7.073229447 | 0.147450998 | 43.90089 | 0 | 0 | ENSG00000163395 | IGFN1 | 91156 |
| ENSG00000105427.10 | 28154.68 | 7.024476 | 0.3233832 | 19.86645 | 7.94E-88 | 2.64E-85 | ENSG00000105427 | CNFN | 84518 |
| ENSG00000224769.1 | 75.40948799 | 7.012766329 | 0.574799213 | 11.15653 | 6.65E-29 | 4.26E-27 | ENSG00000224769 | NA | NA |
| ENSG00000287238.1 | 22.6302821 | 7.004452573 | 0.859538793 | 7.451034 | 9.26E-14 | 2.31E-12 | ENSG00000287238 | NA | NA |
| ENSG00000075673.11 | 2263.79102 | 6.996731862 | 0.348585684 | 18.35053 | 3.27E-75 | 8.88E-73 | ENSG00000075673 | ATP12A | 479 |
| ENSG00000243137.8 | 2186.577 | 6.957345 | 0.1405835 | 45.22113 | 0 | 0 | ENSG00000243137 | PSG4 | 5672 |
| ENSG00000181652.19 | 3569.83575 | 6.949229337 | 0.364794886 | 17.40493 | 7.57E-68 | 1.72E-65 | ENSG00000181652 | ATG9B | 285973 |
| ENSG00000229435.2 | 20.16763709 | 6.9283283 | 0.863026352 | 7.332717 | 2.26E-13 | 5.47E-12 | ENSG00000229435 | NA | NA |
| ENSG00000183018.9 | 5056.081111 | 6.875453191 | 0.138924372 | 45.17172 | 0 | 0 | ENSG00000183018 | SPNS2 | 124976 |
| ENSG00000224397.7 | 10.06365 | 6.807857 | 0.9155083 | 6.780776 | 1.20E-11 | 2.49E-10 | ENSG00000224397 | PELATON | 1.01E+08 |
| ENSG00000203688.6 | 23.38073923 | 6.802786647 | 0.838718098 | 7.395556 | 1.41E-13 | 3.47E-12 | ENSG00000203688 | LINC02487 | 441178 |
| ENSG00000158022.6 | 308.1659333 | 6.792045772 | 0.274968346 | 22.51912 | 2.70E-112 | 1.31E-109 | ENSG00000158022 | TRIM63 | 84676 |
| ENSG00000232310.8 | 10.06510585 | 6.789721206 | 0.932423939 | 6.638312 | 3.17E-11 | 6.34E-10 | ENSG00000232310 | LOC10192 | 1.02E+08 |
| ENSG00000235385.2 | 45.20966 | 6.774092 | 1.006512 | 6.134144 | 8.56E-10 | 1.48E-08 | ENSG00000235385 | LINC0215 | 1.1E+08 |
| ENSG00000253227.2 | 87.37885879 | 6.76484505 | 0.494816786 | 12.45884 | 1.25E-35 | 1.06E-33 | ENSG00000253227 | NA | NA |
| ENSG00000167914.11 | 6753.833466 | 6.740899768 | 0.222474655 | 27.60269 | 1.03E-167 | 8.09E-165 | ENSG00000167914 | GSDMA | 284110 |
| ENSG00000166473.17 | 113.0407505 | 6.737898404 | 0.496954831 | 12.35102 | 4.81E-35 | 3.95E-33 | ENSG00000166473 | PKD1L2 | 114780 |
| ENSG00000259828.1 | 303.9164159 | 6.678960202 | 0.621169491 | 9.786315 | 1.29E-22 | 5.68E-21 | ENSG00000259828 | NA | NA |
| ENSG00000196734.8 | 105.4522169 | 6.658582595 | 0.513568626 | 11.79703 | 4.04E-32 | 3.04E-30 | ENSG00000196734 | LCE1B | 353132 |
| ENSG00000189051.5 | 761.9751418 | 6.640663472 | 0.276515595 | 21.84565 | 8.55E-106 | 3.67E-103 | ENSG00000189051 | RNF222 | 643904 |
| ENSG00000258285.1 | 12.42614445 | 6.615958978 | 0.898363545 | 6.696575 | 2.13E-11 | 4.34E-10 | ENSG00000258285 | NA | NA |
| ENSG00000204764.14 | 8.623695548 | 6.614769921 | 0.906631547 | 6.634194 | 3.26E-11 | 6.49E-10 | ENSG00000204764 | RANBP17 | 64901 |
| ENSG00000138109.11 | 103.7293476 | 6.602389915 | 0.579624462 | 10.35565 | 3.94E-25 | 2.02E-23 | ENSG00000138109 | CYP2C9 | 1559 |
| ENSG00000173221.14 | 16195.48237 | 6.59453519 | 0.204996358 | 29.24215 | 5.65E-188 | 5.79E-185 | ENSG00000173221 | GLRX | 2745 |
| ENSG00000165091.17 | 166.2685287 | 6.563358619 | 0.367008662 | 16.24855 | 2.29E-59 | 4.35E-57 | ENSG00000165091 | TMC1 | 117531 |
| ENSG00000232023.3 | 22.37421041 | 6.549897402 | 0.782213043 | 7.606492 | 2.82E-14 | 7.28E-13 | ENSG00000232023 | LINC01807 | 1.02E+08 |
| ENSG00000182261.4 | 16.83236574 | 6.541184157 | 0.870227894 | 6.827159 | 8.66E-12 | 1.83E-10 | ENSG00000182261 | NLRP10 | 338322 |
| ENSG00000287534.1 | 120.2909979 | 6.516119658 | 0.401163493 | 14.7474 | 3.20E-49 | 4.41E-47 | ENSG00000287534 | NA | NA |
| ENSG00000287527.1 | 16.86917184 | 6.513922111 | 0.869385525 | 6.802416 | 1.03E-11 | 2.16E-10 | ENSG00000287527 | NA | NA |
| ENSG00000133661.16 | 60.55514602 | 6.502118578 | 0.574787486 | 10.26835 | 9.79E-25 | 4.85E-23 | ENSG00000133661 | SFTPD | 6441 |
| ENSG00000231811.3 | 65.74324725 | 6.473491355 | 0.615202174 | 9.547254 | 1.33E-21 | 5.59E-20 | ENSG00000231811 | LOC10192 | 1.02E+08 |
| ENSG00000124939.6 | 15.27223169 | 6.461014912 | 0.869439574 | 6.741141 | 1.57E-11 | 3.24E-10 | ENSG00000124939 | SCGB2A1 | 4246 |
| ENSG00000125355.15 | 48.98436 | 6.458291 | 0.5683206 | 10.30807 | 6.48E-25 | 3.25E-23 | ENSG00000125355 | TMEM255A | 55026 |
| ENSG00000116996.9 | 552.2930168 | 6.454504713 | 0.317611678 | 18.4329 | 7.15E-76 | 1.96E-73 | ENSG00000116996 | ZP4 | 57829 |
| ENSG00000224048.6 | 7.828572387 | 6.449203164 | 0.904286871 | 6.468305 | 9.91E-11 | 1.89E-09 | ENSG00000224048 | LINC02612 | 1.02E+08 |
| ENSG00000250038.7 | 529.9610915 | 6.416572933 | 0.482321898 | 12.05952 | 1.73E-33 | 1.36E-31 | ENSG00000250038 | NA | NA |
| ENSG00000128610.12 | 9.790720262 | 6.406633866 | 0.90959641 | 6.383748 | 1.73E-10 | 3.24E-09 | ENSG00000128610 | FEZF1 | 389549 |
| ENSG00000241104.5 | 21.68165 | 6.405237 | 0.7992094 | 7.263725 | 3.77E-13 | 8.92E-12 | ENSG00000241104 | LOC28434 | 284344 |
| ENSG00000255190.2 | 10.02340054 | 6.400224924 | 0.919311645 | 6.309313 | 2.80E-10 | 5.17E-09 | ENSG00000255190 | NA | NA |
| ENSG00000172061.8 | 28.72574568 | 6.384572883 | 0.697143712 | 8.297533 | 1.06E-16 | 3.28E-15 | ENSG00000172061 | LRRC15 | 131578 |
| ENSG00000227096.1 | 74.35158912 | 6.382315521 | 0.460122196 | 12.56691 | 3.21E-36 | 2.77E-34 | ENSG00000227096 | NA | NA |
| ENSG00000167754.13 | 22957.76 | 6.377459 | 0.4169756 | 13.85563 | 1.18E-43 | 1.36E-41 | ENSG00000167754 | KLK5 | 25818 |

| | | | | | | | | | |
|---|---|---|---|---|---|---|---|---|---|
| ENSG00000141255.12 | 7.830624852 | 6.376812877 | 0.921029801 | 6.272124 | 3.56E-10 | 6.49E-09 | ENSG00000141255 | SPATA22 | 84690 |
| ENSG00000103569.9 | 15.7126117 | 6.373080927 | 0.911233486 | 6.335457 | 2.37E-10 | 4.38E-09 | ENSG00000103569 | AQP9 | 366 |
| ENSG00000100427.15 | 10.66767 | 6.348224 | 0.9170455 | 6.268199 | 3.65E-10 | 6.64E-09 | ENSG00000100427 | MLC1 | 23209 |
| ENSG00000256292.2 | 10.35905749 | 6.344042075 | 1.078923963 | 5.323862 | 1.02E-07 | 1.40E-06 | ENSG00000256292 | NA | NA |
| ENSG00000100292.17 | 6878.7 | 6.341098 | 0.2405885 | 23.86273 | 7.47E-126 | 4.06E-123 | ENSG00000100292 | HMOX1 | 3162 |
| ENSG00000248187.1 | 7.426525471 | 6.340402135 | 0.923809157 | 6.21384 | 5.17E-10 | 9.21E-09 | ENSG00000248187 | NA | NA |
| ENSG00000227301.1 | 7.129140022 | 6.338082239 | 0.967861384 | 5.92862 | 3.05E-09 | 4.99E-08 | ENSG00000227301 | NA | NA |
| ENSG00000183668.18 | 142.2977 | 6.330305 | 0.3331695 | 17.19937 | 2.68E-66 | 6.01E-64 | ENSG00000183668 | PSG9 | 5678 |
| ENSG00000167723.14 | 6448.838649 | 6.327947219 | 0.154737669 | 37.01715 | 6.07E-300 | 1.62E-296 | ENSG00000167723 | TRPV3 | 162514 |
| ENSG00000225249.6 | 8.338404633 | 6.327460814 | 0.927471691 | 6.175348 | 6.60E-10 | 1.16E-08 | ENSG00000225249 | NA | NA |
| ENSG00000242641.6 | 7.049905974 | 6.326592871 | 0.915436506 | 6.255587 | 3.96E-10 | 7.15E-09 | ENSG00000242641 | LINC0097[illegible] | 440970 |
| ENSG00000188100.9 | 32.28475971 | 6.305957741 | 0.672876472 | 8.479948 | 2.25E-17 | 7.32E-16 | ENSG00000188100 | FAM25A | 643161 |
| ENSG00000238024.1 | 15.98935098 | 6.305037052 | 0.817550597 | 6.978207 | 2.99E-12 | 6.55E-11 | ENSG00000238024 | NA | NA |
| ENSG00000205076.4 | 1301.311 | 6.298485 | 0.2186921 | 26.05711 | 1.12E-149 | 7.26E-147 | ENSG00000205076 | LGALS7 | 3963 |
| ENSG00000105889.15 | 94.16969031 | 6.28656144 | 0.530542341 | 10.71839 | 8.34E-27 | 4.65E-25 | ENSG00000105889 | STEAP1B | 256227 |
| ENSG00000255556.2 | 9.762527978 | 6.285713339 | 0.908135894 | 6.260862 | 3.83E-10 | 6.93E-09 | ENSG00000255556 | NA | NA |
| ENSG00000073737.16 | 806.8923416 | 6.273598431 | 0.188739592 | 30.06046 | 1.59E-198 | 1.93E-195 | ENSG00000073737 | DHRS9 | 10170 |
| ENSG00000100253.13 | 9.557461 | 6.266906 | 0.9201976 | 6.158358 | 7.35E-10 | 1.29E-08 | ENSG00000100253 | MIOX | 55586 |
| ENSG00000250266.2 | 10.77732445 | 6.261534625 | 0.899129421 | 6.296685 | 3.04E-10 | 5.58E-09 | ENSG00000250266 | LINC0161[illegible] | 1.02E+08 |
| ENSG00000104808.8 | 725.5804 | 6.252024 | 0.5169788 | 10.9328 | 8.03E-28 | 4.78E-26 | ENSG00000104808 | DHDH | 27294 |
| ENSG00000197084.5 | 258.6015044 | 6.247372789 | 0.917650249 | 6.154167 | 7.55E-10 | 1.32E-08 | ENSG00000197084 | LCE1C | 353133 |
| ENSG00000182898.4 | 13.61378716 | 6.241223415 | 0.898642025 | 6.277498 | 3.44E-10 | 6.28E-09 | ENSG00000182898 | TCHHL1 | 126637 |
| ENSG00000160284.15 | 8.591262 | 6.237725 | 0.9686215 | 5.820358 | 5.87E-09 | 9.29E-08 | ENSG00000160284 | SPATC1L | 84221 |
| ENSG00000235481.3 | 244.994891 | 6.225413758 | 0.275087131 | 20.44957 | 6.06E-93 | 2.30E-90 | ENSG00000235481 | NA | NA |
| ENSG00000253141.2 | 14.95603554 | 6.22431973 | 0.846139728 | 6.647034 | 2.99E-11 | 6.00E-10 | ENSG00000253141 | NA | NA |
| ENSG00000170426.2 | 180.2289056 | 6.2148507 | 0.331920015 | 16.91628 | 3.41E-64 | 7.21E-62 | ENSG00000170426 | SDR9C7 | 121214 |
| ENSG00000225226.2 | 7.200979475 | 6.201969416 | 0.931277588 | 6.015359 | 1.79E-09 | 3.00E-08 | ENSG00000225226 | NA | NA |
| ENSG00000109684.15 | 6.483350527 | 6.197996808 | 0.931325104 | 6.010787 | 1.85E-09 | 3.08E-08 | ENSG00000109684 | CLNK | 116449 |
| ENSG00000260239.1 | 35.60365225 | 6.191069838 | 0.645387718 | 8.663118 | 4.59E-18 | 1.56E-16 | ENSG00000260239 | NA | NA |
| ENSG00000204967.11 | 6.662644797 | 6.177935986 | 0.948895816 | 5.878344 | 4.14E-09 | 6.66E-08 | ENSG00000204967 | PCDHA4 | 56144 |
| ENSG00000228264.1 | 6.546270035 | 6.177609086 | 0.940208811 | 5.932309 | 2.99E-09 | 4.89E-08 | ENSG00000228264 | PSMD8P1 | 1E+08 |
| ENSG00000227925.2 | 6.52195521 | 6.176203162 | 0.925235052 | 6.026796 | 1.67E-09 | 2.80E-08 | ENSG00000227925 | NA | NA |
| ENSG00000254038.1 | 9.337395311 | 6.175021987 | 0.912668701 | 6.108484 | 1.01E-09 | 1.73E-08 | ENSG00000254038 | NA | NA |
| ENSG00000174145.8 | 116.6336778 | 6.14480898 | 0.392141432 | 14.13982 | 2.16E-45 | 2.62E-43 | ENSG00000174145 | NWD2 | 57495 |
| ENSG00000140481.15 | 1051.361878 | 6.139834504 | 0.362833598 | 15.26825 | 1.24E-52 | 1.90E-50 | ENSG00000140481 | CCDC33 | 80125 |
| ENSG00000186207.5 | 42.72848177 | 6.123766626 | 0.772116552 | 7.154058 | 8.42E-13 | 1.94E-11 | ENSG00000186207 | LCE5A | 254910 |
| ENSG00000175315.3 | 15255.58951 | 6.117642525 | 0.211404971 | 26.09987 | 3.66E-150 | 2.43E-147 | ENSG00000175315 | CST6 | 1474 |
| ENSG00000185962.1 | 12.82134951 | 6.109124934 | 0.892346707 | 6.173749 | 6.67E-10 | 1.17E-08 | ENSG00000185962 | LCE3A | 353142 |
| ENSG00000229821.1 | 12.15151103 | 6.102974517 | 0.956688956 | 5.752104 | 8.81E-09 | 1.37E-07 | ENSG00000229821 | NA | NA |
| ENSG00000148677.6 | 65.8270137 | 6.089895322 | 0.444172948 | 12.35981 | 4.31E-35 | 3.55E-33 | ENSG00000148677 | ANKRD1 | 27063 |
| ENSG00000236601.2 | 5.953785838 | 6.083070869 | 0.963225401 | 5.692407 | 1.25E-08 | 1.91E-07 | ENSG00000236601 | NA | NA |
| ENSG00000161849.3 | 23.81334558 | 6.074549396 | 0.870328843 | 6.290208 | 3.17E-10 | 5.81E-09 | ENSG00000161849 | KRT84 | 3890 |

| | | | | | | | | | |
|---|---|---|---|---|---|---|---|---|---|
| ENSG00000253968.2 | 6.30573941 | 6.047065759 | 0.985681932 | 5.52619 | 3.27E-08 | 4.76E-07 | ENSG00000253968 | NA | NA |
| ENSG00000260947.1 | 12.46912821 | 6.046088437 | 0.868895257 | 6.267831 | 3.66E-10 | 6.64E-09 | ENSG00000260947 | NA | NA |
| ENSG00000286270.2 | 5.781013 | 6.042294 | 0.9684722 | 5.619464 | 1.92E-08 | 2.87E-07 | ENSG00000286270 | NA | NA |
| ENSG00000109846.8 | 39.73178204 | 6.041115326 | 0.662928001 | 8.207702 | 2.25E-16 | 6.76E-15 | ENSG00000109846 | CRYAB | 1410 |
| ENSG00000136943.11 | 25968.64589 | 6.038722081 | 0.169901759 | 32.01098 | 7.67E-225 | 1.07E-221 | ENSG00000136943 | CTSV | 1515 |
| ENSG00000276430.2 | 5.923063209 | 6.02079824 | 0.92446317 | 5.863725 | 4.53E-09 | 7.25E-08 | ENSG00000276430 | FAM25C | 644054 |
| ENSG00000203837.5 | 4282.945451 | 6.019219197 | 0.33850246 | 16.00939 | 1.10E-57 | 1.95E-55 | ENSG00000203837 | PNLIPRP3 | 119548 |
| ENSG00000109182.12 | 6.347883287 | 5.99805446 | 0.93272039 | 5.787431 | 7.15E-09 | 1.12E-07 | ENSG00000109182 | CWH43 | 80157 |
| ENSG00000110675.13 | 57.82124273 | 5.991603649 | 0.50315633 | 10.71556 | 8.60E-27 | 4.78E-25 | ENSG00000110675 | ELMOD1 | 55531 |
| ENSG00000287063.1 | 5.60191171 | 5.986710073 | 0.987403579 | 5.455429 | 4.89E-08 | 6.93E-07 | ENSG00000287063 | NA | NA |
| ENSG00000179593.16 | 16.95988259 | 5.977379896 | 0.789751489 | 6.808952 | 9.83E-12 | 2.07E-10 | ENSG00000179593 | ALOX15B | 247 |
| ENSG00000273877.4 | 5.88811 | 5.974355 | 0.970449 | 5.538009 | 3.06E-08 | 4.48E-07 | ENSG00000273877 | LOC10537 | 1.05E+08 |
| ENSG00000228917.1 | 29.98560174 | 5.944165907 | 0.622364001 | 8.586881 | 8.94E-18 | 2.98E-16 | ENSG00000228917 | NA | NA |
| ENSG00000232352.1 | 5.71214297 | 5.940886588 | 0.95236679 | 5.608014 | 2.05E-08 | 3.06E-07 | ENSG00000232352 | SEMA3B-A | 1.02E+08 |
| ENSG00000179057.13 | 52.66079424 | 5.926697431 | 0.512286927 | 10.39788 | 2.54E-25 | 1.32E-23 | ENSG00000179057 | IGSF22 | 283284 |
| ENSG00000253484.1 | 6.760670356 | 5.923163504 | 0.954716389 | 5.575649 | 2.47E-08 | 3.65E-07 | ENSG00000253484 | NA | NA |
| ENSG00000215512.9 | 6.965136823 | 5.917857035 | 0.944792295 | 5.628599 | 1.82E-08 | 2.73E-07 | ENSG00000215512 | LOC64466 | 644669 |
| ENSG00000242837.1 | 5.735581036 | 5.898795691 | 1.040462884 | 5.092729 | 3.53E-07 | 4.52E-06 | ENSG00000242837 | RPL21P13 | 319144 |
| ENSG00000189058.9 | 20.0261099 | 5.894642108 | 0.851576624 | 6.217458 | 5.05E-10 | 9.02E-09 | ENSG00000189058 | APOD | 347 |
| ENSG00000245466.1 | 18.95006786 | 5.874086302 | 0.769734093 | 6.851829 | 7.29E-12 | 1.55E-10 | ENSG00000245466 | LOC10192 | 1.02E+08 |
| ENSG00000236882.8 | 61.78240316 | 5.871777409 | 0.956199808 | 5.513259 | 3.52E-08 | 5.09E-07 | ENSG00000236882 | LINC0155 | 202299 |
| ENSG00000175121.11 | 1069.462 | 5.854569 | 0.4428141 | 11.86631 | 1.77E-32 | 1.36E-30 | ENSG00000175121 | WFDC5 | 149708 |
| ENSG00000141469.18 | 12.75491279 | 5.826896195 | 0.842159047 | 6.206543 | 5.42E-10 | 9.63E-09 | ENSG00000141469 | SLC14A1 | 6563 |
| ENSG00000204539.4 | 11.40631269 | 5.820594449 | 0.830765997 | 6.284073 | 3.30E-10 | 6.03E-09 | ENSG00000204539 | CDSN | 1041 |
| ENSG00000124467.18 | 5.339373 | 5.819321 | 0.9776884 | 5.33843 | 9.38E-08 | 1.30E-06 | ENSG00000124467 | PSG8 | 440533 |
| ENSG00000286046.1 | 4.940350471 | 5.808608083 | 1.164581532 | 4.472515 | 7.73E-06 | 8.34E-05 | ENSG00000286046 | NA | NA |
| ENSG00000165799.5 | 11870.58515 | 5.80393696 | 0.178575197 | 29.14143 | 1.07E-186 | 1.06E-183 | ENSG00000165799 | RNASE7 | 84659 |
| ENSG00000165507.8 | 660.4109789 | 5.787298157 | 0.486138253 | 10.67042 | 1.40E-26 | 7.67E-25 | ENSG00000165507 | DEPP1 | 11067 |
| ENSG00000180316.12 | 31.37196199 | 5.775698015 | 0.603448247 | 8.576871 | 9.75E-18 | 3.24E-16 | ENSG00000180316 | PNPLA1 | 285848 |
| ENSG00000187134.14 | 3250.315744 | 5.758778732 | 0.233602647 | 22.08356 | 4.55E-108 | 2.09E-105 | ENSG00000187134 | AKR1C1 | 1645 |
| ENSG00000287879.1 | 4.978019512 | 5.757455638 | 1.127227783 | 4.575345 | 4.75E-06 | 5.26E-05 | ENSG00000287879 | NA | NA |
| ENSG00000185467.7 | 124.275634 | 5.757014846 | 0.328309074 | 15.7078 | 1.34E-55 | 2.27E-53 | ENSG00000185467 | KPNA7 | 402569 |
| ENSG00000143369.15 | 13404.99105 | 5.748327067 | 0.163067937 | 31.57167 | 9.04E-219 | 1.20E-215 | ENSG00000143369 | ECM1 | 1893 |
| ENSG00000227621.1 | 4.801034488 | 5.742374881 | 1.155374468 | 4.45083 | 8.55E-06 | 9.17E-05 | ENSG00000227621 | NA | NA |
| ENSG00000036672.16 | 720.3617919 | 5.728608044 | 0.307003036 | 16.7054 | 1.20E-62 | 2.45E-60 | ENSG00000036672 | USP2 | 9099 |
| ENSG00000253810.1 | 6.060515021 | 5.714574034 | 0.966957576 | 5.289347 | 1.23E-07 | 1.68E-06 | ENSG00000253810 | NA | NA |
| ENSG00000167755.15 | 4659.913 | 5.714043 | 0.4363934 | 11.71888 | 1.02E-31 | 7.59E-30 | ENSG00000167755 | KLK6 | 5653 |
| ENSG00000269495.1 | 10.96681 | 5.710297 | 0.9414315 | 5.428219 | 5.69E-08 | 8.03E-07 | ENSG00000269495 | NA | NA |
| ENSG00000168703.5 | 5518.575 | 5.7082 | 0.3328594 | 15.34642 | 3.74E-53 | 5.83E-51 | ENSG00000168703 | WFDC12 | 128488 |
| ENSG00000225899.7 | 4.704377429 | 5.707815794 | 1.274818939 | 4.006699 | 6.16E-05 | 0.000574554 | ENSG00000225899 | FRG2B | 441581 |
| ENSG00000152137.6 | 826.5211323 | 5.70485351 | 0.143175603 | 35.65449 | 2.01E-278 | 4.11E-275 | ENSG00000152137 | HSPB8 | 26353 |
| ENSG00000134539.17 | 17.11640923 | 5.702258293 | 0.868475123 | 5.874962 | 4.23E-09 | 6.78E-08 | ENSG00000134539 | KLRD1 | 3824 |

| | | | | | | | | | |
|---|---|---|---|---|---|---|---|---|---|
| ENSG00000224308.1 | 9.426935716 | 5.693021237 | 0.922671191 | 5.519866 | 3.39E-08 | 4.93E-07 | ENSG00000224308 | LINC01527 | 1.02E+08 |
| ENSG00000241479.1 | 4.813248489 | 5.690423383 | 1.068020359 | 4.766223 | 1.88E-06 | 2.19E-05 | ENSG00000241479 | MECOM-A | 1.05E+08 |
| ENSG00000181449.4 | 11.37501924 | 5.689801936 | 0.873069432 | 5.829779 | 5.55E-09 | 8.82E-08 | ENSG00000181449 | SOX2 | 6657 |
| ENSG00000108849.7 | 6.043305346 | 5.668930485 | 1.007615541 | 5.03062 | 4.89E-07 | 6.15E-06 | ENSG00000108849 | PPY | 5539 |
| ENSG00000168367.11 | 44.44952548 | 5.661157639 | 0.544521188 | 9.294694 | 1.48E-20 | 5.87E-19 | ENSG00000168367 | LINC00917 | 732275 |
| ENSG00000213904.9 | 103.3627 | 5.661015 | 0.4020396 | 12.58835 | 2.45E-36 | 2.13E-34 | ENSG00000213904 | LIPE-AS1 | 1.01E+08 |
| ENSG00000184226.15 | 4.607550808 | 5.65524679 | 1.036085604 | 4.879179 | 1.07E-06 | 1.28E-05 | ENSG00000184226 | PCDH9 | 5101 |
| ENSG00000260976.2 | 4.488447931 | 5.65148577 | 1.068238004 | 4.728802 | 2.26E-06 | 2.60E-05 | ENSG00000260976 | NA | NA |
| ENSG00000158786.4 | 174.6659881 | 5.648855807 | 0.280636794 | 17.99071 | 2.30E-72 | 5.90E-70 | ENSG00000158786 | PLA2G2F | 64600 |
| ENSG00000285946.1 | 10.21127211 | 5.640019373 | 0.881793355 | 5.715647 | 1.09E-08 | 1.68E-07 | ENSG00000285946 | NA | NA |
| ENSG00000164120.14 | 114.6023092 | 5.634922123 | 0.397585537 | 12.66375 | 9.39E-37 | 8.33E-35 | ENSG00000164120 | HPGD | 3248 |
| ENSG00000228033.6 | 23.98537226 | 5.610910224 | 0.680321644 | 7.365502 | 1.76E-13 | 4.31E-12 | ENSG00000228033 | NA | NA |
| ENSG00000187479.7 | 70.01298839 | 5.599407306 | 0.371908764 | 13.44256 | 3.40E-41 | 3.57E-39 | ENSG00000187479 | C11orf96 | 387763 |
| ENSG00000237330.3 | 1160.968038 | 5.596755355 | 0.250436704 | 19.95217 | 1.44E-88 | 4.90E-86 | ENSG00000237330 | RNF223 | 401934 |
| ENSG00000228126.1 | 27.50973304 | 5.590702246 | 0.592498458 | 8.423148 | 3.67E-17 | 1.17E-15 | ENSG00000228126 | FALEC | 1.01E+08 |
| ENSG00000284631.1 | 5.433666656 | 5.573933025 | 1.08606103 | 4.579791 | 4.65E-06 | 5.15E-05 | ENSG00000284631 | NA | NA |
| ENSG00000238755.3 | 23.37650729 | 5.569383286 | 0.643469131 | 7.7228 | 1.14E-14 | 3.03E-13 | ENSG00000238755 | NA | NA |
| ENSG00000236151.1 | 8.526472 | 5.568532 | 0.8945175 | 5.554426 | 2.79E-08 | 4.11E-07 | ENSG00000236151 | NA | NA |
| ENSG00000233858.4 | 4.269077308 | 5.56791015 | 1.006173885 | 4.937427 | 7.92E-07 | 9.71E-06 | ENSG00000233858 | NA | NA |
| ENSG00000141506.13 | 5.15245531 | 5.562083303 | 0.980033068 | 5.063179 | 4.12E-07 | 5.24E-06 | ENSG00000141506 | PIK3R5 | 23533 |
| ENSG00000168447.11 | 14.11045214 | 5.561389847 | 0.946651088 | 5.240991 | 1.60E-07 | 2.15E-06 | ENSG00000168447 | SCNN1B | 6338 |
| ENSG00000269919.1 | 20.26398958 | 5.543004839 | 0.67989102 | 7.27029 | 3.59E-13 | 8.51E-12 | ENSG00000269919 | NA | NA |
| ENSG00000113389.16 | 45.72289633 | 5.53609023 | 0.486937643 | 10.13701 | 3.79E-24 | 1.81E-22 | ENSG00000113389 | NPR3 | 4883 |
| ENSG00000278925.1 | 4.397002131 | 5.529420458 | 0.992307453 | 4.967634 | 6.78E-07 | 8.40E-06 | ENSG00000278925 | NA | NA |
| ENSG00000250258.1 | 8.607115486 | 5.528941301 | 0.922569371 | 5.342624 | 9.16E-08 | 1.27E-06 | ENSG00000250258 | NA | NA |
| ENSG00000133958.13 | 4.294994378 | 5.526672327 | 0.976000037 | 5.04782 | 4.47E-07 | 5.64E-06 | ENSG00000133958 | UNC79 | 57578 |
| ENSG00000128849.10 | 457.2007966 | 5.514335268 | 0.420969569 | 11.67385 | 1.73E-31 | 1.28E-29 | ENSG00000128849 | CGNL1 | 84952 |
| ENSG00000235292.2 | 4.172241 | 5.511738 | 1.209046 | 4.06249 | 4.86E-05 | 0.000460639 | ENSG00000235292 | NA | NA |
| ENSG00000265345.1 | 40.20407424 | 5.509489004 | 0.470965907 | 10.4243 | 1.92E-25 | 1.00E-23 | ENSG00000265345 | MIR5188 | 1.01E+08 |
| ENSG00000108684.14 | 4.104210002 | 5.507821878 | 1.052643329 | 4.662379 | 3.13E-06 | 3.52E-05 | ENSG00000108684 | ASIC2 | 40 |
| ENSG00000204941.14 | 44.06277 | 5.500397 | 0.4488617 | 10.91739 | 9.52E-28 | 5.64E-26 | ENSG00000204941 | PSG5 | 5673 |
| ENSG00000105549.10 | 76.62019055 | 5.491170641 | 0.350110106 | 13.97038 | 2.36E-44 | 2.80E-42 | ENSG00000105549 | THEG | 51298 |
| ENSG00000204021.4 | 193.5540888 | 5.485035401 | 0.274555536 | 17.79252 | 8.08E-71 | 1.99E-68 | ENSG00000204021 | LIPK | 643414 |
| ENSG00000260386.6 | 4.119596198 | 5.479766348 | 1.046254735 | 4.664033 | 3.10E-06 | 3.50E-05 | ENSG00000260386 | LDC1P | 149086 |
| ENSG00000256916.1 | 4.008720264 | 5.479428312 | 1.107489394 | 4.405847 | 1.05E-05 | 0.000111067 | ENSG00000256916 | NA | NA |
| ENSG00000214491.8 | 545.5163 | 5.476573 | 0.4726792 | 10.31688 | 5.91E-25 | 2.98E-23 | ENSG00000214491 | SEC14L6 | 730005 |
| ENSG00000100678.18 | 3.906402951 | 5.467051689 | 1.02895429 | 4.730095 | 2.24E-06 | 2.58E-05 | ENSG00000100678 | SLC8A3 | 6547 |
| ENSG00000248441.7 | 3.922789096 | 5.456678364 | 1.291533516 | 3.760397 | 0.00017 | 0.00146775 | ENSG00000248441 | LINC01197 | 400456 |
| ENSG00000012171.19 | 406.9467872 | 5.441129474 | 0.262470901 | 18.44444 | 5.78E-76 | 1.60E-73 | ENSG00000012171 | SEMA3B | 7869 |
| ENSG00000145824.12 | 80.27655274 | 5.437004287 | 0.38265965 | 12.64049 | 1.26E-36 | 1.11E-34 | ENSG00000145824 | CXCL14 | 9547 |
| ENSG00000230442.5 | 4.324617761 | 5.430590112 | 0.972772547 | 4.965796 | 6.84E-07 | 8.47E-06 | ENSG00000230442 | ASB15-AS | 1.03E+08 |
| ENSG00000228476.2 | 3.951701 | 5.430501 | 1.141215 | 4.232771 | 2.31E-05 | 0.000229897 | ENSG00000228476 | NA | NA |

| | | | | | | | | | |
|---|---|---|---|---|---|---|---|---|---|
| ENSG00000151632.17 | 2446.967803 | 5.430346297 | 0.200111952 | 24.13822 | 9.93E-129 | 5.62E-126 | ENSG00000151632 | AKR1C2 | 1646 |
| ENSG00000283227.1 | 160.2735953 | 5.425564395 | 0.325768979 | 14.81284 | 1.21E-49 | 1.69E-47 | ENSG00000283227 | NA | NA |
| ENSG00000125848.10 | 3.776754 | 5.423796 | 0.9959592 | 4.843368 | 1.28E-06 | 1.51E-05 | ENSG00000125848 | FLRT3 | 23767 |
| ENSG00000010310.9 | 846.3764 | 5.415459 | 0.4445084 | 10.83323 | 2.40E-27 | 1.40E-25 | ENSG00000010310 | GIPR | 2696 |
| ENSG00000128510.12 | 4059.182343 | 5.406505894 | 0.430827979 | 11.15644 | 6.66E-29 | 4.26E-27 | ENSG00000128510 | CPA4 | 51200 |
| ENSG00000163687.13 | 34.39928416 | 5.402120612 | 0.505062713 | 9.507969 | 1.94E-21 | 8.06E-20 | ENSG00000163687 | DNASE1L3 | 1776 |
| ENSG00000230587.2 | 3.70919212 | 5.398608688 | 1.006271628 | 4.768701 | 1.85E-06 | 2.16E-05 | ENSG00000230587 | NA | NA |
| ENSG00000287218.1 | 3.698383449 | 5.398441793 | 1.054807563 | 4.549116 | 5.39E-06 | 5.92E-05 | ENSG00000287218 | NA | NA |
| ENSG00000204542.2 | 4.068876831 | 5.397691689 | 1.025995435 | 4.676134 | 2.92E-06 | 3.31E-05 | ENSG00000204542 | C6orf15 | 29113 |
| ENSG00000224367.6 | 33.44765214 | 5.394722619 | 0.580401768 | 8.261041 | 1.44E-16 | 4.42E-15 | ENSG00000224367 | NA | NA |
| ENSG00000138347.15 | 119.056288 | 5.386307463 | 0.468325917 | 10.22004 | 1.61E-24 | 7.88E-23 | ENSG00000138347 | MYPN | 84665 |
| ENSG00000211899.10 | 3.841406889 | 5.381658981 | 1.111048651 | 4.303735 | 1.68E-05 | 0.000171039 | ENSG00000211899 | IGHM | 3507 |
| ENSG00000260186.5 | 25.42525892 | 5.378192109 | 0.603710293 | 7.91471 | 2.48E-15 | 6.98E-14 | ENSG00000260186 | NA | NA |
| ENSG00000225683.6 | 3.721232862 | 5.362068254 | 1.02490761 | 4.646339 | 3.38E-06 | 3.80E-05 | ENSG00000225683 | PACRG-AS | 729658 |
| ENSG00000230908.2 | 3.863634 | 5.360495 | 1.002995 | 4.746282 | 2.07E-06 | 2.39E-05 | ENSG00000230908 | LOC10798 | 1.08E+08 |
| ENSG00000256081.2 | 3.83219569 | 5.35932271 | 1.129969996 | 4.211902 | 2.53E-05 | 0.000250799 | ENSG00000256081 | NA | NA |
| ENSG00000262921.1 | 3.608895545 | 5.358034579 | 1.064154573 | 4.471187 | 7.78E-06 | 8.39E-05 | ENSG00000262921 | NA | NA |
| ENSG00000251638.1 | 5.462744745 | 5.352880975 | 1.007631979 | 4.716882 | 2.39E-06 | 2.74E-05 | ENSG00000251638 | NA | NA |
| ENSG00000232873.1 | 9.247998278 | 5.341648067 | 0.854686187 | 5.547823 | 2.89E-08 | 4.25E-07 | ENSG00000232873 | NA | NA |
| ENSG00000113327.16 | 3.557601396 | 5.341131476 | 1.142834362 | 4.148573 | 3.35E-05 | 0.00032553 | ENSG00000113327 | GABRG2 | 2566 |
| ENSG00000263325.1 | 3.767397584 | 5.338647774 | 1.003828048 | 4.720577 | 2.35E-06 | 2.70E-05 | ENSG00000263325 | NA | NA |
| ENSG00000274428.1 | 19.85180737 | 5.335577937 | 0.64658373 | 7.323998 | 2.41E-13 | 5.81E-12 | ENSG00000274428 | NA | NA |
| ENSG00000278954.1 | 3.521456169 | 5.32819346 | 1.245950949 | 3.794847 | 0.000148 | 0.001291197 | ENSG00000278954 | NA | NA |
| ENSG00000255501.2 | 91.62592915 | 5.32801695 | 0.311159121 | 15.19485 | 3.83E-52 | 5.72E-50 | ENSG00000255501 | CARD18 | 59082 |
| ENSG00000147234.10 | 42.90924 | 5.321751 | 0.5306588 | 8.897904 | 5.69E-19 | 2.04E-17 | ENSG00000147234 | FRMPD3 | 84443 |
| ENSG00000057657.17 | 5645.983192 | 5.311683856 | 0.213491085 | 22.0697 | 6.18E-108 | 2.74E-105 | ENSG00000057657 | PRDM1 | 639 |
| ENSG00000042062.12 | 350.2584 | 5.309195 | 0.5043181 | 9.337747 | 9.84E-21 | 3.96E-19 | ENSG00000042062 | RIPOR3 | 140876 |
| ENSG00000073734.10 | 14.35681958 | 5.306435648 | 0.774078495 | 6.08005 | 1.20E-09 | 2.04E-08 | ENSG00000073734 | ABCB11 | 8647 |
| ENSG00000168952.15 | 386.994125 | 5.305425038 | 0.761371304 | 6.180198 | 6.40E-10 | 1.13E-08 | ENSG00000168952 | STXBP6 | 29091 |
| ENSG00000102760.13 | 66.94566065 | 5.305279652 | 0.428427748 | 10.98267 | 4.63E-28 | 2.83E-26 | ENSG00000102760 | RGCC | 28984 |
| ENSG00000227087.4 | 3.439956104 | 5.292657789 | 1.260813976 | 3.721927 | 0.000198 | 0.001686982 | ENSG00000227087 | NA | NA |
| ENSG00000248498.4 | 24.89923651 | 5.292228417 | 0.842600729 | 5.568745 | 2.57E-08 | 3.79E-07 | ENSG00000248498 | ASNSP1 | 389652 |
| ENSG00000178934.4 | 2468.5 | 5.277645 | 0.2241489 | 20.86847 | 1.04E-96 | 4.24E-94 | ENSG00000178934 | LGALS7 | 3963 |
| ENSG00000189223.15 | 5693.738089 | 5.275509567 | 0.206568773 | 22.63415 | 2.00E-113 | 1.00E-110 | ENSG00000189223 | PAX8-AS1 | 654433 |
| ENSG00000248112.1 | 59.75351212 | 5.271376899 | 0.358737913 | 13.0217 | 9.21E-39 | 8.82E-37 | ENSG00000248112 | NA | NA |
| ENSG00000248400.2 | 10.70960567 | 5.265295209 | 0.788294772 | 5.918212 | 3.25E-09 | 5.30E-08 | ENSG00000248400 | NA | NA |
| ENSG00000154025.15 | 22.06329528 | 5.263243658 | 0.599653215 | 7.776567 | 7.45E-15 | 2.01E-13 | ENSG00000154025 | SLC5A10 | 125206 |
| ENSG00000186723.4 | 73.76421 | 5.258015 | 0.4169109 | 11.17269 | 5.55E-29 | 3.58E-27 | ENSG00000186723 | OR10H1 | 26539 |
| ENSG00000103528.17 | 4.763950773 | 5.257357919 | 0.964693085 | 4.827813 | 1.38E-06 | 1.63E-05 | ENSG00000103528 | SYT17 | 51760 |
| ENSG00000117115.13 | 282.8259643 | 5.243176148 | 0.266262417 | 17.43835 | 4.22E-68 | 9.77E-66 | ENSG00000117115 | PADI2 | 11240 |
| ENSG00000162572.20 | 684.9877846 | 5.242005221 | 0.55501145 | 8.363801 | 6.07E-17 | 1.90E-15 | ENSG00000162572 | SCNN1D | 6339 |
| ENSG00000123612.16 | 2863.157467 | 5.234854524 | 0.242468148 | 19.11531 | 1.88E-81 | 5.63E-79 | ENSG00000123612 | ACVR1C | 130399 |

| | | | | | | | | | |
|---|---|---|---|---|---|---|---|---|---|
| ENSG00000288100.1 | 4.185196935 | 5.231547481 | 1.027283088 | 4.508541 | 6.53E-06 | 7.11E-05 | ENSG00000288100 | NA | NA |
| ENSG00000286818.1 | 9.214286041 | 5.231405522 | 0.912875702 | 5.073424 | 3.91E-07 | 4.98E-06 | ENSG00000286818 | NA | NA |
| ENSG00000159516.9 | 3222.195588 | 5.229972165 | 0.241410052 | 19.17887 | 5.56E-82 | 1.72E-79 | ENSG00000159516 | SPRR2G | 6706 |
| ENSG00000105668.7 | 3.525209 | 5.217557 | 1.045273 | 4.41756 | 9.98E-06 | 0.00010579 | ENSG00000105668 | UPK1A | 11045 |
| ENSG00000198670.11 | 3.393647163 | 5.214356792 | 1.063605437 | 4.33841 | 1.44E-05 | 0.000147975 | ENSG00000198670 | LPA | 4018 |
| ENSG00000082293.13 | 6.5509011 | 5.214028785 | 0.968211733 | 4.765516 | 1.88E-06 | 2.19E-05 | ENSG00000082293 | COL19A1 | 1310 |
| ENSG00000249837.1 | 3.712153081 | 5.204503894 | 1.043569487 | 4.412264 | 1.02E-05 | 0.000108111 | ENSG00000249837 | NA | NA |
| ENSG00000163092.19 | 6.576673322 | 5.203938309 | 0.942216032 | 4.886287 | 1.03E-06 | 1.24E-05 | ENSG00000163092 | XIRP2 | 129446 |
| ENSG00000184106.8 | 4.898193587 | 5.200628568 | 0.949318734 | 4.846242 | 1.26E-06 | 1.49E-05 | ENSG00000184106 | TREML3P | 340206 |
| ENSG00000229782.1 | 7.441404631 | 5.199692773 | 0.902310653 | 5.097682 | 3.44E-07 | 4.41E-06 | ENSG00000229782 | NA | NA |
| ENSG00000179846.9 | 66.44465 | 5.196172 | 0.5196157 | 8.845329 | 9.13E-19 | 3.23E-17 | ENSG00000179846 | NKPD1 | 284353 |
| ENSG00000163273.4 | 3.808506657 | 5.189446333 | 1.028004374 | 4.464423 | 8.03E-06 | 8.64E-05 | ENSG00000163273 | NPPC | 4880 |
| ENSG00000169035.12 | 22865.82 | 5.187539 | 0.4572808 | 10.03221 | 1.10E-23 | 5.17E-22 | ENSG00000169035 | KLK7 | 5650 |
| ENSG00000265107.3 | 41.46230891 | 5.187534156 | 0.455306132 | 10.07571 | 7.07E-24 | 3.38E-22 | ENSG00000265107 | GJA5 | 2702 |
| ENSG00000189056.14 | 29.59825774 | 5.183558761 | 0.506337065 | 9.052386 | 1.40E-19 | 5.21E-18 | ENSG00000189056 | RELN | 5649 |
| ENSG00000107593.17 | 8.83056661 | 5.178351628 | 0.860630402 | 5.319765 | 1.04E-07 | 1.43E-06 | ENSG00000107593 | PKD2L1 | 9033 |
| ENSG00000158528.12 | 3.164854113 | 5.170433334 | 1.00861357 | 4.531402 | 5.86E-06 | 6.42E-05 | ENSG00000158528 | PPP1R9A | 55607 |
| ENSG00000188624.3 | 6.36064 | 5.170362 | 0.9615695 | 4.753024 | 2.00E-06 | 2.32E-05 | ENSG00000188624 | IGFL3 | 388555 |
| ENSG00000178033.6 | 57.69123465 | 5.16991233 | 0.580129735 | 7.877397 | 3.34E-15 | 9.33E-14 | ENSG00000178033 | CALHM5 | 254228 |
| ENSG00000243130.8 | 3.441014 | 5.162917 | 1.019169 | 4.477097 | 7.57E-06 | 8.18E-05 | ENSG00000243130 | PSG11 | 5680 |
| ENSG00000233776.5 | 11.30585382 | 5.159144449 | 0.781423197 | 5.834411 | 5.40E-09 | 8.58E-08 | ENSG00000233776 | LINC0125 | 1.02E+08 |
| ENSG00000234840.2 | 11.88314828 | 5.156294647 | 0.793579175 | 5.741449 | 9.39E-09 | 1.45E-07 | ENSG00000234840 | LINC0123 | 441389 |
| ENSG00000117598.12 | 6.969357886 | 5.146043193 | 0.888714175 | 5.115304 | 3.13E-07 | 4.04E-06 | ENSG00000117598 | PLPPR5 | 163404 |
| ENSG00000169474.4 | 2932.312561 | 5.145863617 | 0.362604433 | 12.5367 | 4.70E-36 | 4.05E-34 | ENSG00000169474 | SPRR1A | 6698 |
| ENSG00000131737.5 | 122.8747633 | 5.143834004 | 0.284263801 | 15.98457 | 1.64E-57 | 2.89E-55 | ENSG00000131737 | KRT34 | 3885 |
| ENSG00000137726.17 | 15.85901791 | 5.134325134 | 0.734169665 | 6.176127 | 6.57E-10 | 1.16E-08 | ENSG00000137726 | FXYD6 | 53826 |
| ENSG00000226715.4 | 7.129216255 | 5.132101711 | 0.93403622 | 4.852169 | 1.22E-06 | 1.46E-05 | ENSG00000226715 | NA | NA |
| ENSG00000288011.1 | 4.553661471 | 5.130199765 | 1.137593113 | 3.982267 | 6.83E-05 | 0.000632305 | ENSG00000288011 | LOC28304 | 283045 |
| ENSG00000163817.16 | 6.349939471 | 5.12455676 | 0.923769918 | 4.897926 | 9.69E-07 | 1.17E-05 | ENSG00000163817 | SLC6A20 | 54716 |
| ENSG00000118946.12 | 3.837150081 | 5.120239795 | 1.431577929 | 3.157523 | 0.001591 | 0.01123007 | ENSG00000118946 | PCDH17 | 27253 |
| ENSG00000251138.7 | 3.066725985 | 5.095169313 | 1.032170242 | 4.355066 | 1.33E-05 | 0.000137907 | ENSG00000251138 | LINC0288 | 1.01E+08 |
| ENSG00000102195.9 | 20.75215 | 5.09179 | 0.6422657 | 6.993663 | 2.68E-12 | 5.89E-11 | ENSG00000102195 | GPR50 | 9248 |
| ENSG00000182836.10 | 7.400007364 | 5.086615604 | 0.94921237 | 4.726672 | 2.28E-06 | 2.62E-05 | ENSG00000182836 | PLCXD3 | 345557 |
| ENSG00000250064.1 | 64.43495294 | 5.084884911 | 0.512123271 | 8.757432 | 2.00E-18 | 6.92E-17 | ENSG00000250064 | NA | NA |
| ENSG00000184678.10 | 287.2122636 | 5.083891442 | 0.31865729 | 14.0712 | 5.71E-45 | 6.88E-43 | ENSG00000184678 | H2BC21 | 8349 |
| ENSG00000214866.8 | 3.561064672 | 5.079744992 | 1.002360721 | 4.469194 | 7.85E-06 | 8.47E-05 | ENSG00000214866 | DCDC2C | 728597 |
| ENSG00000151892.14 | 11.54059092 | 5.072264113 | 0.745730962 | 5.997155 | 2.01E-09 | 3.34E-08 | ENSG00000151892 | GFRA1 | 2674 |
| ENSG00000125775.15 | 1947.652 | 5.0708 | 0.3274628 | 13.65285 | 1.94E-42 | 2.14E-40 | ENSG00000125775 | SDCBP2 | 27111 |
| ENSG00000115602.16 | 298.9890301 | 5.069162755 | 0.209201561 | 21.36295 | 2.96E-101 | 1.23E-98 | ENSG00000115602 | IL1RL1 | 9173 |
| ENSG00000233208.6 | 3.153309464 | 5.067955777 | 1.0654982 | 4.193302 | 2.75E-05 | 0.000270974 | ENSG00000233208 | LINC0064 | 400238 |
| ENSG00000237422.2 | 2.950808996 | 5.065062603 | 1.181058355 | 3.780561 | 0.000156 | 0.001364013 | ENSG00000237422 | NA | NA |
| ENSG00000137747.16 | 1888.62673 | 5.061879202 | 0.360965644 | 12.36095 | 4.25E-35 | 3.51E-33 | ENSG00000137747 | TMPRSS1 | 84000 |

| | | | | | | | | | |
|---|---|---|---|---|---|---|---|---|---|
| ENSG00000143556.9 | 43375.32771 | 5.061758527 | 0.467042358 | 9.55322 | 1.26E-21 | 5.30E-20 | ENSG00000143556 | S100A7 | 6278 |
| ENSG00000220248.1 | 2.936047 | 5.05821 | 1.099735 | 4.053894 | 5.04E-05 | 0.000475871 | ENSG00000220248 | ZNF402P | 266673 |
| ENSG00000006210.7 | 60.54260248 | 5.052351515 | 0.442859431 | 10.05365 | 8.85E-24 | 4.20E-22 | ENSG00000006210 | CX3CL1 | 6376 |
| ENSG00000262097.2 | 5.993515434 | 5.048341237 | 0.987250477 | 4.505788 | 6.61E-06 | 7.20E-05 | ENSG00000262097 | NA | NA |
| ENSG00000136449.14 | 109.9668913 | 5.04513723 | 0.378227911 | 11.75254 | 6.85E-32 | 5.12E-30 | ENSG00000136449 | MYCBPAP | 84073 |
| ENSG00000084734.9 | 5.754570858 | 5.041137682 | 0.998266868 | 4.448848 | 8.63E-06 | 9.25E-05 | ENSG00000084734 | GCKR | 2646 |
| ENSG00000170091.11 | 5.73746892 | 5.040089559 | 1.037289813 | 4.280472 | 1.86E-05 | 0.000188351 | ENSG00000170091 | NSG2 | 51617 |
| ENSG00000279619.1 | 3.157746 | 5.03915 | 1.229 | 3.612003 | 0.000304 | 0.002506404 | ENSG00000279619 | NA | NA |
| ENSG00000228674.4 | 3.271377 | 5.038888 | 1.036385 | 4.283049 | 1.84E-05 | 0.000186749 | ENSG00000228674 | NA | NA |
| ENSG00000196248.5 | 2.898511038 | 5.026037412 | 1.103779193 | 4.009894 | 6.07E-05 | 0.000567233 | ENSG00000196248 | OR10S1 | 219873 |
| ENSG00000174567.8 | 11.98380741 | 5.019211743 | 0.83802877 | 5.273341 | 1.34E-07 | 1.82E-06 | ENSG00000174567 | GOLT1A | 127845 |
| ENSG00000019991.18 | 2.852517728 | 5.018470904 | 1.057937259 | 4.176496 | 2.96E-05 | 0.00029017 | ENSG00000019991 | HGF | 3082 |
| ENSG00000184459.8 | 5.725258 | 5.017431 | 0.9970312 | 4.430584 | 9.40E-06 | 0.000100156 | ENSG00000184459 | BPIFC | 254240 |
| ENSG00000189377.8 | 16.07499 | 5.017374 | 0.6868184 | 6.431648 | 1.26E-10 | 2.39E-09 | ENSG00000189377 | CXCL17 | 284340 |
| ENSG00000108244.16 | 206.9634127 | 5.016979432 | 0.335816186 | 13.15297 | 1.64E-39 | 1.61E-37 | ENSG00000108244 | KRT23 | 25984 |
| ENSG00000178789.9 | 3.304105782 | 5.015933549 | 1.276410141 | 3.459651 | 0.000541 | 0.004255083 | ENSG00000178789 | CD300LB | 124599 |
| ENSG00000226051.8 | 5.98210542 | 5.008465149 | 1.072484739 | 4.110516 | 3.95E-05 | 0.00038065 | ENSG00000226051 | ZNF503-A: | 253264 |
| ENSG00000246363.3 | 8.674305671 | 5.008116161 | 0.849216542 | 5.190803 | 2.09E-07 | 2.76E-06 | ENSG00000246363 | LINC0245ε | 728084 |
| ENSG00000108823.16 | 2.818142787 | 5.008070003 | 1.293599968 | 3.407599 | 0.000655 | 0.005061588 | ENSG00000108823 | SGCA | 6442 |
| ENSG00000198092.5 | 215.909553 | 5.005167611 | 0.345298936 | 12.75755 | 2.83E-37 | 2.58E-35 | ENSG00000198092 | TMPRSS1 | 389208 |
| ENSG00000279283.1 | 76.94190732 | 5.002352405 | 0.453706922 | 9.703075 | 2.93E-22 | 1.27E-20 | ENSG00000279283 | NA | NA |
| ENSG00000249451.1 | 3.464046013 | 4.999420589 | 1.063806626 | 4.135545 | 3.54E-05 | 0.000343682 | ENSG00000249451 | NA | NA |
| ENSG00000204618.8 | 222.005964 | 4.998090375 | 0.257656138 | 17.06961 | 2.50E-65 | 5.45E-63 | ENSG00000204618 | RNF39 | 80352 |
| ENSG00000166603.5 | 13.93364219 | 4.990829622 | 0.840596637 | 5.223468 | 1.76E-07 | 2.35E-06 | ENSG00000166603 | MC4R | 4160 |
| ENSG00000232756.1 | 4.274926869 | 4.989885843 | 1.037726622 | 4.230291 | 2.33E-05 | 0.000232185 | ENSG00000232756 | NA | NA |
| ENSG00000180332.6 | 23.16024968 | 4.989191809 | 0.525238932 | 8.356562 | 6.46E-17 | 2.02E-15 | ENSG00000180332 | KCTD4 | 386618 |
| ENSG00000182752.10 | 17.15150399 | 4.988774265 | 0.669347085 | 6.556799 | 5.50E-11 | 1.07E-09 | ENSG00000182752 | PAPPA | 5069 |
| ENSG00000234699.1 | 2.750794794 | 4.972006489 | 1.381152942 | 3.165476 | 0.001548 | 0.010950768 | ENSG00000234699 | NA | NA |
| ENSG00000204421.3 | 20.72670797 | 4.964051861 | 0.573149289 | 7.614163 | 2.65E-14 | 6.88E-13 | ENSG00000204421 | LY6G6C | 80740 |
| ENSG00000171649.12 | 18.94967 | 4.951234 | 0.7558676 | 5.756609 | 8.58E-09 | 1.34E-07 | ENSG00000171649 | ZIK1 | 284307 |
| ENSG00000260377.1 | 3.763830938 | 4.948668981 | 1.348894679 | 3.223876 | 0.001265 | 0.009153988 | ENSG00000260377 | NA | NA |
| ENSG00000101445.10 | 2.873117 | 4.947523 | 1.058505 | 4.107229 | 4.00E-05 | 0.000385686 | ENSG00000101445 | PPP1R16E | 26051 |
| ENSG00000260846.2 | 2.704706232 | 4.940358686 | 1.136867894 | 3.817821 | 0.000135 | 0.001186448 | ENSG00000260846 | NA | NA |
| ENSG00000239808.3 | 2.789546865 | 4.931397529 | 1.21557529 | 3.563249 | 0.000366 | 0.002980271 | ENSG00000239808 | RN7SL255 | 1.06E+08 |
| ENSG00000284196.3 | 12.56705836 | 4.929047662 | 0.731132038 | 5.92102 | 3.20E-09 | 5.22E-08 | ENSG00000284196 | NA | NA |
| ENSG00000255977.1 | 2.678005081 | 4.927589084 | 1.013006527 | 4.272025 | 1.94E-05 | 0.000195187 | ENSG00000255977 | NA | NA |
| ENSG00000185873.7 | 25.03599778 | 4.925817415 | 0.529014133 | 8.17713 | 2.91E-16 | 8.66E-15 | ENSG00000185873 | TMPRSS1 | 132724 |
| ENSG00000156282.4 | 2.701928 | 4.925317 | 1.05054 | 4.117231 | 3.83E-05 | 0.000370939 | ENSG00000156282 | CLDN17 | 26285 |
| ENSG00000177191.2 | 130.9417 | 4.919628 | 0.5969372 | 7.236318 | 4.61E-13 | 1.09E-11 | ENSG00000177191 | B3GNT8 | 374907 |
| ENSG00000257052.1 | 7.539268289 | 4.914372635 | 0.867741993 | 4.971953 | 6.63E-07 | 8.22E-06 | ENSG00000257052 | NA | NA |
| ENSG00000151812.14 | 30.57117777 | 4.913531156 | 0.668585535 | 6.451727 | 1.11E-10 | 2.10E-09 | ENSG00000151812 | SLC35F4 | 341880 |
| ENSG00000088386.17 | 48.90390924 | 4.912227876 | 0.390716322 | 11.03672 | 2.54E-28 | 1.57E-26 | ENSG00000088386 | SLC15A1 | 6564 |

| | | | | | | | | | |
|---|---|---|---|---|---|---|---|---|---|
| ENSG00000270442.1 | 3.140885252 | 4.904864023 | 1.074301191 | 4.00713 | 6.15E-05 | 0.000573709 | ENSG00000270442 | PPIGP1 | 1E+08 |
| ENSG00000245904.4 | 15.47373811 | 4.902077595 | 0.671115559 | 6.410338 | 1.45E-10 | 2.73E-09 | ENSG00000245904 | NA | NA |
| ENSG00000159337.6 | 467.5910129 | 4.900640771 | 0.484519409 | 8.876096 | 6.92E-19 | 2.47E-17 | ENSG00000159337 | PLA2G4D | 283748 |
| ENSG00000239279.3 | 2.6239284 | 4.900514741 | 1.117238584 | 3.849236 | 0.000118 | 0.001055676 | ENSG00000239279 | RN7SL184 | 1.06E+08 |
| ENSG00000173727.12 | 358.907341 | 4.899130273 | 0.276009747 | 15.57601 | 1.06E-54 | 1.71E-52 | ENSG00000173727 | NA | NA |
| ENSG00000204577.11 | 2.703149 | 4.896778 | 1.052982 | 4.08058 | 4.49E-05 | 0.000428964 | ENSG00000204577 | LILRB3 | 11025 |
| ENSG00000158258.16 | 21.8972358 | 4.895285165 | 0.695112781 | 6.179264 | 6.44E-10 | 1.13E-08 | ENSG00000158258 | CLSTN2 | 64084 |
| ENSG00000197214.5 | 2.692049373 | 4.891961894 | 1.066842037 | 4.023053 | 5.74E-05 | 0.00053833 | ENSG00000197214 | NA | NA |
| ENSG00000286044.1 | 5.82582063 | 4.885147299 | 0.949270964 | 4.514146 | 6.36E-06 | 6.94E-05 | ENSG00000286044 | NA | NA |
| ENSG00000153714.6 | 1283.664038 | 4.884894751 | 0.162701153 | 26.33598 | 7.43E-153 | 5.35E-150 | ENSG00000153714 | LURAP1L | 286343 |
| ENSG00000244119.1 | 142.7538612 | 4.884674196 | 0.264382601 | 16.20634 | 4.55E-59 | 8.41E-57 | ENSG00000244119 | NA | NA |
| ENSG00000049283.18 | 3397.439849 | 4.87927342 | 0.120421468 | 35.5358 | 1.38E-276 | 2.44E-273 | ENSG00000049283 | EPN3 | 55040 |
| ENSG00000198125.13 | 2.901527 | 4.878589 | 1.235212 | 3.46385 | 0.000533 | 0.004194166 | ENSG00000198125 | MB | 4151 |
| ENSG00000200033.1 | 2.908420337 | 4.877804177 | 1.213120561 | 3.526281 | 0.000421 | 0.003384482 | ENSG00000200033 | RNU6-403 | 1.06E+08 |
| ENSG00000133048.13 | 51.06659842 | 4.872531844 | 0.602815746 | 7.087625 | 1.36E-12 | 3.07E-11 | ENSG00000133048 | CHI3L1 | 1116 |
| ENSG00000250127.1 | 2.573194668 | 4.871492938 | 1.095471282 | 3.899229 | 9.65E-05 | 0.000874409 | ENSG00000250127 | LINC0210ε | 1.02E+08 |
| ENSG00000096006.11 | 2.56013842 | 4.870524893 | 1.090404735 | 3.916459 | 8.99E-05 | 0.000817578 | ENSG00000096006 | CRISP3 | 10321 |
| ENSG00000110436.13 | 5.148266913 | 4.869940642 | 0.96388493 | 4.429928 | 9.43E-06 | 0.000100381 | ENSG00000110436 | SLC1A2 | 6506 |
| ENSG00000059804.16 | 395.9245219 | 4.865914089 | 0.535587479 | 7.964925 | 1.65E-15 | 4.72E-14 | ENSG00000059804 | SLC2A3 | 6515 |
| ENSG00000110375.3 | 1783.476621 | 4.864096357 | 0.419935243 | 10.15418 | 3.17E-24 | 1.53E-22 | ENSG00000110375 | UPK2 | 7379 |
| ENSG00000274461.2 | 7.939240293 | 4.860847993 | 0.844247948 | 5.046915 | 4.49E-07 | 5.67E-06 | ENSG00000274461 | LOC10537 | 1.05E+08 |
| ENSG00000267259.1 | 37.12301481 | 4.860322592 | 0.504057653 | 8.452054 | 2.86E-17 | 9.22E-16 | ENSG00000267259 | NA | NA |
| ENSG00000259881.1 | 2.991728023 | 4.8592683 | 1.495305839 | 2.848426 | 0.004394 | 0.027942317 | ENSG00000259881 | LOC10192 | 1.02E+08 |
| ENSG00000095321.17 | 105.913681 | 4.851491527 | 0.77670524 | 5.473752 | 4.41E-08 | 6.29E-07 | ENSG00000095321 | CRAT | 1384 |
| ENSG00000175302.5 | 3.385001 | 4.84941 | 1.036636 | 4.09923 | 4.15E-05 | 0.000397677 | ENSG00000175302 | ANKRD30I | 642460 |
| ENSG00000248208.1 | 3.112830439 | 4.843845877 | 1.191843952 | 3.56074 | 0.00037 | 0.003004312 | ENSG00000248208 | NA | NA |
| ENSG00000002726.20 | 159.3102148 | 4.841619821 | 0.353471592 | 11.99989 | 3.56E-33 | 2.78E-31 | ENSG00000002726 | AOC1 | 26 |
| ENSG00000261553.5 | 88.69398701 | 4.834191407 | 0.279705429 | 15.13804 | 9.09E-52 | 1.34E-49 | ENSG00000261553 | NA | NA |
| ENSG00000171101.13 | 3.321905 | 4.833292 | 1.13209 | 3.739361 | 0.000184 | 0.001583318 | ENSG00000171101 | SIGLEC17 | 284367 |
| ENSG00000225678.2 | 5.224179932 | 4.831105166 | 0.950303391 | 4.452373 | 8.49E-06 | 9.11E-05 | ENSG00000225678 | NA | NA |
| ENSG00000162699.5 | 2.902522101 | 4.828292031 | 1.482334533 | 2.852455 | 0.004338 | 0.02763681 | ENSG00000162699 | NA | NA |
| ENSG00000169469.9 | 31259.20104 | 4.82793633 | 0.391207788 | 10.80739 | 3.18E-27 | 1.83E-25 | ENSG00000169469 | SPRR1B | 6699 |
| ENSG00000131738.10 | 7.418249191 | 4.82690842 | 0.90281442 | 4.681924 | 2.84E-06 | 3.23E-05 | ENSG00000131738 | KRT33B | 3884 |
| ENSG00000213937.4 | 28.51869291 | 4.818576547 | 0.488300853 | 8.639298 | 5.66E-18 | 1.91E-16 | ENSG00000213937 | CLDN9 | 9080 |
| ENSG00000286511.1 | 22.72900247 | 4.817611218 | 0.5779751 | 7.29722 | 2.94E-13 | 7.03E-12 | ENSG00000286511 | NA | NA |
| ENSG00000109610.6 | 8.613846031 | 4.817343189 | 0.869956001 | 4.847766 | 1.25E-06 | 1.48E-05 | ENSG00000109610 | SOD3 | 6649 |
| ENSG00000103034.14 | 3676.773499 | 4.81714504 | 0.335385278 | 12.57403 | 2.93E-36 | 2.54E-34 | ENSG00000103034 | NDRG4 | 65009 |
| ENSG00000228198.3 | 3.054190929 | 4.811369894 | 1.148010964 | 3.668406 | 0.000244 | 0.002052297 | ENSG00000228198 | OR2M3 | 127062 |
| ENSG00000286860.1 | 2.633987615 | 4.811001051 | 1.033850508 | 4.073124 | 4.64E-05 | 0.000441986 | ENSG00000286860 | NA | NA |
| ENSG00000163060.8 | 2.627999475 | 4.808933084 | 1.022991008 | 4.11434 | 3.88E-05 | 0.000374937 | ENSG00000163060 | TEKT4 | 150483 |
| ENSG00000206043.6 | 3.942780166 | 4.805002745 | 1.164193533 | 3.611945 | 0.000304 | 0.002506404 | ENSG00000206043 | C18orf63 | 644041 |
| ENSG00000005001.10 | 2188.395809 | 4.803874933 | 0.344477901 | 12.20361 | 2.97E-34 | 2.36E-32 | ENSG00000005001 | PRSS22 | 64063 |

| | | | | | | | | | |
|---|---|---|---|---|---|---|---|---|---|
| ENSG00000136694.8 | 2.521262394 | 4.800071046 | 1.060215251 | 3.961527 | 7.45E-05 | 0.000686732 | ENSG00000136694 | IL36A | 27179 |
| ENSG00000189052.7 | 43.83209 | 4.799155 | 0.6587651 | 6.374283 | 1.84E-10 | 3.43E-09 | ENSG00000189052 | CGB5 | 93659 |
| ENSG00000258535.1 | 6.233206432 | 4.788989231 | 0.94594338 | 4.428372 | 9.49E-06 | 0.000101026 | ENSG00000258535 | NA | NA |
| ENSG00000259905.7 | 2.420360186 | 4.784143726 | 1.213275631 | 3.448634 | 0.000563 | 0.00440906 | ENSG00000259905 | PWRN1 | 791114 |
| ENSG00000124721.17 | 4.976258138 | 4.782827278 | 0.944622455 | 4.428041 | 9.51E-06 | 0.000101141 | ENSG00000124721 | DNAH8 | 1769 |
| ENSG00000286288.1 | 19.3675 | 4.78216 | 0.6013618 | 6.954482 | 3.54E-12 | 7.72E-11 | ENSG00000286288 | NA | NA |
| ENSG00000286667.1 | 39.09430868 | 4.77855282 | 0.402729506 | 10.37558 | 3.20E-25 | 1.65E-23 | ENSG00000286667 | NA | NA |
| ENSG00000116741.8 | 490.2462606 | 4.777201763 | 0.14129447 | 29.5638 | 4.37E-192 | 5.05E-189 | ENSG00000116741 | RGS2 | 5997 |
| ENSG00000148600.15 | 20.79821577 | 4.775862257 | 0.565372293 | 7.38604 | 1.51E-13 | 3.71E-12 | ENSG00000148600 | CDHR1 | 92211 |
| ENSG00000237859.2 | 2.633982 | 4.77349 | 1.048617 | 3.979996 | 6.89E-05 | 0.000638155 | ENSG00000237859 | EEF1A1P3 | 553820 |
| ENSG00000276241.1 | 2.395678567 | 4.773356168 | 1.259500183 | 3.313502 | 0.000921 | 0.006886107 | ENSG00000276241 | NA | NA |
| ENSG00000286342.1 | 70.49323143 | 4.773104159 | 0.739163392 | 5.645713 | 1.64E-08 | 2.48E-07 | ENSG00000286342 | NA | NA |
| ENSG00000251204.1 | 2.739556678 | 4.770776876 | 1.171027875 | 3.561638 | 0.000369 | 0.002994966 | ENSG00000251204 | NA | NA |
| ENSG00000205632.4 | 2.626897 | 4.770549 | 1.11871 | 3.728 | 0.000193 | 0.001649501 | ENSG00000205632 | LINC0131( | 1E+08 |
| ENSG00000157542.11 | 2.445082 | 4.760241 | 1.414796 | 2.940523 | 0.003277 | 0.02147447 | ENSG00000157542 | KCNJ6 | 3763 |
| ENSG00000250891.2 | 9.020391658 | 4.756176359 | 0.812454716 | 5.115579 | 3.13E-07 | 4.04E-06 | ENSG00000250891 | NA | NA |
| ENSG00000182578.13 | 280.0936998 | 4.754748717 | 0.224802915 | 18.48174 | 2.90E-76 | 8.12E-74 | ENSG00000182578 | CSF1R | 1436 |
| ENSG00000283142.1 | 60.92805 | 4.744582 | 0.3703967 | 11.18958 | 4.59E-29 | 3.00E-27 | ENSG00000283142 | NA | NA |
| ENSG00000182885.17 | 2.452578197 | 4.733235573 | 1.315391918 | 3.142208 | 0.001677 | 0.011778207 | ENSG00000182885 | ADGRG3 | 222487 |
| ENSG00000146250.7 | 69.52365859 | 4.728454368 | 0.376981693 | 10.95134 | 6.55E-28 | 3.93E-26 | ENSG00000146250 | PRSS35 | 167681 |
| ENSG00000258483.4 | 15.04226974 | 4.724945054 | 0.610943373 | 6.751763 | 1.46E-11 | 3.02E-10 | ENSG00000258483 | LINC0225 | 1.05E+08 |
| ENSG00000270575.1 | 2.320304668 | 4.723638591 | 1.110650006 | 3.712816 | 0.000205 | 0.001744441 | ENSG00000270575 | NA | NA |
| ENSG00000005381.7 | 538.6385394 | 4.723183657 | 0.302114291 | 13.64776 | 2.08E-42 | 2.29E-40 | ENSG00000005381 | MPO | 4353 |
| ENSG00000157445.15 | 3.837130016 | 4.721868595 | 1.1032472 | 3.736124 | 0.000187 | 0.001600732 | ENSG00000157445 | CACNA2D | 55799 |
| ENSG00000277981.1 | 2.310648942 | 4.721850901 | 1.084019813 | 3.802376 | 0.000143 | 0.001255043 | ENSG00000277981 | NA | NA |
| ENSG00000285572.2 | 2.335077595 | 4.71868594 | 1.064724976 | 3.86831 | 0.00011 | 0.000984003 | ENSG00000285572 | NA | NA |
| ENSG00000227744.4 | 168.8711806 | 4.716706859 | 0.211201182 | 19.49187 | 1.29E-84 | 4.13E-82 | ENSG00000227744 | LINC0194( | 401039 |
| ENSG00000231683.6 | 2.327151949 | 4.715171294 | 1.108531959 | 3.712271 | 0.000205 | 0.001746523 | ENSG00000231683 | LOC10192 | 1.02E+08 |
| ENSG00000287927.1 | 2.32676049 | 4.714014359 | 1.365373349 | 3.013106 | 0.002586 | 0.017444074 | ENSG00000287927 | LOC10798 | 1.08E+08 |
| ENSG00000169302.16 | 2.298671531 | 4.713868532 | 1.066109492 | 3.858767 | 0.000114 | 0.001020125 | ENSG00000169302 | STK32A | 202374 |
| ENSG00000127743.6 | 4.640879249 | 4.698032037 | 0.953612782 | 4.297375 | 1.73E-05 | 0.000175549 | ENSG00000127743 | IL17B | 27190 |
| ENSG00000109101.7 | 78.42423892 | 4.696739654 | 0.364115909 | 11.2512 | 2.28E-29 | 1.53E-27 | ENSG00000109101 | FOXN1 | 8456 |
| ENSG00000167751.13 | 12.09668 | 4.695584 | 0.7781109 | 5.263496 | 1.41E-07 | 1.92E-06 | ENSG00000167751 | KLK2 | 3817 |
| ENSG00000257194.2 | 4.526617193 | 4.689432931 | 1.094450047 | 3.736519 | 0.000187 | 0.001599254 | ENSG00000257194 | NA | NA |
| ENSG00000196460.14 | 4.565755435 | 4.687964534 | 1.305738856 | 3.130767 | 0.001744 | 0.012224262 | ENSG00000196460 | RFX8 | 731220 |
| ENSG00000251574.8 | 4.651967473 | 4.687092529 | 0.98992226 | 4.1287 | 3.65E-05 | 0.000353429 | ENSG00000251574 | LOC10537 | 1.05E+08 |
| ENSG00000096088.16 | 4.629049301 | 4.68642196 | 1.166745577 | 3.50241 | 0.000461 | 0.003673921 | ENSG00000096088 | PGC | 5225 |
| ENSG00000226608.3 | 31.8725 | 4.683805 | 0.5915249 | 6.903861 | 5.06E-12 | 1.09E-10 | ENSG00000226608 | NA | NA |
| ENSG00000132437.18 | 4.527552027 | 4.675475049 | 0.991826338 | 4.109061 | 3.97E-05 | 0.000382778 | ENSG00000132437 | DDC | 1644 |
| ENSG00000254768.6 | 2.251012391 | 4.670835215 | 1.044790871 | 3.896316 | 9.77E-05 | 0.000883784 | ENSG00000254768 | NA | NA |
| ENSG00000287331.1 | 18.53432871 | 4.670753821 | 0.684891611 | 5.943647 | 2.79E-09 | 4.58E-08 | ENSG00000287331 | LOC10537 | 1.05E+08 |
| ENSG00000225854.2 | 8.440584879 | 4.669796678 | 0.827811196 | 4.916334 | 8.82E-07 | 1.08E-05 | ENSG00000225854 | RPL34P4 | 1E+08 |

| | | | | | | | | | |
|---|---|---|---|---|---|---|---|---|---|
| ENSG00000277494.2 | 2.217555256 | 4.663355476 | 1.090682877 | 3.725515 | 0.000195 | 0.001664766 | ENSG00000277494 | GPIHBP1 | 338328 |
| ENSG00000268100.1 | 2.226885 | 4.660662 | 1.067723 | 3.803104 | 0.000143 | 0.001252184 | ENSG00000268100 | NA | NA |
| ENSG00000253510.1 | 2.366436528 | 4.660441053 | 1.087247384 | 3.734606 | 0.000188 | 0.001609899 | ENSG00000253510 | NA | NA |
| ENSG00000215899.3 | 2.210115587 | 4.659209149 | 1.153367591 | 3.519441 | 0.000432 | 0.003462519 | ENSG00000215899 | LOC44188 | 441880 |
| ENSG00000178146.9 | 2.453002 | 4.658714 | 1.336264 | 3.037359 | 0.002387 | 0.01620411 | ENSG00000178146 | NA | NA |
| ENSG00000282048.1 | 2.221205581 | 4.658275014 | 1.349638686 | 3.006934 | 0.002639 | 0.017758993 | ENSG00000282048 | NA | NA |
| ENSG00000186832.9 | 94018.80973 | 4.655909871 | 0.362995924 | 11.17343 | 5.50E-29 | 3.56E-27 | ENSG00000186832 | KRT16 | 3868 |
| ENSG00000164746.14 | 112.3040672 | 4.650356744 | 0.310431813 | 13.04749 | 6.57E-39 | 6.31E-37 | ENSG00000164746 | C7orf57 | 136288 |
| ENSG00000281491.1 | 7.820731848 | 4.648908905 | 0.91699849 | 4.415393 | 1.01E-05 | 0.000106685 | ENSG00000281491 | DNAJB5-D | 1.02E+08 |
| ENSG00000286772.1 | 2.342712 | 4.648024138 | 1.251891541 | 3.233526 | 0.001223 | 0.008884083 | ENSG00000286772 | NA | NA |
| ENSG00000164867.11 | 202.0641905 | 4.645684436 | 0.258429136 | 15.65491 | 3.08E-55 | 5.09E-53 | ENSG00000164867 | NOS3 | 4846 |
| ENSG00000258791.9 | 425.6308937 | 4.645011069 | 0.278625095 | 14.51776 | 9.35E-48 | 1.23E-45 | ENSG00000258791 | LINC0052( | 645687 |
| ENSG00000285634.2 | 2.260679228 | 4.637824962 | 1.090232613 | 3.703636 | 0.000213 | 0.001800764 | ENSG00000285634 | NA | NA |
| ENSG00000254966.1 | 5.806315043 | 4.635652323 | 0.874820091 | 4.613123 | 3.97E-06 | 4.43E-05 | ENSG00000254966 | NA | NA |
| ENSG00000276649.1 | 46.35939 | 4.633834 | 0.3838669 | 10.50842 | 7.90E-26 | 4.22E-24 | ENSG00000276649 | NA | NA |
| ENSG00000265313.7 | 2.199020225 | 4.632591061 | 1.11915193 | 3.603256 | 0.000314 | 0.002582133 | ENSG00000265313 | NA | NA |
| ENSG00000139151.15 | 4.412096576 | 4.629586162 | 0.962485812 | 4.186645 | 2.83E-05 | 0.000278115 | ENSG00000139151 | PLCZ1 | 89869 |
| ENSG00000266952.2 | 2.673960734 | 4.628496965 | 1.167901699 | 3.449346 | 0.000562 | 0.004398746 | ENSG00000266952 | LINC0153٤ | 400654 |
| ENSG00000064787.13 | 4.375548 | 4.626713 | 1.021207 | 3.943092 | 8.04E-05 | 0.00073791 | ENSG00000064787 | BCAS1 | 8537 |
| ENSG00000243478.9 | 19.10665836 | 4.626251692 | 0.556127664 | 7.239798 | 4.49E-13 | 1.06E-11 | ENSG00000243478 | AOX2P | 344454 |
| ENSG00000228211.2 | 2.677074565 | 4.62480651 | 1.216148562 | 3.309469 | 0.000935 | 0.006976275 | ENSG00000228211 | NA | NA |
| ENSG00000203782.6 | 13.64347349 | 4.620311007 | 0.786055005 | 5.114542 | 3.15E-07 | 4.06E-06 | ENSG00000203782 | LORICRIN | 4014 |
| ENSG00000261268.1 | 7.758336385 | 4.619308735 | 0.83399922 | 4.81932 | 1.44E-06 | 1.70E-05 | ENSG00000261268 | NA | NA |
| ENSG00000146809.13 | 4.396616065 | 4.613962178 | 0.950654199 | 4.222316 | 2.42E-05 | 0.00024002 | ENSG00000146809 | ASB15 | 142685 |
| ENSG00000172602.11 | 63.38975801 | 4.613550609 | 0.428771799 | 9.360575 | 7.93E-21 | 3.21E-19 | ENSG00000172602 | RND1 | 27289 |
| ENSG00000185056.10 | 2.271744711 | 4.613478278 | 1.082162216 | 3.708758 | 0.000208 | 0.001768669 | ENSG00000185056 | C5orf47 | 133491 |
| ENSG00000050628.20 | 2.14319278 | 4.612824625 | 1.110576485 | 3.613281 | 0.000302 | 0.002495066 | ENSG00000050628 | PTGER3 | 5733 |
| ENSG00000240023.1 | 2.860011953 | 4.60863209 | 1.181271568 | 3.393489 | 0.00069 | 0.005311187 | ENSG00000240023 | RPLP0P3 | 122589 |
| ENSG00000230284.1 | 2.463520855 | 4.606613717 | 1.156748661 | 3.463686 | 0.000533 | 0.004195495 | ENSG00000230284 | NA | NA |
| ENSG00000171954.13 | 43.73278338 | 4.603064261 | 0.447254039 | 8.950314 | 3.54E-19 | 1.29E-17 | ENSG00000171954 | CYP4F22 | 126410 |
| ENSG00000279243.3 | 2.156894257 | 4.602355644 | 1.256992695 | 3.184072 | 0.001452 | 0.010350771 | ENSG00000279243 | NA | NA |
| ENSG00000256422.6 | 2.18985854 | 4.602229384 | 1.248397047 | 3.205895 | 0.001346 | 0.009679918 | ENSG00000256422 | LINC0255ž | 1.03E+08 |
| ENSG00000242593.7 | 2.324574038 | 4.597407349 | 1.128699907 | 3.541603 | 0.000398 | 0.003210325 | ENSG00000242593 | NA | NA |
| ENSG00000249884.8 | 36.99770179 | 4.596131866 | 0.401815063 | 9.945202 | 2.65E-23 | 1.23E-21 | ENSG00000249884 | RNF103-C | 1.01E+08 |
| ENSG00000167552.14 | 3974.61736 | 4.593386402 | 0.103558356 | 38.5617 | 0 | 0 | ENSG00000167552 | TUBA1A | 7846 |
| ENSG00000105880.7 | 12.11185666 | 4.593086595 | 0.678477528 | 5.885363 | 3.97E-09 | 6.40E-08 | ENSG00000105880 | DLX5 | 1749 |
| ENSG00000271815.1 | 8.018871224 | 4.592299102 | 0.820934486 | 4.863115 | 1.16E-06 | 1.38E-05 | ENSG00000271815 | NA | NA |
| ENSG00000138678.11 | 2303.662694 | 4.588373573 | 0.212604215 | 18.75962 | 1.62E-78 | 4.68E-76 | ENSG00000138678 | GPAT3 | 84803 |
| ENSG00000272256.1 | 3.025297989 | 4.585517863 | 1.092332507 | 3.648631 | 0.000264 | 0.002198832 | ENSG00000272256 | NA | NA |
| ENSG00000233030.2 | 13.79607009 | 4.585291815 | 0.811989104 | 4.908061 | 9.20E-07 | 1.12E-05 | ENSG00000233030 | NA | NA |
| ENSG00000223394.2 | 2.403341732 | 4.584507061 | 1.237090328 | 3.22087 | 0.001278 | 0.009245512 | ENSG00000223394 | TRBV29OI | 6958 |
| ENSG00000266278.1 | 14.54312658 | 4.582323951 | 0.660515095 | 6.029119 | 1.65E-09 | 2.77E-08 | ENSG00000266278 | LINC0191( | 1.01E+08 |

| | | | | | | | | | | |
|---|---|---|---|---|---|---|---|---|---|---|
| ENSG00000177243.3 | 2.565831293 | 4.580999679 | 1.257374376 | 3.166121 | 0.001545 | 0.010940469 | ENSG00000177243 | DEFB103E | | 55894 |
| ENSG00000129455.15 | 700.4175 | 4.578877 | 0.2962939 | 13.42882 | 4.10E-41 | 4.28E-39 | ENSG00000129455 | KLK8 | | 11202 |
| ENSG00000052344.16 | 4095.858624 | 4.567046459 | 0.179749609 | 22.06985 | 6.16E-108 | 2.74E-105 | ENSG00000052344 | PRSS8 | | 5652 |
| ENSG00000236013.7 | 2.178745543 | 4.566680556 | 1.081483217 | 3.667815 | 0.000245 | 0.002055742 | ENSG00000236013 | LOC10050 | | 1.01E+08 |
| ENSG00000286125.1 | 2.122425 | 4.566545 | 1.120215 | 3.540879 | 0.000399 | 0.003217201 | ENSG00000286125 | ZIM2-AS1 | | 1.02E+08 |
| ENSG00000160360.13 | 2435.824406 | 4.566479062 | 0.256529567 | 15.46207 | 6.26E-54 | 9.85E-52 | ENSG00000160360 | GPSM1 | | 26086 |
| ENSG00000286329.1 | 2.079113004 | 4.564362716 | 1.13719097 | 3.486101 | 0.00049 | 0.003886762 | ENSG00000286329 | LOC10537 | | 1.05E+08 |
| ENSG00000224568.2 | 2.089094149 | 4.562244246 | 1.281091293 | 3.092866 | 0.001982 | 0.013682572 | ENSG00000224568 | LINC0188( | | 1.05E+08 |
| ENSG00000245164.8 | 2.069955013 | 4.559918117 | 1.213911085 | 3.262115 | 0.001106 | 0.008121296 | ENSG00000245164 | LINC0086 | | 1E+08 |
| ENSG00000184313.20 | 2.082826327 | 4.557628089 | 1.104873222 | 3.581975 | 0.000341 | 0.002786458 | ENSG00000184313 | MROH7 | | 374977 |
| ENSG00000248954.1 | 5.996900518 | 4.555095568 | 0.927004956 | 4.266531 | 1.99E-05 | 0.000199827 | ENSG00000248954 | NA | NA | |
| ENSG00000240535.8 | 2.480522728 | 4.553270452 | 1.213510307 | 3.257715 | 0.001123 | 0.008241789 | ENSG00000240535 | NA | NA | |
| ENSG00000261645.7 | 2.066215779 | 4.547945582 | 1.215128111 | 3.248995 | 0.001158 | 0.008460987 | ENSG00000261645 | DISC1FP1 | | 1.02E+08 |
| ENSG00000250230.3 | 2.065891355 | 4.547609547 | 1.173152964 | 3.364957 | 0.000766 | 0.005834701 | ENSG00000250230 | LOC10192 | | 1.02E+08 |
| ENSG00000267420.1 | 4.379142088 | 4.545011744 | 1.057454619 | 3.730668 | 0.000191 | 0.001633703 | ENSG00000267420 | NA | NA | |
| ENSG00000084628.10 | 2.08430199 | 4.543010475 | 1.318531621 | 2.990456 | 0.002786 | 0.018623465 | ENSG00000084628 | NKAIN1 | | 79570 |
| ENSG00000230316.7 | 15.92539158 | 4.540158006 | 0.811009877 | 4.858335 | 1.18E-06 | 1.42E-05 | ENSG00000230316 | FEZF1-AS | | 154860 |
| ENSG00000243323.7 | 2.048026335 | 4.539576555 | 1.111377762 | 3.544768 | 0.000393 | 0.003175878 | ENSG00000243323 | PTPRVP | | 148713 |
| ENSG00000157399.14 | 2.081472 | 4.538891 | 1.201081 | 3.279456 | 0.00104 | 0.007684952 | ENSG00000157399 | ARSL | | 415 |
| ENSG00000287754.1 | 2.081353 | 4.53878 | 1.316794 | 2.991189 | 0.002779 | 0.01858349 | ENSG00000287754 | NA | NA | |
| ENSG00000232368.1 | 4.670514 | 4.535915 | 0.9112107 | 4.319435 | 1.56E-05 | 0.000160049 | ENSG00000232368 | NA | NA | |
| ENSG00000095932.6 | 2.067182461 | 4.531183875 | 1.314153428 | 2.991419 | 0.002777 | 0.018574126 | ENSG00000095932 | SMIM24 | | 284422 |
| ENSG00000132821.12 | 18.60228 | 4.526997 | 0.5581739 | 7.035437 | 1.99E-12 | 4.43E-11 | ENSG00000132821 | VSTM2L | | 128434 |
| ENSG00000111405.9 | 19.076911 | 4.525147311 | 0.694705459 | 5.650088 | 1.60E-08 | 2.42E-07 | ENSG00000111405 | ENDOU | | 8909 |
| ENSG00000115488.3 | 2.602328905 | 4.519049468 | 1.106723296 | 3.541129 | 0.000398 | 0.003215132 | ENSG00000115488 | NEU2 | | 4759 |
| ENSG00000148935.11 | 37.4137064 | 4.517989521 | 0.379705868 | 10.31849 | 5.81E-25 | 2.94E-23 | ENSG00000148935 | GAS2 | | 2620 |
| ENSG00000267069.1 | 47.75111574 | 4.515182448 | 0.345854279 | 11.32032 | 1.04E-29 | 7.05E-28 | ENSG00000267069 | NA | NA | |
| ENSG00000249661.1 | 20.16910988 | 4.514718328 | 0.58715276 | 6.667291 | 2.61E-11 | 5.25E-10 | ENSG00000249661 | NA | NA | |
| ENSG00000228863.8 | 2.032371223 | 4.51443988 | 1.1626156 | 3.366925 | 0.00076 | 0.005799863 | ENSG00000228863 | NA | NA | |
| ENSG00000267938.1 | 3.066872807 | 4.513657925 | 1.014825784 | 3.856483 | 0.000115 | 0.00102832 | ENSG00000267938 | NA | NA | |
| ENSG00000124107.5 | 42995.19 | 4.509716 | 0.327881 | 11.92419 | 8.85E-33 | 6.83E-31 | ENSG00000124107 | SLPI | | 6590 |
| ENSG00000200579.1 | 6.101533906 | 4.5075095 | 0.900583318 | 4.338865 | 1.43E-05 | 0.000147783 | ENSG00000200579 | NA | NA | |
| ENSG00000236049.1 | 4.062867811 | 4.506357682 | 1.025630051 | 3.808739 | 0.00014 | 0.001225717 | ENSG00000236049 | LINC0192( | | 1.02E+08 |
| ENSG00000232359.1 | 4.964947067 | 4.505916832 | 0.927163552 | 4.212759 | 2.52E-05 | 0.000249942 | ENSG00000232359 | NA | NA | |
| ENSG00000175264.8 | 15.3434934 | 4.505506707 | 0.699854011 | 5.580459 | 2.40E-08 | 3.56E-07 | ENSG00000175264 | CHST1 | | 8534 |
| ENSG00000041515.16 | 716.908255 | 4.495850708 | 0.292747986 | 13.30787 | 2.08E-40 | 2.11E-38 | ENSG00000041515 | MYO16 | | 23026 |
| ENSG00000254066.2 | 1.974196315 | 4.49399723 | 1.161093194 | 3.353734 | 0.000797 | 0.0060575 | ENSG00000254066 | NA | NA | |
| ENSG00000161640.15 | 2.201721 | 4.491756 | 1.135952 | 3.425988 | 0.000613 | 0.004760012 | ENSG00000161640 | SIGLEC11 | | 114132 |
| ENSG00000131668.14 | 16.05259861 | 4.490554679 | 0.684220266 | 5.686114 | 1.30E-08 | 1.98E-07 | ENSG00000131668 | BARX1 | | 56033 |
| ENSG00000130518.17 | 71.8722 | 4.490542 | 0.3915381 | 9.936558 | 2.89E-23 | 1.33E-21 | ENSG00000130518 | IQCN | | 80726 |
| ENSG00000174125.8 | 96.98544242 | 4.488351339 | 0.361229775 | 10.7642 | 5.08E-27 | 2.87E-25 | ENSG00000174125 | TLR1 | | 7096 |
| ENSG00000280268.1 | 1.960914775 | 4.486409074 | 1.165280189 | 3.335171 | 0.000852 | 0.006425373 | ENSG00000280268 | NA | NA | |

| | | | | | | | | | |
|---|---|---|---|---|---|---|---|---|---|
| ENSG00000151117.9 | 804.3521107 | 4.486267255 | 0.215223939 | 18.05685 | 6.97E-73 | 1.80E-70 | ENSG00000151117 | TMEM86A | 144110 |
| ENSG00000007062.11 | 3.936167143 | 4.485181923 | 1.125454274 | 3.452101 | 0.000556 | 0.004363053 | ENSG00000007062 | PROM1 | 8842 |
| ENSG00000111249.14 | 1.963189592 | 4.483265171 | 1.113818886 | 3.486442 | 0.000489 | 0.003882966 | ENSG00000111249 | CUX2 | 23316 |
| ENSG00000039987.6 | 8.899550593 | 4.481411707 | 0.829617186 | 4.678558 | 2.89E-06 | 3.28E-05 | ENSG00000039987 | BEST2 | 54831 |
| ENSG00000236453.5 | 23.98108559 | 4.48046071 | 0.591516502 | 6.56019 | 5.37E-11 | 1.05E-09 | ENSG00000236453 | NA | NA |
| ENSG00000163209.15 | 2427.69107 | 4.480288289 | 0.387580602 | 10.01156 | 1.36E-23 | 6.34E-22 | ENSG00000163209 | SPRR3 | 6707 |
| ENSG00000172900.12 | 1.955933215 | 4.478737616 | 1.403812604 | 2.763002 | 0.005727 | 0.035228843 | ENSG00000172900 | ACTE1P | 399923 |
| ENSG00000143816.8 | 678.5099846 | 4.478689941 | 0.256235888 | 15.13718 | 9.21E-52 | 1.35E-49 | ENSG00000143816 | WNT9A | 7483 |
| ENSG00000184108.7 | 2.310402455 | 4.477846267 | 1.22458906 | 3.166651 | 0.001542 | 0.01092694 | ENSG00000184108 | TRIML1 | 339976 |
| ENSG00000101605.12 | 58.96856447 | 4.474088387 | 0.45218552 | 8.567476 | 1.06E-17 | 3.50E-16 | ENSG00000101605 | MYOM1 | 8736 |
| ENSG00000279676.1 | 10.38844 | 4.473851 | 0.7751437 | 4.997591 | 5.81E-07 | 7.25E-06 | ENSG00000279676 | NA | NA |
| ENSG00000287495.1 | 16.47583556 | 4.472900914 | 0.569627438 | 6.799007 | 1.05E-11 | 2.21E-10 | ENSG00000287495 | NA | NA |
| ENSG00000184012.12 | 94.655 | 4.471569 | 0.2534924 | 15.27292 | 1.16E-52 | 1.78E-50 | ENSG00000184012 | TMPRSS2 | 7113 |
| ENSG00000225746.12 | 1.96163343 | 4.468283129 | 1.181968904 | 3.272745 | 0.001065 | 0.007850131 | ENSG00000225746 | NA | NA |
| ENSG00000251169.2 | 139.3866053 | 4.467594819 | 0.290267371 | 13.32425 | 1.67E-40 | 1.70E-38 | ENSG00000251169 | LINC0184З | 1.02E+08 |
| ENSG00000108551.5 | 51.23299467 | 4.466609732 | 0.452914183 | 8.53718 | 1.38E-17 | 4.51E-16 | ENSG00000108551 | RASD1 | 51655 |
| ENSG00000227742.2 | 3.927432448 | 4.46322151 | 1.01181956 | 3.818093 | 0.000134 | 0.001185927 | ENSG00000227742 | NA | NA |
| ENSG00000205325.2 | 23.84626426 | 4.460281809 | 0.573047041 | 6.736413 | 1.62E-11 | 3.34E-10 | ENSG00000205325 | NA | NA |
| ENSG00000287682.1 | 1.986463389 | 4.458340337 | 1.310308012 | 2.944606 | 0.003234 | 0.021250714 | ENSG00000287682 | NA | NA |
| ENSG00000287865.1 | 2.119108991 | 4.456078419 | 1.258101214 | 3.064999 | 0.002177 | 0.014923612 | ENSG00000287865 | NA | NA |
| ENSG00000182965.6 | 2.22583151 | 4.448871085 | 1.2749982 | 3.018727 | 0.002538 | 0.017142854 | ENSG00000182965 | NA | NA |
| ENSG00000273225.4 | 2.108832036 | 4.444447382 | 1.2164321 | 3.160429 | 0.001575 | 0.011130437 | ENSG00000273225 | NA | NA |
| ENSG00000203356.2 | 7.262095877 | 4.443590522 | 0.822850728 | 4.671067 | 3.00E-06 | 3.39E-05 | ENSG00000203356 | LINC0156[illegible] | 1.05E+08 |
| ENSG00000085831.15 | 500.4007628 | 4.442484281 | 0.310219095 | 12.38636 | 3.10E-35 | 2.59E-33 | ENSG00000085831 | TTC39A | 22996 |
| ENSG00000229009.7 | 5.358188415 | 4.441079052 | 1.065168325 | 3.606077 | 0.000311 | 0.002558979 | ENSG00000229009 | TMPRSS1 | 644759 |
| ENSG00000066032.18 | 5.661719153 | 4.434554137 | 0.900975579 | 4.256002 | 2.08E-05 | 0.000208915 | ENSG00000066032 | CTNNA2 | 1496 |
| ENSG00000237787.5 | 3.849752452 | 4.432246105 | 0.976432879 | 3.924741 | 8.68E-05 | 0.000792391 | ENSG00000237787 | LINC0287[illegible] | 152118 |
| ENSG00000105695.15 | 36.39814 | 4.428139 | 0.5693696 | 6.723469 | 1.77E-11 | 3.64E-10 | ENSG00000105695 | MAG | 4099 |
| ENSG00000261026.1 | 28.04651476 | 4.425411261 | 0.429496923 | 8.906726 | 5.26E-19 | 1.89E-17 | ENSG00000261026 | NA | NA |
| ENSG00000270111.3 | 1.897606671 | 4.424237577 | 1.421287953 | 2.690685 | 0.007131 | 0.042706346 | ENSG00000270111 | NA | NA |
| ENSG00000280285.1 | 1.900024006 | 4.424079085 | 1.373166216 | 2.784862 | 0.005355 | 0.033231224 | ENSG00000280285 | NA | NA |
| ENSG00000196167.10 | 2.088280567 | 4.424067712 | 1.297525341 | 2.947201 | 0.003207 | 0.021093999 | ENSG00000196167 | COLCA1 | 399948 |
| ENSG00000144130.11 | 17.80554475 | 4.420632089 | 0.544815304 | 7.012711 | 2.34E-12 | 5.17E-11 | ENSG00000144130 | NT5DC4 | 284958 |
| ENSG00000215875.4 | 3.817881188 | 4.420275974 | 1.008129085 | 3.789471 | 0.000151 | 0.001318596 | ENSG00000215875 | NA | NA |
| ENSG00000142609.18 | 36.75633085 | 4.420115527 | 0.465405771 | 8.20814 | 2.25E-16 | 6.74E-15 | ENSG00000142609 | CFAP74 | 85452 |
| ENSG00000166589.13 | 263.6799703 | 4.412068087 | 0.237976587 | 16.01867 | 9.47E-58 | 1.69E-55 | ENSG00000166589 | CDH16 | 1014 |
| ENSG00000281721.1 | 1.876921219 | 4.411863831 | 1.139425775 | 3.345425 | 0.000822 | 0.006225932 | ENSG00000281721 | NA | NA |
| ENSG00000164744.13 | 201.4733111 | 4.406847659 | 0.363572177 | 10.47068 | 1.18E-25 | 6.25E-24 | ENSG00000164744 | SUN3 | 256979 |
| ENSG00000236700.6 | 3.79481208 | 4.405798799 | 1.007532285 | 3.777347 | 0.000159 | 0.001378119 | ENSG00000236700 | LINC0101[illegible] | 154092 |
| ENSG00000138311.17 | 247.5211265 | 4.404756743 | 0.239750578 | 15.86965 | 1.03E-56 | 1.77E-54 | ENSG00000138311 | ZNF365 | 22891 |
| ENSG00000196549.10 | 10.82449854 | 4.404630477 | 0.708656551 | 5.368793 | 7.93E-08 | 1.10E-06 | ENSG00000196549 | MME | 4311 |
| ENSG00000144010.9 | 2.024766831 | 4.401965392 | 1.339557792 | 2.838224 | 0.004537 | 0.028686818 | ENSG00000144010 | TRIM43B | 653192 |

| | | | | | | | | | |
|---|---|---|---|---|---|---|---|---|---|
| ENSG00000229291.1 | 5.448672944 | 4.399901078 | 1.048422634 | 3.624398 | 0.00029 | 0.002400845 | ENSG00000229291 | NA | NA |
| ENSG00000164256.10 | 3.220883164 | 4.39710282 | 1.086668992 | 3.494259 | 0.000475 | 0.003781165 | ENSG00000164256 | PRDM9 | 56979 |
| ENSG00000237596.7 | 30.19951023 | 4.389986234 | 0.416175512 | 9.106702 | 8.49E-20 | 3.20E-18 | ENSG00000237596 | LOC64413 | 644135 |
| ENSG00000127831.10 | 1.928993706 | 4.389745576 | 1.16401182 | 3.255762 | 0.001131 | 0.008291493 | ENSG00000127831 | VIL1 | 7429 |
| ENSG00000176194.17 | 1.924392633 | 4.388251284 | 1.335038332 | 2.83756 | 0.004546 | 0.028734102 | ENSG00000176194 | CIDEA | 1149 |
| ENSG00000170961.7 | 662.59247 | 4.387572217 | 0.212794571 | 17.7992 | 7.17E-71 | 1.78E-68 | ENSG00000170961 | HAS2 | 3037 |
| ENSG00000170608.3 | 7.4491 | 4.386422 | 0.7643513 | 4.953772 | 7.28E-07 | 8.98E-06 | ENSG00000170608 | FOXA3 | 3171 |
| ENSG00000213022.6 | 3.374443 | 4.384833 | 0.9929891 | 3.811556 | 0.000138 | 0.001212921 | ENSG00000213022 | KLK9 | 284366 |
| ENSG00000145936.9 | 1.865842982 | 4.384258234 | 1.359058845 | 2.78447 | 0.005362 | 0.033263741 | ENSG00000145936 | KCNMB1 | 3779 |
| ENSG00000254544.1 | 1.917281216 | 4.383409933 | 1.39096387 | 2.719992 | 0.006528 | 0.039526489 | ENSG00000254544 | PCNAP4 | 390102 |
| ENSG00000228358.6 | 25.93735317 | 4.379532188 | 0.48071262 | 7.862353 | 3.77E-15 | 1.05E-13 | ENSG00000228358 | LINC0226 | 1.05E+08 |
| ENSG00000237254.2 | 1.859007419 | 4.379511506 | 1.256144114 | 3.00882 | 0.002623 | 0.017671487 | ENSG00000237254 | TRBV30 | 28557 |
| ENSG00000204361.9 | 54.4654558 | 4.37935116 | 0.330995184 | 11.41815 | 3.39E-30 | 2.35E-28 | ENSG00000204361 | NXPE2 | 120406 |
| ENSG00000113303.11 | 2.990020326 | 4.378891807 | 1.048014351 | 3.605763 | 0.000311 | 0.002559697 | ENSG00000113303 | BTNL8 | 79908 |
| ENSG00000198576.4 | 615.7754723 | 4.378634849 | 0.465054852 | 8.125138 | 4.47E-16 | 1.32E-14 | ENSG00000198576 | ARC | 23237 |
| ENSG00000230790.2 | 2.239664663 | 4.376641176 | 1.218678739 | 3.098964 | 0.001942 | 0.013438941 | ENSG00000230790 | NA | NA |
| ENSG00000287000.1 | 3.667870324 | 4.3694712 | 1.034160516 | 3.644958 | 0.000267 | 0.002229777 | ENSG00000287000 | NA | NA |
| ENSG00000166828.3 | 52.56910195 | 4.368733616 | 0.583247322 | 6.461639 | 1.04E-10 | 1.97E-09 | ENSG00000166828 | SCNN1G | 6340 |
| ENSG00000056291.17 | 2.78847962 | 4.368450632 | 1.158891192 | 3.251773 | 0.001147 | 0.008385655 | ENSG00000056291 | NPFFR2 | 10886 |
| ENSG00000251478.1 | 2.229381963 | 4.367255029 | 1.311375701 | 2.87275 | 0.004069 | 0.026084528 | ENSG00000251478 | TCP1P2 | 6952 |
| ENSG00000286460.1 | 1.943745208 | 4.359825762 | 1.277255654 | 2.943675 | 0.003243 | 0.021293675 | ENSG00000286460 | NA | NA |
| ENSG00000120129.6 | 6448.034293 | 4.359802972 | 0.210014198 | 17.90261 | 1.13E-71 | 2.85E-69 | ENSG00000120129 | DUSP1 | 1843 |
| ENSG00000270882.2 | 22.4932307 | 4.358714963 | 0.516535822 | 7.276775 | 3.42E-13 | 8.13E-12 | ENSG00000270882 | H4C14 | 8370 |
| ENSG00000286141.1 | 40.3265822 | 4.354975815 | 0.576719141 | 6.510926 | 7.47E-11 | 1.44E-09 | ENSG00000286141 | NA | NA |
| ENSG00000112936.18 | 1.9363643 | 4.353401422 | 1.300895665 | 2.885244 | 0.003911 | 0.025162255 | ENSG00000112936 | C7 | 730 |
| ENSG00000263648.1 | 2.032429261 | 4.351431379 | 1.370871837 | 2.73653 | 0.006209 | 0.037825784 | ENSG00000263648 | NA | NA |
| ENSG00000280139.1 | 2.03177467 | 4.350722546 | 1.221093843 | 3.071609 | 0.002129 | 0.014619686 | ENSG00000280139 | NA | NA |
| ENSG00000266560.5 | 1.814968 | 4.349874 | 1.217031 | 3.081167 | 0.002062 | 0.01418771 | ENSG00000266560 | MAGEA10 | 1.01E+08 |
| ENSG00000228328.2 | 2.033777 | 4.347414 | 1.287349 | 2.910954 | 0.003603 | 0.02337385 | ENSG00000228328 | NA | NA |
| ENSG00000156194.18 | 1.81023943 | 4.346183479 | 1.246261956 | 3.005936 | 0.002648 | 0.017808413 | ENSG00000156194 | PPEF2 | 5470 |
| ENSG00000185792.9 | 1.921722 | 4.345803 | 1.386611 | 2.701409 | 0.006905 | 0.04154952 | ENSG00000185792 | NLRP9 | 338321 |
| ENSG00000267686.2 | 5.827272713 | 4.343394858 | 0.85442321 | 4.381195 | 1.18E-05 | 0.000123417 | ENSG00000267686 | NA | NA |
| ENSG00000253903.2 | 1.844921357 | 4.339103339 | 1.419856851 | 2.633437 | 0.008453 | 0.049499325 | ENSG00000253903 | NA | NA |
| ENSG00000174640.13 | 223.4313835 | 4.33774445 | 0.532974631 | 7.012988 | 2.33E-12 | 5.17E-11 | ENSG00000174640 | SLCO2A1 | 6578 |
| ENSG00000236790.6 | 3.791787939 | 4.337539878 | 1.315278778 | 2.841633 | 0.004488 | 0.028442698 | ENSG00000236790 | LINC0029 | 339789 |
| ENSG00000159961.2 | 1.829782913 | 4.333961352 | 1.336979703 | 2.792833 | 0.005225 | 0.03249913 | ENSG00000159961 | OR3A3 | 8392 |
| ENSG00000220008.3 | 1.835172692 | 4.333491806 | 1.139073957 | 3.277655 | 0.001047 | 0.007729802 | ENSG00000220008 | LINGO3 | 645191 |
| ENSG00000196542.8 | 23.47028285 | 4.330013705 | 0.440112893 | 8.475129 | 2.35E-17 | 7.61E-16 | ENSG00000196542 | SPTSSB | 165679 |
| ENSG00000268041.2 | 6.82461 | 4.328568 | 0.8219886 | 4.536034 | 5.73E-06 | 6.29E-05 | ENSG00000268041 | ERFL | 390937 |
| ENSG00000261543.1 | 2.130689065 | 4.322661464 | 1.183996621 | 3.144149 | 0.001666 | 0.011715844 | ENSG00000261543 | NA | NA |
| ENSG00000265125.5 | 2.009451862 | 4.319903916 | 1.230546782 | 3.022968 | 0.002503 | 0.016930162 | ENSG00000265125 | NA | NA |
| ENSG00000115648.14 | 19.38410466 | 4.316921727 | 0.488454077 | 7.609562 | 2.75E-14 | 7.12E-13 | ENSG00000115648 | MLPH | 79083 |

| | | | | | | | | | |
|---|---|---|---|---|---|---|---|---|---|
| ENSG00000273472.1 | 68.71649428 | 4.315842936 | 0.403991534 | 9.197824 | 3.65E-20 | 1.41E-18 | ENSG00000273472 | NA | NA |
| ENSG00000177238.14 | 7.889484738 | 4.307114236 | 0.875501031 | 4.234277 | 2.29E-05 | 0.000228704 | ENSG00000177238 | TRIM72 | 493829 |
| ENSG00000283422.1 | 1.944343435 | 4.305254903 | 1.241254624 | 2.985088 | 0.002835 | 0.018910645 | ENSG00000283422 | NA | NA |
| ENSG00000226358.1 | 1.935446494 | 4.300124367 | 1.202237847 | 3.077697 | 0.002086 | 0.014339077 | ENSG00000226358 | KRT8P38 | 1E+08 |
| ENSG00000242220.7 | 23.53481 | 4.299125 | 0.5792884 | 6.385635 | 1.71E-10 | 3.20E-09 | ENSG00000242220 | TCP10L | 140290 |
| ENSG00000232489.1 | 3.961628923 | 4.298924275 | 0.992846996 | 3.725573 | 0.000195 | 0.001664766 | ENSG00000232489 | NA | NA |
| ENSG00000078295.16 | 1.727572871 | 4.298684616 | 1.327537257 | 2.786125 | 0.005334 | 0.033117515 | ENSG00000078295 | ADCY2 | 108 |
| ENSG00000171431.3 | 3.080050367 | 4.297494426 | 1.233714467 | 2.997042 | 0.002726 | 0.018257863 | ENSG00000171431 | KRT20 | 54474 |
| ENSG00000287603.1 | 10.15659 | 4.296902 | 0.7563505 | 4.887816 | 1.02E-06 | 1.23E-05 | ENSG00000287603 | NA | NA |
| ENSG00000272079.2 | 65.80707844 | 4.294633464 | 0.347141165 | 10.64303 | 1.88E-26 | 1.02E-24 | ENSG00000272079 | NA | NA |
| ENSG00000021645.18 | 2.473973479 | 4.293281705 | 1.043516402 | 3.539266 | 0.000401 | 0.003234967 | ENSG00000021645 | NRXN3 | 9369 |
| ENSG00000240710.1 | 1.717657497 | 4.290813058 | 1.184985743 | 3.114648 | 0.001842 | 0.012827928 | ENSG00000240710 | NA | NA |
| ENSG00000163207.7 | 2372.1391 | 4.287774321 | 0.335508972 | 10.99158 | 4.20E-28 | 2.57E-26 | ENSG00000163207 | IVL | 3713 |
| ENSG00000188508.11 | 2900.031 | 4.286556 | 0.3994981 | 9.227968 | 2.76E-20 | 1.07E-18 | ENSG00000188508 | KRTDAP | 388533 |
| ENSG00000271109.2 | 16.61972 | 4.28316 | 0.5399725 | 6.821014 | 9.04E-12 | 1.91E-10 | ENSG00000271109 | NA | NA |
| ENSG00000261235.1 | 41.58421603 | 4.277686043 | 0.436331508 | 8.428651 | 3.50E-17 | 1.12E-15 | ENSG00000261235 | NA | NA |
| ENSG00000129988.6 | 1.696936 | 4.276871 | 1.346211 | 2.731274 | 0.006309 | 0.03833786 | ENSG00000129988 | LBP | 3929 |
| ENSG00000203446.2 | 1.716146076 | 4.276763713 | 1.204409736 | 3.052752 | 0.002268 | 0.01548649 | ENSG00000203446 | NA | NA |
| ENSG00000237166.2 | 49.69263075 | 4.276750834 | 0.385824173 | 9.529602 | 1.58E-21 | 6.59E-20 | ENSG00000237166 | NA | NA |
| ENSG00000135144.7 | 27.88899248 | 4.276544146 | 1.168322712 | 3.146857 | 0.00165 | 0.01161402 | ENSG00000135144 | DTX1 | 1840 |
| ENSG00000170290.4 | 11.91214669 | 4.27445018 | 0.924686287 | 3.973726 | 7.08E-05 | 0.000654059 | ENSG00000170290 | SLN | 6588 |
| ENSG00000278869.1 | 1.742138 | 4.273862 | 1.316727 | 2.790146 | 0.005268 | 0.03273188 | ENSG00000278869 | NA | NA |
| ENSG00000149654.10 | 1.708102 | 4.272339 | 1.363378 | 2.693558 | 0.007069 | 0.04239721 | ENSG00000149654 | CDH22 | 64405 |
| ENSG00000280105.1 | 1.733182064 | 4.268771031 | 1.372613258 | 2.672837 | 0.007521 | 0.044744511 | ENSG00000280105 | NA | NA |
| ENSG00000257294.2 | 1.702784038 | 4.268363373 | 1.321512014 | 2.775883 | 0.005505 | 0.034043962 | ENSG00000257294 | NA | NA |
| ENSG00000249857.1 | 2.517682381 | 4.268355259 | 1.031518981 | 3.556265 | 0.000376 | 0.003050332 | ENSG00000249857 | NA | NA |
| ENSG00000163221.9 | 414.2158889 | 4.266875848 | 0.475822322 | 7.706397 | 1.29E-14 | 3.42E-13 | ENSG00000163221 | S100A12 | 6283 |
| ENSG00000176490.5 | 3.46587799 | 4.260960396 | 1.02825468 | 3.560363 | 0.00037 | 0.003006783 | ENSG00000176490 | DIRAS1 | 148252 |
| ENSG00000142583.17 | 51.11312506 | 4.254418352 | 0.506994499 | 7.208004 | 5.68E-13 | 1.33E-11 | ENSG00000142583 | SLC2A5 | 6518 |
| ENSG00000184502.4 | 72.84924724 | 4.251839066 | 0.263915508 | 13.83715 | 1.52E-43 | 1.75E-41 | ENSG00000184502 | GAST | 2520 |
| ENSG00000101331.16 | 3.490049 | 4.249687 | 1.077739 | 3.386431 | 0.000708 | 0.005440274 | ENSG00000101331 | CCM2L | 140706 |
| ENSG00000206384.10 | 42.57717121 | 4.248172629 | 0.432396361 | 8.437103 | 3.25E-17 | 1.05E-15 | ENSG00000206384 | COL6A6 | 131873 |
| ENSG00000236375.3 | 20.95777722 | 4.247930125 | 0.4703129 | 7.75639 | 8.74E-15 | 2.35E-13 | ENSG00000236375 | NA | NA |
| ENSG00000227012.3 | 6.368255104 | 4.244909881 | 0.890334398 | 4.093866 | 4.24E-05 | 0.000406418 | ENSG00000227012 | NA | NA |
| ENSG00000255201.1 | 2.152220539 | 4.243614159 | 1.140704204 | 3.19418 | 0.001402 | 0.010038106 | ENSG00000255201 | NA | NA |
| ENSG00000253892.1 | 2.152647756 | 4.242154254 | 1.202700445 | 3.028314 | 0.002459 | 0.016661473 | ENSG00000253892 | NA | NA |
| ENSG00000160862.13 | 76.49377839 | 4.241988122 | 0.754943806 | 4.824184 | 1.41E-06 | 1.66E-05 | ENSG00000160862 | AZGP1 | 563 |
| ENSG00000263429.3 | 2.147389517 | 4.241927248 | 1.278111916 | 2.849459 | 0.004379 | 0.027878315 | ENSG00000263429 | TMEM238l | 1E+08 |
| ENSG00000228216.1 | 3.759661166 | 4.238065888 | 0.94987272 | 3.830056 | 0.000128 | 0.001133483 | ENSG00000228216 | NA | NA |
| ENSG00000205231.1 | 1.692065981 | 4.235100423 | 1.381837164 | 2.630629 | 0.008523 | 0.04985533 | ENSG00000205231 | TTLL10-AS | 1.01E+08 |
| ENSG00000173966.7 | 2.376901453 | 4.233193783 | 1.174666305 | 3.092958 | 0.001982 | 0.013682572 | ENSG00000173966 | NA | NA |
| ENSG00000140873.16 | 1.950023011 | 4.23236479 | 1.274860498 | 2.849225 | 0.004383 | 0.027885548 | ENSG00000140873 | ADAMTS1 | 170692 |

| | | | | | | | | | |
|---|---|---|---|---|---|---|---|---|---|
| ENSG00000129682.16 | 6.061176 | 4.232262 | 0.9489502 | 3.827663 | 0.000129 | 0.001143795 | ENSG00000129682 | FGF13 | 2258 |
| ENSG00000237125.10 | 1.684213115 | 4.231039498 | 1.212801929 | 2.993926 | 0.002754 | 0.018436089 | ENSG00000237125 | HAND2-AS | 79804 |
| ENSG00000153162.9 | 27.77077268 | 4.228184434 | 0.560636701 | 6.471543 | 9.70E-11 | 1.85E-09 | ENSG00000153162 | BMP6 | 654 |
| ENSG00000047936.10 | 3.469903181 | 4.227314098 | 1.034027989 | 3.507946 | 0.000452 | 0.003601556 | ENSG00000047936 | ROS1 | 6098 |
| ENSG00000267696.6 | 1.637968 | 4.223415 | 1.373884 | 2.637351 | 0.008356 | 0.04898561 | ENSG00000267696 | NA | NA |
| ENSG00000286257.2 | 1.63686551 | 4.216880629 | 1.328545367 | 2.722437 | 0.00648 | 0.039253035 | ENSG00000286257 | NA | NA |
| ENSG00000187045.18 | 36.43588 | 4.21155 | 0.3727096 | 9.689984 | 3.33E-22 | 1.44E-20 | ENSG00000187045 | TMPRSS6 | 164656 |
| ENSG00000196139.14 | 815.2940881 | 4.205015455 | 0.130354078 | 27.65556 | 2.39E-168 | 1.93E-165 | ENSG00000196139 | AKR1C3 | 8644 |
| ENSG00000253775.2 | 6.251324473 | 4.204640517 | 0.827473906 | 4.356198 | 1.32E-05 | 0.000137249 | ENSG00000253775 | NA | NA |
| ENSG00000188984.12 | 1.763677381 | 4.20055791 | 1.113349449 | 3.233987 | 0.001221 | 0.008877023 | ENSG00000188984 | AADACL3 | 126767 |
| ENSG00000161681.15 | 3.245316 | 4.196947 | 1.168348 | 3.078662 | 0.002079 | 0.01429645 | ENSG00000161681 | SHANK1 | 50944 |
| ENSG00000232628.6 | 1.751687082 | 4.194727069 | 1.162392404 | 3.092525 | 0.001985 | 0.01369479 | ENSG00000232628 | LOC10192 | 1.02E+08 |
| ENSG00000261398.1 | 1.868529431 | 4.193657028 | 1.140909262 | 3.149818 | 0.001634 | 0.011506054 | ENSG00000261398 | LOC11226 | 1.12E+08 |
| ENSG00000163823.4 | 4.797408145 | 4.191752288 | 0.889593062 | 4.037523 | 5.40E-05 | 0.000507977 | ENSG00000163823 | CCR1 | 1230 |
| ENSG00000267196.1 | 1.669556986 | 4.189467294 | 1.360593922 | 2.638162 | 0.008336 | 0.048890127 | ENSG00000267196 | NA | NA |
| ENSG00000231561.2 | 1.670787 | 4.188594 | 1.26345 | 2.840313 | 0.004507 | 0.02853718 | ENSG00000231561 | NA | NA |
| ENSG00000144366.16 | 498.9773938 | 4.185046983 | 0.307715925 | 11.65051 | 2.28E-31 | 1.67E-29 | ENSG00000144366 | GULP1 | 51454 |
| ENSG00000125144.13 | 61.981752 | 4.184378806 | 0.318793368 | 11.24358 | 2.49E-29 | 1.66E-27 | ENSG00000125144 | MT1G | 4495 |
| ENSG00000249007.1 | 9.072868236 | 4.184149336 | 0.690009824 | 5.194345 | 2.05E-07 | 2.71E-06 | ENSG00000249007 | NA | NA |
| ENSG00000160951.4 | 3.235238479 | 4.184047652 | 1.104322145 | 3.245473 | 0.001173 | 0.008552278 | ENSG00000160951 | PTGER1 | 5731 |
| ENSG00000178372.7 | 208.8146318 | 4.183923252 | 0.473309188 | 7.572055 | 3.67E-14 | 9.37E-13 | ENSG00000178372 | CALML5 | 51806 |
| ENSG00000158406.4 | 101.5815657 | 4.182080723 | 0.248883074 | 14.39262 | 5.76E-47 | 7.33E-45 | ENSG00000158406 | H4C8 | 8365 |
| ENSG00000186526.12 | 1.655438 | 4.179278 | 1.149457 | 3.113886 | 0.001846 | 0.01285434 | ENSG00000186526 | CYP4F8 | 11283 |
| ENSG00000244541.6 | 17.51127308 | 4.175492744 | 0.639556325 | 5.590583 | 2.26E-08 | 3.36E-07 | ENSG00000244541 | LINC0121: | 1.02E+08 |
| ENSG00000198580.7 | 1.651017208 | 4.174103962 | 1.318395131 | 2.710951 | 0.006709 | 0.040500838 | ENSG00000198580 | NA | NA |
| ENSG00000184148.4 | 54.91101826 | 4.17241894 | 0.468956245 | 7.617809 | 2.58E-14 | 6.69E-13 | ENSG00000184148 | SPRR4 | 163778 |
| ENSG00000283283.2 | 6.354589789 | 4.17012237 | 0.829429765 | 4.304309 | 1.68E-05 | 0.000170661 | ENSG00000283283 | NA | NA |
| ENSG00000223458.2 | 3.292121492 | 4.169212425 | 1.01620261 | 3.512304 | 0.000444 | 0.003551516 | ENSG00000223458 | LMO7DN-I | 1.04E+08 |
| ENSG00000112139.16 | 569.8397801 | 4.166432798 | 0.434548978 | 8.207206 | 2.26E-16 | 6.78E-15 | ENSG00000112139 | MDGA1 | 266727 |
| ENSG00000005108.16 | 20.62398204 | 4.166431518 | 0.455022174 | 7.837929 | 4.58E-15 | 1.26E-13 | ENSG00000005108 | THSD7A | 221981 |
| ENSG00000224251.6 | 13.52395592 | 4.161458136 | 0.589188921 | 6.044679 | 1.50E-09 | 2.52E-08 | ENSG00000224251 | NA | NA |
| ENSG00000225647.2 | 2.272060796 | 4.159330986 | 1.322404484 | 2.69156 | 0.007112 | 0.042623194 | ENSG00000225647 | NA | NA |
| ENSG00000146038.12 | 6.0197278 | 4.157750336 | 0.867545327 | 4.100939 | 4.11E-05 | 0.000395179 | ENSG00000146038 | DCDC2 | 51473 |
| ENSG00000154493.19 | 2.972793461 | 4.156604471 | 1.072760117 | 3.315377 | 0.000915 | 0.006846597 | ENSG00000154493 | C10orf90 | 118611 |
| ENSG00000085741.13 | 5.232206081 | 4.152724214 | 0.912347453 | 3.894047 | 9.86E-05 | 0.000891791 | ENSG00000085741 | WNT11 | 7481 |
| ENSG00000115165.10 | 29.53712974 | 4.152215623 | 0.412682891 | 8.607615 | 7.46E-18 | 2.50E-16 | ENSG00000115165 | CYTIP | 9595 |
| ENSG00000108759.3 | 3.180178132 | 4.146786287 | 1.039287536 | 3.412709 | 0.000643 | 0.004976287 | ENSG00000108759 | KRT32 | 3882 |
| ENSG00000255916.1 | 2.456572831 | 4.142864949 | 1.332965333 | 2.657882 | 0.007863 | 0.046540176 | ENSG00000255916 | LOC10192 | 1.02E+08 |
| ENSG00000204020.5 | 1.549595829 | 4.14188647 | 1.215417674 | 2.914131 | 0.003567 | 0.023159845 | ENSG00000204020 | LIPN | 643418 |
| ENSG00000140678.16 | 39.56374349 | 4.141487083 | 0.342994972 | 10.32519 | 5.42E-25 | 2.76E-23 | ENSG00000140678 | ITGAX | 3687 |
| ENSG00000165970.11 | 1.545029339 | 4.137297614 | 1.193521591 | 2.963748 | 0.003039 | 0.02012154 | ENSG00000165970 | SLC6A5 | 9152 |
| ENSG00000197585.10 | 1.559968433 | 4.136094989 | 1.154490153 | 3.062906 | 0.002192 | 0.015020618 | ENSG00000197585 | NA | NA |

| | | | | | | | | | |
|---|---|---|---|---|---|---|---|---|---|
| ENSG00000176092.15 | 2071.141466 | 4.13388288 | 0.390226817 | 9.055971 | 1.35E-19 | 5.05E-18 | ENSG00000176092 | CRYBG2 | 55057 |
| ENSG00000136267.13 | 1.602705515 | 4.131297668 | 1.332722601 | 2.649687 | 0.008057 | 0.047512772 | ENSG00000136267 | DGKB | 1607 |
| ENSG00000258550.1 | 2.158873942 | 4.129335398 | 1.192846517 | 2.958751 | 0.003089 | 0.02041013 | ENSG00000258550 | OR7E106F | 81115 |
| ENSG00000163121.10 | 35.64835098 | 4.128700084 | 0.51677818 | 6.828268 | 8.59E-12 | 1.82E-10 | ENSG00000163121 | NEURL3 | 93082 |
| ENSG00000162571.13 | 21.07169098 | 4.127359531 | 0.668226031 | 5.278692 | 1.30E-07 | 1.77E-06 | ENSG00000162571 | TTLL10 | 254173 |
| ENSG00000283052.1 | 3.420945798 | 4.125532407 | 1.038183902 | 3.395865 | 0.000684 | 0.005271394 | ENSG00000283052 | NA | NA |
| ENSG00000140450.9 | 4186.442603 | 4.119203618 | 0.119169015 | 29.5312 | 1.15E-191 | 1.27E-188 | ENSG00000140450 | ARRDC4 | 91947 |
| ENSG00000237417.1 | 20.96942275 | 4.118489377 | 0.536451204 | 6.558825 | 5.42E-11 | 1.06E-09 | ENSG00000237417 | NA | NA |
| ENSG00000256704.3 | 4.800078846 | 4.118485972 | 1.03952185 | 3.384716 | 0.000713 | 0.005469639 | ENSG00000256704 | NA | NA |
| ENSG00000235314.1 | 4.524457611 | 4.116874043 | 0.898846208 | 3.912654 | 9.13E-05 | 0.000829155 | ENSG00000235314 | NA | NA |
| ENSG00000263293.2 | 1.57067247 | 4.116424772 | 1.131044064 | 3.109008 | 0.001877 | 0.01303782 | ENSG00000263293 | THCAT158 | 1.03E+08 |
| ENSG00000100095.19 | 1.566621 | 4.112276 | 1.325829 | 2.649116 | 0.00807 | 0.04755343 | ENSG00000100095 | SEZ6L | 23544 |
| ENSG00000163141.19 | 349.0599686 | 4.110046603 | 0.424711413 | 8.264545 | 1.40E-16 | 4.30E-15 | ENSG00000163141 | BNIPL | 149428 |
| ENSG00000206820.1 | 7.2612327 | 4.108941159 | 0.747900254 | 4.691723 | 2.71E-06 | 3.09E-05 | ENSG00000206820 | RNU1-138 | 1.06E+08 |
| ENSG00000187185.4 | 15.42956136 | 4.103922312 | 0.551511306 | 6.35331 | 2.11E-10 | 3.92E-09 | ENSG00000187185 | LOC38828 | 388282 |
| ENSG00000167100.14 | 3.123455877 | 4.103345343 | 1.061273894 | 3.301076 | 0.000963 | 0.007168287 | ENSG00000167100 | SAMD14 | 201191 |
| ENSG00000213199.8 | 85.62670621 | 4.103061554 | 0.305133135 | 11.48044 | 1.65E-30 | 1.17E-28 | ENSG00000213199 | ASIC3 | 9311 |
| ENSG00000174343.6 | 37.45574 | 4.102684902 | 0.341424458 | 10.25903 | 1.08E-24 | 5.31E-23 | ENSG00000174343 | CHRNA9 | 55584 |
| ENSG00000137868.19 | 1033.018457 | 4.099743331 | 0.376329973 | 9.299667 | 1.41E-20 | 5.62E-19 | ENSG00000137868 | STRA6 | 64220 |
| ENSG00000130222.11 | 34.57892444 | 4.097231257 | 0.384923838 | 9.085515 | 1.03E-19 | 3.86E-18 | ENSG00000130222 | GADD45G | 10912 |
| ENSG00000241956.10 | 2.171388294 | 4.090283041 | 1.186462878 | 2.941755 | 0.003264 | 0.021399765 | ENSG00000241956 | LOC10254 | 1.03E+08 |
| ENSG00000148346.12 | 7118.852559 | 4.087778333 | 0.413591556 | 8.432905 | 3.37E-17 | 1.08E-15 | ENSG00000148346 | LCN2 | 3934 |
| ENSG00000249125.2 | 2.068540915 | 4.08749108 | 1.204376549 | 2.895682 | 0.003783 | 0.024391971 | ENSG00000249125 | NA | NA |
| ENSG00000287448.1 | 68.95776927 | 4.086772463 | 0.285346352 | 12.21944 | 2.45E-34 | 1.95E-32 | ENSG00000287448 | NA | NA |
| ENSG00000173110.8 | 95.77218675 | 4.083691569 | 0.353893648 | 9.843894 | 7.28E-23 | 3.27E-21 | ENSG00000173110 | HSPA6 | 3310 |
| ENSG00000218561.1 | 2.15691712 | 4.080957766 | 1.149065994 | 3.02938 | 0.002451 | 0.016612892 | ENSG00000218561 | NA | NA |
| ENSG00000233680.4 | 24.86951 | 4.080178 | 0.4707771 | 7.392411 | 1.44E-13 | 3.55E-12 | ENSG00000233680 | HNRNPA1 | 664721 |
| ENSG00000229876.1 | 2.166323 | 4.076095 | 1.101028 | 3.157135 | 0.001593 | 0.01124201 | ENSG00000229876 | CASC20 | 1.02E+08 |
| ENSG00000105642.15 | 1.52656 | 4.075714 | 1.19588 | 2.906406 | 0.003656 | 0.02367593 | ENSG00000105642 | KCNN1 | 3780 |
| ENSG00000235885.8 | 3.042572669 | 4.07427264 | 1.131686349 | 3.069996 | 0.002141 | 0.01469128 | ENSG00000235885 | LOC10192 | 1.02E+08 |
| ENSG00000066056.14 | 2.258898036 | 4.072701704 | 1.228385939 | 2.827044 | 0.004698 | 0.029602278 | ENSG00000066056 | TIE1 | 7075 |
| ENSG00000237512.6 | 47.52256121 | 4.070932622 | 0.349738032 | 9.924379 | 3.26E-23 | 1.49E-21 | ENSG00000237512 | UNC5B-AS | 728978 |
| ENSG00000286647.1 | 3.008857399 | 4.066134344 | 1.069258506 | 3.241624 | 0.001189 | 0.008652011 | ENSG00000286647 | NA | NA |
| ENSG00000159674.12 | 82.9252306 | 4.063243308 | 0.283220878 | 12.22807 | 2.20E-34 | 1.76E-32 | ENSG00000159674 | SPON2 | 10417 |
| ENSG00000143217.9 | 2916.620093 | 4.060339724 | 0.34052184 | 10.16187 | 2.93E-24 | 1.42E-22 | ENSG00000143217 | NECTIN4 | 81607 |
| ENSG00000087086.15 | 150279.6 | 4.057283 | 0.2504365 | 13.80503 | 2.38E-43 | 2.70E-41 | ENSG00000087086 | FTL | 2512 |
| ENSG00000241912.1 | 10.52760577 | 4.053471337 | 0.7591986 | 4.548838 | 5.39E-06 | 5.93E-05 | ENSG00000241912 | NA | NA |
| ENSG00000287052.1 | 1.781519533 | 4.053351044 | 1.25025623 | 2.762115 | 0.005743 | 0.035316619 | ENSG00000287052 | NA | NA |
| ENSG00000279255.1 | 1.452818698 | 4.053197423 | 1.165827217 | 2.962015 | 0.003056 | 0.020220093 | ENSG00000279255 | NA | NA |
| ENSG00000259602.1 | 1.773212678 | 4.052968643 | 1.295329326 | 2.665707 | 0.007683 | 0.04558229 | ENSG00000259602 | NA | NA |
| ENSG00000187867.9 | 5.81371319 | 4.051929286 | 0.863236165 | 3.998824 | 6.37E-05 | 0.000592554 | ENSG00000187867 | PALM3 | 342979 |
| ENSG00000224511.2 | 2.160048147 | 4.051130555 | 1.077355728 | 3.203334 | 0.001358 | 0.00975586 | ENSG00000224511 | LINC0036 | 1.01E+08 |

| | | | | | | | | | |
|---|---|---|---|---|---|---|---|---|---|
| ENSG00000251728.1 | 3.115511324 | 4.05059689 | 0.993192528 | 3.474248 | 0.000512 | 0.004046922 | ENSG00000251728 | NA | NA |
| ENSG00000260604.2 | 27.17242438 | 4.044580632 | 0.42740154 | 8.059355 | 7.67E-16 | 2.23E-14 | ENSG00000260604 | NA | NA |
| ENSG00000171403.10 | 156.3844026 | 4.041593492 | 0.188791422 | 18.22961 | 3.00E-74 | 8.00E-72 | ENSG00000171403 | KRT9 | 3857 |
| ENSG00000228208.7 | 1.450312979 | 4.040918785 | 1.210878856 | 2.841671 | 0.004488 | 0.028442698 | ENSG00000228208 | LINC0286 | 440714 |
| ENSG00000224144.1 | 1.443933 | 4.036463 | 1.223485 | 2.808749 | 0.004973 | 0.03112432 | ENSG00000224144 | NA | NA |
| ENSG00000177169.10 | 8521.521482 | 4.036069669 | 0.252106922 | 13.62941 | 2.68E-42 | 2.92E-40 | ENSG00000177169 | ULK1 | 8408 |
| ENSG00000120279.6 | 15.6041002 | 4.035486984 | 0.52420922 | 6.553656 | 5.61E-11 | 1.09E-09 | ENSG00000120279 | MYCT1 | 80177 |
| ENSG00000163431.13 | 2.974978792 | 4.034478334 | 1.042309727 | 3.295065 | 0.000984 | 0.007305007 | ENSG00000163431 | LMOD1 | 25802 |
| ENSG00000206769.1 | 1.596684193 | 4.031160861 | 1.286077082 | 2.667928 | 0.007632 | 0.045312405 | ENSG00000206769 | RNU6-116 | 1.06E+08 |
| ENSG00000188505.5 | 1509.258 | 4.030936 | 0.2566854 | 13.36631 | 9.51E-41 | 9.74E-39 | ENSG00000188505 | NCCRP1 | 342897 |
| ENSG00000251209.9 | 8.168741968 | 4.028125904 | 0.768641881 | 4.459978 | 8.20E-06 | 8.82E-05 | ENSG00000251209 | LINC0092 | 91948 |
| ENSG00000185634.12 | 113.1589114 | 4.024895297 | 0.327333379 | 10.46302 | 1.28E-25 | 6.75E-24 | ENSG00000185634 | SHC4 | 399694 |
| ENSG00000244036.3 | 10.81569854 | 4.024569404 | 0.636557787 | 5.379825 | 7.46E-08 | 1.04E-06 | ENSG00000244036 | NA | NA |
| ENSG00000268833.1 | 2.173448 | 4.0245 | 1.161751 | 2.947706 | 0.003201 | 0.02106476 | ENSG00000268833 | NA | NA |
| ENSG00000159904.11 | 2.965237441 | 4.023096622 | 1.221226986 | 2.802998 | 0.005063 | 0.031625341 | ENSG00000159904 | ZNF890P | 645700 |
| ENSG00000251211.1 | 1.518764842 | 4.020101767 | 1.295319232 | 2.640354 | 0.008282 | 0.048682217 | ENSG00000251211 | LOC64585 | 645853 |
| ENSG00000233681.1 | 1.581754 | 4.019001 | 1.270757 | 2.690523 | 0.007134 | 0.04271746 | ENSG00000233681 | CEACAMF | 1090 |
| ENSG00000166670.10 | 583.9398155 | 4.018103819 | 0.180883109 | 18.89676 | 1.21E-79 | 3.55E-77 | ENSG00000166670 | MMP10 | 4319 |
| ENSG00000223722.3 | 38.05038396 | 4.015256894 | 0.460369276 | 7.418516 | 1.18E-13 | 2.94E-12 | ENSG00000223722 | NA | NA |
| ENSG00000254714.1 | 1.489924651 | 4.014688945 | 1.242937828 | 2.747273 | 0.006009 | 0.036709514 | ENSG00000254714 | NA | NA |
| ENSG00000274414.1 | 474.4944 | 4.00916 | 0.277866 | 12.26908 | 1.33E-34 | 1.07E-32 | ENSG00000274414 | NA | NA |
| ENSG00000170345.10 | 1357.582112 | 4.007228542 | 0.195056043 | 17.46795 | 2.51E-68 | 5.87E-66 | ENSG00000170345 | FOS | 2353 |
| ENSG00000267134.3 | 14.09647308 | 4.003437434 | 0.662862498 | 5.134455 | 2.83E-07 | 3.67E-06 | ENSG00000267134 | LINC0192 | 284294 |
| ENSG00000267462.1 | 8.099238205 | 3.999735193 | 0.764684559 | 4.445932 | 8.75E-06 | 9.36E-05 | ENSG00000267462 | NA | NA |
| ENSG00000171346.16 | 13081.8575 | 3.99817217 | 0.31737614 | 10.70708 | 9.43E-27 | 5.23E-25 | ENSG00000171346 | KRT15 | 3866 |
| ENSG00000085265.11 | 2.970411475 | 3.995567319 | 1.2047136 | 2.818568 | 0.004824 | 0.030280643 | ENSG00000085265 | FCN1 | 2219 |
| ENSG00000082482.13 | 3.323419279 | 3.987029543 | 0.959600495 | 3.529625 | 0.000416 | 0.003346039 | ENSG00000082482 | KCNK2 | 3776 |
| ENSG00000256843.1 | 17.99365938 | 3.98474668 | 0.464334853 | 7.289452 | 3.11E-13 | 7.42E-12 | ENSG00000256843 | NA | NA |
| ENSG00000167261.14 | 3.147653449 | 3.98343844 | 1.136345382 | 2.977474 | 0.002906 | 0.019323863 | ENSG00000167261 | DPEP2 | 64174 |
| ENSG00000231605.6 | 3.063387221 | 3.980791202 | 1.045816716 | 3.23268 | 0.001226 | 0.008903137 | ENSG00000231605 | LINC0136 | 1.02E+08 |
| ENSG00000134258.17 | 25.54565548 | 3.979448223 | 0.54680363 | 6.180369 | 6.40E-10 | 1.13E-08 | ENSG00000134258 | VTCN1 | 79679 |
| ENSG00000217801.10 | 460.9605603 | 3.978478443 | 0.245768847 | 13.74657 | 5.34E-43 | 5.97E-41 | ENSG00000217801 | LOC10028 | 1E+08 |
| ENSG00000183625.15 | 41.20848834 | 3.977448848 | 0.533412353 | 6.331778 | 2.42E-10 | 4.48E-09 | ENSG00000183625 | CCR3 | 1232 |
| ENSG00000150337.13 | 4.296370756 | 3.975171137 | 0.988599917 | 3.414092 | 0.00064 | 0.004952539 | ENSG00000150337 | FCGR1A | 2209 |
| ENSG00000125831.10 | 2.032795 | 3.973257 | 1.058921 | 3.185559 | 0.001445 | 0.01030324 | ENSG00000125831 | CST11 | 140880 |
| ENSG00000259342.1 | 4.698262032 | 3.972314013 | 1.004220787 | 3.35814 | 0.000785 | 0.005968565 | ENSG00000259342 | NA | NA |
| ENSG00000128965.13 | 852.0652479 | 3.972155902 | 0.089557083 | 37.6537 | 0 | 0 | ENSG00000128965 | CHAC1 | 79094 |
| ENSG00000240694.9 | 18.50746581 | 3.97112262 | 0.503028117 | 6.701658 | 2.06E-11 | 4.20E-10 | ENSG00000240694 | PNMA2 | 10687 |
| ENSG00000163638.13 | 7.566120263 | 3.969791704 | 0.872033644 | 3.864291 | 0.000111 | 0.000998324 | ENSG00000163638 | ADAMTS9 | 56999 |
| ENSG00000164330.17 | 2.807004985 | 3.961409905 | 1.061796722 | 3.165775 | 0.001547 | 0.010945745 | ENSG00000164330 | EBF1 | 1879 |
| ENSG00000175155.9 | 405.9095119 | 3.961204862 | 0.244008725 | 13.77494 | 3.61E-43 | 4.07E-41 | ENSG00000175155 | YPEL2 | 388403 |
| ENSG00000158055.15 | 3125.55085 | 3.955514524 | 0.362203349 | 9.264173 | 1.97E-20 | 7.77E-19 | ENSG00000158055 | GRHL3 | 57822 |

| | | | | | | | | | |
|---|---|---|---|---|---|---|---|---|---|
| ENSG00000167232.14 | 1.432448 | 3.954477 | 1.176454 | 2.851344 | 0.004353 | 0.02772687 | ENSG00000167232 | ZNF91 | 7644 |
| ENSG00000103089.9 | 86.88928752 | 3.949975709 | 0.68587081 | 4.884266 | 1.04E-06 | 1.25E-05 | ENSG00000103089 | FA2H | 79152 |
| ENSG00000173578.7 | 1.602239238 | 3.949958079 | 1.262021146 | 2.654439 | 0.007944 | 0.046913547 | ENSG00000173578 | XCR1 | 2829 |
| ENSG00000268307.1 | 17.96283 | 3.949928 | 0.4880667 | 6.863667 | 6.71E-12 | 1.44E-10 | ENSG00000268307 | LINC0256( | 1.11E+08 |
| ENSG00000230601.6 | 31.28455372 | 3.949036853 | 0.376998581 | 8.88342 | 6.48E-19 | 2.31E-17 | ENSG00000230601 | TEX48 | 1.01E+08 |
| ENSG00000258162.4 | 1.963539748 | 3.947803633 | 1.132346551 | 2.956519 | 0.003111 | 0.020538049 | ENSG00000258162 | NA | NA |
| ENSG00000179388.9 | 644.4283493 | 3.944674538 | 0.249181592 | 13.42264 | 4.46E-41 | 4.62E-39 | ENSG00000179388 | EGR3 | 1960 |
| ENSG00000186007.10 | 24.55781648 | 3.942371872 | 0.414401678 | 8.065537 | 7.29E-16 | 2.13E-14 | ENSG00000186007 | LEMD1 | 93273 |
| ENSG00000186517.14 | 234.6562986 | 3.94222718 | 0.295932597 | 11.29388 | 1.41E-29 | 9.48E-28 | ENSG00000186517 | ARHGAP3 | 257106 |
| ENSG00000079393.20 | 11.98201274 | 3.941966548 | 0.802332027 | 4.165316 | 3.11E-05 | 0.000304091 | ENSG00000079393 | DUSP13 | 51207 |
| ENSG00000286708.1 | 3.737269278 | 3.941926877 | 0.999882568 | 3.342319 | 0.000831 | 0.006290652 | ENSG00000286708 | NA | NA |
| ENSG00000112280.16 | 1.787045831 | 3.941382551 | 1.245612204 | 2.682522 | 0.007307 | 0.04367423 | ENSG00000112280 | COL9A1 | 1297 |
| ENSG00000173210.19 | 1303.34311 | 3.94107269 | 0.270914849 | 12.33256 | 6.05E-35 | 4.93E-33 | ENSG00000173210 | ABLIM3 | 22885 |
| ENSG00000245848.3 | 464.3124 | 3.939633 | 0.2886005 | 11.57182 | 5.73E-31 | 4.10E-29 | ENSG00000245848 | CEBPA | 1050 |
| ENSG00000163285.8 | 5.32024551 | 3.937365151 | 0.923076328 | 3.615481 | 0.0003 | 0.002476269 | ENSG00000163285 | GABRG1 | 2565 |
| ENSG00000260599.1 | 1.393205 | 3.936264 | 1.240806 | 2.688787 | 0.007171 | 0.04292088 | ENSG00000260599 | LOC10192 | 1.02E+08 |
| ENSG00000144158.4 | 12.92147246 | 3.936118806 | 0.596266456 | 5.595013 | 2.21E-08 | 3.28E-07 | ENSG00000144158 | NA | NA |
| ENSG00000269993.1 | 6.347295 | 3.932553 | 0.9846722 | 3.384429 | 0.000713 | 0.005473784 | ENSG00000269993 | NA | NA |
| ENSG00000182759.3 | 29.2713025 | 3.930861598 | 0.491117146 | 6.782214 | 1.18E-11 | 2.47E-10 | ENSG00000182759 | MAFA | 389692 |
| ENSG00000182183.15 | 2.849836671 | 3.929503235 | 1.179741284 | 2.822232 | 0.004769 | 0.029982605 | ENSG00000182183 | SHISAL2A | 348378 |
| ENSG00000227471.9 | 129.3023726 | 3.927750994 | 0.300553282 | 11.07208 | 1.71E-28 | 1.08E-26 | ENSG00000227471 | AKR1B15 | 441282 |
| ENSG00000122133.17 | 4.323398515 | 3.923745211 | 1.03890851 | 3.199267 | 0.001378 | 0.009881247 | ENSG00000122133 | PAEP | 5047 |
| ENSG00000199150.3 | 1.948818034 | 3.923495213 | 1.248059324 | 2.66293 | 0.007746 | 0.045919196 | ENSG00000199150 | MIRLET7G | 406890 |
| ENSG00000137674.4 | 2.74477024 | 3.922976293 | 1.021245123 | 3.253848 | 0.001139 | 0.008336085 | ENSG00000137674 | MMP20 | 9313 |
| ENSG00000282950.1 | 2.775188165 | 3.917275742 | 1.038854342 | 3.193206 | 0.001407 | 0.010069315 | ENSG00000282950 | NA | NA |
| ENSG00000136842.14 | 69.2200584 | 3.912021151 | 0.260601389 | 12.70915 | 5.26E-37 | 4.72E-35 | ENSG00000136842 | TMOD1 | 7111 |
| ENSG00000287629.1 | 8.886754142 | 3.910804031 | 0.827426467 | 4.001327 | 6.30E-05 | 0.000586729 | ENSG00000287629 | NA | NA |
| ENSG00000139364.10 | 10.96469205 | 3.909384538 | 0.717409411 | 4.612965 | 3.97E-06 | 4.43E-05 | ENSG00000139364 | TMEM132I | 114795 |
| ENSG00000136689.18 | 4267.638 | 3.907772585 | 0.355818876 | 9.296226 | 1.46E-20 | 5.79E-19 | ENSG00000136689 | IL1RN | 3557 |
| ENSG00000271830.1 | 2.861569137 | 3.899259616 | 1.144106877 | 2.883699 | 0.00393 | 0.025267723 | ENSG00000271830 | NA | NA |
| ENSG00000130720.13 | 33.17538263 | 3.895432725 | 0.51772449 | 6.365225 | 1.95E-10 | 3.64E-09 | ENSG00000130720 | FIBCD1 | 84929 |
| ENSG00000203930.12 | 5.105313 | 3.895256 | 0.9268523 | 3.55532 | 0.000378 | 0.003059463 | ENSG00000203930 | LINC0063 | 286411 |
| ENSG00000132746.14 | 397.4683457 | 3.891294573 | 0.449832977 | 7.316704 | 2.54E-13 | 6.11E-12 | ENSG00000132746 | ALDH3B2 | 222 |
| ENSG00000187689.10 | 151.7949908 | 3.89075473 | 0.529430379 | 6.215651 | 5.11E-10 | 9.11E-09 | ENSG00000187689 | AMTN | 401138 |
| ENSG00000170498.9 | 55.59817021 | 3.886930497 | 0.388713177 | 8.455928 | 2.77E-17 | 8.95E-16 | ENSG00000170498 | KISS1 | 3814 |
| ENSG00000225434.3 | 32.52567594 | 3.886768435 | 0.508063909 | 6.469203 | 9.85E-11 | 1.88E-09 | ENSG00000225434 | LINC0150 | 1.01E+08 |
| ENSG00000147168.12 | 7.72789 | 3.88181 | 0.7693281 | 4.265813 | 1.99E-05 | 0.00020032 | ENSG00000147168 | IL2RG | 3561 |
| ENSG00000183186.8 | 207.834757 | 3.881621692 | 0.260443831 | 12.60011 | 2.11E-36 | 1.85E-34 | ENSG00000183186 | C2CD4C | 126567 |
| ENSG00000133115.11 | 7.420051511 | 3.880055518 | 0.751500627 | 4.364674 | 1.27E-05 | 0.000132602 | ENSG00000133115 | STOML3 | 161003 |
| ENSG00000105289.15 | 202.3096225 | 3.879823914 | 0.235711043 | 13.9146 | 5.17E-44 | 6.03E-42 | ENSG00000105289 | TJP3 | 27134 |
| ENSG00000253214.2 | 1.289147507 | 3.878053178 | 1.196824155 | 2.73896 | 0.006163 | 0.03756452 | ENSG00000253214 | LOC10537 | 1.05E+08 |
| ENSG00000184270.5 | 6.619944038 | 3.872981981 | 0.980350116 | 3.338585 | 0.000842 | 0.006359549 | ENSG00000184270 | H2AC21 | 317772 |

| | | | | | | | | | |
|---|---|---|---|---|---|---|---|---|---|
| ENSG00000133985.3 | 1217.478157 | 3.871418748 | 0.251663536 | 12.99918 | 1.24E-38 | 1.17E-36 | ENSG00000133985 | TTC9 | 23508 |
| ENSG00000253706.5 | 7.139077709 | 3.869341102 | 0.851171892 | 3.840988 | 0.000123 | 0.001088146 | ENSG00000253706 | NA | NA |
| ENSG00000228874.1 | 1.866496683 | 3.863568822 | 1.109185069 | 2.942312 | 0.003258 | 0.021371815 | ENSG00000228874 | RPS24P1( | 1E+08 |
| ENSG00000283646.2 | 683.1902891 | 3.860678786 | 0.430602725 | 7.57236 | 3.67E-14 | 9.35E-13 | ENSG00000283646 | LINC0200( | 1.05E+08 |
| ENSG00000185352.8 | 249.9221432 | 3.859623729 | 0.296365926 | 10.99865 | 3.88E-28 | 2.39E-26 | ENSG00000185352 | HS6ST3 | 266722 |
| ENSG00000267453.7 | 199.9409 | 3.859452 | 0.2952887 | 11.03819 | 2.50E-28 | 1.55E-26 | ENSG00000267453 | CLEC4O | 1.03E+08 |
| ENSG00000078328.20 | 1.304323517 | 3.858114882 | 1.212686856 | 2.686691 | 0.007216 | 0.043181419 | ENSG00000078328 | RBFOX1 | 54715 |
| ENSG00000225532.1 | 1.279930756 | 3.856892108 | 1.202980665 | 2.707352 | 0.006782 | 0.040905423 | ENSG00000225532 | NA | NA |
| ENSG00000223573.7 | 693.9068645 | 3.854973216 | 0.25579692 | 12.72483 | 4.30E-37 | 3.88E-35 | ENSG00000223573 | TINCR | 257000 |
| ENSG00000167653.5 | 116.9445888 | 3.849644186 | 0.323253613 | 10.05292 | 8.92E-24 | 4.22E-22 | ENSG00000167653 | PSCA | 8000 |
| ENSG00000162444.12 | 22.11151498 | 3.840937178 | 0.527948793 | 6.138734 | 8.32E-10 | 1.44E-08 | ENSG00000162444 | RBP7 | 116362 |
| ENSG00000174171.6 | 8.5864912 | 3.840484075 | 0.635182704 | 5.101657 | 3.37E-07 | 4.33E-06 | ENSG00000174171 | LOC10537 | 1.05E+08 |
| ENSG00000249378.6 | 19.59741197 | 3.840188229 | 0.467827782 | 6.926028 | 4.33E-12 | 9.37E-11 | ENSG00000249378 | LINC0106( | 401164 |
| ENSG00000029559.7 | 28.12460835 | 3.837775124 | 0.451231017 | 7.175427 | 7.21E-13 | 1.67E-11 | ENSG00000029559 | IBSP | 3381 |
| ENSG00000166317.12 | 2.778862854 | 3.836002525 | 1.198326953 | 2.700434 | 0.006925 | 0.041656562 | ENSG00000166317 | SYNPO2L | 79933 |
| ENSG00000178150.10 | 458.7043 | 3.832828 | 0.2043067 | 15.82341 | 2.15E-56 | 3.66E-54 | ENSG00000178150 | ZNF114 | 163071 |
| ENSG00000248323.7 | 761.9960376 | 3.832409505 | 0.246781004 | 13.09829 | 3.37E-39 | 3.26E-37 | ENSG00000248323 | LUCAT1 | 1.01E+08 |
| ENSG00000256576.2 | 10.95362258 | 3.830698863 | 0.876080449 | 3.687674 | 0.000226 | 0.001912066 | ENSG00000256576 | LINC0236 | 1.01E+08 |
| ENSG00000253125.1 | 101.3894276 | 3.825951423 | 0.241799842 | 13.34141 | 1.33E-40 | 1.36E-38 | ENSG00000253125 | NA | NA |
| ENSG00000280420.1 | 4.899586223 | 3.824584739 | 0.932362139 | 3.458511 | 0.000543 | 0.004271868 | ENSG00000280420 | NA | NA |
| ENSG00000163220.11 | 39676.34467 | 3.822264735 | 0.386438568 | 8.338362 | 7.53E-17 | 2.34E-15 | ENSG00000163220 | S100A9 | 6280 |
| ENSG00000183379.9 | 2.573855062 | 3.82128181 | 1.050356106 | 3.066847 | 0.002163 | 0.014839266 | ENSG00000183379 | SYNDIG1L | 646658 |
| ENSG00000279489.1 | 2.70721618 | 3.819961208 | 1.081083707 | 2.978457 | 0.002897 | 0.019274621 | ENSG00000279489 | NA | NA |
| ENSG00000242578.1 | 2.549153871 | 3.818266309 | 1.118984754 | 2.876059 | 0.004027 | 0.025843767 | ENSG00000242578 | NA | NA |
| ENSG00000267665.2 | 2.555016944 | 3.81427165 | 1.073956444 | 2.992926 | 0.002763 | 0.018491995 | ENSG00000267665 | LOC40062 | 400622 |
| ENSG00000286507.1 | 3.62178413 | 3.803952009 | 0.96012482 | 3.337016 | 0.000847 | 0.006390113 | ENSG00000286507 | NA | NA |
| ENSG00000147481.17 | 1.802232123 | 3.799267202 | 1.105283623 | 2.894521 | 0.003797 | 0.024466012 | ENSG00000147481 | SNTG1 | 54212 |
| ENSG00000115705.21 | 2.60128129 | 3.799144107 | 1.040585738 | 3.074369 | 0.002109 | 0.014492579 | ENSG00000115705 | TPO | 7173 |
| ENSG00000287868.1 | 2.790355483 | 3.798854718 | 1.065893288 | 3.001102 | 0.00269 | 0.018061616 | ENSG00000287868 | NA | NA |
| ENSG00000255750.6 | 2.795076932 | 3.796862843 | 1.109117167 | 2.882349 | 0.003947 | 0.025363972 | ENSG00000255750 | NA | NA |
| ENSG00000183615.6 | 136.1179678 | 3.795382346 | 0.236503437 | 13.51093 | 1.35E-41 | 1.42E-39 | ENSG00000183615 | FAM167B | 84734 |
| ENSG00000220154.2 | 3.426815488 | 3.791794882 | 1.095393645 | 2.913834 | 0.00357 | 0.02317627 | ENSG00000220154 | NA | NA |
| ENSG00000229938.1 | 7.45135708 | 3.786839493 | 0.744485849 | 4.280591 | 1.86E-05 | 0.000188351 | ENSG00000229938 | NA | NA |
| ENSG00000203801.8 | 6.961319695 | 3.784152476 | 0.73596768 | 4.326484 | 1.52E-05 | 0.000155432 | ENSG00000203801 | LINC0022 | 387111 |
| ENSG00000204866.8 | 44.97531 | 3.783499 | 0.441324 | 7.213518 | 5.45E-13 | 1.28E-11 | ENSG00000204866 | IGFL2 | 147920 |
| ENSG00000167861.16 | 157.6543452 | 3.780989831 | 0.511288608 | 6.221515 | 4.92E-10 | 8.81E-09 | ENSG00000167861 | HID1 | 283987 |
| ENSG00000154553.15 | 186.4463439 | 3.778610824 | 0.374534127 | 8.486839 | 2.12E-17 | 6.92E-16 | ENSG00000154553 | PDLIM3 | 27295 |
| ENSG00000286241.1 | 1.20213761 | 3.777128605 | 1.201566124 | 2.644156 | 0.008189 | 0.048202616 | ENSG00000286241 | NA | NA |
| ENSG00000226835.3 | 2.488728657 | 3.776591848 | 1.041642567 | 3.049599 | 0.002291 | 0.015625921 | ENSG00000226835 | ARHGAP2 | 1.08E+08 |
| ENSG00000113805.8 | 2.61174818 | 3.776160626 | 1.09375173 | 2.903914 | 0.003685 | 0.023813104 | ENSG00000113805 | CNTN3 | 5067 |
| ENSG00000229660.1 | 3.470808787 | 3.770123524 | 1.140841238 | 2.77876 | 0.005457 | 0.033798973 | ENSG00000229660 | NA | NA |
| ENSG00000260828.1 | 1.21399049 | 3.769342876 | 1.196270805 | 2.649352 | 0.008065 | 0.047541257 | ENSG00000260828 | HMGB3P3 | 1.01E+08 |

| | | | | | | | | | |
|---|---|---|---|---|---|---|---|---|---|
| ENSG00000214128.11 | 2.486418031 | 3.767563275 | 1.084607048 | 2.920471 | 0.003495 | 0.022766004 | ENSG00000214128 | TMEM213 | 155006 |
| ENSG00000095370.20 | 4.635303884 | 3.764622306 | 0.865787538 | 3.655195 | 0.000257 | 0.002146688 | ENSG00000095370 | SH2D3C | 10044 |
| ENSG00000235899.2 | 13.48977354 | 3.764035626 | 0.651935169 | 4.853298 | 1.21E-06 | 1.45E-05 | ENSG00000235899 | LINC0156 | 1.02E+08 |
| ENSG00000204581.3 | 16.65316513 | 3.763578978 | 0.512623489 | 6.17135 | 6.77E-10 | 1.19E-08 | ENSG00000204581 | ACOXL-AS | 400997 |
| ENSG00000166592.12 | 1444.119625 | 3.761755878 | 0.345272669 | 9.157272 | 5.32E-20 | 2.03E-18 | ENSG00000166592 | RRAD | 6236 |
| ENSG00000232164.1 | 6.894186429 | 3.750909716 | 0.715169921 | 4.40582 | 1.05E-05 | 0.000111067 | ENSG00000232164 | LINC0187 | 729348 |
| ENSG00000228839.5 | 8.710601 | 3.749408 | 0.6214265 | 5.068029 | 4.02E-07 | 5.11E-06 | ENSG00000228839 | PIK3IP1-D | 1.02E+08 |
| ENSG00000286939.1 | 14.48127 | 3.747647 | 0.6261544 | 5.02695 | 4.98E-07 | 6.26E-06 | ENSG00000286939 | NA | NA |
| ENSG00000106018.14 | 4.917557612 | 3.746313442 | 0.904168893 | 3.479785 | 0.000502 | 0.003974177 | ENSG00000106018 | VIPR2 | 7434 |
| ENSG00000248690.8 | 23.37686174 | 3.746311708 | 0.443804244 | 7.089413 | 1.35E-12 | 3.04E-11 | ENSG00000248690 | NA | NA |
| ENSG00000174792.10 | 5.43890156 | 3.745749131 | 0.861132932 | 3.653035 | 0.000259 | 0.002163473 | ENSG00000174792 | ODAPH | 152816 |
| ENSG00000155886.11 | 38.95659999 | 3.743608972 | 0.434055902 | 7.242406 | 4.41E-13 | 1.04E-11 | ENSG00000155886 | SLC24A2 | 25769 |
| ENSG00000285269.2 | 3.424127373 | 3.741271213 | 1.02835617 | 3.054653 | 0.002253 | 0.015396555 | ENSG00000285269 | SLC35D2 | 11046 |
| ENSG00000175287.19 | 5.716820465 | 3.734668762 | 0.834992747 | 3.754127 | 0.000174 | 0.001501559 | ENSG00000175287 | PHYHD1 | 254295 |
| ENSG00000276900.1 | 87.44737538 | 3.733158042 | 0.278674233 | 11.24308 | 2.50E-29 | 1.66E-27 | ENSG00000276900 | NA | NA |
| ENSG00000111344.11 | 226.3934237 | 3.73282523 | 0.27056464 | 11.57884 | 5.28E-31 | 3.80E-29 | ENSG00000111344 | RASAL1 | 8437 |
| ENSG00000083067.22 | 4.200591791 | 3.732073545 | 0.906876824 | 3.453692 | 0.000553 | 0.004341229 | ENSG00000083067 | TRPM3 | 80036 |
| ENSG00000240541.3 | 4.226571832 | 3.731939608 | 0.907705002 | 3.450394 | 0.00056 | 0.004385576 | ENSG00000240541 | TM4SF1-A | 1.01E+08 |
| ENSG00000188732.11 | 4.732518422 | 3.730262046 | 0.903900281 | 3.463061 | 0.000534 | 0.004203999 | ENSG00000188732 | FAM221A | 340277 |
| ENSG00000102837.7 | 27.15876781 | 3.730159726 | 0.353096232 | 8.864891 | 7.66E-19 | 2.73E-17 | ENSG00000102837 | OLFM4 | 10562 |
| ENSG00000163995.20 | 87.7076955 | 3.729519689 | 0.362777351 | 8.626558 | 6.32E-18 | 2.13E-16 | ENSG00000163995 | ABLIM2 | 84448 |
| ENSG00000071282.12 | 59.07858466 | 3.729070759 | 0.316445243 | 9.88819 | 4.68E-23 | 2.13E-21 | ENSG00000071282 | LMCD1 | 29995 |
| ENSG00000167614.13 | 4.561301 | 3.72884 | 0.8877057 | 3.524637 | 0.000424 | 0.003401451 | ENSG00000167614 | TTYH1 | 57348 |
| ENSG00000198535.5 | 12.46739853 | 3.725237694 | 0.547734636 | 5.705751 | 1.16E-08 | 1.77E-07 | ENSG00000198535 | C2CD4A | 145741 |
| ENSG00000136153.20 | 11078.21911 | 3.72282786 | 0.18207955 | 17.1509 | 6.19E-66 | 1.37E-63 | ENSG00000136153 | LMO7 | 4008 |
| ENSG00000189090.7 | 2.415876336 | 3.721992487 | 1.079393454 | 2.892358 | 0.003824 | 0.024617255 | ENSG00000189090 | FAM25G | 1E+08 |
| ENSG00000279217.1 | 3.459294 | 3.721309 | 1.000656 | 3.119263 | 0.001813 | 0.01264521 | ENSG00000279217 | NA | NA |
| ENSG00000224596.8 | 17.93378037 | 3.719170662 | 0.45731529 | 6.820613 | 9.07E-12 | 1.92E-10 | ENSG00000224596 | ZMIZ1-AS | 283050 |
| ENSG00000185168.5 | 32.61519168 | 3.717425596 | 0.323399318 | 9.639555 | 5.44E-22 | 2.34E-20 | ENSG00000185168 | NA | NA |
| ENSG00000150681.10 | 9.625640585 | 3.71720087 | 0.708826075 | 4.397695 | 1.09E-05 | 0.000114942 | ENSG00000150681 | RGS18 | 64407 |
| ENSG00000069535.14 | 32.19353 | 3.7153 | 0.4312772 | 7.223428 | 5.07E-13 | 1.19E-11 | ENSG00000069535 | MAOB | 4129 |
| ENSG00000279311.1 | 1.691520432 | 3.712546157 | 1.124178628 | 2.768729 | 0.005628 | 0.034692436 | ENSG00000279311 | NA | NA |
| ENSG00000262075.3 | 2.413454067 | 3.711678562 | 1.105959694 | 2.813555 | 0.0049 | 0.030727862 | ENSG00000262075 | DKFZP434 | 26102 |
| ENSG00000273443.1 | 32.6508584 | 3.710562328 | 0.410414856 | 7.579069 | 3.48E-14 | 8.92E-13 | ENSG00000273443 | NA | NA |
| ENSG00000253877.6 | 2.803598858 | 3.70987349 | 1.066275561 | 2.916576 | 0.003539 | 0.022990334 | ENSG00000253877 | LINC0160 | 1.02E+08 |
| ENSG00000231574.6 | 3.873877596 | 3.705225468 | 1.069236322 | 2.904153 | 0.003682 | 0.023806189 | ENSG00000231574 | LINC0201 | 1.03E+08 |
| ENSG00000110934.11 | 2.583033846 | 3.704090885 | 1.162815694 | 2.669461 | 0.007597 | 0.045136289 | ENSG00000110934 | BIN2 | 51411 |
| ENSG00000286699.1 | 290.8066697 | 3.699310495 | 0.201831142 | 15.35596 | 3.23E-53 | 5.06E-51 | ENSG00000286699 | NA | NA |
| ENSG00000167549.18 | 529.9551773 | 3.695096019 | 0.472464976 | 6.550953 | 5.72E-11 | 1.11E-09 | ENSG00000167549 | CORO6 | 84940 |
| ENSG00000164825.4 | 16.03415261 | 3.694048615 | 0.628461047 | 4.923215 | 8.51E-07 | 1.04E-05 | ENSG00000164825 | DEFB1 | 1672 |
| ENSG00000123201.14 | 7.688221311 | 3.692834004 | 0.759482516 | 4.072291 | 4.66E-05 | 0.000443252 | ENSG00000123201 | GUCY1B2 | 2974 |
| ENSG00000208028.1 | 18.9579507 | 3.692675347 | 0.450790746 | 6.860556 | 6.86E-12 | 1.47E-10 | ENSG00000208028 | MIR616 | 693201 |

| | | | | | | | | | |
|---|---|---|---|---|---|---|---|---|---|
| ENSG00000124731.13 | 226.0536881 | 3.69073249 | 0.301177967 | 10.26215 | 1.04E-24 | 5.15E-23 | ENSG00000124731 | TREM1 | 54210 |
| ENSG00000016391.11 | 19.12616213 | 3.689884626 | 0.469645633 | 6.579183 | 4.73E-11 | 9.28E-10 | ENSG00000016391 | CHDH | 55349 |
| ENSG00000173335.4 | 1.756516 | 3.687943 | 1.087584 | 2.83927 | 0.004522 | 0.0285998 | ENSG00000173335 | CST9 | 128822 |
| ENSG00000167757.14 | 916.0556 | 3.687599 | 0.522976 | 5.903903 | 3.55E-09 | 5.76E-08 | ENSG00000167757 | KLK11 | 11012 |
| ENSG00000123405.14 | 160.601494 | 3.686296873 | 0.282765651 | 10.91468 | 9.81E-28 | 5.80E-26 | ENSG00000123405 | NFE2 | 4778 |
| ENSG00000269855.2 | 40.93464 | 3.68613 | 0.3961773 | 7.78977 | 6.71E-15 | 1.82E-13 | ENSG00000269855 | RNF225 | 646862 |
| ENSG00000198074.10 | 214.018621 | 3.684937545 | 0.220794595 | 13.97198 | 2.31E-44 | 2.75E-42 | ENSG00000198074 | AKR1B10 | 57016 |
| ENSG00000246225.7 | 6.565416849 | 3.684752945 | 0.699428515 | 4.410391 | 1.03E-05 | 0.000108964 | ENSG00000246225 | NA | NA |
| ENSG00000114166.8 | 4160.276907 | 3.684455568 | 0.218237929 | 14.13345 | 2.36E-45 | 2.86E-43 | ENSG00000114166 | KAT2B | 8850 |
| ENSG00000235244.4 | 4.281009 | 3.680511 | 0.8482732 | 3.631508 | 0.000282 | 0.002339767 | ENSG00000235244 | DANT2 | 642776 |
| ENSG00000266401.2 | 2.325770011 | 3.668545251 | 1.122730503 | 2.733109 | 0.006274 | 0.03814229 | ENSG00000266401 | LOC10537 | 1.05E+08 |
| ENSG00000182557.8 | 7.898905703 | 3.66479402 | 0.705859477 | 4.341932 | 1.41E-05 | 0.000145961 | ENSG00000182557 | SPNS3 | 201305 |
| ENSG00000234199.3 | 38.23590087 | 3.66418975 | 0.363756473 | 8.423739 | 3.65E-17 | 1.17E-15 | ENSG00000234199 | LINC0119 | 440900 |
| ENSG00000125788.6 | 6.776785 | 3.663965 | 0.7119994 | 4.303325 | 1.68E-05 | 0.000171291 | ENSG00000125788 | DEFB126 | 81623 |
| ENSG00000163239.12 | 8.945395448 | 3.663932248 | 0.655971187 | 4.670834 | 3.00E-06 | 3.40E-05 | ENSG00000163239 | TDRD10 | 126668 |
| ENSG00000162772.17 | 992.3374985 | 3.663702676 | 0.208649849 | 14.68346 | 8.23E-49 | 1.12E-46 | ENSG00000162772 | ATF3 | 467 |
| ENSG00000066813.14 | 1.632795833 | 3.659154962 | 1.132987232 | 2.700079 | 0.006932 | 0.041678356 | ENSG00000066813 | ACSM2B | 348158 |
| ENSG00000233081.1 | 8.888889653 | 3.658206552 | 0.774235196 | 3.949971 | 7.82E-05 | 0.000718506 | ENSG00000233081 | NA | NA |
| ENSG00000286042.1 | 352.5097959 | 3.657798234 | 0.163379591 | 18.71591 | 3.67E-78 | 1.05E-75 | ENSG00000286042 | LCAL1 | 80078 |
| ENSG00000285681.2 | 3.256408197 | 3.657706368 | 1.076660198 | 2.839992 | 0.004511 | 0.028555458 | ENSG00000285681 | NA | NA |
| ENSG00000162520.14 | 204.971728 | 3.657425944 | 0.235155321 | 13.00173 | 1.20E-38 | 1.13E-36 | ENSG00000162520 | SYNC | 81493 |
| ENSG00000158373.8 | 265.8589542 | 3.657324034 | 0.168181537 | 18.17871 | 7.61E-74 | 1.99E-71 | ENSG00000158373 | H2BC5 | 3017 |
| ENSG00000094796.5 | 10.77823088 | 3.653553335 | 0.566000931 | 5.394962 | 6.85E-08 | 9.61E-07 | ENSG00000094796 | KRT31 | 3881 |
| ENSG00000236039.3 | 5.232504034 | 3.652922785 | 0.812835909 | 3.755891 | 0.000173 | 0.001492475 | ENSG00000236039 | LINC0288 | 1.02E+08 |
| ENSG00000286782.1 | 2.346736936 | 3.641434697 | 1.074581038 | 2.830345 | 0.00465 | 0.0293253 | ENSG00000286782 | NA | NA |
| ENSG00000137709.10 | 103.2936135 | 3.640099338 | 0.358697138 | 8.475393 | 2.34E-17 | 7.61E-16 | ENSG00000137709 | POU2F3 | 25833 |
| ENSG00000258592.1 | 6.33542734 | 3.637873151 | 0.72686606 | 4.179413 | 2.92E-05 | 0.000286791 | ENSG00000258592 | NA | NA |
| ENSG00000249069.8 | 3.278420333 | 3.636767439 | 1.037887193 | 2.925913 | 0.003434 | 0.022398948 | ENSG00000249069 | LINC0103 | 1.04E+08 |
| ENSG00000149575.6 | 5.366592264 | 3.627736081 | 0.902026113 | 3.356595 | 0.000789 | 0.005998593 | ENSG00000149575 | SCN2B | 6327 |
| ENSG00000280924.1 | 5.408702791 | 3.62750161 | 0.917188503 | 3.30085 | 0.000964 | 0.007172049 | ENSG00000280924 | LINC0062 | 127841 |
| ENSG00000259039.3 | 6.84641964 | 3.625481117 | 0.882108713 | 3.429828 | 0.000604 | 0.004701384 | ENSG00000259039 | NA | NA |
| ENSG00000182489.9 | 129.0703 | 3.625252 | 0.2668752 | 11.33583 | 8.72E-30 | 5.95E-28 | ENSG00000182489 | XKRX | 402415 |
| ENSG00000272398.6 | 16954.84693 | 3.623779646 | 0.222965077 | 13.56167 | 6.76E-42 | 7.22E-40 | ENSG00000272398 | CD24 | 1E+08 |
| ENSG00000167748.11 | 61.65935 | 3.620929 | 0.4307656 | 7.01293 | 2.33E-12 | 5.17E-11 | ENSG00000167748 | KLK1 | 3816 |
| ENSG00000274180.1 | 726.1323249 | 3.61601307 | 0.236387254 | 12.75878 | 2.79E-37 | 2.56E-35 | ENSG00000274180 | NATD1 | 256302 |
| ENSG00000278611.2 | 8.471642892 | 3.614090077 | 0.676585923 | 4.454852 | 8.40E-06 | 9.01E-05 | ENSG00000278611 | ZNF426-D | 1.02E+08 |
| ENSG00000229463.2 | 2.219130728 | 3.609774984 | 1.140255821 | 2.639561 | 0.008301 | 0.048753228 | ENSG00000229463 | NA | NA |
| ENSG00000169962.5 | 73.48612237 | 3.608421091 | 0.265674886 | 11.32369 | 1.00E-29 | 6.82E-28 | ENSG00000169962 | TAS1R3 | 83756 |
| ENSG00000178538.10 | 2.237692901 | 3.605927225 | 1.073732187 | 2.799513 | 0.005118 | 0.031901368 | ENSG00000178538 | CA8 | 767 |
| ENSG00000080573.7 | 365.1952829 | 3.604088244 | 0.206761077 | 14.52927 | 7.91E-48 | 1.04E-45 | ENSG00000080573 | COL5A3 | 50509 |
| ENSG00000285244.1 | 11.06276301 | 3.603206814 | 0.612093234 | 4.906453 | 9.27E-07 | 1.13E-05 | ENSG00000285244 | DINOL | 1.09E+08 |
| ENSG00000126233.2 | 18.20046478 | 3.598672936 | 0.629395531 | 4.76437 | 1.89E-06 | 2.20E-05 | ENSG00000126233 | SLURP1 | 57152 |

| | | | | | | | | | |
|---|---|---|---|---|---|---|---|---|---|
| ENSG00000219387.2 | 2.438709696 | 3.597048244 | 1.067595553 | 2.807288 | 0.004996 | 0.031251166 | ENSG00000219387 | NA | NA |
| ENSG00000142623.11 | 43.16099861 | 3.594460005 | 0.336702057 | 8.893501 | 5.92E-19 | 2.12E-17 | ENSG00000142623 | PADI1 | 29943 |
| ENSG00000140519.14 | 7743.337585 | 3.594193447 | 0.450669032 | 6.643885 | 3.06E-11 | 6.12E-10 | ENSG00000140519 | RHCG | 51458 |
| ENSG00000013297.11 | 6.404577228 | 3.590759337 | 0.765749638 | 3.905662 | 9.40E-05 | 0.000852918 | ENSG00000013297 | CLDN11 | 5010 |
| ENSG00000115414.19 | 36221.80151 | 3.589484295 | 0.130116382 | 22.97546 | 8.20E-117 | 4.20E-114 | ENSG00000115414 | FN1 | 2335 |
| ENSG00000145428.14 | 23.02432274 | 3.589137243 | 0.491563688 | 6.080875 | 1.20E-09 | 2.03E-08 | ENSG00000145428 | RNF175 | 285533 |
| ENSG00000226690.8 | 25.71756739 | 3.588960611 | 0.513780313 | 5.817585 | 5.97E-09 | 9.44E-08 | ENSG00000226690 | LOC10272 | 1.03E+08 |
| ENSG00000249898.8 | 5.220406282 | 3.588594153 | 0.98007877 | 3.049341 | 0.002293 | 0.015631338 | ENSG00000249898 | NA | NA |
| ENSG00000228288.7 | 37.35392513 | 3.588042603 | 0.341336388 | 8.753953 | 2.06E-18 | 7.13E-17 | ENSG00000228288 | PCAT6 | 1.01E+08 |
| ENSG00000011465.17 | 24.46158705 | 3.586294785 | 0.425319453 | 7.021298 | 2.20E-12 | 4.88E-11 | ENSG00000011465 | DCN | 1634 |
| ENSG00000178734.5 | 5.906092245 | 3.584178096 | 0.767044731 | 3.890488 | 0.0001 | 0.000903747 | ENSG00000178734 | LMO7DN | 729420 |
| ENSG00000173926.6 | 385.0402799 | 3.583569253 | 0.30501228 | 9.7818 | 1.35E-22 | 5.93E-21 | ENSG00000173926 | MARCHF3 | 115123 |
| ENSG00000184697.7 | 11.3072877 | 3.582434025 | 0.60653632 | 4.917157 | 8.78E-07 | 1.07E-05 | ENSG00000184697 | CLDN6 | 9074 |
| ENSG00000197540.8 | 2.246475297 | 3.580606375 | 1.050693139 | 2.8368 | 0.004557 | 0.028787729 | ENSG00000197540 | GZMM | 3004 |
| ENSG00000251432.7 | 181.1695704 | 3.57994482 | 0.187590518 | 15.88537 | 8.00E-57 | 1.38E-54 | ENSG00000251432 | LINC0261! | 1.01E+08 |
| ENSG00000168477.19 | 42.38466146 | 3.578796243 | 0.395085714 | 7.53962 | 4.71E-14 | 1.20E-12 | ENSG00000168477 | TNXB | 7148 |
| ENSG00000258689.1 | 34.35952701 | 3.578159264 | 0.355818195 | 8.36989 | 5.77E-17 | 1.81E-15 | ENSG00000258689 | LINC0126( | 1.04E+08 |
| ENSG00000228069.1 | 3.072948822 | 3.577070813 | 0.999567956 | 2.978358 | 0.002898 | 0.019274621 | ENSG00000228069 | MTCO3P2 | 1.07E+08 |
| ENSG00000187837.3 | 422.0105445 | 3.575972811 | 0.219955055 | 13.52991 | 1.04E-41 | 1.11E-39 | ENSG00000187837 | H1-2 | 3006 |
| ENSG00000137033.11 | 72.47946661 | 3.57383578 | 0.299459382 | 9.930682 | 3.06E-23 | 1.41E-21 | ENSG00000137033 | IL33 | 90865 |
| ENSG00000253796.2 | 4.86194061 | 3.573459471 | 0.840438633 | 3.537985 | 0.000403 | 0.003248727 | ENSG00000253796 | NA | NA |
| ENSG00000104879.5 | 18.34355 | 3.571813 | 0.4848099 | 6.129852 | 8.80E-10 | 1.52E-08 | ENSG00000104879 | CKM | 1158 |
| ENSG00000004948.15 | 16.30954911 | 3.571328805 | 0.458564033 | 6.479638 | 9.19E-11 | 1.76E-09 | ENSG00000004948 | CALCR | 799 |
| ENSG00000185860.13 | 3.765202805 | 3.570963697 | 1.070950181 | 2.774138 | 0.005535 | 0.034195386 | ENSG00000185860 | CCDC190 | 339512 |
| ENSG00000287343.1 | 2.199409588 | 3.569108592 | 1.079538981 | 2.750349 | 0.005953 | 0.036425139 | ENSG00000287343 | NA | NA |
| ENSG00000012223.12 | 10.96424681 | 3.56802969 | 0.921826287 | 3.219728 | 0.001283 | 0.009277383 | ENSG00000012223 | LTF | 4057 |
| ENSG00000207568.1 | 7.682067752 | 3.564290382 | 0.732756371 | 4.045397 | 5.22E-05 | 0.000492244 | ENSG00000207568 | MIR203A | 406986 |
| ENSG00000160868.15 | 3.060499796 | 3.562014026 | 0.991898756 | 2.986206 | 0.002825 | 0.018846394 | ENSG00000160868 | CYP3A4 | 1576 |
| ENSG00000160712.13 | 3618.093171 | 3.560654437 | 0.329935636 | 8.97343 | 2.87E-19 | 1.05E-17 | ENSG00000160712 | IL6R | 3570 |
| ENSG00000225614.3 | 4.24641666 | 3.553168323 | 0.899946573 | 3.281493 | 0.001033 | 0.007633891 | ENSG00000225614 | ZNF469 | 84627 |
| ENSG00000153930.12 | 15.19826214 | 3.553140879 | 0.541081467 | 5.457849 | 4.82E-08 | 6.85E-07 | ENSG00000153930 | ANKFN1 | 162282 |
| ENSG00000147883.12 | 1921.085844 | 3.553054913 | 0.120790828 | 24.44768 | 5.33E-132 | 3.22E-129 | ENSG00000147883 | CDKN2B | 1030 |
| ENSG00000185933.6 | 3.998780406 | 3.543855462 | 0.877892022 | 3.353323 | 0.000798 | 0.006063091 | ENSG00000185933 | CALHM1 | 255022 |
| ENSG00000184163.3 | 9.148098139 | 3.5389742 | 0.782915831 | 3.753883 | 0.000174 | 0.001502537 | ENSG00000184163 | C1QTNF1: | 388581 |
| ENSG00000174502.19 | 4.763859875 | 3.538785417 | 1.06812661 | 2.751346 | 0.005935 | 0.036356241 | ENSG00000174502 | SLC26A9 | 115019 |
| ENSG00000278958.1 | 2.91959007 | 3.533848321 | 1.032727223 | 2.840874 | 0.004499 | 0.028503678 | ENSG00000278958 | NA | NA |
| ENSG00000166920.12 | 804.6648288 | 3.532286779 | 0.375108873 | 7.817162 | 5.40E-15 | 1.48E-13 | ENSG00000166920 | C15orf48 | 84419 |
| ENSG00000251442.6 | 264.0971738 | 3.530637044 | 0.215627661 | 13.59119 | 4.52E-42 | 4.85E-40 | ENSG00000251442 | NA | NA |
| ENSG00000171345.13 | 22.64980419 | 3.529650944 | 0.438096654 | 6.687225 | 2.27E-11 | 4.61E-10 | ENSG00000171345 | KRT19 | 3880 |
| ENSG00000166046.11 | 1228.628931 | 3.52408329 | 0.225621174 | 12.96015 | 2.06E-38 | 1.93E-36 | ENSG00000166046 | TCP11L2 | 255394 |
| ENSG00000255346.10 | 19.06458902 | 3.52048237 | 0.440785403 | 6.625633 | 3.46E-11 | 6.87E-10 | ENSG00000255346 | NOX5 | 79400 |
| ENSG00000189221.9 | 473.498 | 3.520472 | 0.1668252 | 17.50618 | 1.29E-68 | 3.03E-66 | ENSG00000189221 | MAOA | 4128 |

| | | | | | | | | | |
|---|---|---|---|---|---|---|---|---|---|
| ENSG00000090238.11 | 1359.981181 | 3.519775696 | 0.224352355 | 13.01424 | 1.02E-38 | 9.69E-37 | ENSG00000090238 | YPEL3 | 83719 |
| ENSG00000118785.14 | 2.268344381 | 3.517724965 | 1.090984333 | 2.674397 | 0.007486 | 0.044556778 | ENSG00000118785 | SPP1 | 6696 |
| ENSG00000213755.3 | 7.141361748 | 3.517645778 | 0.759671623 | 3.840667 | 0.000123 | 0.001089207 | ENSG00000213755 | NA | NA |
| ENSG00000249669.10 | 4.115149952 | 3.517487459 | 1.001811581 | 2.912212 | 0.003589 | 0.023285605 | ENSG00000249669 | NA | NA |
| ENSG00000125740.14 | 355.9102 | 3.517389 | 0.1869153 | 15.60808 | 6.41E-55 | 1.05E-52 | ENSG00000125740 | FOSB | 2354 |
| ENSG00000168589.15 | 5.044316791 | 3.515453949 | 0.863586695 | 3.375983 | 0.000736 | 0.005631137 | ENSG00000168589 | DYNLRB2 | 83657 |
| ENSG00000258183.6 | 3.918978241 | 3.515429005 | 0.972716734 | 2.997202 | 0.002725 | 0.018252863 | ENSG00000258183 | LINC0239 | 1.05E+08 |
| ENSG00000129009.13 | 44.12346671 | 3.514708743 | 0.575742724 | 5.062519 | 4.14E-07 | 5.25E-06 | ENSG00000129009 | ISLR | 3671 |
| ENSG00000171476.22 | 255.4961297 | 3.512875519 | 0.16621073 | 17.5252 | 9.20E-69 | 2.19E-66 | ENSG00000171476 | HOPX | 84525 |
| ENSG00000123342.16 | 564.9037418 | 3.51077152 | 0.462008326 | 6.300258 | 2.97E-10 | 5.47E-09 | ENSG00000123342 | MMP19 | 4327 |
| ENSG00000142549.9 | 3.922947 | 3.509877 | 0.8739788 | 3.32946 | 0.00087 | 0.00654565 | ENSG00000142549 | IGLON5 | 402665 |
| ENSG00000100344.10 | 8.30067 | 3.507599 | 0.6620303 | 4.391943 | 1.12E-05 | 0.000117979 | ENSG00000100344 | PNPLA3 | 80339 |
| ENSG00000139988.9 | 13.79057953 | 3.501706002 | 0.5863282 | 4.948945 | 7.46E-07 | 9.20E-06 | ENSG00000139988 | RDH12 | 145226 |
| ENSG00000088726.16 | 7863.251317 | 3.499391888 | 0.39170505 | 7.401977 | 1.34E-13 | 3.32E-12 | ENSG00000088726 | TMEM40 | 55287 |
| ENSG00000077984.6 | 11.43186 | 3.498184 | 0.6601043 | 4.390494 | 1.13E-05 | 0.000118675 | ENSG00000077984 | CST7 | 8530 |
| ENSG00000187492.9 | 6.836218351 | 3.496619286 | 0.671995582 | 4.310474 | 1.63E-05 | 0.000166291 | ENSG00000187492 | CDHR4 | 389118 |
| ENSG00000259518.2 | 18.01263337 | 3.492191001 | 0.526913038 | 5.488934 | 4.04E-08 | 5.79E-07 | ENSG00000259518 | LINC0158 | 1.02E+08 |
| ENSG00000249679.2 | 3.221312936 | 3.487862293 | 1.04581819 | 2.761343 | 0.005756 | 0.03537571 | ENSG00000249679 | LOC10537 | 1.05E+08 |
| ENSG00000104765.16 | 12299.14121 | 3.487574252 | 0.243606009 | 11.85346 | 2.06E-32 | 1.57E-30 | ENSG00000104765 | BNIP3L | 665 |
| ENSG00000129990.15 | 2.839204 | 3.485536 | 1.079303 | 2.673517 | 0.007506 | 0.04466383 | ENSG00000129990 | SYT5 | 6861 |
| ENSG00000144476.6 | 3.509484337 | 3.484498838 | 0.975963541 | 2.95554 | 0.003121 | 0.020588033 | ENSG00000144476 | ACKR3 | 57007 |
| ENSG00000236750.1 | 3.560323195 | 3.48330752 | 0.952189921 | 3.02808 | 0.002461 | 0.016663307 | ENSG00000236750 | NA | NA |
| ENSG00000223361.5 | 2.918889901 | 3.482213296 | 1.012404472 | 2.846899 | 0.004415 | 0.028063303 | ENSG00000223361 | FTH1P10 | 2502 |
| ENSG00000158008.9 | 8.025673153 | 3.477566661 | 1.060586273 | 2.713185 | 0.006664 | 0.040292703 | ENSG00000158008 | EXTL1 | 2134 |
| ENSG00000164946.19 | 9.848640968 | 3.475862291 | 0.577104659 | 4.98326 | 6.25E-07 | 7.78E-06 | ENSG00000164946 | FREM1 | 158326 |
| ENSG00000230408.3 | 11.33546721 | 3.475786195 | 0.566377031 | 5.077512 | 3.82E-07 | 4.88E-06 | ENSG00000230408 | BZW1-AS | 1.02E+08 |
| ENSG00000164488.12 | 90.02556844 | 3.473559045 | 0.384822574 | 7.467231 | 8.19E-14 | 2.05E-12 | ENSG00000164488 | DACT2 | 168002 |
| ENSG00000229272.2 | 2.81775723 | 3.473399014 | 1.021722196 | 2.812309 | 0.004919 | 0.030803585 | ENSG00000229272 | NA | NA |
| ENSG00000249706.1 | 2.057017408 | 3.473293267 | 1.062953094 | 2.703123 | 0.006869 | 0.041354563 | ENSG00000249706 | NA | NA |
| ENSG00000283265.1 | 18.23665931 | 3.468721388 | 0.449953368 | 6.375597 | 1.82E-10 | 3.41E-09 | ENSG00000283265 | NA | NA |
| ENSG00000198483.13 | 17.8400273 | 3.468022968 | 0.520874955 | 5.506164 | 3.67E-08 | 5.28E-07 | ENSG00000198483 | ANKRD35 | 148741 |
| ENSG00000226440.7 | 4.955706967 | 3.463924832 | 0.843277511 | 3.396183 | 0.000683 | 0.005266796 | ENSG00000226440 | NA | NA |
| ENSG00000241336.2 | 3.745157184 | 3.46167208 | 0.891646722 | 3.209424 | 0.00133 | 0.009582577 | ENSG00000241336 | LINC0148 | 1.02E+08 |
| ENSG00000261371.6 | 53.15010852 | 3.459345014 | 0.246455304 | 11.60188 | 4.03E-31 | 2.91E-29 | ENSG00000261371 | PECAM1 | 5175 |
| ENSG00000286089.1 | 5.491866373 | 3.458515922 | 0.759056982 | 3.765878 | 0.000166 | 0.001438723 | ENSG00000286089 | NA | NA |
| ENSG00000119686.10 | 53.83294379 | 3.458394182 | 0.276147394 | 10.35097 | 4.14E-25 | 2.12E-23 | ENSG00000119686 | FLVCR2 | 55640 |
| ENSG00000206538.9 | 2061.884301 | 3.457625302 | 0.324843003 | 8.796943 | 1.41E-18 | 4.91E-17 | ENSG00000206538 | VGLL3 | 389136 |
| ENSG00000203785.9 | 5185.74017 | 3.457160411 | 0.425345778 | 6.717265 | 1.85E-11 | 3.79E-10 | ENSG00000203785 | SPRR2E | 6704 |
| ENSG00000285743.1 | 8.947576583 | 3.456795606 | 0.63717057 | 4.483565 | 7.34E-06 | 7.95E-05 | ENSG00000285743 | NA | NA |
| ENSG00000171604.12 | 984.3696549 | 3.456237129 | 0.097012979 | 29.4418 | 1.60E-190 | 1.71E-187 | ENSG00000171604 | CXXC5 | 51523 |
| ENSG00000277406.2 | 16.00353542 | 3.45618887 | 0.647263203 | 4.412716 | 1.02E-05 | 0.000107928 | ENSG00000277406 | NA | NA |
| ENSG00000168481.9 | 8.261957991 | 3.45587405 | 0.679455786 | 4.203179 | 2.63E-05 | 0.000260183 | ENSG00000168481 | LGI3 | 203190 |

| | | | | | | | | | |
|---|---|---|---|---|---|---|---|---|---|
| ENSG00000237292.1 | 7.482275778 | 3.452407272 | 0.638630077 | 4.466447 | 7.95E-06 | 8.56E-05 | ENSG00000237292 | LINC01732 | 1.02E+08 |
| ENSG00000114115.9 | 428.7024028 | 3.452214294 | 0.318268585 | 8.961658 | 3.20E-19 | 1.17E-17 | ENSG00000114115 | RBP1 | 5947 |
| ENSG00000120328.6 | 5.752537102 | 3.44888352 | 0.861178588 | 3.308122 | 0.000939 | 0.007007973 | ENSG00000120328 | PCDHB12 | 56124 |
| ENSG00000268941.2 | 6.666303 | 3.44806 | 0.7205565 | 3.952584 | 7.73E-05 | 0.000711193 | ENSG00000268941 | LINC0171 | 79160 |
| ENSG00000236324.1 | 9.500728985 | 3.44789417 | 0.580487337 | 4.90604 | 9.29E-07 | 1.13E-05 | ENSG00000236324 | NA | NA |
| ENSG00000116701.14 | 4020.161139 | 3.445092186 | 0.315330018 | 9.022586 | 1.84E-19 | 6.79E-18 | ENSG00000116701 | NCF2 | 4688 |
| ENSG00000166535.20 | 3423.252514 | 3.443072893 | 0.423970238 | 6.705831 | 2.00E-11 | 4.09E-10 | ENSG00000166535 | A2ML1 | 144568 |
| ENSG00000220867.1 | 14.16782054 | 3.441906943 | 0.47339259 | 6.003277 | 1.93E-09 | 3.22E-08 | ENSG00000220867 | HSPE1P26 | 1.06E+08 |
| ENSG00000226453.2 | 57.3294457 | 3.441199947 | 0.27774226 | 10.22963 | 1.46E-24 | 7.16E-23 | ENSG00000226453 | NA | NA |
| ENSG00000150722.10 | 247.9951345 | 3.436221876 | 0.227364516 | 12.47434 | 1.03E-35 | 8.77E-34 | ENSG00000150722 | PPP1R1C | 151242 |
| ENSG00000143382.14 | 1071.442446 | 3.436137352 | 0.27159885 | 10.44238 | 1.59E-25 | 8.34E-24 | ENSG00000143382 | ADAMTSL | 54507 |
| ENSG00000166396.13 | 1479.117769 | 3.431336783 | 0.115001111 | 24.62008 | 7.70E-134 | 4.77E-131 | ENSG00000166396 | SERPINB7 | 8710 |
| ENSG00000180573.9 | 757.7283027 | 3.428652186 | 0.218494175 | 12.94612 | 2.47E-38 | 2.31E-36 | ENSG00000180573 | H2AC6 | 8334 |
| ENSG00000161031.13 | 8.70603951 | 3.422120612 | 0.670199959 | 4.210864 | 2.54E-05 | 0.000251767 | ENSG00000161031 | PGLYRP2 | 114770 |
| ENSG00000170807.12 | 11.91083069 | 3.420670429 | 0.577968139 | 4.880322 | 1.06E-06 | 1.28E-05 | ENSG00000170807 | LMOD2 | 442721 |
| ENSG00000099139.13 | 72.69050822 | 3.419245509 | 0.294533257 | 9.571909 | 1.05E-21 | 4.43E-20 | ENSG00000099139 | PCSK5 | 5125 |
| ENSG00000106302.9 | 20.93053877 | 3.417149022 | 0.397162662 | 7.093187 | 1.31E-12 | 2.97E-11 | ENSG00000106302 | HYAL4 | 23553 |
| ENSG00000197506.7 | 223.8784651 | 3.415057916 | 0.346876945 | 8.115437 | 4.84E-16 | 1.42E-14 | ENSG00000197506 | SLC28A3 | 64078 |
| ENSG00000179477.11 | 168.0496196 | 3.414993722 | 0.476041526 | 5.913336 | 3.35E-09 | 5.46E-08 | ENSG00000179477 | ALOX12B | 242 |
| ENSG00000253220.2 | 2.279393371 | 3.412027238 | 1.02772307 | 2.736172 | 0.006216 | 0.037858238 | ENSG00000253220 | NA | NA |
| ENSG00000270547.6 | 88.16137828 | 3.410098317 | 0.386797571 | 7.265036 | 3.73E-13 | 8.84E-12 | ENSG00000270547 | LINC0123 | 401492 |
| ENSG00000213144.2 | 19.77662044 | 3.409068273 | 0.535434887 | 5.24633 | 1.55E-07 | 2.09E-06 | ENSG00000213144 | NA | NA |
| ENSG00000033327.13 | 493.7556518 | 3.408508961 | 0.193496917 | 14.51449 | 9.81E-48 | 1.28E-45 | ENSG00000033327 | GAB2 | 9846 |
| ENSG00000137573.14 | 15.09476657 | 3.406865876 | 0.521303567 | 5.384321 | 7.27E-08 | 1.02E-06 | ENSG00000137573 | SULF1 | 23213 |
| ENSG00000133135.14 | 98.00922 | 3.40559 | 0.3381685 | 8.296427 | 1.07E-16 | 3.30E-15 | ENSG00000133135 | RNF128 | 79589 |
| ENSG00000141431.12 | 3.35207035 | 3.401157076 | 1.061767301 | 2.638202 | 0.008335 | 0.048890127 | ENSG00000141431 | ASXL3 | 80816 |
| ENSG00000102886.15 | 197.757175 | 3.399053603 | 0.19765093 | 14.1616 | 1.58E-45 | 1.96E-43 | ENSG00000102886 | GDPD3 | 79153 |
| ENSG00000139287.13 | 2.7802599 | 3.398093929 | 1.007529659 | 2.777183 | 0.005483 | 0.033939707 | ENSG00000139287 | TPH2 | 121278 |
| ENSG00000178404.9 | 4.703775643 | 3.397206278 | 0.933150008 | 2.997596 | 0.002721 | 0.018238521 | ENSG00000178404 | CEP295NL | 1.01E+08 |
| ENSG00000227218.7 | 13.97582871 | 3.397009887 | 0.480956135 | 5.81552 | 6.04E-09 | 9.55E-08 | ENSG00000227218 | NA | NA |
| ENSG00000169583.12 | 466.2928784 | 3.396087351 | 0.310591173 | 9.002469 | 2.21E-19 | 8.14E-18 | ENSG00000169583 | CLIC3 | 9022 |
| ENSG00000227154.4 | 10.87453133 | 3.395649804 | 0.740041513 | 3.777693 | 0.000158 | 0.001377391 | ENSG00000227154 | NA | NA |
| ENSG00000188488.14 | 16.14698851 | 3.391226133 | 0.521287732 | 5.354483 | 8.58E-08 | 1.19E-06 | ENSG00000188488 | SERPINA5 | 5104 |
| ENSG00000183019.7 | 5.692594449 | 3.390359482 | 0.741767036 | 3.761773 | 0.000169 | 0.001460167 | ENSG00000183019 | MCEMP1 | 199675 |
| ENSG00000174473.16 | 2.763057133 | 3.387329079 | 0.986567934 | 2.825278 | 0.004724 | 0.029751865 | ENSG00000174473 | GALNTL6 | 442117 |
| ENSG00000238133.6 | 73.01864735 | 3.386946848 | 0.315656212 | 8.829058 | 1.06E-18 | 3.73E-17 | ENSG00000238133 | MAP3K20- | 339751 |
| ENSG00000124391.4 | 13.59423191 | 3.378923489 | 0.543878292 | 5.109458 | 3.23E-07 | 4.17E-06 | ENSG00000124391 | IL17C | 27189 |
| ENSG00000138316.11 | 118.6901947 | 3.378054012 | 0.282604697 | 9.830176 | 8.35E-23 | 3.73E-21 | ENSG00000138316 | ADAMTS1 | 140766 |
| ENSG00000233397.3 | 4.11003353 | 3.373495989 | 1.026157464 | 2.702798 | 0.006876 | 0.041385737 | ENSG00000233397 | NA | NA |
| ENSG00000168907.13 | 13.33927632 | 3.373189537 | 0.517918764 | 5.354487 | 8.58E-08 | 1.19E-06 | ENSG00000168907 | PLA2G4F | 255189 |
| ENSG00000115616.2 | 29.29417502 | 3.367847601 | 0.386033605 | 7.169965 | 7.50E-13 | 1.74E-11 | ENSG00000115616 | SLC9A2 | 6549 |
| ENSG00000225329.3 | 2.799602716 | 3.366416518 | 1.03094869 | 2.68337 | 0.007288 | 0.043573489 | ENSG00000225329 | LHFPL3-A | 723809 |

| | | | | | | | | | |
|---|---|---|---|---|---|---|---|---|---|
| ENSG00000228451.4 | 114.6444813 | 3.363787685 | 0.335792693 | 8.230637 | 1.86E-16 | 5.65E-15 | ENSG00000228451 | SDAD1P1 | 157489 |
| ENSG00000188460.4 | 3.710827472 | 3.362659833 | 1.002987213 | 2.754432 | 0.005879 | 0.036056619 | ENSG00000188460 | NA | NA |
| ENSG00000226421.1 | 54.47545398 | 3.362591925 | 0.254777434 | 10.84316 | 2.15E-27 | 1.26E-25 | ENSG00000226421 | SLC25A5F | 442525 |
| ENSG00000171711.2 | 64.47317494 | 3.36240563 | 0.384230543 | 7.189448 | 6.51E-13 | 1.51E-11 | ENSG00000171711 | DEFB4A | 1673 |
| ENSG00000108242.13 | 3.830984464 | 3.362362825 | 0.963316457 | 2.867555 | 0.004137 | 0.026478426 | ENSG00000108242 | CYP2C18 | 1562 |
| ENSG00000171658.8 | 111.5096775 | 3.357941402 | 0.289894332 | 9.513609 | 1.84E-21 | 7.66E-20 | ENSG00000171658 | NMRAL2P | 344887 |
| ENSG00000114013.16 | 23.10702026 | 3.356851207 | 0.525163965 | 5.249506 | 1.53E-07 | 2.06E-06 | ENSG00000114013 | CD86 | 942 |
| ENSG00000123496.8 | 274.5715 | 3.356644 | 0.4757245 | 5.794623 | 6.85E-09 | 1.08E-07 | ENSG00000123496 | IL13RA2 | 3598 |
| ENSG00000272508.1 | 45.86747697 | 3.35490975 | 0.34959516 | 7.880286 | 3.27E-15 | 9.13E-14 | ENSG00000272508 | NA | NA |
| ENSG00000123358.20 | 285.7725592 | 3.353768661 | 0.243690544 | 11.30027 | 1.31E-29 | 8.84E-28 | ENSG00000123358 | NR4A1 | 3164 |
| ENSG00000134755.16 | 20681.66396 | 3.352487179 | 0.208446932 | 13.20474 | 8.24E-40 | 8.21E-38 | ENSG00000134755 | DSC2 | 1824 |
| ENSG00000129451.12 | 4439.79 | 3.350911 | 0.4480641 | 6.139547 | 8.28E-10 | 1.44E-08 | ENSG00000129451 | KLK10 | 5655 |
| ENSG00000157514.16 | 1238.703 | 3.349104 | 0.08143526 | 33.75815 | 8.12E-250 | 1.20E-246 | ENSG00000157514 | TSC22D3 | 1831 |
| ENSG00000237222.4 | 7.274713355 | 3.345767262 | 0.712813593 | 3.852013 | 0.000117 | 0.00104482 | ENSG00000237222 | LINC0196ℓ | 1.01E+08 |
| ENSG00000113073.15 | 18.74237544 | 3.34546943 | 0.565089737 | 4.858466 | 1.18E-06 | 1.41E-05 | ENSG00000113073 | SLC4A9 | 83697 |
| ENSG00000229373.9 | 27.35614687 | 3.345364957 | 0.369370576 | 7.432549 | 1.07E-13 | 2.65E-12 | ENSG00000229373 | LINC0045፡ | 643365 |
| ENSG00000069667.16 | 261.9668266 | 3.337303555 | 0.291787271 | 9.381162 | 6.52E-21 | 2.66E-19 | ENSG00000069667 | RORA | 6095 |
| ENSG00000177575.12 | 5.351219413 | 3.337035408 | 0.87444019 | 3.130043 | 0.001748 | 0.012251216 | ENSG00000177575 | CD163 | 9332 |
| ENSG00000236212.2 | 21.1876217 | 3.331854935 | 0.382956159 | 7.133597 | 9.78E-13 | 2.24E-11 | ENSG00000236212 | NA | NA |
| ENSG00000197588.9 | 13.42182 | 3.330865 | 0.550178 | 4.963603 | 6.92E-07 | 8.55E-06 | ENSG00000197588 | KLKP1 | 606293 |
| ENSG00000081277.12 | 31595.51622 | 3.329541385 | 0.329774501 | 8.276993 | 1.26E-16 | 3.87E-15 | ENSG00000081277 | PKP1 | 5317 |
| ENSG00000130775.16 | 730.0521437 | 3.328856459 | 0.218347503 | 12.49777 | 7.68E-36 | 6.59E-34 | ENSG00000130775 | THEMIS2 | 9473 |
| ENSG00000153531.13 | 475.2296084 | 3.3239406 | 0.420239311 | 6.48188 | 9.06E-11 | 1.73E-09 | ENSG00000153531 | ADPRHL1 | 113622 |
| ENSG00000233143.2 | 4.422913317 | 3.323769166 | 0.850594693 | 3.202194 | 0.001364 | 0.009786645 | ENSG00000233143 | NA | NA |
| ENSG00000279588.2 | 6.203475837 | 3.323470289 | 0.74906572 | 3.635823 | 0.000277 | 0.002303602 | ENSG00000279588 | NA | NA |
| ENSG00000164949.8 | 306.4643139 | 3.317634591 | 0.260437355 | 10.43489 | 1.72E-25 | 9.00E-24 | ENSG00000164949 | GEM | 2669 |
| ENSG00000086544.3 | 5229.588 | 3.309574 | 0.2086809 | 12.98429 | 1.50E-38 | 1.41E-36 | ENSG00000086544 | ITPKC | 80271 |
| ENSG00000271369.1 | 8.644315399 | 3.308987516 | 0.598742817 | 4.524459 | 6.06E-06 | 6.63E-05 | ENSG00000271369 | NA | NA |
| ENSG00000260220.6 | 3.580750404 | 3.304804982 | 0.893869153 | 3.025952 | 0.002479 | 0.016776788 | ENSG00000260220 | CCDC187 | 399693 |
| ENSG00000235277.1 | 24.45846 | 3.302962 | 0.4047 | 6.678928 | 2.41E-11 | 4.87E-10 | ENSG00000235277 | NA | NA |
| ENSG00000250286.2 | 13.57041447 | 3.302569287 | 0.708596106 | 3.813977 | 0.000137 | 0.001203474 | ENSG00000250286 | LOC10537 | 1.05E+08 |
| ENSG00000146276.11 | 22.29470862 | 3.30201491 | 0.426479607 | 6.335625 | 2.36E-10 | 4.38E-09 | ENSG00000146276 | GABRR1 | 2569 |
| ENSG00000249094.2 | 4.481995722 | 3.301918386 | 0.833143813 | 3.24304 | 0.001183 | 0.008613857 | ENSG00000249094 | NA | NA |
| ENSG00000154856.13 | 5.387048557 | 3.297770237 | 0.746622843 | 3.613297 | 0.000302 | 0.002495066 | ENSG00000154856 | APCDD1 | 147495 |
| ENSG00000162241.12 | 114.9737693 | 3.294077792 | 0.181347371 | 14.8559 | 6.37E-50 | 8.97E-48 | ENSG00000162241 | SLC25A45 | 283130 |
| ENSG00000283849.1 | 9.772182989 | 3.293849696 | 0.582712566 | 4.622948 | 3.78E-06 | 4.24E-05 | ENSG00000283849 | NA | NA |
| ENSG00000166831.9 | 5.156504556 | 3.289877814 | 0.902742763 | 2.979673 | 0.002886 | 0.019219304 | ENSG00000166831 | RBPMS2 | 348093 |
| ENSG00000219951.4 | 3.435925267 | 3.287677469 | 0.888625022 | 3.024535 | 0.00249 | 0.016847002 | ENSG00000219951 | NA | NA |
| ENSG00000178075.20 | 1370.087992 | 3.287536555 | 0.276894544 | 9.705993 | 2.84E-22 | 1.24E-20 | ENSG00000178075 | GRAMD1C | 54762 |
| ENSG00000166869.3 | 3.42775096 | 3.284017944 | 0.966780449 | 2.776244 | 0.005499 | 0.034014186 | ENSG00000166869 | CHP2 | 63928 |
| ENSG00000130751.9 | 75.16458 | 3.281233 | 0.3667489 | 7.310812 | 2.66E-13 | 6.37E-12 | ENSG00000130751 | NPAS1 | 4861 |
| ENSG00000144227.5 | 5.640068179 | 3.280595747 | 0.816211386 | 3.284193 | 0.001023 | 0.007569533 | ENSG00000144227 | NXPH2 | 11249 |

| | | | | | | | | | |
|---|---|---|---|---|---|---|---|---|---|
| ENSG00000157423.18 | 3.592946592 | 3.280408362 | 0.988131637 | 2.712603 | 0.006676 | 0.040336094 | ENSG00000157423 | HYDIN | 54768 |
| ENSG00000137449.15 | 1515.451647 | 3.279461462 | 0.165208113 | 16.2187 | 3.72E-59 | 6.97E-57 | ENSG00000137449 | CPEB2 | 132864 |
| ENSG00000224945.1 | 34.1621965 | 3.276070537 | 0.451238714 | 5.930499 | 3.02E-09 | 4.94E-08 | ENSG00000224945 | NA | NA |
| ENSG00000167077.13 | 23.89328 | 3.276068 | 0.6390598 | 4.187508 | 2.82E-05 | 0.000277264 | ENSG00000167077 | MEI1 | 150365 |
| ENSG00000125571.9 | 6.587196025 | 3.273765893 | 0.688645305 | 3.882646 | 0.000103 | 0.000931193 | ENSG00000125571 | IL37 | 27178 |
| ENSG00000168903.8 | 15.87710423 | 3.271135153 | 0.624491674 | 4.277295 | 1.89E-05 | 0.000190913 | ENSG00000168903 | BTNL3 | 10917 |
| ENSG00000214814.7 | 9.203246646 | 3.267418458 | 0.557518642 | 4.784447 | 1.71E-06 | 2.01E-05 | ENSG00000214814 | FER1L6 | 654463 |
| ENSG00000231924.9 | 7.058215 | 3.262909 | 0.6562226 | 4.057935 | 4.95E-05 | 0.000468712 | ENSG00000231924 | PSG1 | 5669 |
| ENSG00000179148.9 | 1260.787633 | 3.254721852 | 0.226570536 | 11.71698 | 1.04E-31 | 7.74E-30 | ENSG00000179148 | ALOXE3 | 59344 |
| ENSG00000082175.14 | 59.62170324 | 3.253855083 | 0.423778799 | 6.262359 | 3.79E-10 | 6.87E-09 | ENSG00000082175 | PGR | 5241 |
| ENSG00000277075.2 | 39.11328661 | 3.253039961 | 0.43610531 | 6.083485 | 1.18E-09 | 2.00E-08 | ENSG00000277075 | H2AC8 | 3012 |
| ENSG00000186487.19 | 5.160241279 | 3.248346822 | 0.795877473 | 3.327581 | 0.000876 | 0.00658808 | ENSG00000186487 | MYT1L | 23040 |
| ENSG00000273117.1 | 102.7304615 | 3.239329644 | 0.174042272 | 15.16488 | 6.04E-52 | 8.98E-50 | ENSG00000273117 | NA | NA |
| ENSG00000137094.14 | 787.6104504 | 3.237322015 | 0.222435936 | 11.85655 | 1.99E-32 | 1.52E-30 | ENSG00000137094 | DNAJB5 | 25822 |
| ENSG00000250444.1 | 3.980671283 | 3.236213404 | 0.898179251 | 2.935064 | 0.003335 | 0.021829066 | ENSG00000250444 | CCT5P1 | 402226 |
| ENSG00000233559.2 | 123.9573437 | 3.23619604 | 0.287701733 | 9.162948 | 5.05E-20 | 1.93E-18 | ENSG00000233559 | NA | NA |
| ENSG00000188672.18 | 13.63493889 | 3.2361332 | 0.478138595 | 5.513324 | 3.52E-08 | 5.09E-07 | ENSG00000188672 | RHCE | 6006 |
| ENSG00000163216.7 | 4393.592385 | 3.233915506 | 0.400718959 | 6.572975 | 4.93E-11 | 9.65E-10 | ENSG00000163216 | SPRR2D | 6703 |
| ENSG00000225689.2 | 3.067672 | 3.232941 | 0.9661877 | 2.725083 | 0.006429 | 0.03895755 | ENSG00000225689 | NA | NA |
| ENSG00000184669.9 | 6.002822214 | 3.230197441 | 0.825927827 | 3.184537 | 0.00145 | 0.010336947 | ENSG00000184669 | OR7E14P | 10819 |
| ENSG00000125046.15 | 42.08046578 | 3.228849682 | 0.295891408 | 8.884508 | 6.42E-19 | 2.29E-17 | ENSG00000125046 | SSUH2 | 51066 |
| ENSG00000269374.1 | 2.714148 | 3.22852 | 0.9569314 | 2.746822 | 0.006018 | 0.03675152 | ENSG00000269374 | LOC73222 | 732229 |
| ENSG00000183044.12 | 74.90510298 | 3.226327615 | 0.212940656 | 12.33361 | 5.97E-35 | 4.88E-33 | ENSG00000183044 | ABAT | 18 |
| ENSG00000279339.1 | 3.127238119 | 3.225447675 | 0.889859107 | 2.950408 | 0.003174 | 0.020896871 | ENSG00000279339 | NA | NA |
| ENSG00000185112.5 | 863.2473887 | 3.222399343 | 0.213150047 | 12.30307 | 8.72E-35 | 7.08E-33 | ENSG00000185112 | FAM43A | 131583 |
| ENSG00000255221.3 | 21.90272129 | 3.218942588 | 0.475978206 | 5.502232 | 3.75E-08 | 5.40E-07 | ENSG00000255221 | CARD17 | 440068 |
| ENSG00000015568.13 | 3.395231962 | 3.217457492 | 0.945665868 | 2.767846 | 0.005643 | 0.034741656 | ENSG00000015568 | RGPD5 | 84220 |
| ENSG00000176046.8 | 64.29168494 | 3.217282086 | 0.239197228 | 10.94194 | 7.26E-28 | 4.34E-26 | ENSG00000176046 | NUPR1 | 26471 |
| ENSG00000108381.11 | 15.42908173 | 3.217005524 | 0.610798333 | 4.284566 | 1.83E-05 | 0.000185621 | ENSG00000108381 | ASPA | 443 |
| ENSG00000261592.1 | 10.37021952 | 3.216840539 | 0.727739697 | 3.595847 | 0.000323 | 0.002651849 | ENSG00000261592 | NA | NA |
| ENSG00000133107.14 | 19.53392561 | 3.215140098 | 0.565307265 | 4.626051 | 3.73E-06 | 4.18E-05 | ENSG00000133107 | TRPC4 | 7223 |
| ENSG00000280587.1 | 35.62933766 | 3.214651091 | 0.320755693 | 8.151534 | 3.59E-16 | 1.06E-14 | ENSG00000280587 | NA | NA |
| ENSG00000143631.10 | 126.1627391 | 3.213112277 | 0.262274246 | 9.963282 | 2.21E-23 | 1.03E-21 | ENSG00000143631 | FLG | 2312 |
| ENSG00000011347.9 | 653.2961624 | 3.212469096 | 0.130848019 | 19.96568 | 1.10E-88 | 3.79E-86 | ENSG00000011347 | SYT7 | 9066 |
| ENSG00000229021.2 | 30.58628425 | 3.210692313 | 0.299277734 | 8.72331 | 2.70E-18 | 9.29E-17 | ENSG00000229021 | NA | NA |
| ENSG00000249942.1 | 9.641667869 | 3.207986346 | 0.69382708 | 3.758842 | 0.000171 | 0.001476419 | ENSG00000249942 | NA | NA |
| ENSG00000213730.3 | 41.94977289 | 3.206097681 | 0.313309563 | 8.317964 | 8.95E-17 | 2.77E-15 | ENSG00000213730 | NA | NA |
| ENSG00000235350.1 | 6.28248 | 3.203684 | 0.6432677 | 4.04759 | 5.17E-05 | 0.000487828 | ENSG00000235350 | NA | NA |
| ENSG00000187122.17 | 26.22886837 | 3.20035687 | 0.48808246 | 5.3277 | 9.95E-08 | 1.37E-06 | ENSG00000187122 | SLIT1 | 6585 |
| ENSG00000184368.16 | 19.68565 | 3.20027 | 0.5130662 | 5.068099 | 4.02E-07 | 5.11E-06 | ENSG00000184368 | MAP7D2 | 256714 |
| ENSG00000179799.9 | 10.64437406 | 3.198227334 | 0.528685987 | 4.9145 | 8.90E-07 | 1.09E-05 | ENSG00000179799 | NA | NA |
| ENSG00000251000.1 | 7.393986014 | 3.19792823 | 0.642701293 | 4.042202 | 5.30E-05 | 0.00049812 | ENSG00000251000 | NA | NA |

| | | | | | | | | | |
|---|---|---|---|---|---|---|---|---|---|
| ENSG00000143850.15 | 833.0220959 | 3.19698874 | 0.136182854 | 19.06987 | 4.49E-81 | 1.33E-78 | ENSG00000143850 | PLEKHA6 | 22874 |
| ENSG00000255282.6 | 19.62912468 | 3.194312031 | 0.483954285 | 5.360655 | 8.29E-08 | 1.15E-06 | ENSG00000255282 | WTAPP1 | 1E+08 |
| ENSG00000204241.8 | 11.51042142 | 3.194250676 | 0.517092041 | 5.017 | 5.25E-07 | 6.58E-06 | ENSG00000204241 | LINC0273 | 1E+08 |
| ENSG00000170899.11 | 74.7029742 | 3.189535441 | 0.279622018 | 9.260842 | 2.03E-20 | 8.00E-19 | ENSG00000170899 | GSTA4 | 2941 |
| ENSG00000271880.1 | 28.68573376 | 3.187745376 | 0.41154307 | 6.287909 | 3.22E-10 | 5.89E-09 | ENSG00000271880 | AGAP11 | 119385 |
| ENSG00000186480.13 | 8730.001252 | 3.187181767 | 0.161893577 | 15.98076 | 1.74E-57 | 3.05E-55 | ENSG00000186480 | INSIG1 | 3638 |
| ENSG00000105369.9 | 41.61481 | 3.186897 | 0.573822 | 4.508187 | 6.54E-06 | 7.12E-05 | ENSG00000105369 | CD79A | 973 |
| ENSG00000162755.14 | 8.109179404 | 3.186102188 | 0.612220863 | 4.224133 | 2.40E-05 | 0.000238181 | ENSG00000162755 | KLHDC9 | 126823 |
| ENSG00000178860.8 | 24.82651582 | 3.18427166 | 0.386943997 | 6.678671 | 2.41E-11 | 4.87E-10 | ENSG00000178860 | MSC | 9242 |
| ENSG00000136535.15 | 3.547358187 | 3.183694336 | 0.965603642 | 2.67573 | 0.007457 | 0.044429638 | ENSG00000136535 | TBR1 | 10716 |
| ENSG00000137877.10 | 603.6264468 | 3.183397368 | 0.394603079 | 6.546825 | 5.88E-11 | 1.14E-09 | ENSG00000137877 | SPTBN5 | 51332 |
| ENSG00000257732.1 | 6.594308089 | 3.180403492 | 0.971686099 | 2.655594 | 0.007917 | 0.046794746 | ENSG00000257732 | NA | NA |
| ENSG00000261168.1 | 4.672741982 | 3.178245091 | 0.928174443 | 2.777759 | 0.005474 | 0.033895315 | ENSG00000261168 | NA | NA |
| ENSG00000067064.11 | 10887.27938 | 3.175218481 | 0.126636129 | 20.33557 | 6.23E-92 | 2.27E-89 | ENSG00000067064 | IDI1 | 3422 |
| ENSG00000101460.13 | 25.05903 | 3.174596 | 0.5798718 | 4.43994 | 9.00E-06 | 9.62E-05 | ENSG00000101460 | MAP1LC3 | 84557 |
| ENSG00000170054.14 | 4.702344125 | 3.171850981 | 0.827827475 | 3.106748 | 0.001892 | 0.013121323 | ENSG00000170054 | SERPINA9 | 327657 |
| ENSG00000250722.6 | 29.88625547 | 3.171667228 | 0.380596274 | 6.756943 | 1.41E-11 | 2.91E-10 | ENSG00000250722 | SELENOP | 6414 |
| ENSG00000155269.12 | 31.57683726 | 3.167457903 | 0.307450566 | 8.350799 | 6.78E-17 | 2.11E-15 | ENSG00000155269 | GPR78 | 27201 |
| ENSG00000171502.15 | 8.485100836 | 3.164191091 | 0.651905084 | 3.933381 | 8.38E-05 | 0.000766005 | ENSG00000171502 | COL24A1 | 255631 |
| ENSG00000273951.1 | 2.967626 | 3.162438 | 0.9315789 | 2.75064 | 0.005948 | 0.03642514 | ENSG00000273951 | NA | NA |
| ENSG00000196136.17 | 9.591198246 | 3.162297813 | 0.76136356 | 3.365406 | 0.000764 | 0.005826878 | ENSG00000196136 | SERPINA3 | 12 |
| ENSG00000052802.13 | 7346.309456 | 3.161016634 | 0.071117227 | 36.0112 | 5.59E-284 | 1.35E-280 | ENSG00000052802 | MSMO1 | 6307 |
| ENSG00000158246.8 | 768.1204145 | 3.160895444 | 0.359254088 | 7.128368 | 1.02E-12 | 2.32E-11 | ENSG00000158246 | TENT5B | 115572 |
| ENSG00000118322.14 | 35.63383088 | 3.157884377 | 0.49152969 | 5.203926 | 1.95E-07 | 2.59E-06 | ENSG00000118322 | ATP10B | 23120 |
| ENSG00000260996.1 | 6.116257908 | 3.157498374 | 0.723912833 | 3.532882 | 0.000411 | 0.003309458 | ENSG00000260996 | NA | NA |
| ENSG00000163359.15 | 323.4708955 | 3.156875537 | 0.157640348 | 16.21968 | 3.66E-59 | 6.91E-57 | ENSG00000163359 | COL6A3 | 1293 |
| ENSG00000168140.5 | 13.23730187 | 3.154881439 | 0.595006415 | 4.293872 | 1.76E-05 | 0.000178208 | ENSG00000168140 | VASN | 114990 |
| ENSG00000168427.9 | 35.93545055 | 3.146191717 | 0.464193478 | 5.485195 | 4.13E-08 | 5.91E-07 | ENSG00000168427 | KLHL30 | 377007 |
| ENSG00000125823.12 | 16.63083 | 3.14611 | 0.5466098 | 4.658002 | 3.19E-06 | 3.59E-05 | ENSG00000125823 | CSTL1 | 128817 |
| ENSG00000154783.11 | 4.355922184 | 3.144854327 | 0.82254369 | 3.093883 | 0.001976 | 0.013653455 | ENSG00000154783 | FGD5 | 152273 |
| ENSG00000149564.12 | 34.85074488 | 3.142038436 | 0.393441278 | 6.461036 | 1.04E-10 | 1.97E-09 | ENSG00000149564 | ESAM | 90952 |
| ENSG00000136235.16 | 128.448829 | 3.139597683 | 0.282685331 | 8.983833 | 2.61E-19 | 9.60E-18 | ENSG00000136235 | GPNMB | 10457 |
| ENSG00000254186.3 | 4.657405156 | 3.136418506 | 0.770215695 | 3.293128 | 0.000991 | 0.007346063 | ENSG00000254186 | NA | NA |
| ENSG00000287292.1 | 4.691470504 | 3.13473143 | 0.938732323 | 2.700164 | 0.006931 | 0.041677085 | ENSG00000287292 | NA | NA |
| ENSG00000086696.11 | 60.22259134 | 3.134119281 | 0.246699805 | 10.27208 | 9.42E-25 | 4.69E-23 | ENSG00000086696 | HSD17B2 | 3294 |
| ENSG00000163618.18 | 6.160542559 | 3.131586827 | 0.696102906 | 3.6368 | 0.000276 | 0.002297252 | ENSG00000163618 | CADPS | 8618 |
| ENSG00000259341.2 | 24.70989243 | 3.125614484 | 0.478913297 | 5.273636 | 1.34E-07 | 1.82E-06 | ENSG00000259341 | NA | NA |
| ENSG00000142156.14 | 2791.556 | 3.123717 | 0.3805401 | 6.631935 | 3.31E-11 | 6.59E-10 | ENSG00000142156 | COL6A1 | 1291 |
| ENSG00000171408.14 | 12.50661637 | 3.121184945 | 0.515584005 | 4.88996 | 1.01E-06 | 1.22E-05 | ENSG00000171408 | PDE7B | 27115 |
| ENSG00000215853.3 | 13.1245636 | 3.121032981 | 0.729103932 | 3.457714 | 0.000545 | 0.004281993 | ENSG00000215853 | RPTN | 126638 |
| ENSG00000104081.14 | 89.90540652 | 3.119668829 | 0.20350201 | 12.38154 | 3.29E-35 | 2.74E-33 | ENSG00000104081 | BMF | 90427 |
| ENSG00000116774.12 | 15.89504064 | 3.118784737 | 0.506171065 | 4.976153 | 6.49E-07 | 8.05E-06 | ENSG00000116774 | OLFML3 | 56944 |

| | | | | | | | | | |
|---|---|---|---|---|---|---|---|---|---|
| ENSG00000237781.3 | 17.978652 | 3.116777943 | 0.415124927 | 6.0627 | 1.34E-09 | 2.27E-08 | ENSG00000237781 | NA | NA |
| ENSG00000251281.1 | 20.10298831 | 3.116229979 | 0.373044786 | 6.745115 | 1.53E-11 | 3.16E-10 | ENSG00000251281 | NA | NA |
| ENSG00000250697.2 | 46.62067178 | 3.115500831 | 0.245007993 | 10.26702 | 9.92E-25 | 4.91E-23 | ENSG00000250697 | NA | NA |
| ENSG00000131981.16 | 12915.00384 | 3.114586105 | 0.111432912 | 22.56592 | 9.37E-113 | 4.62E-110 | ENSG00000131981 | LGALS3 | 3958 |
| ENSG00000139433.10 | 11963.72788 | 3.114539682 | 0.201514245 | 12.47822 | 9.82E-36 | 8.38E-34 | ENSG00000139433 | GLTP | 51228 |
| ENSG00000082684.15 | 8.813756256 | 3.112318798 | 0.723392751 | 3.472967 | 0.000515 | 0.004065084 | ENSG00000082684 | SEMA5B | 54437 |
| ENSG00000170465.10 | 3095.188556 | 3.112014793 | 0.243807385 | 10.30328 | 6.81E-25 | 3.41E-23 | ENSG00000170465 | KRT6C | 286887 |
| ENSG00000222032.1 | 15.8413591 | 3.107048261 | 0.430082831 | 5.829222 | 5.57E-09 | 8.84E-08 | ENSG00000222032 | NA | NA |
| ENSG00000129910.7 | 8.853345358 | 3.10677795 | 0.726029715 | 3.452721 | 0.000555 | 0.004355608 | ENSG00000129910 | CDH15 | 1013 |
| ENSG00000259354.5 | 10.70921625 | 3.105977635 | 0.712120369 | 3.519037 | 0.000433 | 0.003466759 | ENSG00000259354 | NA | NA |
| ENSG00000269506.2 | 9.282049891 | 3.10244004 | 0.637587192 | 3.924859 | 8.68E-05 | 0.000792273 | ENSG00000269506 | NA | NA |
| ENSG00000272666.1 | 6.368397 | 3.100875 | 0.8407992 | 2.974402 | 0.002936 | 0.01949892 | ENSG00000272666 | KLHDC7B- | 1.05E+08 |
| ENSG00000135218.19 | 14.36617223 | 3.095717616 | 0.608427198 | 4.101917 | 4.10E-05 | 0.000393887 | ENSG00000135218 | CD36 | 948 |
| ENSG00000144045.14 | 134.2397281 | 3.095092636 | 0.234208731 | 10.65329 | 1.68E-26 | 9.18E-25 | ENSG00000144045 | DQX1 | 165545 |
| ENSG00000122733.12 | 21.16781315 | 3.094721542 | 0.445814648 | 5.595872 | 2.20E-08 | 3.27E-07 | ENSG00000122733 | PHF24 | 23349 |
| ENSG00000147113.17 | 3.474259 | 3.094645 | 0.883381 | 2.823974 | 0.004743 | 0.02985785 | ENSG00000147113 | DIPK2B | 79742 |
| ENSG00000102970.10 | 3.889021015 | 3.089026415 | 0.82607912 | 3.013061 | 0.002586 | 0.017444074 | ENSG00000102970 | CCL17 | 6361 |
| ENSG00000066248.15 | 69.64887035 | 3.088997959 | 0.296975202 | 8.381164 | 5.24E-17 | 1.65E-15 | ENSG00000066248 | NGEF | 25791 |
| ENSG00000131459.13 | 2983.401566 | 3.0856314 | 0.069633706 | 35.69581 | 4.59E-279 | 1.02E-275 | ENSG00000131459 | GFPT2 | 9945 |
| ENSG00000197635.10 | 29.08026644 | 3.083967355 | 0.471484071 | 5.268401 | 1.38E-07 | 1.87E-06 | ENSG00000197635 | DPP4 | 1803 |
| ENSG00000105523.3 | 78.1082 | 3.080737 | 0.202581 | 12.24566 | 1.77E-34 | 1.42E-32 | ENSG00000105523 | FAM83E | 54854 |
| ENSG00000215386.13 | 4.408348 | 3.080155 | 0.8423102 | 2.944467 | 0.003235 | 0.02125496 | ENSG00000215386 | MIR99AHC | 388815 |
| ENSG00000186204.14 | 20.93101 | 3.079857 | 0.3994829 | 6.207666 | 5.38E-10 | 9.56E-09 | ENSG00000186204 | CYP4F12 | 66002 |
| ENSG00000180596.7 | 161.2958479 | 3.078965859 | 0.269245712 | 9.207076 | 3.35E-20 | 1.29E-18 | ENSG00000180596 | H2BC4 | 8347 |
| ENSG00000171119.2 | 5.995911341 | 3.074493081 | 0.851628378 | 2.905602 | 0.003665 | 0.023731092 | ENSG00000171119 | NRTN | 4902 |
| ENSG00000247372.2 | 6.547931397 | 3.073128332 | 0.730325476 | 3.386337 | 0.000708 | 0.005440557 | ENSG00000247372 | NA | NA |
| ENSG00000253865.1 | 4.46660048 | 3.071920187 | 0.767983926 | 3.218713 | 0.001288 | 0.009307732 | ENSG00000253865 | NA | NA |
| ENSG00000178573.7 | 27.23602658 | 3.071770219 | 0.371880147 | 6.646685 | 3.00E-11 | 6.01E-10 | ENSG00000178573 | MAF | 4094 |
| ENSG00000258640.3 | 5.097759531 | 3.070921432 | 0.860036851 | 2.873041 | 0.004065 | 0.026066792 | ENSG00000258640 | NA | NA |
| ENSG00000093134.14 | 23.25415924 | 3.06743962 | 0.647237526 | 3.812263 | 0.000138 | 0.001209854 | ENSG00000093134 | VNN3 | 55350 |
| ENSG00000285879.1 | 9.748316561 | 3.065971116 | 0.663753415 | 3.715192 | 0.000203 | 0.001729236 | ENSG00000285879 | NA | NA |
| ENSG00000230400.3 | 4.737287 | 3.065784 | 0.8643786 | 2.852666 | 0.004335 | 0.02762505 | ENSG00000230400 | NA | NA |
| ENSG00000168350.8 | 6.003584468 | 3.064900578 | 0.645348006 | 3.819491 | 0.000134 | 0.001179621 | ENSG00000168350 | DEGS2 | 123099 |
| ENSG00000078804.13 | 518.1578 | 3.063922 | 0.2259421 | 10.9051 | 1.09E-27 | 6.40E-26 | ENSG00000078804 | TP53INP2 | 58476 |
| ENSG00000066185.13 | 5.832158708 | 3.063831129 | 0.76189246 | 3.233831 | 0.001221 | 0.008879469 | ENSG00000066185 | ZMYND12 | 84217 |
| ENSG00000285130.2 | 22.99167649 | 3.059713923 | 0.377520775 | 6.51544 | 7.25E-11 | 1.40E-09 | ENSG00000285130 | NA | NA |
| ENSG00000110203.9 | 12.09098451 | 3.05867477 | 0.47919307 | 5.130865 | 2.88E-07 | 3.74E-06 | ENSG00000110203 | FOLR3 | 2352 |
| ENSG00000287872.1 | 4.139828804 | 3.05144511 | 0.88994085 | 2.754616 | 0.005876 | 0.036044651 | ENSG00000287872 | NA | NA |
| ENSG00000127585.12 | 157.1927532 | 3.051333951 | 0.140591134 | 17.43591 | 4.41E-68 | 1.01E-65 | ENSG00000127585 | FBXL16 | 146330 |
| ENSG00000164442.10 | 1718.445613 | 3.047679209 | 0.233221038 | 10.4951 | 9.10E-26 | 4.84E-24 | ENSG00000164442 | CITED2 | 10370 |
| ENSG00000162998.5 | 17.80770566 | 3.04261323 | 0.553777534 | 4.41082 | 1.03E-05 | 0.000108791 | ENSG00000162998 | FRZB | 2487 |
| ENSG00000233221.7 | 29.11487867 | 3.042478052 | 0.41388065 | 5.901407 | 3.60E-09 | 5.84E-08 | ENSG00000233221 | NA | NA |

| | | | | | | | | | |
|---|---|---|---|---|---|---|---|---|---|
| ENSG00000164379.7 | 155.515328 | 3.042049871 | 0.315575839 | 7.738393 | 1.01E-14 | 2.69E-13 | ENSG00000164379 | FOXQ1 | 94234 |
| ENSG00000160471.13 | 101.6315 | 3.041233 | 0.2854779 | 8.551389 | 1.22E-17 | 4.02E-16 | ENSG00000160471 | COX6B2 | 125965 |
| ENSG00000100385.14 | 5.366158 | 3.040469 | 0.7526332 | 3.242574 | 0.001185 | 0.008625591 | ENSG00000100385 | IL2RB | 3560 |
| ENSG00000187091.13 | 1014.533886 | 3.032748485 | 0.266346052 | 9.133788 | 6.61E-20 | 2.51E-18 | ENSG00000187091 | PLCD1 | 5333 |
| ENSG00000256443.1 | 12.6699061 | 3.027650752 | 0.507624099 | 4.782379 | 1.73E-06 | 2.03E-05 | ENSG00000256443 | NA | NA |
| ENSG00000283167.2 | 7.875542095 | 3.025817004 | 0.642738412 | 3.77419 | 0.000161 | 0.001395228 | ENSG00000283167 | NA | NA |
| ENSG00000227110.7 | 17.85565812 | 3.024419315 | 0.407413984 | 5.950751 | 2.67E-09 | 4.40E-08 | ENSG00000227110 | LMCD1-AS | 1E+08 |
| ENSG00000110195.13 | 4.073466161 | 3.023426664 | 0.776517344 | 3.120892 | 0.001803 | 0.012588668 | ENSG00000110195 | FOLR1 | 2348 |
| ENSG00000288029.1 | 6.181356555 | 3.021620812 | 0.645186664 | 3.753365 | 0.000174 | 0.001504671 | ENSG00000288029 | NA | NA |
| ENSG00000170323.8 | 19.02436121 | 3.019600219 | 0.615343351 | 3.932114 | 8.42E-05 | 0.00076979 | ENSG00000170323 | FABP4 | 2167 |
| ENSG00000172478.18 | 268.1859742 | 3.014518193 | 0.290387742 | 8.314808 | 9.19E-17 | 2.84E-15 | ENSG00000172478 | MAB21L4 | 79919 |
| ENSG00000118515.11 | 3988.265402 | 3.014433166 | 0.142395882 | 16.95578 | 1.74E-64 | 3.74E-62 | ENSG00000118515 | SGK1 | 6446 |
| ENSG00000220920.1 | 8.11739109 | 3.01223289 | 0.650386922 | 3.70892 | 0.000208 | 0.001768107 | ENSG00000220920 | RPS12P12 | 1E+08 |
| ENSG00000261092.1 | 42.11243241 | 3.012016355 | 0.274008702 | 8.8027 | 1.34E-18 | 4.67E-17 | ENSG00000261092 | LINC0217 | 1.05E+08 |
| ENSG00000203780.10 | 125.2114844 | 3.010562126 | 0.245736782 | 9.809529 | 1.02E-22 | 4.55E-21 | ENSG00000203780 | FANK1 | 92565 |
| ENSG00000261787.2 | 22.77305059 | 3.009114286 | 0.39802346 | 6.052694 | 1.42E-09 | 2.40E-08 | ENSG00000261787 | TCF24 | 1E+08 |
| ENSG00000095585.16 | 18.45346403 | 3.000475207 | 0.550823748 | 4.357973 | 1.31E-05 | 0.0001363 | ENSG00000095585 | BLNK | 29760 |
| ENSG00000279608.1 | 7.920285655 | 2.997372821 | 0.63714089 | 3.762704 | 0.000168 | 0.001455213 | ENSG00000279608 | NA | NA |
| ENSG00000258116.1 | 7.833122677 | 2.996472817 | 0.599469289 | 3.997657 | 6.40E-05 | 0.000595273 | ENSG00000258116 | PPIAP45 | 1.01E+08 |
| ENSG00000177464.5 | 5.727743 | 2.993653 | 0.649525 | 3.685237 | 0.000228 | 0.001928619 | ENSG00000177464 | GPR4 | 2828 |
| ENSG00000221947.7 | 5.652612335 | 2.993054797 | 0.690207132 | 3.467155 | 0.000526 | 0.004146626 | ENSG00000221947 | XKR9 | 389668 |
| ENSG00000163083.6 | 4.632969557 | 2.989237262 | 0.785405908 | 3.042041 | 0.00235 | 0.015974532 | ENSG00000163083 | INHBB | 3625 |
| ENSG00000078018.19 | 149.0532422 | 2.989072397 | 0.256917028 | 9.299004 | 1.42E-20 | 5.65E-19 | ENSG00000078018 | MAP2 | 4133 |
| ENSG00000214514.8 | 10.34368649 | 2.988445144 | 0.550142703 | 4.341501 | 1.42E-05 | 0.000146191 | ENSG00000214514 | KRT42P | 284116 |
| ENSG00000189433.7 | 1928.637702 | 2.986689541 | 0.207786403 | 11.48626 | 1.55E-30 | 1.09E-28 | ENSG00000189433 | GJB4 | 127534 |
| ENSG00000262823.1 | 6.992853083 | 2.985777324 | 0.840088677 | 2.839911 | 0.004513 | 0.028555895 | ENSG00000262823 | NA | NA |
| ENSG00000136155.17 | 2196.553487 | 2.985043011 | 0.447288545 | 5.332225 | 9.70E-08 | 1.34E-06 | ENSG00000136155 | SCEL | 8796 |
| ENSG00000244242.2 | 3.147368879 | 2.983529189 | 0.903260128 | 2.638807 | 0.00832 | 0.048840258 | ENSG00000244242 | IFITM10 | 402778 |
| ENSG00000158525.15 | 6.391682926 | 2.983155382 | 0.785516585 | 3.03387 | 0.002414 | 0.016384322 | ENSG00000158525 | CPA5 | 93979 |
| ENSG00000254486.1 | 4.999415589 | 2.982061978 | 0.717142545 | 3.321602 | 0.000895 | 0.006719489 | ENSG00000254486 | LINC0254 | 1.05E+08 |
| ENSG00000186529.16 | 575.5783 | 2.978745 | 0.2912028 | 8.168689 | 3.12E-16 | 9.26E-15 | ENSG00000186529 | CYP4F3 | 4051 |
| ENSG00000224899.1 | 14.51448476 | 2.97638602 | 0.452709505 | 5.249251 | 1.53E-07 | 2.06E-06 | ENSG00000224899 | NA | NA |
| ENSG00000140600.17 | 21.06612799 | 2.975494208 | 0.381256616 | 6.230696 | 4.64E-10 | 8.34E-09 | ENSG00000140600 | SH3GL3 | 6457 |
| ENSG00000103522.16 | 4.288550485 | 2.975408073 | 0.745407321 | 3.186725 | 0.001439 | 0.010278333 | ENSG00000103522 | IL21R | 50615 |
| ENSG00000229951.5 | 9.79172659 | 2.973073112 | 0.648036813 | 3.661942 | 0.00025 | 0.002098826 | ENSG00000229951 | FLJ31356 | 403150 |
| ENSG00000115226.10 | 368.5198044 | 2.971952115 | 0.151092244 | 15.6987 | 1.54E-55 | 2.60E-53 | ENSG00000115226 | FNDC4 | 64838 |
| ENSG00000287264.1 | 18.66748985 | 2.971273211 | 0.554914985 | 4.273219 | 1.93E-05 | 0.000194218 | ENSG00000287264 | NA | NA |
| ENSG00000272970.2 | 23.94798164 | 2.968911945 | 0.374247227 | 6.329805 | 2.45E-10 | 4.54E-09 | ENSG00000272970 | LOC10798 | 1.08E+08 |
| ENSG00000230415.1 | 12.53735655 | 2.961079029 | 0.680905331 | 3.467558 | 0.000525 | 0.004141626 | ENSG00000230415 | NA | NA |
| ENSG00000235643.2 | 6.81853432 | 2.959469592 | 0.658809413 | 3.581415 | 0.000342 | 0.002791585 | ENSG00000235643 | LINC0164 | 1.02E+08 |
| ENSG00000163435.16 | 1292.020725 | 2.95804159 | 0.332544453 | 7.090906 | 1.33E-12 | 3.01E-11 | ENSG00000163435 | ELF3 | 1999 |
| ENSG00000137731.14 | 74.12721843 | 2.95743937 | 0.198575887 | 11.87173 | 1.66E-32 | 1.28E-30 | ENSG00000137731 | FXYD2 | 486 |

| | | | | | | | | | |
|---|---|---|---|---|---|---|---|---|---|
| ENSG00000255983.1 | 7.79553505 | 2.956838046 | 0.667879446 | 3.528838 | 0.000417 | 0.003353715 | ENSG00000255983 | NA | NA |
| ENSG00000259207.7 | 50.33088257 | 2.956556151 | 0.378359214 | 6.228357 | 4.71E-10 | 8.45E-09 | ENSG00000259207 | ITGB3 | 3690 |
| ENSG00000231367.6 | 6.237258942 | 2.952437467 | 0.622726574 | 3.777641 | 0.000158 | 0.001377391 | ENSG00000231367 | LINC0261: | 1.02E+08 |
| ENSG00000166257.9 | 28.85756695 | 2.950523134 | 0.354186839 | 6.636393 | 3.21E-11 | 6.42E-10 | ENSG00000166257 | SCN3B | 55800 |
| ENSG00000267709.1 | 3.516366371 | 2.947795328 | 0.839703865 | 2.79598 | 0.005174 | 0.032206944 | ENSG00000267709 | LOC10192 | 1.02E+08 |
| ENSG00000263823.1 | 29.15410239 | 2.946175817 | 0.437564942 | 5.361892 | 8.24E-08 | 1.15E-06 | ENSG00000263823 | NA | NA |
| ENSG00000110324.10 | 11.44710042 | 2.944027594 | 0.757731371 | 3.093481 | 0.001978 | 0.013664893 | ENSG00000110324 | IL10RA | 3587 |
| ENSG00000091010.6 | 8.853184103 | 2.943861558 | 0.551492563 | 4.250033 | 2.14E-05 | 0.000214157 | ENSG00000091010 | POU4F3 | 5459 |
| ENSG00000143515.17 | 125.8276997 | 2.941523885 | 0.217338695 | 10.77362 | 4.59E-27 | 2.60E-25 | ENSG00000143515 | ATP8B2 | 57198 |
| ENSG00000204103.4 | 285.921 | 2.940232 | 0.4919632 | 4.756924 | 1.97E-06 | 2.28E-05 | ENSG00000204103 | MAFB | 9935 |
| ENSG00000184838.15 | 4.056989338 | 2.939664653 | 0.791406453 | 2.956338 | 0.003113 | 0.020545012 | ENSG00000184838 | PRR16 | 51334 |
| ENSG00000166268.10 | 7.525582112 | 2.939083139 | 0.633534574 | 3.692116 | 0.000222 | 0.001881355 | ENSG00000166268 | MYRFL | 196446 |
| ENSG00000104237.10 | 72.59536187 | 2.936088908 | 0.316092704 | 7.390518 | 1.46E-13 | 3.60E-12 | ENSG00000104237 | RP1 | 6101 |
| ENSG00000118402.6 | 1231.65187 | 2.934508836 | 0.156975007 | 14.87185 | 5.02E-50 | 7.11E-48 | ENSG00000118402 | ELOVL4 | 6785 |
| ENSG00000170214.5 | 7.516757624 | 2.933553045 | 0.607026546 | 3.844236 | 0.000121 | 0.001075991 | ENSG00000170214 | ADRA1B | 147 |
| ENSG00000182809.10 | 461.0104535 | 2.933273636 | 0.183191173 | 12.73682 | 3.69E-37 | 3.34E-35 | ENSG00000182809 | CRIP2 | 1397 |
| ENSG00000182580.3 | 550.4019621 | 2.932208463 | 0.386159417 | 6.039497 | 1.55E-09 | 2.60E-08 | ENSG00000182580 | EPHB3 | 2049 |
| ENSG00000100033.16 | 116.8226 | 2.931339 | 0.2564108 | 9.092199 | 9.71E-20 | 3.64E-18 | ENSG00000100033 | PRODH | 5625 |
| ENSG00000225431.1 | 14.93037 | 2.93126 | 0.556032 | 4.192673 | 2.76E-05 | 0.000271626 | ENSG00000225431 | LINC0167 | 1.02E+08 |
| ENSG00000185847.8 | 8.794983347 | 2.931228703 | 0.669965533 | 3.479625 | 0.000502 | 0.003974827 | ENSG00000185847 | LINC0140 | 1E+08 |
| ENSG00000266553.2 | 15.69851247 | 2.928633272 | 0.419412763 | 5.552128 | 2.82E-08 | 4.16E-07 | ENSG00000266553 | RN7SL356 | 1.06E+08 |
| ENSG00000261888.1 | 5.471074317 | 2.927720541 | 0.676635487 | 3.44014 | 0.000581 | 0.004540454 | ENSG00000261888 | NA | NA |
| ENSG00000273010.1 | 4.789811842 | 2.927360187 | 0.745885804 | 3.120263 | 0.001807 | 0.012608945 | ENSG00000273010 | NA | NA |
| ENSG00000287426.1 | 5.436249401 | 2.925674211 | 0.677965565 | 3.430372 | 0.000603 | 0.004693335 | ENSG00000287426 | NA | NA |
| ENSG00000170959.14 | 42.42698515 | 2.924868702 | 0.247455507 | 9.395098 | 5.72E-21 | 2.33E-19 | ENSG00000170959 | DCDC1 | 341019 |
| ENSG00000185338.5 | 163.7511993 | 2.924372393 | 0.365516949 | 6.359137 | 2.03E-10 | 3.78E-09 | ENSG00000185338 | SOCS1 | 8651 |
| ENSG00000234899.10 | 30.58899595 | 2.921797574 | 0.382375009 | 6.072043 | 1.26E-09 | 2.15E-08 | ENSG00000234899 | SOX9-AS1 | 400618 |
| ENSG00000187753.13 | 10.90647213 | 2.919686265 | 0.528555103 | 4.388731 | 1.14E-05 | 0.000119546 | ENSG00000187753 | C9orf153 | 389766 |
| ENSG00000115590.14 | 1449.041382 | 2.917507723 | 0.096203973 | 24.08952 | 3.22E-128 | 1.79E-125 | ENSG00000115590 | IL1R2 | 7850 |
| ENSG00000274038.1 | 8.308932498 | 2.911513854 | 0.56970855 | 4.057362 | 4.96E-05 | 0.000469528 | ENSG00000274038 | NA | NA |
| ENSG00000130203.10 | 275.461 | 2.910721 | 0.4590191 | 5.034041 | 4.80E-07 | 6.04E-06 | ENSG00000130203 | APOE | 348 |
| ENSG00000181333.11 | 1079.881735 | 2.910417998 | 0.240641655 | 9.601073 | 7.91E-22 | 3.38E-20 | ENSG00000181333 | HEPHL1 | 341208 |
| ENSG00000231290.6 | 9.565361 | 2.909893 | 0.5896814 | 3.917188 | 8.96E-05 | 0.000815389 | ENSG00000231290 | APCDD1L- | 149773 |
| ENSG00000129757.13 | 65.51179625 | 2.904548342 | 0.462570304 | 4.98205 | 6.29E-07 | 7.83E-06 | ENSG00000129757 | CDKN1C | 1028 |
| ENSG00000176928.6 | 157.9494147 | 2.903984798 | 0.167814094 | 13.72939 | 6.77E-43 | 7.54E-41 | ENSG00000176928 | GCNT4 | 51301 |
| ENSG00000253406.1 | 7.422780154 | 2.90358945 | 0.55518763 | 4.149209 | 3.34E-05 | 0.000324746 | ENSG00000253406 | NA | NA |
| ENSG00000287728.1 | 4.061616509 | 2.902396728 | 0.840908965 | 2.737986 | 0.006182 | 0.037667335 | ENSG00000287728 | NA | NA |
| ENSG00000100100.12 | 488.2826 | 2.90234 | 0.1191239 | 19.32728 | 3.17E-83 | 1.00E-80 | ENSG00000100100 | PIK3IP1 | 113791 |
| ENSG00000103742.12 | 37.19766159 | 2.901290625 | 0.31044616 | 7.412849 | 1.24E-13 | 3.06E-12 | ENSG00000103742 | IGDCC4 | 57722 |
| ENSG00000261652.2 | 40.42194756 | 2.8987851 | 0.437349788 | 5.256171 | 1.47E-07 | 1.99E-06 | ENSG00000261652 | C15orf65 | 145788 |
| ENSG00000115718.17 | 41.34091241 | 2.895354051 | 0.287925195 | 7.97205 | 1.56E-15 | 4.47E-14 | ENSG00000115718 | PROC | 5624 |
| ENSG00000072657.8 | 8.186778419 | 2.892354435 | 0.568650458 | 4.031219 | 5.55E-05 | 0.00052143 | ENSG00000072657 | TRHDE | 29953 |

| | | | | | | | | | |
|---|---|---|---|---|---|---|---|---|---|
| ENSG00000260231.2 | 329.710363 | 2.891577667 | 0.271801759 | 8.431063 | 3.43E-17 | 1.10E-15 | ENSG00000260231 | KDM7A-D⁻ | 1E+08 |
| ENSG00000272595.2 | 5.645652745 | 2.890225558 | 0.701327942 | 3.265556 | 0.001092 | 0.008034362 | ENSG00000272595 | NA | NA |
| ENSG00000244268.1 | 4.849878913 | 2.890220448 | 0.777844454 | 2.944317 | 0.003237 | 0.021260066 | ENSG00000244268 | NA | NA |
| ENSG00000163803.13 | 32.76831065 | 2.88671604 | 0.331452535 | 6.899075 | 5.23E-12 | 1.13E-10 | ENSG00000163803 | PLB1 | 151056 |
| ENSG00000255364.1 | 49.15103072 | 2.884506589 | 0.279118592 | 8.184717 | 2.73E-16 | 8.16E-15 | ENSG00000255364 | NA | NA |
| ENSG00000144671.10 | 15.68126434 | 2.883953929 | 0.541746948 | 4.215905 | 2.49E-05 | 0.000246665 | ENSG00000144671 | SLC22A14 | 9389 |
| ENSG00000251562.8 | 79789.59429 | 2.881236168 | 0.110198849 | 20.70109 | 3.39E-95 | 1.33E-92 | ENSG00000251562 | MALAT1 | 378938 |
| ENSG00000230454.1 | 14.12008859 | 2.880687656 | 0.549483271 | 4.150604 | 3.32E-05 | 0.00032289 | ENSG00000230454 | NA | NA |
| ENSG00000250208.6 | 14.1224219 | 2.87690437 | 0.576991133 | 3.946169 | 7.94E-05 | 0.000728999 | ENSG00000250208 | FZD10-AS | 440119 |
| ENSG00000155324.9 | 5148.27983 | 2.876679127 | 0.138605516 | 16.4256 | 1.25E-60 | 2.49E-58 | ENSG00000155324 | GRAMD2E | 65983 |
| ENSG00000112972.15 | 5847.383692 | 2.876248448 | 0.086749294 | 26.23939 | 9.45E-152 | 6.62E-149 | ENSG00000112972 | HMGCS1 | 3157 |
| ENSG00000118194.20 | 8.072451933 | 2.875707427 | 0.777206256 | 2.928061 | 0.003411 | 0.02226112 | ENSG00000118194 | TNNT2 | 7139 |
| ENSG00000111432.4 | 78.72860152 | 2.873034495 | 0.281339136 | 8.07934 | 6.51E-16 | 1.91E-14 | ENSG00000111432 | FZD10 | 11211 |
| ENSG00000135373.12 | 6426.190487 | 2.870312782 | 0.325004911 | 6.985472 | 2.84E-12 | 6.24E-11 | ENSG00000135373 | EHF | 26298 |
| ENSG00000232656.8 | 163.2587758 | 2.867541085 | 0.232002001 | 9.7738 | 1.46E-22 | 6.39E-21 | ENSG00000232656 | IDI2-AS1 | 55853 |
| ENSG00000263718.2 | 13.30984744 | 2.867222935 | 0.714518073 | 3.17308 | 0.001508 | 0.010716368 | ENSG00000263718 | SEPTIN9-[ | 1.05E+08 |
| ENSG00000169418.10 | 13.03292329 | 2.865180152 | 0.45353891 | 4.994456 | 5.90E-07 | 7.36E-06 | ENSG00000169418 | NPR1 | 4881 |
| ENSG00000162105.19 | 72.79857101 | 2.864708289 | 0.274510635 | 8.249984 | 1.58E-16 | 4.81E-15 | ENSG00000162105 | SHANK2 | 22941 |
| ENSG00000137462.8 | 1741.254826 | 2.864450024 | 0.08858067 | 25.5637 | 3.87E-144 | 2.45E-141 | ENSG00000137462 | TLR2 | 7097 |
| ENSG00000275989.2 | 111.17655 | 2.858367262 | 0.39895299 | 5.660735 | 1.51E-08 | 2.28E-07 | ENSG00000275989 | NA | NA |
| ENSG00000184545.11 | 86.81351277 | 2.857356669 | 0.19691916 | 11.46337 | 2.02E-30 | 1.42E-28 | ENSG00000184545 | DUSP8 | 1850 |
| ENSG00000164199.18 | 492.823851 | 2.857042885 | 0.201736298 | 11.18809 | 4.66E-29 | 3.04E-27 | ENSG00000164199 | ADGRV1 | 84059 |
| ENSG00000222022.1 | 14.0736142 | 2.856018125 | 0.45113903 | 5.000716 | 5.71E-07 | 7.14E-06 | ENSG00000222022 | NA | NA |
| ENSG00000008226.19 | 28.39546862 | 2.855684777 | 0.322501381 | 6.994341 | 2.67E-12 | 5.87E-11 | ENSG00000008226 | DLEC1 | 9940 |
| ENSG00000092068.20 | 513.0044524 | 2.854391891 | 0.110394978 | 20.42115 | 1.08E-92 | 4.01E-90 | ENSG00000092068 | SLC7A8 | 23428 |
| ENSG00000230006.7 | 5.403939798 | 2.854132249 | 0.798546995 | 2.822792 | 0.004761 | 0.029947248 | ENSG00000230006 | ANKRD36I | 645784 |
| ENSG00000261040.7 | 91.39423478 | 2.852494192 | 0.271980193 | 8.281832 | 1.21E-16 | 3.72E-15 | ENSG00000261040 | WFDC21P | 645638 |
| ENSG00000138764.15 | 3838.071114 | 2.852333945 | 0.246706731 | 9.129601 | 6.88E-20 | 2.60E-18 | ENSG00000138764 | CCNG2 | 901 |
| ENSG00000284669.1 | 82.00215141 | 2.851415623 | 0.258271646 | 8.717239 | 2.85E-18 | 9.77E-17 | ENSG00000284669 | NA | NA |
| ENSG00000168824.14 | 285.7930861 | 2.849166366 | 0.223763577 | 10.05153 | 9.05E-24 | 4.28E-22 | ENSG00000168824 | NSG1 | 27065 |
| ENSG00000134830.5 | 7.45291 | 2.847663 | 0.5907013 | 3.805075 | 0.000142 | 0.001243482 | ENSG00000134830 | C5AR2 | 27202 |
| ENSG00000274750.2 | 82.12201226 | 2.847182738 | 0.255732036 | 8.787255 | 1.53E-18 | 5.33E-17 | ENSG00000274750 | H3C6 | 8353 |
| ENSG00000261357.1 | 5.4621129 | 2.845502768 | 0.708788541 | 3.168086 | 0.001534 | 0.010881855 | ENSG00000261357 | NA | NA |
| ENSG00000112033.14 | 7089.59441 | 2.845072312 | 0.211019111 | 10.63919 | 1.96E-26 | 1.06E-24 | ENSG00000112033 | PPARD | 5467 |
| ENSG00000274173.1 | 124.8039 | 2.844594 | 0.1928469 | 11.63925 | 2.60E-31 | 1.90E-29 | ENSG00000274173 | NA | NA |
| ENSG00000147394.18 | 7987.049 | 2.843765 | 0.4059624 | 5.527027 | 3.26E-08 | 4.75E-07 | ENSG00000147394 | ZNF185 | 7739 |
| ENSG00000226468.2 | 30.76224618 | 2.84270573 | 0.317280576 | 7.068525 | 1.57E-12 | 3.51E-11 | ENSG00000226468 | NA | NA |
| ENSG00000128918.15 | 40.94504819 | 2.840802202 | 0.279737674 | 8.01037 | 1.14E-15 | 3.30E-14 | ENSG00000128918 | ALDH1A2 | 8854 |
| ENSG00000107954.10 | 300.5978381 | 2.837725228 | 0.138565266 | 16.14925 | 1.15E-58 | 2.10E-56 | ENSG00000107954 | NEURL1 | 9148 |
| ENSG00000008056.14 | 5.573972 | 2.836893 | 0.6778686 | 3.299891 | 0.000967 | 0.007190572 | ENSG00000008056 | SYN1 | 6853 |
| ENSG00000100167.20 | 114.2279 | 2.836438 | 0.2205807 | 10.13886 | 3.71E-24 | 1.78E-22 | ENSG00000100167 | SEPTIN3 | 55964 |
| ENSG00000068976.14 | 11.04739075 | 2.830635951 | 0.549992735 | 4.055755 | 5.00E-05 | 0.000472432 | ENSG00000068976 | PYGM | 5837 |

| | | | | | | | | | |
|---|---|---|---|---|---|---|---|---|---|
| ENSG00000111305.19 | 5.141488935 | 2.829698676 | 0.749815974 | 2.973661 | 0.002943 | 0.019541158 | ENSG00000111305 | GSG1 | 83445 |
| ENSG00000156500.15 | 230.5752 | 2.826459 | 0.2909075 | 7.653492 | 1.96E-14 | 5.11E-13 | ENSG00000156500 | FAM122C | 159091 |
| ENSG00000287175.1 | 7.048676504 | 2.826172616 | 0.802533303 | 2.773932 | 0.005538 | 0.034209143 | ENSG00000287175 | NA | NA |
| ENSG00000204869.8 | 9.612514 | 2.825378 | 0.4962302 | 4.484568 | 7.31E-06 | 7.92E-05 | ENSG00000204869 | IGFL4 | 444882 |
| ENSG00000160325.14 | 60.61599528 | 2.824997252 | 0.299090027 | 7.439222 | 1.01E-13 | 2.52E-12 | ENSG00000160325 | CACFD1 | 11094 |
| ENSG00000264448.5 | 13.91213799 | 2.823987294 | 0.645565785 | 3.44502 | 0.000571 | 0.004464492 | ENSG00000264448 | NA | NA |
| ENSG00000102265.12 | 15980.73 | 2.823486 | 0.1375353 | 16.16666 | 8.67E-59 | 1.59E-56 | ENSG00000102265 | TIMP1 | 7076 |
| ENSG00000272192.1 | 4.52012792 | 2.823450331 | 0.741316387 | 2.999327 | 0.002706 | 0.018139742 | ENSG00000272192 | NA | NA |
| ENSG00000170289.12 | 103.9932333 | 2.82270816 | 0.191546988 | 11.60398 | 3.93E-31 | 2.85E-29 | ENSG00000170289 | CNGB3 | 54714 |
| ENSG00000234828.8 | 8.917942533 | 2.819595767 | 0.646015978 | 3.435822 | 0.000591 | 0.004610729 | ENSG00000234828 | IQCM | 285423 |
| ENSG00000265907.2 | 5.10589899 | 2.817846074 | 0.760409341 | 2.916648 | 0.003538 | 0.022990334 | ENSG00000265907 | NA | NA |
| ENSG00000149131.15 | 279.17815 | 2.817809679 | 0.258291636 | 8.586456 | 8.97E-18 | 2.99E-16 | ENSG00000149131 | SERPING1 | 710 |
| ENSG00000092969.12 | 67.99881458 | 2.811713827 | 0.338437241 | 6.535078 | 6.36E-11 | 1.23E-09 | ENSG00000092969 | TGFB2 | 7042 |
| ENSG00000166183.16 | 28.21418587 | 2.810625191 | 0.440937887 | 5.013462 | 5.35E-07 | 6.69E-06 | ENSG00000166183 | ASPG | 374569 |
| ENSG00000132874.14 | 5.327487939 | 2.809179858 | 0.81539111 | 2.70935 | 0.006742 | 0.040687537 | ENSG00000132874 | SLC14A2 | 8170 |
| ENSG00000267583.5 | 9.550436081 | 2.807477204 | 0.627407247 | 3.518412 | 0.000434 | 0.003473885 | ENSG00000267583 | NA | NA |
| ENSG00000259230.1 | 172.8399564 | 2.806499518 | 0.161802854 | 13.63696 | 2.41E-42 | 2.64E-40 | ENSG00000259230 | LINC0232 | 1.05E+08 |
| ENSG00000087074.8 | 6751.355 | 2.80618 | 0.2237653 | 9.859349 | 6.25E-23 | 2.82E-21 | ENSG00000087074 | PPP1R15A | 23645 |
| ENSG00000253315.1 | 15.80352422 | 2.805542859 | 0.404330363 | 5.454804 | 4.90E-08 | 6.95E-07 | ENSG00000253315 | LINC0193 | 1.05E+08 |
| ENSG00000100302.7 | 21.24881 | 2.80243 | 0.5517223 | 3.991917 | 6.55E-05 | 0.000609233 | ENSG00000100302 | RASD2 | 23551 |
| ENSG00000213626.13 | 78.53405016 | 2.802328008 | 0.280085162 | 7.863066 | 3.75E-15 | 1.04E-13 | ENSG00000213626 | LBH | 81606 |
| ENSG00000284882.1 | 6.101817435 | 2.798396853 | 0.637057451 | 3.450861 | 0.000559 | 0.004381052 | ENSG00000284882 | LOC10537 | 1.05E+08 |
| ENSG00000123836.15 | 7886.109124 | 2.797912236 | 0.215855655 | 10.18232 | 2.38E-24 | 1.16E-22 | ENSG00000123836 | PFKFB2 | 5208 |
| ENSG00000227070.1 | 9.293179061 | 2.797041258 | 0.586016365 | 3.749112 | 0.000177 | 0.001527937 | ENSG00000227070 | NA | NA |
| ENSG00000158825.6 | 2509.327644 | 2.794830204 | 0.12300374 | 17.84361 | 3.24E-71 | 8.14E-69 | ENSG00000158825 | CDA | 978 |
| ENSG00000261644.2 | 6.945386151 | 2.790980822 | 0.641526342 | 3.415262 | 0.000637 | 0.00493274 | ENSG00000261644 | NA | NA |
| ENSG00000146090.16 | 4.979585058 | 2.789115435 | 0.717008016 | 3.053125 | 0.002265 | 0.015471175 | ENSG00000146090 | RASGEF1 | 255426 |
| ENSG00000080031.10 | 125.254 | 2.786019 | 0.2706293 | 8.077542 | 6.61E-16 | 1.93E-14 | ENSG00000080031 | PTPRH | 5794 |
| ENSG00000228561.2 | 8.59623043 | 2.784364775 | 0.523320058 | 4.174051 | 2.99E-05 | 0.000293087 | ENSG00000228561 | NA | NA |
| ENSG00000143839.15 | 21.78958345 | 2.784349583 | 0.467242955 | 4.674976 | 2.94E-06 | 3.33E-05 | ENSG00000143839 | REN | 5972 |
| ENSG00000109220.11 | 1419.376163 | 2.783643167 | 0.15705916 | 13.90332 | 6.05E-44 | 7.03E-42 | ENSG00000109220 | CHIC2 | 26511 |
| ENSG00000285820.1 | 117.4485184 | 2.783055994 | 0.201702158 | 10.82317 | 2.67E-27 | 1.54E-25 | ENSG00000285820 | NA | NA |
| ENSG00000102313.8 | 23.20858 | 2.779989 | 0.3720443 | 5.859488 | 4.64E-09 | 7.42E-08 | ENSG00000102313 | ITIH6 | 347365 |
| ENSG00000092929.11 | 748.0417657 | 2.776969256 | 0.202602003 | 10.74505 | 6.25E-27 | 3.50E-25 | ENSG00000092929 | UNC13D | 201294 |
| ENSG00000135549.15 | 97.4483037 | 2.773723882 | 0.303560221 | 7.160767 | 8.02E-13 | 1.85E-11 | ENSG00000135549 | PKIB | 5570 |
| ENSG00000287978.1 | 20.07860713 | 2.770529905 | 0.340081837 | 6.382375 | 1.74E-10 | 3.27E-09 | ENSG00000287978 | LOC10099 | 1.01E+08 |
| ENSG00000255150.2 | 53.72028346 | 2.769829347 | 0.243364878 | 8.915951 | 4.84E-19 | 1.74E-17 | ENSG00000255150 | EID3 | 493861 |
| ENSG00000176040.13 | 10.55733445 | 2.768346388 | 0.562364655 | 3.855766 | 0.000115 | 0.001030992 | ENSG00000176040 | TMPRSS7 | 344805 |
| ENSG00000266467.2 | 5.440479235 | 2.766777935 | 0.786438663 | 2.755177 | 0.005866 | 0.035991141 | ENSG00000266467 | RN7SL220 | 1.06E+08 |
| ENSG00000196730.13 | 2182.968056 | 2.764701686 | 0.156596853 | 13.8234 | 1.84E-43 | 2.10E-41 | ENSG00000196730 | DAPK1 | 1612 |
| ENSG00000076864.19 | 586.8298917 | 2.764215254 | 0.245455933 | 8.817123 | 1.17E-18 | 4.12E-17 | ENSG00000076864 | RAP1GAP | 5909 |
| ENSG00000125735.10 | 172.7885975 | 2.763348397 | 0.567824262 | 3.809891 | 0.000139 | 0.001220715 | ENSG00000125735 | TNFSF14 | 8740 |

| | | | | | | | | | |
|---|---|---|---|---|---|---|---|---|---|
| ENSG00000159527.3 | 167.582485 | 2.762793137 | 0.554457648 | 3.900736 | 9.59E-05 | 0.000869276 | ENSG00000159527 | PGLYRP3 | 114771 |
| ENSG00000258279.4 | 41.50769626 | 2.761489578 | 0.261679268 | 8.260072 | 1.46E-16 | 4.45E-15 | ENSG00000258279 | LINC00592 | 283404 |
| ENSG00000086548.9 | 816.5081 | 2.756327 | 0.3854483 | 5.594336 | 2.21E-08 | 3.29E-07 | ENSG00000086548 | CEACAM6 | 4680 |
| ENSG00000242428.6 | 7.353336159 | 2.754713251 | 0.680237209 | 3.167591 | 0.001537 | 0.01089748 | ENSG00000242428 | CFAP20D( | 1.02E+08 |
| ENSG00000166278.15 | 4.003435912 | 2.754432959 | 0.796140565 | 2.706096 | 0.006808 | 0.041041841 | ENSG00000166278 | C2 | 717 |
| ENSG00000125845.7 | 594.4171 | 2.753819 | 0.2468964 | 8.723576 | 2.70E-18 | 9.28E-17 | ENSG00000125845 | BMP2 | 650 |
| ENSG00000267430.1 | 10.36860293 | 2.75101927 | 0.520722922 | 4.130833 | 3.61E-05 | 0.000350421 | ENSG00000267430 | LOC10042 | 1E+08 |
| ENSG00000099998.17 | 4.353419 | 2.747162 | 0.7602532 | 2.824272 | 0.004739 | 0.02983835 | ENSG00000099998 | GGT5 | 2687 |
| ENSG00000124171.9 | 1530.925 | 2.747103 | 0.200699 | 10.69813 | 1.04E-26 | 5.74E-25 | ENSG00000124171 | PARD6B | 84612 |
| ENSG00000257137.7 | 5.583829588 | 2.746112806 | 0.659319022 | 3.255045 | 0.001134 | 0.008305608 | ENSG00000257137 | NA | NA |
| ENSG00000198515.13 | 135.6812542 | 2.744661233 | 0.19897274 | 10.77867 | 4.34E-27 | 2.46E-25 | ENSG00000198515 | CNGA1 | 1259 |
| ENSG00000162814.11 | 54.01921986 | 2.74402067 | 0.297555731 | 7.205442 | 5.79E-13 | 1.35E-11 | ENSG00000162814 | SPATA17 | 128153 |
| ENSG00000236740.6 | 13.89805774 | 2.74315712 | 0.48122165 | 4.453576 | 8.45E-06 | 9.06E-05 | ENSG00000236740 | LOC10192 | 1.02E+08 |
| ENSG00000099860.9 | 2347.339121 | 2.742151488 | 0.138356366 | 15.48285 | 4.53E-54 | 7.18E-52 | ENSG00000099860 | GADD45B | 4616 |
| ENSG00000277619.1 | 6.819355034 | 2.739831278 | 0.780840313 | 2.740421 | 0.006136 | 0.037415014 | ENSG00000277619 | NA | NA |
| ENSG00000285581.1 | 5.876407706 | 2.738293932 | 0.747192603 | 2.861771 | 0.004213 | 0.026895339 | ENSG00000285581 | NA | NA |
| ENSG00000267308.2 | 7.498096 | 2.73666 | 0.5755263 | 3.712532 | 0.000205 | 0.001745837 | ENSG00000267308 | NA | NA |
| ENSG00000150347.16 | 1484.241811 | 2.735567383 | 0.229195761 | 9.317657 | 1.19E-20 | 4.77E-19 | ENSG00000150347 | ARID5B | 84159 |
| ENSG00000138413.13 | 11062.64163 | 2.729225126 | 0.143364146 | 14.85187 | 6.77E-50 | 9.48E-48 | ENSG00000138413 | IDH1 | 3417 |
| ENSG00000171931.12 | 33.78194261 | 2.729030589 | 0.271762115 | 7.83417 | 4.72E-15 | 1.30E-13 | ENSG00000171931 | FBXW10 | 10517 |
| ENSG00000274507.1 | 3.60882 | 2.728434 | 0.7870785 | 2.704221 | 0.006846 | 0.04125551 | ENSG00000274507 | NA | NA |
| ENSG00000220785.7 | 332.3903837 | 2.72840017 | 0.235659683 | 9.031669 | 1.69E-19 | 6.26E-18 | ENSG00000220785 | NA | NA |
| ENSG00000245522.2 | 69.00483991 | 2.724990335 | 0.290425523 | 7.316817 | 2.54E-13 | 6.11E-12 | ENSG00000245522 | LINC0270( | 440028 |
| ENSG00000245750.9 | 5.453488352 | 2.721899141 | 0.646282405 | 3.283238 | 0.001026 | 0.007590997 | ENSG00000245750 | DRAIC | 145837 |
| ENSG00000229502.5 | 9.035508797 | 2.721575255 | 0.594657122 | 3.567729 | 0.00036 | 0.00293428 | ENSG00000229502 | NA | NA |
| ENSG00000181982.18 | 732.540151 | 2.71991443 | 0.202832419 | 10.45156 | 1.44E-25 | 7.60E-24 | ENSG00000181982 | CCDC149 | 91050 |
| ENSG00000197191.5 | 367.2705806 | 2.716393024 | 0.32147955 | 6.58329 | 4.60E-11 | 9.03E-10 | ENSG00000197191 | CYSRT1 | 375791 |
| ENSG00000205177.6 | 44.581092 | 2.715402301 | 0.251646228 | 8.406255 | 4.23E-17 | 1.34E-15 | ENSG00000205177 | C11orf91 | 1E+08 |
| ENSG00000203722.8 | 258.6816411 | 2.714633605 | 0.202662274 | 10.43427 | 1.73E-25 | 9.04E-24 | ENSG00000203722 | RAET1G | 353091 |
| ENSG00000226564.1 | 79.58948191 | 2.714148258 | 0.195778392 | 10.79868 | 3.49E-27 | 2.00E-25 | ENSG00000226564 | NA | NA |
| ENSG00000223617.2 | 9.117529736 | 2.713979192 | 0.527717903 | 4.005889 | 6.18E-05 | 0.000576124 | ENSG00000223617 | NA | NA |
| ENSG00000160111.13 | 44.96726 | 2.712171 | 0.3231564 | 6.536066 | 6.32E-11 | 1.22E-09 | ENSG00000160111 | CPAMD8 | 27151 |
| ENSG00000079482.13 | 977.1489 | 2.712128 | 0.2472173 | 8.543611 | 1.30E-17 | 4.28E-16 | ENSG00000079482 | OPHN1 | 4983 |
| ENSG00000273802.2 | 155.8721315 | 2.711266268 | 0.30444841 | 6.934726 | 4.07E-12 | 8.83E-11 | ENSG00000273802 | H2BC8 | 8339 |
| ENSG00000267322.2 | 5.568485512 | 2.711031413 | 0.717262607 | 2.943178 | 0.003249 | 0.0213174 | ENSG00000267322 | SCARNA1 | 677769 |
| ENSG00000064205.10 | 12.89804 | 2.710704 | 0.4238711 | 4.97959 | 6.37E-07 | 7.92E-06 | ENSG00000064205 | CCN5 | 8839 |
| ENSG00000102287.19 | 2034.163 | 2.708751 | 0.1292354 | 16.31713 | 7.46E-60 | 1.45E-57 | ENSG00000102287 | GABRE | 2564 |
| ENSG00000143536.7 | 26.88131418 | 2.708615517 | 0.495457998 | 4.255892 | 2.08E-05 | 0.00020894 | ENSG00000143536 | CRNN | 49860 |
| ENSG00000070729.13 | 54.46562725 | 2.704276344 | 0.335596593 | 6.270255 | 3.60E-10 | 6.56E-09 | ENSG00000070729 | CNGB1 | 1258 |
| ENSG00000139112.11 | 2865.748378 | 2.700305915 | 0.213108162 | 9.855586 | 6.48E-23 | 2.92E-21 | ENSG00000139112 | GABARAP | 23710 |
| ENSG00000136630.13 | 51.63379338 | 2.699639315 | 0.355155768 | 5.911883 | 3.38E-09 | 5.49E-08 | ENSG00000136630 | HLX | 3142 |
| ENSG00000249102.3 | 5.419809771 | 2.69897184 | 0.795172907 | 2.639642 | 0.008299 | 0.048753228 | ENSG00000249102 | NA | NA |

| | | | | | | | | | |
|---|---|---|---|---|---|---|---|---|---|
| ENSG00000134317.18 | 1592.418374 | 2.698213131 | 0.262757041 | 7.985374 | 1.40E-15 | 4.03E-14 | ENSG00000134317 | GRHL1 | 29841 |
| ENSG00000197083.11 | 5.328683569 | 2.694487011 | 0.754317139 | 2.776666 | 0.005492 | 0.033977866 | ENSG00000197083 | NA | NA |
| ENSG00000246740.2 | 44.69226797 | 2.694032772 | 0.452672881 | 4.625929 | 3.73E-06 | 4.18E-05 | ENSG00000246740 | PLA2G4E- | 1.02E+08 |
| ENSG00000165030.4 | 1722.695219 | 2.692625014 | 0.217759712 | 9.60979 | 7.27E-22 | 3.12E-20 | ENSG00000165030 | NFIL3 | 4783 |
| ENSG00000272650.1 | 10.67912906 | 2.691346963 | 0.521980287 | 4.006563 | 6.16E-05 | 0.000574683 | ENSG00000272650 | NA | NA |
| ENSG00000164309.15 | 337.3929637 | 2.690422361 | 0.272066932 | 7.683486 | 1.55E-14 | 4.07E-13 | ENSG00000164309 | CMYA5 | 202333 |
| ENSG00000141441.16 | 747.1688302 | 2.689626194 | 0.13696157 | 15.25703 | 1.48E-52 | 2.25E-50 | ENSG00000141441 | GAREM1 | 64762 |
| ENSG00000214145.7 | 26.56996105 | 2.689221295 | 0.347993013 | 6.00363 | 1.93E-09 | 3.21E-08 | ENSG00000214145 | LINC0088 | 1E+08 |
| ENSG00000105856.14 | 2119.57707 | 2.688971131 | 0.151959185 | 13.74692 | 5.31E-43 | 5.97E-41 | ENSG00000105856 | HBP1 | 26959 |
| ENSG00000186474.15 | 41.02618 | 2.687981 | 0.269338 | 7.752271 | 9.03E-15 | 2.42E-13 | ENSG00000186474 | KLK12 | 43849 |
| ENSG00000248242.1 | 6.465725945 | 2.686986312 | 0.607800164 | 3.433672 | 0.000595 | 0.004643368 | ENSG00000248242 | NA | NA |
| ENSG00000176428.6 | 20.13842394 | 2.686233267 | 0.368522443 | 5.661075 | 1.50E-08 | 2.28E-07 | ENSG00000176428 | VPS37D | 155382 |
| ENSG00000103196.12 | 92.60628794 | 2.68596123 | 0.378064989 | 5.517467 | 3.44E-08 | 4.99E-07 | ENSG00000103196 | CRISPLD2 | 83716 |
| ENSG00000144218.18 | 5.5742784 | 2.685726959 | 0.69585186 | 2.997372 | 0.002723 | 0.0182473 | ENSG00000144218 | AFF3 | 3899 |
| ENSG00000257242.8 | 11.71094338 | 2.68471199 | 0.512810354 | 4.065269 | 4.80E-05 | 0.000455834 | ENSG00000257242 | LINC0161 | 256021 |
| ENSG00000184564.11 | 42.25539656 | 2.67678173 | 0.644159834 | 3.224016 | 0.001264 | 0.009151992 | ENSG00000184564 | SLITRK6 | 84189 |
| ENSG00000072952.18 | 14.07860852 | 2.672628281 | 0.402838935 | 5.145055 | 2.67E-07 | 3.49E-06 | ENSG00000072952 | IRAG1 | 10335 |
| ENSG00000280228.1 | 81.82171027 | 2.67252918 | 0.384811245 | 5.385833 | 7.21E-08 | 1.01E-06 | ENSG00000280228 | NA | NA |
| ENSG00000214212.9 | 5.553797778 | 2.671407105 | 0.672409036 | 3.080576 | 0.002066 | 0.014212203 | ENSG00000214212 | C19orf38 | 255809 |
| ENSG00000229950.1 | 114.2054676 | 2.670668532 | 0.185278537 | 11.17598 | 5.35E-29 | 3.47E-27 | ENSG00000229950 | NA | NA |
| ENSG00000124466.9 | 1919.96 | 2.67066 | 0.3672036 | 5.638997 | 1.71E-08 | 2.57E-07 | ENSG00000124466 | LYPD3 | 27076 |
| ENSG00000231971.6 | 6.518236 | 2.668866963 | 0.694487023 | 2.978986 | 0.002892 | 0.019252799 | ENSG00000231971 | CT69 | 1.02E+08 |
| ENSG00000088280.19 | 65.29092321 | 2.668114941 | 0.238316711 | 8.678011 | 4.03E-18 | 1.37E-16 | ENSG00000088280 | ASAP3 | 55616 |
| ENSG00000156265.15 | 164.9808 | 2.66562 | 0.1571428 | 13.14486 | 1.82E-39 | 1.79E-37 | ENSG00000156265 | MAP3K7C | 56911 |
| ENSG00000197948.11 | 5030.066699 | 2.659657454 | 0.350389453 | 5.878195 | 4.15E-09 | 6.66E-08 | ENSG00000197948 | FCHSD1 | 89848 |
| ENSG00000151715.7 | 171.7408564 | 2.659065767 | 0.295628278 | 6.96505 | 3.28E-12 | 7.18E-11 | ENSG00000151715 | TMEM45B | 120224 |
| ENSG00000286373.1 | 7.427910462 | 2.65648245 | 0.650972717 | 3.159092 | 0.001583 | 0.011175713 | ENSG00000286373 | NA | NA |
| ENSG00000087589.16 | 8.14632 | 2.656063 | 0.5934158 | 3.464793 | 0.000531 | 0.004180738 | ENSG00000087589 | CASS4 | 57091 |
| ENSG00000238266.2 | 1200.087599 | 2.653088125 | 0.346281172 | 5.928963 | 3.05E-09 | 4.99E-08 | ENSG00000238266 | LINC0070 | 1.01E+08 |
| ENSG00000124116.19 | 4.039678 | 2.652392 | 0.75908 | 2.703789 | 0.006855 | 0.0412863 | ENSG00000124116 | WFDC3 | 140686 |
| ENSG00000181143.15 | 1042.057253 | 2.650349829 | 0.278367463 | 7.365623 | 1.76E-13 | 4.31E-12 | ENSG00000181143 | MUC16 | 94025 |
| ENSG00000204282.4 | 56.62407302 | 2.646933734 | 0.405485478 | 5.048106 | 4.46E-07 | 5.64E-06 | ENSG00000204282 | NA | NA |
| ENSG00000248588.2 | 32.18849605 | 2.645227519 | 0.354407053 | 5.770843 | 7.89E-09 | 1.24E-07 | ENSG00000248588 | NA | NA |
| ENSG00000197181.12 | 4.980381915 | 2.643287619 | 0.68901249 | 2.965531 | 0.003022 | 0.020012239 | ENSG00000197181 | PIWIL2 | 55124 |
| ENSG00000204540.10 | 38.24070764 | 2.642625233 | 0.466737648 | 4.376388 | 1.21E-05 | 0.000125971 | ENSG00000204540 | PSORS1C | 170679 |
| ENSG00000224116.7 | 46.53814808 | 2.642437286 | 0.285504472 | 7.153784 | 8.44E-13 | 1.94E-11 | ENSG00000224116 | INHBA-AS | 285954 |
| ENSG00000266891.1 | 11.44844851 | 2.640211453 | 0.533395898 | 3.824948 | 0.000131 | 0.001155709 | ENSG00000266891 | NA | NA |
| ENSG00000228903.7 | 6.807545633 | 2.636957245 | 0.666245289 | 3.057368 | 0.002233 | 0.015270648 | ENSG00000228903 | NA | NA |
| ENSG00000205364.4 | 19.15292382 | 2.636027527 | 0.528473435 | 3.852658 | 0.000117 | 0.001043118 | ENSG00000205364 | MT1M | 4499 |
| ENSG00000142173.15 | 289.467 | 2.635941 | 0.4157812 | 4.896664 | 9.75E-07 | 1.18E-05 | ENSG00000142173 | COL6A2 | 1292 |
| ENSG00000112769.19 | 69.26195375 | 2.633601566 | 0.280695408 | 7.244869 | 4.33E-13 | 1.02E-11 | ENSG00000112769 | LAMA4 | 3910 |
| ENSG00000237353.6 | 5.21006331 | 2.63257442 | 0.73919645 | 2.749708 | 0.005965 | 0.036488036 | ENSG00000237353 | PATE4 | 399968 |

| | | | | | | | | | |
|---|---|---|---|---|---|---|---|---|---|
| ENSG00000231160.10 | 25.46320727 | 2.631642098 | 0.368352695 | 5.51548 | 3.48E-08 | 5.04E-07 | ENSG00000231160 | NA | NA |
| ENSG00000149212.11 | 77.21001865 | 2.630940703 | 0.220044158 | 9.229696 | 2.71E-20 | 1.06E-18 | ENSG00000149212 | SESN3 | 143686 |
| ENSG00000259771.1 | 5.411783719 | 2.630349373 | 0.676844758 | 2.999727 | 0.002702 | 0.018125084 | ENSG00000259771 | NA | NA |
| ENSG00000144655.15 | 2017.081409 | 2.627893286 | 0.209898514 | 9.661304 | 4.40E-22 | 1.90E-20 | ENSG00000144655 | CSRNP1 | 64651 |
| ENSG00000128422.17 | 156457.3598 | 2.626233084 | 0.220053962 | 9.207892 | 3.33E-20 | 1.29E-18 | ENSG00000128422 | KRT17 | 3872 |
| ENSG00000198246.9 | 243.8859085 | 2.623175483 | 0.201640403 | 10.03358 | 1.09E-23 | 5.11E-22 | ENSG00000198246 | SLC29A3 | 55315 |
| ENSG00000188037.11 | 9.786383231 | 2.622749554 | 0.541362601 | 3.736404 | 0.000187 | 0.001599466 | ENSG00000188037 | CLCN1 | 1180 |
| ENSG00000232938.2 | 18.55091666 | 2.620591728 | 0.412807324 | 4.894757 | 9.84E-07 | 1.19E-05 | ENSG00000232938 | RPL23AP8 | 388574 |
| ENSG00000184489.11 | 12.22830239 | 2.619421703 | 0.498829784 | 4.048318 | 5.16E-05 | 0.000486485 | ENSG00000184489 | PTP4A3 | 11156 |
| ENSG00000133328.4 | 32.36123307 | 2.618435698 | 0.508422897 | 3.969994 | 7.19E-05 | 0.000663469 | ENSG00000133328 | PLAAT2 | 54979 |
| ENSG00000141854.9 | 15.90727884 | 2.617149611 | 0.462501669 | 4.361389 | 1.29E-05 | 0.000134451 | ENSG00000141854 | MISP3 | 113230 |
| ENSG00000158486.13 | 162.421869 | 2.616597162 | 0.223249785 | 9.032919 | 1.67E-19 | 6.20E-18 | ENSG00000158486 | DNAH3 | 55567 |
| ENSG00000228221.6 | 21.80746529 | 2.612043865 | 0.413065946 | 4.870999 | 1.11E-06 | 1.33E-05 | ENSG00000228221 | LINC0057ε | 1.01E+08 |
| ENSG00000179869.15 | 34.68851577 | 2.610884621 | 0.28098935 | 7.156444 | 8.28E-13 | 1.91E-11 | ENSG00000179869 | ABCA13 | 154664 |
| ENSG00000175197.12 | 502.639463 | 2.610815207 | 0.234841827 | 8.562424 | 1.11E-17 | 3.65E-16 | ENSG00000175197 | DDIT3 | 1649 |
| ENSG00000059728.11 | 4127.871824 | 2.610811119 | 0.223631626 | 8.991622 | 2.44E-19 | 8.97E-18 | ENSG00000059728 | MXD1 | 4084 |
| ENSG00000174844.14 | 7.811820603 | 2.610430046 | 0.656856473 | 3.060684 | 0.002208 | 0.015113089 | ENSG00000174844 | DNAH12 | 201625 |
| ENSG00000163218.15 | 143.4101096 | 2.608339187 | 0.310589312 | 6.466221 | 1.00E-10 | 1.91E-09 | ENSG00000163218 | PGLYRP4 | 57115 |
| ENSG00000114796.16 | 1084.401665 | 2.602606551 | 0.167748575 | 11.93814 | 7.49E-33 | 5.79E-31 | ENSG00000114796 | KLHL24 | 54800 |
| ENSG00000153233.13 | 9.504075618 | 2.600426613 | 0.526811101 | 3.797237 | 0.000146 | 0.001279657 | ENSG00000153233 | PTPRR | 5801 |
| ENSG00000285736.1 | 28.53632924 | 2.600175404 | 0.348010856 | 5.747451 | 9.06E-09 | 1.41E-07 | ENSG00000285736 | NA | NA |
| ENSG00000225339.4 | 26.37740391 | 2.599721753 | 0.345436184 | 5.788976 | 7.08E-09 | 1.11E-07 | ENSG00000225339 | NA | NA |
| ENSG00000105048.17 | 1223.021 | 2.599518 | 0.1854646 | 10.78113 | 4.23E-27 | 2.40E-25 | ENSG00000105048 | TNNT1 | 7138 |
| ENSG00000166596.15 | 28.48210349 | 2.594470413 | 0.361613656 | 5.515473 | 3.48E-08 | 5.04E-07 | ENSG00000166596 | CFAP52 | 146845 |
| ENSG00000272476.1 | 187.4356673 | 2.593120292 | 0.199490952 | 9.991031 | 1.67E-23 | 7.78E-22 | ENSG00000272476 | NA | NA |
| ENSG00000108821.13 | 440.9445453 | 2.592669262 | 0.222536443 | 8.95435 | 3.42E-19 | 1.24E-17 | ENSG00000108821 | COL1A1 | 1277 |
| ENSG00000179031.8 | 10.04963 | 2.592234 | 0.6768367 | 2.943448 | 0.003246 | 0.02130406 | ENSG00000179031 | NA | NA |
| ENSG00000035664.11 | 39.19640754 | 2.590846576 | 0.299828188 | 6.639958 | 3.14E-11 | 6.28E-10 | ENSG00000035664 | DAPK2 | 23604 |
| ENSG00000204947.9 | 56.05819669 | 2.588772384 | 0.236643849 | 8.404074 | 4.31E-17 | 1.37E-15 | ENSG00000204947 | ZNF425 | 155054 |
| ENSG00000124216.4 | 10.82991 | 2.588372 | 0.6017209 | 3.304475 | 0.000952 | 0.007087855 | ENSG00000124216 | SNAI1 | 6615 |
| ENSG00000100170.9 | 694.4764 | 2.587293 | 0.5399021 | 3.680839 | 0.000232 | 0.001960947 | ENSG00000100170 | SLC5A1 | 6523 |
| ENSG00000136059.14 | 506.4011326 | 2.586332401 | 0.240481386 | 8.259818 | 1.46E-16 | 4.45E-15 | ENSG00000136059 | VILL | 50853 |
| ENSG00000130208.9 | 17.74318 | 2.5861 | 0.3905743 | 5.085076 | 3.67E-07 | 4.70E-06 | ENSG00000130208 | APOC1 | 341 |
| ENSG00000072201.13 | 197.8015368 | 2.583959127 | 0.18641468 | 10.64272 | 1.89E-26 | 1.02E-24 | ENSG00000072201 | LNX1 | 84708 |
| ENSG00000237852.1 | 4.412778064 | 2.577759409 | 0.72807754 | 2.716413 | 0.006599 | 0.039929071 | ENSG00000237852 | NA | NA |
| ENSG00000109943.9 | 10.52787217 | 2.576685117 | 0.509834374 | 3.877112 | 0.000106 | 0.000950368 | ENSG00000109943 | CRTAM | 56253 |
| ENSG00000130940.15 | 602.7737624 | 2.576079515 | 0.235977706 | 8.374009 | 5.57E-17 | 1.75E-15 | ENSG00000130940 | CASZ1 | 54897 |
| ENSG00000128016.6 | 3892.328 | 2.575979 | 0.3157013 | 6.259014 | 3.87E-10 | 7.01E-09 | ENSG00000128016 | ZFP36 | 7538 |
| ENSG00000100003.18 | 4014.713 | 2.57463 | 0.1449879 | 13.61927 | 3.08E-42 | 3.33E-40 | ENSG00000100003 | SEC14L2 | 23541 |
| ENSG00000162069.15 | 685.3739895 | 2.5710403 | 0.184638978 | 10.6751 | 1.33E-26 | 7.31E-25 | ENSG00000162069 | BICDL2 | 146439 |
| ENSG00000249364.6 | 9.262781155 | 2.569681822 | 0.515621007 | 3.820019 | 0.000133 | 0.001177879 | ENSG00000249364 | LOC10192 | 1.02E+08 |
| ENSG00000249899.6 | 27.61437172 | 2.566737976 | 0.370243928 | 5.312006 | 1.08E-07 | 1.49E-06 | ENSG00000249899 | NA | NA |

| | | | | | | | | | |
|---|---|---|---|---|---|---|---|---|---|
| ENSG00000124762.13 | 13431.83225 | 2.566657153 | 0.211337053 | 9.305785 | 1.33E-20 | 5.32E-19 | ENSG00000124762 | CDKN1A | 1026 |
| ENSG00000231969.2 | 8.734105847 | 2.565219652 | 0.533991081 | 3.680248 | 0.000233 | 0.001964255 | ENSG00000231969 | MMADHC- | 1.02E+08 |
| ENSG00000119138.4 | 377.4186046 | 2.564948993 | 0.262940968 | 7.472966 | 7.84E-14 | 1.97E-12 | ENSG00000119138 | KLF9 | 687 |
| ENSG00000283554.1 | 50.74132907 | 2.56424594 | 0.3154893 | 6.22603 | 4.78E-10 | 8.56E-09 | ENSG00000283554 | LINC0234 | 1.05E+08 |
| ENSG00000186710.11 | 13.85825544 | 2.563623421 | 0.462867775 | 4.242299 | 2.21E-05 | 0.000221175 | ENSG00000186710 | CFAP73 | 387885 |
| ENSG00000006606.8 | 6.5344612 | 2.561479596 | 0.732811625 | 2.676649 | 0.007436 | 0.044327755 | ENSG00000006606 | CCL26 | 10344 |
| ENSG00000276136.1 | 25.45755189 | 2.560846661 | 0.486717626 | 4.028715 | 5.61E-05 | 0.000526642 | ENSG00000276136 | NA | NA |
| ENSG00000143469.19 | 514.2341548 | 2.55560213 | 0.116112629 | 16.84229 | 1.20E-63 | 2.49E-61 | ENSG00000143469 | SYT14 | 255928 |
| ENSG00000255992.1 | 13.9155675 | 2.555582533 | 0.606122746 | 3.22638 | 0.001254 | 0.009081661 | ENSG00000255992 | NA | NA |
| ENSG00000286952.1 | 4.873525023 | 2.551388905 | 0.684845914 | 2.849384 | 0.00438 | 0.027878315 | ENSG00000286952 | NA | NA |
| ENSG00000120594.17 | 132.3793861 | 2.547124122 | 0.298045837 | 6.532969 | 6.45E-11 | 1.25E-09 | ENSG00000120594 | PLXDC2 | 84898 |
| ENSG00000105419.17 | 84.137 | 2.546001 | 0.1943699 | 10.01184 | 1.35E-23 | 6.34E-22 | ENSG00000105419 | MEIS3 | 56917 |
| ENSG00000173805.15 | 99.50830693 | 2.544835202 | 0.223011405 | 8.720788 | 2.76E-18 | 9.48E-17 | ENSG00000173805 | HAP1 | 9001 |
| ENSG00000112053.13 | 4.803320979 | 2.544831126 | 0.682622869 | 2.849057 | 0.004385 | 0.02789368 | ENSG00000112053 | SLC26A8 | 116369 |
| ENSG00000133805.15 | 2489.395536 | 2.543671925 | 0.177106162 | 10.97461 | 5.06E-28 | 3.07E-26 | ENSG00000133805 | AMPD3 | 272 |
| ENSG00000112378.12 | 73155.79312 | 2.54262121 | 0.132560447 | 14.65461 | 1.26E-48 | 1.68E-46 | ENSG00000112378 | PERP | 64065 |
| ENSG00000165795.23 | 1373.683924 | 2.541365159 | 0.164152045 | 11.82663 | 2.84E-32 | 2.15E-30 | ENSG00000165795 | NDRG2 | 57447 |
| ENSG00000229585.4 | 17.33957735 | 2.540918447 | 0.345591526 | 5.616221 | 1.95E-08 | 2.92E-07 | ENSG00000229585 | NA | NA |
| ENSG00000108840.15 | 1100.130769 | 2.5405251 | 0.176621018 | 10.98694 | 4.42E-28 | 2.70E-26 | ENSG00000108840 | HDAC5 | 10014 |
| ENSG00000206559.8 | 8.549961174 | 2.539823642 | 0.50575815 | 3.835477 | 0.000125 | 0.001111727 | ENSG00000206559 | ZCWPW2 | 152098 |
| ENSG00000253395.1 | 6.587435735 | 2.539385655 | 0.577121878 | 3.360444 | 0.000778 | 0.005920702 | ENSG00000253395 | NA | NA |
| ENSG00000135924.15 | 1937.108858 | 2.538847495 | 0.08743032 | 22.17592 | 5.87E-109 | 2.74E-106 | ENSG00000135924 | DNAJB2 | 3300 |
| ENSG00000225511.7 | 10.15407826 | 2.537875822 | 0.583797011 | 3.319434 | 0.000902 | 0.006768053 | ENSG00000225511 | NA | NA |
| ENSG00000246223.9 | 24.40820661 | 2.537448873 | 0.340255065 | 5.694107 | 1.24E-08 | 1.89E-07 | ENSG00000246223 | LINC0155 | 388011 |
| ENSG00000284968.1 | 160.4229682 | 2.535286128 | 0.172827889 | 11.19777 | 4.18E-29 | 2.74E-27 | ENSG00000284968 | NA | NA |
| ENSG00000175984.15 | 2139.52544 | 2.534856338 | 0.188210629 | 10.28027 | 8.65E-25 | 4.32E-23 | ENSG00000175984 | DENND2C | 163259 |
| ENSG00000166250.12 | 1417.864155 | 2.533521055 | 0.207455854 | 9.320157 | 1.16E-20 | 4.66E-19 | ENSG00000166250 | CLMP | 79827 |
| ENSG00000140297.13 | 51.89941394 | 2.532677057 | 0.27997302 | 6.903083 | 5.09E-12 | 1.10E-10 | ENSG00000140297 | GCNT3 | 9245 |
| ENSG00000254109.6 | 14.04488876 | 2.531040472 | 0.430256334 | 4.488116 | 7.19E-06 | 7.80E-05 | ENSG00000254109 | RBPMS-A | 1E+08 |
| ENSG00000144452.15 | 2852.714952 | 2.530942657 | 0.150599815 | 12.82168 | 1.24E-37 | 1.14E-35 | ENSG00000144452 | ABCA12 | 26154 |
| ENSG00000138669.9 | 76.77633799 | 2.528886017 | 0.194547812 | 9.914715 | 3.59E-23 | 1.64E-21 | ENSG00000138669 | PRKG2 | 5593 |
| ENSG00000137558.9 | 172.5558823 | 2.52830736 | 0.219392858 | 8.78929 | 1.51E-18 | 5.24E-17 | ENSG00000137558 | PI15 | 51050 |
| ENSG00000188089.13 | 1022.444114 | 2.527976031 | 0.502846802 | 3.834122 | 0.000126 | 0.0011169 | ENSG00000188089 | PLA2G4E | 123745 |
| ENSG00000133687.16 | 780.7114321 | 2.527866669 | 0.111443472 | 17.29905 | 4.78E-67 | 1.08E-64 | ENSG00000133687 | TMTC1 | 83857 |
| ENSG00000183171.5 | 5.354065 | 2.526486 | 0.6970624 | 2.763721 | 0.005715 | 0.03516761 | ENSG00000183171 | LOC64301 | 643015 |
| ENSG00000175040.6 | 2880.503514 | 2.526201032 | 0.210554918 | 9.148212 | 5.79E-20 | 2.20E-18 | ENSG00000175040 | CHST2 | 9435 |
| ENSG00000130592.15 | 27.86976151 | 2.523986027 | 0.283257707 | 6.792352 | 1.10E-11 | 2.30E-10 | ENSG00000130592 | LSP1 | 4046 |
| ENSG00000173227.14 | 362.2158105 | 2.521357702 | 0.233912584 | 8.213999 | 2.14E-16 | 6.45E-15 | ENSG00000173227 | SYT12 | 91683 |
| ENSG00000198892.6 | 27.80303428 | 2.519192324 | 0.317460481 | 6.045453 | 1.49E-09 | 2.51E-08 | ENSG00000198892 | SHISA4 | 149345 |
| ENSG00000165495.16 | 7.035889733 | 2.515586821 | 0.564350058 | 3.394324 | 0.000688 | 0.005298081 | ENSG00000165495 | PKNOX2 | 63876 |
| ENSG00000232187.1 | 8.845500663 | 2.510709547 | 0.611949855 | 3.12233 | 0.001794 | 0.012540474 | ENSG00000232187 | NA | NA |
| ENSG00000138131.4 | 60.92780053 | 2.508617958 | 0.307936069 | 6.198098 | 5.71E-10 | 1.01E-08 | ENSG00000138131 | LOXL4 | 84171 |

| | | | | | | | | | |
|---|---|---|---|---|---|---|---|---|---|
| ENSG00000146411.5 | 141.004925 | 2.50860336 | 0.174970747 | 10.90813 | 1.05E-27 | 6.21E-26 | ENSG00000146411 | SLC2A12 | 154091 |
| ENSG00000164463.12 | 2549.303685 | 2.508243365 | 0.154649268 | 12.33917 | 5.57E-35 | 4.56E-33 | ENSG00000164463 | CREBRF | 153222 |
| ENSG00000246526.2 | 5.486869624 | 2.506746005 | 0.680347458 | 2.802606 | 0.005069 | 0.031656339 | ENSG00000246526 | LINC0248 | 339988 |
| ENSG00000205129.8 | 15.4280533 | 2.500480381 | 0.385752919 | 4.926678 | 8.36E-07 | 1.02E-05 | ENSG00000205129 | C4orf47 | 441054 |
| ENSG00000203814.6 | 18.13842689 | 2.498483868 | 0.575833975 | 3.296929 | 0.000977 | 0.007260744 | ENSG00000203814 | H2BC18 | 440689 |
| ENSG00000244459.2 | 14.18145331 | 2.498240347 | 0.391901244 | 4.84367 | 1.27E-06 | 1.51E-05 | ENSG00000244459 | NA | NA |
| ENSG00000262096.2 | 4.118214007 | 2.496722225 | 0.710128272 | 2.670957 | 0.007564 | 0.044965669 | ENSG00000262096 | NA | NA |
| ENSG00000219986.2 | 6.395419753 | 2.495502114 | 0.649163689 | 2.919914 | 0.003501 | 0.02279557 | ENSG00000219986 | NA | NA |
| ENSG00000139971.15 | 50.45295897 | 2.490499147 | 0.211411383 | 8.942277 | 3.81E-19 | 1.38E-17 | ENSG00000139971 | ARMH4 | 145407 |
| ENSG00000287390.1 | 38.67474922 | 2.489017955 | 0.312474416 | 6.045352 | 1.49E-09 | 2.51E-08 | ENSG00000287390 | NA | NA |
| ENSG00000146674.15 | 288.8011295 | 2.488463534 | 0.190324868 | 9.922316 | 3.33E-23 | 1.52E-21 | ENSG00000146674 | IGFBP3 | 3486 |
| ENSG00000280162.1 | 22.2467 | 2.487627 | 0.4492693 | 4.201549 | 2.65E-05 | 0.000261966 | ENSG00000280162 | NA | NA |
| ENSG00000167306.20 | 2211.205089 | 2.487176904 | 0.219868773 | 8.583197 | 9.23E-18 | 3.07E-16 | ENSG00000167306 | MYO5B | 4645 |
| ENSG00000141738.14 | 3955.20459 | 2.4867102 | 0.23548349 | 8.01207 | 1.13E-15 | 3.25E-14 | ENSG00000141738 | GRB7 | 2886 |
| ENSG00000124920.13 | 353.5421583 | 2.485027915 | 0.43468778 | 4.33651 | 1.45E-05 | 0.000149086 | ENSG00000124920 | MYRF | 745 |
| ENSG00000175482.9 | 452.8470559 | 2.484183811 | 0.204234701 | 9.225581 | 2.82E-20 | 1.10E-18 | ENSG00000175482 | POLD4 | 57804 |
| ENSG00000225442.2 | 8.831092196 | 2.483324034 | 0.5631482 | 3.344278 | 0.000825 | 0.006248186 | ENSG00000225442 | NA | NA |
| ENSG00000258602.1 | 95.90801218 | 2.48312405 | 0.201903379 | 9.326858 | 1.09E-20 | 4.39E-19 | ENSG00000258602 | LINC0162 | 1.05E+08 |
| ENSG00000122877.16 | 10.28531045 | 2.481335678 | 0.677322923 | 2.777605 | 0.005476 | 0.033903494 | ENSG00000122877 | EGR2 | 1959 |
| ENSG00000272686.1 | 402.2376284 | 2.480633226 | 0.291871645 | 6.443357 | 1.17E-10 | 2.21E-09 | ENSG00000272686 | NA | NA |
| ENSG00000235448.3 | 16.29772589 | 2.47886553 | 0.363431876 | 5.169787 | 2.34E-07 | 3.08E-06 | ENSG00000235448 | LURAP1L- | 1.02E+08 |
| ENSG00000167767.14 | 7894.996712 | 2.477388238 | 0.189105009 | 9.927755 | 3.15E-23 | 1.45E-21 | ENSG00000167767 | KRT80 | 144501 |
| ENSG00000178226.10 | 29.22608755 | 2.47645983 | 0.330173666 | 5.683251 | 1.32E-08 | 2.01E-07 | ENSG00000178226 | PRSS36 | 146547 |
| ENSG00000254349.6 | 61.27681044 | 2.474885423 | 0.279990569 | 6.696245 | 2.14E-11 | 4.35E-10 | ENSG00000254349 | MIR2052H | 441355 |
| ENSG00000119801.13 | 3949.08902 | 2.474185999 | 0.134337736 | 13.9513 | 3.09E-44 | 3.64E-42 | ENSG00000119801 | YPEL5 | 51646 |
| ENSG00000085662.14 | 23201.89175 | 2.472605298 | 0.05428204 | 34.4977 | 8.69E-261 | 1.36E-257 | ENSG00000085662 | AKR1B1 | 231 |
| ENSG00000067082.15 | 20919.03492 | 2.472572475 | 0.031006579 | 60.39275 | 0 | 0 | ENSG00000067082 | KLF6 | 1316 |
| ENSG00000070731.10 | 330.3537042 | 2.468399641 | 0.285615275 | 6.541666 | 6.08E-11 | 1.18E-09 | ENSG00000070731 | ST6GALN | 10610 |
| ENSG00000003436.16 | 74.93462006 | 2.468149234 | 0.239494204 | 7.800394 | 6.17E-15 | 1.68E-13 | ENSG00000003436 | TFPI | 7035 |
| ENSG00000132518.6 | 12.47614542 | 2.46737157 | 0.43081138 | 4.334546 | 1.46E-05 | 0.000150365 | ENSG00000132518 | GUCY2D | 3000 |
| ENSG00000225190.11 | 976.7699942 | 2.464882402 | 0.17461918 | 10.67971 | 1.27E-26 | 6.97E-25 | ENSG00000225190 | PLEKHM1 | 9842 |
| ENSG00000089163.4 | 18.24386674 | 2.464216248 | 0.412309939 | 4.521395 | 6.14E-06 | 6.71E-05 | ENSG00000089163 | SIRT4 | 23409 |
| ENSG00000161011.20 | 17184.87644 | 2.464173741 | 0.215312372 | 8.657996 | 4.80E-18 | 1.63E-16 | ENSG00000161011 | SQSTM1 | 8878 |
| ENSG00000137878.17 | 597.2968196 | 2.460933277 | 0.171765675 | 10.83414 | 2.37E-27 | 1.38E-25 | ENSG00000137878 | GCOM1 | 145781 |
| ENSG00000109654.15 | 1476.956753 | 2.460040577 | 0.132221847 | 14.06757 | 6.01E-45 | 7.21E-43 | ENSG00000109654 | TRIM2 | 23321 |
| ENSG00000185156.5 | 26.63777828 | 2.459744746 | 0.473985446 | 3.923633 | 8.72E-05 | 0.000795502 | ENSG00000185156 | MFSD6L | 162387 |
| ENSG00000203497.2 | 19.16518414 | 2.459548535 | 0.384333721 | 4.838369 | 1.31E-06 | 1.55E-05 | ENSG00000203497 | NA | NA |
| ENSG00000281327.2 | 46.40113906 | 2.458928058 | 0.244728355 | 7.595883 | 3.06E-14 | 7.86E-13 | ENSG00000281327 | NA | NA |
| ENSG00000249395.3 | 285.1572228 | 2.456630492 | 0.208547977 | 8.902654 | 5.45E-19 | 1.96E-17 | ENSG00000249395 | CASC9 | 1.02E+08 |
| ENSG00000173156.7 | 6546.460909 | 2.455994615 | 0.115540261 | 16.06362 | 4.59E-58 | 8.26E-56 | ENSG00000173156 | RHOD | 29984 |
| ENSG00000155980.12 | 13.36575502 | 2.455590148 | 0.390067994 | 4.757094 | 1.96E-06 | 2.28E-05 | ENSG00000155980 | KIF5A | 3798 |
| ENSG00000113494.17 | 9.190830415 | 2.455226417 | 0.528475221 | 3.510527 | 0.000447 | 0.003573195 | ENSG00000113494 | PRLR | 5618 |

| | | | | | | | | | |
|---|---|---|---|---|---|---|---|---|---|
| ENSG00000075223.14 | 14305.55619 | 2.455124756 | 0.136119103 | 13.62869 | 2.70E-42 | 2.94E-40 | ENSG00000075223 | SEMA3C | 10512 |
| ENSG00000231233.1 | 11.92644168 | 2.45456736 | 0.509466224 | 3.640217 | 0.000272 | 0.002268545 | ENSG00000231233 | NA | NA |
| ENSG00000287067.1 | 9.648062616 | 2.447314647 | 0.675565548 | 2.734471 | 0.006248 | 0.038019581 | ENSG00000287067 | NA | NA |
| ENSG00000224034.1 | 6.306800502 | 2.446332647 | 0.692155852 | 2.66751 | 0.007642 | 0.045358675 | ENSG00000224034 | LINC0256' | 1.11E+08 |
| ENSG00000156804.7 | 1522.601621 | 2.445396419 | 0.185452491 | 9.950777 | 2.50E-23 | 1.16E-21 | ENSG00000156804 | FBXO32 | 114907 |
| ENSG00000012822.16 | 2847.800599 | 2.439750269 | 0.206107513 | 8.926168 | 4.41E-19 | 1.60E-17 | ENSG00000012822 | CALCOCC | 57658 |
| ENSG00000180438.15 | 36.61559815 | 2.438503608 | 0.354985996 | 5.179088 | 2.23E-07 | 2.94E-06 | ENSG00000180438 | TPRXL | 348825 |
| ENSG00000182732.18 | 16.86139196 | 2.435740218 | 0.41636047 | 4.409017 | 1.04E-05 | 0.00010957 | ENSG00000182732 | RGS6 | 9628 |
| ENSG00000182574.8 | 13.1835 | 2.433557 | 0.511259 | 3.586356 | 0.000335 | 0.002745978 | ENSG00000182574 | CSGALNA | 286453 |
| ENSG00000167895.14 | 24.43657099 | 2.426160428 | 0.459856754 | 3.971151 | 7.15E-05 | 0.000660484 | ENSG00000167895 | TMC8 | 147138 |
| ENSG00000163898.10 | 106.5460286 | 2.425805858 | 0.231696701 | 7.880155 | 3.27E-15 | 9.13E-14 | ENSG00000163898 | LIPH | 200879 |
| ENSG00000255587.9 | 13.96667951 | 2.424333469 | 0.387186323 | 4.711771 | 2.46E-06 | 2.81E-05 | ENSG00000255587 | RAB44 | 401258 |
| ENSG00000181016.9 | 95.83474363 | 2.424114736 | 0.328023825 | 5.560921 | 2.68E-08 | 3.96E-07 | ENSG00000181016 | LSMEM1 | 286006 |
| ENSG00000166546.14 | 25.09137561 | 2.423103552 | 0.527575821 | 3.455624 | 0.000549 | 0.004312788 | ENSG00000166546 | BEAN1 | 146227 |
| ENSG00000241458.1 | 7.65429332 | 2.422102516 | 0.596376327 | 3.05529 | 0.002248 | 0.015371788 | ENSG00000241458 | RPL7P19 | 1E+08 |
| ENSG00000140470.14 | 51.71052722 | 2.417892022 | 0.235445966 | 7.721058 | 1.15E-14 | 3.07E-13 | ENSG00000140470 | ADAMTS1 | 170691 |
| ENSG00000111644.8 | 13.93825504 | 2.41588515 | 0.48379927 | 3.753385 | 0.000174 | 0.001504671 | ENSG00000111644 | ACRBP | 84519 |
| ENSG00000166016.6 | 628.88663 | 2.414176292 | 0.117093605 | 15.49338 | 3.85E-54 | 6.13E-52 | ENSG00000166016 | ABTB2 | 25841 |
| ENSG00000159871.15 | 809.1436 | 2.413437 | 0.4094516 | 4.42894 | 9.47E-06 | 0.000100801 | ENSG00000159871 | LYPD5 | 284348 |
| ENSG00000119508.18 | 16.84507375 | 2.412844976 | 0.586592002 | 3.09047 | 0.001998 | 0.013782748 | ENSG00000119508 | NR4A3 | 8013 |
| ENSG00000269927.1 | 34.3900424 | 2.411716348 | 0.480806312 | 3.768079 | 0.000165 | 0.001427491 | ENSG00000269927 | NA | NA |
| ENSG00000215861.6 | 11.92774384 | 2.411051356 | 0.42896769 | 4.221883 | 2.42E-05 | 0.000240392 | ENSG00000215861 | NA | NA |
| ENSG00000154258.17 | 10.31541368 | 2.406804466 | 0.582405139 | 3.102315 | 0.00192 | 0.013294606 | ENSG00000154258 | ABCA9 | 10350 |
| ENSG00000042832.12 | 4.988925994 | 2.405409785 | 0.673330205 | 2.681314 | 0.007333 | 0.043822408 | ENSG00000042832 | TG | 7038 |
| ENSG00000131069.20 | 2485.96 | 2.404655 | 0.07707699 | 23.41367 | 3.10E-121 | 1.65E-118 | ENSG00000131069 | ACSS2 | 55902 |
| ENSG00000197614.11 | 112.8236702 | 2.404557921 | 0.205748916 | 8.77068 | 1.78E-18 | 6.17E-17 | ENSG00000197614 | MFAP5 | 8076 |
| ENSG00000140254.12 | 470.0689612 | 2.402575693 | 0.246392954 | 7.315857 | 2.56E-13 | 6.14E-12 | ENSG00000140254 | DUOXA1 | 90527 |
| ENSG00000167972.14 | 81.07930043 | 2.402232467 | 0.375642602 | 4.797732 | 1.60E-06 | 1.88E-05 | ENSG00000167972 | ABCA3 | 21 |
| ENSG00000150471.16 | 68.03978359 | 2.400701541 | 0.233347348 | 7.716829 | 1.19E-14 | 3.17E-13 | ENSG00000150471 | ADGRL3 | 23284 |
| ENSG00000132000.13 | 52.50667502 | 2.400671026 | 0.324507777 | 5.54893 | 2.87E-08 | 4.23E-07 | ENSG00000132000 | PODNL1 | 79883 |
| ENSG00000260278.1 | 12.10051358 | 2.400027702 | 0.677697738 | 2.656092 | 0.007905 | 0.046756766 | ENSG00000260278 | NA | NA |
| ENSG00000184792.16 | 574.4577 | 2.399529 | 0.2115401 | 8.5068 | 1.79E-17 | 5.84E-16 | ENSG00000184792 | OSBP2 | 23762 |
| ENSG00000006016.11 | 61.02501 | 2.39437 | 0.2492107 | 7.200212 | 6.01E-13 | 1.41E-11 | ENSG00000006016 | CRLF1 | 9244 |
| ENSG00000248196.1 | 7.190143821 | 2.393867787 | 0.625383245 | 2.86843 | 0.004125 | 0.026411677 | ENSG00000248196 | LOC10041 | 1E+08 |
| ENSG00000253368.4 | 1217.586138 | 2.393474062 | 0.218660419 | 8.202097 | 2.36E-16 | 7.07E-15 | ENSG00000253368 | TRNP1 | 388610 |
| ENSG00000268601.1 | 38.98053 | 2.391481 | 0.3053303 | 5.867355 | 4.43E-09 | 7.09E-08 | ENSG00000268601 | NA | NA |
| ENSG00000174460.4 | 5.985645 | 2.389537 | 0.6376628 | 2.8064 | 0.00501 | 0.03133003 | ENSG00000174460 | ZCCHC12 | 170261 |
| ENSG00000145879.11 | 72.53967463 | 2.389522381 | 0.3286436 | 5.445176 | 5.18E-08 | 7.31E-07 | ENSG00000145879 | SPINK7 | 84651 |
| ENSG00000259744.1 | 11.86855045 | 2.38903107 | 0.478077675 | 3.742135 | 0.000182 | 0.001567962 | ENSG00000259744 | NA | NA |
| ENSG00000188833.9 | 5.681546974 | 2.388430186 | 0.674441096 | 2.651722 | 0.008008 | 0.047250796 | ENSG00000188833 | ENTPD8 | 377841 |
| ENSG00000227533.6 | 23.64139792 | 2.38784983 | 0.323501303 | 5.526561 | 3.27E-08 | 4.76E-07 | ENSG00000227533 | SLC2A1-A | 440584 |
| ENSG00000171451.14 | 62.77376627 | 2.385901677 | 0.4044986 | 4.4151 | 1.01E-05 | 0.000106788 | ENSG00000171451 | DSEL | 92126 |

| | | | | | | | | | |
|---|---|---|---|---|---|---|---|---|---|
| ENSG00000230387.4 | 9.983121 | 2.384361 | 0.6162265 | 2.895625 | 0.003784 | 0.02439197 | ENSG00000230387 | LOC10050 | 1.01E+08 |
| ENSG00000162545.6 | 6535.537807 | 2.381104827 | 0.208534449 | 8.541058 | 1.33E-17 | 4.37E-16 | ENSG00000162545 | CAMK2N1 | 55450 |
| ENSG00000116717.13 | 6679.181996 | 2.380374779 | 0.096091014 | 18.528 | 1.23E-76 | 3.48E-74 | ENSG00000116717 | GADD45A | 1647 |
| ENSG00000130584.11 | 121.4759 | 2.380109 | 0.2088603 | 8.522966 | 1.56E-17 | 5.10E-16 | ENSG00000130584 | ZBTB46 | 140685 |
| ENSG00000160588.10 | 2522.341601 | 2.379265405 | 0.132151501 | 13.46383 | 2.55E-41 | 2.69E-39 | ENSG00000160588 | MPZL3 | 196264 |
| ENSG00000197956.10 | 78706.08242 | 2.379108094 | 0.16229755 | 10.96201 | 5.82E-28 | 3.50E-26 | ENSG00000197956 | S100A6 | 6277 |
| ENSG00000287270.1 | 7.625089901 | 2.375516875 | 0.640920101 | 2.770262 | 0.005601 | 0.034556864 | ENSG00000287270 | NA | NA |
| ENSG00000002079.14 | 290.5715553 | 2.374322719 | 0.195730799 | 9.065118 | 1.24E-19 | 4.65E-18 | ENSG00000002079 | NA | NA |
| ENSG00000102243.13 | 36.37926 | 2.373648 | 0.4129433 | 4.295136 | 1.75E-05 | 0.000177263 | ENSG00000102243 | VGLL1 | 51442 |
| ENSG00000105245.9 | 792.8628 | 2.372019 | 0.2025107 | 8.750251 | 2.13E-18 | 7.35E-17 | ENSG00000105245 | NUMBL | 9253 |
| ENSG00000250381.1 | 12.64264981 | 2.371942334 | 0.460564789 | 3.847325 | 0.000119 | 0.001063585 | ENSG00000250381 | NA | NA |
| ENSG00000167123.19 | 949.3448573 | 2.370565771 | 0.117257172 | 15.09985 | 1.62E-51 | 2.37E-49 | ENSG00000167123 | CERCAM | 51148 |
| ENSG00000143507.18 | 937.0770671 | 2.370270314 | 0.092467943 | 19.14469 | 1.07E-81 | 3.24E-79 | ENSG00000143507 | DUSP10 | 11221 |
| ENSG00000234719.8 | 27.23151613 | 2.369470289 | 0.379253217 | 4.66567 | 3.08E-06 | 3.47E-05 | ENSG00000234719 | NPIPB2 | 729978 |
| ENSG00000287566.1 | 8.556939455 | 2.36890971 | 0.580206746 | 3.048758 | 0.002298 | 0.015653697 | ENSG00000287566 | LOC10537 | 1.05E+08 |
| ENSG00000249992.1 | 101.4042152 | 2.367670916 | 0.217416961 | 8.130327 | 4.28E-16 | 1.26E-14 | ENSG00000249992 | TMEM158 | 25907 |
| ENSG00000163354.15 | 10.27659673 | 2.366332431 | 0.52117183 | 3.389156 | 0.000701 | 0.005389607 | ENSG00000163354 | DCST2 | 127579 |
| ENSG00000133639.5 | 2532.282036 | 2.365046447 | 0.130581594 | 13.51681 | 1.24E-41 | 1.32E-39 | ENSG00000133639 | BTG1 | 694 |
| ENSG00000285737.1 | 20.81009867 | 2.364881193 | 0.363973258 | 4.848931 | 1.24E-06 | 1.48E-05 | ENSG00000285737 | NA | NA |
| ENSG00000130522.5 | 5397.16 | 2.364466 | 0.1606612 | 10.98253 | 4.64E-28 | 2.83E-26 | ENSG00000130522 | JUND | 3727 |
| ENSG00000035862.12 | 499.629732 | 2.363602862 | 0.26820365 | 6.575611 | 4.85E-11 | 9.49E-10 | ENSG00000035862 | TIMP2 | 7077 |
| ENSG00000235531.10 | 24.11930998 | 2.356594185 | 0.362861778 | 4.840946 | 1.29E-06 | 1.53E-05 | ENSG00000235531 | MSC-AS1 | 1E+08 |
| ENSG00000287910.1 | 8.763145733 | 2.356421214 | 0.549134535 | 3.198526 | 0.001381 | 0.009903984 | ENSG00000287910 | NA | NA |
| ENSG00000091129.20 | 522.7271024 | 2.356320143 | 0.37170285 | 4.725065 | 2.30E-06 | 2.64E-05 | ENSG00000091129 | NRCAM | 4897 |
| ENSG00000197279.4 | 1401.23791 | 2.355484129 | 0.345237357 | 5.084861 | 3.68E-07 | 4.70E-06 | ENSG00000197279 | ZNF165 | 7718 |
| ENSG00000111913.19 | 43.19713924 | 2.354509277 | 0.222778019 | 7.875594 | 3.39E-15 | 9.45E-14 | ENSG00000111913 | RIPOR2 | 9750 |
| ENSG00000117122.13 | 345.0433539 | 2.35398178 | 0.164962272 | 10.63262 | 2.10E-26 | 1.14E-24 | ENSG00000117122 | MFAP2 | 4237 |
| ENSG00000165171.11 | 35.95081804 | 2.352922468 | 0.29074136 | 6.029147 | 1.65E-09 | 2.77E-08 | ENSG00000165171 | METTL27 | 155368 |
| ENSG00000170537.13 | 426.8340367 | 2.352558097 | 0.170522441 | 10.27758 | 8.89E-25 | 4.43E-23 | ENSG00000170537 | TMC7 | 79905 |
| ENSG00000273199.1 | 16.74231 | 2.352481 | 0.43596 | 4.019822 | 5.82E-05 | 0.000545192 | ENSG00000273199 | NA | NA |
| ENSG00000169116.11 | 43.32941527 | 2.351308421 | 0.387327058 | 4.521524 | 6.14E-06 | 6.71E-05 | ENSG00000169116 | PARM1 | 25849 |
| ENSG00000121797.10 | 13.50412291 | 2.344339708 | 0.493781602 | 3.532614 | 0.000411 | 0.003311173 | ENSG00000121797 | CCRL2 | 9034 |
| ENSG00000130340.15 | 11386.33035 | 2.34297806 | 0.214848336 | 8.112597 | 4.95E-16 | 1.45E-14 | ENSG00000130340 | SNX9 | 51429 |
| ENSG00000230863.3 | 68.17341344 | 2.341507465 | 0.195825812 | 8.893146 | 5.94E-19 | 2.13E-17 | ENSG00000230863 | NA | NA |
| ENSG00000233818.1 | 15.11724 | 2.339041 | 0.4178818 | 4.161561 | 3.16E-05 | 0.000308908 | ENSG00000233818 | NA | NA |
| ENSG00000140057.9 | 74.45751444 | 2.338757883 | 0.22040587 | 7.888891 | 3.05E-15 | 8.54E-14 | ENSG00000140057 | AK7 | 122481 |
| ENSG00000154262.13 | 32.97269428 | 2.338086017 | 0.35129566 | 4.947644 | 7.51E-07 | 9.25E-06 | ENSG00000154262 | ABCA6 | 23460 |
| ENSG00000272123.1 | 60.31195264 | 2.335082394 | 0.275275648 | 6.303073 | 2.92E-10 | 5.38E-09 | ENSG00000272123 | NA | NA |
| ENSG00000066923.17 | 47.0268793 | 2.334890573 | 0.241199896 | 7.19275 | 6.35E-13 | 1.48E-11 | ENSG00000066923 | STAG3 | 10734 |
| ENSG00000251411.1 | 32.62863823 | 2.334579956 | 0.338380116 | 5.126128 | 2.96E-07 | 3.83E-06 | ENSG00000251411 | NA | NA |
| ENSG00000048052.21 | 1777.444769 | 2.333617618 | 0.152004581 | 11.40504 | 3.95E-30 | 2.71E-28 | ENSG00000048052 | HDAC9 | 9734 |
| ENSG00000135604.9 | 18.96473267 | 2.331304416 | 0.44885013 | 3.857199 | 0.000115 | 0.001025775 | ENSG00000135604 | STX11 | 8676 |

| | | | | | | | | | |
|---|---|---|---|---|---|---|---|---|---|
| ENSG00000175294.5 | 41.48311174 | 2.330210592 | 0.385077314 | 4.493151 | 7.02E-06 | 7.63E-05 | ENSG00000175294 | CATSPER | 117144 |
| ENSG00000240764.3 | 23.01351492 | 2.32941847 | 0.434029971 | 3.98456 | 6.76E-05 | 0.000627104 | ENSG00000240764 | PCDHGC5 | 56097 |
| ENSG00000146147.15 | 7.697695621 | 2.326855372 | 0.593683838 | 2.908712 | 0.003629 | 0.023530638 | ENSG00000146147 | MLIP | 90523 |
| ENSG00000091536.19 | 14.53421885 | 2.326156155 | 0.401340384 | 4.300978 | 1.70E-05 | 0.000172957 | ENSG00000091536 | MYO15A | 51168 |
| ENSG00000180667.10 | 8075.102918 | 2.325967712 | 0.189697698 | 9.098517 | 9.16E-20 | 3.45E-18 | ENSG00000180667 | YOD1 | 55432 |
| ENSG00000251467.1 | 10.71746804 | 2.322872393 | 0.444662112 | 3.874565 | 0.000107 | 0.000960039 | ENSG00000251467 | NA | NA |
| ENSG00000118689.15 | 5921.188319 | 2.320426649 | 0.131151614 | 13.11785 | 2.60E-39 | 2.55E-37 | ENSG00000118689 | FOXO3 | 2309 |
| ENSG00000227676.4 | 11.9202531 | 2.319531874 | 0.476508852 | 3.608604 | 0.000308 | 0.002534968 | ENSG00000227676 | LINC0106 | 1.04E+08 |
| ENSG00000232053.7 | 6.710585104 | 2.318872928 | 0.57544989 | 2.987007 | 0.002817 | 0.01881122 | ENSG00000232053 | NA | NA |
| ENSG00000274290.2 | 11.86835655 | 2.316350736 | 0.445802261 | 3.850027 | 0.000118 | 0.001052623 | ENSG00000274290 | H2BC6 | 8344 |
| ENSG00000237264.4 | 8.847412404 | 2.31489219 | 0.646041883 | 2.65446 | 0.007944 | 0.046913547 | ENSG00000237264 | NA | NA |
| ENSG00000197632.9 | 13551.69415 | 2.314798687 | 0.250100346 | 6.856443 | 7.06E-12 | 1.51E-10 | ENSG00000197632 | SERPINB2 | 5055 |
| ENSG00000237772.2 | 6.168654893 | 2.314088546 | 0.587772417 | 2.916245 | 0.003543 | 0.023009088 | ENSG00000237772 | NA | NA |
| ENSG00000278959.1 | 6.073954476 | 2.310950083 | 0.578024398 | 2.959996 | 0.003076 | 0.020337881 | ENSG00000278959 | NA | NA |
| ENSG00000065361.15 | 3292.84711 | 2.309803201 | 0.159756431 | 10.70256 | 9.90E-27 | 5.48E-25 | ENSG00000065361 | ERBB3 | 2065 |
| ENSG00000177427.13 | 234.0882453 | 2.309710173 | 0.292933383 | 5.836515 | 5.33E-09 | 8.48E-08 | ENSG00000177427 | MIEF2 | 125170 |
| ENSG00000132185.16 | 14.81317947 | 2.309523087 | 0.414870475 | 4.120619 | 3.78E-05 | 0.000365792 | ENSG00000132185 | FCRLA | 84824 |
| ENSG00000215018.9 | 169.0366651 | 2.307600014 | 0.174031056 | 9.812042 | 9.99E-23 | 4.45E-21 | ENSG00000215018 | COL28A1 | 340267 |
| ENSG00000134324.11 | 5223.996378 | 2.306271576 | 0.185456275 | 9.200398 | 3.57E-20 | 1.38E-18 | ENSG00000134324 | LPIN1 | 23175 |
| ENSG00000256351.1 | 10.27018183 | 2.305486729 | 0.620705273 | 2.747659 | 0.006002 | 0.036674637 | ENSG00000256351 | NA | NA |
| ENSG00000059377.17 | 13.77052611 | 2.30380204 | 0.562520105 | 3.028873 | 0.002455 | 0.016636563 | ENSG00000059377 | TBXAS1 | 6916 |
| ENSG00000185567.7 | 5872.688707 | 2.303714359 | 0.207546211 | 8.208843 | 2.23E-16 | 6.71E-15 | ENSG00000185567 | AHNAK2 | 113146 |
| ENSG00000167768.4 | 9.096641296 | 2.302814944 | 0.481008792 | 3.540091 | 0.0004 | 0.003225842 | ENSG00000167768 | KRT1 | 3848 |
| ENSG00000255008.3 | 12.32143213 | 2.301546708 | 0.454385491 | 3.744721 | 0.000181 | 0.001553406 | ENSG00000255008 | NA | NA |
| ENSG00000140545.15 | 1179.976298 | 2.300805245 | 0.147027981 | 11.5679 | 5.99E-31 | 4.27E-29 | ENSG00000140545 | MFGE8 | 4240 |
| ENSG00000134107.4 | 7147.204556 | 2.30034446 | 0.1007174 | 16.88233 | 6.07E-64 | 1.27E-61 | ENSG00000134107 | BHLHE40 | 8553 |
| ENSG00000106772.18 | 11.23490216 | 2.300302225 | 0.507303398 | 3.351648 | 0.000803 | 0.006096362 | ENSG00000106772 | PRUNE2 | 158471 |
| ENSG00000253553.7 | 26.30322341 | 2.29606501 | 0.286870491 | 5.912302 | 3.37E-09 | 5.49E-08 | ENSG00000253553 | NA | NA |
| ENSG00000232006.9 | 51.16493058 | 2.296033166 | 0.265811544 | 6.380585 | 1.76E-10 | 3.30E-09 | ENSG00000232006 | NA | NA |
| ENSG00000280047.1 | 118.244114 | 2.29593516 | 0.181592086 | 9.339257 | 9.70E-21 | 3.91E-19 | ENSG00000280047 | NA | NA |
| ENSG00000183696.14 | 15113.68414 | 2.295182687 | 0.242665168 | 6.985686 | 2.83E-12 | 6.23E-11 | ENSG00000183696 | UPP1 | 7378 |
| ENSG00000166173.10 | 1219.548975 | 2.295123976 | 0.180190904 | 9.407378 | 5.09E-21 | 2.08E-19 | ENSG00000166173 | LARP6 | 55323 |
| ENSG00000109107.14 | 233.0322183 | 2.294583581 | 0.15693944 | 10.79769 | 3.53E-27 | 2.02E-25 | ENSG00000109107 | ALDOC | 230 |
| ENSG00000125744.12 | 396.3374 | 2.294581 | 0.1439297 | 11.77367 | 5.34E-32 | 4.00E-30 | ENSG00000125744 | RTN2 | 6253 |
| ENSG00000231721.7 | 219.0200458 | 2.294157076 | 0.157740933 | 10.74012 | 6.60E-27 | 3.68E-25 | ENSG00000231721 | LINC-PINT | 378805 |
| ENSG00000258393.1 | 37.64243394 | 2.288931346 | 0.301022532 | 5.610648 | 2.02E-08 | 3.02E-07 | ENSG00000258393 | NA | NA |
| ENSG00000102271.14 | 47.82612 | 2.287576 | 0.3661298 | 4.609229 | 4.04E-06 | 4.50E-05 | ENSG00000102271 | KLHL4 | 56062 |
| ENSG00000106211.9 | 7398.395125 | 2.284295053 | 0.145083427 | 11.60915 | 3.70E-31 | 2.69E-29 | ENSG00000106211 | HSPB1 | 3315 |
| ENSG00000228952.1 | 18.00250465 | 2.281418709 | 0.41884473 | 4.01442 | 5.96E-05 | 0.000556838 | ENSG00000228952 | NA | NA |
| ENSG00000237989.1 | 44.74528 | 2.276703 | 0.2526634 | 6.636112 | 3.22E-11 | 6.42E-10 | ENSG00000237989 | LINC0167 | 1.02E+08 |
| ENSG00000184481.16 | 464.2811 | 2.276053 | 0.145192 | 11.5437 | 7.94E-31 | 5.62E-29 | ENSG00000184481 | FOXO4 | 4303 |
| ENSG00000141579.7 | 1083.834524 | 2.273035619 | 0.442040285 | 3.784804 | 0.000154 | 0.001342269 | ENSG00000141579 | ZNF750 | 79755 |

| | | | | | | | | | |
|---|---|---|---|---|---|---|---|---|---|
| ENSG00000236345.1 | 169.6369038 | 2.272655415 | 0.178397882 | 9.375982 | 6.85E-21 | 2.78E-19 | ENSG00000236345 | SCAT8 | 1.13E+08 |
| ENSG00000183023.18 | 19.4146825 | 2.271851028 | 0.342916064 | 4.875394 | 1.09E-06 | 1.31E-05 | ENSG00000183023 | SLC8A1 | 6546 |
| ENSG00000173846.13 | 3076.332604 | 2.271774444 | 0.181330969 | 9.219465 | 2.99E-20 | 1.16E-18 | ENSG00000173846 | PLK3 | 1263 |
| ENSG00000286733.1 | 19.62945572 | 2.270490307 | 0.422547013 | 3.953383 | 7.71E-05 | 0.000709066 | ENSG00000286733 | LINC0234ε | 1.02E+08 |
| ENSG00000243225.2 | 51.74557522 | 2.26962758 | 0.236782681 | 7.051308 | 1.77E-12 | 3.97E-11 | ENSG00000243225 | NA | NA |
| ENSG00000169908.12 | 3307.647828 | 2.269186955 | 0.17478941 | 9.549703 | 1.30E-21 | 5.47E-20 | ENSG00000169908 | TM4SF1 | 4071 |
| ENSG00000105499.14 | 47.08833 | 2.268203 | 0.2129673 | 7.833142 | 4.76E-15 | 1.31E-13 | ENSG00000105499 | PLA2G4C | 8605 |
| ENSG00000122367.19 | 24.42993096 | 2.266403733 | 0.356306208 | 4.676887 | 2.91E-06 | 3.30E-05 | ENSG00000122367 | LDB3 | 11155 |
| ENSG00000073150.13 | 100.1289 | 2.265936 | 0.3230935 | 5.156205 | 2.52E-07 | 3.30E-06 | ENSG00000073150 | PANX2 | 56666 |
| ENSG00000107968.10 | 674.6651332 | 2.26572198 | 0.235628867 | 7.069261 | 1.56E-12 | 3.50E-11 | ENSG00000107968 | MAP3K8 | 1326 |
| ENSG00000277449.1 | 25.18846 | 2.265588 | 0.3335362 | 4.993726 | 5.92E-07 | 7.38E-06 | ENSG00000277449 | CEBPB-AS | 1.02E+08 |
| ENSG00000164938.14 | 845.915281 | 2.264465195 | 0.154232651 | 10.79191 | 3.76E-27 | 2.14E-25 | ENSG00000164938 | TP53INP1 | 94241 |
| ENSG00000263627.1 | 6.286438269 | 2.264401668 | 0.585417855 | 2.8431 | 0.004468 | 0.028352596 | ENSG00000263627 | PPP4R1-A | 1.02E+08 |
| ENSG00000141837.20 | 10.29418416 | 2.262432368 | 0.484559606 | 3.430811 | 0.000602 | 0.004687123 | ENSG00000141837 | CACNA1A | 773 |
| ENSG00000188277.9 | 199.1675388 | 2.259486867 | 0.21586597 | 7.68758 | 1.50E-14 | 3.95E-13 | ENSG00000188277 | C15orf62 | 643338 |
| ENSG00000125531.7 | 53.06685 | 2.257675 | 0.3231844 | 5.129194 | 2.91E-07 | 3.77E-06 | ENSG00000125531 | FNDC11 | 79025 |
| ENSG00000262903.1 | 19.19574384 | 2.256195493 | 0.40663365 | 4.072943 | 4.64E-05 | 0.000442172 | ENSG00000262903 | NA | NA |
| ENSG00000167996.16 | 10920.54777 | 2.248321284 | 0.139596973 | 11.80772 | 3.56E-32 | 2.69E-30 | ENSG00000167996 | FTH1 | 2495 |
| ENSG00000128590.5 | 1208.152577 | 2.248237235 | 0.207303099 | 7.950857 | 1.85E-15 | 5.26E-14 | ENSG00000128590 | DNAJB9 | 4189 |
| ENSG00000156453.13 | 1994.81916 | 2.247008003 | 0.216950119 | 7.591644 | 3.16E-14 | 8.12E-13 | ENSG00000156453 | PCDH1 | 5097 |
| ENSG00000011198.9 | 4431.061101 | 2.24540674 | 0.24260124 | 6.782351 | 1.18E-11 | 2.46E-10 | ENSG00000011198 | ABHD5 | 51099 |
| ENSG00000183508.5 | 40.5376582 | 2.241976992 | 0.266838657 | 6.153445 | 7.58E-10 | 1.32E-08 | ENSG00000183508 | TENT5C | 54855 |
| ENSG00000143878.9 | 5296.261451 | 2.241503405 | 0.160714663 | 10.21378 | 1.72E-24 | 8.39E-23 | ENSG00000143878 | RHOB | 388 |
| ENSG00000163191.6 | 35410.2603 | 2.237064115 | 0.152141723 | 10.76013 | 5.31E-27 | 2.99E-25 | ENSG00000163191 | S100A11 | 6282 |
| ENSG00000222009.8 | 363.628915 | 2.236560826 | 0.128621672 | 12.72383 | 4.36E-37 | 3.92E-35 | ENSG00000222009 | BTBD19 | 149478 |
| ENSG00000250920.3 | 2266.556229 | 2.235635174 | 0.133229997 | 12.27678 | 1.21E-34 | 9.77E-33 | ENSG00000250920 | NA | NA |
| ENSG00000285106.2 | 338.1136567 | 2.233496113 | 0.182736867 | 8.939062 | 3.92E-19 | 1.42E-17 | ENSG00000285106 | NA | NA |
| ENSG00000215009.5 | 16.14503147 | 2.233363713 | 0.516205153 | 3.164176 | 0.001555 | 0.010993979 | ENSG00000215009 | ACSM4 | 341392 |
| ENSG00000248794.1 | 36.64176978 | 2.230846603 | 0.31379922 | 5.197102 | 2.02E-07 | 2.68E-06 | ENSG00000248794 | NA | NA |
| ENSG00000118292.9 | 27.19416651 | 2.229716525 | 0.28685683 | 5.681289 | 1.34E-08 | 2.03E-07 | ENSG00000118292 | C1orf54 | 79630 |
| ENSG00000184731.6 | 5235.235633 | 2.228107299 | 0.337099011 | 4.82976 | 1.37E-06 | 1.62E-05 | ENSG00000184731 | FAM110C | 642273 |
| ENSG00000004399.12 | 220.8952127 | 2.22628433 | 0.166443554 | 9.770786 | 1.50E-22 | 6.57E-21 | ENSG00000004399 | PLXND1 | 23129 |
| ENSG00000148180.19 | 3341.773085 | 2.223034101 | 0.17847834 | 9.093731 | 9.57E-20 | 3.60E-18 | ENSG00000148180 | GSN | 2934 |
| ENSG00000057149.16 | 1814.012526 | 2.221019012 | 0.384035989 | 4.221008 | 2.43E-05 | 0.000241237 | ENSG00000057149 | SERPINB3 | 6317 |
| ENSG00000169306.9 | 6.296148 | 2.219399 | 0.5362543 | 3.019834 | 0.002529 | 0.01709331 | ENSG00000169306 | IL1RAPL1 | 11141 |
| ENSG00000227376.1 | 8.148040266 | 2.217992289 | 0.537878407 | 3.0081 | 0.002629 | 0.017704419 | ENSG00000227376 | NA | NA |
| ENSG00000172183.15 | 518.7882706 | 2.215781199 | 0.293799196 | 5.499611 | 3.81E-08 | 5.47E-07 | ENSG00000172183 | ISG20 | 3669 |
| ENSG00000215160.3 | 8.513642464 | 2.213265643 | 0.508990952 | 3.169537 | 0.001527 | 0.010830538 | ENSG00000215160 | NA | NA |
| ENSG00000246560.2 | 64.72703995 | 2.212567583 | 0.220006193 | 7.329646 | 2.31E-13 | 5.59E-12 | ENSG00000246560 | UBE2D3-A | 1.05E+08 |
| ENSG00000121742.19 | 196.2667183 | 2.211893724 | 0.201936007 | 7.982201 | 1.44E-15 | 4.13E-14 | ENSG00000121742 | GJB6 | 10804 |
| ENSG00000268223.5 | 104.3701087 | 2.208153761 | 0.249579767 | 6.443446 | 1.17E-10 | 2.21E-09 | ENSG00000268223 | ARL14EPL | 644100 |
| ENSG00000226747.6 | 13.20827614 | 2.208016131 | 0.438569721 | 3.666501 | 0.000246 | 0.002064379 | ENSG00000226747 | FSIP2-AS2 | 1.02E+08 |

| | | | | | | | | | |
|---|---|---|---|---|---|---|---|---|---|
| ENSG00000127920.5 | 253.4027841 | 2.207619827 | 0.098593576 | 16.30552 | 9.02E-60 | 1.73E-57 | ENSG00000127920 | GNG11 | 2791 |
| ENSG00000135926.15 | 18165.3494 | 2.206972604 | 0.105661734 | 15.20865 | 3.10E-52 | 4.66E-50 | ENSG00000135926 | TMBIM1 | 64114 |
| ENSG00000232692.1 | 12.98141 | 2.205309 | 0.4651948 | 3.450831 | 0.000559 | 0.004381052 | ENSG00000232692 | NA | NA |
| ENSG00000171621.14 | 1859.329373 | 2.20435555 | 0.146269987 | 10.96845 | 5.42E-28 | 3.28E-26 | ENSG00000171621 | SPSB1 | 80176 |
| ENSG00000260630.7 | 66.75723982 | 2.202922788 | 0.196282036 | 8.166426 | 3.18E-16 | 9.43E-15 | ENSG00000260630 | SNAI3-AS1 | 197187 |
| ENSG00000258102.4 | 50.86758889 | 2.202232423 | 0.300715029 | 5.328076 | 9.93E-08 | 1.37E-06 | ENSG00000258102 | MAP1LC3I | 643246 |
| ENSG00000010404.18 | 495.9931 | 2.202085 | 0.1265979 | 12.65491 | 1.05E-36 | 9.30E-35 | ENSG00000010404 | IDS | 3423 |
| ENSG00000101096.20 | 12.87774 | 2.200125 | 0.6014883 | 2.660277 | 0.007808 | 0.04625168 | ENSG00000101096 | NFATC2 | 4773 |
| ENSG00000057704.13 | 3282.240855 | 2.199981444 | 0.207031111 | 7.728217 | 1.09E-14 | 2.91E-13 | ENSG00000057704 | TMCC3 | 57458 |
| ENSG00000124766.6 | 1053.575896 | 2.199693221 | 0.114956559 | 13.91563 | 5.09E-44 | 5.97E-42 | ENSG00000124766 | SOX4 | 6659 |
| ENSG00000134242.16 | 8.489268883 | 2.198836267 | 0.513621734 | 3.112867 | 0.001853 | 0.012892072 | ENSG00000134242 | PTPN22 | 26191 |
| ENSG00000181444.13 | 53.41891742 | 2.198139679 | 0.35601205 | 4.489004 | 7.16E-06 | 7.77E-05 | ENSG00000181444 | ZNF467 | 168544 |
| ENSG00000117114.19 | 447.7697846 | 2.194336717 | 0.193169766 | 8.253552 | 1.54E-16 | 4.68E-15 | ENSG00000117114 | ADGRL2 | 23266 |
| ENSG00000100284.21 | 1316.513 | 2.18903 | 0.2157339 | 7.365695 | 1.76E-13 | 4.31E-12 | ENSG00000100284 | TOM1 | 10043 |
| ENSG00000171860.4 | 26.56560226 | 2.186317843 | 0.376979789 | 4.207965 | 2.58E-05 | 0.000254828 | ENSG00000171860 | C3AR1 | 719 |
| ENSG00000089199.10 | 75.66611 | 2.186223 | 0.2634634 | 6.020658 | 1.74E-09 | 2.90E-08 | ENSG00000089199 | CHGB | 1114 |
| ENSG00000137872.16 | 10.80003708 | 2.18353111 | 0.539714817 | 2.934015 | 0.003346 | 0.021892231 | ENSG00000137872 | SEMA6D | 80031 |
| ENSG00000254560.6 | 107.0645728 | 2.182622947 | 0.200514952 | 7.892793 | 2.95E-15 | 8.30E-14 | ENSG00000254560 | BBOX1-AS | 1.04E+08 |
| ENSG00000151303.11 | 13.54491343 | 2.18146029 | 0.503867256 | 3.138645 | 0.001697 | 0.01191295 | ENSG00000151303 | NA | NA |
| ENSG00000252711.1 | 12.85350533 | 2.180851976 | 0.458615018 | 3.447013 | 0.000567 | 0.004432994 | ENSG00000252711 | RN7SKP1 | 1.06E+08 |
| ENSG00000134757.5 | 13700.51576 | 2.180616372 | 0.20349548 | 7.767329 | 8.02E-15 | 2.16E-13 | ENSG00000134757 | DSG3 | 1830 |
| ENSG00000179088.14 | 7.537211718 | 2.179332689 | 0.481950209 | 3.276962 | 0.001049 | 0.007742395 | ENSG00000179088 | C12orf42 | 374470 |
| ENSG00000198915.11 | 32.95533109 | 2.178029795 | 0.280792034 | 5.619924 | 1.91E-08 | 2.87E-07 | ENSG00000198915 | RASGEF1 | 221002 |
| ENSG00000185640.5 | 20.72957661 | 2.177842942 | 0.376831621 | 4.18713 | 2.83E-05 | 0.000277624 | ENSG00000185640 | KRT79 | 338785 |
| ENSG00000146278.10 | 1627.769577 | 2.176187252 | 0.120244305 | 13.10821 | 2.96E-39 | 2.88E-37 | ENSG00000146278 | PNRC1 | 10957 |
| ENSG00000229334.1 | 94.42502243 | 2.171410152 | 0.201670454 | 7.79197 | 6.60E-15 | 1.79E-13 | ENSG00000229334 | NA | NA |
| ENSG00000124635.8 | 55.14788387 | 2.170371936 | 0.476197542 | 3.297732 | 0.000975 | 0.007244058 | ENSG00000124635 | H2BC11 | 8970 |
| ENSG00000174718.12 | 8907.473287 | 2.166021288 | 0.204558454 | 7.655618 | 1.92E-14 | 5.04E-13 | ENSG00000174718 | RESF1 | 55196 |
| ENSG00000274070.1 | 98.07849948 | 2.163928755 | 0.164795235 | 9.490133 | 2.31E-21 | 9.52E-20 | ENSG00000274070 | CASTOR2 | 729438 |
| ENSG00000125398.8 | 3079.497186 | 2.163825182 | 0.123118627 | 12.70178 | 5.78E-37 | 5.16E-35 | ENSG00000125398 | SOX9 | 6662 |
| ENSG00000235411.1 | 8.140082938 | 2.161706625 | 0.523408746 | 2.983723 | 0.002848 | 0.018985733 | ENSG00000235411 | NA | NA |
| ENSG00000185345.21 | 19.57120795 | 2.157932265 | 0.479857969 | 3.246653 | 0.001168 | 0.008521573 | ENSG00000185345 | PRKN | 5071 |
| ENSG00000092421.16 | 365.9654737 | 2.154636725 | 0.094684076 | 16.4192 | 1.39E-60 | 2.75E-58 | ENSG00000092421 | SEMA6A | 57556 |
| ENSG00000012124.17 | 31.54456 | 2.15052 | 0.3969457 | 3.906127 | 9.38E-05 | 0.00085157 | ENSG00000012124 | CD22 | 933 |
| ENSG00000185950.9 | 1726.110769 | 2.147318341 | 0.218280748 | 7.088662 | 1.35E-12 | 3.06E-11 | ENSG00000185950 | IRS2 | 8660 |
| ENSG00000225774.2 | 17.79605 | 2.147207 | 0.4095103 | 3.778188 | 0.000158 | 0.001375269 | ENSG00000225774 | NA | NA |
| ENSG00000038945.15 | 12.54198446 | 2.146613297 | 0.57106265 | 2.708308 | 0.006763 | 0.040797064 | ENSG00000038945 | MSR1 | 4481 |
| ENSG00000069482.7 | 1149.838612 | 2.142349523 | 0.086960284 | 17.73625 | 2.20E-70 | 5.38E-68 | ENSG00000069482 | GAL | 51083 |
| ENSG00000163472.19 | 823.8873842 | 2.141930102 | 0.246865963 | 6.246021 | 4.21E-10 | 7.59E-09 | ENSG00000163472 | TMEM79 | 84283 |
| ENSG00000162645.13 | 753.6672412 | 2.140562053 | 0.103111626 | 14.94072 | 1.79E-50 | 2.56E-48 | ENSG00000162645 | GBP2 | 2634 |
| ENSG00000256915.2 | 41.14701916 | 2.140345843 | 0.249087656 | 6.183951 | 6.25E-10 | 1.10E-08 | ENSG00000256915 | NA | NA |
| ENSG00000122042.9 | 2122.275001 | 2.139605986 | 0.156078526 | 9.864304 | 5.94E-23 | 2.69E-21 | ENSG00000122042 | UBL3 | 5412 |

| | | | | | | | | | |
|---|---|---|---|---|---|---|---|---|---|
| ENSG00000253771.6 | 9.335800093 | 2.139505249 | 0.511763246 | 3.008237 | 0.002628 | 0.017700931 | ENSG00000253771 | NA | NA |
| ENSG00000133710.15 | 288.021796 | 2.138901172 | 0.233956683 | 6.577718 | 4.78E-11 | 9.37E-10 | ENSG00000133710 | SPINK5 | 11005 |
| ENSG00000167741.10 | 564.3710388 | 2.137875335 | 0.292877883 | 5.25091 | 1.51E-07 | 2.04E-06 | ENSG00000167741 | GGT6 | 124975 |
| ENSG00000092330.18 | 1142.58872 | 2.135757726 | 0.181682901 | 8.452957 | 2.84E-17 | 9.16E-16 | ENSG00000092330 | TINF2 | 26277 |
| ENSG00000165879.8 | 138.7087764 | 2.134542391 | 0.194165229 | 7.903281 | 2.72E-15 | 7.64E-14 | ENSG00000165879 | FRAT1 | 10023 |
| ENSG00000274818.1 | 11.56899317 | 2.133711426 | 0.488011938 | 3.142774 | 0.001674 | 0.011761665 | ENSG00000274818 | NA | NA |
| ENSG00000268621.6 | 588.7946 | 2.131803 | 0.13251 | 11.55991 | 6.58E-31 | 4.67E-29 | ENSG00000268621 | IGFL2-AS1 | 645553 |
| ENSG00000238120.1 | 24.65139 | 2.131467 | 0.417842 | 3.665182 | 0.000247 | 0.002074394 | ENSG00000238120 | LINC0158( | 1.01E+08 |
| ENSG00000007516.13 | 34.40454355 | 2.129606267 | 0.293642587 | 5.209075 | 1.90E-07 | 2.53E-06 | ENSG00000007516 | BAIAP3 | 8938 |
| ENSG00000095752.7 | 398.8876 | 2.129224 | 0.3154378 | 4.847942 | 1.25E-06 | 1.48E-05 | ENSG00000095752 | IL11 | 3589 |
| ENSG00000182511.12 | 23.58017343 | 2.126797018 | 0.297828878 | 5.126424 | 2.95E-07 | 3.82E-06 | ENSG00000182511 | FES | 2242 |
| ENSG00000219507.4 | 11.90188 | 2.126013 | 0.4343546 | 3.513288 | 0.000443 | 0.003540508 | ENSG00000219507 | NA | NA |
| ENSG00000287262.1 | 22.49078675 | 2.125563894 | 0.53616277 | 2.845337 | 0.004436 | 0.028167654 | ENSG00000287262 | NA | NA |
| ENSG00000171124.13 | 442.4180724 | 2.124403445 | 0.219430014 | 6.947105 | 3.73E-12 | 8.12E-11 | ENSG00000171124 | FUT3 | 2525 |
| ENSG00000256417.1 | 31.12443765 | 2.122400125 | 0.342237532 | 4.448373 | 8.65E-06 | 9.26E-05 | ENSG00000256417 | NA | NA |
| ENSG00000237672.1 | 8.08151157 | 2.121134806 | 0.513328327 | 2.963279 | 0.003044 | 0.020147269 | ENSG00000237672 | NA | NA |
| ENSG00000145349.17 | 1597.076904 | 2.120824747 | 0.165801676 | 9.172554 | 4.62E-20 | 1.77E-18 | ENSG00000145349 | CAMK2D | 817 |
| ENSG00000094755.17 | 13.70150715 | 2.119009742 | 0.441516672 | 3.440436 | 0.000581 | 0.004536819 | ENSG00000094755 | GABRP | 2568 |
| ENSG00000285458.1 | 11.72757714 | 2.1188318 | 0.40816665 | 3.721107 | 0.000198 | 0.001691387 | ENSG00000285458 | C4orf36 | 132989 |
| ENSG00000011083.9 | 14.07476328 | 2.117061911 | 0.405508035 | 3.741139 | 0.000183 | 0.001573173 | ENSG00000011083 | SLC6A7 | 6534 |
| ENSG00000163683.12 | 1676.782779 | 2.11684789 | 0.215352649 | 7.043553 | 1.87E-12 | 4.19E-11 | ENSG00000163683 | SMIM14 | 201895 |
| ENSG00000255120.5 | 26.38575611 | 2.115538478 | 0.455511337 | 3.327115 | 0.000878 | 0.006595389 | ENSG00000255120 | OVOL1-AS | 1.02E+08 |
| ENSG00000138606.19 | 50.61212091 | 2.114798286 | 0.306149277 | 4.947907 | 7.50E-07 | 9.25E-06 | ENSG00000138606 | SHF | 90525 |
| ENSG00000145362.18 | 68.3951083 | 2.113181657 | 0.324748354 | 4.659551 | 3.17E-06 | 3.57E-05 | ENSG00000145362 | ANK2 | 287 |
| ENSG00000072163.19 | 19.83309356 | 2.112496637 | 0.457258666 | 3.307748 | 0.00094 | 0.007015352 | ENSG00000072163 | LIMS2 | 55679 |
| ENSG00000147509.14 | 3324.401718 | 2.112157368 | 0.174112516 | 8.684944 | 3.79E-18 | 1.29E-16 | ENSG00000147509 | RGS20 | 8601 |
| ENSG00000197641.11 | 4574.117413 | 2.112129971 | 0.254923126 | 5.93171 | 3.00E-09 | 4.91E-08 | ENSG00000197641 | SERPINB1 | 5275 |
| ENSG00000241749.4 | 1067.077735 | 2.11139968 | 0.096456765 | 15.66919 | 2.46E-55 | 4.11E-53 | ENSG00000241749 | RPSAP52 | 204010 |
| ENSG00000186806.5 | 168.336 | 2.108793 | 0.1936662 | 7.790688 | 6.66E-15 | 1.81E-13 | ENSG00000186806 | VSIG10L | 147645 |
| ENSG00000245532.9 | 22445.54604 | 2.1014412 | 0.140890807 | 10.65677 | 1.62E-26 | 8.86E-25 | ENSG00000245532 | NEAT1 | 283131 |
| ENSG00000103888.17 | 285.7824414 | 2.095649522 | 0.286933458 | 5.212531 | 1.86E-07 | 2.48E-06 | ENSG00000103888 | CEMIP | 57214 |
| ENSG00000175130.7 | 2199.098181 | 2.094895834 | 0.188740936 | 7.920358 | 2.37E-15 | 6.68E-14 | ENSG00000175130 | MARCKSL | 65108 |
| ENSG00000214274.9 | 42.30998527 | 2.093196168 | 0.31802648 | 4.695194 | 2.66E-06 | 3.04E-05 | ENSG00000214274 | ANG | 283 |
| ENSG00000188487.11 | 74.33049324 | 2.092963495 | 0.226523638 | 6.590762 | 4.38E-11 | 8.61E-10 | ENSG00000188487 | INSC | 387755 |
| ENSG00000109472.14 | 42.64642465 | 2.092513261 | 0.286286527 | 5.213355 | 1.85E-07 | 2.47E-06 | ENSG00000109472 | CPE | 1363 |
| ENSG00000166033.13 | 7830.29271 | 2.09082591 | 0.086964776 | 17.14287 | 7.11E-66 | 1.56E-63 | ENSG00000166033 | HTRA1 | 5654 |
| ENSG00000137193.14 | 2026.247443 | 2.089530388 | 0.17563031 | 8.481055 | 2.23E-17 | 7.26E-16 | ENSG00000137193 | PIM1 | 5292 |
| ENSG00000140941.13 | 9743.801952 | 2.089310618 | 0.158768476 | 9.380392 | 6.57E-21 | 2.67E-19 | ENSG00000140941 | MAP1LC3I | 81631 |
| ENSG00000127954.12 | 248.0335985 | 2.086407971 | 0.304051927 | 4.888665 | 1.02E-06 | 1.23E-05 | ENSG00000127954 | STEAP4 | 79689 |
| ENSG00000272690.6 | 7.089222178 | 2.086042236 | 0.50359209 | 2.950885 | 0.003169 | 0.020869823 | ENSG00000272690 | LINC0201{ | 1.08E+08 |
| ENSG00000196352.15 | 2565.087399 | 2.085933843 | 0.240748643 | 6.172138 | 6.74E-10 | 1.18E-08 | ENSG00000196352 | CD55 | 1604 |
| ENSG00000174015.10 | 26.69578812 | 2.084527396 | 0.384874703 | 3.857171 | 0.000115 | 0.001025775 | ENSG00000174015 | CBY2 | 220082 |

| | | | | | | | | | |
|---|---|---|---|---|---|---|---|---|---|
| ENSG00000251577.5 | 12.30070462 | 2.084460621 | 0.483031895 | 3.073214 | 0.002118 | 0.014544995 | ENSG00000251577 | NA | NA |
| ENSG00000170485.17 | 175.6104661 | 2.081047532 | 0.21342027 | 6.939582 | 3.93E-12 | 8.54E-11 | ENSG00000170485 | NPAS2 | 4862 |
| ENSG00000167578.18 | 13.31708 | 2.079433 | 0.4069108 | 3.635766 | 0.000277 | 0.002303602 | ENSG00000167578 | RAB4B | 53916 |
| ENSG00000166311.10 | 493.871889 | 2.078612116 | 0.159924957 | 9.245662 | 2.34E-20 | 9.15E-19 | ENSG00000166311 | SMPD1 | 6609 |
| ENSG00000148516.21 | 297.5709901 | 2.077219506 | 0.139552548 | 10.5854 | 3.48E-26 | 1.88E-24 | ENSG00000148516 | ZEB1 | 6935 |
| ENSG00000230438.7 | 18.62910822 | 2.077116526 | 0.384306827 | 3.843586 | 0.000121 | 0.001077762 | ENSG00000230438 | NA | NA |
| ENSG00000169291.10 | 25.60551287 | 2.074520893 | 0.33330827 | 4.423895 | 9.69E-06 | 0.000102979 | ENSG00000169291 | SHE | 126669 |
| ENSG00000085063.16 | 22815.08384 | 2.073520891 | 0.073900927 | 19.93914 | 1.86E-88 | 6.28E-86 | ENSG00000085063 | CD59 | 966 |
| ENSG00000196154.12 | 567.2123496 | 2.070687176 | 0.108194583 | 13.59298 | 4.41E-42 | 4.75E-40 | ENSG00000196154 | S100A4 | 6275 |
| ENSG00000163689.20 | 1879.104067 | 2.070197873 | 0.127081355 | 11.56895 | 5.92E-31 | 4.23E-29 | ENSG00000163689 | CFAP20D( | 200844 |
| ENSG00000114626.18 | 908.3281885 | 2.069072752 | 0.20624863 | 7.122824 | 1.06E-12 | 2.42E-11 | ENSG00000114626 | ABTB1 | 80325 |
| ENSG00000116991.10 | 202.3770607 | 2.068702249 | 0.242615092 | 6.053631 | 1.42E-09 | 2.39E-08 | ENSG00000116991 | SIPA1L2 | 57568 |
| ENSG00000248787.2 | 11.03564875 | 2.068299793 | 0.445329696 | 3.297107 | 0.000977 | 0.007258167 | ENSG00000248787 | NA | NA |
| ENSG00000176809.10 | 131.7432046 | 2.066659569 | 0.154594424 | 9.487144 | 2.37E-21 | 9.79E-20 | ENSG00000176809 | LRRC37A3 | 374819 |
| ENSG00000211448.12 | 31.29591902 | 2.065842908 | 0.336718472 | 4.353319 | 1.34E-05 | 0.000138957 | ENSG00000211448 | DIO2 | 1734 |
| ENSG00000123095.6 | 160.8620283 | 2.065150057 | 0.157877114 | 9.280319 | 1.69E-20 | 6.69E-19 | ENSG00000123095 | BHLHE41 | 79365 |
| ENSG00000070540.13 | 139.9189355 | 2.064946552 | 0.121377203 | 12.06937 | 1.53E-33 | 1.21E-31 | ENSG00000070540 | WIPI1 | 55062 |
| ENSG00000167508.12 | 1832.62469 | 2.063142568 | 0.070410586 | 20.78015 | 6.55E-96 | 2.64E-93 | ENSG00000167508 | MVD | 4597 |
| ENSG00000145703.16 | 373.1026992 | 2.058725312 | 0.148222687 | 9.841444 | 7.46E-23 | 3.34E-21 | ENSG00000145703 | IQGAP2 | 10788 |
| ENSG00000283959.1 | 15.69418575 | 2.058102929 | 0.370628759 | 3.934133 | 8.35E-05 | 0.000763873 | ENSG00000283959 | NA | NA |
| ENSG00000166922.8 | 78.82124578 | 2.056547042 | 0.194562786 | 7.486257 | 7.09E-14 | 1.78E-12 | ENSG00000166922 | SCG5 | 6447 |
| ENSG00000123933.17 | 1251.665371 | 2.05458644 | 0.11029952 | 13.1876 | 1.03E-39 | 1.03E-37 | ENSG00000123933 | MXD4 | 10608 |
| ENSG00000204396.10 | 29.89904305 | 2.052116906 | 0.332995635 | 4.360769 | 1.30E-05 | 0.000134728 | ENSG00000204396 | VWA7 | 80737 |
| ENSG00000141232.5 | 3507.939495 | 2.050983662 | 0.233361434 | 6.217753 | 5.04E-10 | 9.01E-09 | ENSG00000141232 | TOB1 | 10140 |
| ENSG00000109436.8 | 2106.23967 | 2.049625957 | 0.230013688 | 6.302346 | 2.93E-10 | 5.40E-09 | ENSG00000109436 | TBC1D9 | 23158 |
| ENSG00000072954.7 | 268.8011 | 2.047751 | 0.1089539 | 13.28774 | 2.73E-40 | 2.75E-38 | ENSG00000072954 | TMEM38A | 79041 |
| ENSG00000243742.5 | 9.37778468 | 2.04749207 | 0.510933033 | 2.833037 | 0.004611 | 0.02909432 | ENSG00000243742 | RPLP0P2 | 113157 |
| ENSG00000260549.1 | 308.6887917 | 2.046229788 | 0.136768286 | 10.57431 | 3.92E-26 | 2.11E-24 | ENSG00000260549 | MT1L | 4500 |
| ENSG00000213462.5 | 520.7178633 | 2.045422524 | 0.139152051 | 10.38736 | 2.83E-25 | 1.46E-23 | ENSG00000213462 | ERV3-1 | 2086 |
| ENSG00000160570.14 | 2581.806 | 2.043619 | 0.1964492 | 7.34856 | 2.00E-13 | 4.88E-12 | ENSG00000160570 | DEDD2 | 162989 |
| ENSG00000259692.6 | 29.47022227 | 2.042646535 | 0.309172821 | 4.666149 | 3.07E-06 | 3.46E-05 | ENSG00000259692 | NA | NA |
| ENSG00000145103.15 | 72.88861955 | 2.042305281 | 0.340980714 | 4.229873 | 2.34E-05 | 0.00023253 | ENSG00000145103 | ILDR1 | 286676 |
| ENSG00000158828.8 | 2086.968962 | 2.040148682 | 0.112884523 | 12.75772 | 2.82E-37 | 2.58E-35 | ENSG00000158828 | PINK1 | 65018 |
| ENSG00000260400.1 | 15.7354077 | 2.039553971 | 0.43431815 | 3.314515 | 0.000918 | 0.006865058 | ENSG00000260400 | NA | NA |
| ENSG00000172901.20 | 55.17887186 | 2.037257077 | 0.302161325 | 4.756588 | 1.97E-06 | 2.28E-05 | ENSG00000172901 | LVRN | 206338 |
| ENSG00000165861.14 | 1132.074415 | 2.034732004 | 0.157896307 | 9.086546 | 1.02E-19 | 3.83E-18 | ENSG00000165861 | ZFYVE1 | 53349 |
| ENSG00000247572.8 | 1017.555177 | 2.032707114 | 0.229755089 | 6.235801 | 4.49E-10 | 8.08E-09 | ENSG00000247572 | CKMT2-AS | 1E+08 |
| ENSG00000114279.13 | 50.49320898 | 2.030942368 | 0.247157911 | 5.789588 | 7.06E-09 | 1.11E-07 | ENSG00000114279 | FGF12 | 2257 |
| ENSG00000130701.4 | 21.23012 | 2.030858 | 0.5168147 | 2.76861 | 0.00563 | 0.03469244 | ENSG00000130701 | RBBP8NL | 140893 |
| ENSG00000108846.16 | 1344.802952 | 2.030119651 | 0.122868003 | 11.63948 | 2.60E-31 | 1.90E-29 | ENSG00000108846 | ABCC3 | 8714 |
| ENSG00000239305.7 | 1185.174976 | 2.029972292 | 0.142057807 | 10.06613 | 7.80E-24 | 3.71E-22 | ENSG00000239305 | RNF103 | 7844 |
| ENSG00000173334.4 | 2545.820456 | 2.028874039 | 0.126986192 | 11.2522 | 2.26E-29 | 1.51E-27 | ENSG00000173334 | TRIB1 | 10221 |

| | | | | | | | | | |
|---|---|---|---|---|---|---|---|---|---|
| ENSG00000136158.12 | 1521.529458 | 2.025987578 | 0.095823027 | 14.88147 | 4.35E-50 | 6.19E-48 | ENSG00000136158 | SPRY2 | 10253 |
| ENSG00000227456.9 | 15.92447 | 2.025175 | 0.4822421 | 2.955311 | 0.003124 | 0.02059475 | ENSG00000227456 | LINC0031( | 114036 |
| ENSG00000134215.16 | 46.61395654 | 2.024569901 | 0.23034362 | 6.184542 | 6.23E-10 | 1.10E-08 | ENSG00000134215 | VAV3 | 10451 |
| ENSG00000157502.13 | 27.67505 | 2.02445 | 0.2905065 | 4.903331 | 9.42E-07 | 1.14E-05 | ENSG00000157502 | PWWP3B | 139221 |
| ENSG00000167759.13 | 16.50101 | 2.024223 | 0.4963306 | 2.869505 | 0.004111 | 0.02633466 | ENSG00000167759 | KLK13 | 26085 |
| ENSG00000261207.1 | 16.42643886 | 2.023733862 | 0.406894229 | 3.499027 | 0.000467 | 0.003718634 | ENSG00000261207 | NA | NA |
| ENSG00000235865.2 | 27.82980434 | 2.02324334 | 0.32306944 | 4.405379 | 1.06E-05 | 0.000111249 | ENSG00000235865 | GSN-AS1 | 57000 |
| ENSG00000260581.1 | 47.49765295 | 2.019396749 | 0.241158076 | 5.885752 | 3.96E-09 | 6.39E-08 | ENSG00000260581 | LOC10537 | 1.05E+08 |
| ENSG00000184330.12 | 625.7345327 | 2.018915805 | 0.275843554 | 5.143915 | 2.69E-07 | 3.51E-06 | ENSG00000184330 | S100A7A | 338324 |
| ENSG00000185909.15 | 255.8641963 | 2.013258856 | 0.110913533 | 12.74199 | 3.45E-37 | 3.14E-35 | ENSG00000185909 | KLHDC8B | 200942 |
| ENSG00000107736.20 | 9.625162625 | 2.011056001 | 0.475482743 | 2.967628 | 0.003001 | 0.019899006 | ENSG00000107736 | CDH23 | 64072 |
| ENSG00000250710.3 | 17.24449147 | 2.010582207 | 0.42829515 | 3.293482 | 0.00099 | 0.007342182 | ENSG00000250710 | OR7E99P | 81419 |
| ENSG00000040531.14 | 1133.903761 | 2.009622706 | 0.184027492 | 7.659848 | 1.86E-14 | 4.88E-13 | ENSG00000040531 | CTNS | 1497 |
| ENSG00000113070.8 | 9644.255481 | 2.008714403 | 0.355098006 | 3.967114 | 7.27E-05 | 0.0006713 | ENSG00000113070 | HBEGF | 1839 |
| ENSG00000234782.3 | 27.25748505 | 2.008079273 | 0.378738391 | 3.717815 | 0.000201 | 0.001712478 | ENSG00000234782 | NA | NA |
| ENSG00000214999.3 | 23.15667468 | 2.0072449 | 0.387853412 | 3.628291 | 0.000285 | 0.00236837 | ENSG00000214999 | LOC10798 | 1.08E+08 |
| ENSG00000196335.13 | 112.6295123 | 2.004836247 | 0.174934762 | 8.030629 | 9.70E-16 | 2.81E-14 | ENSG00000196335 | STK31 | 56164 |
| ENSG00000089723.10 | 71.52390892 | 2.003193434 | 0.292255108 | 4.801262 | 1.58E-06 | 1.85E-05 | ENSG00000089723 | OTUB2 | 78990 |
| ENSG00000108179.14 | 31815.32414 | 2.003175225 | 0.218263798 | 6.428804 | 1.29E-10 | 2.43E-09 | ENSG00000108179 | PPIF | 10105 |
| ENSG00000197019.5 | 1789.229 | 1.998789 | 0.1871198 | 7.475366 | 7.70E-14 | 1.93E-12 | ENSG00000197019 | SERTAD1 | 29950 |
| ENSG00000259868.3 | 11.4620671 | 1.998318176 | 0.459540665 | 3.042861 | 0.002343 | 0.015943324 | ENSG00000259868 | NA | NA |
| ENSG00000101577.9 | 1146.571483 | 1.997416201 | 0.146458134 | 9.541404 | 1.41E-21 | 5.91E-20 | ENSG00000101577 | LPIN2 | 9663 |
| ENSG00000242808.9 | 11.06362928 | 1.994286 | 0.437517485 | 3.186812 | 0.001439 | 0.010277993 | ENSG00000242808 | SOX2-OT | 347689 |
| ENSG00000228436.3 | 19.60828881 | 1.993623382 | 0.439384553 | 3.171762 | 0.001515 | 0.010756491 | ENSG00000228436 | LOC10537 | 1.05E+08 |
| ENSG00000173267.14 | 1533.451902 | 1.991690775 | 0.125561098 | 11.08377 | 1.50E-28 | 9.47E-27 | ENSG00000173267 | SNCG | 6623 |
| ENSG00000139725.8 | 68.64912866 | 1.991377372 | 0.270416125 | 5.14532 | 2.67E-07 | 3.48E-06 | ENSG00000139725 | RHOF | 54509 |
| ENSG00000162413.16 | 7725.654144 | 1.987446928 | 0.177921587 | 7.798081 | 6.29E-15 | 1.71E-13 | ENSG00000162413 | KLHL21 | 9903 |
| ENSG00000272734.1 | 997.1764871 | 1.984211548 | 0.111166456 | 12.4517 | 1.37E-35 | 1.16E-33 | ENSG00000272734 | ADIRF-AS | 1E+08 |
| ENSG00000161249.21 | 10935.92 | 1.983944 | 0.1567715 | 8.827773 | 1.07E-18 | 3.77E-17 | ENSG00000161249 | DMKN | 93099 |
| ENSG00000235897.1 | 122.2324526 | 1.983393795 | 0.272761086 | 5.071815 | 3.94E-07 | 5.02E-06 | ENSG00000235897 | TM4SF19-, | 1.01E+08 |
| ENSG00000250746.1 | 18.98686184 | 1.981814866 | 0.361748074 | 3.819826 | 0.000134 | 0.001178406 | ENSG00000250746 | LOC10028 | 1E+08 |
| ENSG00000170296.10 | 13.49322025 | 1.9811983 | 0.500092078 | 2.761888 | 0.005747 | 0.035332984 | ENSG00000170296 | GABARAP | 11337 |
| ENSG00000226380.9 | 313.4211622 | 1.980293314 | 0.111563758 | 12.37224 | 3.69E-35 | 3.06E-33 | ENSG00000226380 | NA | NA |
| ENSG00000235493.2 | 39.32266948 | 1.978410878 | 0.359876768 | 3.83023 | 0.000128 | 0.001133058 | ENSG00000235493 | NA | NA |
| ENSG00000236675.1 | 14.67757631 | 1.977278156 | 0.477519957 | 2.884232 | 0.003924 | 0.0252311 | ENSG00000236675 | MTX1P1 | 4581 |
| ENSG00000179859.9 | 377.3127126 | 1.975829579 | 0.172884158 | 7.958101 | 1.75E-15 | 4.97E-14 | ENSG00000179859 | RNF227 | 284023 |
| ENSG00000005882.11 | 424.5040887 | 1.975235213 | 0.193337757 | 7.113123 | 1.13E-12 | 2.58E-11 | ENSG00000005882 | PDK2 | 5164 |
| ENSG00000170786.12 | 1983.791266 | 1.974015118 | 0.153477039 | 8.952578 | 3.47E-19 | 1.26E-17 | ENSG00000170786 | SDR16C5 | 195814 |
| ENSG00000188613.7 | 161.4679727 | 1.971804138 | 0.141997015 | 9.660796 | 4.42E-22 | 1.91E-20 | ENSG00000188613 | NANOS1 | 340719 |
| ENSG00000272288.6 | 21.13870753 | 1.971551624 | 0.309954617 | 4.425008 | 9.64E-06 | 0.000102491 | ENSG00000272288 | ILRUN-AS | 1.02E+08 |
| ENSG00000145777.15 | 83.88779243 | 1.969327074 | 0.236604892 | 5.7874 | 7.15E-09 | 1.12E-07 | ENSG00000145777 | TSLP | 85480 |
| ENSG00000042493.16 | 4460.021553 | 1.968538685 | 0.16626704 | 8.230968 | 1.86E-16 | 5.64E-15 | ENSG00000042493 | CAPG | 822 |

| | | | | | | | | | |
|---|---|---|---|---|---|---|---|---|---|
| ENSG00000274317.2 | 19.65691086 | 1.967147556 | 0.432663554 | 3.15984 | 0.001579 | 0.01115001 | ENSG00000274317 | NA | NA |
| ENSG00000148634.15 | 8974.471041 | 1.966963815 | 0.195816147 | 6.980853 | 2.93E-12 | 6.43E-11 | ENSG00000148634 | HERC4 | 26091 |
| ENSG00000239503.1 | 14.97394358 | 1.965916473 | 0.480910422 | 2.840272 | 0.004508 | 0.028537177 | ENSG00000239503 | NA | NA |
| ENSG00000100985.7 | 624.055 | 1.965751 | 0.12354 | 11.05513 | 2.07E-28 | 1.29E-26 | ENSG00000100985 | MMP9 | 4318 |
| ENSG00000258376.2 | 20.21859769 | 1.964582685 | 0.49215043 | 2.772694 | 0.005559 | 0.034315605 | ENSG00000258376 | NA | NA |
| ENSG00000063180.9 | 86.62781 | 1.96226 | 0.1620357 | 8.407161 | 4.20E-17 | 1.33E-15 | ENSG00000063180 | CA11 | 770 |
| ENSG00000123353.10 | 1780.442998 | 1.961649348 | 0.144819038 | 9.402419 | 5.33E-21 | 2.18E-19 | ENSG00000123353 | ORMDL2 | 29095 |
| ENSG00000259728.5 | 25.22519144 | 1.961200139 | 0.317931208 | 4.28143 | 1.86E-05 | 0.000187898 | ENSG00000259728 | LINC0093 | 1.01E+08 |
| ENSG00000198443.6 | 25.68899078 | 1.961159666 | 0.309249041 | 4.4015 | 1.08E-05 | 0.000113078 | ENSG00000198443 | KRTAP4-1 | 85285 |
| ENSG00000171450.5 | 11.22310233 | 1.959955858 | 0.397387304 | 3.422243 | 0.000621 | 0.004819025 | ENSG00000171450 | CDK5R2 | 8941 |
| ENSG00000239521.8 | 132.7299991 | 1.958322627 | 0.20234582 | 6.712877 | 1.91E-11 | 3.90E-10 | ENSG00000239521 | CASTOR3 | 352954 |
| ENSG00000049089.15 | 25.94867198 | 1.957173335 | 0.318577349 | 4.260106 | 2.04E-05 | 0.00020527 | ENSG00000049089 | COL9A2 | 1298 |
| ENSG00000133067.17 | 59.55294461 | 1.954654355 | 0.222725172 | 6.082179 | 1.19E-09 | 2.02E-08 | ENSG00000133067 | LGR6 | 59352 |
| ENSG00000276023.5 | 441.6242122 | 1.95406419 | 0.156955972 | 8.627032 | 6.30E-18 | 2.12E-16 | ENSG00000276023 | DUSP14 | 11072 |
| ENSG00000166394.15 | 109.4933863 | 1.953573341 | 0.324194865 | 4.175184 | 2.98E-05 | 0.000291739 | ENSG00000166394 | CYB5R2 | 51700 |
| ENSG00000164675.11 | 19.86822138 | 1.953169104 | 0.325675494 | 4.154961 | 3.25E-05 | 0.00031703 | ENSG00000164675 | IQUB | 154865 |
| ENSG00000198053.11 | 1884.893 | 1.950781 | 0.1372381 | 9.842607 | 7.38E-23 | 3.31E-21 | ENSG00000198053 | SIRPA | 140885 |
| ENSG00000271605.6 | 43.59369149 | 1.950596573 | 0.36771392 | 3.672955 | 0.00024 | 0.002017365 | ENSG00000271605 | MILR1 | 284021 |
| ENSG00000117016.10 | 407.0224755 | 1.949835812 | 0.422046278 | 3.198312 | 0.001382 | 0.009908665 | ENSG00000117016 | RIMS3 | 9783 |
| ENSG00000130066.16 | 23149.12 | 1.949179 | 0.08805528 | 15.32195 | 5.46E-53 | 8.44E-51 | ENSG00000130066 | SAT1 | 6303 |
| ENSG00000120278.16 | 127.4848455 | 1.947217101 | 0.262694102 | 5.128463 | 2.92E-07 | 3.79E-06 | ENSG00000120278 | PLEKHG1 | 57480 |
| ENSG00000115041.13 | 66.32590146 | 1.946415683 | 0.238524229 | 5.644775 | 1.65E-08 | 2.49E-07 | ENSG00000115041 | KCNIP3 | 30818 |
| ENSG00000095739.11 | 91.3980298 | 1.945878742 | 0.255406434 | 5.269557 | 1.37E-07 | 1.86E-06 | ENSG00000095739 | BAMBI | 25805 |
| ENSG00000125864.14 | 215.6719 | 1.945338 | 0.1242553 | 10.82721 | 2.56E-27 | 1.48E-25 | ENSG00000125864 | BFSP1 | 631 |
| ENSG00000267733.5 | 49.71663733 | 1.94473567 | 0.216386755 | 6.214501 | 5.15E-10 | 9.17E-09 | ENSG00000267733 | NA | NA |
| ENSG00000100271.17 | 22.40327 | 1.94463 | 0.3459463 | 3.886818 | 0.000102 | 0.000916269 | ENSG00000100271 | TTLL1 | 25809 |
| ENSG00000282034.1 | 121.1635883 | 1.944490289 | 0.146585774 | 9.172038 | 4.64E-20 | 1.78E-18 | ENSG00000282034 | NA | NA |
| ENSG00000246273.8 | 199.6137409 | 1.94398677 | 0.197400071 | 6.808441 | 9.87E-12 | 2.08E-10 | ENSG00000246273 | SBF2-AS1 | 283104 |
| ENSG00000131037.15 | 855.9411 | 1.943933 | 0.2332674 | 5.761342 | 8.34E-09 | 1.30E-07 | ENSG00000131037 | EPS8L1 | 54869 |
| ENSG00000105357.17 | 1085.088 | 1.941427 | 0.2788769 | 4.810105 | 1.51E-06 | 1.77E-05 | ENSG00000105357 | MYH14 | 79784 |
| ENSG00000176845.13 | 411.443539 | 1.940278924 | 0.262495889 | 5.105904 | 3.29E-07 | 4.24E-06 | ENSG00000176845 | METRNL | 284207 |
| ENSG00000273174.1 | 18.43547307 | 1.939295861 | 0.373895772 | 3.582003 | 0.000341 | 0.002786458 | ENSG00000273174 | NA | NA |
| ENSG00000173269.14 | 250.3906428 | 1.93896905 | 0.263341087 | 5.084543 | 3.69E-07 | 4.71E-06 | ENSG00000173269 | MMRN2 | 79812 |
| ENSG00000226125.2 | 10.99944594 | 1.937373855 | 0.449181576 | 2.977357 | 0.002907 | 0.019326399 | ENSG00000226125 | LINC0190 | 151475 |
| ENSG00000099338.23 | 18.96952 | 1.936942 | 0.4000214 | 3.342176 | 0.000831 | 0.006292104 | ENSG00000099338 | CATSPER | 57828 |
| ENSG00000235831.7 | 18.22038551 | 1.934885399 | 0.349910992 | 3.814928 | 0.000136 | 0.00119964 | ENSG00000235831 | BHLHE40- | 1.01E+08 |
| ENSG00000182782.7 | 869.5383674 | 1.931738417 | 0.436271301 | 3.052546 | 0.002269 | 0.015493103 | ENSG00000182782 | HCAR2 | 338442 |
| ENSG00000099954.18 | 49.93142 | 1.930756 | 0.2720084 | 4.892334 | 9.96E-07 | 1.21E-05 | ENSG00000099954 | CECR2 | 27443 |
| ENSG00000140279.12 | 42.35806497 | 1.930395639 | 0.223603279 | 5.949804 | 2.68E-09 | 4.42E-08 | ENSG00000140279 | DUOX2 | 50506 |
| ENSG00000233198.3 | 17.493676 | 1.929307458 | 0.486244041 | 2.733828 | 0.00626 | 0.038067868 | ENSG00000233198 | RNF224 | 643596 |
| ENSG00000285933.1 | 37.75258313 | 1.929076624 | 0.305912291 | 4.344633 | 1.40E-05 | 0.000144289 | ENSG00000285933 | NA | NA |
| ENSG00000229512.1 | 20.03310723 | 1.928539758 | 0.424248192 | 3.131515 | 0.001739 | 0.012199586 | ENSG00000229512 | NA | NA |

| | | | | | | | | | |
|---|---|---|---|---|---|---|---|---|---|
| ENSG00000125170.11 | 611.6950136 | 1.926073734 | 0.145277034 | 9.127897 | 6.98E-20 | 2.64E-18 | ENSG00000125170 | DOK4 | 55715 |
| ENSG00000148483.9 | 14.6871287 | 1.92460392 | 0.416814392 | 3.177923 | 0.001483 | 0.010555882 | ENSG00000148483 | TMEM236 | 653567 |
| ENSG00000237232.8 | 10.85638 | 1.923632 | 0.4995536 | 2.64963 | 0.008058 | 0.04751277 | ENSG00000237232 | ZNF295-A[illegible] | 150142 |
| ENSG00000115129.14 | 1038.020466 | 1.921781468 | 0.119206039 | 11.08821 | 1.43E-28 | 9.05E-27 | ENSG00000115129 | TP53I3 | 9540 |
| ENSG00000106952.7 | 146.3504139 | 1.921057386 | 0.122046848 | 10.82418 | 2.64E-27 | 1.53E-25 | ENSG00000106952 | TNFSF8 | 944 |
| ENSG00000151090.19 | 75.07261084 | 1.919683985 | 0.376095942 | 3.508902 | 0.00045 | 0.003592931 | ENSG00000151090 | THRB | 7068 |
| ENSG00000248360.7 | 299.1869138 | 1.917046876 | 0.115376816 | 11.41518 | 3.51E-30 | 2.43E-28 | ENSG00000248360 | LINC0050[illegible] | 201853 |
| ENSG00000167772.12 | 4943.431928 | 1.916654269 | 0.084457807 | 15.58949 | 8.58E-55 | 1.39E-52 | ENSG00000167772 | ANGPTL4 | 51129 |
| ENSG00000176826.15 | 63.42419449 | 1.913799541 | 0.194960994 | 6.738782 | 1.60E-11 | 3.29E-10 | ENSG00000176826 | FKBP9P1 | 360132 |
| ENSG00000153391.15 | 1293.184952 | 1.911744169 | 0.063811887 | 20.55642 | 6.74E-94 | 2.60E-91 | ENSG00000153391 | INO80C | 125476 |
| ENSG00000125657.5 | 1494.041665 | 1.909682367 | 0.092120165 | 14.21711 | 7.18E-46 | 8.97E-44 | ENSG00000125657 | TNFSF9 | 8744 |
| ENSG00000058668.14 | 7062.088545 | 1.909021234 | 0.137785453 | 9.500431 | 2.09E-21 | 8.65E-20 | ENSG00000058668 | ATP2B4 | 493 |
| ENSG00000159588.15 | 74.06826187 | 1.908095509 | 0.179920179 | 7.270421 | 3.58E-13 | 8.51E-12 | ENSG00000159588 | CCDC17 | 149483 |
| ENSG00000256975.1 | 19.22739515 | 1.907332503 | 0.356762467 | 3.664434 | 0.000248 | 0.002079806 | ENSG00000256975 | NA | NA |
| ENSG00000088899.15 | 741.1396 | 1.905829 | 0.1279079 | 10.20914 | 1.80E-24 | 8.78E-23 | ENSG00000088899 | LZTS3 | 9762 |
| ENSG00000274322.1 | 14.16654 | 1.905287 | 0.3607113 | 3.618646 | 0.000296 | 0.002447704 | ENSG00000274322 | NA | NA |
| ENSG00000118898.16 | 11979.64401 | 1.902245601 | 0.209953058 | 6.202556 | 5.56E-10 | 9.86E-09 | ENSG00000118898 | PPL | 5493 |
| ENSG00000204323.5 | 22.19368067 | 1.901768138 | 0.392341624 | 3.317946 | 0.000907 | 0.006794654 | ENSG00000204323 | SMIM5 | 643008 |
| ENSG00000181690.8 | 433.3092425 | 1.900338452 | 0.125135638 | 10.39143 | 2.71E-25 | 1.40E-23 | ENSG00000181690 | PLAG1 | 5324 |
| ENSG00000196611.5 | 9669.313372 | 1.896657479 | 0.345068115 | 3.757686 | 0.000171 | 0.001482294 | ENSG00000196611 | MMP1 | 4312 |
| ENSG00000274565.1 | 17.91949987 | 1.896396232 | 0.357887094 | 3.622361 | 0.000292 | 0.002418066 | ENSG00000274565 | NA | NA |
| ENSG00000240476.1 | 21.79148001 | 1.89387775 | 0.37050142 | 3.492234 | 0.000479 | 0.003805397 | ENSG00000240476 | NA | NA |
| ENSG00000162591.16 | 551.7471865 | 1.884884578 | 0.238174766 | 5.394713 | 6.86E-08 | 9.62E-07 | ENSG00000162591 | MEGF6 | 1953 |
| ENSG00000115963.13 | 7098.715741 | 1.884819944 | 0.114837395 | 11.18817 | 4.66E-29 | 3.04E-27 | ENSG00000115963 | RND3 | 390 |
| ENSG00000100439.10 | 610.2284577 | 1.884251559 | 0.114172195 | 11.24837 | 2.36E-29 | 1.57E-27 | ENSG00000100439 | ABHD4 | 63874 |
| ENSG00000169504.15 | 22165.93764 | 1.883501369 | 0.115541061 | 11.10862 | 1.14E-28 | 7.24E-27 | ENSG00000169504 | CLIC4 | 25932 |
| ENSG00000231991.4 | 180.5524531 | 1.8807309 | 0.135414692 | 9.457843 | 3.14E-21 | 1.29E-19 | ENSG00000231991 | NA | NA |
| ENSG00000068078.18 | 1532.121273 | 1.87937645 | 0.363189276 | 3.522616 | 0.000427 | 0.003424394 | ENSG00000068078 | FGFR3 | 2261 |
| ENSG00000159713.11 | 24.92977797 | 1.876972016 | 0.323587289 | 3.946298 | 7.94E-05 | 0.000728856 | ENSG00000159713 | TPPP3 | 51673 |
| ENSG00000273062.1 | 11.52824647 | 1.874889768 | 0.393636047 | 3.238753 | 0.001201 | 0.008737202 | ENSG00000273062 | NA | NA |
| ENSG00000070808.15 | 18.20167918 | 1.872161489 | 0.437463505 | 2.90804 | 0.003637 | 0.02356977 | ENSG00000070808 | CAMK2A | 815 |
| ENSG00000224621.1 | 53.57097713 | 1.870507164 | 0.259837522 | 4.889622 | 1.01E-06 | 1.22E-05 | ENSG00000224621 | NA | NA |
| ENSG00000287958.1 | 14.5635589 | 1.869999526 | 0.374162617 | 3.394245 | 0.000688 | 0.005298081 | ENSG00000287958 | NA | NA |
| ENSG00000184227.8 | 201.4312393 | 1.868256577 | 0.15999523 | 7.92684 | 2.25E-15 | 6.36E-14 | ENSG00000184227 | ACOT1 | 641371 |
| ENSG00000183496.6 | 46.25218683 | 1.867923838 | 0.239792025 | 5.287598 | 1.24E-07 | 1.69E-06 | ENSG00000183496 | MEX3B | 84206 |
| ENSG00000070610.14 | 4292.268584 | 1.866403087 | 0.22999451 | 5.506232 | 3.67E-08 | 5.28E-07 | ENSG00000070610 | GBA2 | 57704 |
| ENSG00000018280.17 | 12.60696776 | 1.865879315 | 0.44229872 | 2.862046 | 0.004209 | 0.026882899 | ENSG00000018280 | SLC11A1 | 6556 |
| ENSG00000283148.1 | 15.08191277 | 1.865615488 | 0.420787869 | 3.007728 | 0.002632 | 0.017721643 | ENSG00000283148 | NA | NA |
| ENSG00000224080.1 | 66.92890674 | 1.864930555 | 0.17345247 | 7.292664 | 3.04E-13 | 7.26E-12 | ENSG00000224080 | NA | NA |
| ENSG00000178750.3 | 18.24304662 | 1.863378875 | 0.404838111 | 3.120701 | 0.001804 | 0.012593508 | ENSG00000178750 | STX19 | 415117 |
| ENSG00000230524.8 | 27.82369919 | 1.861308171 | 0.34813343 | 3.62306 | 0.000291 | 0.002412292 | ENSG00000230524 | COL6A4P[illegible] | 344875 |
| ENSG00000173546.7 | 199.4135406 | 1.861217405 | 0.223249015 | 5.649375 | 1.61E-08 | 2.43E-07 | ENSG00000173546 | CSPG4 | 1464 |

| | | | | | | | | | |
|---|---|---|---|---|---|---|---|---|---|
| ENSG00000213344.2 | 63.79588442 | 1.860759269 | 0.210992366 | 5.975379 | 2.30E-09 | 3.80E-08 | ENSG00000213344 | NA | NA |
| ENSG00000262152.7 | 120.8681214 | 1.858947434 | 0.275443521 | 4.570619 | 4.86E-06 | 5.37E-05 | ENSG00000262152 | NA | NA |
| ENSG00000139626.16 | 17.61979333 | 1.858794738 | 0.436421811 | 2.884353 | 0.003922 | 0.025227437 | ENSG00000139626 | ITGB7 | 3695 |
| ENSG00000100994.12 | 19487.14 | 1.858561 | 0.1360071 | 9.253637 | 2.17E-20 | 8.53E-19 | ENSG00000100994 | PYGB | 5834 |
| ENSG00000131773.14 | 20.55294857 | 1.857806791 | 0.336946972 | 3.732952 | 0.000189 | 0.001619995 | ENSG00000131773 | KHDRBS3 | 10656 |
| ENSG00000167183.3 | 13.40684214 | 1.857679863 | 0.417968257 | 3.009032 | 0.002621 | 0.017668103 | ENSG00000167183 | PRR15L | 79170 |
| ENSG00000130783.13 | 21.91569368 | 1.856793531 | 0.32062827 | 3.919784 | 8.86E-05 | 0.000806931 | ENSG00000130783 | CCDC62 | 84660 |
| ENSG00000278558.4 | 18.23458 | 1.853099 | 0.3218813 | 3.893046 | 9.90E-05 | 0.000894873 | ENSG00000278558 | TMEM191I | 728229 |
| ENSG00000196689.12 | 110.2931313 | 1.852985269 | 0.262754235 | 4.768659 | 1.85E-06 | 2.16E-05 | ENSG00000196689 | TRPV1 | 7442 |
| ENSG00000164171.11 | 71467.81679 | 1.852152081 | 0.130369452 | 9.604643 | 7.64E-22 | 3.27E-20 | ENSG00000164171 | ITGA2 | 3673 |
| ENSG00000132003.9 | 325.8028627 | 1.850771468 | 0.157240263 | 7.954524 | 1.80E-15 | 5.11E-14 | ENSG00000132003 | ZSWIM4 | 65249 |
| ENSG00000074181.9 | 1076.803356 | 1.849545162 | 0.262610246 | 4.758174 | 1.95E-06 | 2.27E-05 | ENSG00000074181 | NOTCH3 | 4854 |
| ENSG00000284969.1 | 10.82620252 | 1.847780983 | 0.457261249 | 2.728814 | 0.006356 | 0.038589763 | ENSG00000284969 | NA | NA |
| ENSG00000249755.2 | 10.43618802 | 1.846982421 | 0.451574934 | 2.761408 | 0.005755 | 0.03537571 | ENSG00000249755 | NA | NA |
| ENSG00000145451.13 | 10.52127687 | 1.846480521 | 0.457130498 | 2.72675 | 0.006396 | 0.038787773 | ENSG00000145451 | GLRA3 | 8001 |
| ENSG00000255325.2 | 30.40640801 | 1.845036827 | 0.30458834 | 4.087605 | 4.36E-05 | 0.000417231 | ENSG00000255325 | NA | NA |
| ENSG00000184599.13 | 111.7401135 | 1.843956387 | 0.145981638 | 8.521321 | 1.58E-17 | 5.17E-16 | ENSG00000184599 | TAFA3 | 284467 |
| ENSG00000136425.13 | 393.3576831 | 1.843893029 | 0.178498348 | 6.968653 | 3.20E-12 | 7.00E-11 | ENSG00000136425 | CIB2 | 10518 |
| ENSG00000145107.15 | 162.1483455 | 1.840944376 | 0.265025806 | 4.682353 | 2.84E-06 | 3.22E-05 | ENSG00000145107 | TM4SF19 | 116211 |
| ENSG00000122641.11 | 6524.089603 | 1.840515271 | 0.077214471 | 16.06584 | 4.43E-58 | 8.02E-56 | ENSG00000122641 | INHBA | 3624 |
| ENSG00000149809.14 | 199.4913677 | 1.840338653 | 0.1811592 | 6.846678 | 7.56E-12 | 1.61E-10 | ENSG00000149809 | TM7SF2 | 7108 |
| ENSG00000130724.8 | 3071.631 | 1.837735 | 0.160951 | 7.690136 | 1.47E-14 | 3.88E-13 | ENSG00000130724 | CHMP2A | 27243 |
| ENSG00000099822.3 | 81.80502528 | 1.837645982 | 0.148030782 | 8.360734 | 6.23E-17 | 1.95E-15 | ENSG00000099822 | HCN2 | 610 |
| ENSG00000164096.13 | 3254.940356 | 1.836189704 | 0.159473522 | 7.751692 | 9.07E-15 | 2.43E-13 | ENSG00000164096 | C4orf3 | 401152 |
| ENSG00000260618.1 | 12.49134187 | 1.835727188 | 0.408832172 | 3.022578 | 0.002506 | 0.016947697 | ENSG00000260618 | NA | NA |
| ENSG00000253819.2 | 60.30299397 | 1.835532919 | 0.237279322 | 5.207082 | 1.92E-07 | 2.55E-06 | ENSG00000253819 | NA | NA |
| ENSG00000237575.4 | 45.91298861 | 1.833986389 | 0.275133773 | 4.485041 | 7.29E-06 | 7.91E-05 | ENSG00000237575 | PYY2 | 23615 |
| ENSG00000172738.12 | 84.06891445 | 1.831434998 | 0.204114949 | 6.033047 | 1.61E-09 | 2.70E-08 | ENSG00000172738 | TMEM217 | 221468 |
| ENSG00000214725.9 | 12.44714395 | 1.829232367 | 0.364187984 | 3.375269 | 0.000737 | 0.005641386 | ENSG00000214725 | CDIPTOSI | 440356 |
| ENSG00000198910.13 | 499.2172 | 1.828849 | 0.1541998 | 7.969201 | 1.60E-15 | 4.57E-14 | ENSG00000198910 | L1CAM | 3897 |
| ENSG00000196998.19 | 855.3027 | 1.827667 | 0.1477719 | 8.307851 | 9.75E-17 | 3.01E-15 | ENSG00000196998 | WDR45 | 11152 |
| ENSG00000267121.6 | 158.4204555 | 1.826903292 | 0.220769508 | 5.557395 | 2.74E-08 | 4.04E-07 | ENSG00000267121 | LOC33919 | 339192 |
| ENSG00000119782.14 | 62.85942588 | 1.826077405 | 0.279920549 | 4.380091 | 1.19E-05 | 0.000123995 | ENSG00000119782 | FKBP1B | 2281 |
| ENSG00000150991.15 | 82495.92767 | 1.825576787 | 0.123341675 | 9.936437 | 2.89E-23 | 1.33E-21 | ENSG00000150991 | UBC | 7316 |
| ENSG00000255202.1 | 14.80054885 | 1.825428641 | 0.356968947 | 3.432872 | 0.000597 | 0.004655727 | ENSG00000255202 | NA | NA |
| ENSG00000257906.3 | 55.03021938 | 1.825089116 | 0.199349809 | 6.145424 | 7.98E-10 | 1.39E-08 | ENSG00000257906 | NA | NA |
| ENSG00000172493.20 | 7267.977405 | 1.824357676 | 0.127443276 | 9.607079 | 7.46E-22 | 3.19E-20 | ENSG00000172493 | AFF1 | 4299 |
| ENSG00000141682.11 | 6985.728697 | 1.824128184 | 0.417766142 | 2.930176 | 0.003388 | 0.022131893 | ENSG00000141682 | PMAIP1 | 5366 |
| ENSG00000267577.1 | 18.1533 | 1.823758 | 0.3697497 | 3.309694 | 0.000934 | 0.006972639 | ENSG00000267577 | NA | NA |
| ENSG00000182795.13 | 8775.157613 | 1.821092235 | 0.233083786 | 5.238855 | 1.62E-07 | 2.17E-06 | ENSG00000182795 | C1orf116 | 79098 |
| ENSG00000047346.12 | 729.2174041 | 1.820459926 | 0.167312611 | 7.294489 | 3.00E-13 | 7.16E-12 | ENSG00000047346 | FAM214A | 56204 |
| ENSG00000261104.1 | 16.82682893 | 1.819738984 | 0.406276952 | 3.002235 | 0.00268 | 0.018003605 | ENSG00000261104 | NA | NA |

| | | | | | | | | | |
|---|---|---|---|---|---|---|---|---|---|
| ENSG00000113161.16 | 15975.6271 | 1.819596436 | 0.092044516 | 13.25007 | 4.51E-40 | 4.53E-38 | ENSG00000113161 | HMGCR | 3156 |
| ENSG00000258655.2 | 96.16645103 | 1.81911964 | 0.251463127 | 4.848105 | 1.25E-06 | 1.48E-05 | ENSG00000258655 | NA | NA |
| ENSG00000213859.6 | 1512.195161 | 1.818144875 | 0.169565464 | 7.183921 | 6.77E-13 | 1.57E-11 | ENSG00000213859 | KCTD11 | 147040 |
| ENSG00000185022.12 | 2321.935 | 1.817423 | 0.2020757 | 6.024588 | 1.70E-09 | 2.84E-08 | ENSG00000185022 | MAFF | 23764 |
| ENSG00000160213.7 | 10013.35 | 1.816954 | 0.1436126 | 8.473864 | 2.37E-17 | 7.69E-16 | ENSG00000160213 | CSTB | 1476 |
| ENSG00000120051.15 | 32.80687927 | 1.816408407 | 0.264977799 | 4.590605 | 4.42E-06 | 4.90E-05 | ENSG00000120051 | CFAP58 | 159686 |
| ENSG00000177294.7 | 12.12993589 | 1.816406734 | 0.416319704 | 2.921809 | 0.00348 | 0.022679512 | ENSG00000177294 | FBXO39 | 162517 |
| ENSG00000108602.18 | 395.0387494 | 1.815678108 | 0.169390549 | 7.176776 | 7.14E-13 | 1.65E-11 | ENSG00000108602 | ALDH3A1 | 218 |
| ENSG00000228363.2 | 30.76679298 | 1.81542851 | 0.246121367 | 4.93833 | 7.88E-07 | 9.67E-06 | ENSG00000228363 | NA | NA |
| ENSG00000010278.14 | 19870.2864 | 1.813648214 | 0.111225811 | 10.91157 | 1.01E-27 | 5.99E-26 | ENSG00000010278 | CD9 | 928 |
| ENSG00000260118.1 | 19.35482 | 1.807716 | 0.3225358 | 3.74444 | 0.000181 | 0.001554642 | ENSG00000260118 | NA | NA |
| ENSG00000008118.10 | 15.50258378 | 1.80663935 | 0.378678019 | 3.186452 | 0.00144 | 0.010282514 | ENSG00000008118 | CAMK1G | 57172 |
| ENSG00000163810.11 | 13.50468216 | 1.805830284 | 0.411732207 | 2.928676 | 0.003404 | 0.022233479 | ENSG00000163810 | TGM4 | 7047 |
| ENSG00000079385.22 | 418.6491 | 1.80436 | 0.3849755 | 3.128407 | 0.001758 | 0.01231312 | ENSG00000079385 | CEACAM1 | 634 |
| ENSG00000175182.15 | 776.332998 | 1.80220145 | 0.196041008 | 6.132398 | 8.66E-10 | 1.49E-08 | ENSG00000175182 | FAM131A | 131408 |
| ENSG00000205890.3 | 97.97705686 | 1.801229472 | 0.310839915 | 3.864463 | 0.000111 | 0.000998291 | ENSG00000205890 | LOC10012 | 1E+08 |
| ENSG00000158201.10 | 1819.494751 | 1.801077031 | 0.125330296 | 9.583294 | 9.40E-22 | 3.99E-20 | ENSG00000158201 | ABHD3 | 171586 |
| ENSG00000214176.9 | 977.2071 | 1.800091252 | 0.177277627 | 6.769558 | 1.29E-11 | 2.68E-10 | ENSG00000214176 | NA | NA |
| ENSG00000198732.10 | 1207.575146 | 1.797694364 | 0.200465963 | 5.974552 | 2.31E-09 | 3.82E-08 | ENSG00000198732 | SMOC1 | 64093 |
| ENSG00000106299.8 | 6321.044485 | 1.797491565 | 0.196547783 | 6.092623 | 1.11E-09 | 1.90E-08 | ENSG00000106299 | WASL | 8976 |
| ENSG00000234997.2 | 47.82016125 | 1.796622249 | 0.220272186 | 5.432471 | 5.56E-08 | 7.84E-07 | ENSG00000234997 | NA | NA |
| ENSG00000021355.13 | 19168.12058 | 1.795040635 | 0.227395639 | 5.255337 | 1.48E-07 | 2.00E-06 | ENSG00000021355 | SERPINB1 | 1992 |
| ENSG00000284543.1 | 15.11467672 | 1.793906259 | 0.349528184 | 3.415765 | 0.000636 | 0.004925072 | ENSG00000284543 | NA | NA |
| ENSG00000074416.14 | 1282.786864 | 1.792657162 | 0.277302364 | 4.300927 | 1.70E-05 | 0.000172957 | ENSG00000074416 | MGLL | 11343 |
| ENSG00000077782.20 | 279.7627761 | 1.791534217 | 0.202031724 | 5.897758 | 3.68E-09 | 5.97E-08 | ENSG00000077782 | FGFR1 | 2260 |
| ENSG00000139899.10 | 51.50348429 | 1.791045989 | 0.20919619 | 5.69344 | 1.25E-08 | 1.90E-07 | ENSG00000139899 | CBLN3 | 643866 |
| ENSG00000197892.13 | 2531.681659 | 1.790116021 | 0.20016324 | 5.945727 | 2.75E-09 | 4.53E-08 | ENSG00000197892 | KIF13B | 23303 |
| ENSG00000180914.10 | 127.6435786 | 1.789309295 | 0.403756757 | 2.945608 | 0.003223 | 0.02118718 | ENSG00000180914 | OXTR | 5021 |
| ENSG00000130766.5 | 515.771032 | 1.787963637 | 0.183398959 | 6.477483 | 9.33E-11 | 1.78E-09 | ENSG00000130766 | SESN2 | 83667 |
| ENSG00000258763.5 | 13.00364078 | 1.783903698 | 0.397505581 | 2.978332 | 0.002898 | 0.019274621 | ENSG00000258763 | NA | NA |
| ENSG00000182718.16 | 171512.9404 | 1.783455774 | 0.097394573 | 12.15115 | 5.66E-34 | 4.48E-32 | ENSG00000182718 | ANXA2 | 302 |
| ENSG00000283511.1 | 124.4795677 | 1.780518673 | 0.177321308 | 6.657512 | 2.79E-11 | 5.60E-10 | ENSG00000283511 | NA | NA |
| ENSG00000286355.1 | 21.41788804 | 1.780495266 | 0.377954843 | 3.123376 | 0.001788 | 0.012505827 | ENSG00000286355 | NA | NA |
| ENSG00000138641.18 | 32.04269533 | 1.780331391 | 0.3584006 | 3.29333 | 0.00099 | 0.0073441 | ENSG00000138641 | HERC3 | 8916 |
| ENSG00000279118.1 | 81.49347558 | 1.778413622 | 0.254082988 | 4.637908 | 3.52E-06 | 3.95E-05 | ENSG00000279118 | NA | NA |
| ENSG00000136867.11 | 79.44711907 | 1.77813957 | 0.206390417 | 5.708306 | 1.14E-08 | 1.75E-07 | ENSG00000136867 | SLC31A2 | 1318 |
| ENSG00000026751.17 | 112.4097862 | 1.777859365 | 0.177488861 | 6.636244 | 3.22E-11 | 6.42E-10 | ENSG00000026751 | SLAMF7 | 57823 |
| ENSG00000101670.12 | 1021.455408 | 1.777135924 | 0.126914996 | 9.274995 | 1.78E-20 | 7.03E-19 | ENSG00000101670 | LIPG | 9388 |
| ENSG00000026025.16 | 9610.401468 | 1.776604382 | 0.12582567 | 9.351068 | 8.68E-21 | 3.51E-19 | ENSG00000026025 | VIM | 7431 |
| ENSG00000137767.14 | 2531.646334 | 1.775624743 | 0.24871597 | 4.726776 | 2.28E-06 | 2.62E-05 | ENSG00000137767 | SQOR | 58472 |
| ENSG00000240163.1 | 28.80584239 | 1.775523372 | 0.352017028 | 3.339393 | 0.00084 | 0.00634526 | ENSG00000240163 | NA | NA |
| ENSG00000135127.11 | 397.4006363 | 1.77438068 | 0.155434275 | 7.555481 | 4.17E-14 | 1.06E-12 | ENSG00000135127 | BICDL1 | 92558 |

| | | | | | | | | | |
|---|---|---|---|---|---|---|---|---|---|
| ENSG00000181467.4 | 6010.233136 | 1.773439672 | 0.126915182 | 9.245857 | 2.33E-20 | 9.15E-19 | ENSG00000181467 | RAP2B | 5912 |
| ENSG00000232027.2 | 12.21052702 | 1.770953255 | 0.413335381 | 2.832937 | 0.004612 | 0.02909647 | ENSG00000232027 | NA | NA |
| ENSG00000213889.10 | 29.11013 | 1.770895 | 0.2686655 | 4.358188 | 1.31E-05 | 0.00013622 | ENSG00000213889 | PPM1N | 147699 |
| ENSG00000165935.9 | 67.07457863 | 1.7701703 | 0.280104521 | 4.17762 | 2.95E-05 | 0.000288846 | ENSG00000165935 | SMCO2 | 341346 |
| ENSG00000143590.14 | 69.89695337 | 1.768724948 | 0.219265496 | 5.330182 | 9.81E-08 | 1.35E-06 | ENSG00000143590 | EFNA3 | 1944 |
| ENSG00000263006.6 | 168.603871 | 1.768156254 | 0.165027145 | 7.07857 | 1.46E-12 | 3.27E-11 | ENSG00000263006 | ROCK1P1 | 727758 |
| ENSG00000160145.15 | 61.42386526 | 1.766617593 | 0.269981515 | 4.321102 | 1.55E-05 | 0.000158966 | ENSG00000160145 | KALRN | 8997 |
| ENSG00000198835.4 | 90.37043827 | 1.766034472 | 0.176059775 | 6.622947 | 3.52E-11 | 6.98E-10 | ENSG00000198835 | GJC2 | 57165 |
| ENSG00000205517.12 | 13.74606681 | 1.765844734 | 0.360987601 | 3.229598 | 0.00124 | 0.008992325 | ENSG00000205517 | RGL3 | 57139 |
| ENSG00000160752.14 | 5383.679244 | 1.764876506 | 0.042004998 | 27.73185 | 2.88E-169 | 2.40E-166 | ENSG00000160752 | FDPS | 2224 |
| ENSG00000006459.11 | 2176.077286 | 1.763303734 | 0.134947068 | 8.620445 | 6.67E-18 | 2.24E-16 | ENSG00000006459 | KDM7A | 80853 |
| ENSG00000145390.11 | 4189.115825 | 1.763099937 | 0.070344069 | 16.53444 | 2.07E-61 | 4.18E-59 | ENSG00000145390 | USP53 | 54532 |
| ENSG00000137266.14 | 509.8653485 | 1.762486069 | 0.185767291 | 6.257754 | 3.91E-10 | 7.06E-09 | ENSG00000137266 | SLC22A23 | 63027 |
| ENSG00000168491.10 | 11.53855009 | 1.758710696 | 0.392037385 | 2.955613 | 0.00312 | 0.020588033 | ENSG00000168491 | CCDC110 | 256309 |
| ENSG00000170919.16 | 509.9512472 | 1.757524083 | 0.230965486 | 5.011676 | 5.40E-07 | 6.75E-06 | ENSG00000170919 | NA | NA |
| ENSG00000165917.10 | 13.83606158 | 1.757088272 | 0.437643939 | 2.643903 | 0.008196 | 0.048227987 | ENSG00000165917 | RAPSN | 5913 |
| ENSG00000170873.18 | 2241.748933 | 1.754055748 | 0.104373874 | 11.05694 | 2.03E-28 | 1.27E-26 | ENSG00000170873 | MTSS1 | 9788 |
| ENSG00000165948.11 | 209.0392776 | 1.753973424 | 0.211481993 | 5.456604 | 4.85E-08 | 6.89E-07 | ENSG00000165948 | IFI27L1 | 122509 |
| ENSG00000172361.6 | 144.9017013 | 1.753654407 | 0.131042976 | 8.803634 | 1.32E-18 | 4.64E-17 | ENSG00000172361 | CFAP53 | 220136 |
| ENSG00000271447.6 | 169.3369291 | 1.753504895 | 0.211603998 | 5.451243 | 5.00E-08 | 7.08E-07 | ENSG00000271447 | MMP28 | 79148 |
| ENSG00000188643.11 | 29982.47026 | 1.753100477 | 0.120454144 | 9.572942 | 1.04E-21 | 4.40E-20 | ENSG00000188643 | S100A16 | 140576 |
| ENSG00000139714.12 | 33.30422238 | 1.749462077 | 0.379577485 | 3.028267 | 0.00246 | 0.016661473 | ENSG00000139714 | MORN3 | 283385 |
| ENSG00000246695.8 | 218.9062434 | 1.746092097 | 0.136241621 | 8.412202 | 4.02E-17 | 1.28E-15 | ENSG00000246695 | NA | NA |
| ENSG00000158458.20 | 88.94755265 | 1.742958898 | 0.194411807 | 5.879061 | 4.13E-09 | 6.64E-08 | ENSG00000158458 | NRG2 | 9542 |
| ENSG00000176170.13 | 5091.78545 | 1.742244598 | 0.123674365 | 9.235904 | 2.56E-20 | 1.00E-18 | ENSG00000176170 | SPHK1 | 8877 |
| ENSG00000153292.16 | 558.0772275 | 1.741187159 | 0.221612111 | 5.14948 | 2.61E-07 | 3.41E-06 | ENSG00000153292 | ADGRF1 | 266977 |
| ENSG00000138756.17 | 1729.949784 | 1.740904415 | 0.149552867 | 7.62877 | 2.37E-14 | 6.17E-13 | ENSG00000138756 | BMP2K | 55589 |
| ENSG00000249180.1 | 15.41513525 | 1.739764134 | 0.392654717 | 2.902713 | 0.003699 | 0.023892951 | ENSG00000249180 | NA | NA |
| ENSG00000248727.6 | 39.78257274 | 1.739559446 | 0.219005407 | 5.203339 | 1.96E-07 | 2.60E-06 | ENSG00000248727 | LINC0194[illegible] | 1.02E+08 |
| ENSG00000153064.12 | 30.34810554 | 1.738571451 | 0.316006473 | 3.603 | 0.000315 | 0.002583879 | ENSG00000153064 | BANK1 | 55024 |
| ENSG00000241112.1 | 14.23510265 | 1.737933888 | 0.344611679 | 3.302076 | 0.00096 | 0.007144771 | ENSG00000241112 | NA | NA |
| ENSG00000185561.10 | 9.99899503 | 1.737025229 | 0.40072149 | 2.837445 | 0.004548 | 0.028736447 | ENSG00000185561 | TLCD2 | 727910 |
| ENSG00000128342.5 | 1465.859 | 1.736581 | 0.253596 | 4.481857 | 7.40E-06 | 8.01E-05 | ENSG00000128342 | LIF | 3976 |
| ENSG00000248740.6 | 57.022272 | 1.735800912 | 0.307835491 | 3.689636 | 0.000225 | 0.001899082 | ENSG00000248740 | NA | NA |
| ENSG00000178773.15 | 63.24528147 | 1.735624453 | 0.219894774 | 5.164399 | 2.41E-07 | 3.16E-06 | ENSG00000178773 | CPNE7 | 27132 |
| ENSG00000179674.3 | 64.29162773 | 1.735546922 | 0.279667752 | 4.060343 | 4.90E-05 | 0.000464069 | ENSG00000179674 | ARL14 | 80117 |
| ENSG00000108830.10 | 17.65810563 | 1.735308359 | 0.349236285 | 3.250832 | 0.001151 | 0.008408838 | ENSG00000108830 | RND2 | 8153 |
| ENSG00000138166.6 | 4817.47811 | 1.734635739 | 0.199275694 | 5.693799 | 1.24E-08 | 1.90E-07 | ENSG00000138166 | DUSP5 | 1847 |
| ENSG00000243708.10 | 26.31970532 | 1.734114371 | 0.408899068 | 2.77358 | 0.005544 | 0.034230241 | ENSG00000243708 | PLA2G4B | 1E+08 |
| ENSG00000143669.13 | 2174.413914 | 1.733800535 | 0.168507922 | 6.72847 | 1.71E-11 | 3.52E-10 | ENSG00000143669 | LYST | 1130 |
| ENSG00000259158.4 | 15.71122136 | 1.73217111 | 0.383100334 | 2.955286 | 0.003124 | 0.020594746 | ENSG00000259158 | ADAM20P | 317760 |
| ENSG00000069011.16 | 6648.653192 | 1.73206818 | 0.180775361 | 6.262292 | 3.79E-10 | 6.87E-09 | ENSG00000069011 | PITX1 | 5307 |

| | | | | | | | | | |
|---|---|---|---|---|---|---|---|---|---|
| ENSG00000227359.1 | 13.08059793 | 1.731911255 | 0.346668949 | 3.265107 | 0.001094 | 0.008044879 | ENSG00000227359 | NA | NA |
| ENSG00000197372.9 | 367.3658 | 1.731904 | 0.1824046 | 6.205455 | 5.45E-10 | 9.69E-09 | ENSG00000197372 | ZNF675 | 171392 |
| ENSG00000172137.19 | 122.3464486 | 1.731754246 | 0.159606898 | 7.090886 | 1.33E-12 | 3.01E-11 | ENSG00000172137 | CALB2 | 794 |
| ENSG00000275964.1 | 37.0846115 | 1.728123789 | 0.322023805 | 3.503231 | 0.00046 | 0.003663718 | ENSG00000275964 | NA | NA |
| ENSG00000260992.1 | 33.02999814 | 1.72742773 | 0.315107577 | 3.577914 | 0.000346 | 0.002827497 | ENSG00000260992 | NA | NA |
| ENSG00000128578.10 | 2441.183171 | 1.727070383 | 0.233964393 | 4.817273 | 1.46E-06 | 1.71E-05 | ENSG00000128578 | STRIP2 | 57464 |
| ENSG00000197140.15 | 23.01892949 | 1.726618587 | 0.343505659 | 3.279767 | 0.001039 | 0.007678605 | ENSG00000197140 | ADAM32 | 203102 |
| ENSG00000225855.7 | 318.6022556 | 1.725778276 | 0.151439642 | 7.433841 | 1.05E-13 | 2.62E-12 | ENSG00000225855 | RUSC1-AS | 284618 |
| ENSG00000175505.11 | 681.5953892 | 1.724985073 | 0.147541465 | 7.624874 | 2.44E-14 | 6.35E-13 | ENSG00000175505 | CLCF1 | 23529 |
| ENSG00000240086.7 | 22.29577884 | 1.721416588 | 0.321163657 | 3.491729 | 0.00048 | 0.003810322 | ENSG00000240086 | NA | NA |
| ENSG00000248161.5 | 16.64533034 | 1.720648026 | 0.384230144 | 2.916606 | 0.003539 | 0.022990334 | ENSG00000248161 | NA | NA |
| ENSG00000154227.13 | 936.6796659 | 1.719904371 | 0.218088913 | 5.135082 | 2.82E-07 | 3.67E-06 | ENSG00000154227 | CERS3 | 204219 |
| ENSG00000238279.1 | 123.8927011 | 1.713707157 | 0.20252782 | 5.499033 | 3.82E-08 | 5.48E-07 | ENSG00000238279 | NA | NA |
| ENSG00000120885.21 | 258.8734545 | 1.71247279 | 0.182191846 | 6.106051 | 1.02E-09 | 1.75E-08 | ENSG00000120885 | CLU | 1191 |
| ENSG00000107738.20 | 1322.489956 | 1.711871489 | 0.219612624 | 5.062876 | 4.13E-07 | 5.24E-06 | ENSG00000107738 | VSIR | 64115 |
| ENSG00000126005.17 | 478.0073 | 1.708598 | 0.215283 | 5.149493 | 2.61E-07 | 3.41E-06 | ENSG00000126005 | MMP24OS | 1.01E+08 |
| ENSG00000159348.13 | 2386.724572 | 1.708500037 | 0.144356535 | 7.678904 | 1.60E-14 | 4.22E-13 | ENSG00000159348 | CYB5R1 | 51706 |
| ENSG00000010295.19 | 46.98290285 | 1.707534662 | 0.242654402 | 4.564247 | 5.01E-06 | 5.52E-05 | ENSG00000010295 | IFFO1 | 25900 |
| ENSG00000034713.8 | 4094.577213 | 1.70453933 | 0.161421987 | 6.842558 | 7.78E-12 | 1.65E-10 | ENSG00000034713 | GABARAP | 11345 |
| ENSG00000167106.12 | 1226.221388 | 1.70399156 | 0.175301937 | 6.297658 | 3.02E-10 | 5.56E-09 | ENSG00000167106 | FAM102A | 399665 |
| ENSG00000124357.13 | 1813.673564 | 1.702824856 | 0.166508937 | 6.623217 | 3.51E-11 | 6.97E-10 | ENSG00000124357 | NAGK | 55577 |
| ENSG00000155918.7 | 146.9709254 | 1.701132423 | 0.298783914 | 3.685381 | 0.000228 | 0.001928142 | ENSG00000155918 | RAET1L | 154064 |
| ENSG00000186594.14 | 119.855779 | 1.700841569 | 0.138933183 | 7.923532 | 2.31E-15 | 6.52E-14 | ENSG00000186594 | MIR22HG | 84981 |
| ENSG00000204758.8 | 62.39437062 | 1.700518522 | 0.228183697 | 4.82295 | 1.41E-06 | 1.67E-05 | ENSG00000204758 | LOC10026 | 1E+08 |
| ENSG00000254614.2 | 67.64440566 | 1.69939041 | 0.235566965 | 4.666997 | 3.06E-06 | 3.45E-05 | ENSG00000254614 | LOC72897 | 728975 |
| ENSG00000253618.1 | 12.76229276 | 1.699330687 | 0.403230898 | 2.726306 | 0.006405 | 0.038822327 | ENSG00000253618 | GRPEL2-A | 1.06E+08 |
| ENSG00000197822.11 | 682.0382985 | 1.699167706 | 0.210872109 | 5.212485 | 1.86E-07 | 2.48E-06 | ENSG00000197822 | OCLN | 1.01E+08 |
| ENSG00000244675.2 | 197.5960695 | 1.698345982 | 0.183461774 | 5.986784 | 2.14E-09 | 3.55E-08 | ENSG00000244675 | NA | NA |
| ENSG00000256268.2 | 317.2091049 | 1.698034351 | 0.130934218 | 8.386153 | 5.02E-17 | 1.58E-15 | ENSG00000256268 | LINC0245 | 1.05E+08 |
| ENSG00000171385.9 | 20.91422176 | 1.697368437 | 0.355485018 | 3.086961 | 0.002022 | 0.013935651 | ENSG00000171385 | KCND3 | 3752 |
| ENSG00000166165.13 | 90.42403575 | 1.694143895 | 0.250190289 | 4.373247 | 1.22E-05 | 0.000127748 | ENSG00000166165 | CKB | 1152 |
| ENSG00000182175.14 | 73.29585066 | 1.693438766 | 0.306458029 | 3.567989 | 0.00036 | 0.002932268 | ENSG00000182175 | RGMA | 56963 |
| ENSG00000204386.10 | 751.2969924 | 1.693240465 | 0.191155021 | 5.71913 | 1.07E-08 | 1.65E-07 | ENSG00000204386 | NEU1 | 4758 |
| ENSG00000177628.16 | 1531.611457 | 1.690965473 | 0.149714957 | 7.28695 | 3.17E-13 | 7.56E-12 | ENSG00000177628 | GBA | 2629 |
| ENSG00000008735.14 | 75.17847 | 1.690854 | 0.300742 | 3.627209 | 0.000287 | 0.00237757 | ENSG00000008735 | MAPK8IP2 | 23542 |
| ENSG00000273218.1 | 37.83406381 | 1.689225672 | 0.259885458 | 4.191176 | 2.78E-05 | 0.000273121 | ENSG00000273218 | NA | NA |
| ENSG00000243927.6 | 3037.186 | 1.688988 | 0.1216082 | 8.954887 | 3.40E-19 | 1.24E-17 | ENSG00000243927 | MRPS6 | 64968 |
| ENSG00000150782.12 | 6200.238757 | 1.688217905 | 0.136482764 | 7.973299 | 1.54E-15 | 4.43E-14 | ENSG00000150782 | IL18 | 3606 |
| ENSG00000250072.6 | 167.6903803 | 1.687730554 | 0.179229565 | 6.068924 | 1.29E-09 | 2.19E-08 | ENSG00000250072 | SH3TC2-D | 255187 |
| ENSG00000258526.7 | 45.25385417 | 1.686489593 | 0.227334606 | 4.779253 | 1.76E-06 | 2.05E-05 | ENSG00000258526 | NA | NA |
| ENSG00000162783.10 | 11435.45225 | 1.686207537 | 0.153184312 | 7.090854 | 1.33E-12 | 3.01E-11 | ENSG00000162783 | IER5 | 51278 |
| ENSG00000119411.11 | 334.693207 | 1.68537491 | 0.301294817 | 3.602368 | 0.000315 | 0.002589372 | ENSG00000119411 | BSPRY | 54836 |

| | | | | | | | | | |
|---|---|---|---|---|---|---|---|---|---|
| ENSG00000269899.1 | 16.4520722 | 1.684322467 | 0.399398725 | 2.714887 | 0.00663 | 0.040104405 | ENSG00000269899 | NA | NA |
| ENSG00000172382.10 | 64.21903479 | 1.680808195 | 0.334287633 | 3.233168 | 0.001224 | 0.008892794 | ENSG00000172382 | PRSS27 | 83886 |
| ENSG00000117472.10 | 4336.751825 | 1.678829451 | 0.211849176 | 5.092441 | 3.53E-07 | 4.53E-06 | ENSG00000117472 | TSPAN1 | 10103 |
| ENSG00000237686.7 | 30.65408902 | 1.678553328 | 0.24682366 | 4.369732 | 1.24E-05 | 0.00012972 | ENSG00000237686 | LOC10192 | 1.02E+08 |
| ENSG00000173801.17 | 20967.94023 | 1.678268464 | 0.194508674 | 5.54355 | 2.96E-08 | 4.35E-07 | ENSG00000173801 | JUP | 3728 |
| ENSG00000164683.17 | 49.79250024 | 1.677635516 | 0.250730357 | 4.297986 | 1.72E-05 | 0.000175133 | ENSG00000164683 | HEY1 | 23462 |
| ENSG00000273345.5 | 43.17377049 | 1.676763562 | 0.28567376 | 3.769207 | 0.000164 | 0.001421521 | ENSG00000273345 | NA | NA |
| ENSG00000255521.2 | 28.04657952 | 1.676564553 | 0.311297146 | 3.458318 | 0.000544 | 0.004273666 | ENSG00000255521 | LOC10050 | 1.01E+08 |
| ENSG00000198814.12 | 262.7754 | 1.675318 | 0.273621 | 3.929955 | 8.50E-05 | 0.0007762 | ENSG00000198814 | GK | 2710 |
| ENSG00000136536.15 | 9529.945571 | 1.673464375 | 0.089819513 | 11.95135 | 6.39E-33 | 4.96E-31 | ENSG00000136536 | MARCHF7 | 64844 |
| ENSG00000249484.9 | 17.62740888 | 1.672362959 | 0.33433544 | 3.207446 | 0.001339 | 0.009635658 | ENSG00000249484 | LINC0147( | 1.02E+08 |
| ENSG00000179361.17 | 929.9739229 | 1.669286595 | 0.212946158 | 5.021394 | 5.13E-07 | 6.44E-06 | ENSG00000179361 | ARID3B | 10620 |
| ENSG00000011028.14 | 1206.308162 | 1.668221252 | 0.086143764 | 12.40045 | 2.60E-35 | 2.18E-33 | ENSG00000011028 | MRC2 | 9902 |
| ENSG00000214309.5 | 48.41439122 | 1.668084715 | 0.223836386 | 4.771721 | 1.83E-06 | 2.13E-05 | ENSG00000214309 | MBLAC1 | 255374 |
| ENSG00000111540.16 | 6728.634288 | 1.666039265 | 0.115653143 | 9.217555 | 3.04E-20 | 1.18E-18 | ENSG00000111540 | RAB5B | 5869 |
| ENSG00000175416.13 | 24329.8366 | 1.665758738 | 0.169847047 | 6.274815 | 3.50E-10 | 6.39E-09 | ENSG00000175416 | CLTB | 1212 |
| ENSG00000188051.7 | 34.55831 | 1.665213 | 0.3254264 | 3.273283 | 0.001063 | 0.007837368 | ENSG00000188051 | TMEM221 | 1E+08 |
| ENSG00000196208.14 | 67.10119941 | 1.663607319 | 0.370356625 | 2.871846 | 0.004081 | 0.026152964 | ENSG00000196208 | GREB1 | 9687 |
| ENSG00000135643.5 | 15.10787768 | 1.663543469 | 0.331416769 | 3.209082 | 0.001332 | 0.009591388 | ENSG00000135643 | KCNMB4 | 27345 |
| ENSG00000253846.2 | 309.491841 | 1.663380802 | 0.196862208 | 5.40165 | 6.60E-08 | 9.27E-07 | ENSG00000253846 | PCDHGA1 | 56106 |
| ENSG00000255823.4 | 30.61345336 | 1.663243788 | 0.354438091 | 2.999801 | 0.002702 | 0.018125084 | ENSG00000255823 | MTRNR2L | 1E+08 |
| ENSG00000003402.20 | 6079.474703 | 1.662347269 | 0.181593272 | 5.850147 | 4.91E-09 | 7.84E-08 | ENSG00000003402 | CFLAR | 8837 |
| ENSG00000064393.16 | 3380.936918 | 1.662245402 | 0.163315367 | 6.504259 | 7.81E-11 | 1.50E-09 | ENSG00000064393 | HIPK2 | 28996 |
| ENSG00000132357.14 | 1339.283098 | 1.662183508 | 0.111609027 | 9.517004 | 1.78E-21 | 7.43E-20 | ENSG00000132357 | CARD6 | 84674 |
| ENSG00000107372.13 | 6253.267613 | 1.661647166 | 0.1238397 | 8.572753 | 1.01E-17 | 3.35E-16 | ENSG00000107372 | ZFAND5 | 7763 |
| ENSG00000198853.12 | 717.4625229 | 1.660663157 | 0.222798599 | 4.760637 | 1.93E-06 | 2.24E-05 | ENSG00000198853 | RUSC2 | 9853 |
| ENSG00000272081.1 | 11.87878516 | 1.659677634 | 0.372830224 | 2.842252 | 0.00448 | 0.028421331 | ENSG00000272081 | NA | NA |
| ENSG00000196428.12 | 3545.396154 | 1.658864563 | 0.18556683 | 5.706109 | 1.16E-08 | 1.77E-07 | ENSG00000196428 | TSC22D2 | 9819 |
| ENSG00000135253.13 | 104.341224 | 1.65881569 | 0.273437934 | 3.872234 | 0.000108 | 0.000968943 | ENSG00000135253 | KCP | 375616 |
| ENSG00000180152.3 | 17.49777222 | 1.658458546 | 0.372450614 | 2.841876 | 0.004485 | 0.028442698 | ENSG00000180152 | XIAPP3 | 1E+08 |
| ENSG00000281406.2 | 175.8394276 | 1.658000254 | 0.207209789 | 5.105938 | 3.29E-07 | 4.24E-06 | ENSG00000281406 | NA | NA |
| ENSG00000106089.12 | 822.6462332 | 1.657546672 | 0.092337176 | 11.4531 | 2.27E-30 | 1.58E-28 | ENSG00000106089 | STX1A | 6804 |
| ENSG00000153006.16 | 9116.977568 | 1.656765554 | 0.152237081 | 6.941578 | 3.88E-12 | 8.43E-11 | ENSG00000153006 | SREK1IP1 | 285672 |
| ENSG00000137819.13 | 1985.381097 | 1.654967028 | 0.084787396 | 12.4425 | 1.54E-35 | 1.29E-33 | ENSG00000137819 | PAQR5 | 54852 |
| ENSG00000241322.10 | 34.49219038 | 1.651977812 | 0.240689036 | 4.370693 | 1.24E-05 | 0.000129201 | ENSG00000241322 | CDRT1 | 374286 |
| ENSG00000287277.1 | 50.03611295 | 1.651512941 | 0.381862434 | 2.753643 | 0.005894 | 0.036126679 | ENSG00000287277 | NA | NA |
| ENSG00000120327.6 | 88.79044434 | 1.650149022 | 0.156875375 | 6.694161 | 2.17E-11 | 4.40E-10 | ENSG00000120327 | PCDHB14 | 56122 |
| ENSG00000119915.5 | 18.98507999 | 1.649117322 | 0.32989352 | 3.18017 | 0.001472 | 0.010477174 | ENSG00000119915 | ELOVL3 | 83401 |
| ENSG00000084110.11 | 62.31369322 | 1.648007827 | 0.220572555 | 4.751307 | 2.02E-06 | 2.34E-05 | ENSG00000084110 | HAL | 3034 |
| ENSG00000164411.11 | 140.0219045 | 1.647434305 | 0.11336688 | 9.239333 | 2.48E-20 | 9.70E-19 | ENSG00000164411 | GJB7 | 375519 |
| ENSG00000102996.5 | 913.7808026 | 1.646465481 | 0.341334509 | 3.065806 | 0.002171 | 0.014887192 | ENSG00000102996 | MMP15 | 4324 |
| ENSG00000150593.18 | 2531.605612 | 1.643479089 | 0.100512208 | 10.38162 | 3.01E-25 | 1.55E-23 | ENSG00000150593 | PDCD4 | 27250 |

| | | | | | | | | | |
|---|---|---|---|---|---|---|---|---|---|
| ENSG00000067715.14 | 278.381487 | 1.641913124 | 0.148043918 | 7.037865 | 1.95E-12 | 4.36E-11 | ENSG00000067715 | SYT1 | 6857 |
| ENSG00000172985.10 | 112.7983494 | 1.641410508 | 0.245930096 | 4.234579 | 2.29E-05 | 0.000228482 | ENSG00000172985 | SH3RF3 | 344558 |
| ENSG00000186591.12 | 10516.83478 | 1.640890964 | 0.200142087 | 5.20076 | 1.98E-07 | 2.63E-06 | ENSG00000186591 | UBE2H | 7328 |
| ENSG00000185262.9 | 3229.060253 | 1.638997387 | 0.187279357 | 5.547848 | 2.89E-08 | 4.25E-07 | ENSG00000185262 | UBALD2 | 283991 |
| ENSG00000149972.11 | 131.4557356 | 1.638837807 | 0.176721244 | 5.878398 | 4.14E-09 | 6.66E-08 | ENSG00000149972 | CNTN5 | 53942 |
| ENSG00000176973.8 | 113.9728569 | 1.637203262 | 0.171914594 | 6.033247 | 1.61E-09 | 2.70E-08 | ENSG00000176973 | FAM89B | 23625 |
| ENSG00000099377.14 | 84.73814074 | 1.636871932 | 0.29102236 | 3.56286 | 0.000367 | 0.002982523 | ENSG00000099377 | HSD3B7 | 80270 |
| ENSG00000119714.11 | 144.0747306 | 1.636713648 | 0.170573551 | 6.07781 | 1.22E-09 | 2.07E-08 | ENSG00000119714 | GPR68 | 8111 |
| ENSG00000136854.21 | 1522.628467 | 1.635285154 | 0.12173952 | 8.504101 | 1.83E-17 | 5.97E-16 | ENSG00000136854 | STXBP1 | 6812 |
| ENSG00000189045.13 | 44.43858121 | 1.634613757 | 0.230838559 | 4.48198 | 7.40E-06 | 8.01E-05 | ENSG00000189045 | ANKDD1B | 728780 |
| ENSG00000275880.1 | 56.73736481 | 1.634461705 | 0.209441667 | 4.93914 | 7.85E-07 | 9.64E-06 | ENSG00000275880 | NA | NA |
| ENSG00000171365.17 | 1339.485 | 1.633575 | 0.1723823 | 5.99583 | 2.02E-09 | 3.36E-08 | ENSG00000171365 | CLCN5 | 1184 |
| ENSG00000133112.16 | 113525.083 | 1.632175123 | 0.173868777 | 5.936518 | 2.91E-09 | 4.77E-08 | ENSG00000133112 | TPT1 | 7178 |
| ENSG00000127220.6 | 233.3052 | 1.628336 | 0.2402066 | 4.28105 | 1.86E-05 | 0.000188148 | ENSG00000127220 | ABHD8 | 79575 |
| ENSG00000119321.9 | 5879.636785 | 1.626642284 | 0.126446194 | 8.119203 | 4.69E-16 | 1.38E-14 | ENSG00000119321 | FKBP15 | 23307 |
| ENSG00000287906.1 | 61.13919916 | 1.624615387 | 0.255948002 | 4.003217 | 6.25E-05 | 0.000582264 | ENSG00000287906 | NA | NA |
| ENSG00000214318.3 | 18.97075803 | 1.620882311 | 0.339701378 | 3.005235 | 0.002654 | 0.017840514 | ENSG00000214318 | NA | NA |
| ENSG00000145685.14 | 2935.269677 | 1.619432693 | 0.071843288 | 14.18967 | 1.06E-45 | 1.32E-43 | ENSG00000145685 | LHFPL2 | 10184 |
| ENSG00000263155.6 | 181.7389622 | 1.616902319 | 0.159296304 | 6.383716 | 1.73E-10 | 3.24E-09 | ENSG00000263155 | MYZAP | 1.01E+08 |
| ENSG00000255112.2 | 2689.475678 | 1.615737077 | 0.172421309 | 5.891018 | 3.84E-09 | 6.20E-08 | ENSG00000255112 | CHMP1B | 57132 |
| ENSG00000131730.16 | 88.93379773 | 1.614452291 | 0.249574686 | 4.064724 | 4.81E-05 | 0.000456737 | ENSG00000131730 | CKMT2 | 1160 |
| ENSG00000143845.15 | 218.9517955 | 1.613651598 | 0.32505944 | 3.118358 | 0.001819 | 0.012677479 | ENSG00000143845 | ETNK2 | 55224 |
| ENSG00000114378.17 | 283.6346851 | 1.613515252 | 0.137114709 | 7.391733 | 1.45E-13 | 3.57E-12 | ENSG00000114378 | HYAL1 | 3373 |
| ENSG00000120738.8 | 6882.571724 | 1.610254778 | 0.251219254 | 4.021407 | 5.79E-05 | 0.00054182 | ENSG00000120738 | EGR1 | 1958 |
| ENSG00000266869.2 | 72.15478784 | 1.60830807 | 0.177690468 | 5.67452 | 1.39E-08 | 2.11E-07 | ENSG00000266869 | NA | NA |
| ENSG00000157954.15 | 4898.610226 | 1.607634314 | 0.128056469 | 7.868672 | 3.58E-15 | 9.97E-14 | ENSG00000157954 | WIPI2 | 26100 |
| ENSG00000272078.1 | 21.96317798 | 1.607038692 | 0.339054942 | 2.970134 | 0.002977 | 0.019757071 | ENSG00000272078 | NA | NA |
| ENSG00000270681.1 | 41.54915852 | 1.605857615 | 0.295296809 | 3.40626 | 0.000659 | 0.005085011 | ENSG00000270681 | NA | NA |
| ENSG00000198513.11 | 105.9533156 | 1.603743446 | 0.165587374 | 6.061715 | 1.35E-09 | 2.28E-08 | ENSG00000198513 | ATL1 | 51062 |
| ENSG00000147251.15 | 316.822 | 1.603729 | 0.1266854 | 7.923003 | 2.32E-15 | 6.55E-14 | ENSG00000147251 | DOCK11 | 139818 |
| ENSG00000164733.21 | 31511.89941 | 1.603137479 | 0.136984019 | 7.323026 | 2.42E-13 | 5.85E-12 | ENSG00000164733 | CTSB | 1508 |
| ENSG00000104419.14 | 12051.64348 | 1.601820301 | 0.17965191 | 5.576452 | 2.45E-08 | 3.64E-07 | ENSG00000104419 | NDRG1 | 10397 |
| ENSG00000160211.18 | 2272.159 | 1.600545 | 0.06780124 | 14.75703 | 2.77E-49 | 3.84E-47 | ENSG00000160211 | G6PD | 2539 |
| ENSG00000158793.14 | 1273.856628 | 1.600022135 | 0.149778795 | 6.67666 | 2.44E-11 | 4.93E-10 | ENSG00000158793 | NIT1 | 4817 |
| ENSG00000267750.6 | 24.3548495 | 1.599602377 | 0.298808865 | 3.34529 | 0.000822 | 0.006227193 | ENSG00000267750 | NA | NA |
| ENSG00000266094.7 | 1203.111253 | 1.598707713 | 0.188607644 | 5.29516 | 1.19E-07 | 1.63E-06 | ENSG00000266094 | RASSF5 | 83593 |
| ENSG00000074410.14 | 175.8489122 | 1.597466111 | 0.360107326 | 2.769913 | 0.005607 | 0.0345859 | ENSG00000074410 | CA12 | 771 |
| ENSG00000266714.9 | 185.4478855 | 1.596912834 | 0.236824826 | 4.209495 | 2.56E-05 | 0.000253203 | ENSG00000266714 | MYO15B | 80022 |
| ENSG00000197580.11 | 40.27210581 | 1.593669284 | 0.261428584 | 3.800921 | 0.000144 | 0.001262024 | ENSG00000197580 | BCO2 | 83875 |
| ENSG00000267510.1 | 16.30952282 | 1.592467241 | 0.297045385 | 3.34113 | 0.000834 | 0.006312277 | ENSG00000267510 | NA | NA |
| ENSG00000197747.9 | 19255.39104 | 1.591324685 | 0.113290936 | 8.750256 | 2.13E-18 | 7.35E-17 | ENSG00000197747 | S100A10 | 6281 |
| ENSG00000090013.10 | 1374.296 | 1.587948 | 0.2022535 | 4.884699 | 1.04E-06 | 1.25E-05 | ENSG00000090013 | BLVRB | 645 |

| | | | | | | | | | |
|---|---|---|---|---|---|---|---|---|---|
| ENSG00000108828.16 | 10497.17716 | 1.587820688 | 0.119641537 | 8.256503 | 1.50E-16 | 4.57E-15 | ENSG00000108828 | VAT1 | 10493 |
| ENSG00000087266.17 | 1791.79673 | 1.584154881 | 0.118441096 | 8.309235 | 9.63E-17 | 2.98E-15 | ENSG00000087266 | SH3BP2 | 6452 |
| ENSG00000104722.14 | 388.9719923 | 1.583874511 | 0.13800332 | 7.129354 | 1.01E-12 | 2.31E-11 | ENSG00000104722 | NEFM | 4741 |
| ENSG00000259240.1 | 19.81989872 | 1.582939469 | 0.305902623 | 3.213243 | 0.001312 | 0.009473995 | ENSG00000259240 | MIR4713H | 1.1E+08 |
| ENSG00000168300.14 | 3388.41343 | 1.580805279 | 0.158427919 | 6.190861 | 5.98E-10 | 1.06E-08 | ENSG00000168300 | PCMTD1 | 115294 |
| ENSG00000143434.15 | 170.1452022 | 1.579497652 | 0.152610413 | 6.418288 | 1.38E-10 | 2.60E-09 | ENSG00000143434 | SEMA6C | 10500 |
| ENSG00000123091.5 | 6136.270501 | 1.579240053 | 0.157180552 | 6.230033 | 4.66E-10 | 8.37E-09 | ENSG00000123091 | RNF11 | 26994 |
| ENSG00000250602.6 | 42.9871639 | 1.578021888 | 0.243012735 | 4.024571 | 5.71E-05 | 0.000535625 | ENSG00000250602 | NA | NA |
| ENSG00000143514.17 | 4328.652294 | 1.577357639 | 0.146516678 | 6.670624 | 2.55E-11 | 5.14E-10 | ENSG00000143514 | TP53BP2 | 7159 |
| ENSG00000079435.10 | 159.8064 | 1.577341 | 0.1963295 | 4.978066 | 6.42E-07 | 7.98E-06 | ENSG00000079435 | LIPE | 3991 |
| ENSG00000121005.9 | 124.9724794 | 1.575174542 | 0.144259819 | 6.759849 | 1.38E-11 | 2.86E-10 | ENSG00000121005 | CRISPLD1 | 83690 |
| ENSG00000181458.10 | 138.0880217 | 1.572191609 | 0.155137935 | 6.266627 | 3.69E-10 | 6.69E-09 | ENSG00000181458 | TMEM45A | 55076 |
| ENSG00000152380.10 | 186.9498423 | 1.57193504 | 0.192318872 | 5.053768 | 4.33E-07 | 5.48E-06 | ENSG00000152380 | FAM151B | 167555 |
| ENSG00000240522.1 | 20.16538 | 1.57 | 0.3026909 | 3.204589 | 0.001353 | 0.009716073 | ENSG00000240522 | RPL7AP1( | 161406 |
| ENSG00000265298.1 | 103.5187019 | 1.569797026 | 0.234179792 | 4.14125 | 3.45E-05 | 0.000335611 | ENSG00000265298 | LOC10192 | 1.02E+08 |
| ENSG00000179397.18 | 31.53004057 | 1.568302645 | 0.255934564 | 3.783399 | 0.000155 | 0.001348984 | ENSG00000179397 | CATSPER | 257044 |
| ENSG00000188396.3 | 23.57945287 | 1.568072007 | 0.280203882 | 3.454884 | 0.000551 | 0.004323357 | ENSG00000188396 | TCTEX1D4 | 343521 |
| ENSG00000092148.13 | 19156.76523 | 1.566195018 | 0.159556022 | 6.055522 | 1.40E-09 | 2.37E-08 | ENSG00000092148 | HECTD1 | 25831 |
| ENSG00000251026.2 | 39.7456089 | 1.564905277 | 0.237564908 | 4.061649 | 4.87E-05 | 0.000461972 | ENSG00000251026 | NA | NA |
| ENSG00000172940.12 | 18.33981028 | 1.564178029 | 0.331892195 | 2.905094 | 0.003671 | 0.023752311 | ENSG00000172940 | SLC22A13 | 9390 |
| ENSG00000119632.4 | 658.1524444 | 1.563783168 | 0.174604769 | 5.519799 | 3.39E-08 | 4.93E-07 | ENSG00000119632 | IFI27L2 | 83982 |
| ENSG00000102882.12 | 2049.243008 | 1.563074203 | 0.168546219 | 5.714007 | 1.10E-08 | 1.70E-07 | ENSG00000102882 | MAPK3 | 5595 |
| ENSG00000121552.4 | 10491.92378 | 1.562218355 | 0.207397621 | 4.639486 | 3.49E-06 | 3.92E-05 | ENSG00000121552 | CSTA | 1475 |
| ENSG00000112782.17 | 17.48726723 | 1.561739337 | 0.350353499 | 2.745054 | 0.00605 | 0.036924695 | ENSG00000112782 | CLIC5 | 53405 |
| ENSG00000205420.11 | 161140.934 | 1.561365703 | 0.192527709 | 4.993389 | 5.93E-07 | 7.39E-06 | ENSG00000205420 | KRT6A | 3853 |
| ENSG00000165169.11 | 2214.606 | 1.55931 | 0.1374671 | 6.97847 | 2.98E-12 | 6.54E-11 | ENSG00000165169 | DYNLT3 | 6990 |
| ENSG00000141449.14 | 28.68487984 | 1.559238632 | 0.333874605 | 2.873051 | 0.004065 | 0.026066792 | ENSG00000141449 | GREB1L | 80000 |
| ENSG00000224939.1 | 33.24384878 | 1.554374752 | 0.268034245 | 3.560645 | 0.00037 | 0.003004479 | ENSG00000224939 | LINC0018 | 1E+08 |
| ENSG00000278709.2 | 28.36904 | 1.554243 | 0.3475792 | 2.745398 | 0.006044 | 0.03689451 | ENSG00000278709 | NKILA | 1.05E+08 |
| ENSG00000180155.20 | 83.60945019 | 1.553417237 | 0.250651627 | 3.803754 | 0.000143 | 0.00124931 | ENSG00000180155 | LYNX1 | 66004 |
| ENSG00000174705.13 | 918.2446069 | 1.553270793 | 0.220630317 | 4.32067 | 1.56E-05 | 0.000159216 | ENSG00000174705 | SH3PXD2I | 285590 |
| ENSG00000271614.1 | 26.09198476 | 1.552410318 | 0.331933284 | 2.869282 | 0.004114 | 0.026346901 | ENSG00000271614 | ATP2B1-A | 338758 |
| ENSG00000171368.12 | 100.6664634 | 1.55208304 | 0.164747031 | 5.77906 | 7.51E-09 | 1.18E-07 | ENSG00000171368 | TPPP | 11076 |
| ENSG00000181634.8 | 3755.550763 | 1.551237569 | 0.105547396 | 9.012421 | 2.02E-19 | 7.44E-18 | ENSG00000181634 | TNFSF15 | 9966 |
| ENSG00000095397.14 | 197.2487205 | 1.549843871 | 0.18498602 | 5.134679 | 2.83E-07 | 3.67E-06 | ENSG00000095397 | WHRN | 25861 |
| ENSG00000237441.10 | 969.2225538 | 1.549305279 | 0.134708954 | 7.047084 | 1.83E-12 | 4.09E-11 | ENSG00000237441 | RGL2 | 5863 |
| ENSG00000176834.14 | 1729.107742 | 1.547025663 | 0.127635936 | 7.419742 | 1.17E-13 | 2.91E-12 | ENSG00000176834 | VSIG10 | 54621 |
| ENSG00000101311.15 | 10368.11 | 1.546833 | 0.1176497 | 8.047899 | 8.42E-16 | 2.45E-14 | ENSG00000101311 | FERMT1 | 55612 |
| ENSG00000113578.18 | 49.87809534 | 1.544888702 | 0.240619523 | 3.9269 | 8.60E-05 | 0.000785855 | ENSG00000113578 | FGF1 | 2246 |
| ENSG00000142227.11 | 2449.556 | 1.541465 | 0.05339709 | 17.63139 | 1.41E-69 | 3.42E-67 | ENSG00000142227 | EMP3 | 2014 |
| ENSG00000139998.15 | 286.1660831 | 1.539268656 | 0.182836266 | 5.137212 | 2.79E-07 | 3.63E-06 | ENSG00000139998 | RAB15 | 376267 |
| ENSG00000176597.12 | 6496.294744 | 1.53885261 | 0.081944766 | 11.45714 | 2.17E-30 | 1.51E-28 | ENSG00000176597 | B3GNT5 | 84002 |

| | | | | | | | | | |
|---|---|---|---|---|---|---|---|---|---|
| ENSG00000107341.5 | 7444.131588 | 1.537692559 | 0.165254857 | 5.674221 | 1.39E-08 | 2.11E-07 | ENSG00000107341 | UBE2R2 | 54926 |
| ENSG00000165006.14 | 4919.32402 | 1.537509898 | 0.084778889 | 11.05829 | 2.00E-28 | 1.25E-26 | ENSG00000165006 | UBAP1 | 51271 |
| ENSG00000119979.18 | 2434.367516 | 1.536753158 | 0.086731707 | 10.80058 | 3.42E-27 | 1.96E-25 | ENSG00000119979 | DENND10 | 404636 |
| ENSG00000139173.10 | 377.7409489 | 1.536578845 | 0.117627775 | 7.962225 | 1.69E-15 | 4.82E-14 | ENSG00000139173 | TMEM117 | 84216 |
| ENSG00000141542.11 | 210.9596896 | 1.536528058 | 0.138509121 | 6.76149 | 1.37E-11 | 2.83E-10 | ENSG00000141542 | RAB40B | 10966 |
| ENSG00000183401.12 | 122.9527674 | 1.535705553 | 0.152248975 | 6.145891 | 7.95E-10 | 1.38E-08 | ENSG00000183401 | CCDC159 | 126075 |
| ENSG00000219891.2 | 527.0181858 | 1.533828596 | 0.287126801 | 3.252321 | 0.001145 | 0.008371783 | ENSG00000219891 | NA | NA |
| ENSG00000161267.12 | 708.1704746 | 1.5295449 | 0.192020422 | 4.840865 | 1.29E-06 | 1.53E-05 | ENSG00000161267 | BDH1 | 622 |
| ENSG00000116661.11 | 3407.040549 | 1.527872058 | 0.08860431 | 10.47209 | 1.16E-25 | 6.17E-24 | ENSG00000116661 | FBXO2 | 26232 |
| ENSG00000104863.12 | 36.37105 | 1.526577 | 0.2411002 | 3.843121 | 0.000121 | 0.001079449 | ENSG00000104863 | LIN7B | 64130 |
| ENSG00000196597.12 | 336.098416 | 1.52445287 | 0.193530606 | 4.776779 | 1.78E-06 | 2.08E-05 | ENSG00000196597 | ZNF782 | 158431 |
| ENSG00000143819.12 | 526.1882196 | 1.523976776 | 0.134871366 | 6.8508 | 7.34E-12 | 1.56E-10 | ENSG00000143819 | EPHX1 | 2052 |
| ENSG00000254122.2 | 28.65370913 | 1.523530338 | 0.319311823 | 2.892252 | 0.003825 | 0.024619603 | ENSG00000254122 | PCDHGB7 | 56099 |
| ENSG00000079459.13 | 6173.952156 | 1.523336127 | 0.094700772 | 9.750038 | 1.84E-22 | 8.05E-21 | ENSG00000079459 | FDFT1 | 2222 |
| ENSG00000143786.8 | 36.95573481 | 1.521882142 | 0.309775552 | 2.975968 | 0.002921 | 0.019409292 | ENSG00000143786 | CNIH3 | 149111 |
| ENSG00000255145.3 | 16.89370206 | 1.521860129 | 0.31066257 | 2.9674 | 0.003003 | 0.019908821 | ENSG00000255145 | NA | NA |
| ENSG00000147174.11 | 113.0729 | 1.521749 | 0.1815153 | 5.078076 | 3.81E-07 | 4.86E-06 | ENSG00000147174 | GCNA | 93953 |
| ENSG00000169174.10 | 1175.944452 | 1.52157804 | 0.100992019 | 9.125256 | 7.16E-20 | 2.70E-18 | ENSG00000169174 | PCSK9 | 255738 |
| ENSG00000106460.19 | 4022.427519 | 1.521488651 | 0.082278058 | 11.19969 | 4.09E-29 | 2.69E-27 | ENSG00000106460 | TMEM106I | 54664 |
| ENSG00000162944.10 | 56.7806292 | 1.521000174 | 0.261449895 | 3.522664 | 0.000427 | 0.003424394 | ENSG00000162944 | RFTN2 | 130132 |
| ENSG00000275894.1 | 43.03158 | 1.517757 | 0.2654589 | 3.457248 | 0.000546 | 0.004288138 | ENSG00000275894 | NA | NA |
| ENSG00000160888.7 | 2628.120466 | 1.517737531 | 0.198853688 | 4.61514 | 3.93E-06 | 4.39E-05 | ENSG00000160888 | IER2 | 9592 |
| ENSG00000142661.19 | 275.4216408 | 1.517441964 | 0.171571026 | 5.347301 | 8.93E-08 | 1.24E-06 | ENSG00000142661 | MYOM3 | 127294 |
| ENSG00000214456.8 | 26.90527548 | 1.517385683 | 0.302534221 | 3.032337 | 0.002427 | 0.01646359 | ENSG00000214456 | PLIN5 | 440503 |
| ENSG00000178531.6 | 110.3667459 | 1.515201654 | 0.172198037 | 5.31482 | 1.07E-07 | 1.47E-06 | ENSG00000178531 | CTXN1 | 404217 |
| ENSG00000142669.15 | 4428.523828 | 1.513898599 | 0.070769199 | 12.91379 | 3.76E-38 | 3.50E-36 | ENSG00000142669 | SH3BGRL: | 83442 |
| ENSG00000120324.8 | 28.02892795 | 1.513512048 | 0.305349695 | 2.991691 | 0.002774 | 0.018562244 | ENSG00000120324 | PCDHB10 | 56126 |
| ENSG00000130164.13 | 18773.16864 | 1.512762902 | 0.11116442 | 8.210927 | 2.19E-16 | 6.60E-15 | ENSG00000130164 | LDLR | 3949 |
| ENSG00000139636.15 | 896.3455606 | 1.512716831 | 0.117275403 | 7.782679 | 7.10E-15 | 1.92E-13 | ENSG00000139636 | LMBR1L | 55716 |
| ENSG00000274512.5 | 50.60223091 | 1.512701473 | 0.184720625 | 4.940983 | 7.77E-07 | 9.56E-06 | ENSG00000274512 | TBC1D3 | 729873 |
| ENSG00000180481.11 | 17.12940825 | 1.511942774 | 0.334417454 | 2.726959 | 0.006392 | 0.038771998 | ENSG00000180481 | GLIPR1L2 | 144321 |
| ENSG00000099194.6 | 55792.76984 | 1.511445034 | 0.060504814 | 15.06401 | 2.79E-51 | 4.06E-49 | ENSG00000099194 | SCD | 6319 |
| ENSG00000273350.1 | 25.3967 | 1.510727 | 0.3122568 | 2.916596 | 0.003539 | 0.02299033 | ENSG00000273350 | NA | NA |
| ENSG00000167656.5 | 1165.370641 | 1.510389486 | 0.334993201 | 2.717636 | 0.006575 | 0.039799942 | ENSG00000167656 | LY6D | 8581 |
| ENSG00000076258.10 | 69.84316223 | 1.508818982 | 0.259172047 | 3.506624 | 0.000454 | 0.003618404 | ENSG00000076258 | FMO4 | 2329 |
| ENSG00000104549.12 | 8454.95571 | 1.507856789 | 0.112636418 | 8.060064 | 7.63E-16 | 2.22E-14 | ENSG00000104549 | SQLE | 6713 |
| ENSG00000163788.14 | 2339.201872 | 1.50731176 | 0.195092929 | 4.650665 | 3.31E-06 | 3.72E-05 | ENSG00000163788 | SNRK | 54861 |
| ENSG00000268858.2 | 481.11 | 1.506218 | 0.1812781 | 4.999046 | 5.76E-07 | 7.19E-06 | ENSG00000268858 | LOC11226 | 1.12E+08 |
| ENSG00000249673.7 | 1028.849678 | 1.504971425 | 0.173777829 | 5.207635 | 1.91E-07 | 2.54E-06 | ENSG00000249673 | NA | NA |
| ENSG00000120437.8 | 1573.414412 | 1.503804449 | 0.086781779 | 10.41468 | 2.13E-25 | 1.11E-23 | ENSG00000120437 | ACAT2 | 39 |
| ENSG00000171631.14 | 53.8598499 | 1.499877306 | 0.251830068 | 3.573351 | 0.000352 | 0.002875481 | ENSG00000171631 | P2RY6 | 5031 |
| ENSG00000132823.11 | 3537.234 | 1.499864 | 0.173402 | 5.189467 | 2.11E-07 | 2.78E-06 | ENSG00000132823 | OSER1 | 51526 |

| | | | | | | | | | |
|---|---|---|---|---|---|---|---|---|---|
| ENSG00000185010.15 | 233.8745 | 1.499349 | 0.2270479 | 3.961056 | 7.46E-05 | 0.000687848 | ENSG00000185010 | F8 | 2157 |
| ENSG00000138944.8 | 41.05357 | 1.496705 | 0.2610967 | 3.434378 | 0.000594 | 0.00463264 | ENSG00000138944 | SHISAL1 | 85352 |
| ENSG00000198598.6 | 140.8389836 | 1.49606252 | 0.147875174 | 6.059587 | 1.36E-09 | 2.31E-08 | ENSG00000198598 | MMP17 | 4326 |
| ENSG00000163346.17 | 739.5523794 | 1.49277987 | 0.149017883 | 5.991092 | 2.08E-09 | 3.46E-08 | ENSG00000163346 | PBXIP1 | 57326 |
| ENSG00000137177.20 | 7226.920788 | 1.491665687 | 0.168982949 | 5.276661 | 1.32E-07 | 1.79E-06 | ENSG00000137177 | KIF13A | 63971 |
| ENSG00000237978.6 | 66.23210842 | 1.49045951 | 0.216300755 | 4.116766 | 3.84E-05 | 0.000371419 | ENSG00000237978 | NA | NA |
| ENSG00000047365.12 | 3278.52231 | 1.489999901 | 0.123673634 | 7.196359 | 6.18E-13 | 1.44E-11 | ENSG00000047365 | ARAP2 | 116984 |
| ENSG00000253982.2 | 68.53000776 | 1.489104547 | 0.174221683 | 5.103294 | 3.34E-07 | 4.29E-06 | ENSG00000253982 | LOC10192 | 1.02E+08 |
| ENSG00000143797.12 | 7778.232179 | 1.488992149 | 0.096651018 | 9.19796 | 3.65E-20 | 1.41E-18 | ENSG00000143797 | MBOAT2 | 129642 |
| ENSG00000166946.14 | 2934.400371 | 1.485575761 | 0.115443949 | 7.671045 | 1.71E-14 | 4.48E-13 | ENSG00000166946 | CCNDBP1 | 23582 |
| ENSG00000241288.8 | 79.59182324 | 1.484773674 | 0.192359771 | 4.599577 | 4.23E-06 | 4.70E-05 | ENSG00000241288 | LINC0261 | 1.02E+08 |
| ENSG00000241889.1 | 41.01376987 | 1.482496936 | 0.275746282 | 3.200395 | 0.001372 | 0.009845299 | ENSG00000241889 | NA | NA |
| ENSG00000187957.8 | 301.4194121 | 1.481093224 | 0.134891003 | 6.53189 | 6.49E-11 | 1.26E-09 | ENSG00000187957 | DNER | 92737 |
| ENSG00000127507.18 | 865.1989541 | 1.480646229 | 0.147851882 | 5.956273 | 2.58E-09 | 4.26E-08 | ENSG00000127507 | ADGRE2 | 30817 |
| ENSG00000141524.15 | 1085.606628 | 1.480530816 | 0.229811985 | 3.831527 | 0.000127 | 0.001127852 | ENSG00000141524 | TMC6 | 11322 |
| ENSG00000277310.1 | 19.70956746 | 1.480255252 | 0.330995352 | 2.659419 | 0.007828 | 0.046359357 | ENSG00000277310 | NA | NA |
| ENSG00000249476.2 | 42.08412054 | 1.480225882 | 0.249836714 | 3.523205 | 0.000426 | 0.003418858 | ENSG00000249476 | LOC28563 | 285638 |
| ENSG00000143933.18 | 40080.04942 | 1.476495695 | 0.190863798 | 4.592257 | 4.38E-06 | 4.87E-05 | ENSG00000143933 | CALM2 | 805 |
| ENSG00000144802.11 | 2013.587314 | 1.475457807 | 0.204609932 | 4.278667 | 1.88E-05 | 0.000189812 | ENSG00000144802 | NFKBIZ | 64332 |
| ENSG00000262655.4 | 162.5636336 | 1.474613245 | 0.200742138 | 4.356899 | 1.32E-05 | 0.000136864 | ENSG00000262655 | SPON1 | 10418 |
| ENSG00000135241.17 | 1975.196928 | 1.473327895 | 0.127642271 | 6.841996 | 7.81E-12 | 1.66E-10 | ENSG00000135241 | PNPLA8 | 50640 |
| ENSG00000001630.17 | 423.4096584 | 1.472709107 | 0.127316245 | 6.854656 | 7.15E-12 | 1.52E-10 | ENSG00000001630 | CYP51A1 | 1595 |
| ENSG00000171223.6 | 7279.257541 | 1.471966103 | 0.208472558 | 4.182642 | 2.88E-05 | 0.000282954 | ENSG00000171223 | JUNB | 3726 |
| ENSG00000178585.15 | 2215.788289 | 1.471393254 | 0.172266597 | 5.058399 | 4.23E-07 | 5.36E-06 | ENSG00000178585 | CTNNBIP1 | 56998 |
| ENSG00000273270.1 | 117.3378427 | 1.469813702 | 0.204830184 | 4.246511 | 2.17E-05 | 0.000217468 | ENSG00000273270 | NA | NA |
| ENSG00000178381.11 | 688.6278887 | 1.468578575 | 0.121363921 | 7.15681 | 8.26E-13 | 1.91E-11 | ENSG00000178381 | ZFAND2A | 90637 |
| ENSG00000101928.13 | 809.1203 | 1.46762 | 0.14153 | 6.130291 | 8.77E-10 | 1.51E-08 | ENSG00000101928 | MOSPD1 | 56180 |
| ENSG00000186417.14 | 58.41609078 | 1.466748259 | 0.248711352 | 3.484957 | 0.000492 | 0.003902263 | ENSG00000186417 | GLDN | 342035 |
| ENSG00000126460.11 | 303.9842 | 1.46488 | 0.1827394 | 4.732857 | 2.21E-06 | 2.55E-05 | ENSG00000126460 | PRRG2 | 5639 |
| ENSG00000105520.10 | 740.1139483 | 1.464857829 | 0.124437003 | 6.950166 | 3.65E-12 | 7.96E-11 | ENSG00000105520 | PLPPR2 | 64748 |
| ENSG00000227959.1 | 56.96129333 | 1.463996774 | 0.194203173 | 4.448932 | 8.63E-06 | 9.25E-05 | ENSG00000227959 | NA | NA |
| ENSG00000236347.1 | 22.87614833 | 1.462494093 | 0.326091284 | 2.644947 | 0.00817 | 0.048100788 | ENSG00000236347 | NA | NA |
| ENSG00000139044.11 | 779.3119972 | 1.461877519 | 0.087875724 | 9.807914 | 1.04E-22 | 4.61E-21 | ENSG00000139044 | B4GALNT | 283358 |
| ENSG00000179178.11 | 247.5898676 | 1.461748519 | 0.244825114 | 3.519853 | 0.000432 | 0.00345818 | ENSG00000179178 | TMEM125 | 128218 |
| ENSG00000169193.12 | 252.139155 | 1.461484508 | 0.121074056 | 7.115352 | 1.12E-12 | 2.55E-11 | ENSG00000169193 | CCDC126 | 90693 |
| ENSG00000102547.19 | 882.3513192 | 1.461331314 | 0.09732338 | 8.850199 | 8.74E-19 | 3.10E-17 | ENSG00000102547 | CAB39L | 81617 |
| ENSG00000125637.15 | 3760.449158 | 1.460905805 | 0.107349989 | 8.019617 | 1.06E-15 | 3.06E-14 | ENSG00000125637 | PSD4 | 23550 |
| ENSG00000089041.17 | 30.51947735 | 1.459028942 | 0.303109813 | 2.834052 | 0.004596 | 0.02901117 | ENSG00000089041 | P2RX7 | 5027 |
| ENSG00000109814.12 | 2153.217446 | 1.458935 | 0.144942559 | 5.926037 | 3.10E-09 | 5.07E-08 | ENSG00000109814 | UGDH | 7358 |
| ENSG00000106537.8 | 560.6925781 | 1.458402662 | 0.213833977 | 4.014342 | 5.96E-05 | 0.000556838 | ENSG00000106537 | TSPAN13 | 27075 |
| ENSG00000132196.14 | 643.5696723 | 1.457485029 | 0.082075342 | 10.44754 | 1.50E-25 | 7.91E-24 | ENSG00000132196 | HSD17B7 | 51478 |
| ENSG00000074276.10 | 59.7729101 | 1.457251529 | 0.301518085 | 2.843118 | 0.004467 | 0.028352596 | ENSG00000074276 | CDHR2 | 54825 |

| | | | | | | | | | |
|---|---|---|---|---|---|---|---|---|---|
| ENSG00000137474.22 | 213.50527 | 1.456485141 | 0.144024878 | 5.946786 | 2.73E-09 | 4.50E-08 | ENSG00000137474 | MYO7A | 4647 |
| ENSG00000226824.7 | 55.52280519 | 1.456022638 | 0.212731816 | 4.023952 | 5.72E-05 | 0.00053669 | ENSG00000226824 | NA | NA |
| ENSG00000111653.20 | 625.509495 | 1.454389734 | 0.149871057 | 5.700832 | 1.19E-08 | 1.82E-07 | ENSG00000111653 | ING4 | 51147 |
| ENSG00000114423.22 | 467.4981201 | 1.454124292 | 0.163324915 | 5.229602 | 1.70E-07 | 2.27E-06 | ENSG00000114423 | CBLB | 868 |
| ENSG00000102312.22 | 765.6042 | 1.453828 | 0.1957126 | 4.362662 | 1.28E-05 | 0.000133723 | ENSG00000102312 | PORCN | 64840 |
| ENSG00000164850.15 | 50.07485194 | 1.453141923 | 0.222108143 | 3.841111 | 0.000122 | 0.001087966 | ENSG00000164850 | GPER1 | 2852 |
| ENSG00000166415.15 | 45.04942079 | 1.453082317 | 0.245382487 | 3.476541 | 0.000508 | 0.004014849 | ENSG00000166415 | WDR72 | 256764 |
| ENSG00000260428.3 | 101.5194622 | 1.452516223 | 0.177237349 | 4.810026 | 1.51E-06 | 1.77E-05 | ENSG00000260428 | SCX | 642658 |
| ENSG00000160323.18 | 71.83750424 | 1.452508059 | 0.189932847 | 4.488471 | 7.17E-06 | 7.79E-05 | ENSG00000160323 | ADAMTS1 | 11093 |
| ENSG00000154710.17 | 179.0225455 | 1.449952356 | 0.176298149 | 4.821108 | 1.43E-06 | 1.68E-05 | ENSG00000154710 | RABGEF1 | 27342 |
| ENSG00000169231.13 | 245.6429228 | 1.442846036 | 0.103573172 | 8.137687 | 4.03E-16 | 1.19E-14 | ENSG00000169231 | THBS3 | 7059 |
| ENSG00000232618.2 | 34.6633557 | 1.44227302 | 0.269553833 | 3.124693 | 0.00178 | 0.012456535 | ENSG00000232618 | NA | NA |
| ENSG00000168032.9 | 230.3510124 | 1.440859015 | 0.211140183 | 3.982468 | 6.82E-05 | 0.000631991 | ENSG00000168032 | ENTPD3 | 956 |
| ENSG00000198939.8 | 38.13676369 | 1.440100241 | 0.298639162 | 2.813095 | 0.004907 | 0.030757405 | ENSG00000198939 | ZFP2 | 80108 |
| ENSG00000245975.2 | 30.31664455 | 1.440042928 | 0.291575007 | 2.881053 | 0.003963 | 0.025462395 | ENSG00000245975 | LOC10192 | 1.02E+08 |
| ENSG00000248712.7 | 27.1754712 | 1.440027667 | 0.260425884 | 3.225592 | 0.001257 | 0.009104231 | ENSG00000248712 | CCDC153 | 283152 |
| ENSG00000196705.8 | 1830.201 | 1.438488 | 0.1913192 | 4.382664 | 1.17E-05 | 0.000122636 | ENSG00000196705 | ZNF431 | 170959 |
| ENSG00000113732.9 | 10523.56562 | 1.438125866 | 0.094630857 | 8.856793 | 8.23E-19 | 2.92E-17 | ENSG00000113732 | ATP6V0E1 | 8992 |
| ENSG00000119048.7 | 8063.738382 | 1.437445653 | 0.093508499 | 8.955824 | 3.37E-19 | 1.23E-17 | ENSG00000119048 | UBE2B | 7320 |
| ENSG00000162641.20 | 48.60023807 | 1.435732615 | 0.248292447 | 3.36592 | 0.000763 | 0.005817698 | ENSG00000162641 | AKNAD1 | 254268 |
| ENSG00000138772.13 | 22523.23983 | 1.435651541 | 0.137335532 | 6.084744 | 1.17E-09 | 1.99E-08 | ENSG00000138772 | ANXA3 | 306 |
| ENSG00000217624.2 | 23.07999 | 1.434639 | 0.3015387 | 2.767933 | 0.005641 | 0.03474044 | ENSG00000217624 | NA | NA |
| ENSG00000121104.8 | 227.1168311 | 1.434046882 | 0.136983484 | 6.088667 | 1.14E-09 | 1.94E-08 | ENSG00000121104 | FAM117A | 81558 |
| ENSG00000111596.12 | 6798.701019 | 1.431300043 | 0.135287746 | 6.144681 | 8.01E-10 | 1.39E-08 | ENSG00000111596 | CNOT2 | 4848 |
| ENSG00000068971.14 | 721.7318255 | 1.429965502 | 0.121947138 | 6.805945 | 1.00E-11 | 2.11E-10 | ENSG00000068971 | PPP2R5B | 5526 |
| ENSG00000169946.14 | 93.26571213 | 1.429415127 | 0.245897451 | 3.373012 | 0.000744 | 0.005682924 | ENSG00000169946 | ZFPM2 | 23414 |
| ENSG00000214960.10 | 57.26912591 | 1.428784371 | 0.202121989 | 4.100417 | 4.12E-05 | 0.000395928 | ENSG00000214960 | CRPPA | 729920 |
| ENSG00000115919.15 | 2364.695336 | 1.428379783 | 0.181175723 | 4.572245 | 4.83E-06 | 5.33E-05 | ENSG00000115919 | KYNU | 8942 |
| ENSG00000224152.1 | 34.4359032 | 1.428361886 | 0.244819487 | 3.383562 | 0.000716 | 0.005487927 | ENSG00000224152 | NA | NA |
| ENSG00000236423.6 | 40.7597938 | 1.428056601 | 0.21012988 | 3.940689 | 8.12E-05 | 0.000745084 | ENSG00000236423 | LINC0113 | 1E+08 |
| ENSG00000188994.13 | 3207.947107 | 1.424667999 | 0.121359224 | 6.795264 | 1.08E-11 | 2.26E-10 | ENSG00000188994 | ZNF292 | 23036 |
| ENSG00000136002.18 | 2275.064817 | 1.424646736 | 0.151185458 | 5.454537 | 4.91E-08 | 6.96E-07 | ENSG00000136002 | ARHGEF4 | 50649 |
| ENSG00000215769.8 | 32.58854373 | 1.424112567 | 0.283688189 | 2.904994 | 0.003673 | 0.023754108 | ENSG00000215769 | ARHGAP2 | 1.09E+08 |
| ENSG00000177337.8 | 53.70742339 | 1.422486008 | 0.23413413 | 3.512884 | 0.000443 | 0.003544838 | ENSG00000177337 | NA | NA |
| ENSG00000023287.13 | 9874.036729 | 1.420016838 | 0.104703398 | 7.831807 | 4.81E-15 | 1.32E-13 | ENSG00000023287 | RB1CC1 | 9821 |
| ENSG00000125618.17 | 391.9413945 | 1.418568856 | 0.126480182 | 6.471914 | 9.68E-11 | 1.85E-09 | ENSG00000125618 | PAX8 | 7849 |
| ENSG00000214049.8 | 10033.53 | 1.415616 | 0.1475159 | 5.529002 | 3.22E-08 | 4.70E-07 | ENSG00000214049 | UCA1 | 652995 |
| ENSG00000196605.7 | 238.493835 | 1.415502529 | 0.125123588 | 6.517576 | 7.15E-11 | 1.38E-09 | ENSG00000196605 | ZNF846 | 162993 |
| ENSG00000172216.5 | 3035.428 | 1.414242 | 0.1512717 | 5.382645 | 7.34E-08 | 1.03E-06 | ENSG00000172216 | CEBPB | 1051 |
| ENSG00000242147.2 | 612.73819 | 1.413502072 | 0.07127141 | 11.41414 | 3.55E-30 | 2.45E-28 | ENSG00000242147 | LINC0267 | 1.03E+08 |
| ENSG00000116209.11 | 6016.50467 | 1.413083677 | 0.097977024 | 8.298718 | 1.05E-16 | 3.25E-15 | ENSG00000116209 | TMEM59 | 9528 |
| ENSG00000237356.7 | 41.85967283 | 1.412477597 | 0.228244288 | 3.559684 | 0.000371 | 0.003012732 | ENSG00000237356 | NA | NA |

| | | | | | | | | | |
|---|---|---|---|---|---|---|---|---|---|
| ENSG00000261780.3 | 59.9367015 | 1.411939712 | 0.254043236 | 3.196069 | 0.001393 | 0.009977971 | ENSG00000261780 | LINC0258[illegible] | 1.01E+08 |
| ENSG00000133818.14 | 11153.0368 | 1.411385535 | 0.156973198 | 5.168943 | 2.35E-07 | 3.09E-06 | ENSG00000133818 | RRAS2 | 22800 |
| ENSG00000132824.14 | 9798.793 | 1.40913 | 0.1030629 | 7.850831 | 4.13E-15 | 1.14E-13 | ENSG00000132824 | SERINC3 | 10955 |
| ENSG00000285966.1 | 35.55123351 | 1.40765674 | 0.278430691 | 2.900746 | 0.003723 | 0.024031802 | ENSG00000285966 | NA | NA |
| ENSG00000181019.13 | 13192.23966 | 1.407565912 | 0.077731265 | 10.3892 | 2.78E-25 | 1.44E-23 | ENSG00000181019 | NQO1 | 1728 |
| ENSG00000117475.14 | 3150.468208 | 1.407087504 | 0.147271369 | 5.480274 | 4.25E-08 | 6.07E-07 | ENSG00000117475 | BLZF1 | 8548 |
| ENSG00000231346.5 | 235.2100413 | 1.407001828 | 0.138167386 | 5.840755 | 5.20E-09 | 8.27E-08 | ENSG00000231346 | NA | NA |
| ENSG00000053524.12 | 130.6159853 | 1.406867029 | 0.251333177 | 3.210348 | 0.001326 | 0.009558972 | ENSG00000053524 | MCF2L2 | 23101 |
| ENSG00000223551.1 | 128.1952176 | 1.406188137 | 0.19052245 | 4.23146 | 2.32E-05 | 0.000231068 | ENSG00000223551 | TMSB4XP[illegible] | 7118 |
| ENSG00000171552.13 | 5390.445 | 1.405808 | 0.1255711 | 6.417147 | 1.39E-10 | 2.62E-09 | ENSG00000171552 | BCL2L1 | 598 |
| ENSG00000168243.11 | 185.3598342 | 1.404196795 | 0.187873672 | 4.280519 | 1.86E-05 | 0.000188351 | ENSG00000168243 | GNG4 | 2786 |
| ENSG00000124788.18 | 414.9459038 | 1.402750513 | 0.107061204 | 7.498052 | 6.48E-14 | 1.63E-12 | ENSG00000124788 | ATXN1 | 6310 |
| ENSG00000152763.17 | 163.5542457 | 1.402470358 | 0.129073939 | 6.217137 | 5.06E-10 | 9.03E-09 | ENSG00000152763 | DNAI4 | 79819 |
| ENSG00000087842.11 | 326.1662 | 1.402372 | 0.09875778 | 8.124648 | 4.49E-16 | 1.32E-14 | ENSG00000087842 | PIR | 8544 |
| ENSG00000188039.14 | 38.25842 | 1.402192 | 0.2359245 | 3.400207 | 0.000673 | 0.005195913 | ENSG00000188039 | NWD1 | 284434 |
| ENSG00000165555.9 | 25.73775103 | 1.401162499 | 0.284222332 | 2.818788 | 0.004821 | 0.030267045 | ENSG00000165555 | NOXRED1 | 122945 |
| ENSG00000164509.15 | 466.1424547 | 1.399558511 | 0.279243246 | 2.863305 | 0.004192 | 0.026797592 | ENSG00000164509 | IL31RA | 133396 |
| ENSG00000118640.11 | 6253.237463 | 1.398826346 | 0.139903144 | 5.709853 | 1.13E-08 | 1.74E-07 | ENSG00000118640 | VAMP8 | 8673 |
| ENSG00000123595.7 | 1337.963 | 1.396847 | 0.1260538 | 6.321481 | 2.59E-10 | 4.78E-09 | ENSG00000123595 | RAB9A | 9367 |
| ENSG00000109929.10 | 3061.694859 | 1.396455105 | 0.065602202 | 12.14068 | 6.43E-34 | 5.08E-32 | ENSG00000109929 | SC5D | 6309 |
| ENSG00000262155.2 | 27.43866242 | 1.394788534 | 0.284823529 | 2.79046 | 0.005263 | 0.032707823 | ENSG00000262155 | LINC0217[illegible] | 283887 |
| ENSG00000058091.17 | 862.8247694 | 1.391251067 | 0.055173444 | 14.34116 | 1.21E-46 | 1.52E-44 | ENSG00000058091 | CDK14 | 5218 |
| ENSG00000186469.8 | 54.31072888 | 1.391236177 | 0.291726272 | 2.712255 | 0.006683 | 0.040350877 | ENSG00000186469 | GNG2 | 54331 |
| ENSG00000058085.14 | 118808.2227 | 1.390678841 | 0.077358826 | 10.22093 | 1.60E-24 | 7.82E-23 | ENSG00000058085 | LAMC2 | 3918 |
| ENSG00000080561.13 | 399.7165 | 1.389628 | 0.09846291 | 8.019552 | 1.06E-15 | 3.06E-14 | ENSG00000080561 | MID2 | 11043 |
| ENSG00000101782.15 | 6005.67472 | 1.387569423 | 0.118633433 | 6.63868 | 3.17E-11 | 6.33E-10 | ENSG00000101782 | RIOK3 | 8780 |
| ENSG00000183092.16 | 89.57057003 | 1.387396351 | 0.149518151 | 5.266226 | 1.39E-07 | 1.89E-06 | ENSG00000183092 | BEGAIN | 57596 |
| ENSG00000182612.10 | 121.5870162 | 1.385232513 | 0.204875882 | 3.832723 | 0.000127 | 0.001122754 | ENSG00000182612 | TSPAN10 | 83882 |
| ENSG00000109046.15 | 5929.668139 | 1.385096913 | 0.194213913 | 4.042434 | 5.29E-05 | 0.000497802 | ENSG00000109046 | WSB1 | 26118 |
| ENSG00000113163.16 | 7885.444005 | 1.384620161 | 0.140239225 | 5.59487 | 2.21E-08 | 3.29E-07 | ENSG00000113163 | CERT1 | 10087 |
| ENSG00000186577.14 | 447.6080553 | 1.384247637 | 0.181490951 | 4.321139 | 1.55E-05 | 0.000158966 | ENSG00000186577 | SMIM29 | 221491 |
| ENSG00000173013.5 | 120.6262804 | 1.384049602 | 0.160820047 | 4.875323 | 1.09E-06 | 1.31E-05 | ENSG00000173013 | CCDC96 | 257236 |
| ENSG00000174808.12 | 145.9600108 | 1.38146763 | 0.263275872 | 2.968246 | 0.002995 | 0.019864976 | ENSG00000174808 | BTC | 685 |
| ENSG00000115738.10 | 175.6357228 | 1.378374939 | 0.212895785 | 3.656131 | 0.000256 | 0.002142228 | ENSG00000115738 | ID2 | 3398 |
| ENSG00000124313.16 | 386.6621 | 1.378274 | 0.1536823 | 5.064176 | 4.10E-07 | 5.21E-06 | ENSG00000124313 | IQSEC2 | 23096 |
| ENSG00000074964.17 | 540.2810348 | 1.376149081 | 0.164619246 | 4.714814 | 2.42E-06 | 2.77E-05 | ENSG00000074964 | ARHGEF1 | 55160 |
| ENSG00000162576.16 | 287.3076246 | 1.375562292 | 0.144134724 | 5.380815 | 7.41E-08 | 1.04E-06 | ENSG00000162576 | MXRA8 | 54587 |
| ENSG00000143622.11 | 992.9559668 | 1.374991273 | 0.140032271 | 5.534376 | 3.12E-08 | 4.56E-07 | ENSG00000143622 | RIT1 | 6016 |
| ENSG00000227486.2 | 52.74291 | 1.37075 | 0.2434642 | 3.165764 | 0.001547 | 0.01094574 | ENSG00000227486 | NA | NA |
| ENSG00000152926.14 | 860.849898 | 1.370152173 | 0.156248935 | 4.929008 | 8.26E-07 | 1.01E-05 | ENSG00000152926 | ZNF117 | 51351 |
| ENSG00000107362.13 | 1379.415251 | 1.369297159 | 0.201498109 | 3.817888 | 0.000135 | 0.001186448 | ENSG00000107362 | ABHD17B | 51104 |
| ENSG00000113742.14 | 2870.242752 | 1.368415519 | 0.130552282 | 5.885883 | 3.96E-09 | 6.39E-08 | ENSG00000113742 | CPEB4 | 80315 |

| | | | | | | | | | |
|---|---|---|---|---|---|---|---|---|---|
| ENSG00000153294.11 | 585.2771849 | 1.36746609 | 0.223802823 | 3.429206 | 0.000605 | 0.004709409 | ENSG00000153294 | ADGRF4 | 221393 |
| ENSG00000198743.7 | 6551.743 | 1.367099 | 0.07560467 | 10.14619 | 3.45E-24 | 1.66E-22 | ENSG00000198743 | SLC5A3 | 6526 |
| ENSG00000005238.19 | 1215.361351 | 1.365420878 | 0.14167412 | 5.402687 | 6.57E-08 | 9.22E-07 | ENSG00000005238 | FAM214B | 80256 |
| ENSG00000170260.8 | 914.6161992 | 1.364698048 | 0.169600796 | 4.508812 | 6.52E-06 | 7.10E-05 | ENSG00000170260 | ZNF212 | 7988 |
| ENSG00000206140.11 | 56.88415 | 1.362835 | 0.2452276 | 3.110722 | 0.001866 | 0.01297931 | ENSG00000206140 | TMEM191( | 645426 |
| ENSG00000051108.15 | 1872.209476 | 1.359569682 | 0.052474224 | 14.4751 | 1.74E-47 | 2.25E-45 | ENSG00000051108 | HERPUD1 | 9709 |
| ENSG00000107731.12 | 605.4830647 | 1.358809996 | 0.199748138 | 3.798834 | 0.000145 | 0.001271859 | ENSG00000107731 | UNC5B | 219699 |
| ENSG00000166900.17 | 2766.870586 | 1.358638592 | 0.147827605 | 5.131914 | 2.87E-07 | 3.72E-06 | ENSG00000166900 | STX3 | 6809 |
| ENSG00000265982.1 | 61.58093931 | 1.358173613 | 0.277180219 | 2.735309 | 0.006232 | 0.037940275 | ENSG00000265982 | NA | NA |
| ENSG00000112561.18 | 288.6422552 | 1.357756413 | 0.127453436 | 5.945359 | 2.76E-09 | 4.53E-08 | ENSG00000112561 | TFEB | 7942 |
| ENSG00000112679.14 | 1572.157272 | 1.355339847 | 0.15375308 | 4.912681 | 8.98E-07 | 1.10E-05 | ENSG00000112679 | DUSP22 | 56940 |
| ENSG00000145040.4 | 80.78952207 | 1.353998627 | 0.176960776 | 4.260823 | 2.04E-05 | 0.00020469 | ENSG00000145040 | UCN2 | 90226 |
| ENSG00000198668.12 | 25167.00017 | 1.353964052 | 0.158513706 | 4.75646 | 1.97E-06 | 2.28E-05 | ENSG00000198668 | CALM1 | 801 |
| ENSG00000276952.1 | 58.69171 | 1.350946 | 0.2020082 | 3.717404 | 0.000201 | 0.001714713 | ENSG00000276952 | NA | NA |
| ENSG00000254750.1 | 63.6004197 | 1.350879992 | 0.249390583 | 3.010859 | 0.002605 | 0.017566569 | ENSG00000254750 | NA | NA |
| ENSG00000070081.16 | 2456.508431 | 1.349674457 | 0.182663792 | 4.104122 | 4.06E-05 | 0.000390623 | ENSG00000070081 | NUCB2 | 4925 |
| ENSG00000028277.21 | 160.5475 | 1.348287 | 0.2395664 | 3.123504 | 0.001787 | 0.01250371 | ENSG00000028277 | POU2F2 | 5452 |
| ENSG00000276853.1 | 27.38886696 | 1.348279302 | 0.252633223 | 2.96192 | 0.003057 | 0.020221315 | ENSG00000276853 | NA | NA |
| ENSG00000110429.14 | 3879.850076 | 1.345784952 | 0.101399391 | 7.354925 | 1.91E-13 | 4.67E-12 | ENSG00000110429 | FBXO3 | 26273 |
| ENSG00000248994.2 | 98.50321145 | 1.344880275 | 0.254481302 | 2.927053 | 0.003422 | 0.022322459 | ENSG00000248994 | LOC10537 | 1.05E+08 |
| ENSG00000105376.5 | 52.86433753 | 1.344619247 | 0.201298822 | 3.699074 | 0.000216 | 0.001831691 | ENSG00000105376 | ICAM5 | 7087 |
| ENSG00000187609.16 | 174.9054672 | 1.344577359 | 0.202439016 | 3.678033 | 0.000235 | 0.001980135 | ENSG00000187609 | EXD3 | 54932 |
| ENSG00000167703.14 | 33.37142584 | 1.343882456 | 0.272681359 | 2.728028 | 0.006371 | 0.038673006 | ENSG00000167703 | SLC43A2 | 124935 |
| ENSG00000267284.2 | 127.1343479 | 1.3430259 | 0.135606979 | 5.47926 | 4.27E-08 | 6.10E-07 | ENSG00000267284 | NA | NA |
| ENSG00000154917.11 | 388.737635 | 1.34188703 | 0.114376824 | 6.486341 | 8.79E-11 | 1.69E-09 | ENSG00000154917 | RAB6B | 51560 |
| ENSG00000137267.6 | 1853.419975 | 1.34148501 | 0.104445466 | 7.099255 | 1.25E-12 | 2.85E-11 | ENSG00000137267 | TUBB2A | 7280 |
| ENSG00000104055.15 | 72.79073389 | 1.340769745 | 0.261877468 | 2.828688 | 0.004674 | 0.029464615 | ENSG00000104055 | TGM5 | 9333 |
| ENSG00000114270.17 | 3260.311799 | 1.340767971 | 0.173651067 | 4.265842 | 1.99E-05 | 0.00020032 | ENSG00000114270 | COL7A1 | 1294 |
| ENSG00000126467.11 | 28.47161 | 1.339638 | 0.2725494 | 2.713775 | 0.006652 | 0.04023016 | ENSG00000126467 | TSKS | 60385 |
| ENSG00000185761.11 | 589.1574133 | 1.338248653 | 0.119397256 | 6.183129 | 6.28E-10 | 1.11E-08 | ENSG00000185761 | ADAMTSL | 339366 |
| ENSG00000038002.9 | 357.963473 | 1.337644923 | 0.152500056 | 4.837014 | 1.32E-06 | 1.56E-05 | ENSG00000038002 | AGA | 175 |
| ENSG00000163795.14 | 413.2500998 | 1.334944933 | 0.146170982 | 5.027981 | 4.96E-07 | 6.23E-06 | ENSG00000163795 | ZNF513 | 130557 |
| ENSG00000197780.9 | 3302.176764 | 1.334624247 | 0.107057787 | 6.861941 | 6.79E-12 | 1.45E-10 | ENSG00000197780 | TAF13 | 6884 |
| ENSG00000175793.12 | 40185.29544 | 1.333983805 | 0.176835276 | 4.150664 | 3.32E-05 | 0.00032289 | ENSG00000175793 | SFN | 2810 |
| ENSG00000099804.9 | 3538.943857 | 1.333847829 | 0.149456577 | 4.910107 | 9.10E-07 | 1.11E-05 | ENSG00000099804 | CDC34 | 997 |
| ENSG00000111266.8 | 411.9488413 | 1.333286506 | 0.118837412 | 6.170502 | 6.81E-10 | 1.19E-08 | ENSG00000111266 | DUSP16 | 80824 |
| ENSG00000083817.9 | 293.2492 | 1.333133 | 0.1397657 | 5.245442 | 1.56E-07 | 2.10E-06 | ENSG00000083817 | ZNF416 | 55659 |
| ENSG00000206075.14 | 23239.79185 | 1.332453586 | 0.084824107 | 8.63497 | 5.87E-18 | 1.98E-16 | ENSG00000206075 | SERPINB5 | 5268 |
| ENSG00000197043.14 | 397.1979358 | 1.33232779 | 0.137160643 | 5.339198 | 9.34E-08 | 1.29E-06 | ENSG00000197043 | ANXA6 | 309 |
| ENSG00000108786.10 | 250.9630836 | 1.331577304 | 0.132819403 | 5.50806 | 3.63E-08 | 5.24E-07 | ENSG00000108786 | HSD17B1 | 3292 |
| ENSG00000141644.17 | 5850.692898 | 1.331570717 | 0.055366461 | 13.21325 | 7.36E-40 | 7.36E-38 | ENSG00000141644 | MBD1 | 4152 |
| ENSG00000189077.11 | 675.5954956 | 1.331055932 | 0.077448411 | 9.439263 | 3.75E-21 | 1.54E-19 | ENSG00000189077 | TMEM120/ | 83862 |

| | | | | | | | | | |
|---|---|---|---|---|---|---|---|---|---|
| ENSG00000163064.6 | 35.926638 | 1.330861581 | 0.219190499 | 3.334367 | 0.000855 | 0.006440336 | ENSG00000163064 | EN1 | 2019 |
| ENSG00000106052.13 | 8161.28603 | 1.329122385 | 0.105819987 | 6.890214 | 5.57E-12 | 1.20E-10 | ENSG00000106052 | TAX1BP1 | 8887 |
| ENSG00000001084.13 | 1908.874568 | 1.327711961 | 0.06141581 | 11.84894 | 2.18E-32 | 1.65E-30 | ENSG00000001084 | GCLC | 2729 |
| ENSG00000224687.2 | 75.69831412 | 1.327333211 | 0.264082679 | 2.754187 | 0.005884 | 0.036075233 | ENSG00000224687 | RASAL2-A | 1E+08 |
| ENSG00000131711.15 | 204.2658978 | 1.327088728 | 0.182345317 | 3.987427 | 6.68E-05 | 0.000620439 | ENSG00000131711 | MAP1B | 4131 |
| ENSG00000110025.13 | 25.05277181 | 1.326616491 | 0.254981763 | 2.84968 | 0.004376 | 0.027865678 | ENSG00000110025 | SNX15 | 29907 |
| ENSG00000179546.4 | 44.34651451 | 1.323110443 | 0.245623291 | 2.943982 | 0.00324 | 0.021277853 | ENSG00000179546 | HTR1D | 3352 |
| ENSG00000118620.13 | 707.3451 | 1.322307 | 0.1236293 | 5.842522 | 5.14E-09 | 8.19E-08 | ENSG00000118620 | ZNF430 | 80264 |
| ENSG00000164040.17 | 3993.938173 | 1.32210889 | 0.135345291 | 5.335309 | 9.54E-08 | 1.32E-06 | ENSG00000164040 | PGRMC2 | 10424 |
| ENSG00000165474.8 | 5229.837605 | 1.321881369 | 0.165403453 | 4.364367 | 1.27E-05 | 0.000132737 | ENSG00000165474 | GJB2 | 2706 |
| ENSG00000173077.15 | 44.1382639 | 1.321219313 | 0.224782634 | 3.208519 | 0.001334 | 0.00960758 | ENSG00000173077 | DELEC1 | 50514 |
| ENSG00000119139.20 | 382.5132475 | 1.320973159 | 0.176767847 | 4.078644 | 4.53E-05 | 0.000432241 | ENSG00000119139 | TJP2 | 9414 |
| ENSG00000090674.16 | 875.2961812 | 1.319956021 | 0.131098844 | 5.491704 | 3.98E-08 | 5.70E-07 | ENSG00000090674 | MCOLN1 | 57192 |
| ENSG00000149260.18 | 73.4759878 | 1.319546797 | 0.163544886 | 4.39969 | 1.08E-05 | 0.00011398 | ENSG00000149260 | CAPN5 | 726 |
| ENSG00000136560.13 | 3089.212444 | 1.318696281 | 0.090292742 | 7.959624 | 1.73E-15 | 4.92E-14 | ENSG00000136560 | TANK | 10010 |
| ENSG00000262454.4 | 136.1568035 | 1.315980972 | 0.127909199 | 5.597572 | 2.17E-08 | 3.24E-07 | ENSG00000262454 | NA | NA |
| ENSG00000269994.3 | 87.9517982 | 1.315899858 | 0.188391644 | 3.800062 | 0.000145 | 0.00126599 | ENSG00000269994 | LOC44017 | 440173 |
| ENSG00000110492.15 | 488.9769793 | 1.315369978 | 0.189072379 | 3.783577 | 0.000155 | 0.00134846 | ENSG00000110492 | MDK | 4192 |
| ENSG00000172765.17 | 2615.458712 | 1.314435553 | 0.067684253 | 10.55542 | 4.80E-26 | 2.57E-24 | ENSG00000172765 | TMCC1 | 23023 |
| ENSG00000115295.20 | 5266.834405 | 1.313983948 | 0.086805035 | 8.225144 | 1.95E-16 | 5.90E-15 | ENSG00000115295 | CLIP4 | 79745 |
| ENSG00000184205.14 | 474.5316 | 1.313273 | 0.1519819 | 4.693141 | 2.69E-06 | 3.07E-05 | ENSG00000184205 | TSPYL2 | 64061 |
| ENSG00000198843.13 | 7297.95186 | 1.313151711 | 0.121128494 | 5.887564 | 3.92E-09 | 6.33E-08 | ENSG00000198843 | SELENOT | 51714 |
| ENSG00000223745.8 | 271.9320554 | 1.312632709 | 0.11316143 | 6.297488 | 3.03E-10 | 5.56E-09 | ENSG00000223745 | NA | NA |
| ENSG00000167103.12 | 178.6828873 | 1.31199863 | 0.136921943 | 5.200033 | 1.99E-07 | 2.64E-06 | ENSG00000167103 | PIP5KL1 | 138429 |
| ENSG00000189050.16 | 347.5004334 | 1.310523313 | 0.093779804 | 7.576507 | 3.55E-14 | 9.09E-13 | ENSG00000189050 | RNFT1 | 51136 |
| ENSG00000184349.13 | 149.1193216 | 1.310168945 | 0.134053098 | 5.297669 | 1.17E-07 | 1.61E-06 | ENSG00000184349 | EFNA5 | 1946 |
| ENSG00000003249.13 | 142.5296781 | 1.309630064 | 0.204070198 | 3.477382 | 0.000506 | 0.004003465 | ENSG00000003249 | DBNDD1 | 79007 |
| ENSG00000154845.15 | 18733.63486 | 1.309382568 | 0.110060956 | 6.445361 | 1.15E-10 | 2.19E-09 | ENSG00000154845 | PPP4R1 | 9989 |
| ENSG00000126391.14 | 2148.363025 | 1.308642737 | 0.102603226 | 6.906632 | 4.96E-12 | 1.07E-10 | ENSG00000126391 | FRMD8 | 83786 |
| ENSG00000226476.4 | 285.6461136 | 1.308413769 | 0.162542463 | 4.35833 | 1.31E-05 | 0.000136184 | ENSG00000226476 | LINC0174ℓ | 1.05E+08 |
| ENSG00000100767.16 | 131.0681064 | 1.307033561 | 0.180311665 | 3.921175 | 8.81E-05 | 0.000802836 | ENSG00000100767 | PAPLN | 89932 |
| ENSG00000205336.11 | 3240.027972 | 1.305880397 | 0.132329851 | 5.334249 | 9.59E-08 | 1.33E-06 | ENSG00000205336 | ADGRG1 | 9289 |
| ENSG00000110046.13 | 2167.999766 | 1.304953611 | 0.200828025 | 3.510235 | 0.000448 | 0.003576042 | ENSG00000110046 | ATG2A | 23130 |
| ENSG00000163602.10 | 535.678968 | 1.300485891 | 0.170609263 | 4.105791 | 4.03E-05 | 0.000387954 | ENSG00000163602 | RYBP | 23429 |
| ENSG00000077585.14 | 502.1061225 | 1.298046919 | 0.100890677 | 6.918845 | 4.55E-12 | 9.84E-11 | ENSG00000077585 | GPR137B | 7107 |
| ENSG00000159314.11 | 638.8136508 | 1.297810868 | 0.20063499 | 3.478012 | 0.000505 | 0.003996446 | ENSG00000159314 | ARHGAP2 | 201176 |
| ENSG00000136888.7 | 5485.84855 | 1.297430625 | 0.128196729 | 5.440315 | 5.32E-08 | 7.51E-07 | ENSG00000136888 | ATP6V1G' | 9550 |
| ENSG00000181649.8 | 878.999226 | 1.297220131 | 0.205319723 | 3.395778 | 0.000684 | 0.005271551 | ENSG00000181649 | PHLDA2 | 7262 |
| ENSG00000137968.16 | 937.6721353 | 1.29685062 | 0.121714504 | 5.725288 | 1.03E-08 | 1.59E-07 | ENSG00000137968 | SLC44A5 | 204962 |
| ENSG00000183876.9 | 592.4413093 | 1.296097958 | 0.190314992 | 3.65761 | 0.000255 | 0.002130583 | ENSG00000183876 | ARSI | 340075 |
| ENSG00000171159.5 | 420.5721224 | 1.295495021 | 0.199029461 | 3.494433 | 0.000475 | 0.003779836 | ENSG00000171159 | C9orf16 | 79095 |
| ENSG00000135452.10 | 909.3932812 | 1.295428688 | 0.132962599 | 5.230258 | 1.69E-07 | 2.27E-06 | ENSG00000135452 | TSPAN31 | 6302 |

| | | | | | | | | | |
|---|---|---|---|---|---|---|---|---|---|
| ENSG00000229809.8 | 229.7262516 | 1.294592204 | 0.165754175 | 4.190496 | 2.78E-05 | 0.00027384 | ENSG00000229809 | ZNF688 | 146542 |
| ENSG00000287299.1 | 86.66310227 | 1.294081262 | 0.178049739 | 3.898244 | 9.69E-05 | 0.000877674 | ENSG00000287299 | LOC10192 | 1.02E+08 |
| ENSG00000155252.13 | 2496.158637 | 1.293996991 | 0.104213003 | 6.659409 | 2.75E-11 | 5.53E-10 | ENSG00000155252 | PI4K2A | 55361 |
| ENSG00000280138.1 | 490.3676454 | 1.292916427 | 0.181389683 | 3.820043 | 0.000133 | 0.001177879 | ENSG00000280138 | NA | NA |
| ENSG00000197256.10 | 1304.619926 | 1.292575961 | 0.124381249 | 5.56817 | 2.57E-08 | 3.81E-07 | ENSG00000197256 | KANK2 | 25959 |
| ENSG00000112406.5 | 2172.052484 | 1.292563057 | 0.134196743 | 5.160804 | 2.46E-07 | 3.22E-06 | ENSG00000112406 | HECA | 51696 |
| ENSG00000259030.7 | 35.75345076 | 1.292276194 | 0.241623446 | 2.865104 | 0.004169 | 0.026652234 | ENSG00000259030 | FPGT-TNN | 1.01E+08 |
| ENSG00000075785.12 | 14677.2124 | 1.291008288 | 0.154212942 | 4.480871 | 7.43E-06 | 8.04E-05 | ENSG00000075785 | RAB7A | 7879 |
| ENSG00000157045.9 | 1446.262643 | 1.290732735 | 0.079362089 | 8.70356 | 3.22E-18 | 1.10E-16 | ENSG00000157045 | NTAN1 | 123803 |
| ENSG00000187372.11 | 83.00297712 | 1.290261741 | 0.247299763 | 2.791195 | 0.005251 | 0.032648882 | ENSG00000187372 | PCDHB13 | 56123 |
| ENSG00000167705.12 | 66.10473467 | 1.289517355 | 0.222206536 | 3.103047 | 0.001915 | 0.013265231 | ENSG00000167705 | RILP | 83547 |
| ENSG00000104946.13 | 746.8667 | 1.289182 | 0.07619926 | 9.044474 | 1.50E-19 | 5.58E-18 | ENSG00000104946 | TBC1D17 | 79735 |
| ENSG00000150636.17 | 57.98660101 | 1.288630086 | 0.257113131 | 2.678316 | 0.007399 | 0.044167141 | ENSG00000150636 | CCDC102E | 79839 |
| ENSG00000240038.6 | 124.8733533 | 1.284410019 | 0.160968478 | 4.251826 | 2.12E-05 | 0.000212689 | ENSG00000240038 | AMY2B | 280 |
| ENSG00000083223.18 | 3062.731362 | 1.284286503 | 0.090454134 | 7.565011 | 3.88E-14 | 9.87E-13 | ENSG00000083223 | TUT7 | 79670 |
| ENSG00000172638.13 | 86.81286882 | 1.284124113 | 0.167925933 | 4.073963 | 4.62E-05 | 0.000440552 | ENSG00000172638 | EFEMP2 | 30008 |
| ENSG00000160445.11 | 1435.401219 | 1.284012284 | 0.128878237 | 5.30743 | 1.11E-07 | 1.52E-06 | ENSG00000160445 | ZER1 | 10444 |
| ENSG00000131016.17 | 13799.34895 | 1.283892865 | 0.074805336 | 9.142301 | 6.11E-20 | 2.32E-18 | ENSG00000131016 | AKAP12 | 9590 |
| ENSG00000111897.7 | 9403.977994 | 1.283169755 | 0.101880189 | 6.705619 | 2.01E-11 | 4.09E-10 | ENSG00000111897 | SERINC1 | 57515 |
| ENSG00000176485.11 | 497.238755 | 1.283130554 | 0.171049359 | 3.993763 | 6.50E-05 | 0.000604718 | ENSG00000176485 | PLAAT3 | 11145 |
| ENSG00000084764.11 | 248.0712462 | 1.282579992 | 0.184299807 | 3.703639 | 0.000213 | 0.001800764 | ENSG00000084764 | MAPRE3 | 22924 |
| ENSG00000005020.13 | 2438.303031 | 1.2787499 | 0.156756848 | 4.329954 | 1.49E-05 | 0.000153357 | ENSG00000005020 | SKAP2 | 8935 |
| ENSG00000102901.13 | 1379.898671 | 1.278225248 | 0.137094619 | 4.947133 | 7.53E-07 | 9.27E-06 | ENSG00000102901 | CENPT | 80152 |
| ENSG00000156384.14 | 584.9065192 | 1.275388988 | 0.136071912 | 4.963471 | 6.92E-07 | 8.55E-06 | ENSG00000156384 | SFR1 | 119392 |
| ENSG00000251003.9 | 588.0465792 | 1.27228129 | 0.190294222 | 3.532852 | 0.000411 | 0.003309458 | ENSG00000251003 | NA | NA |
| ENSG00000060971.18 | 1998.191028 | 1.271583514 | 0.150364044 | 4.466384 | 7.96E-06 | 8.56E-05 | ENSG00000060971 | ACAA1 | 30 |
| ENSG00000213722.9 | 211.8183885 | 1.271322961 | 0.19265996 | 3.484497 | 0.000493 | 0.003907814 | ENSG00000213722 | DDAH2 | 23564 |
| ENSG00000132002.8 | 11889.3652 | 1.269293043 | 0.116295734 | 5.755095 | 8.66E-09 | 1.35E-07 | ENSG00000132002 | DNAJB1 | 3337 |
| ENSG00000279159.1 | 164.4528 | 1.269 | 0.1172634 | 5.705105 | 1.16E-08 | 1.78E-07 | ENSG00000279159 | NA | NA |
| ENSG00000034510.6 | 97831.84196 | 1.267579608 | 0.108785247 | 6.136674 | 8.43E-10 | 1.46E-08 | ENSG00000034510 | TMSB10 | 9168 |
| ENSG00000140403.12 | 2705.470766 | 1.266729943 | 0.103916985 | 6.415986 | 1.40E-10 | 2.64E-09 | ENSG00000140403 | DNAJA4 | 55466 |
| ENSG00000197329.12 | 846.5362843 | 1.265846833 | 0.125153193 | 5.320254 | 1.04E-07 | 1.42E-06 | ENSG00000197329 | PELI1 | 57162 |
| ENSG00000117305.15 | 697.6416456 | 1.26549893 | 0.144129285 | 4.617375 | 3.89E-06 | 4.34E-05 | ENSG00000117305 | HMGCL | 3155 |
| ENSG00000152223.15 | 4508.092255 | 1.265402254 | 0.152902864 | 4.351797 | 1.35E-05 | 0.000139762 | ENSG00000152223 | EPG5 | 57724 |
| ENSG00000075399.14 | 387.6678725 | 1.264914552 | 0.127499592 | 5.215033 | 1.84E-07 | 2.45E-06 | ENSG00000075399 | VPS9D1 | 9605 |
| ENSG00000197183.14 | 865.9513 | 1.26336 | 0.121456 | 5.461731 | 4.72E-08 | 6.71E-07 | ENSG00000197183 | NOL4L | 140688 |
| ENSG00000163082.10 | 1160.013864 | 1.262416159 | 0.250956716 | 2.639563 | 0.008301 | 0.048753228 | ENSG00000163082 | SGPP2 | 130367 |
| ENSG00000160055.19 | 270.4014976 | 1.262351263 | 0.093276067 | 7.100978 | 1.24E-12 | 2.81E-11 | ENSG00000160055 | TMEM234 | 56063 |
| ENSG00000129657.16 | 5302.951645 | 1.261709106 | 0.121939635 | 5.42653 | 5.75E-08 | 8.09E-07 | ENSG00000129657 | SEC14L1 | 6397 |
| ENSG00000167207.13 | 39.89064871 | 1.261071462 | 0.209608068 | 3.153846 | 0.001611 | 0.011363265 | ENSG00000167207 | NOD2 | 64127 |
| ENSG00000140682.19 | 721.161944 | 1.26025817 | 0.066993678 | 9.855529 | 6.49E-23 | 2.92E-21 | ENSG00000140682 | TGFB1I1 | 7041 |
| ENSG00000182208.14 | 1667.730777 | 1.260093055 | 0.128878639 | 5.121819 | 3.03E-07 | 3.91E-06 | ENSG00000182208 | MOB2 | 81532 |

| | | | | | | | | | |
|---|---|---|---|---|---|---|---|---|---|
| ENSG00000129465.16 | 58.80386782 | 1.25830757 | 0.248589191 | 2.648175 | 0.008093 | 0.047675547 | ENSG00000129465 | RIPK3 | 11035 |
| ENSG00000102796.11 | 53.91791858 | 1.257806457 | 0.235291794 | 2.795705 | 0.005179 | 0.032226832 | ENSG00000102796 | DHRS12 | 79758 |
| ENSG00000170871.12 | 4195.536121 | 1.257704767 | 0.076918643 | 8.550655 | 1.22E-17 | 4.04E-16 | ENSG00000170871 | KIAA0232 | 9778 |
| ENSG00000162782.16 | 384.9536322 | 1.25737667 | 0.119223219 | 5.513831 | 3.51E-08 | 5.08E-07 | ENSG00000162782 | TDRD5 | 163589 |
| ENSG00000131100.13 | 6249.669 | 1.257073 | 0.1168897 | 5.621313 | 1.90E-08 | 2.84E-07 | ENSG00000131100 | ATP6V1E1 | 529 |
| ENSG00000259456.4 | 46.06476 | 1.254045 | 0.2011485 | 3.251552 | 0.001148 | 0.008389871 | ENSG00000259456 | ADNP-AS1 | 1.02E+08 |
| ENSG00000164296.7 | 351.4625706 | 1.253565365 | 0.086023464 | 7.597524 | 3.02E-14 | 7.77E-13 | ENSG00000164296 | TIGD6 | 81789 |
| ENSG00000129518.9 | 1629.36017 | 1.252875209 | 0.100786965 | 6.477774 | 9.31E-11 | 1.78E-09 | ENSG00000129518 | EAPP | 55837 |
| ENSG00000198431.16 | 14056.24187 | 1.251917181 | 0.095782797 | 6.806203 | 1.00E-11 | 2.11E-10 | ENSG00000198431 | TXNRD1 | 7296 |
| ENSG00000051620.11 | 4725.909938 | 1.251850697 | 0.118317426 | 5.509338 | 3.60E-08 | 5.20E-07 | ENSG00000051620 | HEBP2 | 23593 |
| ENSG00000143367.16 | 1441.375788 | 1.251501609 | 0.120763054 | 5.394875 | 6.86E-08 | 9.61E-07 | ENSG00000143367 | TUFT1 | 7286 |
| ENSG00000119655.10 | 7764.308346 | 1.250355662 | 0.080319292 | 8.097129 | 5.63E-16 | 1.65E-14 | ENSG00000119655 | NPC2 | 10577 |
| ENSG00000268879.1 | 57.6745 | 1.250302 | 0.2411628 | 2.696528 | 0.007007 | 0.04208729 | ENSG00000268879 | NA | NA |
| ENSG00000132950.19 | 878.8869015 | 1.249987349 | 0.105921246 | 6.136515 | 8.44E-10 | 1.46E-08 | ENSG00000132950 | ZMYM5 | 9205 |
| ENSG00000168306.13 | 47.14961167 | 1.24745858 | 0.215812462 | 3.000098 | 0.002699 | 0.018112125 | ENSG00000168306 | ACOX2 | 8309 |
| ENSG00000111962.8 | 191.292376 | 1.247103329 | 0.158974934 | 4.070474 | 4.69E-05 | 0.000446405 | ENSG00000111962 | UST | 10090 |
| ENSG00000154146.13 | 140.6401077 | 1.246991643 | 0.177242901 | 3.650311 | 0.000262 | 0.002185185 | ENSG00000154146 | NRGN | 4900 |
| ENSG00000111863.12 | 669.2529362 | 1.246802587 | 0.08469574 | 7.636778 | 2.23E-14 | 5.81E-13 | ENSG00000111863 | ADTRP | 84830 |
| ENSG00000137331.12 | 1487.94822 | 1.246578122 | 0.195025826 | 3.315346 | 0.000915 | 0.006846597 | ENSG00000137331 | IER3 | 8870 |
| ENSG00000116815.16 | 764.8592688 | 1.243530424 | 0.090585056 | 7.104157 | 1.21E-12 | 2.75E-11 | ENSG00000116815 | CD58 | 965 |
| ENSG00000133101.10 | 370.1931096 | 1.24344913 | 0.201221752 | 3.197712 | 0.001385 | 0.009926655 | ENSG00000133101 | CCNA1 | 8900 |
| ENSG00000184500.15 | 341.312246 | 1.243269037 | 0.11191813 | 5.747675 | 9.05E-09 | 1.41E-07 | ENSG00000184500 | PROS1 | 5627 |
| ENSG00000065717.15 | 57.05125296 | 1.243007773 | 0.179067647 | 3.590865 | 0.00033 | 0.002700565 | ENSG00000065717 | TLE2 | 7089 |
| ENSG00000019995.6 | 1450.111088 | 1.240690331 | 0.118718927 | 5.396699 | 6.79E-08 | 9.53E-07 | ENSG00000019995 | ZRANB1 | 54764 |
| ENSG00000143409.15 | 501.2624625 | 1.240587267 | 0.199508501 | 3.210827 | 0.001324 | 0.009546242 | ENSG00000143409 | MINDY1 | 55793 |
| ENSG00000137364.5 | 987.7359836 | 1.240482218 | 0.148504088 | 4.312893 | 1.61E-05 | 0.000164607 | ENSG00000137364 | TPMT | 7172 |
| ENSG00000111319.13 | 966.6982274 | 1.240359692 | 0.189945695 | 3.371278 | 0.000748 | 0.005715548 | ENSG00000111319 | SCNN1A | 6337 |
| ENSG00000157107.14 | 4661.024303 | 1.240339535 | 0.133109891 | 4.810608 | 1.50E-06 | 1.77E-05 | ENSG00000157107 | FCHO2 | 115548 |
| ENSG00000075391.16 | 3911.885762 | 1.240076833 | 0.126944696 | 5.042171 | 4.60E-07 | 5.80E-06 | ENSG00000075391 | RASAL2 | 9462 |
| ENSG00000139597.18 | 45.02411287 | 1.240013358 | 0.20567371 | 3.11179 | 0.00186 | 0.012935832 | ENSG00000139597 | N4BP2L1 | 90634 |
| ENSG00000243244.6 | 44.94226705 | 1.239834388 | 0.196941898 | 3.248848 | 0.001159 | 0.008463033 | ENSG00000243244 | STON1 | 11037 |
| ENSG00000235109.7 | 203.3954614 | 1.235694733 | 0.153510056 | 4.141062 | 3.46E-05 | 0.000335762 | ENSG00000235109 | ZSCAN31 | 64288 |
| ENSG00000105472.13 | 37.70785 | 1.234893 | 0.2404071 | 2.640908 | 0.008268 | 0.0486134 | ENSG00000105472 | CLEC11A | 6320 |
| ENSG00000116954.8 | 1145.800366 | 1.234466013 | 0.147269326 | 4.308202 | 1.65E-05 | 0.000167879 | ENSG00000116954 | RRAGC | 64121 |
| ENSG00000165516.11 | 1554.427347 | 1.230585545 | 0.115759192 | 5.447391 | 5.11E-08 | 7.23E-07 | ENSG00000165516 | KLHDC2 | 23588 |
| ENSG00000159363.18 | 3826.994136 | 1.230231555 | 0.134634202 | 4.681066 | 2.85E-06 | 3.24E-05 | ENSG00000159363 | ATP13A2 | 23400 |
| ENSG00000180354.16 | 1735.202357 | 1.229787604 | 0.070260496 | 8.963609 | 3.14E-19 | 1.15E-17 | ENSG00000180354 | MTURN | 222166 |
| ENSG00000138834.12 | 2499.229393 | 1.229539506 | 0.122268983 | 5.148808 | 2.62E-07 | 3.42E-06 | ENSG00000138834 | MAPK8IP3 | 23162 |
| ENSG00000196110.7 | 162.7220116 | 1.229340759 | 0.214922997 | 2.928215 | 0.003409 | 0.022261016 | ENSG00000196110 | ZNF699 | 374879 |
| ENSG00000127663.15 | 1840.697292 | 1.228322975 | 0.12864221 | 4.884268 | 1.04E-06 | 1.25E-05 | ENSG00000127663 | KDM4B | 23030 |
| ENSG00000261087.1 | 70.8707109 | 1.228107336 | 0.192189187 | 3.268172 | 0.001082 | 0.007973674 | ENSG00000261087 | ZNNT1 | 1.16E+08 |
| ENSG00000260916.7 | 264.0308063 | 1.22766076 | 0.095041212 | 6.60409 | 4.00E-11 | 7.90E-10 | ENSG00000260916 | CCPG1 | 9236 |

| | | | | | | | | | |
|---|---|---|---|---|---|---|---|---|---|
| ENSG00000211459.2 | 45938.49 | 1.226926 | 0.05334569 | 11.75213 | 6.89E-32 | 5.13E-30 | ENSG00000211459 | NA | NA |
| ENSG00000183977.13 | 33.24487443 | 1.226083078 | 0.2243591 | 2.79054 | 0.005262 | 0.032707313 | ENSG00000183977 | PP2D1 | 151649 |
| ENSG00000241170.3 | 51.11325215 | 1.225150877 | 0.215057854 | 2.906896 | 0.00365 | 0.023650379 | ENSG00000241170 | RPL31P46 | 1E+08 |
| ENSG00000109944.10 | 256.7537076 | 1.223002835 | 0.160019033 | 3.893305 | 9.89E-05 | 0.000894223 | ENSG00000109944 | JHY | 79864 |
| ENSG00000109458.8 | 1045.731054 | 1.220706001 | 0.082949957 | 7.482897 | 7.27E-14 | 1.83E-12 | ENSG00000109458 | GAB1 | 2549 |
| ENSG00000140961.13 | 174.9064497 | 1.220085509 | 0.113639135 | 5.456619 | 4.85E-08 | 6.89E-07 | ENSG00000140961 | OSGIN1 | 29948 |
| ENSG00000110921.14 | 1181.889632 | 1.219665468 | 0.066771929 | 9.280329 | 1.69E-20 | 6.69E-19 | ENSG00000110921 | MVK | 4598 |
| ENSG00000256683.7 | 122.3122 | 1.218993 | 0.1489408 | 4.155966 | 3.24E-05 | 0.000315756 | ENSG00000256683 | ZNF350 | 59348 |
| ENSG00000270629.6 | 606.7934205 | 1.21850635 | 0.166094513 | 3.723822 | 0.000196 | 0.00167544 | ENSG00000270629 | NBPF14 | 25832 |
| ENSG00000178038.17 | 4407.156079 | 1.217715672 | 0.118554276 | 5.210404 | 1.88E-07 | 2.51E-06 | ENSG00000178038 | ALS2CL | 259173 |
| ENSG00000178026.13 | 153.7335 | 1.212243 | 0.1487077 | 4.117088 | 3.84E-05 | 0.000371035 | ENSG00000178026 | LRRC75B | 388886 |
| ENSG00000160991.16 | 489.0165712 | 1.210588036 | 0.096280387 | 6.34177 | 2.27E-10 | 4.22E-09 | ENSG00000160991 | ORAI2 | 80228 |
| ENSG00000227051.6 | 467.7377278 | 1.210259366 | 0.113256004 | 5.388318 | 7.11E-08 | 9.96E-07 | ENSG00000227051 | C14orf132 | 56967 |
| ENSG00000148572.16 | 1484.402957 | 1.208904831 | 0.088471193 | 6.882521 | 5.88E-12 | 1.26E-10 | ENSG00000148572 | NRBF2 | 29982 |
| ENSG00000151743.11 | 287.2253077 | 1.208174997 | 0.224226445 | 2.712325 | 0.006681 | 0.040350877 | ENSG00000151743 | AMN1 | 196394 |
| ENSG00000108479.11 | 239.9562859 | 1.206284698 | 0.122112838 | 4.964955 | 6.87E-07 | 8.50E-06 | ENSG00000108479 | GALK1 | 2584 |
| ENSG00000174307.6 | 3159.848109 | 1.206231412 | 0.132566629 | 4.573032 | 4.81E-06 | 5.31E-05 | ENSG00000174307 | PHLDA3 | 23612 |
| ENSG00000261189.1 | 71.04741003 | 1.205370698 | 0.189999891 | 3.186163 | 0.001442 | 0.010284902 | ENSG00000261189 | NA | NA |
| ENSG00000113369.9 | 1952.131994 | 1.204827594 | 0.084255923 | 7.178458 | 7.05E-13 | 1.63E-11 | ENSG00000113369 | ARRDC3 | 57561 |
| ENSG00000159335.16 | 4092.219422 | 1.204517457 | 0.063363967 | 9.540398 | 1.42E-21 | 5.95E-20 | ENSG00000159335 | PTMS | 5763 |
| ENSG00000096717.12 | 2034.79463 | 1.203814135 | 0.175974805 | 3.431253 | 0.000601 | 0.004682073 | ENSG00000096717 | SIRT1 | 23411 |
| ENSG00000173715.16 | 1296.806764 | 1.203518513 | 0.144103757 | 4.188083 | 2.81E-05 | 0.000276665 | ENSG00000173715 | C11orf80 | 79703 |
| ENSG00000156675.15 | 1819.997168 | 1.200998165 | 0.121752788 | 4.936217 | 7.97E-07 | 9.77E-06 | ENSG00000156675 | RAB11FIP | 80223 |
| ENSG00000187583.10 | 550.3801911 | 1.200953083 | 0.139013248 | 4.322991 | 1.54E-05 | 0.000157792 | ENSG00000187583 | PLEKHN1 | 84069 |
| ENSG00000250748.6 | 82.25336376 | 1.200687978 | 0.188336077 | 3.189447 | 0.001425 | 0.010195694 | ENSG00000250748 | LOC10536 | 1.05E+08 |
| ENSG00000048140.18 | 9072.607231 | 1.200032078 | 0.164138359 | 3.655648 | 0.000257 | 0.002144919 | ENSG00000048140 | TSPAN17 | 26262 |
| ENSG00000007255.10 | 98.44576 | 1.198863 | 0.1488379 | 4.023595 | 5.73E-05 | 0.000537281 | ENSG00000007255 | TRAPPC6/ | 79090 |
| ENSG00000143412.10 | 127.5584025 | 1.198799119 | 0.162943634 | 3.674885 | 0.000238 | 0.002003441 | ENSG00000143412 | ANXA9 | 8416 |
| ENSG00000122378.14 | 3458.735321 | 1.19848093 | 0.086114829 | 6.949801 | 3.66E-12 | 7.97E-11 | ENSG00000122378 | PRXL2A | 84293 |
| ENSG00000187642.9 | 181.8697744 | 1.19762395 | 0.141421919 | 4.225823 | 2.38E-05 | 0.000236488 | ENSG00000187642 | PERM1 | 84808 |
| ENSG00000088538.13 | 255.0917834 | 1.197168423 | 0.203592942 | 2.933149 | 0.003355 | 0.021947967 | ENSG00000088538 | DOCK3 | 1795 |
| ENSG00000125971.16 | 390.4959 | 1.195902 | 0.1023487 | 5.822275 | 5.81E-09 | 9.19E-08 | ENSG00000125971 | DYNLRB1 | 83658 |
| ENSG00000089356.18 | 114.4707 | 1.195401 | 0.1821688 | 3.268403 | 0.001082 | 0.007969362 | ENSG00000089356 | FXYD3 | 5349 |
| ENSG00000176903.4 | 1375.334371 | 1.194424291 | 0.070790082 | 8.397 | 4.58E-17 | 1.45E-15 | ENSG00000176903 | PNMA1 | 9240 |
| ENSG00000225663.7 | 995.6294375 | 1.19393155 | 0.158751334 | 3.74127 | 0.000183 | 0.001572862 | ENSG00000225663 | MCRIP1 | 348262 |
| ENSG00000147010.18 | 20166.58 | 1.192646 | 0.08648092 | 6.852912 | 7.24E-12 | 1.54E-10 | ENSG00000147010 | SH3KBP1 | 30011 |
| ENSG00000254682.1 | 55.35404571 | 1.192400444 | 0.169671858 | 3.491448 | 0.00048 | 0.003813203 | ENSG00000254682 | NA | NA |
| ENSG00000177606.6 | 4110.658777 | 1.192117945 | 0.102974275 | 5.750154 | 8.92E-09 | 1.39E-07 | ENSG00000177606 | JUN | 3725 |
| ENSG00000065833.9 | 1921.264644 | 1.191578898 | 0.104651798 | 5.652831 | 1.58E-08 | 2.38E-07 | ENSG00000065833 | ME1 | 4199 |
| ENSG00000110852.4 | 330.0861146 | 1.189366713 | 0.169422864 | 3.478673 | 0.000504 | 0.003987787 | ENSG00000110852 | CLEC2B | 9976 |
| ENSG00000105227.14 | 180.9361 | 1.18855 | 0.146604 | 4.014553 | 5.96E-05 | 0.000556729 | ENSG00000105227 | PRX | 57716 |
| ENSG00000279530.2 | 84.54490551 | 1.18625471 | 0.208281829 | 2.814718 | 0.004882 | 0.030624071 | ENSG00000279530 | NA | NA |

| | | | | | | | | | |
|---|---|---|---|---|---|---|---|---|---|
| ENSG00000205542.11 | 30208.17 | 1.185934 | 0.05716249 | 10.25033 | 1.18E-24 | 5.80E-23 | ENSG00000205542 | TMSB4X | 7114 |
| ENSG00000175137.11 | 1914.33287 | 1.185074619 | 0.117608206 | 4.974777 | 6.53E-07 | 8.10E-06 | ENSG00000175137 | SH3BP5L | 80851 |
| ENSG00000164211.13 | 4087.952811 | 1.18463418 | 0.068628958 | 8.518768 | 1.61E-17 | 5.27E-16 | ENSG00000164211 | STARD4 | 134429 |
| ENSG00000242802.9 | 1517.279527 | 1.184196657 | 0.150644625 | 3.877979 | 0.000105 | 0.000947311 | ENSG00000242802 | AP5Z1 | 9907 |
| ENSG00000173599.14 | 713.6096641 | 1.182556555 | 0.1000756 | 5.821165 | 5.84E-09 | 9.25E-08 | ENSG00000173599 | PC | 5091 |
| ENSG00000196569.12 | 120.9776766 | 1.182476956 | 0.132870849 | 4.383783 | 1.17E-05 | 0.000122055 | ENSG00000196569 | LAMA2 | 3908 |
| ENSG00000185813.10 | 1042.982174 | 1.18199191 | 0.066584556 | 8.740644 | 2.32E-18 | 7.99E-17 | ENSG00000185813 | PCYT2 | 5833 |
| ENSG00000134955.11 | 2553.852013 | 1.180189583 | 0.128308941 | 4.521817 | 6.13E-06 | 6.71E-05 | ENSG00000134955 | SLC37A2 | 219855 |
| ENSG00000152377.14 | 303.8629349 | 1.179873298 | 0.207141911 | 2.799401 | 0.00512 | 0.031904955 | ENSG00000152377 | SPOCK1 | 6695 |
| ENSG00000147689.16 | 12510.10743 | 1.179788467 | 0.130690142 | 4.43636 | 9.15E-06 | 9.77E-05 | ENSG00000147689 | FAM83A | 84985 |
| ENSG00000148154.10 | 6318.071485 | 1.178125581 | 0.185913976 | 3.10964 | 0.001873 | 0.013013332 | ENSG00000148154 | UGCG | 7357 |
| ENSG00000108448.21 | 922.6988392 | 1.177804699 | 0.110775449 | 5.216 | 1.83E-07 | 2.44E-06 | ENSG00000108448 | TRIM16L | 147166 |
| ENSG00000275216.2 | 6203.151444 | 1.177288165 | 0.093775414 | 6.156072 | 7.46E-10 | 1.30E-08 | ENSG00000275216 | NA | NA |
| ENSG00000223891.6 | 121.3743 | 1.174981 | 0.2054874 | 2.798132 | 0.00514 | 0.03202305 | ENSG00000223891 | OSER1-D1 | 1.01E+08 |
| ENSG00000143443.9 | 96.45135288 | 1.174474398 | 0.164252391 | 3.49751 | 0.00047 | 0.003737609 | ENSG00000143443 | C1orf56 | 54964 |
| ENSG00000166128.13 | 1797.32029 | 1.174241764 | 0.14711211 | 3.90343 | 9.48E-05 | 0.000860534 | ENSG00000166128 | RAB8B | 51762 |
| ENSG00000147155.11 | 946.0458 | 1.174033 | 0.09669058 | 5.936803 | 2.91E-09 | 4.77E-08 | ENSG00000147155 | EBP | 10682 |
| ENSG00000158079.16 | 1455.46179 | 1.17152455 | 0.150577314 | 3.795555 | 0.000147 | 0.001287939 | ENSG00000158079 | PTPDC1 | 138639 |
| ENSG00000254531.1 | 57.25305364 | 1.168872339 | 0.208341033 | 2.730486 | 0.006324 | 0.03842083 | ENSG00000254531 | FLJ20021 | 90024 |
| ENSG00000173482.16 | 182.827446 | 1.168027607 | 0.10422501 | 5.450013 | 5.04E-08 | 7.12E-07 | ENSG00000173482 | PTPRM | 5797 |
| ENSG00000137203.12 | 6383.337083 | 1.167377484 | 0.14539049 | 3.902439 | 9.52E-05 | 0.000863771 | ENSG00000137203 | TFAP2A | 7020 |
| ENSG00000145687.16 | 271.5233417 | 1.167297611 | 0.08896031 | 6.376974 | 1.81E-10 | 3.38E-09 | ENSG00000145687 | SSBP2 | 23635 |
| ENSG00000213445.10 | 698.1780611 | 1.167201375 | 0.129262417 | 4.387984 | 1.14E-05 | 0.000119863 | ENSG00000213445 | SIPA1 | 6494 |
| ENSG00000134531.10 | 20495.99153 | 1.166843068 | 0.103663148 | 5.468125 | 4.55E-08 | 6.47E-07 | ENSG00000134531 | EMP1 | 2012 |
| ENSG00000134202.10 | 1238.01963 | 1.165044926 | 0.057282762 | 9.864136 | 5.95E-23 | 2.69E-21 | ENSG00000134202 | GSTM3 | 2947 |
| ENSG00000123240.17 | 9277.903899 | 1.164525897 | 0.130827571 | 4.315038 | 1.60E-05 | 0.000163142 | ENSG00000123240 | OPTN | 10133 |
| ENSG00000255874.2 | 343.2735878 | 1.164235935 | 0.116348686 | 4.849526 | 1.24E-06 | 1.47E-05 | ENSG00000255874 | PRECSIT | 283487 |
| ENSG00000116017.11 | 796.8925648 | 1.16421765 | 0.178902367 | 3.153774 | 0.001612 | 0.011363265 | ENSG00000116017 | ARID3A | 1820 |
| ENSG00000240891.7 | 1075.191602 | 1.164083442 | 0.211093045 | 2.672203 | 0.007536 | 0.044819098 | ENSG00000240891 | PLCXD2 | 257068 |
| ENSG00000253276.3 | 1390.903093 | 1.162623968 | 0.208751518 | 2.695185 | 0.007035 | 0.042220054 | ENSG00000253276 | CCDC71L | 168455 |
| ENSG00000162804.14 | 82.73199841 | 1.1625642 | 0.184510505 | 3.048955 | 0.002296 | 0.015647425 | ENSG00000162804 | SNED1 | 25992 |
| ENSG00000070404.10 | 2698.62963 | 1.16186153 | 0.073064215 | 7.689969 | 1.47E-14 | 3.88E-13 | ENSG00000070404 | FSTL3 | 10272 |
| ENSG00000204344.14 | 354.5153158 | 1.161548131 | 0.134756592 | 4.167129 | 3.08E-05 | 0.000301794 | ENSG00000204344 | STK19 | 8859 |
| ENSG00000100554.12 | 5199.16528 | 1.158019595 | 0.13794814 | 4.04514 | 5.23E-05 | 0.000492434 | ENSG00000100554 | ATP6V1D | 51382 |
| ENSG00000110925.6 | 1371.220906 | 1.15691678 | 0.124161691 | 4.485416 | 7.28E-06 | 7.89E-05 | ENSG00000110925 | CSRNP2 | 81566 |
| ENSG00000132471.12 | 1899.354396 | 1.156453697 | 0.162175478 | 3.431183 | 0.000601 | 0.004682073 | ENSG00000132471 | WBP2 | 23558 |
| ENSG00000285278.1 | 134.8996356 | 1.156011533 | 0.186998989 | 2.97334 | 0.002946 | 0.019556757 | ENSG00000285278 | TFAP2A-A | 1.1E+08 |
| ENSG00000160255.18 | 154.5182 | 1.154879 | 0.1214203 | 4.569901 | 4.88E-06 | 5.38E-05 | ENSG00000160255 | ITGB2 | 3689 |
| ENSG00000163703.17 | 453.7557524 | 1.153878926 | 0.193828394 | 2.857574 | 0.004269 | 0.027227497 | ENSG00000163703 | CRELD1 | 78987 |
| ENSG00000036054.13 | 4729.122611 | 1.151955613 | 0.121469437 | 4.543988 | 5.52E-06 | 6.06E-05 | ENSG00000036054 | TBC1D23 | 55773 |
| ENSG00000147408.14 | 102.4426389 | 1.150709966 | 0.173355782 | 3.176761 | 0.001489 | 0.010592574 | ENSG00000147408 | CSGALNA | 55790 |
| ENSG00000198933.9 | 222.0361614 | 1.149420808 | 0.130415744 | 4.212841 | 2.52E-05 | 0.000249942 | ENSG00000198933 | TBKBP1 | 9755 |

| | | | | | | | | | |
|---|---|---|---|---|---|---|---|---|---|
| ENSG00000196878.15 | 44544.61024 | 1.148884341 | 0.074970276 | 7.32136 | 2.45E-13 | 5.92E-12 | ENSG00000196878 | LAMB3 | 3914 |
| ENSG00000181004.10 | 255.0400141 | 1.146323437 | 0.166847639 | 3.274385 | 0.001059 | 0.007809033 | ENSG00000181004 | BBS12 | 166379 |
| ENSG00000144566.11 | 3905.250129 | 1.145676511 | 0.145795395 | 3.742755 | 0.000182 | 0.001564598 | ENSG00000144566 | RAB5A | 5868 |
| ENSG00000189171.14 | 1190.710763 | 1.144893066 | 0.079979447 | 6.812914 | 9.56E-12 | 2.02E-10 | ENSG00000189171 | S100A13 | 6284 |
| ENSG00000132334.16 | 1876.546031 | 1.144315474 | 0.054618183 | 9.965829 | 2.15E-23 | 1.00E-21 | ENSG00000132334 | PTPRE | 5791 |
| ENSG00000188112.9 | 2031.088549 | 1.144110753 | 0.161619202 | 3.366622 | 0.000761 | 0.005804582 | ENSG00000188112 | C6orf132 | 647024 |
| ENSG00000153246.13 | 1324.121127 | 1.143787962 | 0.11717372 | 4.64087 | 3.47E-06 | 3.90E-05 | ENSG00000153246 | PLA2R1 | 22925 |
| ENSG00000013563.14 | 176.5581 | 1.14295 | 0.1758794 | 3.087058 | 0.002021 | 0.01393472 | ENSG00000013563 | DNASE1L1 | 1774 |
| ENSG00000176531.10 | 253.9411 | 1.142739 | 0.1671857 | 3.246322 | 0.001169 | 0.008529154 | ENSG00000176531 | PHLDB3 | 653583 |
| ENSG00000111145.8 | 12685.74726 | 1.141952803 | 0.091570358 | 5.918431 | 3.25E-09 | 5.30E-08 | ENSG00000111145 | ELK3 | 2004 |
| ENSG00000002822.15 | 75.14797366 | 1.140826476 | 0.170474079 | 3.172485 | 0.001511 | 0.010732615 | ENSG00000002822 | MAD1L1 | 8379 |
| ENSG00000188177.14 | 491.4460949 | 1.140187801 | 0.139832846 | 3.863097 | 0.000112 | 0.001002544 | ENSG00000188177 | ZC3H6 | 376940 |
| ENSG00000153814.13 | 273.4046482 | 1.140100178 | 0.138581682 | 3.897342 | 9.73E-05 | 0.000880648 | ENSG00000153814 | JAZF1 | 221895 |
| ENSG00000186352.9 | 85.96022666 | 1.13981593 | 0.186555992 | 2.893587 | 0.003809 | 0.024533025 | ENSG00000186352 | ANKRD37 | 353322 |
| ENSG00000119950.21 | 732.7670464 | 1.139130167 | 0.076159301 | 7.07898 | 1.45E-12 | 3.27E-11 | ENSG00000119950 | MXI1 | 4601 |
| ENSG00000143882.12 | 77.33394056 | 1.138365471 | 0.176395351 | 3.052039 | 0.002273 | 0.015515346 | ENSG00000143882 | ATP6V1C2 | 245973 |
| ENSG00000177000.12 | 632.613101 | 1.138321092 | 0.190910569 | 2.819755 | 0.004806 | 0.030183081 | ENSG00000177000 | MTHFR | 4524 |
| ENSG00000134590.13 | 860.5529 | 1.138198 | 0.0980038 | 5.491601 | 3.98E-08 | 5.70E-07 | ENSG00000134590 | RTL8C | 8933 |
| ENSG00000204949.9 | 472.5482627 | 1.138086158 | 0.119271662 | 4.511433 | 6.44E-06 | 7.02E-05 | ENSG00000204949 | FAM83A-A | 1E+08 |
| ENSG00000104892.17 | 510.6971 | 1.137343 | 0.1903632 | 2.822726 | 0.004762 | 0.02994725 | ENSG00000104892 | KLC3 | 147700 |
| ENSG00000104361.10 | 2450.995332 | 1.137333506 | 0.072005615 | 7.462383 | 8.50E-14 | 2.12E-12 | ENSG00000104361 | NIPAL2 | 79815 |
| ENSG00000178980.15 | 3789.516 | 1.137066 | 0.08311613 | 6.461635 | 1.04E-10 | 1.97E-09 | ENSG00000178980 | SELENOW | 6415 |
| ENSG00000165434.8 | 688.2896233 | 1.13527408 | 0.09290571 | 5.761477 | 8.34E-09 | 1.30E-07 | ENSG00000165434 | PGM2L1 | 283209 |
| ENSG00000136802.11 | 8000.876134 | 1.134980175 | 0.108788883 | 4.9176 | 8.76E-07 | 1.07E-05 | ENSG00000136802 | LRRC8A | 56262 |
| ENSG00000090530.10 | 98.82772491 | 1.134591028 | 0.131989591 | 4.050251 | 5.12E-05 | 0.000482655 | ENSG00000090530 | P3H2 | 55214 |
| ENSG00000101844.18 | 722.1112 | 1.134076 | 0.1075594 | 4.965411 | 6.86E-07 | 8.48E-06 | ENSG00000101844 | ATG4A | 115201 |
| ENSG00000125733.18 | 5093.946514 | 1.133347504 | 0.202187491 | 2.637886 | 0.008342 | 0.048919203 | ENSG00000125733 | TRIP10 | 9322 |
| ENSG00000119878.5 | 1181.931312 | 1.133111351 | 0.129562515 | 4.114704 | 3.88E-05 | 0.000374483 | ENSG00000119878 | CRIPT | 9419 |
| ENSG00000074319.13 | 4578.722382 | 1.131031437 | 0.105392289 | 5.038618 | 4.69E-07 | 5.91E-06 | ENSG00000074319 | TSG101 | 7251 |
| ENSG00000259153.2 | 835.8684909 | 1.130465133 | 0.116542613 | 4.551684 | 5.32E-06 | 5.85E-05 | ENSG00000259153 | LOC10050 | 1.01E+08 |
| ENSG00000162852.14 | 1740.879251 | 1.129566274 | 0.127907002 | 4.140245 | 3.47E-05 | 0.000336839 | ENSG00000162852 | CNST | 163882 |
| ENSG00000143322.21 | 5536.04484 | 1.128681198 | 0.112177346 | 4.712905 | 2.44E-06 | 2.79E-05 | ENSG00000143322 | ABL2 | 27 |
| ENSG00000170175.11 | 502.5311015 | 1.127704485 | 0.18608547 | 2.835818 | 0.004571 | 0.028869565 | ENSG00000170175 | CHRNB1 | 1140 |
| ENSG00000117069.15 | 73.0329586 | 1.127388762 | 0.190530213 | 2.768006 | 0.00564 | 0.034740436 | ENSG00000117069 | ST6GALNA | 81849 |
| ENSG00000168461.13 | 6413.752037 | 1.126747119 | 0.12009661 | 4.386028 | 1.15E-05 | 0.00012085 | ENSG00000168461 | RAB31 | 11031 |
| ENSG00000147383.11 | 1694.735 | 1.125789 | 0.06003085 | 8.758653 | 1.98E-18 | 6.86E-17 | ENSG00000147383 | NSDHL | 50814 |
| ENSG00000105404.11 | 218.6966 | 1.125664 | 0.150455 | 3.493827 | 0.000476 | 0.003786162 | ENSG00000105404 | RABAC1 | 10567 |
| ENSG00000213694.5 | 484.4940736 | 1.124381734 | 0.116092297 | 4.516938 | 6.27E-06 | 6.85E-05 | ENSG00000213694 | S1PR3 | 1903 |
| ENSG00000116663.11 | 455.0781454 | 1.12326955 | 0.140604143 | 3.72158 | 0.000198 | 0.001688763 | ENSG00000116663 | FBXO6 | 26270 |
| ENSG00000007866.21 | 1639.223229 | 1.121390636 | 0.189712461 | 2.74832 | 0.00599 | 0.036634412 | ENSG00000007866 | TEAD3 | 7005 |
| ENSG00000138613.14 | 754.0569899 | 1.121289358 | 0.102244251 | 5.098471 | 3.42E-07 | 4.40E-06 | ENSG00000138613 | APH1B | 83464 |
| ENSG00000006704.10 | 2084.136619 | 1.119702599 | 0.127273172 | 4.083363 | 4.44E-05 | 0.000424313 | ENSG00000006704 | GTF2IRD1 | 9569 |

| | | | | | | | | | |
|---|---|---|---|---|---|---|---|---|---|
| ENSG00000169228.14 | 245.5118148 | 1.119007536 | 0.197230824 | 2.631473 | 0.008502 | 0.049742586 | ENSG00000169228 | RAB24 | 53917 |
| ENSG00000165782.10 | 1191.756799 | 1.11804118 | 0.119666801 | 4.32903 | 1.50E-05 | 0.000153942 | ENSG00000165782 | PIP4P1 | 90809 |
| ENSG00000188001.10 | 81.78961817 | 1.117799281 | 0.153294422 | 3.377809 | 0.000731 | 0.005597577 | ENSG00000188001 | TPRG1 | 285386 |
| ENSG00000122912.15 | 1084.065666 | 1.117695011 | 0.105599374 | 4.902444 | 9.47E-07 | 1.15E-05 | ENSG00000122912 | SLC25A16 | 8034 |
| ENSG00000285533.2 | 76.06511983 | 1.117157344 | 0.189158448 | 2.73399 | 0.006257 | 0.038057786 | ENSG00000285533 | RELA-DT | 1.05E+08 |
| ENSG00000137947.12 | 1779.347492 | 1.117022723 | 0.093873765 | 5.507638 | 3.64E-08 | 5.24E-07 | ENSG00000137947 | GTF2B | 2959 |
| ENSG00000286133.2 | 84.2165127 | 1.116553785 | 0.186168351 | 2.77466 | 0.005526 | 0.034148542 | ENSG00000286133 | NA | NA |
| ENSG00000214530.9 | 2703.841578 | 1.116405195 | 0.109148593 | 4.731213 | 2.23E-06 | 2.57E-05 | ENSG00000214530 | STARD10 | 10809 |
| ENSG00000267106.7 | 120.5691547 | 1.116161128 | 0.163013311 | 3.166374 | 0.001544 | 0.01093444 | ENSG00000267106 | NA | NA |
| ENSG00000251247.11 | 68.48215 | 1.115834 | 0.1555808 | 3.315537 | 0.000915 | 0.006846597 | ENSG00000251247 | ZNF345 | 25850 |
| ENSG00000134278.15 | 2511.271063 | 1.114651006 | 0.097518328 | 5.27748 | 1.31E-07 | 1.78E-06 | ENSG00000134278 | SPIRE1 | 56907 |
| ENSG00000152904.11 | 762.9374234 | 1.114024434 | 0.089961143 | 5.71385 | 1.10E-08 | 1.70E-07 | ENSG00000152904 | GGPS1 | 9453 |
| ENSG00000168026.18 | 98.55791897 | 1.113423537 | 0.180358184 | 2.846688 | 0.004418 | 0.028075177 | ENSG00000168026 | TTC21A | 199223 |
| ENSG00000172936.14 | 3069.491468 | 1.113320421 | 0.14674303 | 3.498091 | 0.000469 | 0.003730598 | ENSG00000172936 | MYD88 | 4615 |
| ENSG00000198624.13 | 2631.406741 | 1.113295616 | 0.142571955 | 3.600257 | 0.000318 | 0.002608885 | ENSG00000198624 | CCDC69 | 26112 |
| ENSG00000173575.21 | 7356.018386 | 1.112289641 | 0.093642721 | 5.470683 | 4.48E-08 | 6.39E-07 | ENSG00000173575 | CHD2 | 1106 |
| ENSG00000196544.7 | 172.7165333 | 1.110572993 | 0.121544908 | 4.200694 | 2.66E-05 | 0.000262859 | ENSG00000196544 | BORCS6 | 54785 |
| ENSG00000162458.13 | 6021.969508 | 1.108647418 | 0.162053871 | 3.138755 | 0.001697 | 0.011911602 | ENSG00000162458 | FBLIM1 | 54751 |
| ENSG00000169247.12 | 1849.221073 | 1.108476106 | 0.088122087 | 5.770132 | 7.92E-09 | 1.24E-07 | ENSG00000169247 | SH3TC2 | 79628 |
| ENSG00000124882.4 | 15949.02181 | 1.108180968 | 0.10345719 | 4.911993 | 9.02E-07 | 1.10E-05 | ENSG00000124882 | EREG | 2069 |
| ENSG00000247081.8 | 173.7967106 | 1.108016212 | 0.140983511 | 3.603373 | 0.000314 | 0.002581767 | ENSG00000247081 | LOC10536 | 1.05E+08 |
| ENSG00000221994.10 | 81.75571 | 1.107955 | 0.1816617 | 2.796157 | 0.005171 | 0.03219686 | ENSG00000221994 | ZNF630 | 57232 |
| ENSG00000172893.15 | 3391.273995 | 1.107254194 | 0.143747807 | 3.528779 | 0.000417 | 0.003353715 | ENSG00000172893 | DHCR7 | 1717 |
| ENSG00000184209.14 | 861.2772551 | 1.106810005 | 0.111322307 | 4.552637 | 5.30E-06 | 5.83E-05 | ENSG00000184209 | SNRNP35 | 11066 |
| ENSG00000131732.12 | 3957.513453 | 1.104767165 | 0.147745368 | 3.416467 | 0.000634 | 0.004915259 | ENSG00000131732 | ZCCHC9 | 84240 |
| ENSG00000171219.9 | 1390.946068 | 1.1031212 | 0.156672066 | 3.211301 | 0.001321 | 0.009533082 | ENSG00000171219 | CDC42BP( | 55561 |
| ENSG00000140548.10 | 1395.787085 | 1.102118186 | 0.17828057 | 2.816449 | 0.004856 | 0.030466728 | ENSG00000140548 | ZNF710 | 374655 |
| ENSG00000113558.18 | 17127.21297 | 1.101994146 | 0.083310392 | 6.025589 | 1.68E-09 | 2.82E-08 | ENSG00000113558 | SKP1 | 6500 |
| ENSG00000184292.6 | 19225.50504 | 1.101152695 | 0.135998062 | 3.684999 | 0.000229 | 0.00192981 | ENSG00000184292 | TACSTD2 | 4070 |
| ENSG00000165813.20 | 2121.338499 | 1.100842839 | 0.11507595 | 4.352281 | 1.35E-05 | 0.000139508 | ENSG00000165813 | CCDC186 | 55088 |
| ENSG00000100417.12 | 826.7582 | 1.100395 | 0.120284 | 4.160112 | 3.18E-05 | 0.000310761 | ENSG00000100417 | PMM1 | 5372 |
| ENSG00000154803.13 | 851.9986068 | 1.09911611 | 0.175634194 | 2.841794 | 0.004486 | 0.028442698 | ENSG00000154803 | FLCN | 201163 |
| ENSG00000172086.8 | 1185.135964 | 1.097606819 | 0.126834647 | 3.923272 | 8.74E-05 | 0.000796421 | ENSG00000172086 | KRCC1 | 51315 |
| ENSG00000181191.11 | 1509.043 | 1.097131 | 0.08537241 | 5.823081 | 5.78E-09 | 9.16E-08 | ENSG00000181191 | PJA1 | 64219 |
| ENSG00000006756.16 | 354.7404 | 1.096317 | 0.1374877 | 3.609906 | 0.000306 | 0.002523153 | ENSG00000006756 | ARSD | 414 |
| ENSG00000230487.8 | 71.36046912 | 1.09612216 | 0.153430455 | 3.233531 | 0.001223 | 0.008884083 | ENSG00000230487 | PSMG3-A! | 114796 |
| ENSG00000141458.13 | 5851.587044 | 1.095970293 | 0.133750912 | 3.708164 | 0.000209 | 0.001772261 | ENSG00000141458 | NPC1 | 4864 |
| ENSG00000125868.16 | 21313.12 | 1.095962 | 0.05407894 | 9.171074 | 4.68E-20 | 1.79E-18 | ENSG00000125868 | DSTN | 11034 |
| ENSG00000135404.11 | 17740.31652 | 1.095440566 | 0.095661021 | 5.179127 | 2.23E-07 | 2.94E-06 | ENSG00000135404 | CD63 | 967 |
| ENSG00000231483.1 | 61.20199253 | 1.095264107 | 0.156167812 | 3.171358 | 0.001517 | 0.010765718 | ENSG00000231483 | NA | NA |
| ENSG00000286048.1 | 154.0293798 | 1.093946403 | 0.127783278 | 3.865501 | 0.000111 | 0.00099439 | ENSG00000286048 | NA | NA |
| ENSG00000172663.9 | 564.5321744 | 1.093037754 | 0.158703552 | 3.106659 | 0.001892 | 0.013121323 | ENSG00000172663 | TMEM134 | 80194 |

| | | | | | | | | | |
|---|---|---|---|---|---|---|---|---|---|
| ENSG00000161682.14 | 65.34445071 | 1.091141696 | 0.167303488 | 2.935633 | 0.003329 | 0.021794391 | ENSG00000161682 | FAM171A2 | 284069 |
| ENSG00000138593.9 | 2368.088458 | 1.089404225 | 0.115387954 | 4.241381 | 2.22E-05 | 0.000221999 | ENSG00000138593 | SECISBP2 | 9728 |
| ENSG00000089876.11 | 3372.306675 | 1.087123249 | 0.088540388 | 5.501707 | 3.76E-08 | 5.41E-07 | ENSG00000089876 | DHX32 | 55760 |
| ENSG00000167468.16 | 2241.494726 | 1.086773632 | 0.089097001 | 5.463412 | 4.67E-08 | 6.65E-07 | ENSG00000167468 | GPX4 | 2879 |
| ENSG00000213625.9 | 3300.70921 | 1.086638543 | 0.081697847 | 5.956565 | 2.58E-09 | 4.25E-08 | ENSG00000213625 | LEPROT | 54741 |
| ENSG00000231389.7 | 215.8263779 | 1.086113862 | 0.104859831 | 4.635844 | 3.55E-06 | 3.99E-05 | ENSG00000231389 | HLA-DPA1 | 3113 |
| ENSG00000135821.18 | 8393.352839 | 1.084931251 | 0.093255014 | 5.200056 | 1.99E-07 | 2.64E-06 | ENSG00000135821 | GLUL | 2752 |
| ENSG00000160685.13 | 2778.092204 | 1.084314104 | 0.138033122 | 3.50868 | 0.00045 | 0.003594786 | ENSG00000160685 | ZBTB7B | 51043 |
| ENSG00000166275.15 | 401.3321128 | 1.083293965 | 0.104640989 | 4.618591 | 3.86E-06 | 4.32E-05 | ENSG00000166275 | BORCS7 | 119032 |
| ENSG00000126524.10 | 6414.500537 | 1.080730538 | 0.078667206 | 6.11094 | 9.90E-10 | 1.70E-08 | ENSG00000126524 | SBDS | 51119 |
| ENSG00000126458.4 | 1294.649 | 1.08052 | 0.08770578 | 5.478776 | 4.28E-08 | 6.11E-07 | ENSG00000126458 | RRAS | 6237 |
| ENSG00000081087.15 | 3041.76573 | 1.080234394 | 0.108357879 | 4.431929 | 9.34E-06 | 9.96E-05 | ENSG00000081087 | OSTM1 | 28962 |
| ENSG00000139722.7 | 2156.964278 | 1.080079933 | 0.130898051 | 3.667587 | 0.000245 | 0.002056931 | ENSG00000139722 | VPS37B | 79720 |
| ENSG00000236753.7 | 238.5509193 | 1.07946558 | 0.146743858 | 3.267364 | 0.001086 | 0.007994263 | ENSG00000236753 | MKLN1-AS | 1.01E+08 |
| ENSG00000085433.15 | 1861.485668 | 1.078718666 | 0.097540648 | 4.907889 | 9.21E-07 | 1.12E-05 | ENSG00000085433 | WDR47 | 22911 |
| ENSG00000204054.14 | 332.6188828 | 1.076700602 | 0.101026591 | 4.718566 | 2.38E-06 | 2.72E-05 | ENSG00000204054 | NA | NA |
| ENSG00000156273.16 | 3742.039 | 1.076059 | 0.05643519 | 8.435502 | 3.30E-17 | 1.06E-15 | ENSG00000156273 | BACH1 | 571 |
| ENSG00000234465.11 | 193.4192 | 1.071245 | 0.102791 | 4.584492 | 4.55E-06 | 5.04E-05 | ENSG00000234465 | PINLYP | 390940 |
| ENSG00000272888.7 | 1348.979405 | 1.070650946 | 0.085702687 | 5.491671 | 3.98E-08 | 5.70E-07 | ENSG00000272888 | NA | NA |
| ENSG00000143178.13 | 74.05061163 | 1.068584758 | 0.156222658 | 2.999467 | 0.002705 | 0.018135962 | ENSG00000143178 | TBX19 | 9095 |
| ENSG00000073910.21 | 116.6003164 | 1.067174932 | 0.145524293 | 3.210288 | 0.001326 | 0.009558972 | ENSG00000073910 | FRY | 10129 |
| ENSG00000182621.18 | 306.7789 | 1.06422 | 0.1276678 | 3.636159 | 0.000277 | 0.002301532 | ENSG00000182621 | PLCB1 | 23236 |
| ENSG00000269743.3 | 110.8032 | 1.063876 | 0.1462181 | 3.172495 | 0.001511 | 0.01073262 | ENSG00000269743 | SLC25A53 | 401612 |
| ENSG00000121039.10 | 1269.543678 | 1.063532746 | 0.089318406 | 5.189667 | 2.11E-07 | 2.78E-06 | ENSG00000121039 | RDH10 | 157506 |
| ENSG00000141279.16 | 7541.78483 | 1.06342981 | 0.123962602 | 3.738465 | 0.000185 | 0.001588462 | ENSG00000141279 | NPEPPS | 9520 |
| ENSG00000140939.14 | 367.3641194 | 1.062073648 | 0.115808486 | 3.989981 | 6.61E-05 | 0.000614012 | ENSG00000140939 | NOL3 | 8996 |
| ENSG00000204438.11 | 191.656777 | 1.061272683 | 0.137739176 | 3.348885 | 0.000811 | 0.006152208 | ENSG00000204438 | GPANK1 | 7918 |
| ENSG00000168890.14 | 179.2059784 | 1.05911533 | 0.109423129 | 4.19578 | 2.72E-05 | 0.000268426 | ENSG00000168890 | TMEM150A | 129303 |
| ENSG00000130958.13 | 375.8610899 | 1.058058733 | 0.133734859 | 3.425126 | 0.000615 | 0.004772347 | ENSG00000130958 | SLC35D2 | 11046 |
| ENSG00000164609.10 | 4975.513607 | 1.057633322 | 0.122142199 | 3.746726 | 0.000179 | 0.001541544 | ENSG00000164609 | SLU7 | 10569 |
| ENSG00000271643.2 | 453.6495927 | 1.057107141 | 0.121407548 | 3.765064 | 0.000167 | 0.00144295 | ENSG00000271643 | NA | NA |
| ENSG00000269929.3 | 295.6261012 | 1.056658994 | 0.119728681 | 3.814115 | 0.000137 | 0.001203199 | ENSG00000269929 | NA | NA |
| ENSG00000197363.9 | 237.1812371 | 1.054875284 | 0.11958146 | 3.803895 | 0.000142 | 0.001249013 | ENSG00000197363 | ZNF517 | 340385 |
| ENSG00000129038.16 | 70.78710409 | 1.052083843 | 0.159731097 | 2.830281 | 0.004651 | 0.0293253 | ENSG00000129038 | LOXL1 | 4016 |
| ENSG00000068745.15 | 2569.198942 | 1.051728149 | 0.088854797 | 5.083892 | 3.70E-07 | 4.72E-06 | ENSG00000068745 | IP6K2 | 51447 |
| ENSG00000184574.10 | 315.7482428 | 1.049280012 | 0.105899795 | 4.242501 | 2.21E-05 | 0.000221059 | ENSG00000184574 | LPAR5 | 57121 |
| ENSG00000144063.3 | 173.0519362 | 1.048904375 | 0.167847649 | 2.674475 | 0.007485 | 0.044556318 | ENSG00000144063 | MALL | 7851 |
| ENSG00000128524.5 | 1915.148205 | 1.046989682 | 0.118252123 | 3.779972 | 0.000157 | 0.001366795 | ENSG00000128524 | ATP6V1F | 9296 |
| ENSG00000115216.14 | 6075.49008 | 1.04634178 | 0.075141494 | 5.940017 | 2.85E-09 | 4.68E-08 | ENSG00000115216 | NRBP1 | 29959 |
| ENSG00000103942.13 | 1196.528364 | 1.046108058 | 0.108525358 | 4.110634 | 3.95E-05 | 0.000380592 | ENSG00000103942 | HOMER2 | 9455 |
| ENSG00000141429.13 | 5802.548675 | 1.043059381 | 0.06763221 | 6.551011 | 5.71E-11 | 1.11E-09 | ENSG00000141429 | GALNT1 | 2589 |
| ENSG00000113916.18 | 406.3330969 | 1.042131171 | 0.100404433 | 4.403503 | 1.07E-05 | 0.000112083 | ENSG00000113916 | BCL6 | 604 |

| | | | | | | | | | |
|---|---|---|---|---|---|---|---|---|---|
| ENSG00000166579.15 | 3555.519025 | 1.039238796 | 0.123488622 | 3.556917 | 0.000375 | 0.003043704 | ENSG00000166579 | NDEL1 | 81565 |
| ENSG00000285867.1 | 266.9056197 | 1.037202613 | 0.102985254 | 4.245293 | 2.18E-05 | 0.00021857 | ENSG00000285867 | NA | NA |
| ENSG00000087884.14 | 100.4702117 | 1.034981816 | 0.158301515 | 2.747806 | 0.006 | 0.036674637 | ENSG00000087884 | AAMDC | 28971 |
| ENSG00000107864.15 | 280.2375863 | 1.03446735 | 0.140768508 | 3.086396 | 0.002026 | 0.013954946 | ENSG00000107864 | CPEB3 | 22849 |
| ENSG00000157483.8 | 6740.15583 | 1.034463857 | 0.136584587 | 3.180914 | 0.001468 | 0.010455041 | ENSG00000157483 | MYO1E | 4643 |
| ENSG00000070190.13 | 1672.094648 | 1.032378255 | 0.145773791 | 2.966091 | 0.003016 | 0.019983841 | ENSG00000070190 | DAPP1 | 27071 |
| ENSG00000280693.2 | 765.3845433 | 1.032165628 | 0.097748205 | 4.421213 | 9.81E-06 | 0.000104141 | ENSG00000280693 | SH3PXD2/ | 1.01E+08 |
| ENSG00000204991.11 | 231.7357842 | 1.031131691 | 0.141690761 | 3.042765 | 0.002344 | 0.015944313 | ENSG00000204991 | SPIRE2 | 84501 |
| ENSG00000067113.17 | 348.2096011 | 1.03078232 | 0.09135434 | 4.71551 | 2.41E-06 | 2.76E-05 | ENSG00000067113 | PLPP1 | 8611 |
| ENSG00000225548.7 | 372.0292784 | 1.027780726 | 0.109278562 | 3.91459 | 9.06E-05 | 0.000823373 | ENSG00000225548 | LINC0198( | 1.05E+08 |
| ENSG00000134108.13 | 4926.980578 | 1.027496729 | 0.119552197 | 3.575817 | 0.000349 | 0.002849389 | ENSG00000134108 | ARL8B | 55207 |
| ENSG00000172831.12 | 1756.714132 | 1.027164029 | 0.132919682 | 3.2137 | 0.00131 | 0.009463002 | ENSG00000172831 | CES2 | 8824 |
| ENSG00000174080.11 | 336.7275018 | 1.025448157 | 0.138065593 | 3.081493 | 0.00206 | 0.014175825 | ENSG00000174080 | CTSF | 8722 |
| ENSG00000043143.20 | 24112.93462 | 1.025127066 | 0.049397783 | 8.606197 | 7.55E-18 | 2.53E-16 | ENSG00000043143 | JADE2 | 23338 |
| ENSG00000198961.10 | 10711.6689 | 1.024829341 | 0.096135736 | 4.419057 | 9.91E-06 | 0.000105114 | ENSG00000198961 | PJA2 | 9867 |
| ENSG00000155085.15 | 267.2720567 | 1.023068309 | 0.107168698 | 3.947685 | 7.89E-05 | 0.000724897 | ENSG00000155085 | AK9 | 221264 |
| ENSG00000160948.14 | 1974.029287 | 1.021803962 | 0.129615958 | 3.25426 | 0.001137 | 0.008326305 | ENSG00000160948 | VPS28 | 51160 |
| ENSG00000144115.16 | 217.0771786 | 1.021670401 | 0.145069727 | 2.906674 | 0.003653 | 0.023661433 | ENSG00000144115 | THNSL2 | 55258 |
| ENSG00000116266.11 | 2561.53816 | 1.020195017 | 0.098695715 | 4.25748 | 2.07E-05 | 0.000207618 | ENSG00000116266 | STXBP3 | 6814 |
| ENSG00000162236.11 | 1463.731031 | 1.019770317 | 0.120549643 | 3.482137 | 0.000497 | 0.003941244 | ENSG00000162236 | STX5 | 6811 |
| ENSG00000168615.12 | 9405.652348 | 1.0197596 | 0.060651848 | 6.920805 | 4.49E-12 | 9.71E-11 | ENSG00000168615 | ADAM9 | 8754 |
| ENSG00000179832.17 | 2139.864648 | 1.019469507 | 0.130831904 | 3.206171 | 0.001345 | 0.009673224 | ENSG00000179832 | MROH1 | 727957 |
| ENSG00000086666.19 | 3837.466712 | 1.018842003 | 0.089219212 | 4.694527 | 2.67E-06 | 3.05E-05 | ENSG00000086666 | ZFAND6 | 54469 |
| ENSG00000169155.10 | 1752.156868 | 1.018113662 | 0.087800798 | 4.762071 | 1.92E-06 | 2.23E-05 | ENSG00000169155 | ZBTB43 | 23099 |
| ENSG00000168894.9 | 1963.665129 | 1.016905286 | 0.10210331 | 4.083171 | 4.44E-05 | 0.000424512 | ENSG00000168894 | RNF181 | 51255 |
| ENSG00000170776.21 | 3447.3421 | 1.016628905 | 0.110588909 | 3.767366 | 0.000165 | 0.001430642 | ENSG00000170776 | AKAP13 | 11214 |
| ENSG00000130772.14 | 212.7821657 | 1.015706255 | 0.125770318 | 3.305281 | 0.000949 | 0.007072728 | ENSG00000130772 | MED18 | 54797 |
| ENSG00000136021.18 | 6144.18562 | 1.015587398 | 0.111356569 | 3.732042 | 0.00019 | 0.001625335 | ENSG00000136021 | SCYL2 | 55681 |
| ENSG00000117139.17 | 5565.318082 | 1.015576038 | 0.079958074 | 5.197424 | 2.02E-07 | 2.68E-06 | ENSG00000117139 | KDM5B | 10765 |
| ENSG00000075945.13 | 2179.015722 | 1.015450898 | 0.127212589 | 3.2658 | 0.001092 | 0.008031871 | ENSG00000075945 | KIFAP3 | 22920 |
| ENSG00000170677.6 | 2115.566842 | 1.013123521 | 0.069200785 | 5.969925 | 2.37E-09 | 3.93E-08 | ENSG00000170677 | SOCS6 | 9306 |
| ENSG00000140598.15 | 2763.847022 | 1.011496875 | 0.085917596 | 4.789437 | 1.67E-06 | 1.96E-05 | ENSG00000140598 | EFL1 | 79631 |
| ENSG00000221838.9 | 577.0610597 | 1.009179454 | 0.121998476 | 3.353972 | 0.000797 | 0.006054014 | ENSG00000221838 | AP4M1 | 9179 |
| ENSG00000164924.18 | 105209.182 | 1.009048109 | 0.097511544 | 4.194869 | 2.73E-05 | 0.000269207 | ENSG00000164924 | YWHAZ | 7534 |
| ENSG00000006534.16 | 151.6554917 | 1.008795996 | 0.132045308 | 3.095877 | 0.001962 | 0.013572581 | ENSG00000006534 | ALDH3B1 | 221 |
| ENSG00000159256.13 | 4599.931 | 1.008402 | 0.09551326 | 4.275865 | 1.90E-05 | 0.000192071 | ENSG00000159256 | MORC3 | 23515 |
| ENSG00000075711.21 | 10055.46799 | 1.008245365 | 0.112873657 | 3.616835 | 0.000298 | 0.002464126 | ENSG00000075711 | DLG1 | 1739 |
| ENSG00000101745.17 | 3989.638817 | 1.008239114 | 0.100479192 | 4.062922 | 4.85E-05 | 0.000459951 | ENSG00000101745 | ANKRD12 | 23253 |
| ENSG00000101940.18 | 1969.579 | 1.005373 | 0.08017516 | 5.056089 | 4.28E-07 | 5.42E-06 | ENSG00000101940 | WDR13 | 64743 |
| ENSG00000112511.17 | 810.3443698 | 1.001774423 | 0.098742898 | 4.068894 | 4.72E-05 | 0.00044912 | ENSG00000112511 | PHF1 | 5252 |
| ENSG00000092020.10 | 835.7910277 | 1.000946842 | 0.112792036 | 3.554744 | 0.000378 | 0.003065232 | ENSG00000092020 | PPP2R3C | 55012 |
| ENSG00000247271.7 | 80.70633737 | 0.999074236 | 0.141886446 | 2.812631 | 0.004914 | 0.030791051 | ENSG00000247271 | ZBED5-AS | 729013 |

| | | | | | | | | | |
|---|---|---|---|---|---|---|---|---|---|
| ENSG00000041353.10 | 862.2171015 | 0.99845664 | 0.137264958 | 2.902829 | 0.003698 | 0.023889968 | ENSG00000041353 | RAB27B | 5874 |
| ENSG00000168216.13 | 2043.999243 | 0.99843877 | 0.108829268 | 3.661136 | 0.000251 | 0.002104112 | ENSG00000168216 | LMBRD1 | 55788 |
| ENSG00000109270.13 | 2171.895745 | 0.997579475 | 0.114374102 | 3.476132 | 0.000509 | 0.004019787 | ENSG00000109270 | LAMTOR3 | 8649 |
| ENSG00000160285.15 | 1929.329 | 0.997054 | 0.08761572 | 4.531767 | 5.85E-06 | 6.41E-05 | ENSG00000160285 | LSS | 4047 |
| ENSG00000104497.14 | 1289.175834 | 0.996758125 | 0.129586081 | 3.061734 | 0.002201 | 0.015071793 | ENSG00000104497 | SNX16 | 64089 |
| ENSG00000183578.7 | 143.7612862 | 0.996464865 | 0.132118812 | 3.000821 | 0.002693 | 0.018073764 | ENSG00000183578 | TNFAIP8L: | 388121 |
| ENSG00000196821.10 | 8638.949373 | 0.994755014 | 0.10242117 | 3.854233 | 0.000116 | 0.001037125 | ENSG00000196821 | ILRUN | 64771 |
| ENSG00000221926.12 | 979.0480168 | 0.992390586 | 0.104228675 | 3.764709 | 0.000167 | 0.001444531 | ENSG00000221926 | TRIM16 | 10626 |
| ENSG00000107262.22 | 2228.437618 | 0.991669301 | 0.107595547 | 3.6402 | 0.000272 | 0.002268545 | ENSG00000107262 | BAG1 | 573 |
| ENSG00000205707.10 | 279.2149517 | 0.991175812 | 0.126498105 | 3.092345 | 0.001986 | 0.013699517 | ENSG00000205707 | ETFRF1 | 144363 |
| ENSG00000166971.17 | 1415.293105 | 0.989507403 | 0.127500881 | 3.054939 | 0.002251 | 0.015385832 | ENSG00000166971 | AKTIP | 64400 |
| ENSG00000164039.15 | 285.0632419 | 0.988624956 | 0.094829192 | 4.098157 | 4.16E-05 | 0.00039938 | ENSG00000164039 | BDH2 | 56898 |
| ENSG00000108590.11 | 469.2806043 | 0.988391929 | 0.065311255 | 5.946784 | 2.73E-09 | 4.50E-08 | ENSG00000108590 | MED31 | 51003 |
| ENSG00000233369.7 | 575.3530258 | 0.986443128 | 0.104739363 | 3.689569 | 0.000225 | 0.001899082 | ENSG00000233369 | GTF2IP4 | 1E+08 |
| ENSG00000148803.12 | 179.8893968 | 0.985206867 | 0.103461983 | 3.723173 | 0.000197 | 0.001679214 | ENSG00000148803 | FUOM | 282969 |
| ENSG00000170310.15 | 914.5090846 | 0.982983954 | 0.107487714 | 3.563049 | 0.000367 | 0.002981637 | ENSG00000170310 | STX8 | 9482 |
| ENSG00000102038.15 | 159.9579 | 0.98172 | 0.1293489 | 2.951088 | 0.003167 | 0.02086128 | ENSG00000102038 | SMARCA1 | 6594 |
| ENSG00000197324.9 | 4897.6306 | 0.981430763 | 0.124683761 | 3.059186 | 0.002219 | 0.015184982 | ENSG00000197324 | LRP10 | 26020 |
| ENSG00000183826.18 | 757.52128 | 0.979485424 | 0.133271326 | 2.847465 | 0.004407 | 0.028020149 | ENSG00000183826 | BTBD9 | 114781 |
| ENSG00000138433.16 | 2045.98177 | 0.979232991 | 0.124270349 | 3.051677 | 0.002276 | 0.015530069 | ENSG00000138433 | CIR1 | 9541 |
| ENSG00000100319.12 | 236.2942 | 0.9788483 | 0.1189047 | 3.186152 | 0.001442 | 0.0102849 | ENSG00000100319 | ZMAT5 | 55954 |
| ENSG00000205155.8 | 414.1631 | 0.9784116 | 0.1145372 | 3.303832 | 0.000954 | 0.007102155 | ENSG00000205155 | PSENEN | 55851 |
| ENSG00000141741.12 | 954.43446 | 0.978315069 | 0.113375771 | 3.336825 | 0.000847 | 0.006392704 | ENSG00000141741 | MIEN1 | 84299 |
| ENSG00000091436.17 | 7797.72717 | 0.977920997 | 0.085760778 | 4.406688 | 1.05E-05 | 0.00011071 | ENSG00000091436 | MAP3K20 | 51776 |
| ENSG00000145022.4 | 306.6716721 | 0.976821739 | 0.114189619 | 3.299965 | 0.000967 | 0.007190572 | ENSG00000145022 | TCTA | 6988 |
| ENSG00000161791.14 | 1028.616418 | 0.976290779 | 0.097396898 | 3.863478 | 0.000112 | 0.001001316 | ENSG00000161791 | FMNL3 | 91010 |
| ENSG00000103363.14 | 5368.942236 | 0.976048611 | 0.063432501 | 5.928327 | 3.06E-09 | 5.00E-08 | ENSG00000103363 | ELOB | 6923 |
| ENSG00000265972.5 | 3074.171954 | 0.975181467 | 0.123811061 | 3.030274 | 0.002443 | 0.016568009 | ENSG00000265972 | TXNIP | 10628 |
| ENSG00000214226.9 | 143.2377419 | 0.973348108 | 0.139550526 | 2.675362 | 0.007465 | 0.044468511 | ENSG00000214226 | C17orf67 | 339210 |
| ENSG00000170185.9 | 2648.01191 | 0.973051985 | 0.093270179 | 3.999692 | 6.34E-05 | 0.000590591 | ENSG00000170185 | USP38 | 84640 |
| ENSG00000117984.14 | 2280.87766 | 0.97271558 | 0.122389155 | 3.045332 | 0.002324 | 0.015820982 | ENSG00000117984 | CTSD | 1509 |
| ENSG00000100485.12 | 1561.188621 | 0.971809406 | 0.085921117 | 4.327334 | 1.51E-05 | 0.000155054 | ENSG00000100485 | SOS2 | 6655 |
| ENSG00000072682.18 | 3782.433626 | 0.971682748 | 0.061828244 | 6.011537 | 1.84E-09 | 3.07E-08 | ENSG00000072682 | P4HA2 | 8974 |
| ENSG00000129521.13 | 187.1815776 | 0.970930768 | 0.138455018 | 2.679071 | 0.007383 | 0.044087434 | ENSG00000129521 | EGLN3 | 112399 |
| ENSG00000106853.20 | 3019.100461 | 0.970819489 | 0.060107127 | 6.16931 | 6.86E-10 | 1.20E-08 | ENSG00000106853 | PTGR1 | 22949 |
| ENSG00000161638.11 | 2985.465325 | 0.970364907 | 0.101709069 | 3.641415 | 0.000271 | 0.002259278 | ENSG00000161638 | ITGA5 | 3678 |
| ENSG00000210082.2 | 324384.7 | 0.9677987 | 0.07688312 | 4.783868 | 1.72E-06 | 2.01E-05 | ENSG00000210082 | NA | NA |
| ENSG00000157625.15 | 4308.711 | 0.967702 | 0.08377276 | 4.389279 | 1.14E-05 | 0.000119292 | ENSG00000157625 | TAB3 | 257397 |
| ENSG00000071859.15 | 1794.756 | 0.9667633 | 0.1091269 | 3.360889 | 0.000777 | 0.005912863 | ENSG00000071859 | FAM50A | 9130 |
| ENSG00000155366.16 | 4852.707315 | 0.965100168 | 0.089027723 | 4.100972 | 4.11E-05 | 0.000395179 | ENSG00000155366 | RHOC | 389 |
| ENSG00000100614.18 | 3696.147542 | 0.96442166 | 0.084077848 | 4.334336 | 1.46E-05 | 0.00015045 | ENSG00000100614 | PPM1A | 5494 |
| ENSG00000185158.12 | 363.1303874 | 0.96433682 | 0.099668861 | 3.655473 | 0.000257 | 0.002145711 | ENSG00000185158 | LRRC37B | 114659 |

| | | | | | | | | | |
|---|---|---|---|---|---|---|---|---|---|
| ENSG00000113595.15 | 2181.0945 | 0.964020486 | 0.118901939 | 3.061519 | 0.002202 | 0.015078773 | ENSG00000113595 | TRIM23 | 373 |
| ENSG00000164615.5 | 3645.039601 | 0.962483699 | 0.110678527 | 3.275104 | 0.001056 | 0.007791346 | ENSG00000164615 | CAMLG | 819 |
| ENSG00000163875.15 | 2825.586192 | 0.961641299 | 0.075648813 | 4.780528 | 1.75E-06 | 2.04E-05 | ENSG00000163875 | MEAF6 | 64769 |
| ENSG00000096060.14 | 5370.373242 | 0.961311328 | 0.099449626 | 3.633109 | 0.00028 | 0.002326738 | ENSG00000096060 | FKBP5 | 2289 |
| ENSG00000169612.4 | 386.7579001 | 0.960078024 | 0.103478181 | 3.479748 | 0.000502 | 0.003974177 | ENSG00000169612 | RAMAC | 83640 |
| ENSG00000115548.17 | 1313.465594 | 0.958314564 | 0.084023539 | 4.264455 | 2.00E-05 | 0.000201466 | ENSG00000115548 | KDM3A | 55818 |
| ENSG00000205763.13 | 388.710267 | 0.956998142 | 0.08136154 | 4.3878 | 1.15E-05 | 0.000119917 | ENSG00000205763 | RP9P | 441212 |
| ENSG00000214655.10 | 1492.542766 | 0.955193335 | 0.135048642 | 2.630114 | 0.008536 | 0.049908923 | ENSG00000214655 | ZSWIM8 | 23053 |
| ENSG00000117614.10 | 2701.803337 | 0.953300394 | 0.089675311 | 3.939773 | 8.16E-05 | 0.00074742 | ENSG00000117614 | SYF2 | 25949 |
| ENSG00000131435.13 | 3369.234091 | 0.952891931 | 0.081553803 | 4.327106 | 1.51E-05 | 0.000155054 | ENSG00000131435 | PDLIM4 | 8572 |
| ENSG00000071054.16 | 28481.27359 | 0.952114078 | 0.076079224 | 4.628255 | 3.69E-06 | 4.13E-05 | ENSG00000071054 | MAP4K4 | 9448 |
| ENSG00000186132.15 | 266.8904615 | 0.952054349 | 0.099845727 | 3.525983 | 0.000422 | 0.00338625 | ENSG00000186132 | C2orf76 | 130355 |
| ENSG00000256235.2 | 1858.769329 | 0.950917429 | 0.096933543 | 3.620186 | 0.000294 | 0.002434698 | ENSG00000256235 | SMIM3 | 85027 |
| ENSG00000170271.11 | 178.9289913 | 0.950503083 | 0.111350835 | 3.147736 | 0.001645 | 0.011582187 | ENSG00000170271 | FAXDC2 | 10826 |
| ENSG00000141401.12 | 1392.257521 | 0.949831619 | 0.104211572 | 3.356936 | 0.000788 | 0.005992896 | ENSG00000141401 | IMPA2 | 3613 |
| ENSG00000127824.14 | 6005.342173 | 0.94982779 | 0.081127046 | 4.312098 | 1.62E-05 | 0.000165137 | ENSG00000127824 | TUBA4A | 7277 |
| ENSG00000121931.16 | 1236.598017 | 0.94820427 | 0.084646208 | 4.113643 | 3.89E-05 | 0.000375936 | ENSG00000121931 | LRIF1 | 55791 |
| ENSG00000102554.14 | 6639.976148 | 0.946376171 | 0.107733883 | 3.215109 | 0.001304 | 0.009420288 | ENSG00000102554 | KLF5 | 688 |
| ENSG00000162976.13 | 580.1696697 | 0.945313203 | 0.108704587 | 3.17662 | 0.00149 | 0.010594898 | ENSG00000162976 | SLC66A3 | 130814 |
| ENSG00000183155.4 | 939.7832681 | 0.94145555 | 0.106380174 | 3.209767 | 0.001328 | 0.009573737 | ENSG00000183155 | RABIF | 5877 |
| ENSG00000158092.7 | 1436.989735 | 0.940988777 | 0.08290164 | 4.113173 | 3.90E-05 | 0.000376566 | ENSG00000158092 | NCK1 | 4690 |
| ENSG00000164023.14 | 2235.945195 | 0.940147255 | 0.067333172 | 5.051704 | 4.38E-07 | 5.54E-06 | ENSG00000164023 | SGMS2 | 166929 |
| ENSG00000166471.11 | 2249.365594 | 0.938035103 | 0.10929242 | 3.092942 | 0.001982 | 0.013682572 | ENSG00000166471 | TMEM41B | 440026 |
| ENSG00000203879.12 | 4490.358 | 0.9359949 | 0.1076282 | 3.121812 | 0.001797 | 0.01255927 | ENSG00000203879 | GDI1 | 2664 |
| ENSG00000170006.12 | 4592.862939 | 0.932723448 | 0.045367397 | 7.333977 | 2.23E-13 | 5.42E-12 | ENSG00000170006 | TMEM154 | 201799 |
| ENSG00000164117.13 | 785.003368 | 0.93149712 | 0.088863689 | 3.7304 | 0.000191 | 0.001634913 | ENSG00000164117 | FBXO8 | 26269 |
| ENSG00000229833.9 | 143.8306085 | 0.930940679 | 0.116546886 | 2.83955 | 0.004518 | 0.028581484 | ENSG00000229833 | PET100 | 1E+08 |
| ENSG00000169241.19 | 1303.313755 | 0.929033689 | 0.093696781 | 3.511686 | 0.000445 | 0.003558713 | ENSG00000169241 | SLC50A1 | 55974 |
| ENSG00000092841.18 | 24548.99965 | 0.928956388 | 0.106581197 | 3.086439 | 0.002026 | 0.013954946 | ENSG00000092841 | MYL6 | 4637 |
| ENSG00000116514.16 | 2084.233625 | 0.927970914 | 0.098302186 | 3.336354 | 0.000849 | 0.006399905 | ENSG00000116514 | RNF19B | 127544 |
| ENSG00000165355.7 | 1719.176656 | 0.927020756 | 0.067116741 | 4.872417 | 1.10E-06 | 1.32E-05 | ENSG00000165355 | FBXO33 | 254170 |
| ENSG00000160058.18 | 3147.387826 | 0.926841843 | 0.113616177 | 2.876719 | 0.004018 | 0.025795989 | ENSG00000160058 | BSDC1 | 55108 |
| ENSG00000165502.6 | 7747.712134 | 0.925605109 | 0.081455565 | 3.997334 | 6.41E-05 | 0.000595878 | ENSG00000165502 | RPL36AL | 6166 |
| ENSG00000065613.14 | 18277.69729 | 0.924932412 | 0.053294959 | 6.09687 | 1.08E-09 | 1.85E-08 | ENSG00000065613 | SLK | 9748 |
| ENSG00000182534.13 | 2672.435034 | 0.924743862 | 0.082132853 | 3.953885 | 7.69E-05 | 0.000707824 | ENSG00000182534 | MXRA7 | 439921 |
| ENSG00000088970.16 | 708.7545 | 0.9237472 | 0.06462568 | 5.009576 | 5.46E-07 | 6.82E-06 | ENSG00000088970 | KIZ | 55857 |
| ENSG00000176720.6 | 1700.822185 | 0.92368886 | 0.121877937 | 2.655845 | 0.007911 | 0.046780722 | ENSG00000176720 | BOK | 666 |
| ENSG00000155368.16 | 6551.180245 | 0.923007664 | 0.044650924 | 7.234065 | 4.69E-13 | 1.10E-11 | ENSG00000155368 | DBI | 1622 |
| ENSG00000168575.10 | 3590.141287 | 0.922585682 | 0.116513459 | 2.768656 | 0.005629 | 0.034692436 | ENSG00000168575 | SLC20A2 | 6575 |
| ENSG00000163110.15 | 6765.475194 | 0.922574717 | 0.060025982 | 5.373918 | 7.70E-08 | 1.07E-06 | ENSG00000163110 | PDLIM5 | 10611 |
| ENSG00000164603.12 | 558.2443232 | 0.921764963 | 0.090140065 | 3.569611 | 0.000358 | 0.002915956 | ENSG00000164603 | BMT2 | 154743 |
| ENSG00000117533.15 | 701.1050827 | 0.920340985 | 0.093690722 | 3.419132 | 0.000628 | 0.004870194 | ENSG00000117533 | VAMP4 | 8674 |

| | | | | | | | | | |
|---|---|---|---|---|---|---|---|---|---|
| ENSG00000227036.8 | 353.757001 | 0.918386363 | 0.106683537 | 2.9844 | 0.002841 | 0.018948498 | ENSG00000227036 | LINC0051 | 400619 |
| ENSG00000140022.13 | 1123.209972 | 0.917673383 | 0.081419422 | 3.90169 | 9.55E-05 | 0.000866151 | ENSG00000140022 | STON2 | 85439 |
| ENSG00000119899.13 | 965.0768709 | 0.915625001 | 0.083502993 | 3.779805 | 0.000157 | 0.001367265 | ENSG00000119899 | SLC17A5 | 26503 |
| ENSG00000005486.17 | 969.7602343 | 0.915276478 | 0.05693407 | 5.537571 | 3.07E-08 | 4.49E-07 | ENSG00000005486 | RHBDD2 | 57414 |
| ENSG00000100532.12 | 664.8853692 | 0.914962896 | 0.100098085 | 3.146543 | 0.001652 | 0.011623427 | ENSG00000100532 | CGRRF1 | 10668 |
| ENSG00000111911.7 | 1203.675594 | 0.914866995 | 0.118669067 | 2.65332 | 0.00797 | 0.047048511 | ENSG00000111911 | HINT3 | 135114 |
| ENSG00000184924.5 | 310.7918287 | 0.913547016 | 0.118047248 | 2.656115 | 0.007905 | 0.046756766 | ENSG00000184924 | PTRHD1 | 391356 |
| ENSG00000179523.5 | 271.5912213 | 0.913320177 | 0.087826595 | 3.567486 | 0.00036 | 0.002936095 | ENSG00000179523 | EIF3J-DT | 645212 |
| ENSG00000154217.15 | 1370.772004 | 0.913247397 | 0.104106828 | 3.008903 | 0.002622 | 0.017671104 | ENSG00000154217 | PITPNC1 | 26207 |
| ENSG00000154813.10 | 1614.663549 | 0.912723293 | 0.088805734 | 3.521431 | 0.000429 | 0.003438699 | ENSG00000154813 | DPH3 | 285381 |
| ENSG00000076513.16 | 3122.945308 | 0.91196814 | 0.105869999 | 2.94671 | 0.003212 | 0.021122314 | ENSG00000076513 | ANKRD13/ | 88455 |
| ENSG00000178761.15 | 583.3021258 | 0.910209504 | 0.089194634 | 3.477894 | 0.000505 | 0.003997012 | ENSG00000178761 | FAM219B | 57184 |
| ENSG00000188321.13 | 203.3082389 | 0.909616649 | 0.098599356 | 3.140149 | 0.001689 | 0.011858204 | ENSG00000188321 | ZNF559 | 84527 |
| ENSG00000152518.8 | 4091.88441 | 0.907223793 | 0.082757752 | 3.712326 | 0.000205 | 0.001746523 | ENSG00000152518 | ZFP36L2 | 678 |
| ENSG00000131584.19 | 1242.414792 | 0.904668609 | 0.064201926 | 4.745475 | 2.08E-06 | 2.40E-05 | ENSG00000131584 | ACAP3 | 116983 |
| ENSG00000130818.12 | 953.4895373 | 0.90431 | 0.092729767 | 3.281686 | 0.001032 | 0.007630779 | ENSG00000130818 | ZNF426 | 79088 |
| ENSG00000169599.12 | 1493.734971 | 0.901995826 | 0.09051518 | 3.336411 | 0.000849 | 0.006399905 | ENSG00000169599 | NFU1 | 27247 |
| ENSG00000137414.6 | 1575.26794 | 0.899726585 | 0.099088834 | 3.024827 | 0.002488 | 0.016835021 | ENSG00000137414 | FAM8A1 | 51439 |
| ENSG00000165424.6 | 171.246356 | 0.89888246 | 0.098068599 | 3.047688 | 0.002306 | 0.015705514 | ENSG00000165424 | ZCCHC24 | 219654 |
| ENSG00000051128.19 | 1422.15 | 0.8985645 | 0.1106001 | 2.699495 | 0.006944 | 0.04173267 | ENSG00000051128 | HOMER3 | 9454 |
| ENSG00000143653.10 | 3181.569969 | 0.898053546 | 0.112668254 | 2.645408 | 0.008159 | 0.048045833 | ENSG00000143653 | SCCPDH | 51097 |
| ENSG00000144746.7 | 2666.412097 | 0.895688494 | 0.066961527 | 4.415797 | 1.01E-05 | 0.000106529 | ENSG00000144746 | ARL6IP5 | 10550 |
| ENSG00000143222.12 | 2562.059337 | 0.892632188 | 0.069493121 | 4.210952 | 2.54E-05 | 0.000251762 | ENSG00000143222 | UFC1 | 51506 |
| ENSG00000075651.16 | 213.8897187 | 0.891158409 | 0.099334994 | 2.931076 | 0.003378 | 0.022084114 | ENSG00000075651 | PLD1 | 5337 |
| ENSG00000188554.14 | 6440.451724 | 0.889338689 | 0.087134855 | 3.320585 | 0.000898 | 0.006742114 | ENSG00000188554 | NBR1 | 4077 |
| ENSG00000163393.13 | 689.5358361 | 0.887982772 | 0.088685032 | 3.247253 | 0.001165 | 0.008505944 | ENSG00000163393 | SLC22A15 | 55356 |
| ENSG00000134996.11 | 1905.047825 | 0.887869152 | 0.10811806 | 2.662545 | 0.007755 | 0.045951422 | ENSG00000134996 | OSTF1 | 26578 |
| ENSG00000149948.13 | 13115.57229 | 0.887496039 | 0.038176376 | 7.530731 | 5.05E-14 | 1.28E-12 | ENSG00000149948 | HMGA2 | 8091 |
| ENSG00000224032.7 | 1599.812171 | 0.887397539 | 0.066241935 | 4.338604 | 1.43E-05 | 0.000147902 | ENSG00000224032 | NA | NA |
| ENSG00000133935.6 | 1267.541269 | 0.886872097 | 0.105420714 | 2.721212 | 0.006504 | 0.039389818 | ENSG00000133935 | ERG28 | 11161 |
| ENSG00000197579.8 | 2731.995125 | 0.886060667 | 0.050512947 | 5.663116 | 1.49E-08 | 2.25E-07 | ENSG00000197579 | TOPORS | 10210 |
| ENSG00000106789.13 | 603.6131027 | 0.882628746 | 0.094050558 | 3.005073 | 0.002655 | 0.017845514 | ENSG00000106789 | CORO2A | 7464 |
| ENSG00000022267.18 | 3503.371 | 0.8824178 | 0.09671953 | 2.919967 | 0.003501 | 0.02279557 | ENSG00000022267 | FHL1 | 2273 |
| ENSG00000158109.15 | 2472.708132 | 0.882129931 | 0.098413915 | 2.866769 | 0.004147 | 0.026525156 | ENSG00000158109 | TPRG1L | 127262 |
| ENSG00000111269.3 | 1920.490951 | 0.881350251 | 0.073801028 | 3.812281 | 0.000138 | 0.001209854 | ENSG00000111269 | CREBL2 | 1389 |
| ENSG00000062716.13 | 9089.490483 | 0.880526897 | 0.088617421 | 3.165595 | 0.001548 | 0.010949184 | ENSG00000062716 | VMP1 | 81671 |
| ENSG00000180098.9 | 1600.05877 | 0.879433438 | 0.084280816 | 3.315505 | 0.000915 | 0.006846597 | ENSG00000180098 | TRNAU1AI | 54952 |
| ENSG00000040487.13 | 529.3754847 | 0.879329309 | 0.095671243 | 2.919679 | 0.003504 | 0.022807192 | ENSG00000040487 | SLC66A1 | 54896 |
| ENSG00000167671.12 | 3062.492058 | 0.879210042 | 0.086054672 | 3.244566 | 0.001176 | 0.008572521 | ENSG00000167671 | UBXN6 | 80700 |
| ENSG00000108799.13 | 1573.721473 | 0.879067458 | 0.105123304 | 2.654668 | 0.007939 | 0.046902533 | ENSG00000108799 | EZH1 | 2145 |
| ENSG00000057935.13 | 2180.386418 | 0.878259063 | 0.092590905 | 3.005253 | 0.002654 | 0.017840514 | ENSG00000057935 | MTA3 | 57504 |
| ENSG00000159346.13 | 5842.60493 | 0.877933029 | 0.079554551 | 3.493616 | 0.000477 | 0.003788022 | ENSG00000159346 | ADIPOR1 | 51094 |

| | | | | | | | | | |
|---|---|---|---|---|---|---|---|---|---|
| ENSG00000115520.8 | 1966.358783 | 0.876246283 | 0.100658496 | 2.744391 | 0.006062 | 0.036990888 | ENSG00000115520 | COQ10B | 80219 |
| ENSG00000172331.12 | 771.9892598 | 0.874102238 | 0.094737621 | 2.893278 | 0.003812 | 0.02455124 | ENSG00000172331 | BPGM | 669 |
| ENSG00000176148.16 | 711.4359645 | 0.873861047 | 0.099657312 | 2.748028 | 0.005995 | 0.036658724 | ENSG00000176148 | TCP11L1 | 55346 |
| ENSG00000177058.12 | 2083.169691 | 0.873802486 | 0.076795509 | 3.565345 | 0.000363 | 0.002957786 | ENSG00000177058 | SLC38A9 | 153129 |
| ENSG00000183726.11 | 5291.816633 | 0.873008739 | 0.062916535 | 4.339221 | 1.43E-05 | 0.000147602 | ENSG00000183726 | TMEM50A | 23585 |
| ENSG00000153561.13 | 3759.382686 | 0.871914547 | 0.036406003 | 7.468948 | 8.08E-14 | 2.02E-12 | ENSG00000153561 | RMND5A | 64795 |
| ENSG00000075413.18 | 6128.070746 | 0.871173424 | 0.052201446 | 5.194749 | 2.05E-07 | 2.71E-06 | ENSG00000075413 | MARK3 | 4140 |
| ENSG00000171867.16 | 16683.71 | 0.8709831 | 0.05263514 | 5.14833 | 2.63E-07 | 3.43E-06 | ENSG00000171867 | PRNP | 5621 |
| ENSG00000108306.13 | 907.8762468 | 0.870243339 | 0.097727219 | 2.765282 | 0.005687 | 0.035007818 | ENSG00000108306 | FBXL20 | 84961 |
| ENSG00000078618.21 | 13180.35016 | 0.869221718 | 0.097771869 | 2.75357 | 0.005895 | 0.036126679 | ENSG00000078618 | NRDC | 4898 |
| ENSG00000139625.13 | 257.2198002 | 0.868982835 | 0.082690533 | 3.252886 | 0.001142 | 0.008357473 | ENSG00000139625 | MAP3K12 | 7786 |
| ENSG00000196387.9 | 302.2074783 | 0.868684019 | 0.093715141 | 2.867029 | 0.004143 | 0.026509725 | ENSG00000196387 | ZNF140 | 7699 |
| ENSG00000145779.8 | 2983.354849 | 0.868336296 | 0.10021542 | 2.677595 | 0.007415 | 0.044245163 | ENSG00000145779 | TNFAIP8 | 25816 |
| ENSG00000286966.1 | 2631.164981 | 0.867246959 | 0.065470043 | 4.081973 | 4.47E-05 | 0.000426553 | ENSG00000286966 | NA | NA |
| ENSG00000115649.16 | 1659.812509 | 0.866432261 | 0.078909952 | 3.376409 | 0.000734 | 0.005624525 | ENSG00000115649 | CNPPD1 | 27013 |
| ENSG00000153558.15 | 589.5370907 | 0.866251278 | 0.082057672 | 3.244685 | 0.001176 | 0.008571286 | ENSG00000153558 | FBXL2 | 25827 |
| ENSG00000159720.12 | 3684.810663 | 0.863978695 | 0.094910721 | 2.781337 | 0.005414 | 0.033555197 | ENSG00000159720 | ATP6V0D1 | 9114 |
| ENSG00000147533.16 | 676.7590293 | 0.861150078 | 0.07727683 | 3.37941 | 0.000726 | 0.005567841 | ENSG00000147533 | GOLGA7 | 51125 |
| ENSG00000114670.14 | 256.3752985 | 0.860588247 | 0.092794277 | 2.808236 | 0.004981 | 0.031166619 | ENSG00000114670 | NEK11 | 79858 |
| ENSG00000115685.15 | 2231.83787 | 0.858638415 | 0.08905085 | 2.90439 | 0.00368 | 0.023794213 | ENSG00000115685 | PPP1R7 | 5510 |
| ENSG00000142546.13 | 4566.984 | 0.8586103 | 0.09594934 | 2.69528 | 0.007033 | 0.04222005 | ENSG00000142546 | NOSIP | 51070 |
| ENSG00000178913.7 | 3303.451368 | 0.856700702 | 0.078623438 | 3.264939 | 0.001095 | 0.008047443 | ENSG00000178913 | TAF7 | 6879 |
| ENSG00000102897.10 | 667.014025 | 0.855340264 | 0.081672649 | 3.126386 | 0.00177 | 0.012391541 | ENSG00000102897 | LYRM1 | 57149 |
| ENSG00000121749.15 | 2250.614126 | 0.855018903 | 0.093275477 | 2.73404 | 0.006256 | 0.038057786 | ENSG00000121749 | TBC1D15 | 64786 |
| ENSG00000136378.15 | 248.15942 | 0.854767784 | 0.093146669 | 2.735125 | 0.006236 | 0.037952839 | ENSG00000136378 | ADAMTS7 | 11173 |
| ENSG00000141985.9 | 3434.837469 | 0.852568617 | 0.081611341 | 3.094774 | 0.00197 | 0.013616087 | ENSG00000141985 | SH3GL1 | 6455 |
| ENSG00000100744.15 | 2092.058045 | 0.852119835 | 0.090830976 | 2.775703 | 0.005508 | 0.034054928 | ENSG00000100744 | GSKIP | 51527 |
| ENSG00000067167.8 | 12286.59762 | 0.85165622 | 0.068847568 | 3.655267 | 0.000257 | 0.002146688 | ENSG00000067167 | TRAM1 | 23471 |
| ENSG00000205352.11 | 1641.645972 | 0.848555358 | 0.084903879 | 2.927491 | 0.003417 | 0.022296495 | ENSG00000205352 | PRR13 | 54458 |
| ENSG00000188706.13 | 1957.54 | 0.8468924 | 0.08177321 | 3.019234 | 0.002534 | 0.01711851 | ENSG00000188706 | ZDHHC9 | 51114 |
| ENSG00000105993.15 | 8186.319028 | 0.84236111 | 0.053160657 | 4.559032 | 5.14E-06 | 5.66E-05 | ENSG00000105993 | DNAJB6 | 10049 |
| ENSG00000165775.18 | 1828.743 | 0.8379001 | 0.05837308 | 4.07551 | 4.59E-05 | 0.00043779 | ENSG00000165775 | FUNDC2 | 65991 |
| ENSG00000103671.9 | 1359.705892 | 0.836054448 | 0.08082562 | 2.92054 | 0.003494 | 0.022766004 | ENSG00000103671 | TRIP4 | 9325 |
| ENSG00000089289.16 | 1900.909 | 0.8332376 | 0.08544882 | 2.729559 | 0.006342 | 0.0385115 | ENSG00000089289 | IGBP1 | 3476 |
| ENSG00000026652.14 | 664.4959823 | 0.832259237 | 0.080688348 | 2.878473 | 0.003996 | 0.025659178 | ENSG00000026652 | AGPAT4 | 56895 |
| ENSG00000196754.11 | 141757.926 | 0.832186587 | 0.062590905 | 3.70959 | 0.000208 | 0.001763997 | ENSG00000196754 | S100A2 | 6273 |
| ENSG00000197712.12 | 2286.538201 | 0.830744364 | 0.078515602 | 2.938835 | 0.003294 | 0.021575862 | ENSG00000197712 | FAM114A1 | 92689 |
| ENSG00000075790.10 | 2634.102601 | 0.830024493 | 0.052557749 | 4.376605 | 1.21E-05 | 0.000125896 | ENSG00000075790 | BCAP29 | 55973 |
| ENSG00000063660.9 | 2680.728175 | 0.827032385 | 0.066968333 | 3.390145 | 0.000699 | 0.005371736 | ENSG00000063660 | GPC1 | 2817 |
| ENSG00000122490.19 | 1721.302792 | 0.826744274 | 0.084771074 | 2.674784 | 0.007478 | 0.044525316 | ENSG00000122490 | SLC66A2 | 80148 |
| ENSG00000266412.5 | 12906.36416 | 0.825860697 | 0.056870362 | 3.971501 | 7.14E-05 | 0.000659742 | ENSG00000266412 | NCOA4 | 8031 |
| ENSG00000166478.10 | 1657.71359 | 0.825419105 | 0.048131684 | 4.683383 | 2.82E-06 | 3.21E-05 | ENSG00000166478 | ZNF143 | 7702 |

| | | | | | | | | | |
|---|---|---|---|---|---|---|---|---|---|
| ENSG00000068305.17 | 2546.731226 | 0.824781801 | 0.061278071 | 3.668226 | 0.000244 | 0.002053092 | ENSG00000068305 | MEF2A | 4205 |
| ENSG00000023697.13 | 1290.465011 | 0.823355573 | 0.074286391 | 3.006682 | 0.002641 | 0.017769237 | ENSG00000023697 | DERA | 51071 |
| ENSG00000131507.11 | 9530.219837 | 0.822867944 | 0.082418373 | 2.704105 | 0.006849 | 0.041260557 | ENSG00000131507 | NDFIP1 | 80762 |
| ENSG00000198911.12 | 6118.867 | 0.8225057 | 0.04826701 | 4.609891 | 4.03E-06 | 4.49E-05 | ENSG00000198911 | SREBF2 | 6721 |
| ENSG00000130449.6 | 1876.944867 | 0.818891231 | 0.068815116 | 3.18086 | 0.001468 | 0.010455041 | ENSG00000130449 | ZSWIM6 | 57688 |
| ENSG00000032219.18 | 1683.522661 | 0.818251496 | 0.077190506 | 2.82744 | 0.004692 | 0.02957277 | ENSG00000032219 | ARID4A | 5926 |
| ENSG00000123395.14 | 1243.161197 | 0.817790109 | 0.079880859 | 2.726437 | 0.006402 | 0.038815752 | ENSG00000123395 | ATG101 | 60673 |
| ENSG00000141447.18 | 1436.700967 | 0.817657935 | 0.053601903 | 4.060638 | 4.89E-05 | 0.000463729 | ENSG00000141447 | OSBPL1A | 114876 |
| ENSG00000170955.10 | 572.3586844 | 0.815845717 | 0.082065383 | 2.630168 | 0.008534 | 0.049908923 | ENSG00000170955 | CAVIN3 | 112464 |
| ENSG00000204178.11 | 1566.072792 | 0.814457429 | 0.070144967 | 3.057346 | 0.002233 | 0.015270648 | ENSG00000204178 | MACO1 | 55219 |
| ENSG00000015153.14 | 899.7928521 | 0.809577539 | 0.073095594 | 2.867171 | 0.004142 | 0.026504227 | ENSG00000015153 | YAF2 | 10138 |
| ENSG00000133961.20 | 5259.037663 | 0.807044726 | 0.048471058 | 4.271512 | 1.94E-05 | 0.000195562 | ENSG00000133961 | NUMB | 8650 |
| ENSG00000182149.20 | 6255.371515 | 0.805744706 | 0.056765119 | 3.624492 | 0.00029 | 0.002400845 | ENSG00000182149 | IST1 | 9798 |
| ENSG00000173545.5 | 1867.514721 | 0.798220883 | 0.066507 | 2.980451 | 0.002878 | 0.019180095 | ENSG00000173545 | ZNF622 | 90441 |
| ENSG00000144048.10 | 2544.706791 | 0.797816386 | 0.071426311 | 2.769517 | 0.005614 | 0.034619989 | ENSG00000144048 | DUSP11 | 8446 |
| ENSG00000169242.12 | 1741.848662 | 0.796698645 | 0.065862616 | 2.986499 | 0.002822 | 0.018837778 | ENSG00000169242 | EFNA1 | 1942 |
| ENSG00000166272.18 | 1918.483944 | 0.793648449 | 0.069535835 | 2.784873 | 0.005355 | 0.033231224 | ENSG00000166272 | WBP1L | 54838 |
| ENSG00000013561.18 | 3908.188608 | 0.791759476 | 0.047323094 | 4.052133 | 5.08E-05 | 0.000479128 | ENSG00000013561 | RNF14 | 9604 |
| ENSG00000250682.6 | 429.1238312 | 0.790527637 | 0.071174859 | 2.676895 | 0.007431 | 0.044305134 | ENSG00000250682 | LINC0049 | 285708 |
| ENSG00000187231.14 | 1085.85373 | 0.788749404 | 0.065985208 | 2.860481 | 0.00423 | 0.026992071 | ENSG00000187231 | SESTD1 | 91404 |
| ENSG00000171503.11 | 937.9139309 | 0.785531691 | 0.063454636 | 2.923848 | 0.003457 | 0.022537038 | ENSG00000171503 | ETFDH | 2110 |
| ENSG00000179532.12 | 565.7702726 | 0.784210991 | 0.069384255 | 2.654939 | 0.007932 | 0.046875205 | ENSG00000179532 | DNHD1 | 144132 |
| ENSG00000109854.13 | 2675.681691 | 0.778844398 | 0.066339969 | 2.695877 | 0.00702 | 0.042160157 | ENSG00000109854 | HTATIP2 | 10553 |
| ENSG00000139624.12 | 2347.394149 | 0.776898707 | 0.052070814 | 3.397272 | 0.000681 | 0.005247404 | ENSG00000139624 | CERS5 | 91012 |
| ENSG00000137575.12 | 14247.657 | 0.774612368 | 0.064864938 | 2.691938 | 0.007104 | 0.042584546 | ENSG00000137575 | SDCBP | 6386 |
| ENSG00000125089.17 | 2797.765857 | 0.771706625 | 0.06198034 | 2.77034 | 0.0056 | 0.034556627 | ENSG00000125089 | SH3TC1 | 54436 |
| ENSG00000150403.18 | 2790.148812 | 0.767015779 | 0.057647281 | 2.897201 | 0.003765 | 0.024299144 | ENSG00000150403 | TMCO3 | 55002 |
| ENSG00000109762.16 | 1561.454719 | 0.765545575 | 0.055537755 | 2.980775 | 0.002875 | 0.019164612 | ENSG00000109762 | SNX25 | 83891 |
| ENSG00000150459.12 | 6530.659569 | 0.753920147 | 0.053146985 | 2.896122 | 0.003778 | 0.024365166 | ENSG00000150459 | SAP18 | 10284 |
| ENSG00000083799.17 | 2532.112749 | 0.753592778 | 0.058300074 | 2.634521 | 0.008426 | 0.049374083 | ENSG00000083799 | CYLD | 1540 |
| ENSG00000107771.17 | 3148.456356 | 0.741428824 | 0.052549384 | 2.691351 | 0.007116 | 0.04264034 | ENSG00000107771 | CCSER2 | 54462 |
| ENSG00000117868.16 | 6301.083668 | 0.737625483 | 0.03811924 | 3.610394 | 0.000306 | 0.002519876 | ENSG00000117868 | ESYT2 | 57488 |
| ENSG00000182022.18 | 2415.289426 | 0.736229856 | 0.047459806 | 2.870426 | 0.004099 | 0.026264429 | ENSG00000182022 | CHST15 | 51363 |
| ENSG00000187164.20 | 2612.854831 | -0.734105397 | 0.044406963 | -3.01992 | 0.002528 | 0.017092902 | ENSG00000187164 | SHTN1 | 57698 |
| ENSG00000162302.13 | 4136.089089 | -0.745735523 | 0.045103921 | -3.23111 | 0.001233 | 0.008947464 | ENSG00000162302 | RPS6KA4 | 8986 |
| ENSG00000108106.14 | 5857.806 | -0.7470433 | 0.04663242 | -3.15324 | 0.001615 | 0.01137795 | ENSG00000108106 | UBE2S | 27338 |
| ENSG00000139289.13 | 18108.07854 | -0.769055752 | 0.04620845 | -3.65855 | 0.000254 | 0.002123481 | ENSG00000139289 | PHLDA1 | 22822 |
| ENSG00000139579.13 | 1653.53051 | -0.775495548 | 0.065373119 | -2.68452 | 0.007263 | 0.043433382 | ENSG00000139579 | NABP2 | 79035 |
| ENSG00000087253.13 | 1405.737116 | -0.775808975 | 0.040953066 | -4.29294 | 1.76E-05 | 0.000178891 | ENSG00000087253 | LPCAT2 | 54947 |
| ENSG00000150938.10 | 5658.992732 | -0.779081067 | 0.056203829 | -3.18628 | 0.001441 | 0.010284902 | ENSG00000150938 | CRIM1 | 51232 |
| ENSG00000140853.15 | 1114.753379 | -0.782299658 | 0.055802776 | -3.26686 | 0.001087 | 0.008004178 | ENSG00000140853 | NLRC5 | 84166 |
| ENSG00000116213.16 | 773.1429238 | -0.78306804 | 0.067656129 | -2.70586 | 0.006813 | 0.041061693 | ENSG00000116213 | WRAP73 | 49856 |

| | | | | | | | | | |
|---|---|---|---|---|---|---|---|---|---|
| ENSG00000165804.16 | 2257.085716 | -0.785104803 | 0.057897818 | -3.19709 | 0.001388 | 0.009945235 | ENSG00000165804 | ZNF219 | 51222 |
| ENSG00000160710.17 | 11153.69294 | -0.785177259 | 0.05234132 | -3.53788 | 0.000403 | 0.00324905 | ENSG00000160710 | ADAR | 103 |
| ENSG00000148468.17 | 561.3179146 | -0.785971096 | 0.069843907 | -2.66267 | 0.007752 | 0.045944876 | ENSG00000148468 | FAM171A1 | 221061 |
| ENSG00000166822.13 | 1065.45014 | -0.78814053 | 0.059123262 | -3.18217 | 0.001462 | 0.010413268 | ENSG00000166822 | TMEM170A | 124491 |
| ENSG00000106462.10 | 1877.938405 | -0.79159285 | 0.064723141 | -2.96019 | 0.003074 | 0.020330089 | ENSG00000106462 | EZH2 | 2146 |
| ENSG00000141480.18 | 1021.07925 | -0.793782541 | 0.061194306 | -3.16668 | 0.001542 | 0.01092694 | ENSG00000141480 | ARRB2 | 409 |
| ENSG00000188157.15 | 9531.710141 | -0.795432694 | 0.070377885 | -2.7769 | 0.005488 | 0.033960832 | ENSG00000188157 | AGRN | 375790 |
| ENSG00000085276.18 | 509.2096506 | -0.803298389 | 0.074219682 | -2.73914 | 0.00616 | 0.037552081 | ENSG00000085276 | MECOM | 2122 |
| ENSG00000235750.10 | 1172.939089 | -0.811169496 | 0.079668984 | -2.65059 | 0.008035 | 0.047399469 | ENSG00000235750 | KIAA0040 | 9674 |
| ENSG00000142089.16 | 5925.022063 | -0.826970448 | 0.047492145 | -4.77912 | 1.76E-06 | 2.05E-05 | ENSG00000142089 | IFITM3 | 10410 |
| ENSG00000181873.13 | 847.4816583 | -0.828660095 | 0.077622146 | -2.94581 | 0.003221 | 0.021178605 | ENSG00000181873 | IBA57 | 200205 |
| ENSG00000213699.9 | 1450.583651 | -0.829066983 | 0.059246994 | -3.86631 | 0.00011 | 0.000991452 | ENSG00000213699 | SLC35F6 | 54978 |
| ENSG00000092010.15 | 6095.989806 | -0.829672526 | 0.049223158 | -4.66594 | 3.07E-06 | 3.47E-05 | ENSG00000092010 | PSME1 | 5720 |
| ENSG00000204264.11 | 2464.341798 | -0.830250689 | 0.073236259 | -3.14394 | 0.001667 | 0.01172095 | ENSG00000204264 | PSMB8 | 5696 |
| ENSG00000156521.14 | 1034.949389 | -0.831135642 | 0.061795093 | -3.74036 | 0.000184 | 0.001577573 | ENSG00000156521 | TYSND1 | 219743 |
| ENSG00000183647.10 | 298.4932 | -0.8314728 | 0.08581438 | -2.69737 | 0.006989 | 0.04199101 | ENSG00000183647 | ZNF530 | 348327 |
| ENSG00000101596.15 | 4995.565502 | -0.833419529 | 0.081787375 | -2.85398 | 0.004318 | 0.027524225 | ENSG00000101596 | SMCHD1 | 23347 |
| ENSG00000067057.17 | 5926.171338 | -0.833501375 | 0.069418162 | -3.36369 | 0.000769 | 0.005856466 | ENSG00000067057 | PFKP | 5214 |
| ENSG00000141577.14 | 407.0495023 | -0.833895009 | 0.083489262 | -2.8015 | 0.005087 | 0.031735482 | ENSG00000141577 | CEP131 | 22994 |
| ENSG00000155506.17 | 24516.71711 | -0.840678162 | 0.080026788 | -3.00747 | 0.002634 | 0.017732214 | ENSG00000155506 | LARP1 | 23367 |
| ENSG00000187764.11 | 338.3626161 | -0.844037781 | 0.079189041 | -3.08171 | 0.002058 | 0.014169086 | ENSG00000187764 | SEMA4D | 10507 |
| ENSG00000169570.10 | 676.1719148 | -0.846067947 | 0.07366532 | -3.34035 | 0.000837 | 0.006328237 | ENSG00000169570 | DTWD2 | 285605 |
| ENSG00000112941.14 | 2744.751194 | -0.847131154 | 0.066730534 | -3.70342 | 0.000213 | 0.001801731 | ENSG00000112941 | TENT4A | 11044 |
| ENSG00000116141.15 | 614.5882359 | -0.847985496 | 0.089810129 | -2.76122 | 0.005759 | 0.035380858 | ENSG00000116141 | MARK1 | 4139 |
| ENSG00000163565.18 | 6529.208235 | -0.848073205 | 0.060216081 | -4.11972 | 3.79E-05 | 0.000367093 | ENSG00000163565 | IFI16 | 3428 |
| ENSG00000198466.12 | 671.8864 | -0.8502265 | 0.09378328 | -2.66814 | 0.007627 | 0.04529452 | ENSG00000198466 | ZNF587 | 84914 |
| ENSG00000275023.4 | 995.1523245 | -0.851786857 | 0.078016546 | -3.22735 | 0.001249 | 0.009053339 | ENSG00000275023 | MLLT6 | 4302 |
| ENSG00000095002.14 | 3212.631409 | -0.852832601 | 0.09468643 | -2.67021 | 0.00758 | 0.045055873 | ENSG00000095002 | MSH2 | 4436 |
| ENSG00000175643.10 | 370.3431635 | -0.853044766 | 0.094707514 | -2.67186 | 0.007543 | 0.044855533 | ENSG00000175643 | RMI2 | 116028 |
| ENSG00000147459.18 | 4537.699368 | -0.853400183 | 0.065341274 | -3.8781 | 0.000105 | 0.000947148 | ENSG00000147459 | DOCK5 | 80005 |
| ENSG00000105976.15 | 6167.895905 | -0.854325842 | 0.076959814 | -3.30466 | 0.000951 | 0.007085227 | ENSG00000105976 | MET | 4233 |
| ENSG00000110583.13 | 2096.174456 | -0.854348216 | 0.083609293 | -3.0421 | 0.002349 | 0.015974532 | ENSG00000110583 | NAA40 | 79829 |
| ENSG00000090621.14 | 9992.01782 | -0.855271371 | 0.090403231 | -2.8237 | 0.004747 | 0.029870737 | ENSG00000090621 | PABPC4 | 8761 |
| ENSG00000174197.16 | 2116.89128 | -0.857464536 | 0.085291941 | -3.01863 | 0.002539 | 0.01714416 | ENSG00000174197 | MGA | 23269 |
| ENSG00000082898.16 | 14117.41195 | -0.858026272 | 0.094612729 | -2.72718 | 0.006388 | 0.038754489 | ENSG00000082898 | XPO1 | 7514 |
| ENSG00000155561.15 | 7055.49583 | -0.858967569 | 0.092074355 | -2.81259 | 0.004914 | 0.030791051 | ENSG00000155561 | NUP205 | 23165 |
| ENSG00000137822.12 | 989.8108846 | -0.86038744 | 0.076908182 | -3.38569 | 0.00071 | 0.005451785 | ENSG00000137822 | TUBGCP4 | 27229 |
| ENSG00000137843.12 | 761.7937359 | -0.861205538 | 0.094600625 | -2.76114 | 0.00576 | 0.035381364 | ENSG00000137843 | PAK6 | 56924 |
| ENSG00000164032.12 | 13753.84665 | -0.863171134 | 0.099591191 | -2.64251 | 0.008229 | 0.048415574 | ENSG00000164032 | H2AZ1 | 3015 |
| ENSG00000139629.16 | 1438.314421 | -0.86406975 | 0.084591821 | -3.12169 | 0.001798 | 0.012561025 | ENSG00000139629 | GALNT6 | 11226 |
| ENSG00000104687.14 | 1573.948838 | -0.865731204 | 0.094822879 | -2.8024 | 0.005072 | 0.031669603 | ENSG00000104687 | GSR | 2936 |
| ENSG00000084444.14 | 367.2113632 | -0.866230498 | 0.080359512 | -3.31299 | 0.000923 | 0.006896712 | ENSG00000084444 | FAM234B | 57613 |

| | | | | | | | | | |
|---|---|---|---|---|---|---|---|---|---|
| ENSG00000123064.12 | 2991.93283 | -0.868145501 | 0.099590197 | -2.69249 | 0.007092 | 0.042523784 | ENSG00000123064 | DDX54 | 79039 |
| ENSG00000099219.14 | 1597.127498 | -0.869958359 | 0.06787591 | -3.97723 | 6.97E-05 | 0.000645161 | ENSG00000099219 | ERMP1 | 79956 |
| ENSG00000132932.17 | 359.0068802 | -0.870515244 | 0.097247963 | -2.78171 | 0.005407 | 0.033524871 | ENSG00000132932 | ATP8A2 | 51761 |
| ENSG00000103353.16 | 2581.595457 | -0.870609615 | 0.070641418 | -3.83075 | 0.000128 | 0.001131043 | ENSG00000103353 | UBFD1 | 56061 |
| ENSG00000105939.13 | 5048.980183 | -0.871299401 | 0.067077534 | -4.04456 | 5.24E-05 | 0.000493472 | ENSG00000105939 | ZC3HAV1 | 56829 |
| ENSG00000135045.7 | 557.4023807 | -0.874360979 | 0.082666371 | -3.31889 | 0.000904 | 0.006779234 | ENSG00000135045 | C9orf40 | 55071 |
| ENSG00000164167.12 | 1318.648894 | -0.874415992 | 0.103895887 | -2.64126 | 0.00826 | 0.048584527 | ENSG00000164167 | LSM6 | 11157 |
| ENSG00000138111.14 | 240.0064029 | -0.874876927 | 0.090371947 | -3.04162 | 0.002353 | 0.015992939 | ENSG00000138111 | MFSD13A | 79847 |
| ENSG00000104290.11 | 1304.264933 | -0.874994965 | 0.083897918 | -3.27773 | 0.001046 | 0.007729802 | ENSG00000104290 | FZD3 | 7976 |
| ENSG00000115368.10 | 3142.70358 | -0.87528668 | 0.101478494 | -2.71276 | 0.006673 | 0.040326227 | ENSG00000115368 | WDR75 | 84128 |
| ENSG00000196700.9 | 858.4031 | -0.8756351 | 0.09413392 | -2.92812 | 0.00341 | 0.02226112 | ENSG00000196700 | ZNF512B | 57473 |
| ENSG00000164916.11 | 2937.266754 | -0.877192254 | 0.090532829 | -3.06179 | 0.0022 | 0.015071793 | ENSG00000164916 | FOXK1 | 221937 |
| ENSG00000168003.16 | 22641.91744 | -0.87783094 | 0.098289945 | -2.82665 | 0.004704 | 0.029632067 | ENSG00000168003 | SLC3A2 | 6520 |
| ENSG00000060749.15 | 4057.437498 | -0.877993157 | 0.096696388 | -2.87491 | 0.004041 | 0.025925725 | ENSG00000060749 | QSER1 | 79832 |
| ENSG00000168393.13 | 1983.173511 | -0.878144052 | 0.104277873 | -2.66734 | 0.007646 | 0.045372142 | ENSG00000168393 | DTYMK | 1841 |
| ENSG00000174547.13 | 1790.011994 | -0.878378089 | 0.103287434 | -2.69518 | 0.007035 | 0.042220054 | ENSG00000174547 | MRPL11 | 65003 |
| ENSG00000100413.17 | 1482.895 | -0.8800275 | 0.1034669 | -2.70644 | 0.006801 | 0.04100812 | ENSG00000100413 | POLR3H | 171568 |
| ENSG00000136699.19 | 3477.19818 | -0.880794279 | 0.087661223 | -3.20318 | 0.001359 | 0.00975861 | ENSG00000136699 | SMPD4 | 55627 |
| ENSG00000185219.17 | 1196.494593 | -0.8808761 | 0.075145543 | -3.73776 | 0.000186 | 0.001591885 | ENSG00000185219 | ZNF445 | 353274 |
| ENSG00000107443.16 | 609.4866392 | -0.883370949 | 0.080006067 | -3.54187 | 0.000397 | 0.003209048 | ENSG00000107443 | CCNJ | 54619 |
| ENSG00000178695.5 | 3899.132516 | -0.885525747 | 0.075881696 | -3.76277 | 0.000168 | 0.001455213 | ENSG00000178695 | KCTD12 | 115207 |
| ENSG00000155096.14 | 8708.927927 | -0.886296175 | 0.049166524 | -5.82299 | 5.78E-09 | 9.16E-08 | ENSG00000155096 | AZIN1 | 51582 |
| ENSG00000163428.4 | 5449.023281 | -0.886937533 | 0.091290927 | -3.14311 | 0.001672 | 0.011751244 | ENSG00000163428 | LRRC58 | 116064 |
| ENSG00000130299.17 | 906.8415 | -0.8889849 | 0.1033203 | -2.79698 | 0.005158 | 0.03212244 | ENSG00000130299 | GTPBP3 | 84705 |
| ENSG00000162408.11 | 1343.697324 | -0.89075953 | 0.098141603 | -2.96265 | 0.00305 | 0.020183225 | ENSG00000162408 | NOL9 | 79707 |
| ENSG00000196367.13 | 4654.214976 | -0.892083029 | 0.084117675 | -3.47231 | 0.000516 | 0.004073761 | ENSG00000196367 | TRRAP | 8295 |
| ENSG00000213066.13 | 762.3606351 | -0.892288155 | 0.075086681 | -3.89268 | 9.91E-05 | 0.000895934 | ENSG00000213066 | CEP43 | 11116 |
| ENSG00000072736.19 | 644.4020216 | -0.89265727 | 0.093993963 | -3.11358 | 0.001848 | 0.012864545 | ENSG00000072736 | NFATC3 | 4775 |
| ENSG00000213965.3 | 1085.648 | -0.8938682 | 0.1110815 | -2.64552 | 0.008157 | 0.04804085 | ENSG00000213965 | NUDT19 | 390916 |
| ENSG00000213024.11 | 1870.883 | -0.8948205 | 0.08712907 | -3.38372 | 0.000715 | 0.005486311 | ENSG00000213024 | NUP62 | 23636 |
| ENSG00000139514.13 | 7830.756935 | -0.895145668 | 0.093639537 | -3.15193 | 0.001622 | 0.011426028 | ENSG00000139514 | SLC7A1 | 6541 |
| ENSG00000129028.9 | 210.9860833 | -0.895426914 | 0.108299262 | -2.72788 | 0.006374 | 0.03868209 | ENSG00000129028 | THAP10 | 56906 |
| ENSG00000179051.14 | 2437.078174 | -0.895513076 | 0.06596308 | -4.47998 | 7.47E-06 | 8.07E-05 | ENSG00000179051 | RCC2 | 55920 |
| ENSG00000197461.13 | 1543.340383 | -0.895581569 | 0.059199885 | -4.99294 | 5.95E-07 | 7.40E-06 | ENSG00000197461 | PDGFA | 5154 |
| ENSG00000164048.14 | 601.9690753 | -0.896916923 | 0.083834447 | -3.54171 | 0.000398 | 0.003210055 | ENSG00000164048 | ZNF589 | 51385 |
| ENSG00000125484.12 | 3406.101759 | -0.899723326 | 0.107380764 | -2.79122 | 0.005251 | 0.032648882 | ENSG00000125484 | GTF3C4 | 9329 |
| ENSG00000163781.13 | 4107.438204 | -0.899916195 | 0.105108962 | -2.85338 | 0.004326 | 0.027569329 | ENSG00000163781 | TOPBP1 | 11073 |
| ENSG00000088247.17 | 7481.474697 | -0.900167851 | 0.107284105 | -2.79788 | 0.005144 | 0.032040779 | ENSG00000088247 | KHSRP | 8570 |
| ENSG00000196924.17 | 49729.68 | -0.9012878 | 0.1011563 | -2.97844 | 0.002897 | 0.01927462 | ENSG00000196924 | FLNA | 2316 |
| ENSG00000145375.8 | 812.732786 | -0.901410541 | 0.075815534 | -3.97558 | 7.02E-05 | 0.000649217 | ENSG00000145375 | SPATA5 | 166378 |
| ENSG00000183814.15 | 782.8147637 | -0.903022127 | 0.085123538 | -3.55979 | 0.000371 | 0.003012409 | ENSG00000183814 | LIN9 | 286826 |
| ENSG00000103653.16 | 2780.762509 | -0.904886808 | 0.082968847 | -3.67471 | 0.000238 | 0.002004147 | ENSG00000103653 | CSK | 1445 |

| | | | | | | | | | |
|---|---|---|---|---|---|---|---|---|---|
| ENSG00000175548.9 | 564.4638007 | -0.906820725 | 0.099071297 | -3.09697 | 0.001955 | 0.013526189 | ENSG00000175548 | ALG10B | 144245 |
| ENSG00000111581.10 | 3483.314769 | -0.907082361 | 0.09398861 | -3.26723 | 0.001086 | 0.007995846 | ENSG00000111581 | NUP107 | 57122 |
| ENSG00000130702.15 | 6780.887 | -0.9071127 | 0.07904513 | -3.88528 | 0.000102 | 0.000921765 | ENSG00000130702 | LAMA5 | 3911 |
| ENSG00000075213.11 | 1889.686507 | -0.90720838 | 0.102781179 | -2.98896 | 0.002799 | 0.018701057 | ENSG00000075213 | SEMA3A | 10371 |
| ENSG00000172009.15 | 3057.946799 | -0.908172414 | 0.106076872 | -2.90518 | 0.00367 | 0.023751552 | ENSG00000172009 | THOP1 | 7064 |
| ENSG00000164818.16 | 1807.468277 | -0.908386796 | 0.113680003 | -2.71276 | 0.006673 | 0.040326227 | ENSG00000164818 | DNAAF5 | 54919 |
| ENSG00000137574.10 | 2599.748832 | -0.909168539 | 0.072211267 | -4.28144 | 1.86E-05 | 0.000187898 | ENSG00000137574 | TGS1 | 96764 |
| ENSG00000042088.13 | 987.6417266 | -0.909395919 | 0.095098338 | -3.25343 | 0.00114 | 0.008346019 | ENSG00000042088 | TDP1 | 55775 |
| ENSG00000239264.8 | 232.9955992 | -0.909784197 | 0.101710244 | -3.04575 | 0.002321 | 0.015802919 | ENSG00000239264 | TXNDC5 | 81567 |
| ENSG00000143537.13 | 4185.061854 | -0.909901013 | 0.083597211 | -3.70707 | 0.00021 | 0.001778771 | ENSG00000143537 | ADAM15 | 8751 |
| ENSG00000112578.10 | 1378.41101 | -0.910196538 | 0.10991318 | -2.8222 | 0.00477 | 0.029982605 | ENSG00000112578 | BYSL | 705 |
| ENSG00000103148.15 | 706.8239104 | -0.91047376 | 0.108346213 | -2.86557 | 0.004163 | 0.026619315 | ENSG00000103148 | NPRL3 | 8131 |
| ENSG00000134987.11 | 6475.467167 | -0.910922595 | 0.115546718 | -2.69088 | 0.007126 | 0.042690675 | ENSG00000134987 | WDR36 | 134430 |
| ENSG00000139083.10 | 980.5224177 | -0.912465565 | 0.101607138 | -3.07523 | 0.002103 | 0.014454398 | ENSG00000139083 | ETV6 | 2120 |
| ENSG00000143641.10 | 2193.05912 | -0.913634132 | 0.111935192 | -2.80193 | 0.00508 | 0.031708278 | ENSG00000143641 | GALNT2 | 2590 |
| ENSG00000163535.18 | 1477.766946 | -0.914883764 | 0.115616891 | -2.72351 | 0.006459 | 0.039134614 | ENSG00000163535 | SGO2 | 151246 |
| ENSG00000140854.13 | 1167.161944 | -0.915809843 | 0.093552461 | -3.37575 | 0.000736 | 0.005633122 | ENSG00000140854 | KATNB1 | 10300 |
| ENSG00000198000.12 | 2474.4153 | -0.916640183 | 0.096216554 | -3.29091 | 0.000999 | 0.007401335 | ENSG00000198000 | NOL8 | 55035 |
| ENSG00000002549.12 | 3358.265138 | -0.917085181 | 0.042860457 | -7.39808 | 1.38E-13 | 3.41E-12 | ENSG00000002549 | LAP3 | 51056 |
| ENSG00000141905.19 | 1732.625336 | -0.917140363 | 0.116105804 | -2.73148 | 0.006305 | 0.038322972 | ENSG00000141905 | NFIC | 4782 |
| ENSG00000181751.10 | 1061.424944 | -0.917346153 | 0.06838296 | -4.64072 | 3.47E-06 | 3.90E-05 | ENSG00000181751 | MACIR | 90355 |
| ENSG00000015133.19 | 1316.592904 | -0.917526276 | 0.085563511 | -3.711 | 0.000206 | 0.001754751 | ENSG00000015133 | CCDC88C | 440193 |
| ENSG00000160352.15 | 609.751 | -0.9193591 | 0.07422196 | -4.30276 | 1.69E-05 | 0.000171665 | ENSG00000160352 | ZNF714 | 148206 |
| ENSG00000160072.19 | 2764.529436 | -0.919825926 | 0.116906521 | -2.73574 | 0.006224 | 0.037899237 | ENSG00000160072 | ATAD3B | 83858 |
| ENSG00000117395.12 | 4724.868595 | -0.920190503 | 0.110533813 | -2.89677 | 0.00377 | 0.024321157 | ENSG00000117395 | EBNA1BP2 | 10969 |
| ENSG00000164430.16 | 1565.927547 | -0.921590065 | 0.041369787 | -7.77355 | 7.63E-15 | 2.06E-13 | ENSG00000164430 | CGAS | 115004 |
| ENSG00000117632.23 | 7393.085722 | -0.921903833 | 0.06188468 | -5.20167 | 1.98E-07 | 2.62E-06 | ENSG00000117632 | STMN1 | 3925 |
| ENSG00000115942.9 | 2126.016088 | -0.922275319 | 0.109056486 | -2.95512 | 0.003125 | 0.020600545 | ENSG00000115942 | ORC2 | 4999 |
| ENSG00000183207.14 | 3797.557 | -0.9225624 | 0.1037142 | -3.11011 | 0.00187 | 0.01299611 | ENSG00000183207 | RUVBL2 | 10856 |
| ENSG00000168061.15 | 395.065545 | -0.922996465 | 0.104074263 | -3.10352 | 0.001912 | 0.013250984 | ENSG00000168061 | SAC3D1 | 29901 |
| ENSG00000129474.16 | 6035.062039 | -0.924015421 | 0.067054609 | -4.83211 | 1.35E-06 | 1.60E-05 | ENSG00000129474 | AJUBA | 84962 |
| ENSG00000116750.13 | 3625.717094 | -0.924730864 | 0.100485186 | -3.23163 | 0.001231 | 0.008933514 | ENSG00000116750 | UCHL5 | 51377 |
| ENSG00000126464.14 | 1320.92 | -0.9249361 | 0.0919833 | -3.53256 | 0.000412 | 0.003311173 | ENSG00000126464 | PRR12 | 57479 |
| ENSG00000162377.6 | 1838.145912 | -0.926857965 | 0.122463832 | -2.66902 | 0.007607 | 0.045185978 | ENSG00000162377 | COA7 | 65260 |
| ENSG00000161800.13 | 4002.145719 | -0.927425613 | 0.107762125 | -3.03841 | 0.002378 | 0.016153914 | ENSG00000161800 | RACGAP1 | 29127 |
| ENSG00000077514.8 | 2010.22256 | -0.928081105 | 0.122892156 | -2.66967 | 0.007593 | 0.045118679 | ENSG00000077514 | POLD3 | 10714 |
| ENSG00000152778.9 | 1712.074591 | -0.928500495 | 0.074852731 | -4.38862 | 1.14E-05 | 0.000119558 | ENSG00000152778 | IFIT5 | 24138 |
| ENSG00000138182.14 | 4722.009662 | -0.928875176 | 0.105947919 | -3.10412 | 0.001908 | 0.013227492 | ENSG00000138182 | KIF20B | 9585 |
| ENSG00000117228.10 | 1148.122559 | -0.930055508 | 0.085674598 | -3.85243 | 0.000117 | 0.001043736 | ENSG00000117228 | GBP1 | 2633 |
| ENSG00000165185.14 | 275.1222045 | -0.930423244 | 0.092340739 | -3.5783 | 0.000346 | 0.002824145 | ENSG00000165185 | KIAA1958 | 158405 |
| ENSG00000005156.11 | 839.4081045 | -0.930762184 | 0.087513354 | -3.77956 | 0.000157 | 0.001368148 | ENSG00000005156 | LIG3 | 3980 |
| ENSG00000162430.17 | 1779.572288 | -0.931821853 | 0.087026844 | -3.81287 | 0.000137 | 0.001208092 | ENSG00000162430 | SELENON | 57190 |

| | | | | | | | | | |
|---|---|---|---|---|---|---|---|---|---|
| ENSG00000100228.12 | 347.7201 | -0.9318272 | 0.1122328 | -2.9566 | 0.003111 | 0.02053805 | ENSG00000100228 | RAB36 | 9609 |
| ENSG00000005810.18 | 4460.412146 | -0.932171175 | 0.117706564 | -2.82203 | 0.004772 | 0.029991271 | ENSG00000005810 | MYCBP2 | 23077 |
| ENSG00000184162.14 | 370.1351 | -0.9332129 | 0.102293 | -3.25744 | 0.001124 | 0.008247265 | ENSG00000184162 | NR2C2AP | 126382 |
| ENSG00000269343.8 | 653.4653 | -0.9333021 | 0.1265258 | -2.63426 | 0.008432 | 0.04940087 | ENSG00000269343 | ZNF587B | 1E+08 |
| ENSG00000124608.5 | 915.1585143 | -0.933528826 | 0.100234172 | -3.3275 | 0.000876 | 0.006588226 | ENSG00000124608 | AARS2 | 57505 |
| ENSG00000155893.13 | 246.9419176 | -0.933615967 | 0.106544564 | -3.13123 | 0.001741 | 0.012208064 | ENSG00000155893 | PXYLP1 | 92370 |
| ENSG00000130119.16 | 2165.013 | -0.9352605 | 0.1136668 | -2.9495 | 0.003183 | 0.02095307 | ENSG00000130119 | GNL3L | 54552 |
| ENSG00000178202.13 | 813.6043143 | -0.936810523 | 0.086070473 | -3.91319 | 9.11E-05 | 0.000827863 | ENSG00000178202 | POGLUT3 | 143888 |
| ENSG00000182179.13 | 775.4676975 | -0.937310305 | 0.08742881 | -3.85811 | 0.000114 | 0.001022512 | ENSG00000182179 | UBA7 | 7318 |
| ENSG00000120833.14 | 1435.300087 | -0.937353494 | 0.062360534 | -5.40973 | 6.31E-08 | 8.88E-07 | ENSG00000120833 | SOCS2 | 8835 |
| ENSG00000151806.14 | 2150.200933 | -0.937605368 | 0.109529117 | -3.08233 | 0.002054 | 0.014143124 | ENSG00000151806 | GUF1 | 60558 |
| ENSG00000107938.18 | 859.0319455 | -0.937693725 | 0.072443247 | -4.66149 | 3.14E-06 | 3.54E-05 | ENSG00000107938 | EDRF1 | 26098 |
| ENSG00000183765.22 | 554.9557 | -0.9398506 | 0.1140308 | -2.98034 | 0.002879 | 0.01918218 | ENSG00000183765 | CHEK2 | 11200 |
| ENSG00000086712.13 | 2721.748 | -0.939997 | 0.1014509 | -3.35134 | 0.000804 | 0.006101308 | ENSG00000086712 | TXLNG | 55787 |
| ENSG00000128829.12 | 4313.42099 | -0.940303895 | 0.094021744 | -3.61942 | 0.000295 | 0.002441184 | ENSG00000128829 | EIF2AK4 | 440275 |
| ENSG00000134490.14 | 290.519263 | -0.940616092 | 0.129535123 | -2.62953 | 0.00855 | 0.049984216 | ENSG00000134490 | TMEM241 | 85019 |
| ENSG00000069399.15 | 608.129 | -0.9406991 | 0.105321 | -3.23486 | 0.001217 | 0.008852249 | ENSG00000069399 | BCL3 | 602 |
| ENSG00000124279.12 | 606.8693414 | -0.941303854 | 0.120984814 | -2.82105 | 0.004787 | 0.030075999 | ENSG00000124279 | FASTKD3 | 79072 |
| ENSG00000104731.14 | 1059.74754 | -0.942323702 | 0.102769169 | -3.331 | 0.000865 | 0.006511466 | ENSG00000104731 | KLHDC4 | 54758 |
| ENSG00000129351.17 | 16923.98537 | -0.942631602 | 0.099672316 | -3.43758 | 0.000587 | 0.004582235 | ENSG00000129351 | ILF3 | 3609 |
| ENSG00000166938.13 | 1325.934512 | -0.943176073 | 0.079306594 | -4.32721 | 1.51E-05 | 0.000155054 | ENSG00000166938 | DIS3L | 115752 |
| ENSG00000075131.10 | 589.0379273 | -0.944153186 | 0.1178805 | -2.91951 | 0.003506 | 0.022814047 | ENSG00000075131 | TIPIN | 54962 |
| ENSG00000136108.15 | 4486.753179 | -0.944713191 | 0.089768788 | -3.84001 | 0.000123 | 0.001091755 | ENSG00000136108 | CKAP2 | 26586 |
| ENSG00000181026.15 | 1467.902791 | -0.945723636 | 0.108444036 | -3.18804 | 0.001432 | 0.010240037 | ENSG00000181026 | AEN | 64782 |
| ENSG00000113356.12 | 2513.70182 | -0.945743627 | 0.103726442 | -3.33323 | 0.000858 | 0.00646499 | ENSG00000113356 | POLR3G | 10622 |
| ENSG00000185730.7 | 443.2716659 | -0.946552695 | 0.114774135 | -3.01943 | 0.002532 | 0.017111671 | ENSG00000185730 | ZNF696 | 79943 |
| ENSG00000236088.10 | 397.7745194 | -0.94672014 | 0.13014784 | -2.66405 | 0.007721 | 0.045787267 | ENSG00000236088 | COX10-AS | 1.01E+08 |
| ENSG00000185379.20 | 416.9714265 | -0.94850557 | 0.106822223 | -3.26248 | 0.001104 | 0.008113041 | ENSG00000185379 | RAD51D | 5892 |
| ENSG00000174231.17 | 12907.09949 | -0.948834745 | 0.109825888 | -3.17625 | 0.001492 | 0.010605512 | ENSG00000174231 | PRPF8 | 10594 |
| ENSG00000168806.7 | 402.917012 | -0.949674395 | 0.093103476 | -3.75576 | 0.000173 | 0.001492764 | ENSG00000168806 | LCMT2 | 9836 |
| ENSG00000053254.15 | 612.5430367 | -0.950440317 | 0.078427827 | -4.46832 | 7.88E-06 | 8.50E-05 | ENSG00000053254 | FOXN3 | 1112 |
| ENSG00000169252.5 | 2697.477408 | -0.950587801 | 0.08666817 | -4.04517 | 5.23E-05 | 0.000492434 | ENSG00000169252 | ADRB2 | 154 |
| ENSG00000123104.12 | 1702.421692 | -0.950964012 | 0.133326474 | -2.63237 | 0.008479 | 0.049633865 | ENSG00000123104 | ITPR2 | 3709 |
| ENSG00000135966.13 | 2326.412916 | -0.951877148 | 0.103015553 | -3.41577 | 0.000636 | 0.004925072 | ENSG00000135966 | TGFBRAP | 9392 |
| ENSG00000277534.1 | 278.877614 | -0.952536856 | 0.115541846 | -3.05116 | 0.00228 | 0.015552771 | ENSG00000277534 | NA | NA |
| ENSG00000253293.5 | 309.7769566 | -0.953142925 | 0.10642948 | -3.31809 | 0.000906 | 0.006792977 | ENSG00000253293 | HOXA10 | 3206 |
| ENSG00000156136.10 | 1556.792142 | -0.953576504 | 0.085016035 | -4.15894 | 3.20E-05 | 0.000312246 | ENSG00000156136 | DCK | 1633 |
| ENSG00000175455.15 | 2950.197448 | -0.953702588 | 0.079695496 | -4.43818 | 9.07E-06 | 9.69E-05 | ENSG00000175455 | CCDC14 | 64770 |
| ENSG00000237289.9 | 294.7699213 | -0.953807346 | 0.107654626 | -3.2865 | 0.001014 | 0.007513966 | ENSG00000237289 | CKMT1B | 1159 |
| ENSG00000126215.14 | 474.5909046 | -0.954219979 | 0.123171513 | -2.87583 | 0.00403 | 0.025856534 | ENSG00000126215 | XRCC3 | 7517 |
| ENSG00000101911.12 | 2185.114 | -0.9559962 | 0.1047083 | -3.39989 | 0.000674 | 0.00520053 | ENSG00000101911 | PRPS2 | 5634 |
| ENSG00000100911.16 | 3875.91653 | -0.956197839 | 0.054750333 | -6.50586 | 7.73E-11 | 1.48E-09 | ENSG00000100911 | PSME2 | 5721 |

| | | | | | | | | | |
|---|---|---|---|---|---|---|---|---|---|
| ENSG00000130726.12 | 10977.41 | -0.9567878 | 0.07611672 | -4.68738 | 2.77E-06 | 3.15E-05 | ENSG00000130726 | TRIM28 | 10155 |
| ENSG00000147874.11 | 2724.615344 | -0.957326193 | 0.108647324 | -3.28886 | 0.001006 | 0.007453336 | ENSG00000147874 | HAUS6 | 54801 |
| ENSG00000097096.9 | 249.623458 | -0.958243626 | 0.081351078 | -4.40367 | 1.06E-05 | 0.000112039 | ENSG00000097096 | SYDE2 | 84144 |
| ENSG00000116830.12 | 1428.951073 | -0.958288156 | 0.086208993 | -4.15604 | 3.24E-05 | 0.000315756 | ENSG00000116830 | TTF2 | 8458 |
| ENSG00000171067.11 | 1021.011383 | -0.959028067 | 0.099122685 | -3.62206 | 0.000292 | 0.002420152 | ENSG00000171067 | C11orf24 | 53838 |
| ENSG00000145332.14 | 1077.469034 | -0.959514811 | 0.095796219 | -3.75291 | 0.000175 | 0.001506414 | ENSG00000145332 | KLHL8 | 57563 |
| ENSG00000113360.16 | 2575.313312 | -0.959968286 | 0.081491761 | -4.41724 | 1.00E-05 | 0.000105907 | ENSG00000113360 | DROSHA | 29102 |
| ENSG00000166788.10 | 1390.917122 | -0.960201712 | 0.130702482 | -2.75589 | 0.005853 | 0.03592105 | ENSG00000166788 | SAAL1 | 113174 |
| ENSG00000187792.5 | 162.3947 | -0.9602773 | 0.1080325 | -3.3349 | 0.000853 | 0.00642989 | ENSG00000187792 | ZNF70 | 7621 |
| ENSG00000146243.14 | 274.5620192 | -0.960562042 | 0.123745149 | -2.91375 | 0.003571 | 0.023177062 | ENSG00000146243 | IRAK1BP1 | 134728 |
| ENSG00000198720.13 | 361.5855978 | -0.961116062 | 0.13584758 | -2.65824 | 0.007855 | 0.046500541 | ENSG00000198720 | ANKRD13I | 124930 |
| ENSG00000005100.13 | 2883.628232 | -0.961393035 | 0.122154524 | -2.95849 | 0.003091 | 0.020422276 | ENSG00000005100 | DHX33 | 56919 |
| ENSG00000174177.13 | 369.0250731 | -0.962539119 | 0.120796967 | -3.00123 | 0.002689 | 0.01805877 | ENSG00000174177 | CTU2 | 348180 |
| ENSG00000077684.16 | 666.4926403 | -0.96263537 | 0.10504175 | -3.4523 | 0.000556 | 0.004361163 | ENSG00000077684 | JADE1 | 79960 |
| ENSG00000124496.12 | 1308.68754 | -0.962865756 | 0.133970495 | -2.70855 | 0.006758 | 0.040776535 | ENSG00000124496 | TRERF1 | 55809 |
| ENSG00000172954.13 | 1011.566325 | -0.963754822 | 0.136972935 | -2.65567 | 0.007915 | 0.046794644 | ENSG00000172954 | LCLAT1 | 253558 |
| ENSG00000138796.16 | 1555.76246 | -0.963847726 | 0.125226825 | -2.90551 | 0.003667 | 0.023732335 | ENSG00000138796 | HADH | 3033 |
| ENSG00000253930.1 | 173.2171363 | -0.963908876 | 0.127152242 | -2.86199 | 0.00421 | 0.026882899 | ENSG00000253930 | NA | NA |
| ENSG00000196365.11 | 3473.883324 | -0.96543204 | 0.082393683 | -4.43519 | 9.20E-06 | 9.82E-05 | ENSG00000196365 | LONP1 | 9361 |
| ENSG00000009413.15 | 1998.123419 | -0.967409485 | 0.093139011 | -3.94474 | 7.99E-05 | 0.000733096 | ENSG00000009413 | REV3L | 5980 |
| ENSG00000100304.13 | 3895.844 | -0.9678765 | 0.1374652 | -2.67614 | 0.007448 | 0.04438496 | ENSG00000100304 | TTLL12 | 23170 |
| ENSG00000185278.15 | 829.0679752 | -0.968373551 | 0.080574762 | -4.57182 | 4.83E-06 | 5.34E-05 | ENSG00000185278 | ZBTB37 | 84614 |
| ENSG00000159055.4 | 1117.46 | -0.9697928 | 0.1205603 | -3.06729 | 0.00216 | 0.01482136 | ENSG00000159055 | MIS18A | 54069 |
| ENSG00000245571.7 | 128.9623971 | -0.970459978 | 0.137480789 | -2.69463 | 0.007047 | 0.042279989 | ENSG00000245571 | FAM111A- | 1.02E+08 |
| ENSG00000112365.4 | 829.372433 | -0.971465016 | 0.104738517 | -3.54659 | 0.00039 | 0.003155865 | ENSG00000112365 | ZBTB24 | 9841 |
| ENSG00000118420.17 | 231.4062231 | -0.971947213 | 0.1326712 | -2.80353 | 0.005055 | 0.031580961 | ENSG00000118420 | UBE3D | 90025 |
| ENSG00000127423.10 | 558.5283042 | -0.973066674 | 0.098954779 | -3.77007 | 0.000163 | 0.001417064 | ENSG00000127423 | AUNIP | 79000 |
| ENSG00000183723.12 | 3904.963171 | -0.97330341 | 0.038507219 | -9.69437 | 3.19E-22 | 1.38E-20 | ENSG00000183723 | CMTM4 | 146223 |
| ENSG00000156853.12 | 438.3662822 | -0.973442566 | 0.1248957 | -2.99004 | 0.002789 | 0.018644431 | ENSG00000156853 | ZNF689 | 115509 |
| ENSG00000144283.21 | 3312.606394 | -0.973577827 | 0.109851992 | -3.40074 | 0.000672 | 0.005187333 | ENSG00000144283 | PKP4 | 8502 |
| ENSG00000133739.16 | 496.7192988 | -0.973992697 | 0.111179896 | -3.36385 | 0.000769 | 0.005854768 | ENSG00000133739 | LRRCC1 | 85444 |
| ENSG00000172878.14 | 356.9558857 | -0.974137282 | 0.107258371 | -3.48819 | 0.000486 | 0.00385886 | ENSG00000172878 | METAP1D | 254042 |
| ENSG00000146648.18 | 14156.83401 | -0.974400752 | 0.074110257 | -5.05194 | 4.37E-07 | 5.53E-06 | ENSG00000146648 | EGFR | 1956 |
| ENSG00000183808.11 | 1357.256325 | -0.975003912 | 0.084140568 | -4.45687 | 8.32E-06 | 8.94E-05 | ENSG00000183808 | RBM12B | 389677 |
| ENSG00000265415.1 | 439.1669177 | -0.975195531 | 0.120698039 | -3.10855 | 0.00188 | 0.01305209 | ENSG00000265415 | NA | NA |
| ENSG00000163389.12 | 873.8910128 | -0.976723267 | 0.134838766 | -2.79388 | 0.005208 | 0.032401741 | ENSG00000163389 | POGLUT1 | 56983 |
| ENSG00000108064.11 | 4600.646173 | -0.976891613 | 0.118229269 | -3.1878 | 0.001434 | 0.010245589 | ENSG00000108064 | TFAM | 7019 |
| ENSG00000008311.15 | 798.404325 | -0.979916299 | 0.124520182 | -3.05104 | 0.00228 | 0.015554991 | ENSG00000008311 | AASS | 10157 |
| ENSG00000067177.15 | 952.2458 | -0.9809364 | 0.05605822 | -6.79537 | 1.08E-11 | 2.26E-10 | ENSG00000067177 | PHKA1 | 5255 |
| ENSG00000116771.6 | 190.893537 | -0.982221608 | 0.130408769 | -2.93095 | 0.003379 | 0.022087628 | ENSG00000116771 | AGMAT | 79814 |
| ENSG00000088205.13 | 6177.538701 | -0.983121198 | 0.10117479 | -3.78673 | 0.000153 | 0.001332368 | ENSG00000088205 | DDX18 | 8886 |
| ENSG00000103995.14 | 805.8351529 | -0.983180284 | 0.107750773 | -3.55617 | 0.000376 | 0.003050487 | ENSG00000103995 | CEP152 | 22995 |

| | | | | | | | | | |
|---|---|---|---|---|---|---|---|---|---|
| ENSG00000087903.13 | 227.7095564 | -0.983557154 | 0.119243886 | -3.21658 | 0.001297 | 0.009374764 | ENSG00000087903 | RFX2 | 5990 |
| ENSG00000204392.11 | 728.2943105 | -0.984511773 | 0.08201173 | -4.6885 | 2.75E-06 | 3.14E-05 | ENSG00000204392 | LSM2 | 57819 |
| ENSG00000129116.19 | 4184.972684 | -0.98544515 | 0.140134337 | -2.75054 | 0.00595 | 0.036425139 | ENSG00000129116 | PALLD | 23022 |
| ENSG00000186230.7 | 473.3209 | -0.9858018 | 0.1374989 | -2.80585 | 0.005018 | 0.03137591 | ENSG00000186230 | ZNF749 | 388567 |
| ENSG00000253304.2 | 330.6432572 | -0.986840584 | 0.097822389 | -3.95452 | 7.67E-05 | 0.000706436 | ENSG00000253304 | TMEM200I | 399474 |
| ENSG00000110092.3 | 7573.798875 | -0.987762392 | 0.102650403 | -3.7775 | 0.000158 | 0.001377695 | ENSG00000110092 | CCND1 | 595 |
| ENSG00000196363.9 | 3270.872659 | -0.988901882 | 0.121331693 | -3.20528 | 0.001349 | 0.009698041 | ENSG00000196363 | WDR5 | 11091 |
| ENSG00000204371.11 | 718.4435784 | -0.989703904 | 0.084623588 | -4.60515 | 4.12E-06 | 4.59E-05 | ENSG00000204371 | EHMT2 | 10919 |
| ENSG00000163006.12 | 580.6145988 | -0.990976371 | 0.114070089 | -3.42751 | 0.000609 | 0.004737548 | ENSG00000163006 | CCDC138 | 165055 |
| ENSG00000132953.16 | 3405.759554 | -0.991247215 | 0.121470916 | -3.22091 | 0.001278 | 0.009245512 | ENSG00000132953 | XPO4 | 64328 |
| ENSG00000021826.15 | 745.5636833 | -0.991783672 | 0.091448157 | -4.28422 | 1.83E-05 | 0.000185842 | ENSG00000021826 | CPS1 | 1373 |
| ENSG00000152818.18 | 1721.551817 | -0.992831593 | 0.133864613 | -2.93454 | 0.00334 | 0.021860289 | ENSG00000152818 | UTRN | 7402 |
| ENSG00000278259.4 | 1961.040015 | -0.992903371 | 0.075529762 | -5.20197 | 1.97E-07 | 2.62E-06 | ENSG00000278259 | MYO19 | 80179 |
| ENSG00000116062.15 | 3965.469021 | -0.993279772 | 0.118002152 | -3.33282 | 0.00086 | 0.006472621 | ENSG00000116062 | MSH6 | 2956 |
| ENSG00000068489.12 | 3251.077018 | -0.993445416 | 0.114038187 | -3.45012 | 0.00056 | 0.004388735 | ENSG00000068489 | PRR11 | 55771 |
| ENSG00000156876.10 | 1007.156591 | -0.994029697 | 0.067298789 | -5.85493 | 4.77E-09 | 7.62E-08 | ENSG00000156876 | SASS6 | 163786 |
| ENSG00000141456.15 | 2203.345147 | -0.996356513 | 0.099458945 | -3.98513 | 6.74E-05 | 0.000625827 | ENSG00000141456 | PELP1 | 27043 |
| ENSG00000196935.9 | 1035.792186 | -0.996988195 | 0.131547255 | -3.01784 | 0.002546 | 0.017180176 | ENSG00000196935 | SRGAP1 | 57522 |
| ENSG00000153574.9 | 819.3026502 | -0.997950493 | 0.131416633 | -3.02816 | 0.00246 | 0.016663191 | ENSG00000153574 | RPIA | 22934 |
| ENSG00000064666.15 | 5798.090696 | -0.998354042 | 0.062217976 | -6.40256 | 1.53E-10 | 2.87E-09 | ENSG00000064666 | CNN2 | 1265 |
| ENSG00000115827.14 | 878.009313 | -0.998538097 | 0.139234089 | -2.86236 | 0.004205 | 0.026864702 | ENSG00000115827 | DCAF17 | 80067 |
| ENSG00000136450.13 | 4565.729726 | -0.998975411 | 0.107307091 | -3.71807 | 0.000201 | 0.00171129 | ENSG00000136450 | SRSF1 | 6426 |
| ENSG00000184990.13 | 1625.401131 | -1.000046741 | 0.122500871 | -3.26566 | 0.001092 | 0.008033501 | ENSG00000184990 | SIVA1 | 10572 |
| ENSG00000186575.17 | 2575.539 | -1.000273 | 0.09458055 | -4.23209 | 2.32E-05 | 0.000230508 | ENSG00000186575 | NF2 | 4771 |
| ENSG00000049449.9 | 1158.984499 | -1.000379143 | 0.109618246 | -3.65249 | 0.00026 | 0.002167427 | ENSG00000049449 | RCN1 | 5954 |
| ENSG00000123219.13 | 2935.849761 | -1.001246262 | 0.145889133 | -2.75035 | 0.005953 | 0.036425139 | ENSG00000123219 | CENPK | 64105 |
| ENSG00000102934.10 | 209.2258206 | -1.001425749 | 0.148470134 | -2.70375 | 0.006856 | 0.041286304 | ENSG00000102934 | PLLP | 51090 |
| ENSG00000145632.15 | 9109.232441 | -1.001584038 | 0.112081131 | -3.58298 | 0.00034 | 0.002778351 | ENSG00000145632 | PLK2 | 10769 |
| ENSG00000170515.14 | 8521.106001 | -1.001871867 | 0.134469763 | -2.98857 | 0.002803 | 0.018720171 | ENSG00000170515 | PA2G4 | 5036 |
| ENSG00000282936.2 | 206.4379843 | -1.002935795 | 0.122937039 | -3.27758 | 0.001047 | 0.007729802 | ENSG00000282936 | NA | NA |
| ENSG00000177706.9 | 1069.804175 | -1.003097171 | 0.119465749 | -3.37417 | 0.00074 | 0.005660801 | ENSG00000177706 | FAM20C | 56975 |
| ENSG00000132361.17 | 6083.142345 | -1.003391863 | 0.137105526 | -2.9422 | 0.003259 | 0.021374304 | ENSG00000132361 | CLUH | 23277 |
| ENSG00000198176.13 | 8364.130829 | -1.007139221 | 0.115376604 | -3.52878 | 0.000417 | 0.003353715 | ENSG00000198176 | TFDP1 | 7027 |
| ENSG00000119403.15 | 2550.879487 | -1.007899176 | 0.097224538 | -4.19543 | 2.72E-05 | 0.000268636 | ENSG00000119403 | PHF19 | 26147 |
| ENSG00000151458.12 | 4230.084028 | -1.009096703 | 0.120045644 | -3.40784 | 0.000655 | 0.005058533 | ENSG00000151458 | ANKRD50 | 57182 |
| ENSG00000197961.11 | 2938.11682 | -1.009585013 | 0.100988529 | -4.05576 | 5.00E-05 | 0.000472432 | ENSG00000197961 | ZNF121 | 7675 |
| ENSG00000101444.13 | 7487.364 | -1.01007 | 0.1139932 | -3.59732 | 0.000322 | 0.002637669 | ENSG00000101444 | AHCY | 191 |
| ENSG00000182158.15 | 879.5652738 | -1.010073674 | 0.07666135 | -5.34916 | 8.84E-08 | 1.23E-06 | ENSG00000182158 | CREB3L2 | 64764 |
| ENSG00000163251.4 | 451.5682942 | -1.011095276 | 0.104356022 | -3.93935 | 8.17E-05 | 0.000748471 | ENSG00000163251 | FZD5 | 7855 |
| ENSG00000180035.12 | 383.5081996 | -1.012269222 | 0.075511433 | -5.45969 | 4.77E-08 | 6.78E-07 | ENSG00000180035 | ZNF48 | 197407 |
| ENSG00000138095.19 | 11765.28161 | -1.012498789 | 0.124417277 | -3.31545 | 0.000915 | 0.006846597 | ENSG00000138095 | LRPPRC | 10128 |
| ENSG00000100065.15 | 4546.266 | -1.012567 | 0.0900569 | -4.58119 | 4.62E-06 | 5.12E-05 | ENSG00000100065 | CARD10 | 29775 |

| | | | | | | | | | |
|---|---|---|---|---|---|---|---|---|---|
| ENSG00000137513.10 | 836.2301544 | -1.014440604 | 0.122806799 | -3.37474 | 0.000739 | 0.005650683 | ENSG00000137513 | NARS2 | 79731 |
| ENSG00000173432.12 | 2113.039111 | -1.0145762 | 0.112947124 | -3.67053 | 0.000242 | 0.002035929 | ENSG00000173432 | SAA1 | 6288 |
| ENSG00000178966.16 | 829.2239133 | -1.015196057 | 0.098950103 | -4.19601 | 2.72E-05 | 0.000268247 | ENSG00000178966 | RMI1 | 80010 |
| ENSG00000073282.13 | 8223.083759 | -1.015285729 | 0.1077838 | -3.85295 | 0.000117 | 0.00104222 | ENSG00000073282 | TP63 | 8626 |
| ENSG00000013810.19 | 4214.839148 | -1.016260178 | 0.053203488 | -7.82393 | 5.12E-15 | 1.40E-13 | ENSG00000013810 | TACC3 | 10460 |
| ENSG00000148840.11 | 2518.262812 | -1.016761405 | 0.118801951 | -3.50804 | 0.000451 | 0.003601426 | ENSG00000148840 | PPRC1 | 23082 |
| ENSG00000110107.9 | 4255.88533 | -1.017098391 | 0.145992528 | -2.85698 | 0.004277 | 0.027271559 | ENSG00000110107 | PRPF19 | 27339 |
| ENSG00000103494.14 | 577.5995266 | -1.018104478 | 0.158745527 | -2.6338 | 0.008443 | 0.04945683 | ENSG00000103494 | RPGRIP1L | 23322 |
| ENSG00000116922.14 | 872.6587018 | -1.018562279 | 0.10773955 | -3.88495 | 0.000102 | 0.000922735 | ENSG00000116922 | C1orf109 | 54955 |
| ENSG00000008441.16 | 1459.883993 | -1.02088748 | 0.148500324 | -2.83425 | 0.004593 | 0.029004442 | ENSG00000008441 | NFIX | 4784 |
| ENSG00000074695.6 | 7293.949031 | -1.023222848 | 0.107529108 | -3.93589 | 8.29E-05 | 0.000759085 | ENSG00000074695 | LMAN1 | 3998 |
| ENSG00000236255.2 | 103.8980553 | -1.026501074 | 0.143380146 | -2.97462 | 0.002934 | 0.019490081 | ENSG00000236255 | NA | NA |
| ENSG00000120889.13 | 7979.403351 | -1.026887888 | 0.113599656 | -3.75783 | 0.000171 | 0.001481936 | ENSG00000120889 | TNFRSF10 | 8795 |
| ENSG00000100297.16 | 3076.297 | -1.027562 | 0.0815685 | -5.24175 | 1.59E-07 | 2.14E-06 | ENSG00000100297 | MCM5 | 4174 |
| ENSG00000167693.17 | 1872.995885 | -1.027764964 | 0.059393636 | -7.2022 | 5.92E-13 | 1.39E-11 | ENSG00000167693 | NXN | 64359 |
| ENSG00000100578.16 | 879.9102076 | -1.029126295 | 0.132268347 | -3.24436 | 0.001177 | 0.008576332 | ENSG00000100578 | KIAA0586 | 9786 |
| ENSG00000140563.15 | 787.0325551 | -1.029430882 | 0.084678746 | -5.07129 | 3.95E-07 | 5.03E-06 | ENSG00000140563 | MCTP2 | 55784 |
| ENSG00000071242.12 | 239.4430011 | -1.030459833 | 0.119380866 | -3.60577 | 0.000311 | 0.002559697 | ENSG00000071242 | RPS6KA2 | 6196 |
| ENSG00000175792.11 | 4088.324881 | -1.031197738 | 0.163327728 | -2.64008 | 0.008289 | 0.048711356 | ENSG00000175792 | RUVBL1 | 8607 |
| ENSG00000030066.13 | 1727.579075 | -1.031500484 | 0.106052652 | -4.06874 | 4.73E-05 | 0.000449261 | ENSG00000030066 | NUP160 | 23279 |
| ENSG00000138442.9 | 2267.4947 | -1.033984281 | 0.158978149 | -2.72984 | 0.006337 | 0.038487911 | ENSG00000138442 | WDR12 | 55759 |
| ENSG00000138134.12 | 866.8744626 | -1.034804874 | 0.085347218 | -5.09454 | 3.50E-07 | 4.48E-06 | ENSG00000138134 | STAMBPL | 57559 |
| ENSG00000138031.14 | 2133.286166 | -1.035917201 | 0.123666388 | -3.52494 | 0.000424 | 0.003398525 | ENSG00000138031 | ADCY3 | 109 |
| ENSG00000132773.12 | 733.4583172 | -1.037687664 | 0.134354663 | -3.2577 | 0.001123 | 0.008241789 | ENSG00000132773 | TOE1 | 114034 |
| ENSG00000160447.7 | 439.1970665 | -1.04039073 | 0.070880199 | -6.21317 | 5.19E-10 | 9.24E-09 | ENSG00000160447 | PKN3 | 29941 |
| ENSG00000164307.13 | 5258.987935 | -1.042504501 | 0.070367465 | -6.28848 | 3.21E-10 | 5.87E-09 | ENSG00000164307 | ERAP1 | 51752 |
| ENSG00000129317.14 | 915.7298056 | -1.042784806 | 0.129234861 | -3.4262 | 0.000612 | 0.004757634 | ENSG00000129317 | PUS7L | 83448 |
| ENSG00000152642.10 | 976.9957444 | -1.044635287 | 0.117245649 | -3.79234 | 0.000149 | 0.001303882 | ENSG00000152642 | GPD1L | 23171 |
| ENSG00000162433.15 | 3101.014962 | -1.044663782 | 0.117360209 | -3.78888 | 0.000151 | 0.001321302 | ENSG00000162433 | AK4 | 205 |
| ENSG00000110057.8 | 1062.422859 | -1.044728224 | 0.060216479 | -7.38549 | 1.52E-13 | 3.73E-12 | ENSG00000110057 | UNC93B1 | 81622 |
| ENSG00000172716.16 | 1462.119765 | -1.045961379 | 0.04960972 | -8.9894 | 2.49E-19 | 9.14E-18 | ENSG00000172716 | SLFN11 | 91607 |
| ENSG00000130038.9 | 422.1661794 | -1.046020559 | 0.156716879 | -2.84603 | 0.004427 | 0.028113366 | ENSG00000130038 | CRACR2A | 84766 |
| ENSG00000133302.13 | 1415.716703 | -1.046158393 | 0.125517843 | -3.55454 | 0.000379 | 0.003066662 | ENSG00000133302 | SLF1 | 84250 |
| ENSG00000166965.12 | 620.734958 | -1.047897575 | 0.076199234 | -5.87798 | 4.15E-09 | 6.66E-08 | ENSG00000166965 | RCCD1 | 91433 |
| ENSG00000213347.10 | 341.4754823 | -1.049313866 | 0.121364465 | -3.70219 | 0.000214 | 0.001809936 | ENSG00000213347 | MXD3 | 83463 |
| ENSG00000166860.3 | 695.1621516 | -1.049702213 | 0.068160716 | -6.59767 | 4.18E-11 | 8.22E-10 | ENSG00000166860 | ZBTB39 | 9880 |
| ENSG00000113569.16 | 3004.855803 | -1.049710566 | 0.130362305 | -3.4497 | 0.000561 | 0.004394308 | ENSG00000113569 | NUP155 | 9631 |
| ENSG00000247092.7 | 113.8406471 | -1.049730515 | 0.147718358 | -3.04451 | 0.002331 | 0.015860035 | ENSG00000247092 | NA | NA |
| ENSG00000128944.13 | 2074.781714 | -1.051005593 | 0.116586787 | -3.86841 | 0.00011 | 0.000983927 | ENSG00000128944 | KNSTRN | 90417 |
| ENSG00000049541.11 | 1908.45194 | -1.051533347 | 0.106390807 | -4.2441 | 2.19E-05 | 0.000219653 | ENSG00000049541 | RFC2 | 5982 |
| ENSG00000082497.12 | 80.95804213 | -1.052927012 | 0.158371858 | -2.8599 | 0.004238 | 0.027035412 | ENSG00000082497 | SERTAD4 | 56256 |
| ENSG00000077348.9 | 716.8752 | -1.053408 | 0.1277426 | -3.54939 | 0.000386 | 0.003125419 | ENSG00000077348 | EXOSC5 | 56915 |

| | | | | | | | | | |
|---|---|---|---|---|---|---|---|---|---|
| ENSG00000119397.16 | 890.2169334 | -1.053476294 | 0.071831113 | -6.31309 | 2.74E-10 | 5.05E-09 | ENSG00000119397 | CNTRL | 11064 |
| ENSG00000174206.12 | 293.6647828 | -1.05379147 | 0.086402102 | -5.25209 | 1.50E-07 | 2.03E-06 | ENSG00000174206 | C12orf66 | 144577 |
| ENSG00000188486.3 | 2035.154742 | -1.05475273 | 0.067947525 | -6.6927 | 2.19E-11 | 4.45E-10 | ENSG00000188486 | H2AX | 3014 |
| ENSG00000163811.12 | 5039.369571 | -1.055061639 | 0.147023725 | -3.09516 | 0.001967 | 0.013601987 | ENSG00000163811 | WDR43 | 23160 |
| ENSG00000127399.15 | 274.6321322 | -1.055179123 | 0.138509641 | -3.28626 | 0.001015 | 0.007518299 | ENSG00000127399 | LRRC61 | 65999 |
| ENSG00000149557.13 | 2572.198349 | -1.056672602 | 0.08646883 | -5.28136 | 1.28E-07 | 1.75E-06 | ENSG00000149557 | FEZ1 | 9638 |
| ENSG00000106004.5 | 84.48166947 | -1.056732115 | 0.170581613 | -2.6775 | 0.007417 | 0.044245163 | ENSG00000106004 | HOXA5 | 3202 |
| ENSG00000109667.12 | 1048.423347 | -1.059213958 | 0.126275081 | -3.63662 | 0.000276 | 0.002298173 | ENSG00000109667 | SLC2A9 | 56606 |
| ENSG00000111196.10 | 875.1192842 | -1.05921583 | 0.1098417 | -4.18071 | 2.91E-05 | 0.000285269 | ENSG00000111196 | MAGOHB | 55110 |
| ENSG00000123146.20 | 757.4072201 | -1.061041886 | 0.111171138 | -4.14714 | 3.37E-05 | 0.000327339 | ENSG00000123146 | ADGRE5 | 976 |
| ENSG00000163219.11 | 128.8749127 | -1.06255665 | 0.144266896 | -3.20626 | 0.001345 | 0.009672981 | ENSG00000163219 | ARHGAP2 | 9938 |
| ENSG00000118523.6 | 453.493424 | -1.063966387 | 0.136014063 | -3.41116 | 0.000647 | 0.005003109 | ENSG00000118523 | CCN2 | 1490 |
| ENSG00000213468.7 | 764.4944 | -1.065301 | 0.1690939 | -2.75173 | 0.005928 | 0.03632199 | ENSG00000213468 | FIRRE | 286467 |
| ENSG00000175920.18 | 502.3078842 | -1.065378247 | 0.079641217 | -5.84343 | 5.11E-09 | 8.15E-08 | ENSG00000175920 | DOK7 | 285489 |
| ENSG00000138642.14 | 1123.864556 | -1.065796149 | 0.176546664 | -2.63837 | 0.00833 | 0.048881103 | ENSG00000138642 | HERC6 | 55008 |
| ENSG00000166845.15 | 792.3644504 | -1.065805898 | 0.103231076 | -4.51226 | 6.41E-06 | 7.00E-05 | ENSG00000166845 | C18orf54 | 162681 |
| ENSG00000173918.15 | 330.2235953 | -1.066308657 | 0.09530243 | -4.89294 | 9.93E-07 | 1.20E-05 | ENSG00000173918 | C1QTNF1 | 114897 |
| ENSG00000162337.12 | 1646.450688 | -1.06685051 | 0.060528212 | -7.71294 | 1.23E-14 | 3.26E-13 | ENSG00000162337 | LRP5 | 4041 |
| ENSG00000104320.13 | 3278.015954 | -1.067907237 | 0.128843719 | -3.63159 | 0.000282 | 0.002339767 | ENSG00000104320 | NBN | 4683 |
| ENSG00000163918.10 | 1379.081664 | -1.069324957 | 0.161254687 | -2.91046 | 0.003609 | 0.023405293 | ENSG00000163918 | RFC4 | 5984 |
| ENSG00000102098.19 | 224.6217 | -1.06962 | 0.1231862 | -3.81228 | 0.000138 | 0.001209854 | ENSG00000102098 | SCML2 | 10389 |
| ENSG00000123416.15 | 12994.52999 | -1.069991675 | 0.148532848 | -3.16423 | 0.001555 | 0.010993979 | ENSG00000123416 | TUBA1B | 10376 |
| ENSG00000125454.12 | 372.3374929 | -1.070470873 | 0.116546359 | -4.03677 | 5.42E-05 | 0.000509429 | ENSG00000125454 | SLC25A19 | 60386 |
| ENSG00000116984.13 | 2238.197951 | -1.070727528 | 0.151430891 | -3.10853 | 0.00188 | 0.01305209 | ENSG00000116984 | MTR | 4548 |
| ENSG00000164308.16 | 724.1795584 | -1.071998717 | 0.067437874 | -6.99902 | 2.58E-12 | 5.68E-11 | ENSG00000164308 | ERAP2 | 64167 |
| ENSG00000187790.11 | 769.7003282 | -1.072608899 | 0.149808215 | -3.15476 | 0.001606 | 0.011330957 | ENSG00000187790 | FANCM | 57697 |
| ENSG00000166881.10 | 1825.038156 | -1.07359537 | 0.072751096 | -6.5098 | 7.52E-11 | 1.45E-09 | ENSG00000166881 | NEMP1 | 23306 |
| ENSG00000170549.4 | 245.5157437 | -1.074045137 | 0.177204422 | -2.67513 | 0.00747 | 0.044489076 | ENSG00000170549 | IRX1 | 79192 |
| ENSG00000105810.9 | 5433.803297 | -1.076094644 | 0.161990395 | -2.93903 | 0.003292 | 0.021567561 | ENSG00000105810 | CDK6 | 1021 |
| ENSG00000152455.15 | 1012.369017 | -1.076571201 | 0.138956584 | -3.42964 | 0.000604 | 0.004703248 | ENSG00000152455 | SUV39H2 | 79723 |
| ENSG00000071794.16 | 1052.122413 | -1.076764781 | 0.106980211 | -4.45657 | 8.33E-06 | 8.95E-05 | ENSG00000071794 | HLTF | 6596 |
| ENSG00000006062.17 | 466.5697446 | -1.076878501 | 0.103638496 | -4.60136 | 4.20E-06 | 4.67E-05 | ENSG00000006062 | MAP3K14 | 9020 |
| ENSG00000160208.13 | 3741.789 | -1.077652 | 0.1231217 | -3.87951 | 0.000105 | 0.000942177 | ENSG00000160208 | RRP1B | 23076 |
| ENSG00000119042.17 | 698.1410808 | -1.08025422 | 0.132626132 | -3.62111 | 0.000293 | 0.002427495 | ENSG00000119042 | SATB2 | 23314 |
| ENSG00000157833.13 | 75.74386863 | -1.080265173 | 0.16327405 | -2.94147 | 0.003267 | 0.021414421 | ENSG00000157833 | GAREM2 | 150946 |
| ENSG00000084112.15 | 3646.98763 | -1.0813462 | 0.104191151 | -4.61984 | 3.84E-06 | 4.30E-05 | ENSG00000084112 | SSH1 | 54434 |
| ENSG00000146909.8 | 2626.541082 | -1.081514052 | 0.125057941 | -3.85033 | 0.000118 | 0.001051684 | ENSG00000146909 | NOM1 | 64434 |
| ENSG00000143643.13 | 746.3601947 | -1.081570923 | 0.1250135 | -3.85215 | 0.000117 | 0.00104458 | ENSG00000143643 | TTC13 | 79573 |
| ENSG00000161847.14 | 644.9638006 | -1.082436771 | 0.10651043 | -4.52948 | 5.91E-06 | 6.48E-05 | ENSG00000161847 | RAVER1 | 125950 |
| ENSG00000213186.8 | 321.9857502 | -1.082459612 | 0.087498429 | -5.51392 | 3.51E-08 | 5.08E-07 | ENSG00000213186 | TRIM59 | 286827 |
| ENSG00000064115.11 | 4296.359541 | -1.082595559 | 0.087006974 | -5.54663 | 2.91E-08 | 4.28E-07 | ENSG00000064115 | TM7SF3 | 51768 |
| ENSG00000247077.7 | 2891.423613 | -1.083325464 | 0.078786127 | -6.13465 | 8.53E-10 | 1.48E-08 | ENSG00000247077 | PGAM5 | 192111 |

| | | | | | | | | | |
|---|---|---|---|---|---|---|---|---|---|
| ENSG00000224934.5 | 57.86064668 | -1.083854657 | 0.173852084 | -2.78314 | 0.005384 | 0.033392622 | ENSG00000224934 | NA | NA |
| ENSG00000120254.15 | 3799.070703 | -1.083878225 | 0.133011593 | -3.63787 | 0.000275 | 0.002288488 | ENSG00000120254 | MTHFD1L | 25902 |
| ENSG00000167034.10 | 669.5135625 | -1.084211844 | 0.13286654 | -3.64435 | 0.000268 | 0.002234373 | ENSG00000167034 | NKX3-1 | 4824 |
| ENSG00000163362.11 | 1885.823905 | -1.085154124 | 0.127379951 | -3.80872 | 0.00014 | 0.001225717 | ENSG00000163362 | INAVA | 55765 |
| ENSG00000132274.16 | 1504.85965 | -1.085204102 | 0.069312831 | -7.00021 | 2.56E-12 | 5.64E-11 | ENSG00000132274 | TRIM22 | 10346 |
| ENSG00000144355.15 | 285.7360074 | -1.085244569 | 0.171834196 | -2.82391 | 0.004744 | 0.029857848 | ENSG00000144355 | DLX1 | 1745 |
| ENSG00000186283.14 | 1304.504955 | -1.086011588 | 0.140022084 | -3.47096 | 0.000519 | 0.004091881 | ENSG00000186283 | TOR3A | 64222 |
| ENSG00000128951.14 | 3229.950268 | -1.087922552 | 0.123358101 | -3.95533 | 7.64E-05 | 0.000704277 | ENSG00000128951 | DUT | 1854 |
| ENSG00000287778.1 | 506.2280254 | -1.090332842 | 0.164192101 | -2.98634 | 0.002823 | 0.018843083 | ENSG00000287778 | NA | NA |
| ENSG00000111602.12 | 2874.211876 | -1.09059702 | 0.104690713 | -4.68616 | 2.78E-06 | 3.17E-05 | ENSG00000111602 | TIMELESS | 8914 |
| ENSG00000165304.8 | 3610.761011 | -1.090824461 | 0.10656142 | -4.60602 | 4.10E-06 | 4.57E-05 | ENSG00000165304 | MELK | 9833 |
| ENSG00000158402.20 | 842.1262826 | -1.091187029 | 0.137768164 | -3.56532 | 0.000363 | 0.002957786 | ENSG00000158402 | CDC25C | 995 |
| ENSG00000137642.13 | 4096.60284 | -1.091381246 | 0.148088067 | -3.31817 | 0.000906 | 0.006792977 | ENSG00000137642 | SORL1 | 6653 |
| ENSG00000182199.11 | 5799.574893 | -1.092061372 | 0.10720486 | -4.58992 | 4.43E-06 | 4.92E-05 | ENSG00000182199 | SHMT2 | 6472 |
| ENSG00000185347.17 | 412.2762973 | -1.092245413 | 0.107746673 | -4.56854 | 4.91E-06 | 5.41E-05 | ENSG00000185347 | TEDC1 | 283643 |
| ENSG00000088305.18 | 130.9387 | -1.092675 | 0.1472197 | -3.34653 | 0.000818 | 0.006202989 | ENSG00000088305 | DNMT3B | 1789 |
| ENSG00000175591.11 | 457.8854222 | -1.092963329 | 0.12367415 | -3.98599 | 6.72E-05 | 0.000623786 | ENSG00000175591 | P2RY2 | 5029 |
| ENSG00000092201.10 | 11126.01312 | -1.092964474 | 0.134447131 | -3.6666 | 0.000246 | 0.002064191 | ENSG00000092201 | SUPT16H | 11198 |
| ENSG00000185838.14 | 180.4978 | -1.093611 | 0.1163117 | -4.24386 | 2.20E-05 | 0.000219806 | ENSG00000185838 | GNB1L | 54584 |
| ENSG00000146143.18 | 999.5916975 | -1.095113012 | 0.093507691 | -5.29489 | 1.19E-07 | 1.63E-06 | ENSG00000146143 | PRIM2 | 5558 |
| ENSG00000126457.21 | 9282.621 | -1.095199 | 0.1593992 | -3.10666 | 0.001892 | 0.01312132 | ENSG00000126457 | PRMT1 | 3276 |
| ENSG00000249815.1 | 376.5026608 | -1.095372039 | 0.102156994 | -4.84913 | 1.24E-06 | 1.48E-05 | ENSG00000249815 | NA | NA |
| ENSG00000136824.19 | 5563.220821 | -1.095835684 | 0.119637546 | -4.14448 | 3.41E-05 | 0.000331033 | ENSG00000136824 | SMC2 | 10592 |
| ENSG00000280202.1 | 134.2972436 | -1.095917007 | 0.124738101 | -3.97567 | 7.02E-05 | 0.000649201 | ENSG00000280202 | NA | NA |
| ENSG00000169220.18 | 504.2829981 | -1.096040846 | 0.175030372 | -2.83403 | 0.004597 | 0.02901117 | ENSG00000169220 | RGS14 | 10636 |
| ENSG00000152990.14 | 1753.158559 | -1.096385366 | 0.142108138 | -3.49301 | 0.000478 | 0.00379547 | ENSG00000152990 | ADGRA3 | 166647 |
| ENSG00000164099.3 | 201.8748184 | -1.096797871 | 0.149846246 | -3.31538 | 0.000915 | 0.006846597 | ENSG00000164099 | PRSS12 | 8492 |
| ENSG00000175764.14 | 222.1211273 | -1.097614336 | 0.102623706 | -4.84892 | 1.24E-06 | 1.48E-05 | ENSG00000175764 | TTLL11 | 158135 |
| ENSG00000196230.13 | 8184.839391 | -1.097862013 | 0.150628507 | -3.30523 | 0.000949 | 0.007072728 | ENSG00000196230 | TUBB | 203068 |
| ENSG00000237950.2 | 48.0929764 | -1.098815553 | 0.187971119 | -2.65368 | 0.007962 | 0.047008494 | ENSG00000237950 | NA | NA |
| ENSG00000171723.15 | 863.7048328 | -1.100742742 | 0.151941487 | -3.29563 | 0.000982 | 0.007292399 | ENSG00000171723 | GPHN | 10243 |
| ENSG00000235173.7 | 1552.738833 | -1.102948686 | 0.121781611 | -4.12992 | 3.63E-05 | 0.000351682 | ENSG00000235173 | HGH1 | 51236 |
| ENSG00000161980.5 | 804.57743 | -1.103532686 | 0.161332467 | -3.12109 | 0.001802 | 0.012583624 | ENSG00000161980 | POLR3K | 51728 |
| ENSG00000180198.16 | 5630.905583 | -1.105793124 | 0.099038285 | -5.10705 | 3.27E-07 | 4.22E-06 | ENSG00000180198 | RCC1 | 1104 |
| ENSG00000182481.9 | 9863.235396 | -1.108796063 | 0.137285703 | -3.70611 | 0.00021 | 0.001784629 | ENSG00000182481 | KPNA2 | 3838 |
| ENSG00000132386.11 | 94.79621634 | -1.110057267 | 0.139366809 | -3.65982 | 0.000252 | 0.002114293 | ENSG00000132386 | SERPINF1 | 5176 |
| ENSG00000100526.20 | 1210.24432 | -1.11310552 | 0.124831563 | -4.11038 | 3.95E-05 | 0.000380731 | ENSG00000100526 | CDKN3 | 1033 |
| ENSG00000158711.13 | 2862.794735 | -1.113144916 | 0.107449804 | -4.77567 | 1.79E-06 | 2.09E-05 | ENSG00000158711 | ELK4 | 2005 |
| ENSG00000121057.13 | 2824.135496 | -1.113877661 | 0.139625296 | -3.68041 | 0.000233 | 0.001963667 | ENSG00000121057 | AKAP1 | 8165 |
| ENSG00000189046.11 | 437.4837583 | -1.114148759 | 0.15843878 | -3.24509 | 0.001174 | 0.008561317 | ENSG00000189046 | ALKBH2 | 121642 |
| ENSG00000143401.15 | 5540.701516 | -1.114592645 | 0.108702537 | -4.73395 | 2.20E-06 | 2.54E-05 | ENSG00000143401 | ANP32E | 81611 |
| ENSG00000096996.15 | 64.94075 | -1.114607 | 0.1862224 | -2.7634 | 0.00572 | 0.03519402 | ENSG00000096996 | IL12RB1 | 3594 |

| | | | | | | | | | |
|---|---|---|---|---|---|---|---|---|---|
| ENSG00000132646.11 | 7030.276 | -1.11604 | 0.1545457 | -3.33908 | 0.000841 | 0.006350089 | ENSG00000132646 | PCNA | 5111 |
| ENSG00000077152.11 | 1207.735499 | -1.116044006 | 0.150870455 | -3.42044 | 0.000625 | 0.004849587 | ENSG00000077152 | UBE2T | 29089 |
| ENSG00000165891.16 | 2674.633301 | -1.116105661 | 0.163388534 | -3.15876 | 0.001584 | 0.011185353 | ENSG00000165891 | E2F7 | 144455 |
| ENSG00000137054.16 | 1837.208711 | -1.116786967 | 0.180539821 | -2.86245 | 0.004204 | 0.026863172 | ENSG00000137054 | POLR1E | 64425 |
| ENSG00000132768.14 | 1017.128382 | -1.117542883 | 0.151081348 | -3.42559 | 0.000613 | 0.004765577 | ENSG00000132768 | DPH2 | 1802 |
| ENSG00000173621.9 | 947.1109886 | -1.118194874 | 0.132047073 | -3.92432 | 8.70E-05 | 0.000793508 | ENSG00000173621 | LRFN4 | 78999 |
| ENSG00000131844.16 | 4835.748996 | -1.118383741 | 0.117194684 | -4.42327 | 9.72E-06 | 0.000103237 | ENSG00000131844 | MCCC2 | 64087 |
| ENSG00000109320.13 | 3922.379054 | -1.118497823 | 0.115840359 | -4.47597 | 7.61E-06 | 8.22E-05 | ENSG00000109320 | NFKB1 | 4790 |
| ENSG00000230836.1 | 70.28785964 | -1.119217379 | 0.188780416 | -2.75038 | 0.005953 | 0.036425139 | ENSG00000230836 | NA | NA |
| ENSG00000000460.17 | 937.5956132 | -1.120270432 | 0.111041502 | -4.68537 | 2.79E-06 | 3.18E-05 | ENSG00000000460 | C1orf112 | 55732 |
| ENSG00000145431.11 | 789.5588745 | -1.120468009 | 0.129342974 | -4.02394 | 5.72E-05 | 0.00053669 | ENSG00000145431 | PDGFC | 56034 |
| ENSG00000111012.10 | 1823.06971 | -1.120488165 | 0.072789639 | -7.15058 | 8.64E-13 | 1.98E-11 | ENSG00000111012 | CYP27B1 | 1594 |
| ENSG00000151849.15 | 1638.588897 | -1.120928314 | 0.1701717 | -3.06119 | 0.002205 | 0.015091325 | ENSG00000151849 | CENPJ | 55835 |
| ENSG00000188191.15 | 1419.502133 | -1.121285185 | 0.11360437 | -4.5886 | 4.46E-06 | 4.95E-05 | ENSG00000188191 | PRKAR1B | 5575 |
| ENSG00000099901.17 | 7897.164 | -1.121596 | 0.1454981 | -3.5849 | 0.000337 | 0.002759684 | ENSG00000099901 | RANBP1 | 5902 |
| ENSG00000188976.11 | 5541.813707 | -1.12172274 | 0.115325556 | -4.52391 | 6.07E-06 | 6.64E-05 | ENSG00000188976 | NOC2L | 26155 |
| ENSG00000105281.12 | 4305.176 | -1.124343 | 0.165795 | -3.1626 | 0.001564 | 0.01105075 | ENSG00000105281 | SLC1A5 | 6510 |
| ENSG00000162437.14 | 1049.109592 | -1.124497802 | 0.101515566 | -5.16667 | 2.38E-07 | 3.13E-06 | ENSG00000162437 | RAVER2 | 55225 |
| ENSG00000141574.7 | 492.9636099 | -1.124771718 | 0.123428296 | -4.25163 | 2.12E-05 | 0.000212794 | ENSG00000141574 | SECTM1 | 6398 |
| ENSG00000076356.7 | 1774.711544 | -1.125910299 | 0.126134017 | -4.16946 | 3.05E-05 | 0.000298839 | ENSG00000076356 | PLXNA2 | 5362 |
| ENSG00000169851.15 | 281.2301251 | -1.126714815 | 0.112806618 | -4.66918 | 3.02E-06 | 3.42E-05 | ENSG00000169851 | PCDH7 | 5099 |
| ENSG00000120696.8 | 356.7488625 | -1.127671498 | 0.10835791 | -4.86971 | 1.12E-06 | 1.34E-05 | ENSG00000120696 | KBTBD7 | 84078 |
| ENSG00000285756.2 | 694.7394 | -1.1277 | 0.09370644 | -5.63141 | 1.79E-08 | 2.69E-07 | ENSG00000285756 | LOC38990 | 389906 |
| ENSG00000111788.10 | 264.275797 | -1.128084832 | 0.174971224 | -3.01812 | 0.002543 | 0.017168281 | ENSG00000111788 | NA | NA |
| ENSG00000114767.7 | 1413.357959 | -1.128397284 | 0.154373518 | -3.42285 | 0.00062 | 0.00480969 | ENSG00000114767 | RRP9 | 9136 |
| ENSG00000137337.15 | 323.3347982 | -1.128465881 | 0.182167768 | -2.90098 | 0.00372 | 0.024019347 | ENSG00000137337 | MDC1 | 9656 |
| ENSG00000205208.5 | 681.9978116 | -1.128800103 | 0.155564011 | -3.39924 | 0.000676 | 0.00521121 | ENSG00000205208 | C4orf46 | 201725 |
| ENSG00000274925.1 | 58.77738033 | -1.131017112 | 0.160216615 | -3.31437 | 0.000918 | 0.006866692 | ENSG00000274925 | NA | NA |
| ENSG00000108561.8 | 9421.295139 | -1.131035361 | 0.168983298 | -3.14253 | 0.001675 | 0.011768316 | ENSG00000108561 | C1QBP | 708 |
| ENSG00000150990.8 | 1447.583175 | -1.131707127 | 0.12746872 | -4.17128 | 3.03E-05 | 0.000296572 | ENSG00000150990 | DHX37 | 57647 |
| ENSG00000171823.7 | 505.7720971 | -1.132629877 | 0.122730294 | -4.33984 | 1.43E-05 | 0.000147243 | ENSG00000171823 | FBXL14 | 144699 |
| ENSG00000263535.2 | 86.64531698 | -1.133527299 | 0.198006621 | -2.69449 | 0.00705 | 0.042288053 | ENSG00000263535 | NA | NA |
| ENSG00000065308.5 | 1678.815532 | -1.134143431 | 0.117571664 | -4.54313 | 5.54E-06 | 6.09E-05 | ENSG00000065308 | TRAM2 | 9697 |
| ENSG00000111335.12 | 1922.56342 | -1.134411593 | 0.188335772 | -2.83755 | 0.004546 | 0.028734102 | ENSG00000111335 | OAS2 | 4939 |
| ENSG00000153044.10 | 2091.68806 | -1.134978172 | 0.104366669 | -5.12595 | 2.96E-07 | 3.83E-06 | ENSG00000153044 | CENPH | 64946 |
| ENSG00000158715.6 | 434.1269278 | -1.135392952 | 0.097181157 | -5.50923 | 3.60E-08 | 5.20E-07 | ENSG00000158715 | SLC45A3 | 85414 |
| ENSG00000095539.15 | 39.44150968 | -1.135518747 | 0.201610164 | -2.65621 | 0.007902 | 0.046756766 | ENSG00000095539 | SEMA4G | 57715 |
| ENSG00000179409.11 | 2071.640575 | -1.135895861 | 0.145271706 | -3.68892 | 0.000225 | 0.001903321 | ENSG00000179409 | GEMIN4 | 50628 |
| ENSG00000128059.8 | 2219.234811 | -1.137137621 | 0.140317985 | -3.828 | 0.000129 | 0.0011426 | ENSG00000128059 | PPAT | 5471 |
| ENSG00000151131.11 | 605.7713847 | -1.137421387 | 0.108516848 | -4.95242 | 7.33E-07 | 9.04E-06 | ENSG00000151131 | C12orf45 | 121053 |
| ENSG00000198039.11 | 227.3746002 | -1.138164375 | 0.100664723 | -5.34611 | 8.99E-08 | 1.25E-06 | ENSG00000198039 | ZNF273 | 10793 |
| ENSG00000112149.10 | 279.7853185 | -1.138841951 | 0.165634688 | -3.2532 | 0.001141 | 0.008350668 | ENSG00000112149 | CD83 | 9308 |

| | | | | | | | | | |
|---|---|---|---|---|---|---|---|---|---|
| ENSG00000136848.17 | 3444.272177 | -1.139069031 | 0.096121394 | -5.60821 | 2.04E-08 | 3.05E-07 | ENSG00000136848 | DAB2IP | 153090 |
| ENSG00000134247.10 | 5923.208575 | -1.139863722 | 0.088410425 | -6.10634 | 1.02E-09 | 1.75E-08 | ENSG00000134247 | PTGFRN | 5738 |
| ENSG00000183853.18 | 4852.426347 | -1.14124195 | 0.16985004 | -3.18659 | 0.00144 | 0.010280471 | ENSG00000183853 | KIRREL1 | 55243 |
| ENSG00000131127.14 | 464.9447131 | -1.141333426 | 0.089292931 | -6.06244 | 1.34E-09 | 2.27E-08 | ENSG00000131127 | ZNF141 | 7700 |
| ENSG00000139880.19 | 278.4723196 | -1.141949444 | 0.117348344 | -4.6183 | 3.87E-06 | 4.33E-05 | ENSG00000139880 | CDH24 | 64403 |
| ENSG00000169255.15 | 382.284025 | -1.14216867 | 0.137802408 | -3.93439 | 8.34E-05 | 0.000763313 | ENSG00000169255 | B3GALNT1 | 8706 |
| ENSG00000169750.9 | 86.55499895 | -1.142533641 | 0.152276876 | -3.56281 | 0.000367 | 0.002982523 | ENSG00000169750 | RAC3 | 5881 |
| ENSG00000048342.16 | 456.4874488 | -1.142799053 | 0.170478574 | -3.18397 | 0.001453 | 0.010351578 | ENSG00000048342 | CC2D2A | 57545 |
| ENSG00000185885.16 | 964.1050241 | -1.143849685 | 0.103227536 | -5.26846 | 1.38E-07 | 1.87E-06 | ENSG00000185885 | IFITM1 | 8519 |
| ENSG00000221829.9 | 1293.016757 | -1.144173873 | 0.087173357 | -6.24243 | 4.31E-10 | 7.76E-09 | ENSG00000221829 | FANCG | 2189 |
| ENSG00000170264.13 | 248.8076677 | -1.144373223 | 0.150966845 | -3.60591 | 0.000311 | 0.002559697 | ENSG00000170264 | FAM161A | 84140 |
| ENSG00000160179.18 | 383.8209 | -1.144445 | 0.1930311 | -2.8205 | 0.004795 | 0.03011999 | ENSG00000160179 | ABCG1 | 9619 |
| ENSG00000204856.12 | 283.7822882 | -1.144897131 | 0.171628451 | -3.17486 | 0.001499 | 0.010653513 | ENSG00000204856 | FAM216A | 29902 |
| ENSG00000151229.13 | 303.3754936 | -1.145759678 | 0.108132364 | -5.04714 | 4.48E-07 | 5.66E-06 | ENSG00000151229 | SLC2A13 | 114134 |
| ENSG00000214357.8 | 469.8876655 | -1.147403381 | 0.141025699 | -3.88159 | 0.000104 | 0.000934945 | ENSG00000214357 | NEURL1B | 54492 |
| ENSG00000204394.13 | 3944.054734 | -1.147417524 | 0.136295278 | -4.01641 | 5.91E-05 | 0.000552755 | ENSG00000204394 | VARS1 | 7407 |
| ENSG00000161692.18 | 630.4167578 | -1.151156925 | 0.129957865 | -4.24104 | 2.22E-05 | 0.00022225 | ENSG00000161692 | DBF4B | 80174 |
| ENSG00000130826.18 | 8385.836 | -1.151288 | 0.1880005 | -2.93238 | 0.003364 | 0.02199735 | ENSG00000130826 | DKC1 | 1736 |
| ENSG00000152056.17 | 356.60237 | -1.151455121 | 0.109760523 | -5.02417 | 5.06E-07 | 6.35E-06 | ENSG00000152056 | AP1S3 | 130340 |
| ENSG00000143815.15 | 4922.914515 | -1.152555773 | 0.135702984 | -4.0718 | 4.67E-05 | 0.000444024 | ENSG00000143815 | LBR | 3930 |
| ENSG00000260260.1 | 549.3975516 | -1.152635143 | 0.1399227 | -3.94957 | 7.83E-05 | 0.000719448 | ENSG00000260260 | SNHG19 | 1.01E+08 |
| ENSG00000107099.15 | 98.63643817 | -1.154377925 | 0.154752784 | -3.58235 | 0.000341 | 0.002784216 | ENSG00000107099 | DOCK8 | 81704 |
| ENSG00000229152.2 | 416.8920796 | -1.154973508 | 0.114121416 | -4.86301 | 1.16E-06 | 1.38E-05 | ENSG00000229152 | NA | NA |
| ENSG00000087299.12 | 900.0460984 | -1.155372912 | 0.114694758 | -4.84218 | 1.28E-06 | 1.52E-05 | ENSG00000087299 | L2HGDH | 79944 |
| ENSG00000261236.8 | 4407.923877 | -1.15541562 | 0.13436139 | -4.13374 | 3.57E-05 | 0.000346261 | ENSG00000261236 | BOP1 | 23246 |
| ENSG00000014138.9 | 260.2150978 | -1.155698395 | 0.14811425 | -3.75182 | 0.000176 | 0.001512488 | ENSG00000014138 | POLA2 | 23649 |
| ENSG00000177084.16 | 3768.316892 | -1.158236279 | 0.115640352 | -4.82735 | 1.38E-06 | 1.63E-05 | ENSG00000177084 | POLE | 5426 |
| ENSG00000091127.14 | 1880.861585 | -1.158361023 | 0.162183559 | -3.44277 | 0.000576 | 0.00450045 | ENSG00000091127 | PUS7 | 54517 |
| ENSG00000163840.10 | 2872.622329 | -1.158918687 | 0.084637683 | -6.60366 | 4.01E-11 | 7.92E-10 | ENSG00000163840 | DTX3L | 151636 |
| ENSG00000065911.12 | 4394.724526 | -1.159698483 | 0.061832105 | -9.05191 | 1.40E-19 | 5.22E-18 | ENSG00000065911 | MTHFD2 | 10797 |
| ENSG00000143674.11 | 805.3175026 | -1.159733388 | 0.10682822 | -5.23956 | 1.61E-07 | 2.16E-06 | ENSG00000143674 | MAP3K21 | 84451 |
| ENSG00000144893.12 | 58.84397827 | -1.160416996 | 0.179752698 | -3.11771 | 0.001823 | 0.012701992 | ENSG00000144893 | MED12L | 116931 |
| ENSG00000166451.13 | 1318.189652 | -1.16044573 | 0.14951966 | -3.74831 | 0.000178 | 0.001532349 | ENSG00000166451 | CENPN | 55839 |
| ENSG00000145819.18 | 533.4640299 | -1.162522483 | 0.100447099 | -5.60019 | 2.14E-08 | 3.20E-07 | ENSG00000145819 | ARHGAP2 | 23092 |
| ENSG00000058335.15 | 143.5594535 | -1.163004371 | 0.13810438 | -4.07666 | 4.57E-05 | 0.000435791 | ENSG00000058335 | RASGRF1 | 5923 |
| ENSG00000162714.12 | 1345.606788 | -1.163124542 | 0.114677694 | -4.9105 | 9.08E-07 | 1.11E-05 | ENSG00000162714 | ZNF496 | 84838 |
| ENSG00000005073.5 | 75.90286162 | -1.16371972 | 0.172738001 | -3.26344 | 0.001101 | 0.008087955 | ENSG00000005073 | HOXA11 | 3207 |
| ENSG00000083857.14 | 20235.04856 | -1.165087622 | 0.194292891 | -2.90843 | 0.003632 | 0.023546009 | ENSG00000083857 | FAT1 | 2195 |
| ENSG00000225313.5 | 180.1857598 | -1.166772396 | 0.199117328 | -2.84642 | 0.004421 | 0.028091765 | ENSG00000225313 | NA | NA |
| ENSG00000176974.20 | 715.9968757 | -1.167126606 | 0.085236289 | -6.65358 | 2.86E-11 | 5.74E-10 | ENSG00000176974 | SHMT1 | 6470 |
| ENSG00000170425.3 | 1193.20618 | -1.168773118 | 0.194858996 | -2.9189 | 0.003513 | 0.022847799 | ENSG00000170425 | ADORA2B | 136 |
| ENSG00000198780.12 | 1725.305388 | -1.168947912 | 0.098788923 | -5.75923 | 8.45E-09 | 1.32E-07 | ENSG00000198780 | FAM169A | 26049 |

| | | | | | | | | | |
|---|---|---|---|---|---|---|---|---|---|
| ENSG00000172197.10 | 222.6531828 | -1.169414712 | 0.120437851 | -4.72787 | 2.27E-06 | 2.61E-05 | ENSG00000172197 | MBOAT1 | 154141 |
| ENSG00000277883.1 | 82.83489 | -1.169771 | 0.1651182 | -3.45069 | 0.000559 | 0.004382106 | ENSG00000277883 | NA | NA |
| ENSG00000130193.8 | 868.0210033 | -1.169886805 | 0.080397578 | -7.08836 | 1.36E-12 | 3.06E-11 | ENSG00000130193 | THEM6 | 51337 |
| ENSG00000130204.13 | 5293.575 | -1.170506 | 0.171678 | -3.32312 | 0.00089 | 0.006686901 | ENSG00000130204 | TOMM40 | 10452 |
| ENSG00000135372.9 | 3757.217109 | -1.171534946 | 0.147901476 | -3.8643 | 0.000111 | 0.000998324 | ENSG00000135372 | NAT10 | 55226 |
| ENSG00000173894.11 | 414.7925228 | -1.171539856 | 0.141966143 | -4.02589 | 5.68E-05 | 0.000532821 | ENSG00000173894 | CBX2 | 84733 |
| ENSG00000138778.11 | 4085.658854 | -1.17232771 | 0.086859287 | -6.58914 | 4.42E-11 | 8.69E-10 | ENSG00000138778 | CENPE | 1062 |
| ENSG00000160190.14 | 490.0892 | -1.173043 | 0.07745267 | -7.39862 | 1.38E-13 | 3.40E-12 | ENSG00000160190 | SLC37A1 | 54020 |
| ENSG00000141040.15 | 49.26012278 | -1.173460014 | 0.208871043 | -2.74552 | 0.006041 | 0.036889021 | ENSG00000141040 | ZNF287 | 57336 |
| ENSG00000177192.13 | 1354.813316 | -1.174357087 | 0.122974344 | -4.67054 | 3.00E-06 | 3.40E-05 | ENSG00000177192 | PUS1 | 80324 |
| ENSG00000166801.15 | 3089.418022 | -1.17467376 | 0.082289312 | -6.98358 | 2.88E-12 | 6.32E-11 | ENSG00000166801 | FAM111A | 63901 |
| ENSG00000173821.19 | 10963.46753 | -1.174942972 | 0.0848745 | -6.77404 | 1.25E-11 | 2.60E-10 | ENSG00000173821 | RNF213 | 57674 |
| ENSG00000160957.13 | 1630.171233 | -1.175332536 | 0.157246037 | -3.6588 | 0.000253 | 0.002122009 | ENSG00000160957 | RECQL4 | 9401 |
| ENSG00000128298.17 | 773.9388 | -1.17536 | 0.1180412 | -4.87423 | 1.09E-06 | 1.31E-05 | ENSG00000128298 | BAIAP2L2 | 80115 |
| ENSG00000109738.11 | 146.8376966 | -1.175732457 | 0.123983644 | -4.64362 | 3.42E-06 | 3.85E-05 | ENSG00000109738 | GLRB | 2743 |
| ENSG00000230084.6 | 80.9226165 | -1.175877687 | 0.171344931 | -3.36093 | 0.000777 | 0.005912863 | ENSG00000230084 | NA | NA |
| ENSG00000204519.11 | 454.6183 | -1.176687 | 0.1567028 | -3.68013 | 0.000233 | 0.001964519 | ENSG00000204519 | ZNF551 | 90233 |
| ENSG00000131470.14 | 392.8542573 | -1.178635918 | 0.186242455 | -3.1069 | 0.001891 | 0.013121049 | ENSG00000131470 | PSMC3IP | 29893 |
| ENSG00000256667.6 | 141.3164219 | -1.178843195 | 0.190768678 | -3.03427 | 0.002411 | 0.016366933 | ENSG00000256667 | NA | NA |
| ENSG00000110104.13 | 1739.150787 | -1.179675158 | 0.171978933 | -3.37062 | 0.00075 | 0.005727623 | ENSG00000110104 | CCDC86 | 79080 |
| ENSG00000171130.18 | 214.7235086 | -1.179747209 | 0.096556142 | -6.00425 | 1.92E-09 | 3.20E-08 | ENSG00000171130 | ATP6V0E2 | 155066 |
| ENSG00000140525.17 | 7646.81255 | -1.180842473 | 0.12200755 | -4.76071 | 1.93E-06 | 2.24E-05 | ENSG00000140525 | FANCI | 55215 |
| ENSG00000102743.15 | 900.3385332 | -1.181181144 | 0.127351085 | -4.56361 | 5.03E-06 | 5.54E-05 | ENSG00000102743 | SLC25A15 | 10166 |
| ENSG00000170779.10 | 2528.22425 | -1.181489358 | 0.161082002 | -3.6099 | 0.000306 | 0.002523153 | ENSG00000170779 | CDCA4 | 55038 |
| ENSG00000138092.11 | 766.5234842 | -1.181885633 | 0.186534445 | -3.11945 | 0.001812 | 0.012640323 | ENSG00000138092 | CENPO | 79172 |
| ENSG00000174442.12 | 2121.17221 | -1.182140031 | 0.147223123 | -3.95413 | 7.68E-05 | 0.000707331 | ENSG00000174442 | ZWILCH | 55055 |
| ENSG00000133401.16 | 320.2154019 | -1.182718459 | 0.140992674 | -4.13297 | 3.58E-05 | 0.000347304 | ENSG00000133401 | PDZD2 | 23037 |
| ENSG00000151725.12 | 1651.098025 | -1.182951842 | 0.115793007 | -5.03443 | 4.79E-07 | 6.04E-06 | ENSG00000151725 | CENPU | 79682 |
| ENSG00000170525.21 | 2617.720874 | -1.182977625 | 0.041212965 | -14.1455 | 1.99E-45 | 2.43E-43 | ENSG00000170525 | PFKFB3 | 5209 |
| ENSG00000168394.11 | 2675.675349 | -1.183797667 | 0.069522334 | -8.39727 | 4.57E-17 | 1.44E-15 | ENSG00000168394 | TAP1 | 6890 |
| ENSG00000188158.15 | 804.0037 | -1.183856 | 0.09103257 | -6.41371 | 1.42E-10 | 2.68E-09 | ENSG00000188158 | NHS | 4810 |
| ENSG00000151576.10 | 1871.827313 | -1.184524948 | 0.143298094 | -4.07908 | 4.52E-05 | 0.000431579 | ENSG00000151576 | QTRT2 | 79691 |
| ENSG00000145990.11 | 621.5568557 | -1.185261828 | 0.08137482 | -7.19217 | 6.38E-13 | 1.49E-11 | ENSG00000145990 | GFOD1 | 54438 |
| ENSG00000254726.3 | 34.33898502 | -1.186161991 | 0.21026819 | -2.78769 | 0.005309 | 0.032973599 | ENSG00000254726 | MEX3A | 92312 |
| ENSG00000119326.15 | 3500.209477 | -1.186216509 | 0.198867207 | -2.94778 | 0.003201 | 0.021064757 | ENSG00000119326 | CTNNAL1 | 8727 |
| ENSG00000072571.20 | 3022.179402 | -1.18840273 | 0.121258237 | -4.85248 | 1.22E-06 | 1.45E-05 | ENSG00000072571 | HMMR | 3161 |
| ENSG00000070718.12 | 998.1938748 | -1.190132736 | 0.12022786 | -4.90845 | 9.18E-07 | 1.12E-05 | ENSG00000070718 | AP3M2 | 10947 |
| ENSG00000235499.1 | 42.47736473 | -1.190190835 | 0.220177157 | -2.68053 | 0.007351 | 0.043915734 | ENSG00000235499 | NA | NA |
| ENSG00000050165.17 | 1197.774881 | -1.190427873 | 0.099659191 | -5.92447 | 3.13E-09 | 5.11E-08 | ENSG00000050165 | DKK3 | 27122 |
| ENSG00000125885.13 | 703.9737 | -1.190739 | 0.1635885 | -3.61113 | 0.000305 | 0.002513555 | ENSG00000125885 | MCM8 | 84515 |
| ENSG00000113739.10 | 1816.72713 | -1.190949645 | 0.169231089 | -3.49197 | 0.000479 | 0.003808047 | ENSG00000113739 | STC2 | 8614 |
| ENSG00000100105.18 | 237.2496 | -1.191301 | 0.1323577 | -4.46745 | 7.92E-06 | 8.53E-05 | ENSG00000100105 | PATZ1 | 23598 |

| | | | | | | | | | |
|---|---|---|---|---|---|---|---|---|---|
| ENSG00000162493.16 | 1523.789123 | -1.193832702 | 0.052842488 | -11.2378 | 2.66E-29 | 1.76E-27 | ENSG00000162493 | PDPN | 10630 |
| ENSG00000073849.15 | 374.4753394 | -1.193909328 | 0.208997307 | -2.84171 | 0.004487 | 0.028442698 | ENSG00000073849 | ST6GAL1 | 6480 |
| ENSG00000169105.7 | 531.3298668 | -1.193961557 | 0.151527563 | -3.91983 | 8.86E-05 | 0.000806931 | ENSG00000169105 | CHST14 | 113189 |
| ENSG00000134461.16 | 210.6757566 | -1.195399853 | 0.128409274 | -4.63674 | 3.54E-06 | 3.97E-05 | ENSG00000134461 | ANKRD16 | 54522 |
| ENSG00000167775.11 | 773.5975603 | -1.198296149 | 0.202508033 | -2.95443 | 0.003132 | 0.020641638 | ENSG00000167775 | CD320 | 51293 |
| ENSG00000116649.10 | 2714.19179 | -1.198619057 | 0.168886256 | -3.54451 | 0.000393 | 0.003178023 | ENSG00000116649 | SRM | 6723 |
| ENSG00000229807.12 | 7983.648 | -1.198799 | 0.108181 | -5.53516 | 3.11E-08 | 4.54E-07 | ENSG00000229807 | XIST | 7503 |
| ENSG00000120539.14 | 1700.150046 | -1.198925325 | 0.112326448 | -5.33201 | 9.71E-08 | 1.34E-06 | ENSG00000120539 | MASTL | 84930 |
| ENSG00000185551.15 | 1165.736551 | -1.199729509 | 0.085332518 | -7.02815 | 2.09E-12 | 4.66E-11 | ENSG00000185551 | NR2F2 | 7026 |
| ENSG00000120253.14 | 2474.317928 | -1.201429128 | 0.102266017 | -5.88103 | 4.08E-09 | 6.56E-08 | ENSG00000120253 | NUP43 | 348995 |
| ENSG00000226950.7 | 737.9684181 | -1.203788076 | 0.145280481 | -4.15602 | 3.24E-05 | 0.000315756 | ENSG00000226950 | DANCR | 57291 |
| ENSG00000175322.11 | 213.5310172 | -1.203979997 | 0.179087765 | -3.37254 | 0.000745 | 0.005691122 | ENSG00000175322 | ZNF519 | 162655 |
| ENSG00000166002.7 | 92.58600904 | -1.204375469 | 0.153394188 | -3.94002 | 8.15E-05 | 0.000746923 | ENSG00000166002 | SMCO4 | 56935 |
| ENSG00000055332.18 | 5026.078313 | -1.204576322 | 0.057351677 | -10.5416 | 5.56E-26 | 2.97E-24 | ENSG00000055332 | EIF2AK2 | 5610 |
| ENSG00000173193.15 | 4930.272677 | -1.206813485 | 0.083459424 | -7.27076 | 3.57E-13 | 8.50E-12 | ENSG00000173193 | PARP14 | 54625 |
| ENSG00000080839.11 | 1103.524 | -1.208111 | 0.1197462 | -5.07833 | 3.81E-07 | 4.86E-06 | ENSG00000080839 | RBL1 | 5933 |
| ENSG00000157111.13 | 416.9128652 | -1.208540808 | 0.105102719 | -5.78996 | 7.04E-09 | 1.11E-07 | ENSG00000157111 | TMEM171 | 134285 |
| ENSG00000164611.13 | 4923.857623 | -1.209015617 | 0.149013047 | -4.087 | 4.37E-05 | 0.000418178 | ENSG00000164611 | PTTG1 | 9232 |
| ENSG00000279348.1 | 192.3641316 | -1.209410687 | 0.156719279 | -3.88855 | 0.000101 | 0.000910067 | ENSG00000279348 | NA | NA |
| ENSG00000121454.6 | 110.2188382 | -1.211118287 | 0.195565359 | -3.12488 | 0.001779 | 0.012451899 | ENSG00000121454 | LHX4 | 89884 |
| ENSG00000156787.17 | 617.4772974 | -1.211971876 | 0.118506361 | -5.16404 | 2.42E-07 | 3.17E-06 | ENSG00000156787 | TBC1D31 | 93594 |
| ENSG00000156802.13 | 6788.927867 | -1.213243204 | 0.074995642 | -8.17705 | 2.91E-16 | 8.66E-15 | ENSG00000156802 | ATAD2 | 29028 |
| ENSG00000214223.4 | 48.7498717 | -1.213623398 | 0.229092262 | -2.6785 | 0.007395 | 0.044152846 | ENSG00000214223 | HNRNPA1 | 664709 |
| ENSG00000257093.7 | 216.2461844 | -1.213707932 | 0.116951754 | -5.24753 | 1.54E-07 | 2.08E-06 | ENSG00000257093 | DENND11 | 57189 |
| ENSG00000119285.11 | 4135.440672 | -1.213928342 | 0.160123606 | -3.83409 | 0.000126 | 0.0011169 | ENSG00000119285 | HEATR1 | 55127 |
| ENSG00000171467.16 | 2723.455081 | -1.215492426 | 0.099324004 | -6.19681 | 5.76E-10 | 1.02E-08 | ENSG00000171467 | ZNF318 | 24149 |
| ENSG00000112759.18 | 1900.354926 | -1.218444057 | 0.119465029 | -5.17678 | 2.26E-07 | 2.97E-06 | ENSG00000112759 | SLC29A1 | 2030 |
| ENSG00000132382.14 | 3601.124451 | -1.219086167 | 0.132555743 | -4.67038 | 3.01E-06 | 3.40E-05 | ENSG00000132382 | MYBBP1A | 10514 |
| ENSG00000184445.12 | 2775.627339 | -1.219346264 | 0.12774417 | -4.84833 | 1.25E-06 | 1.48E-05 | ENSG00000184445 | KNTC1 | 9735 |
| ENSG00000213366.13 | 36.84843018 | -1.219567526 | 0.212428552 | -2.91659 | 0.003539 | 0.022990334 | ENSG00000213366 | GSTM2 | 2946 |
| ENSG00000112312.10 | 1412.422236 | -1.220067993 | 0.126412767 | -4.90511 | 9.34E-07 | 1.13E-05 | ENSG00000112312 | GMNN | 51053 |
| ENSG00000149573.9 | 5253.366079 | -1.220241809 | 0.181345537 | -3.42022 | 0.000626 | 0.004852159 | ENSG00000149573 | MPZL2 | 10205 |
| ENSG00000128011.4 | 47.48789 | -1.220629 | 0.2307693 | -2.68939 | 0.007158 | 0.0428526 | ENSG00000128011 | LRFN1 | 57622 |
| ENSG00000169684.13 | 146.9216705 | -1.220791982 | 0.180337569 | -3.44239 | 0.000577 | 0.004505513 | ENSG00000169684 | CHRNA5 | 1138 |
| ENSG00000076351.13 | 97.46100495 | -1.223314741 | 0.131279958 | -4.74798 | 2.05E-06 | 2.37E-05 | ENSG00000076351 | SLC46A1 | 113235 |
| ENSG00000149136.9 | 10967.38677 | -1.22336994 | 0.152269449 | -4.09386 | 4.24E-05 | 0.000406418 | ENSG00000149136 | SSRP1 | 6749 |
| ENSG00000170689.10 | 98.91681749 | -1.225664522 | 0.203207697 | -3.07894 | 0.002077 | 0.014286735 | ENSG00000170689 | HOXB9 | 3219 |
| ENSG00000169855.20 | 1148.252669 | -1.225723786 | 0.0993743 | -6.29664 | 3.04E-10 | 5.58E-09 | ENSG00000169855 | ROBO1 | 6091 |
| ENSG00000115526.11 | 456.9744645 | -1.225863788 | 0.210854146 | -2.96823 | 0.002995 | 0.019864976 | ENSG00000115526 | CHST10 | 9486 |
| ENSG00000141384.12 | 738.3725999 | -1.226922473 | 0.174904264 | -3.58438 | 0.000338 | 0.002764355 | ENSG00000141384 | TAF4B | 6875 |
| ENSG00000183840.7 | 228.9257191 | -1.227648893 | 0.22409148 | -2.80086 | 0.005097 | 0.031790837 | ENSG00000183840 | GPR39 | 2863 |
| ENSG00000179841.8 | 36.17612343 | -1.228427956 | 0.231336223 | -2.71651 | 0.006597 | 0.039926053 | ENSG00000179841 | AKAP5 | 9495 |

| | | | | | | | | | |
|---|---|---|---|---|---|---|---|---|---|
| ENSG00000105202.9 | 9444.89 | -1.22949 | 0.1444493 | -4.35786 | 1.31E-05 | 0.000136315 | ENSG00000105202 | FBL | 2091 |
| ENSG00000232445.2 | 306.8771614 | -1.2296496 | 0.188662197 | -3.33744 | 0.000846 | 0.006383891 | ENSG00000232445 | NA | NA |
| ENSG00000048991.16 | 2041.139537 | -1.230379447 | 0.175636573 | -3.58911 | 0.000332 | 0.002717946 | ENSG00000048991 | R3HDM1 | 23518 |
| ENSG00000167994.13 | 67.16792435 | -1.230964371 | 0.213172291 | -2.95988 | 0.003078 | 0.020340518 | ENSG00000167994 | RAB3IL1 | 5866 |
| ENSG00000162775.16 | 1229.514258 | -1.231537704 | 0.085566457 | -7.38067 | 1.57E-13 | 3.86E-12 | ENSG00000162775 | RBM15 | 64783 |
| ENSG00000144395.18 | 369.294399 | -1.232446245 | 0.145014013 | -4.36128 | 1.29E-05 | 0.000134468 | ENSG00000144395 | CCDC150 | 284992 |
| ENSG00000153107.13 | 3433.437479 | -1.233123787 | 0.154642082 | -4.09412 | 4.24E-05 | 0.00040625 | ENSG00000153107 | ANAPC1 | 64682 |
| ENSG00000180071.20 | 212.389015 | -1.234352302 | 0.165022929 | -3.84403 | 0.000121 | 0.001076195 | ENSG00000180071 | ANKRD18/ | 253650 |
| ENSG00000172244.9 | 201.6937787 | -1.23544121 | 0.15441512 | -4.11515 | 3.87E-05 | 0.000373896 | ENSG00000172244 | C5orf34 | 375444 |
| ENSG00000101842.13 | 67.85885 | -1.235995 | 0.1686498 | -3.7711 | 0.000163 | 0.001411703 | ENSG00000101842 | VSIG1 | 340547 |
| ENSG00000120868.13 | 1122.418536 | -1.237096071 | 0.130643809 | -4.87659 | 1.08E-06 | 1.30E-05 | ENSG00000120868 | APAF1 | 317 |
| ENSG00000168918.14 | 995.0358667 | -1.237225058 | 0.183564268 | -3.4714 | 0.000518 | 0.004086445 | ENSG00000168918 | INPP5D | 3635 |
| ENSG00000163507.14 | 2377.65665 | -1.237329395 | 0.106441277 | -5.98762 | 2.13E-09 | 3.53E-08 | ENSG00000163507 | CIP2A | 57650 |
| ENSG00000214022.11 | 1321.172109 | -1.238225048 | 0.093389719 | -6.834 | 8.26E-12 | 1.75E-10 | ENSG00000214022 | REPIN1 | 29803 |
| ENSG00000240065.8 | 1393.76011 | -1.238860865 | 0.08688059 | -7.35332 | 1.93E-13 | 4.72E-12 | ENSG00000240065 | PSMB9 | 5698 |
| ENSG00000065150.20 | 14639.03947 | -1.238967248 | 0.150339563 | -4.25016 | 2.14E-05 | 0.000214116 | ENSG00000065150 | IPO5 | 3843 |
| ENSG00000272711.1 | 95.12799462 | -1.24076167 | 0.189803154 | -3.37593 | 0.000736 | 0.005631137 | ENSG00000272711 | NA | NA |
| ENSG00000146858.8 | 320.5211402 | -1.243430553 | 0.16728956 | -3.84621 | 0.00012 | 0.001067725 | ENSG00000146858 | ZC3HAV1L | 92092 |
| ENSG00000183579.15 | 621.2756 | -1.243499 | 0.1147059 | -5.60999 | 2.02E-08 | 3.03E-07 | ENSG00000183579 | ZNRF3 | 84133 |
| ENSG00000130816.16 | 7007.966676 | -1.243663337 | 0.135667641 | -4.74441 | 2.09E-06 | 2.41E-05 | ENSG00000130816 | DNMT1 | 1786 |
| ENSG00000179041.4 | 1388.653609 | -1.244287003 | 0.163039817 | -3.95172 | 7.76E-05 | 0.000713533 | ENSG00000179041 | RRS1 | 23212 |
| ENSG00000164294.14 | 1915.018276 | -1.245171474 | 0.198168029 | -3.25568 | 0.001131 | 0.008291639 | ENSG00000164294 | GPX8 | 493869 |
| ENSG00000136895.19 | 66.80684004 | -1.245272097 | 0.20219276 | -3.19137 | 0.001416 | 0.010130766 | ENSG00000136895 | GARNL3 | 84253 |
| ENSG00000212719.12 | 878.8359336 | -1.247453256 | 0.155502005 | -4.16363 | 3.13E-05 | 0.00030623 | ENSG00000212719 | LINC0269: | 339263 |
| ENSG00000171877.21 | 136.0844431 | -1.249404956 | 0.179328027 | -3.62132 | 0.000293 | 0.002426267 | ENSG00000171877 | FRMD5 | 84978 |
| ENSG00000040275.17 | 3097.749553 | -1.249458708 | 0.163029388 | -3.98369 | 6.79E-05 | 0.000629183 | ENSG00000040275 | SPDL1 | 54908 |
| ENSG00000033170.16 | 514.6550244 | -1.249828785 | 0.181229199 | -3.58567 | 0.000336 | 0.002752324 | ENSG00000033170 | FUT8 | 2530 |
| ENSG00000072274.13 | 12728.27929 | -1.252734854 | 0.186380023 | -3.50217 | 0.000461 | 0.003676114 | ENSG00000072274 | TFRC | 7037 |
| ENSG00000151835.16 | 5470.112496 | -1.25318488 | 0.214942951 | -3.03888 | 0.002375 | 0.016135115 | ENSG00000151835 | SACS | 26278 |
| ENSG00000072501.17 | 10382.15 | -1.253604 | 0.1375406 | -4.75208 | 2.01E-06 | 2.33E-05 | ENSG00000072501 | SMC1A | 8243 |
| ENSG00000140451.13 | 612.2590596 | -1.255016834 | 0.193763171 | -3.3805 | 0.000724 | 0.005547795 | ENSG00000140451 | PIF1 | 80119 |
| ENSG00000287431.1 | 30.96579543 | -1.255798498 | 0.241780157 | -2.71238 | 0.00668 | 0.040350877 | ENSG00000287431 | NA | NA |
| ENSG00000174370.10 | 148.3450698 | -1.256236312 | 0.137137815 | -4.78523 | 1.71E-06 | 2.00E-05 | ENSG00000174370 | C11orf45 | 219833 |
| ENSG00000072135.13 | 367.3290388 | -1.257880027 | 0.107205341 | -6.13663 | 8.43E-10 | 1.46E-08 | ENSG00000072135 | PTPN18 | 26469 |
| ENSG00000131351.15 | 557.4252 | -1.258683 | 0.1525482 | -4.31787 | 1.58E-05 | 0.000161127 | ENSG00000131351 | HAUS8 | 93323 |
| ENSG00000077044.10 | 784.5836335 | -1.258777304 | 0.135554871 | -4.85986 | 1.17E-06 | 1.41E-05 | ENSG00000077044 | DGKD | 8527 |
| ENSG00000234602.7 | 128.5497263 | -1.258921871 | 0.168396768 | -3.91291 | 9.12E-05 | 0.000828548 | ENSG00000234602 | MCIDAS | 345643 |
| ENSG00000198056.14 | 596.7288305 | -1.259031039 | 0.146014451 | -4.51346 | 6.38E-06 | 6.96E-05 | ENSG00000198056 | PRIM1 | 5557 |
| ENSG00000117266.15 | 134.9409938 | -1.261237721 | 0.122776374 | -5.38571 | 7.22E-08 | 1.01E-06 | ENSG00000117266 | CDK18 | 5129 |
| ENSG00000172292.14 | 1078.705797 | -1.262199862 | 0.129129348 | -5.12819 | 2.93E-07 | 3.79E-06 | ENSG00000172292 | CERS6 | 253782 |
| ENSG00000123473.15 | 1867.080827 | -1.262282756 | 0.115292174 | -5.74439 | 9.23E-09 | 1.43E-07 | ENSG00000123473 | STIL | 6491 |
| ENSG00000141448.10 | 168.0636837 | -1.263100658 | 0.134221399 | -4.94035 | 7.80E-07 | 9.59E-06 | ENSG00000141448 | GATA6 | 2627 |

| | | | | | | | | | | |
|---|---|---|---|---|---|---|---|---|---|---|
| ENSG00000173898.13 | 877.7162401 | -1.263698942 | 0.128236981 | -5.17557 | 2.27E-07 | 2.99E-06 | ENSG00000173898 | SPTBN2 | | 6712 |
| ENSG00000011332.19 | 133.3948 | -1.264028 | 0.1797628 | -3.69391 | 0.000221 | 0.001868711 | ENSG00000011332 | DPF1 | | 8193 |
| ENSG00000168398.6 | 1093.770194 | -1.264118689 | 0.064350654 | -10.3203 | 5.70E-25 | 2.89E-23 | ENSG00000168398 | BDKRB2 | | 624 |
| ENSG00000285728.2 | 110.1623205 | -1.264929725 | 0.162075993 | -4.10258 | 4.09E-05 | 0.000393094 | ENSG00000285728 | NA | NA | |
| ENSG00000100311.16 | 541.1731 | -1.26649 | 0.1995414 | -3.34011 | 0.000837 | 0.006331992 | ENSG00000100311 | PDGFB | | 5155 |
| ENSG00000059588.10 | 1308.060417 | -1.266687813 | 0.113159215 | -5.89159 | 3.82E-09 | 6.18E-08 | ENSG00000059588 | TARBP1 | | 6894 |
| ENSG00000138035.15 | 2587.78099 | -1.267060424 | 0.145860006 | -4.57329 | 4.80E-06 | 5.31E-05 | ENSG00000138035 | PNPT1 | | 87178 |
| ENSG00000164778.4 | 429.3419931 | -1.267481913 | 0.224832965 | -2.96879 | 0.00299 | 0.019838739 | ENSG00000164778 | EN2 | | 2020 |
| ENSG00000075886.11 | 23.82849254 | -1.268052821 | 0.238563284 | -2.80032 | 0.005105 | 0.031836921 | ENSG00000075886 | TUBA3D | | 113457 |
| ENSG00000239900.12 | 218.8395 | -1.269534 | 0.1651546 | -4.05399 | 5.04E-05 | 0.000475852 | ENSG00000239900 | ADSL | | 158 |
| ENSG00000083807.10 | 127.6698 | -1.269537 | 0.1795458 | -3.72906 | 0.000192 | 0.001643092 | ENSG00000083807 | SLC27A5 | | 10998 |
| ENSG00000180626.10 | 210.9527698 | -1.269559793 | 0.109543761 | -6.11226 | 9.82E-10 | 1.69E-08 | ENSG00000180626 | ZNF594 | | 84622 |
| ENSG00000104147.9 | 131.5249057 | -1.270350725 | 0.167434336 | -4.00366 | 6.24E-05 | 0.000581369 | ENSG00000104147 | OIP5 | | 11339 |
| ENSG00000227848.1 | 53.41026054 | -1.270433297 | 0.254052518 | -2.63896 | 0.008316 | 0.048829639 | ENSG00000227848 | NA | NA | |
| ENSG00000124126.14 | 48.24589 | -1.270704 | 0.2013197 | -3.33154 | 0.000864 | 0.006500651 | ENSG00000124126 | PREX1 | | 57580 |
| ENSG00000087586.17 | 3093.312 | -1.270808 | 0.1454705 | -4.6113 | 4.00E-06 | 4.46E-05 | ENSG00000087586 | AURKA | | 6790 |
| ENSG00000267374.2 | 156.8323569 | -1.271811701 | 0.209628583 | -3.20477 | 0.001352 | 0.009712522 | ENSG00000267374 | MIR924HG | | 647946 |
| ENSG00000285517.1 | 751.530876 | -1.272233486 | 0.162082837 | -4.14747 | 3.36E-05 | 0.000326984 | ENSG00000285517 | NA | NA | |
| ENSG00000079616.12 | 2602.573284 | -1.272482441 | 0.106148123 | -6.33532 | 2.37E-10 | 4.38E-09 | ENSG00000079616 | KIF22 | | 3835 |
| ENSG00000107815.10 | 1190.279535 | -1.274163144 | 0.171508834 | -3.93078 | 8.47E-05 | 0.000773815 | ENSG00000107815 | TWNK | | 56652 |
| ENSG00000184307.15 | 876.0359758 | -1.276016792 | 0.062404603 | -10.8328 | 2.41E-27 | 1.40E-25 | ENSG00000184307 | ZDHHC23 | | 254887 |
| ENSG00000180385.8 | 140.9254476 | -1.278052868 | 0.177816369 | -3.81322 | 0.000137 | 0.001206771 | ENSG00000180385 | NA | NA | |
| ENSG00000136811.16 | 1631.908814 | -1.27901776 | 0.147719283 | -4.59668 | 4.29E-06 | 4.77E-05 | ENSG00000136811 | ODF2 | | 4957 |
| ENSG00000144381.17 | 35557.74689 | -1.279468052 | 0.170447556 | -3.98638 | 6.71E-05 | 0.000622977 | ENSG00000144381 | HSPD1 | | 3329 |
| ENSG00000064309.14 | 115.7224084 | -1.281213104 | 0.127748346 | -5.33246 | 9.69E-08 | 1.34E-06 | ENSG00000064309 | CDON | | 50937 |
| ENSG00000068654.16 | 3496.483126 | -1.281322261 | 0.151581526 | -4.49476 | 6.96E-06 | 7.57E-05 | ENSG00000068654 | POLR1A | | 25885 |
| ENSG00000120802.13 | 6921.166466 | -1.282468561 | 0.111684436 | -6.11069 | 9.92E-10 | 1.71E-08 | ENSG00000120802 | TMPO | | 7112 |
| ENSG00000174136.12 | 611.7258572 | -1.282498664 | 0.128187843 | -5.32421 | 1.01E-07 | 1.40E-06 | ENSG00000174136 | RGMB | | 285704 |
| ENSG00000279500.1 | 31.79678013 | -1.283094798 | 0.258874304 | -2.63871 | 0.008322 | 0.048843144 | ENSG00000279500 | NA | NA | |
| ENSG00000166123.14 | 726.2048709 | -1.284078097 | 0.10969902 | -6.23595 | 4.49E-10 | 8.08E-09 | ENSG00000166123 | GPT2 | | 84706 |
| ENSG00000186197.14 | 127.7146249 | -1.284279746 | 0.142292775 | -4.80896 | 1.52E-06 | 1.78E-05 | ENSG00000186197 | EDARADD | | 128178 |
| ENSG00000188312.14 | 194.3654563 | -1.284932076 | 0.111331528 | -6.15218 | 7.64E-10 | 1.33E-08 | ENSG00000188312 | CENPP | | 401541 |
| ENSG00000103319.12 | 350.8081997 | -1.285285537 | 0.157438562 | -4.35272 | 1.34E-05 | 0.000139284 | ENSG00000103319 | EEF2K | | 29904 |
| ENSG00000136122.16 | 435.4550464 | -1.285583515 | 0.124059842 | -5.52623 | 3.27E-08 | 4.76E-07 | ENSG00000136122 | BORA | | 79866 |
| ENSG00000076003.5 | 3194.997049 | -1.285923763 | 0.167222479 | -4.10186 | 4.10E-05 | 0.000393887 | ENSG00000076003 | MCM6 | | 4175 |
| ENSG00000051825.14 | 1000.195543 | -1.286827321 | 0.170463769 | -4.02917 | 5.60E-05 | 0.000525812 | ENSG00000051825 | MPHOSPH | | 10198 |
| ENSG00000214826.5 | 294.2453988 | -1.288169301 | 0.186642602 | -3.6871 | 0.000227 | 0.001915798 | ENSG00000214826 | NA | NA | |
| ENSG00000135446.17 | 2449.942085 | -1.291000146 | 0.164822711 | -4.19238 | 2.76E-05 | 0.000271871 | ENSG00000135446 | CDK4 | | 1019 |
| ENSG00000115053.16 | 52674.8746 | -1.291049969 | 0.166176056 | -4.15854 | 3.20E-05 | 0.000312561 | ENSG00000115053 | NCL | | 4691 |
| ENSG00000269609.6 | 180.978267 | -1.29231485 | 0.124888268 | -5.54347 | 2.97E-08 | 4.35E-07 | ENSG00000269609 | RPARP-AS | | 1.01E+08 |
| ENSG00000134285.11 | 449.2366515 | -1.292424928 | 0.153928686 | -4.49835 | 6.85E-06 | 7.45E-05 | ENSG00000134285 | FKBP11 | | 51303 |
| ENSG00000103540.16 | 1276.627713 | -1.296275734 | 0.1218806 | -5.71277 | 1.11E-08 | 1.71E-07 | ENSG00000103540 | CCP110 | | 9738 |

| | | | | | | | | | |
|---|---|---|---|---|---|---|---|---|---|
| ENSG00000058056.9 | 720.270198 | -1.297049727 | 0.223845809 | -3.11397 | 0.001846 | 0.012853948 | ENSG00000058056 | USP13 | 8975 |
| ENSG00000101220.17 | 1013.374 | -1.297433 | 0.182809 | -3.81509 | 0.000136 | 0.001199234 | ENSG00000101220 | C20orf27 | 54976 |
| ENSG00000108679.13 | 7526.652635 | -1.297784526 | 0.12534507 | -5.56691 | 2.59E-08 | 3.83E-07 | ENSG00000108679 | LGALS3BP | 3959 |
| ENSG00000110446.11 | 637.9472336 | -1.298219526 | 0.084836561 | -8.23017 | 1.87E-16 | 5.66E-15 | ENSG00000110446 | SLC15A3 | 51296 |
| ENSG00000111445.14 | 1267.785862 | -1.298689596 | 0.170437897 | -4.09938 | 4.14E-05 | 0.000397564 | ENSG00000111445 | RFC5 | 5985 |
| ENSG00000125319.14 | 325.3895386 | -1.300038059 | 0.171342175 | -4.08561 | 4.40E-05 | 0.000420522 | ENSG00000125319 | HROB | 78995 |
| ENSG00000162139.10 | 788.7199302 | -1.300217581 | 0.12353475 | -5.66818 | 1.44E-08 | 2.19E-07 | ENSG00000162139 | NEU3 | 10825 |
| ENSG00000101945.16 | 502.617 | -1.300232 | 0.145051 | -4.82749 | 1.38E-06 | 1.63E-05 | ENSG00000101945 | SUV39H1 | 6839 |
| ENSG00000285417.1 | 33.55667929 | -1.301275823 | 0.239793674 | -2.9245 | 0.00345 | 0.022495601 | ENSG00000285417 | NA | NA |
| ENSG00000169607.13 | 2246.460284 | -1.302481925 | 0.16452394 | -4.26979 | 1.96E-05 | 0.000197008 | ENSG00000169607 | CKAP2L | 150468 |
| ENSG00000171793.16 | 4998.986793 | -1.303267184 | 0.173539388 | -4.05249 | 5.07E-05 | 0.00047856 | ENSG00000171793 | CTPS1 | 1503 |
| ENSG00000184557.4 | 446.0959125 | -1.303948378 | 0.255371171 | -2.75657 | 0.005841 | 0.035863076 | ENSG00000184557 | SOCS3 | 9021 |
| ENSG00000165138.18 | 1400.490828 | -1.304004976 | 0.120411393 | -5.84666 | 5.02E-09 | 8.00E-08 | ENSG00000165138 | ANKS6 | 203286 |
| ENSG00000235098.8 | 81.09557183 | -1.30486198 | 0.15909515 | -4.43044 | 9.40E-06 | 0.000100181 | ENSG00000235098 | ANKRD65 | 441869 |
| ENSG00000139133.7 | 296.5574911 | -1.304937339 | 0.155422691 | -4.53561 | 5.74E-06 | 6.30E-05 | ENSG00000139133 | ALG10 | 84920 |
| ENSG00000166582.10 | 284.0742567 | -1.305100027 | 0.203272868 | -3.46874 | 0.000523 | 0.004124729 | ENSG00000166582 | CENPV | 201161 |
| ENSG00000279191.1 | 54.82290879 | -1.305415999 | 0.228271268 | -3.09025 | 0.002 | 0.013789243 | ENSG00000279191 | NA | NA |
| ENSG00000174804.4 | 93.54711455 | -1.305715615 | 0.192680302 | -3.66262 | 0.00025 | 0.002093897 | ENSG00000174804 | FZD4 | 8322 |
| ENSG00000214711.10 | 52.66834286 | -1.306944562 | 0.26381277 | -2.67972 | 0.007368 | 0.044011771 | ENSG00000214711 | CAPN14 | 440854 |
| ENSG00000171462.15 | 122.6913308 | -1.309835986 | 0.165815072 | -4.28089 | 1.86E-05 | 0.000188212 | ENSG00000171462 | DLK2 | 65989 |
| ENSG00000187908.19 | 154.0104665 | -1.313505835 | 0.179584116 | -3.9731 | 7.09E-05 | 0.000655551 | ENSG00000187908 | DMBT1 | 1755 |
| ENSG00000196418.12 | 163.8950612 | -1.314843094 | 0.178028947 | -4.01532 | 5.94E-05 | 0.000555119 | ENSG00000196418 | ZNF124 | 7678 |
| ENSG00000148019.13 | 2168.270975 | -1.316390896 | 0.112658586 | -6.35896 | 2.03E-10 | 3.78E-09 | ENSG00000148019 | CEP78 | 84131 |
| ENSG00000157657.14 | 1242.17152 | -1.316580884 | 0.097492223 | -7.35013 | 1.98E-13 | 4.83E-12 | ENSG00000157657 | ZNF618 | 114991 |
| ENSG00000128050.8 | 12755.02688 | -1.317110889 | 0.165315832 | -4.33782 | 1.44E-05 | 0.000148313 | ENSG00000128050 | PAICS | 10606 |
| ENSG00000187240.16 | 2094.563437 | -1.317369224 | 0.187703591 | -3.82182 | 0.000132 | 0.001170085 | ENSG00000187240 | DYNC2H1 | 79659 |
| ENSG00000112379.9 | 428.2812119 | -1.321705986 | 0.189790629 | -3.80264 | 0.000143 | 0.001254104 | ENSG00000112379 | ARFGEF3 | 57221 |
| ENSG00000165271.17 | 3787.877912 | -1.322462036 | 0.113016013 | -6.39256 | 1.63E-10 | 3.06E-09 | ENSG00000165271 | NOL6 | 65083 |
| ENSG00000166197.16 | 10960.04531 | -1.32284384 | 0.167833905 | -4.3069 | 1.66E-05 | 0.000168805 | ENSG00000166197 | NOLC1 | 9221 |
| ENSG00000129195.16 | 277.2740621 | -1.323139818 | 0.20064226 | -3.60413 | 0.000313 | 0.0025751 | ENSG00000129195 | PIMREG | 54478 |
| ENSG00000112742.10 | 2196.152808 | -1.323618716 | 0.127284806 | -5.68504 | 1.31E-08 | 1.99E-07 | ENSG00000112742 | TTK | 7272 |
| ENSG00000182957.16 | 66.9651548 | -1.324071308 | 0.203869725 | -3.55164 | 0.000383 | 0.003099765 | ENSG00000182957 | SPATA13 | 221178 |
| ENSG00000148400.12 | 6158.862031 | -1.324284801 | 0.096609127 | -7.49706 | 6.53E-14 | 1.64E-12 | ENSG00000148400 | NOTCH1 | 4851 |
| ENSG00000103356.15 | 1421.523018 | -1.324714196 | 0.18677152 | -3.88022 | 0.000104 | 0.000939902 | ENSG00000103356 | EARS2 | 124454 |
| ENSG00000171517.6 | 1120.294909 | -1.325027099 | 0.083515685 | -8.68133 | 3.91E-18 | 1.33E-16 | ENSG00000171517 | LPAR3 | 23566 |
| ENSG00000138658.15 | 829.9116152 | -1.325520796 | 0.149588297 | -4.85012 | 1.23E-06 | 1.47E-05 | ENSG00000138658 | ZGRF1 | 55345 |
| ENSG00000168754.15 | 130.6964856 | -1.325934104 | 0.203576039 | -3.56591 | 0.000363 | 0.002952884 | ENSG00000168754 | FAM178B | 51252 |
| ENSG00000051180.17 | 892.4443024 | -1.326278761 | 0.133407879 | -5.44405 | 5.21E-08 | 7.36E-07 | ENSG00000051180 | RAD51 | 5888 |
| ENSG00000165572.7 | 689.5757914 | -1.327026287 | 0.168015184 | -4.32715 | 1.51E-05 | 0.000155054 | ENSG00000165572 | KBTBD6 | 89890 |
| ENSG00000065183.16 | 2750.342296 | -1.32732493 | 0.190095641 | -3.8261 | 0.00013 | 0.001150698 | ENSG00000065183 | WDR3 | 10885 |
| ENSG00000092470.12 | 1020.140673 | -1.327344594 | 0.13407471 | -5.42492 | 5.80E-08 | 8.16E-07 | ENSG00000092470 | WDR76 | 79968 |
| ENSG00000146410.12 | 439.8045911 | -1.329216304 | 0.107124465 | -6.80719 | 9.95E-12 | 2.09E-10 | ENSG00000146410 | MTFR2 | 113115 |

| | | | | | | | | | |
|---|---|---|---|---|---|---|---|---|---|
| ENSG00000119866.21 | 175.5559301 | -1.332253653 | 0.163647752 | -4.47457 | 7.66E-06 | 8.27E-05 | ENSG00000119866 | BCL11A | 53335 |
| ENSG00000104738.18 | 11021.09313 | -1.332548614 | 0.134370846 | -5.45169 | 4.99E-08 | 7.07E-07 | ENSG00000104738 | MCM4 | 4173 |
| ENSG00000128973.13 | 1338.759118 | -1.335085357 | 0.138914903 | -5.29162 | 1.21E-07 | 1.66E-06 | ENSG00000128973 | CLN6 | 54982 |
| ENSG00000160183.16 | 36.99154 | -1.336352 | 0.2553956 | -2.88318 | 0.003937 | 0.02530319 | ENSG00000160183 | TMPRSS3 | 64699 |
| ENSG00000197275.14 | 145.656662 | -1.337612901 | 0.176557134 | -4.17776 | 2.94E-05 | 0.000288778 | ENSG00000197275 | RAD54B | 25788 |
| ENSG00000136159.4 | 2055.900254 | -1.337675875 | 0.099536567 | -7.4111 | 1.25E-13 | 3.10E-12 | ENSG00000136159 | NUDT15 | 55270 |
| ENSG00000272269.1 | 28.03080273 | -1.338244442 | 0.263506827 | -2.80161 | 0.005085 | 0.031731512 | ENSG00000272269 | NUP153-A | 1.05E+08 |
| ENSG00000085552.17 | 213.0249488 | -1.338320835 | 0.129918672 | -5.68295 | 1.32E-08 | 2.01E-07 | ENSG00000085552 | IGSF9 | 57549 |
| ENSG00000118507.17 | 219.7033709 | -1.339719576 | 0.094683388 | -7.81256 | 5.60E-15 | 1.53E-13 | ENSG00000118507 | AKAP7 | 9465 |
| ENSG00000160193.11 | 1029.978 | -1.342896 | 0.1671832 | -4.44361 | 8.85E-06 | 9.46E-05 | ENSG00000160193 | WDR4 | 10785 |
| ENSG00000105270.15 | 36.47547 | -1.344931 | 0.2716202 | -2.74255 | 0.006096 | 0.03719062 | ENSG00000105270 | CLIP3 | 25999 |
| ENSG00000178295.15 | 1754.828803 | -1.345966341 | 0.129262989 | -5.77092 | 7.88E-09 | 1.24E-07 | ENSG00000178295 | GEN1 | 348654 |
| ENSG00000285043.1 | 43.02791059 | -1.346009134 | 0.232934053 | -3.20266 | 0.001362 | 0.009773375 | ENSG00000285043 | LOC11269 | 1.13E+08 |
| ENSG00000126778.11 | 477.7176883 | -1.348486641 | 0.080142487 | -9.33945 | 9.68E-21 | 3.91E-19 | ENSG00000126778 | SIX1 | 6495 |
| ENSG00000168496.4 | 3191.664175 | -1.349408727 | 0.162575419 | -4.60961 | 4.03E-06 | 4.50E-05 | ENSG00000168496 | FEN1 | 2237 |
| ENSG00000241634.1 | 57.06210696 | -1.349764845 | 0.223877593 | -3.34899 | 0.000811 | 0.006151531 | ENSG00000241634 | RPL13P8 | 1E+08 |
| ENSG00000175063.17 | 2267.22 | -1.351314 | 0.1707775 | -4.39937 | 1.09E-05 | 0.000114102 | ENSG00000175063 | UBE2C | 11065 |
| ENSG00000101868.10 | 1847.532 | -1.3517 | 0.1560714 | -4.81639 | 1.46E-06 | 1.72E-05 | ENSG00000101868 | POLA1 | 5422 |
| ENSG00000137273.6 | 432.611098 | -1.352280751 | 0.111677165 | -6.73621 | 1.63E-11 | 3.34E-10 | ENSG00000137273 | FOXF2 | 2295 |
| ENSG00000142731.11 | 1687.768785 | -1.352740703 | 0.149365714 | -5.03958 | 4.67E-07 | 5.88E-06 | ENSG00000142731 | PLK4 | 10733 |
| ENSG00000162614.18 | 134.116684 | -1.353827943 | 0.137786713 | -5.47098 | 4.48E-08 | 6.38E-07 | ENSG00000162614 | NEXN | 91624 |
| ENSG00000121621.7 | 1189.326534 | -1.3547113 | 0.187675374 | -4.02137 | 5.79E-05 | 0.00054182 | ENSG00000121621 | KIF18A | 81930 |
| ENSG00000145934.16 | 5362.462515 | -1.35537099 | 0.110824222 | -6.81594 | 9.37E-12 | 1.98E-10 | ENSG00000145934 | TENM2 | 57451 |
| ENSG00000131187.9 | 452.8568764 | -1.356647008 | 0.130355329 | -5.8045 | 6.46E-09 | 1.02E-07 | ENSG00000131187 | F12 | 2161 |
| ENSG00000187741.15 | 1193.685409 | -1.356986083 | 0.132530997 | -5.71177 | 1.12E-08 | 1.72E-07 | ENSG00000187741 | FANCA | 2175 |
| ENSG00000029534.20 | 744.8349721 | -1.357754722 | 0.234705126 | -3.22854 | 0.001244 | 0.009020752 | ENSG00000029534 | ANK1 | 286 |
| ENSG00000127586.17 | 1097.729444 | -1.359937892 | 0.141326437 | -5.37718 | 7.57E-08 | 1.06E-06 | ENSG00000127586 | CHTF18 | 63922 |
| ENSG00000119922.10 | 1413.157762 | -1.359995336 | 0.243960862 | -3.11523 | 0.001838 | 0.012805776 | ENSG00000119922 | IFIT2 | 3433 |
| ENSG00000185480.11 | 647.0878491 | -1.360589803 | 0.165112422 | -4.6065 | 4.10E-06 | 4.56E-05 | ENSG00000185480 | PARPBP | 55010 |
| ENSG00000084774.14 | 4868.265403 | -1.361750264 | 0.153234346 | -4.97115 | 6.66E-07 | 8.25E-06 | ENSG00000084774 | CAD | 790 |
| ENSG00000156509.14 | 123.1461253 | -1.36325585 | 0.239377782 | -3.1885 | 0.00143 | 0.01022644 | ENSG00000156509 | FBXO43 | 286151 |
| ENSG00000164104.11 | 8369.140499 | -1.363811528 | 0.124520638 | -6.13402 | 8.57E-10 | 1.48E-08 | ENSG00000164104 | HMGB2 | 3148 |
| ENSG00000221963.6 | 2499.327 | -1.364082 | 0.04686069 | -16.3054 | 9.03E-60 | 1.73E-57 | ENSG00000221963 | APOL6 | 80830 |
| ENSG00000160949.17 | 1274.82499 | -1.364145483 | 0.12152936 | -6.28774 | 3.22E-10 | 5.89E-09 | ENSG00000160949 | TONSL | 4796 |
| ENSG00000134256.12 | 58.78490117 | -1.364993453 | 0.180725922 | -4.23289 | 2.31E-05 | 0.000229896 | ENSG00000134256 | CD101 | 9398 |
| ENSG00000131188.12 | 591.4539354 | -1.365298236 | 0.146154487 | -5.23623 | 1.64E-07 | 2.20E-06 | ENSG00000131188 | PRR7 | 80758 |
| ENSG00000174951.11 | 334.6369 | -1.36543 | 0.133289 | -5.74263 | 9.32E-09 | 1.45E-07 | ENSG00000174951 | FUT1 | 2523 |
| ENSG00000139618.15 | 1839.078504 | -1.368401187 | 0.158275656 | -4.85483 | 1.20E-06 | 1.44E-05 | ENSG00000139618 | BRCA2 | 675 |
| ENSG00000109255.11 | 107.8424626 | -1.36852535 | 0.213396809 | -3.60139 | 0.000317 | 0.002598326 | ENSG00000109255 | NMU | 10874 |
| ENSG00000134874.17 | 574.0656769 | -1.36888131 | 0.189825743 | -4.05046 | 5.11E-05 | 0.000482399 | ENSG00000134874 | DZIP1 | 22873 |
| ENSG00000120217.14 | 1217.994644 | -1.369294522 | 0.087240573 | -8.81808 | 1.16E-18 | 4.09E-17 | ENSG00000120217 | CD274 | 29126 |
| ENSG00000260924.2 | 43.53791 | -1.369435 | 0.2426106 | -3.17148 | 0.001517 | 0.01076404 | ENSG00000260924 | NA | NA |

| | | | | | | | | | |
|---|---|---|---|---|---|---|---|---|---|
| ENSG00000273162.1 | 28.59334892 | -1.369752271 | 0.234391922 | -3.28404 | 0.001023 | 0.00757156 | ENSG00000273162 | NA | NA |
| ENSG00000082516.9 | 4720.275907 | -1.371981993 | 0.182035245 | -4.24084 | 2.23E-05 | 0.000222369 | ENSG00000082516 | GEMIN5 | 25929 |
| ENSG00000116711.10 | 188.4384848 | -1.374097205 | 0.13884245 | -5.57536 | 2.47E-08 | 3.66E-07 | ENSG00000116711 | PLA2G4A | 5321 |
| ENSG00000126602.11 | 3692.62347 | -1.374264133 | 0.161939646 | -4.78119 | 1.74E-06 | 2.04E-05 | ENSG00000126602 | TRAP1 | 10131 |
| ENSG00000175305.17 | 328.7192658 | -1.376788484 | 0.135429809 | -5.73573 | 9.71E-09 | 1.50E-07 | ENSG00000175305 | CCNE2 | 9134 |
| ENSG00000137807.15 | 4782.423503 | -1.377751416 | 0.174584134 | -4.45488 | 8.39E-06 | 9.01E-05 | ENSG00000137807 | KIF23 | 9493 |
| ENSG00000097046.13 | 1337.696827 | -1.377969509 | 0.153791427 | -5.0586 | 4.22E-07 | 5.36E-06 | ENSG00000097046 | CDC7 | 8317 |
| ENSG00000076770.14 | 767.2704 | -1.378259 | 0.1472422 | -5.28557 | 1.25E-07 | 1.71E-06 | ENSG00000076770 | MBNL3 | 55796 |
| ENSG00000189337.17 | 756.1848442 | -1.382334922 | 0.079861899 | -9.7961 | 1.17E-22 | 5.17E-21 | ENSG00000189337 | KAZN | 23254 |
| ENSG00000119917.14 | 3128.456332 | -1.383370047 | 0.248844396 | -3.14803 | 0.001644 | 0.011573555 | ENSG00000119917 | IFIT3 | 3437 |
| ENSG00000261373.1 | 144.7500014 | -1.385403708 | 0.209199582 | -3.75433 | 0.000174 | 0.001500845 | ENSG00000261373 | VPS9D1-A | 1E+08 |
| ENSG00000198554.12 | 1867.251339 | -1.387219572 | 0.145407798 | -5.41387 | 6.17E-08 | 8.68E-07 | ENSG00000198554 | WDHD1 | 11169 |
| ENSG00000151692.15 | 96.82462718 | -1.387944248 | 0.166162565 | -4.74201 | 2.12E-06 | 2.44E-05 | ENSG00000151692 | RNF144A | 9781 |
| ENSG00000153904.20 | 821.8702571 | -1.388322843 | 0.085766601 | -9.19149 | 3.87E-20 | 1.49E-18 | ENSG00000153904 | DDAH1 | 23576 |
| ENSG00000100092.23 | 255.4161 | -1.388934 | 0.1033598 | -7.63289 | 2.30E-14 | 5.98E-13 | ENSG00000100092 | SH3BP1 | 23616 |
| ENSG00000188130.14 | 803.9836 | -1.389383 | 0.171759 | -4.59587 | 4.31E-06 | 4.78E-05 | ENSG00000188130 | MAPK12 | 6300 |
| ENSG00000156463.18 | 1370.504658 | -1.389417545 | 0.194554311 | -4.05757 | 4.96E-05 | 0.000469279 | ENSG00000156463 | SH3RF2 | 153769 |
| ENSG00000105852.11 | 37.57837463 | -1.390437765 | 0.221442249 | -3.5695 | 0.000358 | 0.002916314 | ENSG00000105852 | PON3 | 5446 |
| ENSG00000139832.5 | 200.6857668 | -1.390946995 | 0.126635706 | -6.24585 | 4.22E-10 | 7.60E-09 | ENSG00000139832 | RAB20 | 55647 |
| ENSG00000234290.2 | 69.23733036 | -1.393086712 | 0.183395068 | -4.32447 | 1.53E-05 | 0.000156797 | ENSG00000234290 | NA | NA |
| ENSG00000105486.14 | 1157.96 | -1.393393 | 0.1398527 | -5.67306 | 1.40E-08 | 2.13E-07 | ENSG00000105486 | LIG1 | 3978 |
| ENSG00000147454.14 | 5028.015512 | -1.394708132 | 0.119986782 | -6.6233 | 3.51E-11 | 6.97E-10 | ENSG00000147454 | SLC25A37 | 51312 |
| ENSG00000116117.18 | 138.1926217 | -1.394983924 | 0.205603183 | -3.86659 | 0.00011 | 0.000990616 | ENSG00000116117 | PARD3B | 117583 |
| ENSG00000198807.12 | 89.83127197 | -1.395048599 | 0.178342481 | -4.45799 | 8.27E-06 | 8.90E-05 | ENSG00000198807 | PAX9 | 5083 |
| ENSG00000179583.19 | 47.47594329 | -1.395407496 | 0.246338313 | -3.22892 | 0.001243 | 0.009011105 | ENSG00000179583 | CIITA | 4261 |
| ENSG00000111247.14 | 1037.83136 | -1.39663329 | 0.159263938 | -5.00197 | 5.67E-07 | 7.09E-06 | ENSG00000111247 | RAD51AP1 | 10635 |
| ENSG00000168813.17 | 1381.931 | -1.397303 | 0.1682011 | -4.74018 | 2.14E-06 | 2.46E-05 | ENSG00000168813 | ZNF507 | 22847 |
| ENSG00000134222.16 | 1177.688109 | -1.398342375 | 0.150553826 | -5.3027 | 1.14E-07 | 1.56E-06 | ENSG00000134222 | PSRC1 | 84722 |
| ENSG00000181192.12 | 1960.984822 | -1.40027671 | 0.076478306 | -10.4641 | 1.26E-25 | 6.68E-24 | ENSG00000181192 | DHTKD1 | 55526 |
| ENSG00000100342.20 | 1018.183 | -1.400461 | 0.0545944 | -14.662 | 1.13E-48 | 1.52E-46 | ENSG00000100342 | APOL1 | 8542 |
| ENSG00000231466.2 | 18.84384 | -1.400633 | 0.3031475 | -2.64107 | 0.008265 | 0.04860125 | ENSG00000231466 | NA | NA |
| ENSG00000171208.9 | 3542.833712 | -1.403587585 | 0.16137827 | -4.97953 | 6.37E-07 | 7.92E-06 | ENSG00000171208 | NETO2 | 81831 |
| ENSG00000074590.13 | 802.1517792 | -1.405064449 | 0.202101603 | -3.98346 | 6.79E-05 | 0.000629566 | ENSG00000074590 | NUAK1 | 9891 |
| ENSG00000035499.13 | 2861.570702 | -1.407320124 | 0.096348568 | -8.37916 | 5.33E-17 | 1.68E-15 | ENSG00000035499 | DEPDC1B | 55789 |
| ENSG00000146950.13 | 277.7035 | -1.409299 | 0.1852297 | -4.36917 | 1.25E-05 | 0.000130007 | ENSG00000146950 | SHROOM2 | 357 |
| ENSG00000164342.12 | 462.601285 | -1.409402295 | 0.135612459 | -5.9685 | 2.39E-09 | 3.96E-08 | ENSG00000164342 | TLR3 | 7098 |
| ENSG00000100473.18 | 37.82870275 | -1.409540931 | 0.265471183 | -3.04945 | 0.002293 | 0.015629689 | ENSG00000100473 | COCH | 1690 |
| ENSG00000125630.15 | 2718.06007 | -1.409789251 | 0.155292967 | -5.21459 | 1.84E-07 | 2.46E-06 | ENSG00000125630 | POLR1B | 84172 |
| ENSG00000178409.13 | 576.4416389 | -1.41005844 | 0.166773779 | -4.85723 | 1.19E-06 | 1.42E-05 | ENSG00000178409 | BEND3 | 57673 |
| ENSG00000263843.1 | 23.0133707 | -1.411441182 | 0.261484396 | -3.10321 | 0.001914 | 0.01326135 | ENSG00000263843 | LOC10028 | 1E+08 |
| ENSG00000138074.15 | 1223.367792 | -1.412615206 | 0.157233199 | -5.16822 | 2.36E-07 | 3.10E-06 | ENSG00000138074 | SLC5A6 | 8884 |
| ENSG00000170312.15 | 4332.936097 | -1.413405057 | 0.134518502 | -6.04679 | 1.48E-09 | 2.49E-08 | ENSG00000170312 | CDK1 | 983 |

| | | | | | | | | | |
|---|---|---|---|---|---|---|---|---|---|
| ENSG00000198794.12 | 345.9559608 | -1.41396959 | 0.188724003 | -4.31302 | 1.61E-05 | 0.000164579 | ENSG00000198794 | SCAMP5 | 192683 |
| ENSG00000099256.19 | 36.36109729 | -1.414974483 | 0.309519383 | -2.63303 | 0.008463 | 0.049547447 | ENSG00000099256 | PRTFDC1 | 56952 |
| ENSG00000100077.15 | 650.3319 | -1.416579 | 0.07213235 | -11.3206 | 1.04E-29 | 7.05E-28 | ENSG00000100077 | GRK3 | 157 |
| ENSG00000198901.14 | 78.98572124 | -1.416612551 | 0.167252727 | -4.88251 | 1.05E-06 | 1.26E-05 | ENSG00000198901 | PRC1 | 9055 |
| ENSG00000109685.18 | 7782.609366 | -1.417614328 | 0.105461607 | -7.75272 | 8.99E-15 | 2.42E-13 | ENSG00000109685 | NSD2 | 7468 |
| ENSG00000167670.16 | 2244.902916 | -1.417789892 | 0.140937113 | -5.80252 | 6.53E-09 | 1.03E-07 | ENSG00000167670 | CHAF1A | 10036 |
| ENSG00000107105.15 | 564.480027 | -1.418918701 | 0.103551209 | -7.90835 | 2.61E-15 | 7.34E-14 | ENSG00000107105 | ELAVL2 | 1993 |
| ENSG00000017483.15 | 3397.786 | -1.419558 | 0.2039983 | -4.01747 | 5.88E-05 | 0.000550458 | ENSG00000017483 | SLC38A5 | 92745 |
| ENSG00000108839.12 | 273.8143003 | -1.421501233 | 0.131925003 | -6.22703 | 4.75E-10 | 8.52E-09 | ENSG00000108839 | ALOX12 | 239 |
| ENSG00000183072.10 | 344.7958942 | -1.421517601 | 0.163362063 | -5.02882 | 4.94E-07 | 6.20E-06 | ENSG00000183072 | NKX2-5 | 1482 |
| ENSG00000073536.18 | 1214.628889 | -1.422277354 | 0.123719476 | -6.6463 | 3.01E-11 | 6.02E-10 | ENSG00000073536 | NLE1 | 54475 |
| ENSG00000053438.11 | 96.99741 | -1.422786 | 0.1629127 | -5.05047 | 4.41E-07 | 5.57E-06 | ENSG00000053438 | NNAT | 4826 |
| ENSG00000088325.16 | 6603.017 | -1.42959 | 0.1690738 | -4.90667 | 9.26E-07 | 1.13E-05 | ENSG00000088325 | TPX2 | 22974 |
| ENSG00000272455.1 | 126.5859758 | -1.429988478 | 0.166790145 | -4.97624 | 6.48E-07 | 8.05E-06 | ENSG00000272455 | NA | NA |
| ENSG00000281398.4 | 3565.080706 | -1.430443818 | 0.186975672 | -4.44145 | 8.94E-06 | 9.55E-05 | ENSG00000281398 | NA | NA |
| ENSG00000111696.12 | 663.0733584 | -1.431807759 | 0.180746072 | -4.60208 | 4.18E-06 | 4.65E-05 | ENSG00000111696 | NT5DC3 | 51559 |
| ENSG00000171843.16 | 201.7864913 | -1.432128847 | 0.17251154 | -4.82361 | 1.41E-06 | 1.66E-05 | ENSG00000171843 | MLLT3 | 4300 |
| ENSG00000227372.12 | 145.3260784 | -1.432480181 | 0.124452787 | -6.68912 | 2.25E-11 | 4.55E-10 | ENSG00000227372 | TP73-AS1 | 57212 |
| ENSG00000177542.11 | 1157.528971 | -1.432751603 | 0.161601795 | -5.15311 | 2.56E-07 | 3.35E-06 | ENSG00000177542 | SLC25A22 | 79751 |
| ENSG00000013573.17 | 1730.932887 | -1.432774596 | 0.158955599 | -5.23904 | 1.61E-07 | 2.17E-06 | ENSG00000013573 | DDX11 | 1663 |
| ENSG00000154127.10 | 513.2910539 | -1.433155548 | 0.08428698 | -9.88475 | 4.85E-23 | 2.20E-21 | ENSG00000154127 | UBASH3B | 84959 |
| ENSG00000121690.11 | 107.5544309 | -1.43385081 | 0.272202922 | -3.06334 | 0.002189 | 0.015002587 | ENSG00000121690 | DEPDC7 | 91614 |
| ENSG00000003137.8 | 270.8270586 | -1.437319702 | 0.128949526 | -6.49339 | 8.39E-11 | 1.61E-09 | ENSG00000003137 | CYP26B1 | 56603 |
| ENSG00000010030.14 | 173.8504107 | -1.437625577 | 0.15228486 | -5.50039 | 3.79E-08 | 5.45E-07 | ENSG00000010030 | ETV7 | 51513 |
| ENSG00000180801.14 | 732.9498609 | -1.440008428 | 0.231766757 | -3.62437 | 0.00029 | 0.002400845 | ENSG00000180801 | ARSJ | 79642 |
| ENSG00000113810.16 | 7888.684687 | -1.440024481 | 0.128659449 | -6.52905 | 6.62E-11 | 1.28E-09 | ENSG00000113810 | SMC4 | 10051 |
| ENSG00000110328.6 | 630.5787406 | -1.440404468 | 0.119709572 | -7.02036 | 2.21E-12 | 4.91E-11 | ENSG00000110328 | GALNT18 | 374378 |
| ENSG00000054690.13 | 864.6249945 | -1.443058417 | 0.079960474 | -10.5434 | 5.45E-26 | 2.92E-24 | ENSG00000054690 | PLEKHH1 | 57475 |
| ENSG00000104889.6 | 28.54211153 | -1.444437831 | 0.256389609 | -3.29357 | 0.000989 | 0.007341849 | ENSG00000104889 | RNASEH2 | 10535 |
| ENSG00000204267.15 | 78.81318272 | -1.444470212 | 0.15292254 | -5.52221 | 3.35E-08 | 4.86E-07 | ENSG00000204267 | TAP2 | 6891 |
| ENSG00000103257.9 | 74105.57016 | -1.445331559 | 0.082626407 | -10.2308 | 1.44E-24 | 7.09E-23 | ENSG00000103257 | SLC7A5 | 8140 |
| ENSG00000117791.16 | 66.58266366 | -1.446082154 | 0.230071387 | -3.67748 | 0.000236 | 0.001983829 | ENSG00000117791 | MTARC2 | 54996 |
| ENSG00000176890.16 | 982.6922338 | -1.446341289 | 0.225613339 | -3.75129 | 0.000176 | 0.001515209 | ENSG00000176890 | TYMS | 7298 |
| ENSG00000100714.17 | 5324.176483 | -1.448845618 | 0.178528342 | -4.75468 | 1.99E-06 | 2.30E-05 | ENSG00000100714 | MTHFD1 | 4522 |
| ENSG00000143228.13 | 1335.335292 | -1.44896168 | 0.144028935 | -5.89438 | 3.76E-09 | 6.08E-08 | ENSG00000143228 | NUF2 | 83540 |
| ENSG00000184992.11 | 1664.47494 | -1.450898203 | 0.128798746 | -6.60642 | 3.94E-11 | 7.79E-10 | ENSG00000184992 | BRI3BP | 140707 |
| ENSG00000215883.10 | 181.9490208 | -1.453650679 | 0.119994505 | -7.11408 | 1.13E-12 | 2.57E-11 | ENSG00000215883 | CYB5RL | 606495 |
| ENSG00000238105.7 | 480.6335928 | -1.4544644 | 0.112231582 | -7.6134 | 2.67E-14 | 6.91E-13 | ENSG00000238105 | GOLGA2P | 55592 |
| ENSG00000286431.1 | 86.27736 | -1.455115 | 0.1675344 | -5.10411 | 3.32E-07 | 4.28E-06 | ENSG00000286431 | NA | NA |
| ENSG00000116127.18 | 1412.94579 | -1.455396924 | 0.150706369 | -5.67592 | 1.38E-08 | 2.09E-07 | ENSG00000116127 | ALMS1 | 7840 |
| ENSG00000149503.13 | 2559.463008 | -1.456285899 | 0.153462797 | -5.57976 | 2.41E-08 | 3.57E-07 | ENSG00000149503 | INCENP | 3619 |
| ENSG00000076248.10 | 2175.210561 | -1.456676989 | 0.165997828 | -5.16077 | 2.46E-07 | 3.22E-06 | ENSG00000076248 | UNG | 7374 |

| | | | | | | | | | |
|---|---|---|---|---|---|---|---|---|---|
| ENSG00000171115.4 | 35.31178674 | -1.458599823 | 0.236816126 | -3.6256 | 0.000288 | 0.002391705 | ENSG00000171115 | GIMAP8 | 155038 |
| ENSG00000180346.3 | 588.6502628 | -1.461161455 | 0.100321146 | -8.58405 | 9.16E-18 | 3.05E-16 | ENSG00000180346 | TIGD2 | 166815 |
| ENSG00000142207.7 | 2323.263 | -1.461838 | 0.1754057 | -4.9134 | 8.95E-07 | 1.09E-05 | ENSG00000142207 | URB1 | 9875 |
| ENSG00000159259.8 | 924.1225 | -1.462652 | 0.1291081 | -6.68163 | 2.36E-11 | 4.78E-10 | ENSG00000159259 | CHAF1B | 8208 |
| ENSG00000121211.7 | 604.6944334 | -1.464122787 | 0.114095543 | -7.57368 | 3.63E-14 | 9.28E-13 | ENSG00000121211 | MND1 | 84057 |
| ENSG00000091651.9 | 1395.282332 | -1.464142577 | 0.170888171 | -5.05677 | 4.26E-07 | 5.40E-06 | ENSG00000091651 | ORC6 | 23594 |
| ENSG00000064042.18 | 63.25665405 | -1.46474297 | 0.326010615 | -2.6525 | 0.00799 | 0.04715255 | ENSG00000064042 | LIMCH1 | 22998 |
| ENSG00000112159.12 | 4559.105642 | -1.466257494 | 0.156473044 | -5.53615 | 3.09E-08 | 4.52E-07 | ENSG00000112159 | MDN1 | 23195 |
| ENSG00000134326.11 | 341.9544007 | -1.47059341 | 0.310959922 | -2.7997 | 0.005115 | 0.031890707 | ENSG00000134326 | CMPK2 | 129607 |
| ENSG00000231345.3 | 91.80637067 | -1.470874572 | 0.194550773 | -4.47634 | 7.59E-06 | 8.20E-05 | ENSG00000231345 | NA | NA |
| ENSG00000286545.1 | 49.7166823 | -1.471001234 | 0.28048392 | -3.10535 | 0.001901 | 0.013176009 | ENSG00000286545 | NA | NA |
| ENSG00000227473.1 | 70.67105418 | -1.473402599 | 0.18080854 | -4.83054 | 1.36E-06 | 1.61E-05 | ENSG00000227473 | TSSK5P | 648630 |
| ENSG00000132780.17 | 7620.684587 | -1.473411117 | 0.158641236 | -5.50557 | 3.68E-08 | 5.30E-07 | ENSG00000132780 | NASP | 4678 |
| ENSG00000184916.9 | 2136.836032 | -1.474811408 | 0.091386806 | -9.57262 | 1.04E-21 | 4.41E-20 | ENSG00000184916 | JAG2 | 3714 |
| ENSG00000169683.8 | 700.6747463 | -1.475871885 | 0.092286391 | -9.4908 | 2.29E-21 | 9.48E-20 | ENSG00000169683 | LRRC45 | 201255 |
| ENSG00000164109.14 | 3887.394482 | -1.476758341 | 0.168177234 | -5.2133 | 1.86E-07 | 2.47E-06 | ENSG00000164109 | MAD2L1 | 4085 |
| ENSG00000283486.2 | 33.88196777 | -1.476866353 | 0.31464108 | -2.78688 | 0.005322 | 0.033048344 | ENSG00000283486 | FAM95C | 1E+08 |
| ENSG00000134470.21 | 426.8431201 | -1.476923303 | 0.107271505 | -8.1748 | 2.96E-16 | 8.82E-15 | ENSG00000134470 | IL15RA | 3601 |
| ENSG00000157456.8 | 2379.805731 | -1.477072011 | 0.14287065 | -6.13892 | 8.31E-10 | 1.44E-08 | ENSG00000157456 | CCNB2 | 9133 |
| ENSG00000131759.18 | 288.634695 | -1.478768792 | 0.131742039 | -6.67037 | 2.55E-11 | 5.14E-10 | ENSG00000131759 | RARA | 5914 |
| ENSG00000198830.11 | 7831.347587 | -1.47939724 | 0.092470352 | -9.51005 | 1.91E-21 | 7.92E-20 | ENSG00000198830 | HMGN2 | 3151 |
| ENSG00000135119.14 | 90.3356178 | -1.479419993 | 0.154020885 | -5.70975 | 1.13E-08 | 1.74E-07 | ENSG00000135119 | RNFT2 | 84900 |
| ENSG00000247626.4 | 920.0191776 | -1.479507513 | 0.202481792 | -4.34364 | 1.40E-05 | 0.000144888 | ENSG00000247626 | MARS2 | 92935 |
| ENSG00000230330.1 | 79.05479316 | -1.479668608 | 0.172527337 | -5.09872 | 3.42E-07 | 4.39E-06 | ENSG00000230330 | NA | NA |
| ENSG00000134057.15 | 12606.3566 | -1.480182785 | 0.170846796 | -5.15188 | 2.58E-07 | 3.37E-06 | ENSG00000134057 | CCNB1 | 891 |
| ENSG00000186638.16 | 526.9044301 | -1.481754524 | 0.180764713 | -4.87791 | 1.07E-06 | 1.29E-05 | ENSG00000186638 | KIF24 | 347240 |
| ENSG00000112029.10 | 1183.409241 | -1.482385528 | 0.204738315 | -4.30982 | 1.63E-05 | 0.000166718 | ENSG00000112029 | FBXO5 | 26271 |
| ENSG00000086570.12 | 14511.9614 | -1.482523906 | 0.174114294 | -5.06865 | 4.01E-07 | 5.10E-06 | ENSG00000086570 | FAT2 | 2196 |
| ENSG00000154898.15 | 69.12950949 | -1.483090457 | 0.18064099 | -4.88865 | 1.02E-06 | 1.23E-05 | ENSG00000154898 | NA | NA |
| ENSG00000118707.10 | 460.4996 | -1.483269 | 0.1690144 | -5.226 | 1.73E-07 | 2.32E-06 | ENSG00000118707 | TGIF2 | 60436 |
| ENSG00000283341.2 | 39.68258388 | -1.483585325 | 0.297786852 | -2.96717 | 0.003006 | 0.019918523 | ENSG00000283341 | NA | NA |
| ENSG00000100479.13 | 337.0465835 | -1.484526012 | 0.132107243 | -6.69551 | 2.15E-11 | 4.37E-10 | ENSG00000100479 | POLE2 | 5427 |
| ENSG00000260401.1 | 154.3418205 | -1.484800212 | 0.203989096 | -4.33749 | 1.44E-05 | 0.000148482 | ENSG00000260401 | NA | NA |
| ENSG00000171241.9 | 1811.999206 | -1.485306033 | 0.225728893 | -3.92199 | 8.78E-05 | 0.000800404 | ENSG00000171241 | SHCBP1 | 79801 |
| ENSG00000130224.15 | 92.24555 | -1.48534 | 0.1734938 | -5.103 | 3.34E-07 | 4.30E-06 | ENSG00000130224 | LRCH2 | 57631 |
| ENSG00000267102.1 | 19.1491315 | -1.485761481 | 0.307488576 | -2.88063 | 0.003969 | 0.025490229 | ENSG00000267102 | NA | NA |
| ENSG00000205213.14 | 1365.275211 | -1.485905165 | 0.171293272 | -5.17186 | 2.32E-07 | 3.05E-06 | ENSG00000205213 | LGR4 | 55366 |
| ENSG00000075643.6 | 2053.817573 | -1.48784275 | 0.129140047 | -6.87504 | 6.20E-12 | 1.33E-10 | ENSG00000075643 | MOCOS | 55034 |
| ENSG00000126787.13 | 3333.12377 | -1.488009582 | 0.140054595 | -6.34045 | 2.29E-10 | 4.25E-09 | ENSG00000126787 | DLGAP5 | 9787 |
| ENSG00000240449.1 | 17.09208445 | -1.488937525 | 0.300665139 | -2.95657 | 0.003111 | 0.020538049 | ENSG00000240449 | NA | NA |
| ENSG00000059573.9 | 3382.982506 | -1.491706298 | 0.171367779 | -5.20347 | 1.96E-07 | 2.60E-06 | ENSG00000059573 | ALDH18A1 | 5832 |
| ENSG00000261485.1 | 22.7571362 | -1.491734118 | 0.320898242 | -2.77887 | 0.005455 | 0.033795473 | ENSG00000261485 | NA | NA |

| | | | | | | | | | |
|---|---|---|---|---|---|---|---|---|---|
| ENSG00000197905.9 | 834.0451686 | -1.494682115 | 0.13533481 | -6.61088 | 3.82E-11 | 7.56E-10 | ENSG00000197905 | TEAD4 | 7004 |
| ENSG00000149639.15 | 2365.295 | -1.49478 | 0.1062484 | -8.42159 | 3.71E-17 | 1.18E-15 | ENSG00000149639 | SOGA1 | 140710 |
| ENSG00000157240.3 | 164.8273126 | -1.494829042 | 0.208065922 | -4.3007 | 1.70E-05 | 0.000173068 | ENSG00000157240 | FZD1 | 8321 |
| ENSG00000140534.13 | 1306.903386 | -1.496208838 | 0.156511266 | -5.72616 | 1.03E-08 | 1.59E-07 | ENSG00000140534 | TICRR | 90381 |
| ENSG00000178921.14 | 1496.056717 | -1.498181619 | 0.225723592 | -3.97912 | 6.92E-05 | 0.000640283 | ENSG00000178921 | PFAS | 5198 |
| ENSG00000175183.10 | 334.5939212 | -1.499123368 | 0.105213944 | -8.54567 | 1.28E-17 | 4.21E-16 | ENSG00000175183 | CSRP2 | 1466 |
| ENSG00000138646.9 | 1063.820839 | -1.500241096 | 0.203353749 | -4.42697 | 9.56E-06 | 0.000101603 | ENSG00000138646 | HERC5 | 51191 |
| ENSG00000130675.15 | 56.4190329 | -1.500421487 | 0.267223496 | -3.36954 | 0.000753 | 0.005748314 | ENSG00000130675 | MNX1 | 3110 |
| ENSG00000133740.11 | 438.1743904 | -1.502591116 | 0.133492388 | -6.76137 | 1.37E-11 | 2.83E-10 | ENSG00000133740 | E2F5 | 1875 |
| ENSG00000129810.15 | 467.8100955 | -1.505278927 | 0.16354324 | -5.53541 | 3.11E-08 | 4.54E-07 | ENSG00000129810 | SGO1 | 151648 |
| ENSG00000198768.10 | 25.644 | -1.505372 | 0.3435613 | -2.63526 | 0.008407 | 0.04927826 | ENSG00000198768 | APCDD1L | 164284 |
| ENSG00000093217.10 | 181.3137658 | -1.506952972 | 0.180647699 | -5.02056 | 5.15E-07 | 6.46E-06 | ENSG00000093217 | XYLB | 9942 |
| ENSG00000170837.2 | 19.19146386 | -1.506957127 | 0.313070881 | -2.89697 | 0.003768 | 0.024311132 | ENSG00000170837 | GPR27 | 2850 |
| ENSG00000109576.14 | 38.62838084 | -1.507714419 | 0.230707811 | -3.93448 | 8.34E-05 | 0.000763307 | ENSG00000109576 | AADAT | 51166 |
| ENSG00000279278.1 | 85.78237 | -1.50864 | 0.198622 | -4.57472 | 4.77E-06 | 5.27E-05 | ENSG00000279278 | NA | NA |
| ENSG00000138180.15 | 3465.540829 | -1.509752755 | 0.160710626 | -5.66081 | 1.51E-08 | 2.28E-07 | ENSG00000138180 | CEP55 | 55165 |
| ENSG00000277268.2 | 79.32558965 | -1.509753326 | 0.181225191 | -5.02002 | 5.17E-07 | 6.48E-06 | ENSG00000277268 | LHX1-DT | 1.03E+08 |
| ENSG00000272405.1 | 95.95699551 | -1.509956828 | 0.146971577 | -6.19138 | 5.96E-10 | 1.06E-08 | ENSG00000272405 | NA | NA |
| ENSG00000108018.15 | 39.11415997 | -1.513434487 | 0.277378464 | -3.2931 | 0.000991 | 0.007346063 | ENSG00000108018 | SORCS1 | 114815 |
| ENSG00000125347.14 | 1912.92262 | -1.513708499 | 0.092054255 | -9.92576 | 3.22E-23 | 1.47E-21 | ENSG00000125347 | IRF1 | 3659 |
| ENSG00000183914.14 | 22.34710907 | -1.516265624 | 0.280996672 | -3.26077 | 0.001111 | 0.008157666 | ENSG00000183914 | DNAH2 | 146754 |
| ENSG00000165512.5 | 491.5273181 | -1.517984914 | 0.104175215 | -8.81193 | 1.23E-18 | 4.31E-17 | ENSG00000165512 | ZNF22 | 7570 |
| ENSG00000183763.9 | 437.6743377 | -1.519309149 | 0.180420232 | -5.09538 | 3.48E-07 | 4.47E-06 | ENSG00000183763 | TRAIP | 10293 |
| ENSG00000205045.9 | 22.3391203 | -1.519817382 | 0.294566139 | -3.12262 | 0.001793 | 0.012531533 | ENSG00000205045 | SLFN12L | 1.01E+08 |
| ENSG00000164741.15 | 264.0025258 | -1.520339273 | 0.226559514 | -4.06224 | 4.86E-05 | 0.000460967 | ENSG00000164741 | DLC1 | 10395 |
| ENSG00000106785.15 | 1723.719922 | -1.52409302 | 0.104637012 | -8.83142 | 1.03E-18 | 3.65E-17 | ENSG00000106785 | TRIM14 | 9830 |
| ENSG00000083307.12 | 1738.417156 | -1.526244771 | 0.202322748 | -4.57806 | 4.69E-06 | 5.19E-05 | ENSG00000083307 | GRHL2 | 79977 |
| ENSG00000144724.20 | 1771.231561 | -1.527290195 | 0.157813673 | -5.87585 | 4.21E-09 | 6.75E-08 | ENSG00000144724 | PTPRG | 5793 |
| ENSG00000253729.7 | 28906.30456 | -1.527690523 | 0.157354723 | -5.89554 | 3.73E-09 | 6.04E-08 | ENSG00000253729 | PRKDC | 5591 |
| ENSG00000163520.14 | 31.84619763 | -1.527776145 | 0.289231948 | -3.20772 | 0.001338 | 0.009628975 | ENSG00000163520 | FBLN2 | 2199 |
| ENSG00000105538.10 | 313.0422 | -1.529086 | 0.0913857 | -10.1667 | 2.79E-24 | 1.35E-22 | ENSG00000105538 | RASIP1 | 54922 |
| ENSG00000197536.11 | 198.0166439 | -1.53080133 | 0.105861341 | -8.79265 | 1.46E-18 | 5.10E-17 | ENSG00000197536 | NA | NA |
| ENSG00000117650.13 | 1048.263482 | -1.53098402 | 0.1818557 | -5.11936 | 3.07E-07 | 3.96E-06 | ENSG00000117650 | NEK2 | 4751 |
| ENSG00000166508.17 | 7301.72811 | -1.531127564 | 0.148162725 | -6.28449 | 3.29E-10 | 6.01E-09 | ENSG00000166508 | MCM7 | 4176 |
| ENSG00000284946.1 | 170.3999943 | -1.5329542 | 0.198029591 | -4.71119 | 2.46E-06 | 2.82E-05 | ENSG00000284946 | NA | NA |
| ENSG00000080986.13 | 1773.664156 | -1.534524893 | 0.15807529 | -5.9119 | 3.38E-09 | 5.49E-08 | ENSG00000080986 | NDC80 | 10403 |
| ENSG00000062822.14 | 1787.909 | -1.536429 | 0.1263903 | -7.40903 | 1.27E-13 | 3.15E-12 | ENSG00000062822 | POLD1 | 5424 |
| ENSG00000115604.10 | 48.87761368 | -1.537668781 | 0.210526685 | -4.45392 | 8.43E-06 | 9.05E-05 | ENSG00000115604 | IL18R1 | 8809 |
| ENSG00000255624.3 | 79.77832633 | -1.539101554 | 0.156184545 | -6.01277 | 1.82E-09 | 3.05E-08 | ENSG00000255624 | NA | NA |
| ENSG00000132164.10 | 135.68589 | -1.539225437 | 0.17630934 | -5.32715 | 9.98E-08 | 1.37E-06 | ENSG00000132164 | SLC6A11 | 6538 |
| ENSG00000110881.11 | 73.41464948 | -1.539727732 | 0.281254568 | -3.3412 | 0.000834 | 0.006312277 | ENSG00000110881 | ASIC1 | 41 |
| ENSG00000102384.13 | 889.3503 | -1.540224 | 0.1773025 | -5.30294 | 1.14E-07 | 1.56E-06 | ENSG00000102384 | CENPI | 2491 |

| | | | | | | | | | |
|---|---|---|---|---|---|---|---|---|---|
| ENSG00000140263.14 | 728.7103955 | -1.542735295 | 0.114254667 | -8.25118 | 1.57E-16 | 4.77E-15 | ENSG00000140263 | SORD | 6652 |
| ENSG00000269473.1 | 20.25173 | -1.545498 | 0.3056103 | -3.09381 | 0.001976 | 0.01365353 | ENSG00000269473 | NA | NA |
| ENSG00000263513.5 | 37.20977896 | -1.547219905 | 0.27573317 | -3.43528 | 0.000592 | 0.00461865 | ENSG00000263513 | FAM72C | 554282 |
| ENSG00000179240.10 | 213.1237078 | -1.548893319 | 0.129513996 | -7.32657 | 2.36E-13 | 5.71E-12 | ENSG00000179240 | GVQW3 | 1.01E+08 |
| ENSG00000276529.1 | 55.19088 | -1.549327 | 0.1921914 | -4.93949 | 7.83E-07 | 9.62E-06 | ENSG00000276529 | NA | NA |
| ENSG00000058804.12 | 4686.103885 | -1.554304823 | 0.181452015 | -5.25927 | 1.45E-07 | 1.96E-06 | ENSG00000058804 | NDC1 | 55706 |
| ENSG00000138346.15 | 1094.190478 | -1.555670016 | 0.094226 | -10.1423 | 3.58E-24 | 1.72E-22 | ENSG00000138346 | DNA2 | 1763 |
| ENSG00000024526.17 | 1518.348721 | -1.556706893 | 0.150089422 | -6.37425 | 1.84E-10 | 3.43E-09 | ENSG00000024526 | DEPDC1 | 55635 |
| ENSG00000114251.14 | 73.84614808 | -1.557330476 | 0.168973656 | -5.66556 | 1.47E-08 | 2.22E-07 | ENSG00000114251 | WNT5A | 7474 |
| ENSG00000164087.7 | 805.2954363 | -1.557846251 | 0.122777937 | -7.80145 | 6.12E-15 | 1.67E-13 | ENSG00000164087 | POC1A | 25886 |
| ENSG00000178718.6 | 155.6222963 | -1.558457819 | 0.138875477 | -6.90156 | 5.14E-12 | 1.11E-10 | ENSG00000178718 | RPP25 | 54913 |
| ENSG00000204588.5 | 42.99743576 | -1.561721877 | 0.222517135 | -4.32201 | 1.55E-05 | 0.000158433 | ENSG00000204588 | LINC01123 | 440894 |
| ENSG00000071539.14 | 2152.196903 | -1.563005287 | 0.117980959 | -8.16238 | 3.28E-16 | 9.74E-15 | ENSG00000071539 | TRIP13 | 9319 |
| ENSG00000187951.11 | 118.9661053 | -1.563541753 | 0.227617441 | -4.23316 | 2.30E-05 | 0.000229754 | ENSG00000187951 | LOC10028 | 1E+08 |
| ENSG00000259488.2 | 25.91798988 | -1.56354505 | 0.309810164 | -3.11011 | 0.00187 | 0.01299611 | ENSG00000259488 | NA | NA |
| ENSG00000137404.14 | 83.09574408 | -1.565024778 | 0.196569043 | -4.90934 | 9.14E-07 | 1.11E-05 | ENSG00000137404 | NRM | 11270 |
| ENSG00000287001.1 | 21.90545 | -1.566249 | 0.2948415 | -3.27718 | 0.001048 | 0.007738563 | ENSG00000287001 | NA | NA |
| ENSG00000276603.1 | 40.38473 | -1.56737 | 0.2382334 | -4.0606 | 4.89E-05 | 0.000463729 | ENSG00000276603 | NA | NA |
| ENSG00000120800.5 | 2950.935924 | -1.568036607 | 0.176863584 | -5.47335 | 4.42E-08 | 6.30E-07 | ENSG00000120800 | UTP20 | 27340 |
| ENSG00000138376.11 | 749.3965588 | -1.568871674 | 0.175431537 | -5.52279 | 3.34E-08 | 4.85E-07 | ENSG00000138376 | BARD1 | 580 |
| ENSG00000183048.12 | 200.0206393 | -1.569120552 | 0.173668705 | -5.58028 | 2.40E-08 | 3.56E-07 | ENSG00000183048 | SLC25A10 | 1468 |
| ENSG00000250802.7 | 30.21756244 | -1.570629512 | 0.265649725 | -3.65379 | 0.000258 | 0.00215776 | ENSG00000250802 | ZBED3-AS | 728723 |
| ENSG00000122574.10 | 123.0278396 | -1.572011 | 0.145042132 | -6.70158 | 2.06E-11 | 4.20E-10 | ENSG00000122574 | WIPF3 | 644150 |
| ENSG00000179598.6 | 162.3056948 | -1.575329585 | 0.122928858 | -7.9341 | 2.12E-15 | 6.00E-14 | ENSG00000179598 | PLD6 | 201164 |
| ENSG00000172167.8 | 865.0595272 | -1.57785368 | 0.088598883 | -11.0369 | 2.54E-28 | 1.57E-26 | ENSG00000172167 | MTBP | 27085 |
| ENSG00000135763.10 | 1381.36888 | -1.578668407 | 0.188439744 | -5.19354 | 2.06E-07 | 2.72E-06 | ENSG00000135763 | URB2 | 9816 |
| ENSG00000221955.10 | 201.9131518 | -1.581961787 | 0.149649373 | -6.56175 | 5.32E-11 | 1.04E-09 | ENSG00000221955 | SLC12A8 | 84561 |
| ENSG00000090889.12 | 1954.989 | -1.582424 | 0.1879951 | -5.2258 | 1.73E-07 | 2.32E-06 | ENSG00000090889 | KIF4A | 24137 |
| ENSG00000221740.1 | 25.6050735 | -1.584283181 | 0.330379957 | -2.97925 | 0.00289 | 0.019241263 | ENSG00000221740 | SNORD93 | 692210 |
| ENSG00000218336.9 | 322.1067556 | -1.585585346 | 0.134904147 | -7.30582 | 2.76E-13 | 6.60E-12 | ENSG00000218336 | TENM3 | 55714 |
| ENSG00000094804.12 | 2483.931321 | -1.585822005 | 0.209294738 | -4.71021 | 2.47E-06 | 2.83E-05 | ENSG00000094804 | CDC6 | 990 |
| ENSG00000117877.10 | 295.7572 | -1.586292 | 0.1676869 | -5.88174 | 4.06E-09 | 6.54E-08 | ENSG00000117877 | POLR1G | 10849 |
| ENSG00000241484.9 | 181.822 | -1.596623 | 0.1375137 | -7.24745 | 4.25E-13 | 1.00E-11 | ENSG00000241484 | ARHGAP8 | 23779 |
| ENSG00000019582.15 | 1293.286945 | -1.600031481 | 0.118932068 | -8.40843 | 4.16E-17 | 1.32E-15 | ENSG00000019582 | CD74 | 972 |
| ENSG00000126500.3 | 26.11857848 | -1.602318924 | 0.267055809 | -3.75322 | 0.000175 | 0.001505059 | ENSG00000126500 | FLRT1 | 23769 |
| ENSG00000164070.12 | 1825.886068 | -1.602426737 | 0.079006858 | -12.6878 | 6.91E-37 | 6.15E-35 | ENSG00000164070 | HSPA4L | 22824 |
| ENSG00000082438.16 | 1018.941003 | -1.604187214 | 0.13688418 | -7.33604 | 2.20E-13 | 5.35E-12 | ENSG00000082438 | COBLL1 | 22837 |
| ENSG00000115163.15 | 930.8939686 | -1.604330981 | 0.185706064 | -5.40818 | 6.37E-08 | 8.95E-07 | ENSG00000115163 | CENPA | 1058 |
| ENSG00000112118.19 | 6746.87229 | -1.605013899 | 0.181354387 | -5.54171 | 3.00E-08 | 4.39E-07 | ENSG00000112118 | MCM3 | 4172 |
| ENSG00000248432.1 | 60.98947799 | -1.605819522 | 0.174493918 | -5.76421 | 8.20E-09 | 1.28E-07 | ENSG00000248432 | NA | NA |
| ENSG00000146263.11 | 1599.704643 | -1.606378344 | 0.137476524 | -7.32037 | 2.47E-13 | 5.96E-12 | ENSG00000146263 | MMS22L | 253714 |
| ENSG00000012048.22 | 2781.572384 | -1.608591962 | 0.149268797 | -6.75688 | 1.41E-11 | 2.91E-10 | ENSG00000012048 | BRCA1 | 672 |

| | | | | | | | | | |
|---|---|---|---|---|---|---|---|---|---|
| ENSG00000085840.13 | 956.9097496 | -1.609580128 | 0.230534955 | -4.37929 | 1.19E-05 | 0.000124401 | ENSG00000085840 | ORC1 | 4998 |
| ENSG00000118495.19 | 400.7144902 | -1.609795697 | 0.081451568 | -12.3975 | 2.70E-35 | 2.26E-33 | ENSG00000118495 | PLAGL1 | 5325 |
| ENSG00000187498.16 | 2006.484035 | -1.611100063 | 0.260628722 | -3.87947 | 0.000105 | 0.000942177 | ENSG00000187498 | COL4A1 | 1282 |
| ENSG00000153721.19 | 75.39004228 | -1.615133086 | 0.179610528 | -5.65186 | 1.59E-08 | 2.40E-07 | ENSG00000153721 | CNKSR3 | 154043 |
| ENSG00000137124.8 | 1428.417218 | -1.61551863 | 0.18325104 | -5.54168 | 3.00E-08 | 4.39E-07 | ENSG00000137124 | ALDH1B1 | 219 |
| ENSG00000188807.13 | 991.5525903 | -1.616558559 | 0.175069655 | -5.80659 | 6.38E-09 | 1.01E-07 | ENSG00000188807 | TMEM201 | 199953 |
| ENSG00000128284.19 | 1033.311 | -1.621209 | 0.1396593 | -7.31214 | 2.63E-13 | 6.31E-12 | ENSG00000128284 | APOL3 | 80833 |
| ENSG00000105974.12 | 16596.45769 | -1.623999436 | 0.225483802 | -4.54134 | 5.59E-06 | 6.14E-05 | ENSG00000105974 | CAV1 | 857 |
| ENSG00000222112.1 | 18.85860887 | -1.625009632 | 0.382815569 | -2.67755 | 0.007416 | 0.044245163 | ENSG00000222112 | RN7SKP1( | 1.06E+08 |
| ENSG00000120162.10 | 816.9279328 | -1.628740078 | 0.082344966 | -12.4931 | 8.15E-36 | 6.97E-34 | ENSG00000120162 | MOB3B | 79817 |
| ENSG00000205592.14 | 17.7942337 | -1.63033346 | 0.339107197 | -3.03837 | 0.002379 | 0.016153914 | ENSG00000205592 | MUC19 | 283463 |
| ENSG00000130881.14 | 30.49896 | -1.632618 | 0.3566534 | -2.8953 | 0.003788 | 0.02441149 | ENSG00000130881 | LRP3 | 4037 |
| ENSG00000167900.12 | 2415.185321 | -1.634193815 | 0.134138619 | -7.70989 | 1.26E-14 | 3.34E-13 | ENSG00000167900 | TK1 | 7083 |
| ENSG00000101412.13 | 770.2833 | -1.634443 | 0.1876265 | -5.51331 | 3.52E-08 | 5.09E-07 | ENSG00000101412 | E2F1 | 1869 |
| ENSG00000147536.12 | 769.1132238 | -1.635803172 | 0.185390135 | -5.58715 | 2.31E-08 | 3.43E-07 | ENSG00000147536 | GINS4 | 84296 |
| ENSG00000154839.10 | 321.8892381 | -1.635849704 | 0.187365384 | -5.5285 | 3.23E-08 | 4.71E-07 | ENSG00000154839 | SKA1 | 220134 |
| ENSG00000128683.14 | 30.42264089 | -1.636785773 | 0.309263199 | -3.35244 | 0.000801 | 0.00608071 | ENSG00000128683 | GAD1 | 2571 |
| ENSG00000162817.7 | 131.4572882 | -1.636812539 | 0.249319512 | -4.15857 | 3.20E-05 | 0.000312561 | ENSG00000162817 | C1orf115 | 79762 |
| ENSG00000154920.14 | 247.9574521 | -1.637571473 | 0.218912025 | -4.73967 | 2.14E-06 | 2.47E-05 | ENSG00000154920 | EME1 | 146956 |
| ENSG00000169991.11 | 4439.125617 | -1.639415668 | 0.162128392 | -6.41107 | 1.45E-10 | 2.72E-09 | ENSG00000169991 | IFFO2 | 126917 |
| ENSG00000141510.17 | 1378.617477 | -1.641383828 | 0.178286127 | -5.84108 | 5.19E-09 | 8.26E-08 | ENSG00000141510 | TP53 | 7157 |
| ENSG00000144554.10 | 1957.200299 | -1.642991893 | 0.159713843 | -6.53038 | 6.56E-11 | 1.27E-09 | ENSG00000144554 | FANCD2 | 2177 |
| ENSG00000003096.14 | 302.2742 | -1.644718 | 0.0937504 | -11.1436 | 7.69E-29 | 4.90E-27 | ENSG00000003096 | KLHL13 | 90293 |
| ENSG00000182253.14 | 182.7229413 | -1.649496196 | 0.137157643 | -7.65175 | 1.98E-14 | 5.17E-13 | ENSG00000182253 | SYNM | 23336 |
| ENSG00000136167.14 | 267.6166913 | -1.649616699 | 0.193376422 | -5.42784 | 5.70E-08 | 8.04E-07 | ENSG00000136167 | LCP1 | 3936 |
| ENSG00000146918.19 | 4551.211418 | -1.651703352 | 0.149859646 | -7.01792 | 2.25E-12 | 5.00E-11 | ENSG00000146918 | NCAPG2 | 54892 |
| ENSG00000225345.3 | 72.2434457 | -1.652882805 | 0.228293519 | -4.61197 | 3.99E-06 | 4.45E-05 | ENSG00000225345 | NA | NA |
| ENSG00000100162.15 | 415.5317 | -1.654838 | 0.2135487 | -4.93957 | 7.83E-07 | 9.62E-06 | ENSG00000100162 | CENPM | 79019 |
| ENSG00000135451.13 | 965.6321421 | -1.655079636 | 0.197521927 | -5.34158 | 9.21E-08 | 1.28E-06 | ENSG00000135451 | TROAP | 10024 |
| ENSG00000164841.5 | 54.48010855 | -1.655752591 | 0.246965554 | -4.2749 | 1.91E-05 | 0.000192833 | ENSG00000164841 | TMEM74 | 157753 |
| ENSG00000010292.13 | 5973.753307 | -1.656623937 | 0.175408985 | -6.02377 | 1.70E-09 | 2.85E-08 | ENSG00000010292 | NCAPD2 | 9918 |
| ENSG00000183943.5 | 1207.742 | -1.657077 | 0.1212047 | -8.72143 | 2.75E-18 | 9.44E-17 | ENSG00000183943 | PRKX | 5613 |
| ENSG00000102053.12 | 33.27121 | -1.657218 | 0.285266 | -3.70608 | 0.00021 | 0.001784629 | ENSG00000102053 | ZC3H12B | 340554 |
| ENSG00000121068.14 | 19.29491844 | -1.657516301 | 0.298215098 | -3.54615 | 0.000391 | 0.003160196 | ENSG00000121068 | TBX2 | 6909 |
| ENSG00000118193.12 | 2575.163053 | -1.658175607 | 0.172500885 | -6.13432 | 8.55E-10 | 1.48E-08 | ENSG00000118193 | KIF14 | 9928 |
| ENSG00000198121.13 | 572.5776665 | -1.664741404 | 0.135945169 | -7.83214 | 4.80E-15 | 1.32E-13 | ENSG00000198121 | LPAR1 | 1902 |
| ENSG00000139734.18 | 2341.033011 | -1.665101635 | 0.157057366 | -6.78161 | 1.19E-11 | 2.47E-10 | ENSG00000139734 | DIAPH3 | 81624 |
| ENSG00000109805.10 | 3558.622923 | -1.66512071 | 0.186491474 | -5.71136 | 1.12E-08 | 1.72E-07 | ENSG00000109805 | NCAPG | 64151 |
| ENSG00000156398.13 | 316.582596 | -1.665342871 | 0.209296698 | -5.09011 | 3.58E-07 | 4.58E-06 | ENSG00000156398 | SFXN2 | 118980 |
| ENSG00000143476.18 | 1962.018196 | -1.669154609 | 0.252584688 | -4.23286 | 2.31E-05 | 0.000229896 | ENSG00000143476 | DTL | 51514 |
| ENSG00000227896.2 | 39.650308 | -1.669289237 | 0.217450569 | -4.91739 | 8.77E-07 | 1.07E-05 | ENSG00000227896 | NA | NA |
| ENSG00000221164.1 | 18.0079219 | -1.669548203 | 0.332165485 | -3.21993 | 0.001282 | 0.009273496 | ENSG00000221164 | SNORA11I | 1.1E+08 |

| | | | | | | | | | |
|---|---|---|---|---|---|---|---|---|---|
| ENSG00000131620.17 | 1591.549154 | -1.669813951 | 0.138181328 | -7.7421 | 9.78E-15 | 2.61E-13 | ENSG00000131620 | ANO1 | 55107 |
| ENSG00000166851.15 | 3009.678129 | -1.672289073 | 0.192839234 | -5.56053 | 2.69E-08 | 3.97E-07 | ENSG00000166851 | PLK1 | 5347 |
| ENSG00000257489.6 | 29.77679643 | -1.672676526 | 0.25567443 | -4.19548 | 2.72E-05 | 0.000268636 | ENSG00000257489 | NA | NA |
| ENSG00000111110.12 | 29.03561723 | -1.676381532 | 0.230476533 | -4.67024 | 3.01E-06 | 3.40E-05 | ENSG00000111110 | PPM1H | 57460 |
| ENSG00000168078.10 | 2667.784483 | -1.676646244 | 0.160003069 | -6.72891 | 1.71E-11 | 3.51E-10 | ENSG00000168078 | PBK | 55872 |
| ENSG00000164742.16 | 599.7607106 | -1.67761209 | 0.071843716 | -14.9994 | 7.41E-51 | 1.07E-48 | ENSG00000164742 | ADCY1 | 107 |
| ENSG00000160298.17 | 244.1248 | -1.679364 | 0.1755434 | -6.14871 | 7.81E-10 | 1.36E-08 | ENSG00000160298 | C21orf58 | 54058 |
| ENSG00000166073.10 | 224.0579088 | -1.679458123 | 0.107524085 | -10.0392 | 1.02E-23 | 4.84E-22 | ENSG00000166073 | GPR176 | 11245 |
| ENSG00000165905.18 | 622.7284678 | -1.67954093 | 0.09737355 | -11.0866 | 1.46E-28 | 9.19E-27 | ENSG00000165905 | LARGE2 | 120071 |
| ENSG00000171791.13 | 334.8064568 | -1.68073933 | 0.177420923 | -6.09139 | 1.12E-09 | 1.91E-08 | ENSG00000171791 | BCL2 | 596 |
| ENSG00000166803.13 | 777.1816508 | -1.682782659 | 0.177935187 | -6.08526 | 1.16E-09 | 1.98E-08 | ENSG00000166803 | PCLAF | 9768 |
| ENSG00000232233.1 | 22.19394949 | -1.683033948 | 0.325828742 | -3.32394 | 0.000888 | 0.006669138 | ENSG00000232233 | LINC0204: | 1.03E+08 |
| ENSG00000164651.16 | 25.30512252 | -1.683876214 | 0.307383703 | -3.52613 | 0.000422 | 0.003385338 | ENSG00000164651 | SP8 | 221833 |
| ENSG00000143498.18 | 603.1649787 | -1.68492749 | 0.190013303 | -5.70974 | 1.13E-08 | 1.74E-07 | ENSG00000143498 | TAF1A | 9015 |
| ENSG00000118596.12 | 272.3106192 | -1.688363187 | 0.21707663 | -5.01373 | 5.34E-07 | 6.68E-06 | ENSG00000118596 | SLC16A7 | 9194 |
| ENSG00000110427.16 | 178.1555946 | -1.689146044 | 0.146365318 | -7.44128 | 9.97E-14 | 2.49E-12 | ENSG00000110427 | KIAA1549l | 25758 |
| ENSG00000137135.18 | 360.1695658 | -1.689185783 | 0.158375367 | -6.87724 | 6.10E-12 | 1.31E-10 | ENSG00000137135 | ARHGEF3 | 84904 |
| ENSG00000105649.9 | 27.15431 | -1.689383 | 0.2748966 | -3.96288 | 7.41E-05 | 0.000683079 | ENSG00000105649 | RAB3A | 5864 |
| ENSG00000066279.18 | 5509.501738 | -1.689527411 | 0.155394399 | -7.01137 | 2.36E-12 | 5.22E-11 | ENSG00000066279 | ASPM | 259266 |
| ENSG00000176619.13 | 11411.50385 | -1.690488091 | 0.136758535 | -7.97382 | 1.54E-15 | 4.41E-14 | ENSG00000176619 | LMNB2 | 84823 |
| ENSG00000270890.1 | 23.20903788 | -1.69079938 | 0.320066981 | -3.40803 | 0.000654 | 0.005056447 | ENSG00000270890 | NA | NA |
| ENSG00000183856.11 | 1708.58696 | -1.69248499 | 0.171757181 | -6.36064 | 2.01E-10 | 3.75E-09 | ENSG00000183856 | IQGAP3 | 128239 |
| ENSG00000119969.15 | 2050.867672 | -1.693318995 | 0.190581693 | -5.73675 | 9.65E-09 | 1.49E-07 | ENSG00000119969 | HELLS | 3070 |
| ENSG00000151503.12 | 3487.196933 | -1.694150458 | 0.190773492 | -5.73534 | 9.73E-09 | 1.50E-07 | ENSG00000151503 | NCAPD3 | 23310 |
| ENSG00000142945.13 | 3167.892579 | -1.695096749 | 0.18697002 | -5.85707 | 4.71E-09 | 7.53E-08 | ENSG00000142945 | KIF2C | 11004 |
| ENSG00000155016.18 | 240.6593006 | -1.697983033 | 0.195483986 | -5.61674 | 1.95E-08 | 2.92E-07 | ENSG00000155016 | CYP2U1 | 113612 |
| ENSG00000197467.14 | 98.65996619 | -1.6986418 | 0.198430272 | -5.53666 | 3.08E-08 | 4.51E-07 | ENSG00000197467 | COL13A1 | 1305 |
| ENSG00000131153.9 | 1157.170355 | -1.69986881 | 0.221913056 | -4.95631 | 7.18E-07 | 8.87E-06 | ENSG00000131153 | GINS2 | 51659 |
| ENSG00000137310.12 | 308.5876515 | -1.700693785 | 0.180594051 | -6.09485 | 1.10E-09 | 1.88E-08 | ENSG00000137310 | TCF19 | 6941 |
| ENSG00000101447.15 | 1990.305 | -1.702324 | 0.1404833 | -7.84666 | 4.27E-15 | 1.18E-13 | ENSG00000101447 | FAM83D | 81610 |
| ENSG00000101347.9 | 2171.045 | -1.704393 | 0.106777 | -10.343 | 4.50E-25 | 2.30E-23 | ENSG00000101347 | SAMHD1 | 25939 |
| ENSG00000162063.13 | 1282.257596 | -1.704539037 | 0.144228521 | -7.65826 | 1.88E-14 | 4.94E-13 | ENSG00000162063 | CCNF | 899 |
| ENSG00000138160.6 | 4088.446215 | -1.706161355 | 0.176775136 | -6.25745 | 3.91E-10 | 7.07E-09 | ENSG00000138160 | KIF11 | 3832 |
| ENSG00000171790.15 | 19.13030732 | -1.706344064 | 0.298384373 | -3.70778 | 0.000209 | 0.001774372 | ENSG00000171790 | SLFNL1 | 200172 |
| ENSG00000111879.19 | 48.80352176 | -1.708657112 | 0.251738316 | -4.40401 | 1.06E-05 | 0.000111911 | ENSG00000111879 | FAM184A | 79632 |
| ENSG00000188610.12 | 235.3904436 | -1.70868486 | 0.209587843 | -5.28983 | 1.22E-07 | 1.67E-06 | ENSG00000188610 | FAM72B | 653820 |
| ENSG00000181544.15 | 273.5843 | -1.710362 | 0.198399 | -5.59661 | 2.19E-08 | 3.26E-07 | ENSG00000181544 | FANCB | 2187 |
| ENSG00000237649.8 | 1397.507016 | -1.713828092 | 0.180712544 | -6.16354 | 7.11E-10 | 1.25E-08 | ENSG00000237649 | KIFC1 | 3833 |
| ENSG00000215784.6 | 89.46030477 | -1.715950755 | 0.256813114 | -4.34538 | 1.39E-05 | 0.000143854 | ENSG00000215784 | FAM72D | 728833 |
| ENSG00000008853.16 | 953.3464096 | -1.718149949 | 0.1383867 | -8.07989 | 6.48E-16 | 1.90E-14 | ENSG00000008853 | RHOBTB2 | 23221 |
| ENSG00000180592.16 | 40.82823574 | -1.718678807 | 0.273935241 | -4.08373 | 4.43E-05 | 0.000423788 | ENSG00000180592 | SKIDA1 | 387640 |
| ENSG00000145147.20 | 751.4964498 | -1.719985236 | 0.254020667 | -4.40903 | 1.04E-05 | 0.00010957 | ENSG00000145147 | SLIT2 | 9353 |

| | | | | | | | | | |
|---|---|---|---|---|---|---|---|---|---|
| ENSG00000273373.1 | 87.56910722 | -1.721035814 | 0.213675097 | -5.24645 | 1.55E-07 | 2.09E-06 | ENSG00000273373 | NA | NA |
| ENSG00000260589.1 | 21.11401026 | -1.722319863 | 0.291734629 | -3.84706 | 0.00012 | 0.00106439 | ENSG00000260589 | STAM-AS1 | 1.03E+08 |
| ENSG00000134363.12 | 8724.197554 | -1.72241297 | 0.071892636 | -15.6123 | 6.00E-55 | 9.86E-53 | ENSG00000134363 | FST | 10468 |
| ENSG00000121152.10 | 2180.557523 | -1.722881272 | 0.199505458 | -5.62832 | 1.82E-08 | 2.73E-07 | ENSG00000121152 | NCAPH | 23397 |
| ENSG00000102967.12 | 329.5961707 | -1.723177534 | 0.218369349 | -5.14348 | 2.70E-07 | 3.51E-06 | ENSG00000102967 | DHODH | 1723 |
| ENSG00000186205.13 | 76.99664936 | -1.72401543 | 0.183204724 | -6.1353 | 8.50E-10 | 1.47E-08 | ENSG00000186205 | MTARC1 | 64757 |
| ENSG00000285128.1 | 59.66836 | -1.724071 | 0.233437 | -4.81531 | 1.47E-06 | 1.73E-05 | ENSG00000285128 | NA | NA |
| ENSG00000164045.11 | 1624.81933 | -1.724202176 | 0.182853233 | -6.14811 | 7.84E-10 | 1.37E-08 | ENSG00000164045 | CDC25A | 993 |
| ENSG00000183010.16 | 336.4280951 | -1.725264129 | 0.164512849 | -6.83998 | 7.92E-12 | 1.68E-10 | ENSG00000183010 | PYCR1 | 5831 |
| ENSG00000117724.13 | 7238.441843 | -1.727414245 | 0.182158261 | -6.1892 | 6.05E-10 | 1.07E-08 | ENSG00000117724 | CENPF | 1063 |
| ENSG00000149596.6 | 144.2223 | -1.728288 | 0.1742232 | -6.47611 | 9.41E-11 | 1.80E-09 | ENSG00000149596 | JPH2 | 57158 |
| ENSG00000268798.1 | 17.28353347 | -1.728496049 | 0.4214886 | -2.67741 | 0.007419 | 0.044247564 | ENSG00000268798 | NA | NA |
| ENSG00000262692.1 | 14.85825542 | -1.729692927 | 0.351615236 | -3.21287 | 0.001314 | 0.009483841 | ENSG00000262692 | NA | NA |
| ENSG00000172986.12 | 40.92614166 | -1.729751551 | 0.290462598 | -3.88949 | 0.0001 | 0.000906853 | ENSG00000172986 | GXYLT2 | 727936 |
| ENSG00000165490.13 | 645.7263125 | -1.735133995 | 0.197895114 | -5.73604 | 9.69E-09 | 1.50E-07 | ENSG00000165490 | DDIAS | 220042 |
| ENSG00000121236.21 | 99.39730282 | -1.742785647 | 0.170406357 | -6.70624 | 2.00E-11 | 4.08E-10 | ENSG00000121236 | TRIM6 | 117854 |
| ENSG00000143570.18 | 61.10822797 | -1.744507306 | 0.193602653 | -5.91163 | 3.39E-09 | 5.50E-08 | ENSG00000143570 | SLC39A1 | 27173 |
| ENSG00000134690.11 | 1578.359068 | -1.745564425 | 0.184184709 | -6.21965 | 4.98E-10 | 8.91E-09 | ENSG00000134690 | CDCA8 | 55143 |
| ENSG00000196550.10 | 121.6597446 | -1.746286289 | 0.258636901 | -4.43203 | 9.34E-06 | 9.96E-05 | ENSG00000196550 | FAM72A | 729533 |
| ENSG00000179431.6 | 1570.695613 | -1.746386253 | 0.156480482 | -7.32607 | 2.37E-13 | 5.73E-12 | ENSG00000179431 | FJX1 | 24147 |
| ENSG00000198826.11 | 949.2952964 | -1.747377637 | 0.178378677 | -6.43226 | 1.26E-10 | 2.38E-09 | ENSG00000198826 | ARHGAP1 | 9824 |
| ENSG00000184675.11 | 473.9652 | -1.747828 | 0.1639839 | -6.99964 | 2.57E-12 | 5.66E-11 | ENSG00000184675 | AMER1 | 139285 |
| ENSG00000075702.18 | 653.9706 | -1.750943 | 0.2100188 | -5.48019 | 4.25E-08 | 6.07E-07 | ENSG00000075702 | WDR62 | 284403 |
| ENSG00000171056.8 | 200.7738672 | -1.754485966 | 0.120265568 | -9.59947 | 8.04E-22 | 3.42E-20 | ENSG00000171056 | SOX7 | 83595 |
| ENSG00000112984.12 | 4967.098654 | -1.757725769 | 0.202255662 | -5.72407 | 1.04E-08 | 1.61E-07 | ENSG00000112984 | KIF20A | 10112 |
| ENSG00000162062.15 | 306.7327015 | -1.757916937 | 0.245522771 | -4.71613 | 2.40E-06 | 2.75E-05 | ENSG00000162062 | TEDC2 | 80178 |
| ENSG00000272853.1 | 11.18725178 | -1.758934202 | 0.434965504 | -2.66443 | 0.007712 | 0.045745831 | ENSG00000272853 | NA | NA |
| ENSG00000143799.13 | 8509.707603 | -1.761705232 | 0.152796405 | -7.60296 | 2.89E-14 | 7.47E-13 | ENSG00000143799 | PARP1 | 142 |
| ENSG00000152253.9 | 923.6528514 | -1.762932504 | 0.160282409 | -7.25552 | 4.00E-13 | 9.47E-12 | ENSG00000152253 | SPC25 | 57405 |
| ENSG00000176208.9 | 852.3485042 | -1.763270837 | 0.153472587 | -7.57967 | 3.46E-14 | 8.89E-13 | ENSG00000176208 | ATAD5 | 79915 |
| ENSG00000178999.13 | 1578.069104 | -1.766273459 | 0.188945701 | -6.17253 | 6.72E-10 | 1.18E-08 | ENSG00000178999 | AURKB | 9212 |
| ENSG00000073111.14 | 2242.893516 | -1.767499395 | 0.163218983 | -7.15296 | 8.49E-13 | 1.95E-11 | ENSG00000073111 | MCM2 | 4171 |
| ENSG00000228716.7 | 4413.60587 | -1.770198579 | 0.148303212 | -7.89058 | 3.01E-15 | 8.44E-14 | ENSG00000228716 | DHFR | 1719 |
| ENSG00000050344.9 | 1493.478039 | -1.772299282 | 0.057848292 | -20.2651 | 2.62E-91 | 9.29E-89 | ENSG00000050344 | NFE2L3 | 9603 |
| ENSG00000076382.17 | 2120.822473 | -1.773302017 | 0.187147238 | -6.26941 | 3.62E-10 | 6.59E-09 | ENSG00000076382 | SPAG5 | 10615 |
| ENSG00000122952.17 | 3290.111279 | -1.775185301 | 0.176390175 | -6.66242 | 2.69E-11 | 5.42E-10 | ENSG00000122952 | ZWINT | 11130 |
| ENSG00000177602.5 | 577.4902659 | -1.777086662 | 0.195198146 | -6.03021 | 1.64E-09 | 2.75E-08 | ENSG00000177602 | HASPIN | 83903 |
| ENSG00000135842.17 | 314.0609463 | -1.777468134 | 0.163783166 | -7.18919 | 6.52E-13 | 1.52E-11 | ENSG00000135842 | NIBAN1 | 116496 |
| ENSG00000145087.12 | 15.67003353 | -1.779407136 | 0.442888317 | -2.66299 | 0.007745 | 0.045919196 | ENSG00000145087 | STXBP5L | 9515 |
| ENSG00000123485.12 | 1892.551437 | -1.779790871 | 0.206914472 | -5.70183 | 1.19E-08 | 1.81E-07 | ENSG00000123485 | HJURP | 55355 |
| ENSG00000158089.15 | 151.1861524 | -1.780665806 | 0.242967565 | -4.85936 | 1.18E-06 | 1.41E-05 | ENSG00000158089 | GALNT14 | 79623 |
| ENSG00000165244.7 | 734.7966038 | -1.781671465 | 0.181875792 | -6.49713 | 8.19E-11 | 1.57E-09 | ENSG00000165244 | ZNF367 | 195828 |

| | | | | | | | | | |
|---|---|---|---|---|---|---|---|---|---|
| ENSG00000151651.16 | 1465.452434 | -1.782937506 | 0.080642959 | -14.6688 | 1.02E-48 | 1.38E-46 | ENSG00000151651 | ADAM8 | 101 |
| ENSG00000070814.20 | 16309.44443 | -1.785923655 | 0.156609986 | -7.57247 | 3.66E-14 | 9.35E-13 | ENSG00000070814 | TCOF1 | 6949 |
| ENSG00000250899.3 | 525.4397341 | -1.78632194 | 0.171225769 | -6.92841 | 4.26E-12 | 9.22E-11 | ENSG00000250899 | NA | NA |
| ENSG00000043355.12 | 472.5758011 | -1.789318465 | 0.212845879 | -5.5877 | 2.30E-08 | 3.42E-07 | ENSG00000043355 | ZIC2 | 7546 |
| ENSG00000169679.15 | 3765.522864 | -1.790036636 | 0.175544599 | -6.77911 | 1.21E-11 | 2.51E-10 | ENSG00000169679 | BUB1 | 699 |
| ENSG00000133119.13 | 2365.394205 | -1.793206691 | 0.211408947 | -5.64407 | 1.66E-08 | 2.50E-07 | ENSG00000133119 | RFC3 | 5983 |
| ENSG00000011426.11 | 10351.45032 | -1.795504493 | 0.159533766 | -7.49374 | 6.69E-14 | 1.68E-12 | ENSG00000011426 | ANLN | 54443 |
| ENSG00000152284.5 | 101.3865072 | -1.797586745 | 0.152345382 | -7.861 | 3.81E-15 | 1.06E-13 | ENSG00000152284 | TCF7L1 | 83439 |
| ENSG00000136492.8 | 1365.474009 | -1.799883972 | 0.168925415 | -7.10304 | 1.22E-12 | 2.77E-11 | ENSG00000136492 | BRIP1 | 83990 |
| ENSG00000173638.19 | 721.8133 | -1.800059 | 0.2047288 | -5.8617 | 4.58E-09 | 7.33E-08 | ENSG00000173638 | SLC19A1 | 6573 |
| ENSG00000102359.7 | 450.4489 | -1.800798 | 0.1746169 | -6.87676 | 6.12E-12 | 1.31E-10 | ENSG00000102359 | SRPX2 | 27286 |
| ENSG00000092853.14 | 2149.277781 | -1.801110698 | 0.207965426 | -5.77553 | 7.67E-09 | 1.20E-07 | ENSG00000092853 | CLSPN | 63967 |
| ENSG00000077264.15 | 40.22729 | -1.801456 | 0.2773497 | -4.33192 | 1.48E-05 | 0.000152055 | ENSG00000077264 | PAK3 | 5063 |
| ENSG00000168811.7 | 29.5801236 | -1.805105388 | 0.33282988 | -3.62078 | 0.000294 | 0.002429822 | ENSG00000168811 | IL12A | 3592 |
| ENSG00000118971.8 | 8237.251388 | -1.808773456 | 0.0853883 | -14.1562 | 1.71E-45 | 2.11E-43 | ENSG00000118971 | CCND2 | 894 |
| ENSG00000260552.1 | 11.40409145 | -1.810579201 | 0.429558049 | -2.8182 | 0.004829 | 0.030308519 | ENSG00000260552 | COSMOC | 1.02E+08 |
| ENSG00000187244.11 | 1918.065 | -1.812428 | 0.170959 | -7.09192 | 1.32E-12 | 3.00E-11 | ENSG00000187244 | BCAM | 4059 |
| ENSG00000089685.15 | 2527.799126 | -1.812790831 | 0.216203537 | -5.60949 | 2.03E-08 | 3.03E-07 | ENSG00000089685 | BIRC5 | 332 |
| ENSG00000156970.12 | 2572.584364 | -1.81285073 | 0.220685827 | -5.49583 | 3.89E-08 | 5.58E-07 | ENSG00000156970 | BUB1B | 701 |
| ENSG00000272620.1 | 3560.610492 | -1.814138623 | 0.161309509 | -7.52676 | 5.20E-14 | 1.32E-12 | ENSG00000272620 | AFAP1-AS | 84740 |
| ENSG00000204860.5 | 146.2032932 | -1.815722061 | 0.228414382 | -5.32244 | 1.02E-07 | 1.41E-06 | ENSG00000204860 | NA | NA |
| ENSG00000131747.15 | 14231.08666 | -1.815767651 | 0.162044595 | -7.50267 | 6.25E-14 | 1.58E-12 | ENSG00000131747 | TOP2A | 7153 |
| ENSG00000104432.14 | 107.2507187 | -1.816469472 | 0.175992688 | -6.91205 | 4.78E-12 | 1.03E-10 | ENSG00000104432 | IL7 | 3574 |
| ENSG00000242715.7 | 67.5150713 | -1.817796226 | 0.231700577 | -5.25591 | 1.47E-07 | 1.99E-06 | ENSG00000242715 | CCDC169 | 728591 |
| ENSG00000184661.14 | 1870.907293 | -1.820548364 | 0.175405554 | -6.95844 | 3.44E-12 | 7.51E-11 | ENSG00000184661 | CDCA2 | 157313 |
| ENSG00000151224.13 | 16.60609407 | -1.820572808 | 0.415241388 | -2.93943 | 0.003288 | 0.021545081 | ENSG00000151224 | MAT1A | 4143 |
| ENSG00000112218.9 | 325.2560221 | -1.82090382 | 0.260698327 | -4.68321 | 2.82E-06 | 3.21E-05 | ENSG00000112218 | GPR63 | 81491 |
| ENSG00000157168.20 | 3813.472319 | -1.822218066 | 0.206883156 | -5.90777 | 3.47E-09 | 5.63E-08 | ENSG00000157168 | NRG1 | 3084 |
| ENSG00000146281.6 | 1827.854612 | -1.826484681 | 0.084584864 | -14.5 | 1.21E-47 | 1.57E-45 | ENSG00000146281 | PM20D2 | 135293 |
| ENSG00000065675.14 | 38.15320979 | -1.832275278 | 0.27280599 | -4.51704 | 6.27E-06 | 6.85E-05 | ENSG00000065675 | PRKCQ | 5588 |
| ENSG00000064300.9 | 312.1570631 | -1.83322563 | 0.181410553 | -6.79798 | 1.06E-11 | 2.22E-10 | ENSG00000064300 | NGFR | 4804 |
| ENSG00000233122.2 | 20.19486231 | -1.834265015 | 0.294444765 | -4.19184 | 2.77E-05 | 0.000272425 | ENSG00000233122 | CTAGE7P | 119437 |
| ENSG00000165480.16 | 1718.339236 | -1.834308485 | 0.216520713 | -5.70065 | 1.19E-08 | 1.82E-07 | ENSG00000165480 | SKA3 | 221150 |
| ENSG00000225383.8 | 150.3561944 | -1.834359235 | 0.242672984 | -5.08651 | 3.65E-07 | 4.67E-06 | ENSG00000225383 | SFTA1P | 207107 |
| ENSG00000248774.1 | 13.25983583 | -1.83465185 | 0.439010906 | -2.81235 | 0.004918 | 0.030803585 | ENSG00000248774 | NA | NA |
| ENSG00000117399.14 | 5988.830906 | -1.836334544 | 0.197250054 | -6.26785 | 3.66E-10 | 6.64E-09 | ENSG00000117399 | CDC20 | 991 |
| ENSG00000137812.19 | 3008.625499 | -1.840895108 | 0.162110611 | -7.65462 | 1.94E-14 | 5.07E-13 | ENSG00000137812 | KNL1 | 57082 |
| ENSG00000287047.1 | 69.71330492 | -1.842767208 | 0.257604334 | -4.82433 | 1.40E-06 | 1.66E-05 | ENSG00000287047 | NA | NA |
| ENSG00000167513.9 | 952.4796109 | -1.843559146 | 0.148263612 | -8.38749 | 4.97E-17 | 1.57E-15 | ENSG00000167513 | CDT1 | 81620 |
| ENSG00000146670.10 | 2081.07828 | -1.847026274 | 0.187958866 | -6.63457 | 3.25E-11 | 6.48E-10 | ENSG00000146670 | CDCA5 | 113130 |
| ENSG00000261762.1 | 51.27547781 | -1.847647842 | 0.33375593 | -3.7382 | 0.000185 | 0.001589595 | ENSG00000261762 | NA | NA |
| ENSG00000101003.10 | 1768.971 | -1.850324 | 0.12851 | -9.72939 | 2.26E-22 | 9.84E-21 | ENSG00000101003 | GINS1 | 9837 |

| | | | | | | | | | |
|---|---|---|---|---|---|---|---|---|---|
| ENSG00000145386.10 | 3444.115393 | -1.850809378 | 0.183836031 | -6.80394 | 1.02E-11 | 2.14E-10 | ENSG00000145386 | CCNA2 | 890 |
| ENSG00000274849.1 | 42.86035443 | -1.851274524 | 0.232826584 | -5.37428 | 7.69E-08 | 1.07E-06 | ENSG00000274849 | NA | NA |
| ENSG00000105204.14 | 13.36675 | -1.8516 | 0.4397388 | -2.84624 | 0.004424 | 0.02810172 | ENSG00000105204 | DYRK1B | 9149 |
| ENSG00000111665.11 | 825.285622 | -1.852587294 | 0.186471504 | -6.71731 | 1.85E-11 | 3.79E-10 | ENSG00000111665 | CDCA3 | 83461 |
| ENSG00000109674.4 | 645.1184939 | -1.852860845 | 0.16294483 | -7.68887 | 1.48E-14 | 3.91E-13 | ENSG00000109674 | NEIL3 | 55247 |
| ENSG00000128602.11 | 123.3967578 | -1.853356081 | 0.157654441 | -7.95002 | 1.86E-15 | 5.29E-14 | ENSG00000128602 | SMO | 6608 |
| ENSG00000111206.12 | 2670.086474 | -1.853370096 | 0.205286666 | -6.10546 | 1.03E-09 | 1.76E-08 | ENSG00000111206 | FOXM1 | 2305 |
| ENSG00000197299.12 | 1005.630307 | -1.857895282 | 0.153069057 | -8.21783 | 2.07E-16 | 6.25E-15 | ENSG00000197299 | BLM | 641 |
| ENSG00000224999.1 | 166.51887 | -1.860730599 | 0.16020912 | -7.86928 | 3.57E-15 | 9.93E-14 | ENSG00000224999 | VTA1P1 | 1.06E+08 |
| ENSG00000212232.1 | 57.23505 | -1.861125 | 0.2188171 | -5.76338 | 8.24E-09 | 1.29E-07 | ENSG00000212232 | SNORD17 | 692086 |
| ENSG00000266235.1 | 17.46596386 | -1.863298746 | 0.35144057 | -3.59463 | 0.000325 | 0.002662636 | ENSG00000266235 | MIR3176 | 1E+08 |
| ENSG00000138587.6 | 134.0103245 | -1.864999328 | 0.162860342 | -7.76739 | 8.01E-15 | 2.16E-13 | ENSG00000138587 | MNS1 | 55329 |
| ENSG00000182263.14 | 694.6632916 | -1.868450089 | 0.115570534 | -10.9755 | 5.01E-28 | 3.05E-26 | ENSG00000182263 | FIGN | 55137 |
| ENSG00000002587.10 | 912.4502906 | -1.868984554 | 0.202308556 | -6.27252 | 3.55E-10 | 6.48E-09 | ENSG00000002587 | HS3ST1 | 9957 |
| ENSG00000180340.6 | 171.2339446 | -1.869200316 | 0.137143069 | -9.25457 | 2.15E-20 | 8.47E-19 | ENSG00000180340 | FZD2 | 2535 |
| ENSG00000103187.8 | 8044.3643 | -1.869219667 | 0.193510839 | -6.55891 | 5.42E-11 | 1.06E-09 | ENSG00000103187 | COTL1 | 23406 |
| ENSG00000260651.1 | 8.350080066 | -1.870790742 | 0.45814377 | -2.77378 | 0.005541 | 0.034217001 | ENSG00000260651 | NA | NA |
| ENSG00000132613.15 | 275.8114235 | -1.873950547 | 0.117817416 | -10.8129 | 2.99E-27 | 1.72E-25 | ENSG00000132613 | MTSS2 | 92154 |
| ENSG00000254433.1 | 10.36083043 | -1.874097997 | 0.400959496 | -3.17762 | 0.001485 | 0.010563986 | ENSG00000254433 | NA | NA |
| ENSG00000122966.16 | 1695.206693 | -1.87660564 | 0.174760277 | -7.3049 | 2.77E-13 | 6.64E-12 | ENSG00000122966 | CIT | 11113 |
| ENSG00000149591.16 | 516.6451017 | -1.877516001 | 0.220723137 | -5.78787 | 7.13E-09 | 1.12E-07 | ENSG00000149591 | TAGLN | 6876 |
| ENSG00000119408.16 | 879.1266562 | -1.878341463 | 0.174276155 | -7.33515 | 2.21E-13 | 5.38E-12 | ENSG00000119408 | NEK6 | 10783 |
| ENSG00000093009.9 | 1205.546 | -1.878476 | 0.2225032 | -5.74588 | 9.14E-09 | 1.42E-07 | ENSG00000093009 | CDC45 | 8318 |
| ENSG00000110723.12 | 230.5521629 | -1.878858379 | 0.125963827 | -10.1526 | 3.23E-24 | 1.56E-22 | ENSG00000110723 | EXPH5 | 23086 |
| ENSG00000169604.19 | 1903.270964 | -1.881139377 | 0.205523904 | -6.23353 | 4.56E-10 | 8.19E-09 | ENSG00000169604 | ANTXR1 | 84168 |
| ENSG00000185697.16 | 709.4939447 | -1.883755045 | 0.207350845 | -6.19122 | 5.97E-10 | 1.06E-08 | ENSG00000185697 | MYBL1 | 4603 |
| ENSG00000003989.17 | 487.5267376 | -1.883913398 | 0.252351396 | -5.0878 | 3.62E-07 | 4.64E-06 | ENSG00000003989 | SLC7A2 | 6542 |
| ENSG00000119888.10 | 3339.850967 | -1.890180749 | 0.140817012 | -9.16211 | 5.09E-20 | 1.94E-18 | ENSG00000119888 | EPCAM | 4072 |
| ENSG00000119547.6 | 244.8313835 | -1.894280429 | 0.204087929 | -6.34178 | 2.27E-10 | 4.22E-09 | ENSG00000119547 | ONECUT2 | 9480 |
| ENSG00000090447.12 | 720.5124838 | -1.897705904 | 0.167419775 | -7.75121 | 9.10E-15 | 2.44E-13 | ENSG00000090447 | TFAP4 | 7023 |
| ENSG00000167634.12 | 12.82062 | -1.89789 | 0.4668417 | -2.78015 | 0.005433 | 0.03367016 | ENSG00000167634 | NLRP7 | 199713 |
| ENSG00000278861.1 | 9.793768588 | -1.898494824 | 0.416355575 | -3.11872 | 0.001816 | 0.012665386 | ENSG00000278861 | NA | NA |
| ENSG00000186871.7 | 879.6744 | -1.899964 | 0.1962881 | -6.62273 | 3.53E-11 | 6.98E-10 | ENSG00000186871 | ERCC6L | 54821 |
| ENSG00000105011.9 | 1371.138953 | -1.902090872 | 0.182222793 | -7.1456 | 8.96E-13 | 2.06E-11 | ENSG00000105011 | ASF1B | 55723 |
| ENSG00000167889.12 | 27.91604988 | -1.902562224 | 0.355711413 | -3.66185 | 0.00025 | 0.002098908 | ENSG00000167889 | MGAT5B | 146664 |
| ENSG00000127564.17 | 1459.322116 | -1.904651035 | 0.147252234 | -8.85997 | 8.00E-19 | 2.84E-17 | ENSG00000127564 | PKMYT1 | 9088 |
| ENSG00000189057.11 | 1139.480544 | -1.904949873 | 0.179244045 | -7.2803 | 3.33E-13 | 7.93E-12 | ENSG00000189057 | FAM111B | 374393 |
| ENSG00000279965.1 | 15.22208221 | -1.908181207 | 0.466479329 | -2.80437 | 0.005041 | 0.031513111 | ENSG00000279965 | NA | NA |
| ENSG00000224566.2 | 155.0151164 | -1.91177075 | 0.249261277 | -5.26263 | 1.42E-07 | 1.92E-06 | ENSG00000224566 | NA | NA |
| ENSG00000161888.11 | 745.2790529 | -1.913355718 | 0.19257406 | -6.82 | 9.10E-12 | 1.92E-10 | ENSG00000161888 | SPC24 | 147841 |
| ENSG00000233085.5 | 146.2496094 | -1.918285626 | 0.136521933 | -9.65622 | 4.63E-22 | 1.99E-20 | ENSG00000233085 | NA | NA |
| ENSG00000055163.20 | 779.6445461 | -1.918497001 | 0.100638681 | -13.1013 | 3.24E-39 | 3.14E-37 | ENSG00000055163 | CYFIP2 | 26999 |

| | | | | | | | | | |
|---|---|---|---|---|---|---|---|---|---|
| ENSG00000081052.12 | 16.60174064 | -1.921470313 | 0.389542172 | -3.39237 | 0.000693 | 0.00533143 | ENSG00000081052 | COL4A4 | 1286 |
| ENSG00000118257.16 | 781.1054332 | -1.924256896 | 0.128292649 | -10.3222 | 5.60E-25 | 2.84E-23 | ENSG00000118257 | NRP2 | 8828 |
| ENSG00000276672.1 | 117.7586996 | -1.931631513 | 0.314836443 | -4.2296 | 2.34E-05 | 0.000232641 | ENSG00000276672 | NA | NA |
| ENSG00000171320.14 | 786.6656408 | -1.945084327 | 0.1667756 | -8.06523 | 7.31E-16 | 2.13E-14 | ENSG00000171320 | ESCO2 | 157570 |
| ENSG00000270392.2 | 23.77265612 | -1.946344502 | 0.394886635 | -3.40945 | 0.000651 | 0.005031825 | ENSG00000270392 | PFN1P2 | 767846 |
| ENSG00000196584.3 | 1386.176895 | -1.948952013 | 0.188513732 | -7.15572 | 8.32E-13 | 1.92E-11 | ENSG00000196584 | XRCC2 | 7516 |
| ENSG00000257167.2 | 116.6782923 | -1.951493882 | 0.243343343 | -5.55386 | 2.79E-08 | 4.12E-07 | ENSG00000257167 | TMPO-AS1 | 1E+08 |
| ENSG00000149256.15 | 16.82617873 | -1.961766837 | 0.383845989 | -3.54769 | 0.000389 | 0.003143712 | ENSG00000149256 | TENM4 | 26011 |
| ENSG00000163808.17 | 678.7316592 | -1.970352104 | 0.219706109 | -6.23721 | 4.45E-10 | 8.02E-09 | ENSG00000163808 | KIF15 | 56992 |
| ENSG00000047597.7 | 38.2208 | -1.972043 | 0.2578246 | -5.32162 | 1.03E-07 | 1.41E-06 | ENSG00000047597 | XK | 7504 |
| ENSG00000183287.14 | 235.3107211 | -1.979044521 | 0.175403616 | -7.86212 | 3.78E-15 | 1.05E-13 | ENSG00000183287 | CCBE1 | 147372 |
| ENSG00000136982.6 | 666.5842118 | -1.980026363 | 0.173422448 | -7.9576 | 1.75E-15 | 4.99E-14 | ENSG00000136982 | DSCC1 | 79075 |
| ENSG00000258793.1 | 17.63121837 | -1.981248193 | 0.407390407 | -3.39048 | 0.000698 | 0.005366773 | ENSG00000258793 | NA | NA |
| ENSG00000287575.1 | 6.972043034 | -1.981606758 | 0.50126376 | -2.75625 | 0.005847 | 0.035890174 | ENSG00000287575 | NA | NA |
| ENSG00000075218.19 | 1647.758 | -1.9819 | 0.191938 | -7.19972 | 6.03E-13 | 1.41E-11 | ENSG00000075218 | GTSE1 | 51512 |
| ENSG00000204176.13 | 13.00706078 | -1.982845198 | 0.431077423 | -3.20788 | 0.001337 | 0.009626299 | ENSG00000204176 | SYT15 | 83849 |
| ENSG00000134871.18 | 3150.546331 | -1.983391793 | 0.239265341 | -5.78183 | 7.39E-09 | 1.16E-07 | ENSG00000134871 | COL4A2 | 1284 |
| ENSG00000174371.17 | 707.6587501 | -1.9924865 | 0.253270151 | -5.49803 | 3.84E-08 | 5.51E-07 | ENSG00000174371 | EXO1 | 9156 |
| ENSG00000177842.12 | 126.5904926 | -1.997039647 | 0.142384118 | -9.81177 | 1.00E-22 | 4.45E-21 | ENSG00000177842 | ZNF620 | 253639 |
| ENSG00000065328.16 | 1558.565825 | -1.999352518 | 0.233020011 | -6.00529 | 1.91E-09 | 3.18E-08 | ENSG00000065328 | MCM10 | 55388 |
| ENSG00000023171.18 | 55.27125128 | -1.999423375 | 0.3872861 | -3.61341 | 0.000302 | 0.002495066 | ENSG00000023171 | GRAMD1B | 57476 |
| ENSG00000051341.14 | 1889.310686 | -2.00123729 | 0.212376655 | -6.59789 | 4.17E-11 | 8.22E-10 | ENSG00000051341 | POLQ | 10721 |
| ENSG00000230638.4 | 13.70766616 | -2.004614255 | 0.506072377 | -2.77552 | 0.005511 | 0.034066162 | ENSG00000230638 | LOC10041 | 1E+08 |
| ENSG00000129422.14 | 379.7584389 | -2.006227234 | 0.169509851 | -8.29584 | 1.08E-16 | 3.31E-15 | ENSG00000129422 | MTUS1 | 57509 |
| ENSG00000273374.1 | 10.84481862 | -2.006548866 | 0.47430401 | -2.9655 | 0.003022 | 0.020012239 | ENSG00000273374 | NA | NA |
| ENSG00000118513.19 | 201.8193371 | -2.007851522 | 0.130980481 | -10.7486 | 6.02E-27 | 3.38E-25 | ENSG00000118513 | MYB | 4602 |
| ENSG00000154556.18 | 205.6001194 | -2.014417876 | 0.179369121 | -7.88551 | 3.13E-15 | 8.77E-14 | ENSG00000154556 | SORBS2 | 8470 |
| ENSG00000247903.1 | 62.24877392 | -2.01802733 | 0.237576124 | -5.96873 | 2.39E-09 | 3.95E-08 | ENSG00000247903 | NA | NA |
| ENSG00000140807.7 | 51.60046513 | -2.018488535 | 0.202437928 | -7.00703 | 2.43E-12 | 5.38E-11 | ENSG00000140807 | NKD1 | 85407 |
| ENSG00000186340.15 | 65.80933905 | -2.019603476 | 0.196011151 | -7.24246 | 4.41E-13 | 1.04E-11 | ENSG00000186340 | THBS2 | 7058 |
| ENSG00000174607.11 | 246.8201967 | -2.022171729 | 0.170724267 | -8.33023 | 8.07E-17 | 2.51E-15 | ENSG00000174607 | UGT8 | 7368 |
| ENSG00000085999.13 | 605.3083845 | -2.031938101 | 0.213174241 | -6.71722 | 1.85E-11 | 3.79E-10 | ENSG00000085999 | RAD54L | 8438 |
| ENSG00000153132.13 | 13.70335879 | -2.033218504 | 0.366041839 | -3.91545 | 9.02E-05 | 0.000820723 | ENSG00000153132 | CLGN | 1047 |
| ENSG00000152078.10 | 330.5673683 | -2.036702648 | 0.092374815 | -15.553 | 1.52E-54 | 2.44E-52 | ENSG00000152078 | TLCD4 | 148534 |
| ENSG00000145604.15 | 1446.068383 | -2.038800842 | 0.176410293 | -8.15599 | 3.46E-16 | 1.03E-14 | ENSG00000145604 | SKP2 | 6502 |
| ENSG00000115318.12 | 58.05641473 | -2.040021096 | 0.191432327 | -7.52235 | 5.38E-14 | 1.36E-12 | ENSG00000115318 | LOXL3 | 84695 |
| ENSG00000100206.10 | 47.07556 | -2.04433 | 0.3090676 | -4.67319 | 2.97E-06 | 3.36E-05 | ENSG00000100206 | DMC1 | 11144 |
| ENSG00000151150.22 | 388.9697137 | -2.044534162 | 0.143030669 | -10.0995 | 5.55E-24 | 2.65E-22 | ENSG00000151150 | ANK3 | 288 |
| ENSG00000132623.16 | 377.5504 | -2.048915 | 0.1297989 | -11.1628 | 6.20E-29 | 4.00E-27 | ENSG00000132623 | ANKEF1 | 63926 |
| ENSG00000237943.7 | 136.4764414 | -2.051379582 | 0.205580081 | -7.05992 | 1.67E-12 | 3.74E-11 | ENSG00000237943 | PRKCQ-AS | 439949 |
| ENSG00000276043.5 | 3170.480081 | -2.05585975 | 0.214277693 | -6.79427 | 1.09E-11 | 2.27E-10 | ENSG00000276043 | UHRF1 | 29128 |
| ENSG00000164649.20 | 1401.842086 | -2.056224199 | 0.146811043 | -9.91904 | 3.44E-23 | 1.57E-21 | ENSG00000164649 | CDCA7L | 55536 |

| | | | | | | | | | |
|---|---|---|---|---|---|---|---|---|---|
| ENSG00000274356.1 | 8.887821689 | -2.057556586 | 0.501899492 | -2.90408 | 0.003683 | 0.023806189 | ENSG00000274356 | NA | NA |
| ENSG00000261534.1 | 201.7668386 | -2.057778174 | 0.127221692 | -11.4586 | 2.13E-30 | 1.49E-28 | ENSG00000261534 | NA | NA |
| ENSG00000237548.1 | 32.15051008 | -2.059826923 | 0.35584905 | -4.10238 | 4.09E-05 | 0.000393297 | ENSG00000237548 | NA | NA |
| ENSG00000135083.15 | 226.0297219 | -2.060175159 | 0.210695138 | -6.93027 | 4.20E-12 | 9.11E-11 | ENSG00000135083 | CCNJL | 79616 |
| ENSG00000196793.13 | 43.98645164 | -2.062029166 | 0.27921392 | -5.23623 | 1.64E-07 | 2.20E-06 | ENSG00000196793 | ZNF239 | 8187 |
| ENSG00000186185.13 | 1241.804103 | -2.063232002 | 0.196692925 | -7.43917 | 1.01E-13 | 2.52E-12 | ENSG00000186185 | KIF18B | 146909 |
| ENSG00000259330.2 | 458.3907171 | -2.066243587 | 0.09228131 | -15.8888 | 7.57E-57 | 1.32E-54 | ENSG00000259330 | INAFM2 | 1.01E+08 |
| ENSG00000111490.13 | 24.30890556 | -2.066634274 | 0.293658871 | -4.99435 | 5.90E-07 | 7.36E-06 | ENSG00000111490 | TBC1D30 | 23329 |
| ENSG00000049130.16 | 965.9621742 | -2.069200306 | 0.189135971 | -7.76796 | 7.98E-15 | 2.15E-13 | ENSG00000049130 | KITLG | 4254 |
| ENSG00000233056.2 | 270.3322 | -2.071884 | 0.1286482 | -11.4412 | 2.60E-30 | 1.81E-28 | ENSG00000233056 | ERVH48-1 | 90625 |
| ENSG00000276107.1 | 18.95374971 | -2.072344632 | 0.437638656 | -3.36429 | 0.000767 | 0.005847091 | ENSG00000276107 | NA | NA |
| ENSG00000196739.15 | 1307.226564 | -2.073080445 | 0.076930584 | -19.1482 | 1.00E-81 | 3.07E-79 | ENSG00000196739 | COL27A1 | 85301 |
| ENSG00000128714.6 | 22.46339463 | -2.074749879 | 0.35783289 | -4.12134 | 3.77E-05 | 0.000364786 | ENSG00000128714 | HOXD13 | 3239 |
| ENSG00000267500.1 | 37.79917042 | -2.079202085 | 0.224070062 | -6.60152 | 4.07E-11 | 8.03E-10 | ENSG00000267500 | NA | NA |
| ENSG00000171004.18 | 1007.695 | -2.086088 | 0.1304897 | -11.3885 | 4.77E-30 | 3.27E-28 | ENSG00000171004 | HS6ST2 | 90161 |
| ENSG00000186472.20 | 103.371476 | -2.090211754 | 0.185693653 | -8.02511 | 1.01E-15 | 2.94E-14 | ENSG00000186472 | PCLO | 27445 |
| ENSG00000132846.6 | 234.8189171 | -2.096255568 | 0.208425799 | -7.17884 | 7.03E-13 | 1.63E-11 | ENSG00000132846 | ZBED3 | 84327 |
| ENSG00000104332.12 | 199.3511529 | -2.097720497 | 0.337597956 | -4.4364 | 9.15E-06 | 9.77E-05 | ENSG00000104332 | SFRP1 | 6422 |
| ENSG00000152804.11 | 66.00142674 | -2.10004504 | 0.213156604 | -7.03729 | 1.96E-12 | 4.37E-11 | ENSG00000152804 | HHEX | 3087 |
| ENSG00000198133.8 | 595.6682723 | -2.101662553 | 0.076010748 | -19.7559 | 7.13E-87 | 2.34E-84 | ENSG00000198133 | TMEM229l | 161145 |
| ENSG00000171848.15 | 9148.758699 | -2.102765261 | 0.177479477 | -8.46726 | 2.51E-17 | 8.13E-16 | ENSG00000171848 | RRM2 | 6241 |
| ENSG00000258590.5 | 15.38803589 | -2.10383244 | 0.485196446 | -3.09943 | 0.001939 | 0.013421293 | ENSG00000258590 | NA | NA |
| ENSG00000115380.20 | 737.4691886 | -2.105029443 | 0.183068066 | -8.22115 | 2.02E-16 | 6.09E-15 | ENSG00000115380 | EFEMP1 | 2202 |
| ENSG00000231066.3 | 9.576749 | -2.105538 | 0.4344031 | -3.46576 | 0.000529 | 0.00416692 | ENSG00000231066 | NA | NA |
| ENSG00000106066.15 | 300.5071104 | -2.110213377 | 0.288259543 | -5.23908 | 1.61E-07 | 2.17E-06 | ENSG00000106066 | CPVL | 54504 |
| ENSG00000164236.12 | 884.9080949 | -2.112250667 | 0.13798726 | -10.9593 | 5.99E-28 | 3.60E-26 | ENSG00000164236 | ANKRD33l | 651746 |
| ENSG00000136010.14 | 400.0201175 | -2.116506718 | 0.184685527 | -8.21129 | 2.19E-16 | 6.59E-15 | ENSG00000136010 | ALDH1L2 | 160428 |
| ENSG00000157470.12 | 100.7223731 | -2.126115801 | 0.214094308 | -7.12824 | 1.02E-12 | 2.33E-11 | ENSG00000157470 | FAM81A | 145773 |
| ENSG00000227598.1 | 9.619055425 | -2.129053002 | 0.451465842 | -3.38686 | 0.000707 | 0.005433275 | ENSG00000227598 | NA | NA |
| ENSG00000246334.2 | 79.55201396 | -2.132251819 | 0.27678347 | -5.53592 | 3.10E-08 | 4.53E-07 | ENSG00000246334 | NA | NA |
| ENSG00000236824.2 | 62.49213645 | -2.133799755 | 0.263325253 | -5.82473 | 5.72E-09 | 9.08E-08 | ENSG00000236824 | BCYRN1 | 618 |
| ENSG00000128567.17 | 879.0469558 | -2.136239814 | 0.08415152 | -18.2556 | 1.87E-74 | 5.02E-72 | ENSG00000128567 | PODXL | 5420 |
| ENSG00000178919.8 | 194.2294355 | -2.139882294 | 0.196573811 | -7.83361 | 4.74E-15 | 1.30E-13 | ENSG00000178919 | FOXE1 | 2304 |
| ENSG00000152661.9 | 726.6995841 | -2.140387421 | 0.094098571 | -16.3699 | 3.14E-60 | 6.14E-58 | ENSG00000152661 | GJA1 | 2697 |
| ENSG00000157570.11 | 172.6920243 | -2.145784633 | 0.110208837 | -14.026 | 1.08E-44 | 1.29E-42 | ENSG00000157570 | TSPAN18 | 90139 |
| ENSG00000154930.15 | 552.9229 | -2.14593 | 0.1845317 | -8.37759 | 5.40E-17 | 1.70E-15 | ENSG00000154930 | ACSS1 | 84532 |
| ENSG00000146955.10 | 19.10907709 | -2.16358795 | 0.407746213 | -3.83471 | 0.000126 | 0.001114836 | ENSG00000146955 | RAB19 | 401409 |
| ENSG00000250855.1 | 12.69758433 | -2.165039785 | 0.441046859 | -3.54847 | 0.000387 | 0.003135424 | ENSG00000250855 | NA | NA |
| ENSG00000287895.1 | 206.3678115 | -2.167192358 | 0.234435841 | -6.68495 | 2.31E-11 | 4.68E-10 | ENSG00000287895 | NA | NA |
| ENSG00000101057.16 | 2546.683 | -2.167911 | 0.2084059 | -7.52335 | 5.34E-14 | 1.35E-12 | ENSG00000101057 | MYBL2 | 4605 |
| ENSG00000248663.6 | 8.520375263 | -2.169113018 | 0.475392523 | -3.30067 | 0.000965 | 0.007174689 | ENSG00000248663 | NA | NA |
| ENSG00000134802.17 | 2068.477558 | -2.179456903 | 0.26084086 | -6.05525 | 1.40E-09 | 2.37E-08 | ENSG00000134802 | SLC43A3 | 29015 |

| | | | | | | | | | |
|---|---|---|---|---|---|---|---|---|---|
| ENSG00000187140.5 | 9.592927465 | -2.179934229 | 0.528451722 | -2.98974 | 0.002792 | 0.018657673 | ENSG00000187140 | FOXD3 | 27022 |
| ENSG00000214336.4 | 39.8398124 | -2.180054034 | 0.352561665 | -4.48164 | 7.41E-06 | 8.02E-05 | ENSG00000214336 | FOXI3 | 344167 |
| ENSG00000278023.6 | 37.21449578 | -2.18210332 | 0.264053705 | -5.9916 | 2.08E-09 | 3.45E-08 | ENSG00000278023 | RDM1 | 201299 |
| ENSG00000227329.2 | 9.638451 | -2.183368 | 0.4875018 | -3.24792 | 0.001163 | 0.008488294 | ENSG00000227329 | NA | NA |
| ENSG00000137801.10 | 14702.35272 | -2.184660542 | 0.202247606 | -7.83525 | 4.68E-15 | 1.29E-13 | ENSG00000137801 | THBS1 | 7057 |
| ENSG00000157992.12 | 317.4528303 | -2.186780019 | 0.137059759 | -11.5773 | 5.37E-31 | 3.85E-29 | ENSG00000157992 | KRTCAP3 | 200634 |
| ENSG00000100867.14 | 36.25746477 | -2.187067851 | 0.363513195 | -4.36592 | 1.27E-05 | 0.000131903 | ENSG00000100867 | DHRS2 | 10202 |
| ENSG00000170298.15 | 30.91569316 | -2.187354686 | 0.408063986 | -3.88997 | 0.0001 | 0.000905388 | ENSG00000170298 | LGALS9B | 284194 |
| ENSG00000165685.8 | 14.59529932 | -2.198597408 | 0.424266226 | -3.76791 | 0.000165 | 0.001427988 | ENSG00000165685 | TMEM52B | 120939 |
| ENSG00000279347.1 | 37.99971619 | -2.199996147 | 0.466914181 | -3.42675 | 0.000611 | 0.004749515 | ENSG00000279347 | NA | NA |
| ENSG00000224629.1 | 6.169546895 | -2.201943478 | 0.559794148 | -2.86167 | 0.004214 | 0.026897839 | ENSG00000224629 | NA | NA |
| ENSG00000129007.14 | 18.20418869 | -2.206594402 | 0.513834208 | -3.12668 | 0.001768 | 0.012382495 | ENSG00000129007 | CALML4 | 91860 |
| ENSG00000129173.13 | 466.1890602 | -2.207246958 | 0.109145315 | -14.7258 | 4.41E-49 | 6.05E-47 | ENSG00000129173 | E2F8 | 79733 |
| ENSG00000229586.2 | 7.729289408 | -2.213447915 | 0.538873035 | -2.99412 | 0.002752 | 0.018429311 | ENSG00000229586 | TNPO1P3 | 339257 |
| ENSG00000148773.14 | 20710.5115 | -2.214645205 | 0.226839156 | -7.11802 | 1.09E-12 | 2.50E-11 | ENSG00000148773 | MKI67 | 4288 |
| ENSG00000117318.9 | 344.9574043 | -2.216726822 | 0.246343897 | -6.56289 | 5.28E-11 | 1.03E-09 | ENSG00000117318 | ID3 | 3399 |
| ENSG00000285730.1 | 25.6047667 | -2.217684986 | 0.336357055 | -4.80943 | 1.51E-06 | 1.78E-05 | ENSG00000285730 | NA | NA |
| ENSG00000227512.1 | 14.01078125 | -2.217771177 | 0.480397924 | -3.36756 | 0.000758 | 0.005788085 | ENSG00000227512 | NA | NA |
| ENSG00000285904.1 | 29.21630147 | -2.222401302 | 0.459495079 | -3.53083 | 0.000414 | 0.003331772 | ENSG00000285904 | NA | NA |
| ENSG00000077274.9 | 8.903682 | -2.224876 | 0.5775455 | -2.81342 | 0.004902 | 0.0307338 | ENSG00000077274 | CAPN6 | 827 |
| ENSG00000135476.11 | 1661.897032 | -2.231034796 | 0.188564859 | -8.64973 | 5.16E-18 | 1.75E-16 | ENSG00000135476 | ESPL1 | 9700 |
| ENSG00000185760.15 | 374.3173193 | -2.23192201 | 0.163036615 | -10.0095 | 1.38E-23 | 6.46E-22 | ENSG00000185760 | KCNQ5 | 56479 |
| ENSG00000153898.13 | 129.8355702 | -2.233293026 | 0.269485996 | -6.06077 | 1.35E-09 | 2.29E-08 | ENSG00000153898 | MCOLN2 | 255231 |
| ENSG00000111664.10 | 16.38270322 | -2.238375782 | 0.533641141 | -3.07018 | 0.002139 | 0.01468589 | ENSG00000111664 | GNB3 | 2784 |
| ENSG00000163739.4 | 5688.328116 | -2.244045061 | 0.107788927 | -15.2524 | 1.59E-52 | 2.40E-50 | ENSG00000163739 | CXCL1 | 2919 |
| ENSG00000114790.13 | 18.27799651 | -2.248384351 | 0.375435275 | -4.3906 | 1.13E-05 | 0.000118666 | ENSG00000114790 | ARHGEF2 | 26084 |
| ENSG00000162711.17 | 245.8281935 | -2.251105518 | 0.14877501 | -11.098 | 1.28E-28 | 8.13E-27 | ENSG00000162711 | NLRP3 | 114548 |
| ENSG00000101230.6 | 23.34431 | -2.253254 | 0.3516184 | -4.70184 | 2.58E-06 | 2.94E-05 | ENSG00000101230 | ISM1 | 140862 |
| ENSG00000229154.2 | 8.497107367 | -2.25362622 | 0.588064671 | -2.81198 | 0.004924 | 0.030827883 | ENSG00000229154 | KCNQ5-AS | 1.01E+08 |
| ENSG00000103044.11 | 2771.073643 | -2.254552461 | 0.263895385 | -6.26973 | 3.62E-10 | 6.58E-09 | ENSG00000103044 | HAS3 | 3038 |
| ENSG00000168502.17 | 895.2095224 | -2.259876428 | 0.150149101 | -11.0549 | 2.08E-28 | 1.29E-26 | ENSG00000168502 | MTCL1 | 23255 |
| ENSG00000162654.9 | 652.3333623 | -2.264813891 | 0.113444425 | -14.6751 | 9.30E-49 | 1.26E-46 | ENSG00000162654 | GBP4 | 115361 |
| ENSG00000202533.1 | 4.978514375 | -2.269188948 | 0.634190852 | -2.632 | 0.008488 | 0.049676689 | ENSG00000202533 | NA | NA |
| ENSG00000231381.2 | 6.022970432 | -2.269767113 | 0.607697898 | -2.74769 | 0.006002 | 0.036674637 | ENSG00000231381 | NA | NA |
| ENSG00000165029.16 | 669.4269947 | -2.27466652 | 0.180999451 | -9.25233 | 2.20E-20 | 8.62E-19 | ENSG00000165029 | ABCA1 | 19 |
| ENSG00000101871.14 | 502.6074 | -2.276596 | 0.1330989 | -12.5966 | 2.20E-36 | 1.92E-34 | ENSG00000101871 | MID1 | 4281 |
| ENSG00000113368.12 | 7334.884837 | -2.283850121 | 0.209895966 | -8.02231 | 1.04E-15 | 3.00E-14 | ENSG00000113368 | LMNB1 | 4001 |
| ENSG00000043591.5 | 9.114748131 | -2.28583345 | 0.4804853 | -3.50861 | 0.00045 | 0.003594786 | ENSG00000043591 | ADRB1 | 153 |
| ENSG00000213160.10 | 530.3697877 | -2.290150604 | 0.164385139 | -10.2817 | 8.53E-25 | 4.27E-23 | ENSG00000213160 | KLHL23 | 151230 |
| ENSG00000279409.1 | 7.942703434 | -2.292381455 | 0.509952538 | -3.3187 | 0.000904 | 0.006781957 | ENSG00000279409 | NA | NA |
| ENSG00000265763.4 | 80.57659341 | -2.293691649 | 0.243843954 | -6.9458 | 3.76E-12 | 8.18E-11 | ENSG00000265763 | ZNF488 | 118738 |
| ENSG00000274024.1 | 7.153283 | -2.297333 | 0.5258895 | -3.22755 | 0.001249 | 0.009049654 | ENSG00000274024 | NA | NA |

| | | | | | | | | | |
|---|---|---|---|---|---|---|---|---|---|
| ENSG00000229677.1 | 18.85054644 | -2.300025524 | 0.359560936 | -4.72806 | 2.27E-06 | 2.61E-05 | ENSG00000229677 | RPL17P28 | 1E+08 |
| ENSG00000270933.1 | 10.55084933 | -2.301789152 | 0.480920022 | -3.53861 | 0.000402 | 0.00324201 | ENSG00000270933 | NA | NA |
| ENSG00000144354.14 | 1959.12594 | -2.303757339 | 0.242598448 | -7.02295 | 2.17E-12 | 4.83E-11 | ENSG00000144354 | CDCA7 | 83879 |
| ENSG00000218358.2 | 12.92143342 | -2.306982298 | 0.515253841 | -3.3129 | 0.000923 | 0.006897172 | ENSG00000218358 | RAET1K | 646024 |
| ENSG00000256663.1 | 62.18498088 | -2.307007375 | 0.188521767 | -9.0547 | 1.37E-19 | 5.11E-18 | ENSG00000256663 | NA | NA |
| ENSG00000004139.14 | 13.6421351 | -2.307089195 | 0.386303841 | -4.41903 | 9.91E-06 | 0.000105114 | ENSG00000004139 | SARM1 | 23098 |
| ENSG00000148143.13 | 860.8926834 | -2.313346835 | 0.159479826 | -10.7433 | 6.37E-27 | 3.56E-25 | ENSG00000148143 | ZNF462 | 58499 |
| ENSG00000068831.19 | 7.246985868 | -2.319128727 | 0.58663135 | -2.93051 | 0.003384 | 0.022113556 | ENSG00000068831 | RASGRP2 | 10235 |
| ENSG00000100350.14 | 399.884 | -2.333037 | 0.1491052 | -11.6229 | 3.15E-31 | 2.29E-29 | ENSG00000100350 | FOXRED2 | 80020 |
| ENSG00000273706.4 | 498.4693454 | -2.336841296 | 0.168410606 | -10.3131 | 6.15E-25 | 3.09E-23 | ENSG00000273706 | LHX1 | 3975 |
| ENSG00000078081.8 | 124.5198637 | -2.343535487 | 0.358430616 | -4.86436 | 1.15E-06 | 1.38E-05 | ENSG00000078081 | LAMP3 | 27074 |
| ENSG00000134775.15 | 401.2507811 | -2.346302 | 0.187694522 | -9.30396 | 1.35E-20 | 5.41E-19 | ENSG00000134775 | FHOD3 | 80206 |
| ENSG00000273064.1 | 5.331361338 | -2.357939911 | 0.641016524 | -2.74243 | 0.006099 | 0.037195982 | ENSG00000273064 | NA | NA |
| ENSG00000285831.1 | 177.7583114 | -2.358085843 | 0.170382985 | -10.3184 | 5.82E-25 | 2.94E-23 | ENSG00000285831 | NA | NA |
| ENSG00000185483.12 | 52.46166261 | -2.364063978 | 0.370474841 | -4.76163 | 1.92E-06 | 2.23E-05 | ENSG00000185483 | ROR1 | 4919 |
| ENSG00000181418.8 | 38.86461238 | -2.37105173 | 0.377504089 | -4.69148 | 2.71E-06 | 3.09E-05 | ENSG00000181418 | DDN | 23109 |
| ENSG00000140479.17 | 439.4058469 | -2.375827214 | 0.123542722 | -14.3742 | 7.51E-47 | 9.53E-45 | ENSG00000140479 | PCSK6 | 5046 |
| ENSG00000254535.4 | 22.66506939 | -2.379608931 | 0.392336391 | -4.53593 | 5.74E-06 | 6.29E-05 | ENSG00000254535 | PABPC4L | 132430 |
| ENSG00000095303.17 | 314.7056485 | -2.387306106 | 0.123834447 | -14.433 | 3.21E-47 | 4.12E-45 | ENSG00000095303 | PTGS1 | 5742 |
| ENSG00000187123.15 | 59.96994206 | -2.387408635 | 0.218429333 | -8.18301 | 2.77E-16 | 8.26E-15 | ENSG00000187123 | LYPD6 | 130574 |
| ENSG00000261668.1 | 6.414566214 | -2.400933318 | 0.593997646 | -3.03189 | 0.00243 | 0.016483985 | ENSG00000261668 | NA | NA |
| ENSG00000255176.1 | 9.067688794 | -2.408569045 | 0.619532828 | -2.91925 | 0.003509 | 0.022827691 | ENSG00000255176 | NA | NA |
| ENSG00000164061.5 | 8.685747902 | -2.419512729 | 0.558700665 | -3.25669 | 0.001127 | 0.008266826 | ENSG00000164061 | BSN | 8927 |
| ENSG00000100625.9 | 536.4824424 | -2.43272721 | 0.11154816 | -16.4299 | 1.17E-60 | 2.34E-58 | ENSG00000100625 | SIX4 | 51804 |
| ENSG00000008277.14 | 119.0810231 | -2.439238285 | 0.212433878 | -8.65793 | 4.80E-18 | 1.63E-16 | ENSG00000008277 | ADAM22 | 53616 |
| ENSG00000133561.15 | 8.092437358 | -2.443456981 | 0.55756782 | -3.30625 | 0.000946 | 0.007051077 | ENSG00000133561 | GIMAP6 | 474344 |
| ENSG00000185070.11 | 779.0539101 | -2.443914085 | 0.261830424 | -7.0424 | 1.89E-12 | 4.22E-11 | ENSG00000185070 | FLRT2 | 23768 |
| ENSG00000228459.4 | 14.39472 | -2.445673 | 0.6967017 | -2.64916 | 0.008069 | 0.04755343 | ENSG00000228459 | NA | NA |
| ENSG00000079215.14 | 839.8411931 | -2.458439085 | 0.240588974 | -7.72454 | 1.12E-14 | 2.99E-13 | ENSG00000079215 | SLC1A3 | 6507 |
| ENSG00000280027.1 | 53.47017334 | -2.463474521 | 0.230871347 | -8.07148 | 6.94E-16 | 2.03E-14 | ENSG00000280027 | NA | NA |
| ENSG00000178445.9 | 1598.5846 | -2.469305036 | 0.148123144 | -12.6199 | 1.64E-36 | 1.44E-34 | ENSG00000178445 | GLDC | 2731 |
| ENSG00000240303.8 | 4.616921162 | -2.475888934 | 0.694867978 | -2.69963 | 0.006942 | 0.041724759 | ENSG00000240303 | ACAD11 | 84129 |
| ENSG00000135069.14 | 1717.655858 | -2.477668031 | 0.111689198 | -16.8115 | 2.01E-63 | 4.15E-61 | ENSG00000135069 | PSAT1 | 29968 |
| ENSG00000229953.1 | 12.07970184 | -2.484277304 | 0.439560547 | -4.28673 | 1.81E-05 | 0.000183892 | ENSG00000229953 | NA | NA |
| ENSG00000075340.23 | 50.47609622 | -2.486596378 | 0.249798251 | -7.55248 | 4.27E-14 | 1.08E-12 | ENSG00000075340 | ADD2 | 119 |
| ENSG00000229057.1 | 6.568455705 | -2.486611796 | 0.595886487 | -3.16606 | 0.001545 | 0.010940469 | ENSG00000229057 | NA | NA |
| ENSG00000137834.15 | 30.33506733 | -2.494427463 | 0.554127584 | -3.41876 | 0.000629 | 0.004875501 | ENSG00000137834 | SMAD6 | 4091 |
| ENSG00000150556.17 | 26.49261404 | -2.498122157 | 0.328961799 | -5.77004 | 7.93E-09 | 1.24E-07 | ENSG00000150556 | LYPD6B | 130576 |
| ENSG00000104369.5 | 482.4768921 | -2.510919599 | 0.190816041 | -10.0145 | 1.32E-23 | 6.18E-22 | ENSG00000104369 | JPH1 | 56704 |
| ENSG00000231982.1 | 15.64311733 | -2.512105164 | 0.380144443 | -5.02994 | 4.91E-07 | 6.17E-06 | ENSG00000231982 | NA | NA |
| ENSG00000079931.15 | 19.34818457 | -2.520680665 | 0.3565259 | -5.38721 | 7.16E-08 | 1.00E-06 | ENSG00000079931 | MOXD1 | 26002 |
| ENSG00000235427.1 | 7.974026023 | -2.528517601 | 0.693182105 | -2.78212 | 0.0054 | 0.033489667 | ENSG00000235427 | NA | NA |

| | | | | | | | | | |
|---|---|---|---|---|---|---|---|---|---|
| ENSG00000106484.15 | 144.4834016 | -2.530530144 | 0.141037416 | -13.6881 | 1.20E-42 | 1.33E-40 | ENSG00000106484 | MEST | 4232 |
| ENSG00000165887.11 | 112.1620957 | -2.531490798 | 0.171384184 | -11.2699 | 1.85E-29 | 1.24E-27 | ENSG00000165887 | ANKRD2 | 26287 |
| ENSG00000154655.15 | 51.66375364 | -2.532401099 | 0.225385628 | -8.57375 | 1.00E-17 | 3.32E-16 | ENSG00000154655 | L3MBTL4 | 91133 |
| ENSG00000216775.3 | 28.6463882 | -2.542348012 | 0.326053144 | -5.95715 | 2.57E-09 | 4.24E-08 | ENSG00000216775 | LOC73010 | 730101 |
| ENSG00000216867.2 | 5.59669258 | -2.543575835 | 0.608251023 | -3.19535 | 0.001397 | 0.010000121 | ENSG00000216867 | NA | NA |
| ENSG00000262714.1 | 98.15607022 | -2.544340792 | 0.230560993 | -8.43309 | 3.37E-17 | 1.08E-15 | ENSG00000262714 | NA | NA |
| ENSG00000198788.8 | 135.1210455 | -2.545682301 | 0.225455974 | -8.62999 | 6.14E-18 | 2.07E-16 | ENSG00000198788 | MUC2 | 4583 |
| ENSG00000066468.22 | 101.7548717 | -2.561266743 | 0.310198824 | -6.32261 | 2.57E-10 | 4.75E-09 | ENSG00000066468 | FGFR2 | 2263 |
| ENSG00000253500.6 | 83.9048639 | -2.563844537 | 0.297647784 | -6.59788 | 4.17E-11 | 8.22E-10 | ENSG00000253500 | NA | NA |
| ENSG00000134569.10 | 322.2736909 | -2.565103626 | 0.200714587 | -9.79054 | 1.24E-22 | 5.46E-21 | ENSG00000134569 | LRP4 | 4038 |
| ENSG00000138030.13 | 46.66210865 | -2.580197884 | 0.300525063 | -6.58913 | 4.42E-11 | 8.69E-10 | ENSG00000138030 | KHK | 3795 |
| ENSG00000114757.19 | 50.0416271 | -2.584973122 | 0.294371979 | -6.74308 | 1.55E-11 | 3.20E-10 | ENSG00000114757 | PEX5L | 51555 |
| ENSG00000134198.10 | 30.09814535 | -2.589340038 | 0.28322255 | -7.02395 | 2.16E-12 | 4.80E-11 | ENSG00000134198 | TSPAN2 | 10100 |
| ENSG00000007968.7 | 35.42081924 | -2.600942871 | 0.350125574 | -5.71493 | 1.10E-08 | 1.69E-07 | ENSG00000007968 | E2F2 | 1870 |
| ENSG00000230088.1 | 12.81967478 | -2.601950537 | 0.472415011 | -4.23769 | 2.26E-05 | 0.00022542 | ENSG00000230088 | KRT16P5 | 353196 |
| ENSG00000149328.14 | 7.947265269 | -2.606131907 | 0.603785535 | -3.32259 | 0.000892 | 0.006697617 | ENSG00000149328 | GLB1L2 | 89944 |
| ENSG00000272189.1 | 30.53019995 | -2.609615562 | 0.494473205 | -4.06415 | 4.82E-05 | 0.00045769 | ENSG00000272189 | NA | NA |
| ENSG00000154319.16 | 65.11116247 | -2.619565301 | 0.226437767 | -8.91885 | 4.71E-19 | 1.70E-17 | ENSG00000154319 | FAM167A | 83648 |
| ENSG00000112394.17 | 23.46174975 | -2.625089814 | 0.331516923 | -6.10856 | 1.01E-09 | 1.73E-08 | ENSG00000112394 | SLC16A10 | 117247 |
| ENSG00000261804.2 | 113.3011635 | -2.626618091 | 0.173278899 | -11.6957 | 1.34E-31 | 9.91E-30 | ENSG00000261804 | NA | NA |
| ENSG00000164684.13 | 186.9688543 | -2.637172441 | 0.244289862 | -8.33916 | 7.48E-17 | 2.33E-15 | ENSG00000164684 | ZNF704 | 619279 |
| ENSG00000186193.9 | 737.581261 | -2.652521434 | 0.092386415 | -22.2167 | 2.37E-109 | 1.13E-106 | ENSG00000186193 | SAPCD2 | 89958 |
| ENSG00000251563.1 | 6.249076251 | -2.656004516 | 0.608401882 | -3.37935 | 0.000727 | 0.005567841 | ENSG00000251563 | IARS2P1 | 642461 |
| ENSG00000258412.1 | 7.891252541 | -2.660233771 | 0.554715124 | -3.71404 | 0.000204 | 0.001736578 | ENSG00000258412 | NA | NA |
| ENSG00000060982.15 | 1918.808245 | -2.662118125 | 0.153611064 | -13.4243 | 4.36E-41 | 4.53E-39 | ENSG00000060982 | BCAT1 | 586 |
| ENSG00000204767.4 | 191.5359898 | -2.668540642 | 0.154354977 | -13.4012 | 5.95E-41 | 6.14E-39 | ENSG00000204767 | INSYN2B | 1E+08 |
| ENSG00000229544.9 | 157.586256 | -2.684646048 | 0.183286299 | -11.3737 | 5.65E-30 | 3.87E-28 | ENSG00000229544 | NKX1-2 | 390010 |
| ENSG00000249084.1 | 6.216937372 | -2.704734552 | 0.673935479 | -3.12305 | 0.00179 | 0.012516397 | ENSG00000249084 | NA | NA |
| ENSG00000171916.16 | 21.58137253 | -2.715348296 | 0.40717454 | -5.19519 | 2.05E-07 | 2.70E-06 | ENSG00000171916 | LGALS9C | 654346 |
| ENSG00000261115.6 | 6.428405655 | -2.733752675 | 0.638969147 | -3.33937 | 0.00084 | 0.00634526 | ENSG00000261115 | TMEM178I | 1.01E+08 |
| ENSG00000076716.9 | 43.20012 | -2.741332 | 0.2681362 | -7.98599 | 1.39E-15 | 4.01E-14 | ENSG00000076716 | GPC4 | 2239 |
| ENSG00000144810.16 | 285.0068972 | -2.750090671 | 0.207385683 | -10.3676 | 3.48E-25 | 1.79E-23 | ENSG00000144810 | COL8A1 | 1295 |
| ENSG00000197415.12 | 16.86484033 | -2.759976023 | 0.412476856 | -5.2366 | 1.64E-07 | 2.19E-06 | ENSG00000197415 | VEPH1 | 79674 |
| ENSG00000212135.1 | 5.493690124 | -2.769250971 | 0.697415164 | -3.11042 | 0.001868 | 0.012989388 | ENSG00000212135 | SNORD67 | 692108 |
| ENSG00000226887.8 | 270.0091867 | -2.782981143 | 0.244569946 | -8.92579 | 4.43E-19 | 1.60E-17 | ENSG00000226887 | ERVMER3 | 1E+08 |
| ENSG00000189068.11 | 5.55545 | -2.805535 | 0.7764633 | -2.84049 | 0.004504 | 0.02853135 | ENSG00000189068 | VSTM1 | 284415 |
| ENSG00000166402.8 | 447.1309573 | -2.823381618 | 0.187453424 | -11.861 | 1.89E-32 | 1.44E-30 | ENSG00000166402 | TUB | 7275 |
| ENSG00000260765.2 | 16.27383677 | -2.829181886 | 0.547707589 | -4.07002 | 4.70E-05 | 0.000447112 | ENSG00000260765 | NA | NA |
| ENSG00000165238.16 | 41.63303007 | -2.886852488 | 0.302111794 | -7.56956 | 3.74E-14 | 9.54E-13 | ENSG00000165238 | WNK2 | 65268 |
| ENSG00000196188.11 | 8.656688232 | -2.894353673 | 0.596851716 | -3.84409 | 0.000121 | 0.001076195 | ENSG00000196188 | CTSE | 1510 |
| ENSG00000279741.1 | 20.84511802 | -2.897592973 | 0.377204841 | -6.0911 | 1.12E-09 | 1.92E-08 | ENSG00000279741 | NA | NA |
| ENSG00000163923.10 | 8.761559705 | -2.904130999 | 0.56325418 | -4.09075 | 4.30E-05 | 0.000411764 | ENSG00000163923 | RPL39L | 116832 |

| | | | | | | | | | |
|---|---|---|---|---|---|---|---|---|---|
| ENSG00000136244.12 | 45.47415984 | -2.939813448 | 0.520747637 | -4.49318 | 7.02E-06 | 7.63E-05 | ENSG00000136244 | IL6 | 3569 |
| ENSG00000107159.13 | 8.436783978 | -2.954524481 | 0.680202279 | -3.46151 | 0.000537 | 0.00422711 | ENSG00000107159 | CA9 | 768 |
| ENSG00000104213.12 | 9.981496245 | -2.955711312 | 0.727384132 | -3.23861 | 0.001201 | 0.008739275 | ENSG00000104213 | PDGFRL | 5157 |
| ENSG00000152779.14 | 8.678044913 | -2.972718567 | 0.70757289 | -3.35332 | 0.000798 | 0.006063091 | ENSG00000152779 | SLC16A12 | 387700 |
| ENSG00000166106.3 | 42.74422455 | -2.973293666 | 0.485642778 | -4.88691 | 1.02E-06 | 1.24E-05 | ENSG00000166106 | ADAMTS1 | 170689 |
| ENSG00000285744.1 | 48.03047138 | -2.978200945 | 0.460789902 | -5.16114 | 2.45E-07 | 3.22E-06 | ENSG00000285744 | NA | NA |
| ENSG00000081803.16 | 235.7058475 | -2.98271755 | 0.21255635 | -11.2098 | 3.65E-29 | 2.40E-27 | ENSG00000081803 | CADPS2 | 93664 |
| ENSG00000168685.15 | 2358.375386 | -2.987774684 | 0.085205996 | -28.0236 | 8.39E-173 | 7.45E-170 | ENSG00000168685 | IL7R | 3575 |
| ENSG00000196668.4 | 9.848424844 | -2.990781909 | 0.565230863 | -4.22974 | 2.34E-05 | 0.000232577 | ENSG00000196668 | LINC0017: | 1E+08 |
| ENSG00000111799.21 | 1899.739133 | -2.993613864 | 0.140926738 | -16.9848 | 1.06E-64 | 2.30E-62 | ENSG00000111799 | COL12A1 | 1303 |
| ENSG00000229526.2 | 7.153713502 | -3.017020292 | 0.70613283 | -3.4229 | 0.00062 | 0.00480969 | ENSG00000229526 | KRT16P4 | 339244 |
| ENSG00000198542.14 | 6.347575756 | -3.022225187 | 0.737316992 | -3.28519 | 0.001019 | 0.00754494 | ENSG00000198542 | ITGBL1 | 9358 |
| ENSG00000092621.12 | 1083.822503 | -3.026413126 | 0.092964023 | -26.1006 | 3.59E-150 | 2.43E-147 | ENSG00000092621 | PHGDH | 26227 |
| ENSG00000248429.5 | 15.44609153 | -3.033246479 | 0.47330062 | -5.14102 | 2.73E-07 | 3.56E-06 | ENSG00000248429 | FAM198B- | 285505 |
| ENSG00000162409.11 | 4.654593377 | -3.036517481 | 0.748430635 | -3.2555 | 0.001132 | 0.008294521 | ENSG00000162409 | PRKAA2 | 5563 |
| ENSG00000261618.2 | 34.65144904 | -3.038211184 | 0.474070789 | -5.14314 | 2.70E-07 | 3.52E-06 | ENSG00000261618 | NA | NA |
| ENSG00000135747.11 | 5.19619777 | -3.045902809 | 0.717322982 | -3.40977 | 0.00065 | 0.005027396 | ENSG00000135747 | ZNF670-ZI | 1.01E+08 |
| ENSG00000249867.6 | 27.68819236 | -3.051182408 | 0.496072122 | -4.94118 | 7.77E-07 | 9.56E-06 | ENSG00000249867 | LINC0274: | 1.05E+08 |
| ENSG00000184185.10 | 8.445572557 | -3.057951707 | 0.573904778 | -4.28286 | 1.85E-05 | 0.000186839 | ENSG00000184185 | KCNJ12 | 3768 |
| ENSG00000156103.16 | 176.3065346 | -3.058781362 | 0.251439127 | -9.77883 | 1.39E-22 | 6.10E-21 | ENSG00000156103 | MMP16 | 4325 |
| ENSG00000198944.5 | 9.485641087 | -3.089580275 | 0.563777524 | -4.41589 | 1.01E-05 | 0.000106525 | ENSG00000198944 | SOWAHA | 134548 |
| ENSG00000258018.2 | 10.22232577 | -3.139776178 | 0.574335512 | -4.42211 | 9.77E-06 | 0.00010375 | ENSG00000258018 | NA | NA |
| ENSG00000228695.10 | 32.04301175 | -3.215197652 | 0.337996934 | -7.73734 | 1.02E-14 | 2.71E-13 | ENSG00000228695 | CES1P1 | 51716 |
| ENSG00000287707.1 | 15.00604527 | -3.229255476 | 0.497802856 | -5.28172 | 1.28E-07 | 1.74E-06 | ENSG00000287707 | NA | NA |
| ENSG00000175841.8 | 3.254162206 | -3.262376863 | 0.985918266 | -2.7004 | 0.006926 | 0.041656562 | ENSG00000175841 | NA | NA |
| ENSG00000273786.1 | 3.771134741 | -3.267972427 | 0.887986752 | -3.00452 | 0.00266 | 0.017873547 | ENSG00000273786 | NA | NA |
| ENSG00000163508.12 | 92.95178527 | -3.283610203 | 0.206310685 | -13.0076 | 1.11E-38 | 1.05E-36 | ENSG00000163508 | EOMES | 8320 |
| ENSG00000111348.9 | 975.7247431 | -3.32554382 | 0.211820822 | -12.8672 | 6.88E-38 | 6.36E-36 | ENSG00000111348 | ARHGDIB | 397 |
| ENSG00000164292.13 | 515.4296837 | -3.327183108 | 0.233964683 | -11.6564 | 2.13E-31 | 1.57E-29 | ENSG00000164292 | RHOBTB3 | 22836 |
| ENSG00000132182.12 | 1078.291163 | -3.327577386 | 0.100737499 | -27.0761 | 1.88E-161 | 1.39E-158 | ENSG00000132182 | NUP210 | 23225 |
| ENSG00000287618.1 | 2.079676188 | -3.367745742 | 1.040832742 | -2.65916 | 0.007833 | 0.046384007 | ENSG00000287618 | NA | NA |
| ENSG00000271659.1 | 3.672452556 | -3.386034731 | 0.889309737 | -3.13281 | 0.001731 | 0.012149283 | ENSG00000271659 | NA | NA |
| ENSG00000168505.7 | 14.52784799 | -3.392227036 | 0.48438844 | -5.76444 | 8.19E-09 | 1.28E-07 | ENSG00000168505 | GBX2 | 2637 |
| ENSG00000242444.3 | 3.134033607 | -3.463443219 | 1.065911317 | -2.68638 | 0.007223 | 0.043211881 | ENSG00000242444 | RPL18P9 | 1E+08 |
| ENSG00000101938.14 | 5.430744 | -3.481303 | 0.8702589 | -3.31086 | 0.00093 | 0.006945674 | ENSG00000101938 | CHRDL1 | 91851 |
| ENSG00000138829.12 | 6068.26791 | -3.494806325 | 0.198962718 | -14.5495 | 5.88E-48 | 7.83E-46 | ENSG00000138829 | FBN2 | 2201 |
| ENSG00000171435.13 | 17.53601148 | -3.505917222 | 0.445970331 | -6.51594 | 7.22E-11 | 1.39E-09 | ENSG00000171435 | KSR2 | 283455 |
| ENSG00000139800.8 | 85.76300237 | -3.539409462 | 0.222940898 | -13.1847 | 1.07E-39 | 1.06E-37 | ENSG00000139800 | ZIC5 | 85416 |
| ENSG00000198719.9 | 382.9723148 | -3.54038655 | 0.143832284 | -20.4432 | 6.91E-93 | 2.59E-90 | ENSG00000198719 | DLL1 | 28514 |
| ENSG00000105610.5 | 2.483732825 | -3.558216621 | 1.068643072 | -2.7682 | 0.005637 | 0.034728147 | ENSG00000105610 | KLF1 | 10661 |
| ENSG00000205562.3 | 12.09234449 | -3.561442238 | 0.596323367 | -4.96617 | 6.83E-07 | 8.46E-06 | ENSG00000205562 | NA | NA |
| ENSG00000248538.8 | 6.517024049 | -3.605398796 | 0.698149139 | -4.30481 | 1.67E-05 | 0.000170342 | ENSG00000248538 | LOC15727 | 157273 |

| | | | | | | | | | |
|---|---|---|---|---|---|---|---|---|---|
| ENSG00000227200.1 | 2.545530198 | -3.696092861 | 1.122199239 | -2.75895 | 0.005799 | 0.035610966 | ENSG00000227200 | NA | NA |
| ENSG00000144583.5 | 22.74229264 | -3.711335231 | 0.489756141 | -6.35283 | 2.11E-10 | 3.93E-09 | ENSG00000144583 | MARCHF4 | 57574 |
| ENSG00000226153.1 | 2.001622 | -3.715548 | 1.078638 | -2.88841 | 0.003872 | 0.02491634 | ENSG00000226153 | CHP1P3 | 1.17E+08 |
| ENSG00000224940.8 | 4.739754057 | -3.720223148 | 0.853455741 | -3.65599 | 0.000256 | 0.00214276 | ENSG00000224940 | PRRT4 | 401399 |
| ENSG00000226609.2 | 3.746020905 | -3.744934014 | 0.997221853 | -3.1537 | 0.001612 | 0.011363318 | ENSG00000226609 | NA | NA |
| ENSG00000103489.11 | 9.088331741 | -3.745924312 | 0.660366309 | -4.76391 | 1.90E-06 | 2.21E-05 | ENSG00000103489 | XYLT1 | 64131 |
| ENSG00000258952.2 | 5.03829524 | -3.773341075 | 0.882615143 | -3.59538 | 0.000324 | 0.002655743 | ENSG00000258952 | NA | NA |
| ENSG00000164362.20 | 42.53011526 | -3.803457464 | 0.334692672 | -9.57134 | 1.06E-21 | 4.45E-20 | ENSG00000164362 | TERT | 7015 |
| ENSG00000258616.5 | 3.10392153 | -3.8158091 | 0.963588494 | -3.33733 | 0.000846 | 0.006384798 | ENSG00000258616 | LINC0230: | 1.05E+08 |
| ENSG00000243089.2 | 2.859594106 | -3.840136859 | 1.035232038 | -3.12987 | 0.001749 | 0.012255402 | ENSG00000243089 | NA | NA |
| ENSG00000259254.1 | 3.048623896 | -3.894286157 | 1.019055916 | -3.23268 | 0.001226 | 0.008903137 | ENSG00000259254 | NA | NA |
| ENSG00000287466.1 | 1.41987303 | -3.906414026 | 1.177038745 | -2.8091 | 0.004968 | 0.031098186 | ENSG00000287466 | NA | NA |
| ENSG00000164125.15 | 8.577475997 | -4.024398786 | 0.708146895 | -4.83572 | 1.33E-06 | 1.57E-05 | ENSG00000164125 | GASK1B | 51313 |
| ENSG00000255450.1 | 2.387164394 | -4.11270766 | 1.093057094 | -3.21365 | 0.001311 | 0.009463002 | ENSG00000255450 | LOC40167 | 401677 |
| ENSG00000215262.8 | 1.828768337 | -4.120791199 | 1.255722844 | -2.8038 | 0.00505 | 0.031561934 | ENSG00000215262 | KCNU1 | 157855 |
| ENSG00000165168.7 | 1.890706 | -4.146061 | 1.320492 | -2.68541 | 0.007244 | 0.04332784 | ENSG00000165168 | CYBB | 1536 |
| ENSG00000102445.18 | 15.8122342 | -4.274583711 | 0.561944627 | -6.53905 | 6.19E-11 | 1.20E-09 | ENSG00000102445 | RUBCNL | 80183 |
| ENSG00000250292.1 | 3.415143733 | -4.335685771 | 1.083562545 | -3.4476 | 0.000566 | 0.004424741 | ENSG00000250292 | NA | NA |
| ENSG00000139973.16 | 19.17513923 | -4.39929408 | 0.526625089 | -7.21442 | 5.42E-13 | 1.27E-11 | ENSG00000139973 | SYT16 | 83851 |
| ENSG00000171388.12 | 173.0406 | -4.417059 | 0.2658946 | -14.3555 | 9.84E-47 | 1.24E-44 | ENSG00000171388 | APLN | 8862 |
| ENSG00000279970.1 | 17.23787778 | -4.418979305 | 0.584570909 | -6.53296 | 6.45E-11 | 1.25E-09 | ENSG00000279970 | NA | NA |
| ENSG00000179242.16 | 55.00254 | -4.570408 | 0.36297 | -10.9387 | 7.53E-28 | 4.49E-26 | ENSG00000179242 | CDH4 | 1002 |
| ENSG00000232110.8 | 2.387211293 | -4.627838335 | 1.067996305 | -3.7714 | 0.000162 | 0.001410479 | ENSG00000232110 | NA | NA |
| ENSG00000233611.3 | 4.931166863 | -4.694925903 | 1.050781329 | -3.89703 | 9.74E-05 | 0.000881484 | ENSG00000233611 | NA | NA |
| ENSG00000250155.2 | 4.756561457 | -5.611687587 | 0.942231928 | -5.31895 | 1.04E-07 | 1.43E-06 | ENSG00000250155 | LOC10798 | 1.08E+08 |
