## Supplementary File for "Combined analysis of somatic mutations and gene expression reveals nuclear speckles-associated enhanced stemness in gingivobuccal carcinoma under DNA damage response"

[1]IISc Mathematics Initiative
[2]Developmental Biology and Genetics
[3]Computational and Data Science
Indian Institute of Science, Bengaluru 560 012, Karnataka, India

*Corresponding Author

ǂCo-correspondence:


#### Supplementary information

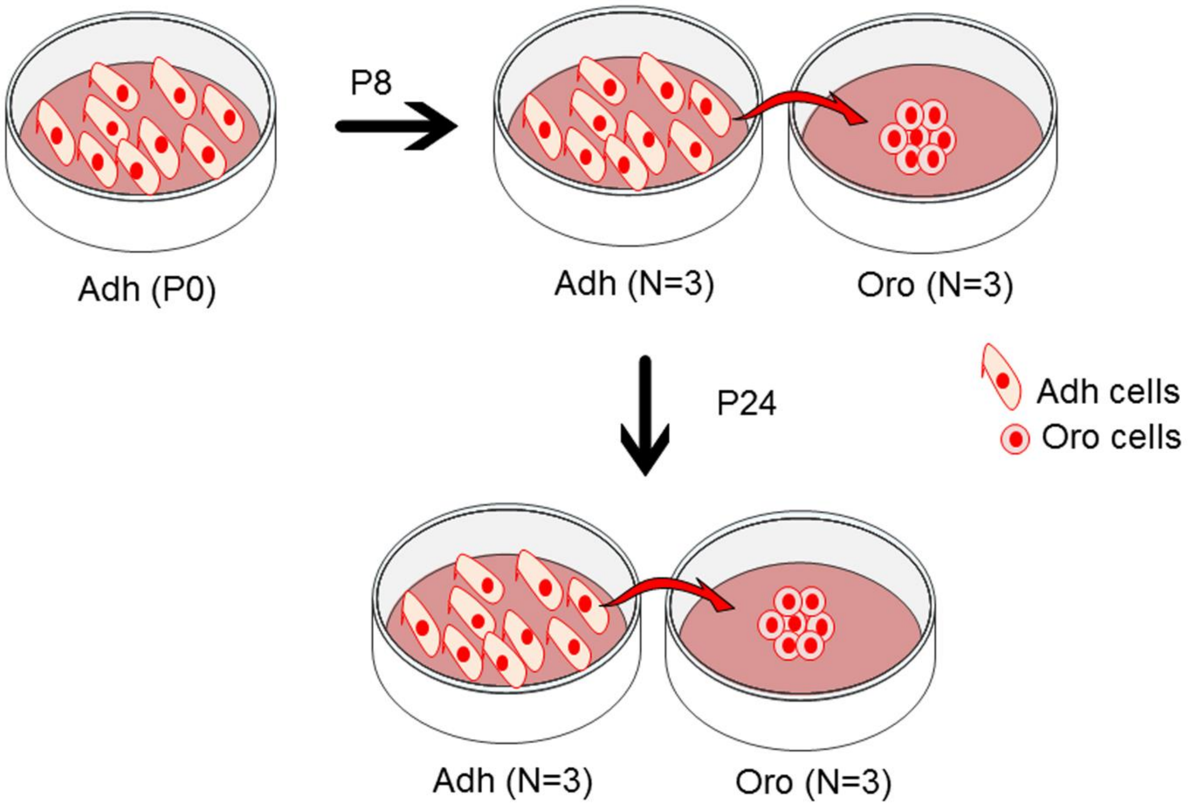


**Figure S1.** Schematic representation of the experimental design for IIOC019 cell line derived RNA-seq. IIOC019 cells were cultured in adherent (Adh) and orosphere (Oro) conditions. Additionally, two different passages, early passage (P8) and late passage (P24), and three technical triplicates (N=3) for each condition were used to generate the RNA-seq data.

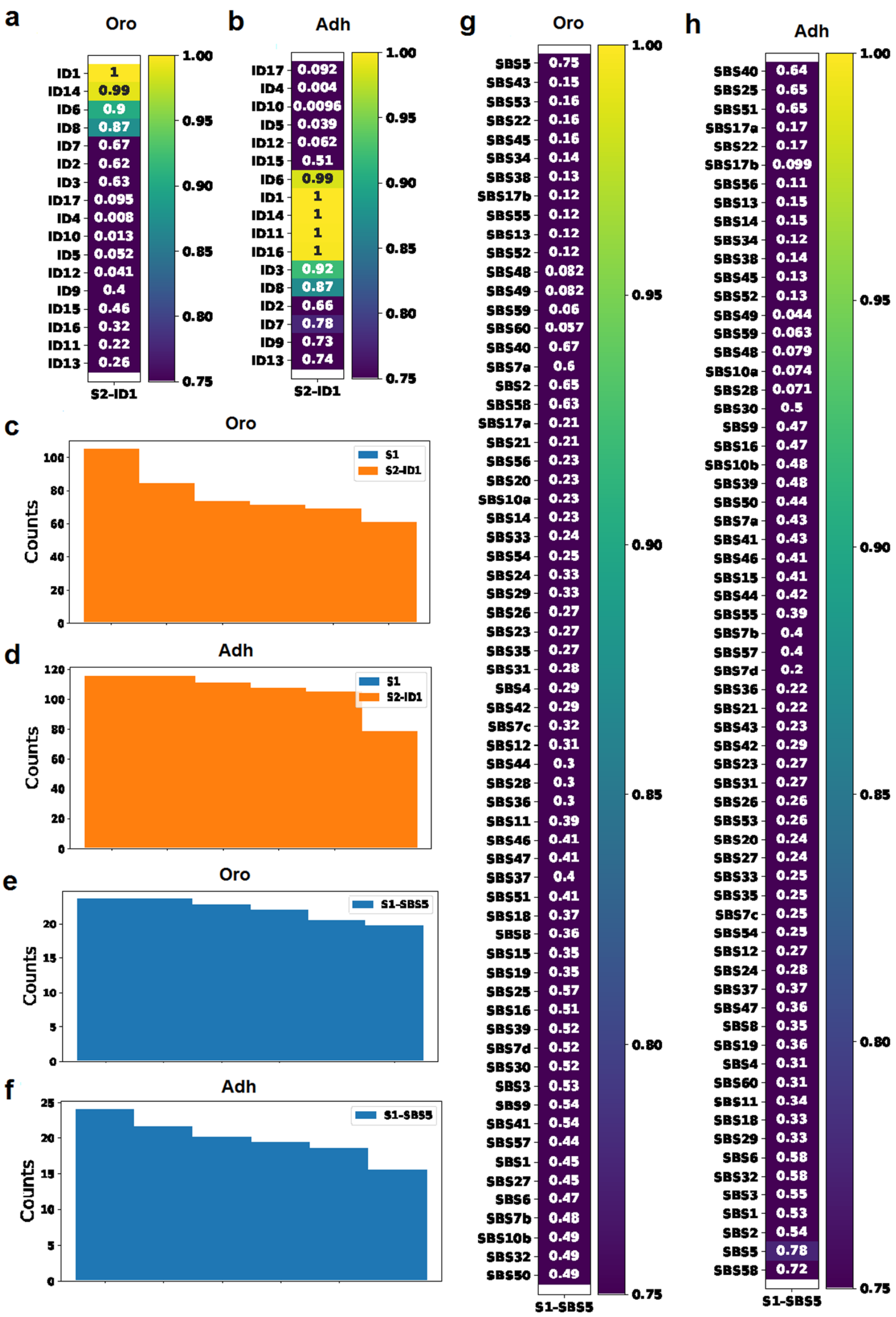
a
Oro
ID1 1
ID14 0.99
ID6 0.9
ID8 0.87
ID7 0.67
ID2 0.62
ID3 0.63
ID17 0.095
ID4 0.008
ID10 0.013
ID5 0.052
ID12 0.041
ID9 0.4
ID15 0.46
ID16 0.32
ID11 0.22
ID13 0.26
S2-ID1
1.00 0.95 0.90 0.85 0.80 0.75
b
Adh
ID17 0.092
ID4 0.004
ID10 0.0096
ID5 0.039
ID12 0.062
ID15 0.51
ID6 0.99
ID1 1
ID14 1
ID11 1
ID16 1
ID3 0.92
ID8 0.87
ID2 0.66
ID7 0.78
ID9 0.73
ID13 0.74
S2-ID1
1.00 0.95 0.90 0.85 0.80 0.75
c
Oro
Counts
S1
S2-ID1
d
Adh
Counts
S1
S2-ID1
e
Oro
Counts
S1-SBS5
f
Adh
Counts
S1-SBS5
g
Oro
SBS5 0.75
SBS43 0.15
SBS53 0.16
SBS22 0.16
SBS45 0.16
SBS34 0.14
SBS38 0.13
SBS17b 0.12
SBS55 0.12
SBS13 0.12
SBS52 0.12
SBS48 0.082
SBS49 0.082
SBS59 0.06
SBS60 0.057
SBS40 0.67
SBS7a 0.6
SBS2 0.65
SBS58 0.63
SBS17a 0.21
SBS21 0.21
SBS56 0.23
SBS20 0.23
SBS10a 0.23
SBS14 0.23
SBS33 0.24
SBS54 0.25
SBS24 0.33
SBS29 0.33
SBS26 0.27
SBS23 0.27
SBS35 0.27
SBS31 0.28
SBS4 0.29
SBS42 0.29
SBS7c 0.32
SBS12 0.31
SBS44 0.3
SBS28 0.3
SBS36 0.3
SBS11 0.39
SBS46 0.41
SBS47 0.41
SBS37 0.4
SBS51 0.41
SBS18 0.37
SBS8 0.36
SBS15 0.35
SBS19 0.35
SBS25 0.57
SBS16 0.51
SBS39 0.52
SBS7d 0.52
SBS30 0.52
SBS3 0.53
SBS9 0.54
SBS41 0.54
SBS57 0.44
SBS1 0.45
SBS27 0.45
SBS6 0.47
SBS7b 0.48
SBS10b 0.49
SBS32 0.49
SBS50 0.49
S1-SBS5
1.00 0.95 0.90 0.85 0.80 0.75
h
Adh
SBS40 0.64
SBS25 0.65
SBS51 0.65
SBS17a 0.17
SBS22 0.17
SBS17b 0.099
SBS56 0.11
SBS13 0.15
SBS14 0.15
SBS34 0.12
SBS38 0.14
SBS45 0.13
SBS52 0.13
SBS49 0.044
SBS59 0.063
SBS48 0.079
SBS10a 0.074
SBS28 0.071
SBS30 0.5
SBS9 0.47
SBS16 0.47
SBS10b 0.48
SBS39 0.48
SBS50 0.44
SBS7a 0.43
SBS41 0.43
SBS46 0.41
SBS15 0.41
SBS44 0.42
SBS55 0.39
SBS7b 0.4
SBS57 0.4
SBS7d 0.2
SBS36 0.22
SBS21 0.22
SBS43 0.23
SBS42 0.29
SBS23 0.27
SBS31 0.27
SBS26 0.26
SBS53 0.26
SBS20 0.24
SBS27 0.24
SBS33 0.25
SBS35 0.25
SBS7c 0.25
SBS54 0.25
SBS12 0.27
SBS24 0.28
SBS37 0.37
SBS47 0.36
SBS8 0.35
SBS19 0.36
SBS4 0.31
SBS60 0.31
SBS11 0.34
SBS18 0.33
SBS29 0.33
SBS6 0.58
SBS32 0.58
SBS3 0.55
SBS1 0.53
SBS2 0.54
SBS5 0.78
SBS58 0.72
S1-SBS5
1.00 0.95 0.90 0.85 0.80 0.75

42

**Figure S2.** Mutational signature profile of IIOC019 Oro (N=6, including P8 and P24) and IIOC019 Adh (N=6, including P8 and P24) using SignatureAnalyzer v0.0.7 python tool showing the most enriched mutational signature (cosine similarity enrichment score≥0.75). **a-b** DNA mismatch repair deficiency (ID1), Defective homologous recombination-based DNA damage repair (ID6), and DNA double-strand breaks-based DNA damage repair (ID8) of both subgroups of IIOC019 based on small insertion and deletion (ID) cosmic3_ID. Additionally, tobacco-associated carcinogens (ID3), defective DNA mismatch repair (ID7), DNA double strand breaks by non-homologous DNA end-joining (ID8) in IIOC019 Adh. Also, additional signatures with unknown importance (ID11, ID16) in IIOC019 Adh and mutational signature (ID14) in both subgroups of IIOC019. The mutational signature contribution (counts) per sample for **c-d** cosmic3_ID and **e-f** cosmic3_exome SBS for both the subgroups of IIOC019 cohorts (N=6). **g-h** Tobacco-associated carcinogens (SBS5) mutational signature profile of both subgroups of IIOC019 based on single base substitution (SBS) cosmic3_exome. Here, S1 exhibits the prevailing signature pattern among the corresponding COSMIC mutational signatures.

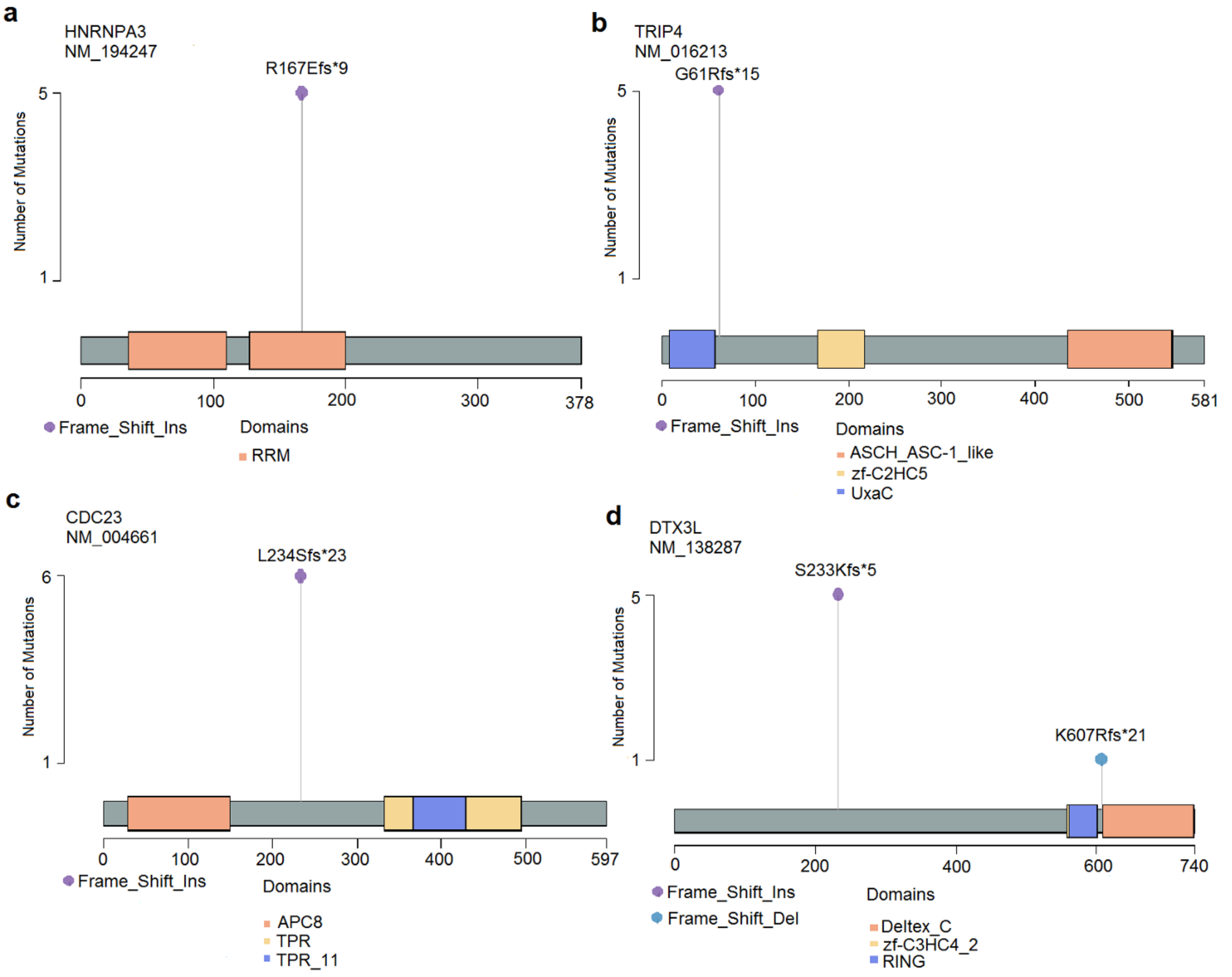

**Figure S3.** The LollipopPlot showing the location of somatic variants within the identified DNA damage response (DDR) genes/proteins linked to functional damage and nonsense-mediated decay (NMD), found in five or six samples in IIOC019 cohorts (N=6, including P8 and P24) using Maftools v2.6.05 R package. Frameshift insertion in **a** HNRNPA3: p.R167Efs*9 (5), **b** TRIP4: p.G61Rfs*15 (5) in IIOC019 Oro, **c** CDC23: p.L234Sfs*23 (6) and **d** DTX3L: p.S233Kfs*5 (5); p.K607Rfs*21 (1) in IIOC019 Adh. Here, the number of patients exhibiting recurring mutations per gene is denoted within the small bracket. The prefix NM_ signifies the RefSeq transcript identifier and various domains on the corresponding protein structure are highlighted using distinct colors.

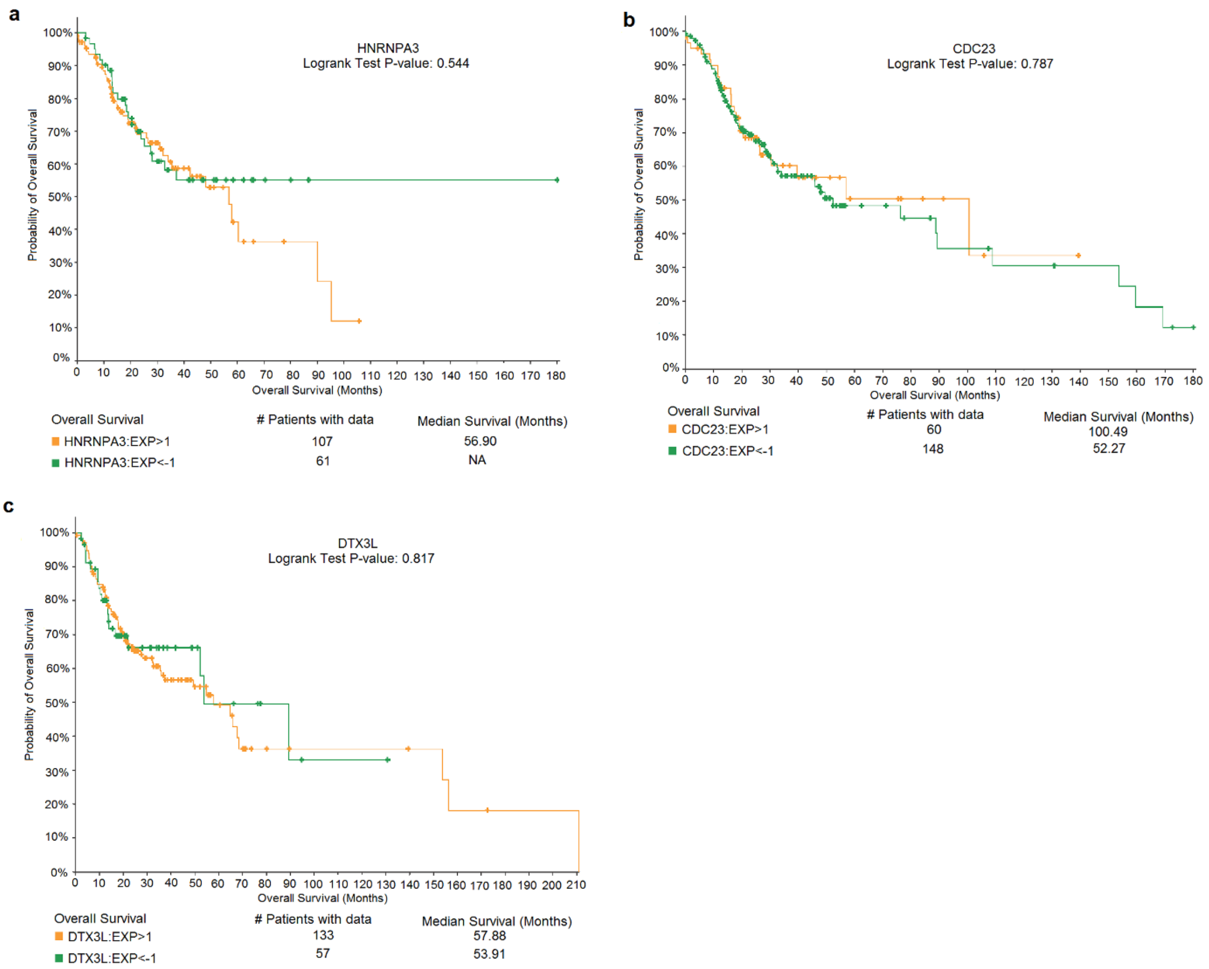


**Figure S4.** The plot showing Kaplan–Meier survival analysis correlating mRNA gene expression (expression Z-score or EXP indicating expression Z-score relative to diploid tumor samples) with overall survival outcome for the identified functionally damaging and NMD frameshift insertion somatic variants associated DNA damage response genes in the subgroups of the IIOC019 cell line cohorts (N=6, including P8 and P24; HNSCC TCGA Firehose Legacy cohort using cBioPortal accessed on 6 Jan 2024) **a** HNRNPA3 with Logrank test p-value (0.544) in IIOC019 Oro, **b** CDC23 with Logrank test p-value (0.787) and **c** DTX3L with Logrank test p-value (0.817) in IIOC019 Adh.

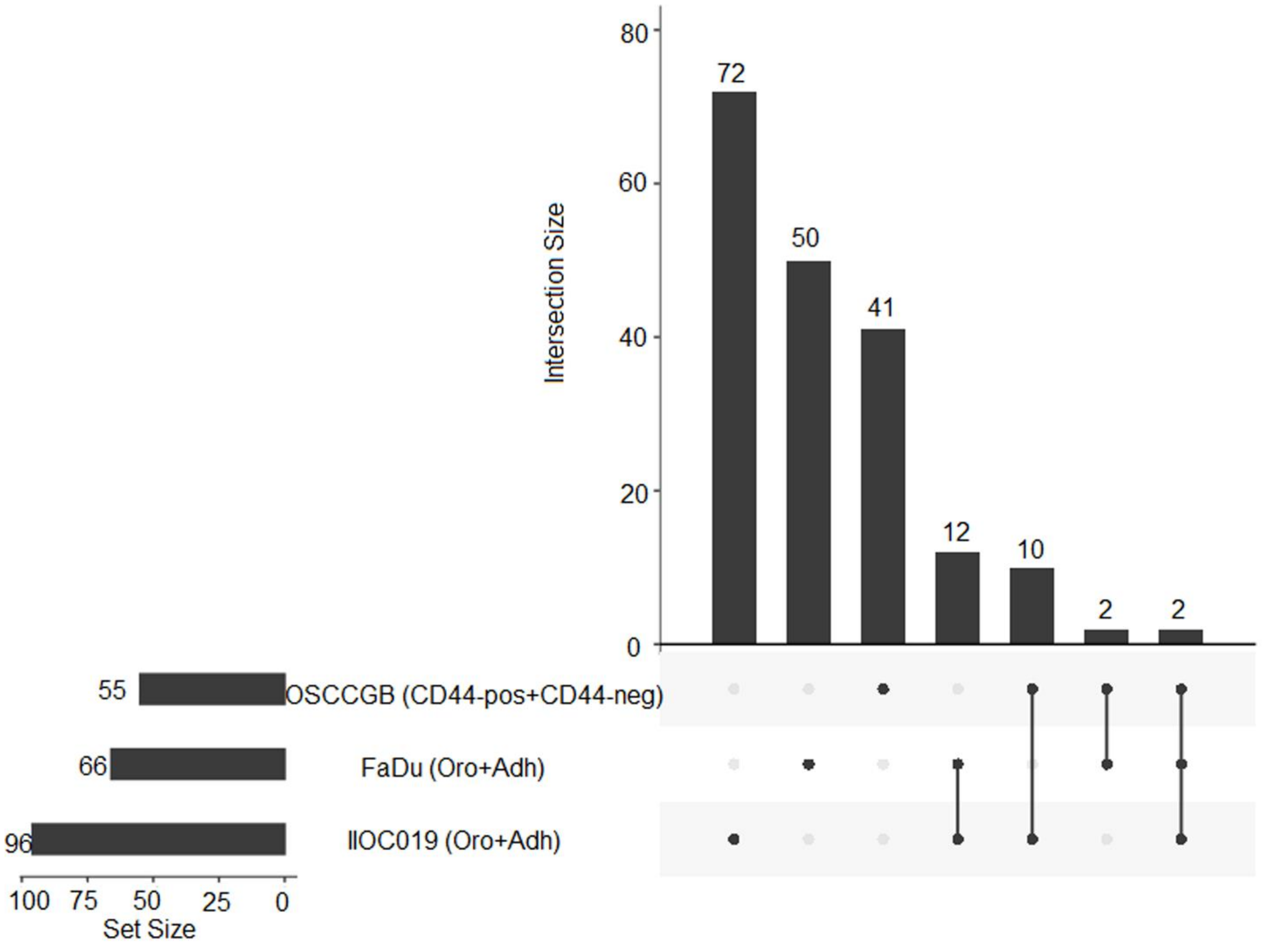


**Figure S5.** UpSet plot showing genomic similarities of the IIOC019 (Oro and Adh) cell line with existing cell FaDu (Oro and Adh) and our previously published OSCC-GB (CD44$^{+}$Lin$^{-}$/CD44-pos and CD44$^{-}$Lin$^{-}$/CD44-neg) primary tumor [1] indicated the relatedness of our novel IIOC019 cell line with OSCC-GB. The boxplot was plotted UpSetR v1.4.0 package in Rstudio R v4.0.5. The line between dots indicated the common genes among the groups.

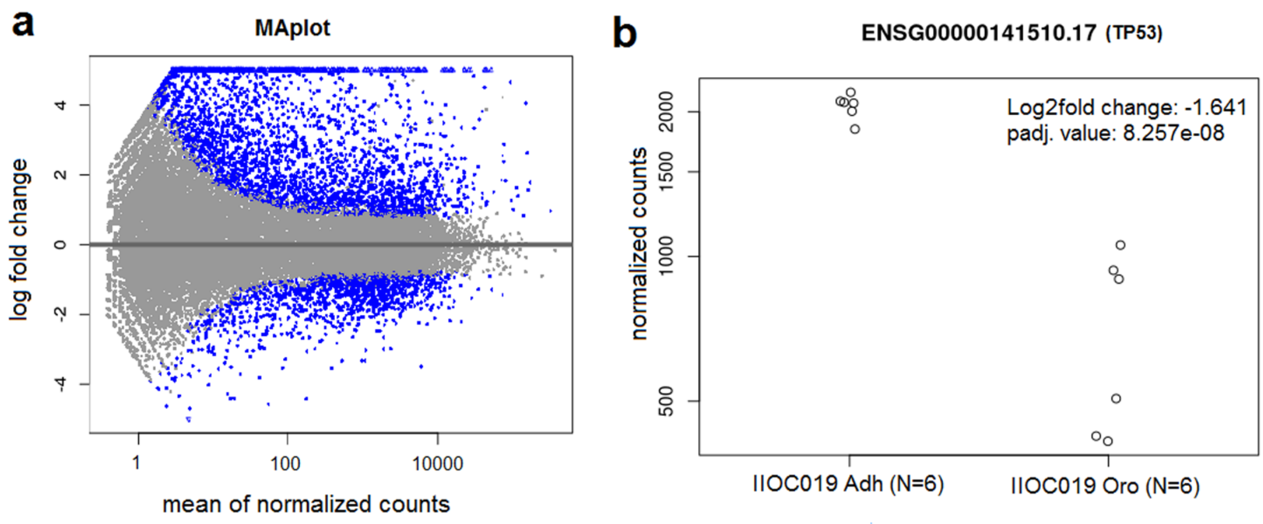


**Figure S6. a** The MAplot shows log2 fold changes over the mean of normalized counts for IIOC019 Oro (N=6, including P8 and P24) and IIOC019 Adh (N=6, including P8 and P24) showing higher up-regulated genes compared to down-regulated genes **b** the comparative normalized read counts for TP53 between IIOC019 Oro (N=6) versus IIOC019 Adh (N=6) with Log2fold change: -1.641 and padj. value: 8.257e-08 calculated using DESeq2 v1.30.1 R package. The blue dots represent the adjusted p-value (padj<0.1), and the points that fall out of the window are plotted as open triangles indicating either up or down.

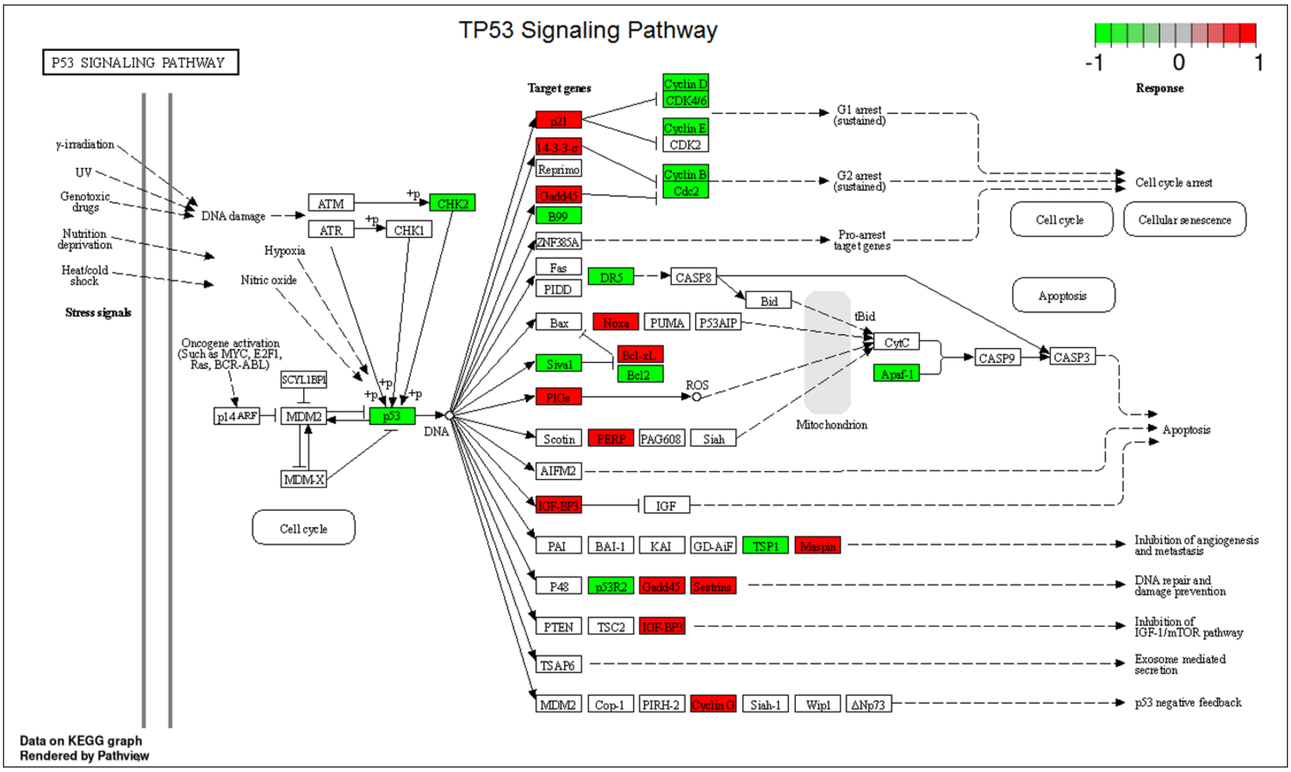


**Figure S7.** The identified differentially and statistically significant TP53 signaling KEGG pathway (p-value<0.05) for IIOC019 Oro (N=6, including P8 and P24) as compared to IIOC019 Adh (N=6, including P8 and P24) cohort using GAGE v2.40.2 and visualized in Pathview v1.30.1 R package.

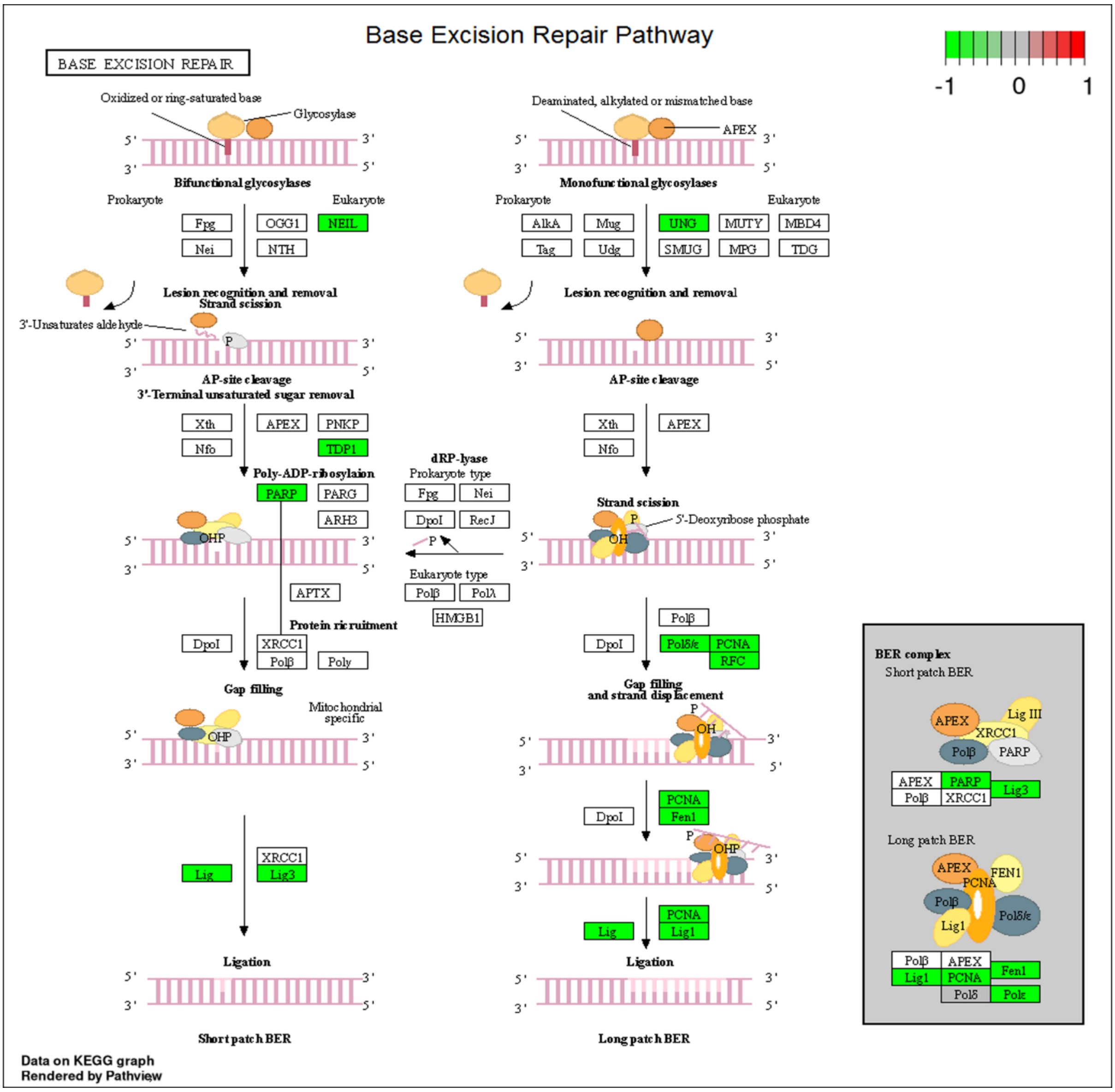


**Figure S8.** The identified differentially and statistically significant Base Excision Repair KEGG pathway (p-value<0.05) for IIOC019 Oro (N=6, including P8 and P24) as compared to IIOC019 Adh (N=6, including P8 and P24) cohort using GAGE v2.40.2 and visualized in Pathview v1.30.1 R package.

### Supplementary information

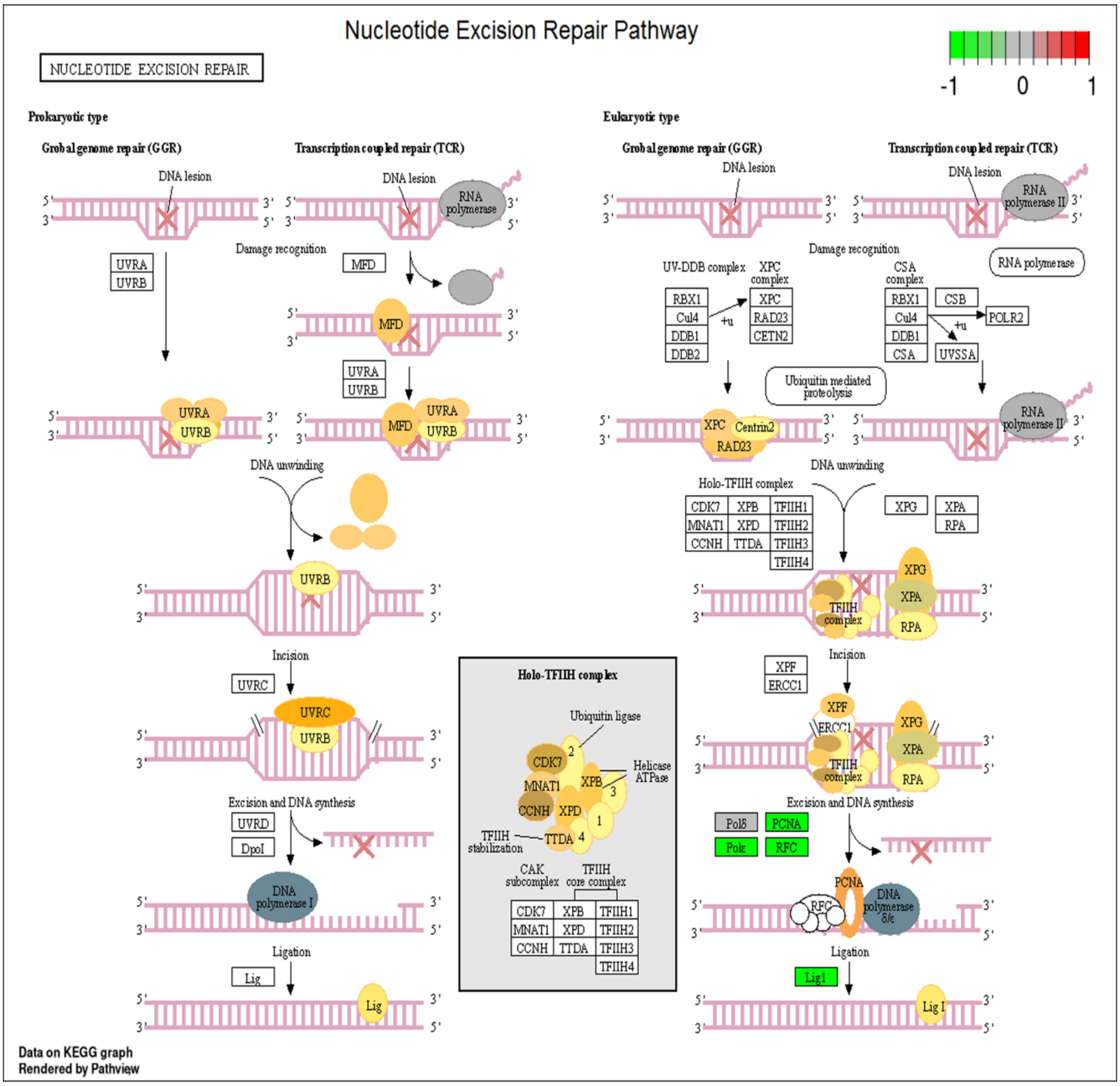


**Figure S9.** The identified differentially and statistically significant Nucleotide Excision Repair KEGG pathway (p-value<0.05) for IIOC019 Oro (N=6, including P8 and P24) as compared to IIOC019 Adh (N=6, including P8 and P24) cohort using GAGE v2.40.2 and visualized in Pathview v1.30.1 R package.

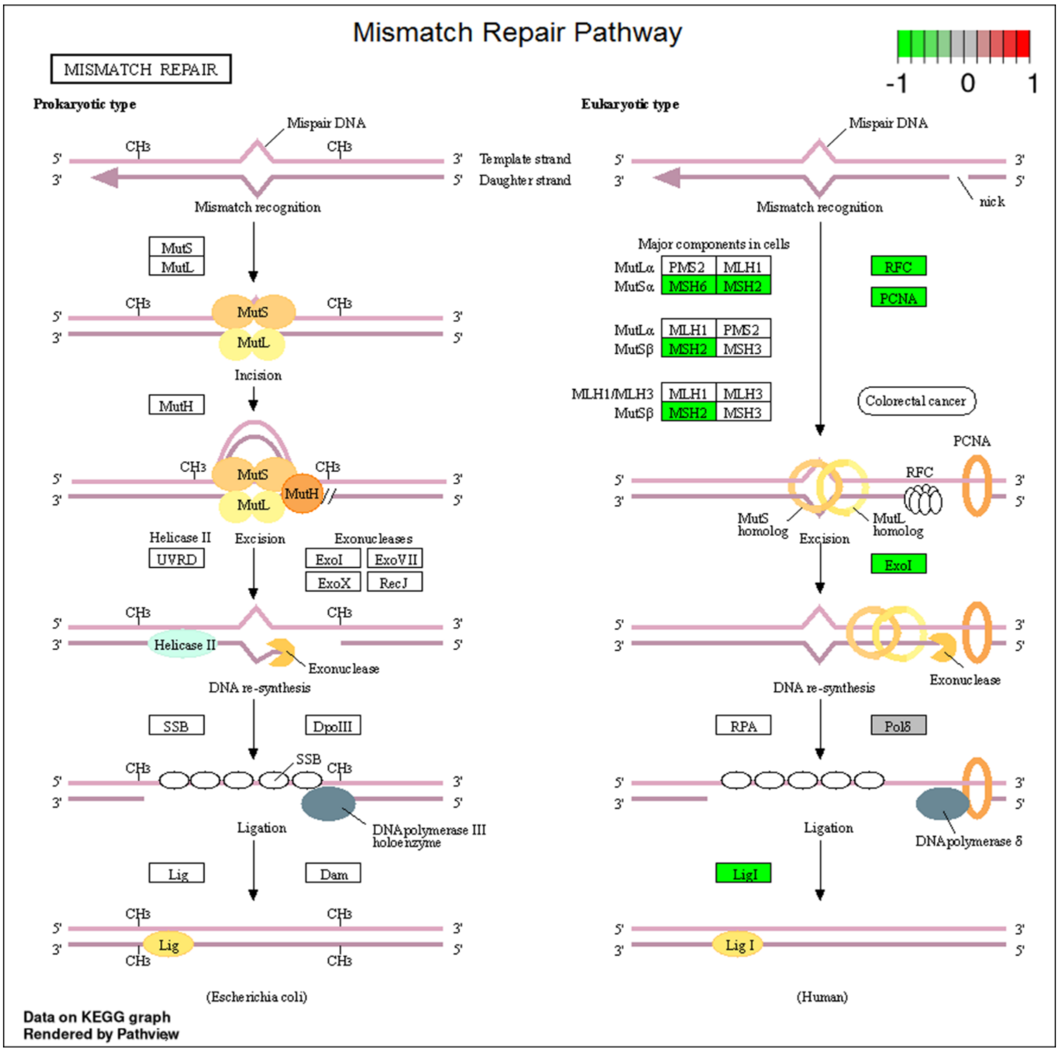


**Figure S10.** The identified differentially and statistically significant Mismatch Repair KEGG pathway (p-value<0.05) for IIOC019 Oro (N=6, including P8 and P24) as compared to IIOC019 Adh (N=6, including P8 and P24) cohort using GAGE v2.40.2 and visualized in Pathview v1.30.1 R package.

#### Supplementary information

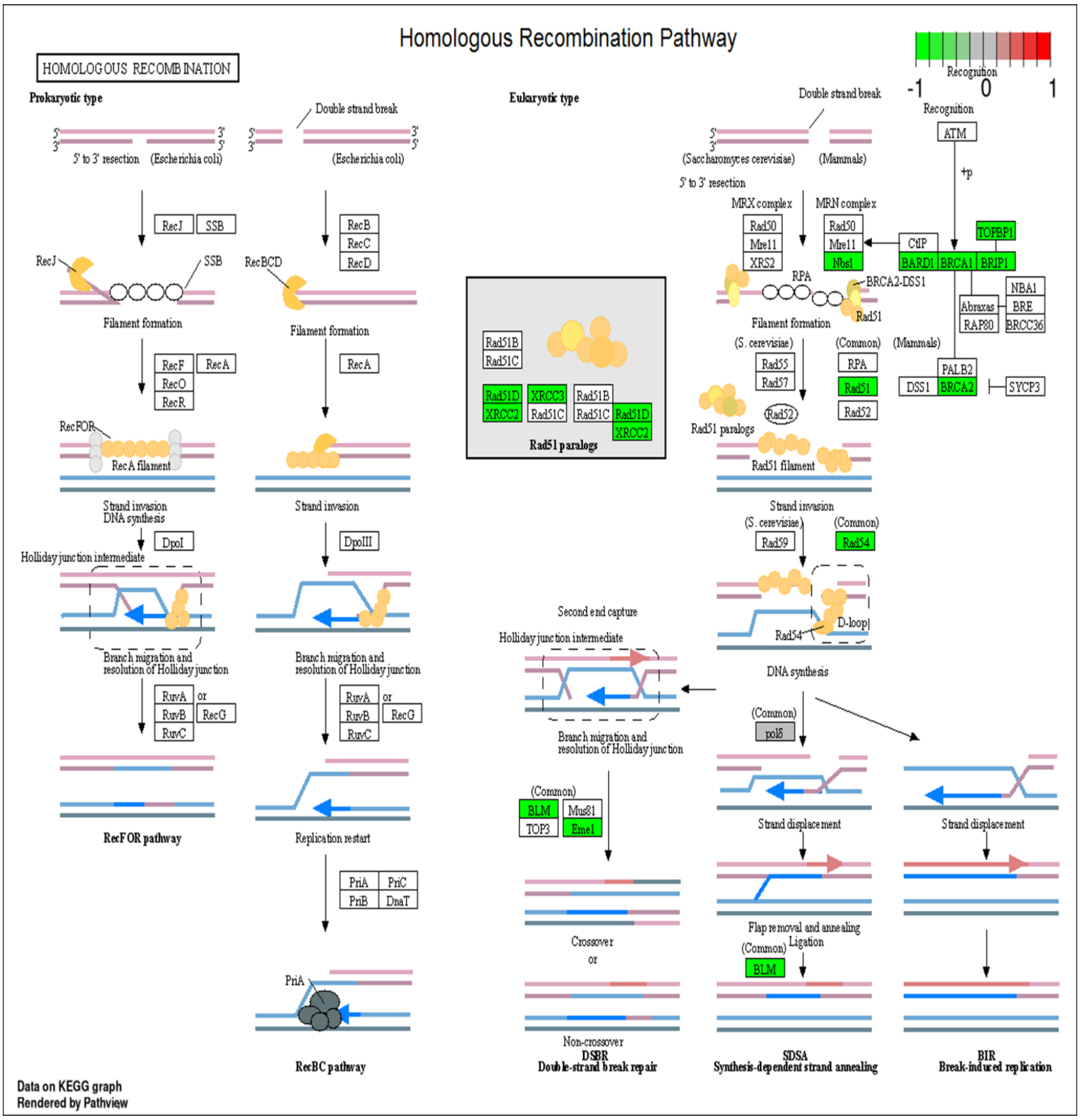


**Figure S11.** The identified differentially and statistically significant Homologous Recombination KEGG pathway (p-value<0.05) IIOC019 Oro (N=6, including P8 and P24) as compared to IIOC019 Adh (N=6, including P8 and P24) cohort using GAGE v2.40.2 and visualized in Pathview v1.30.1 R package.

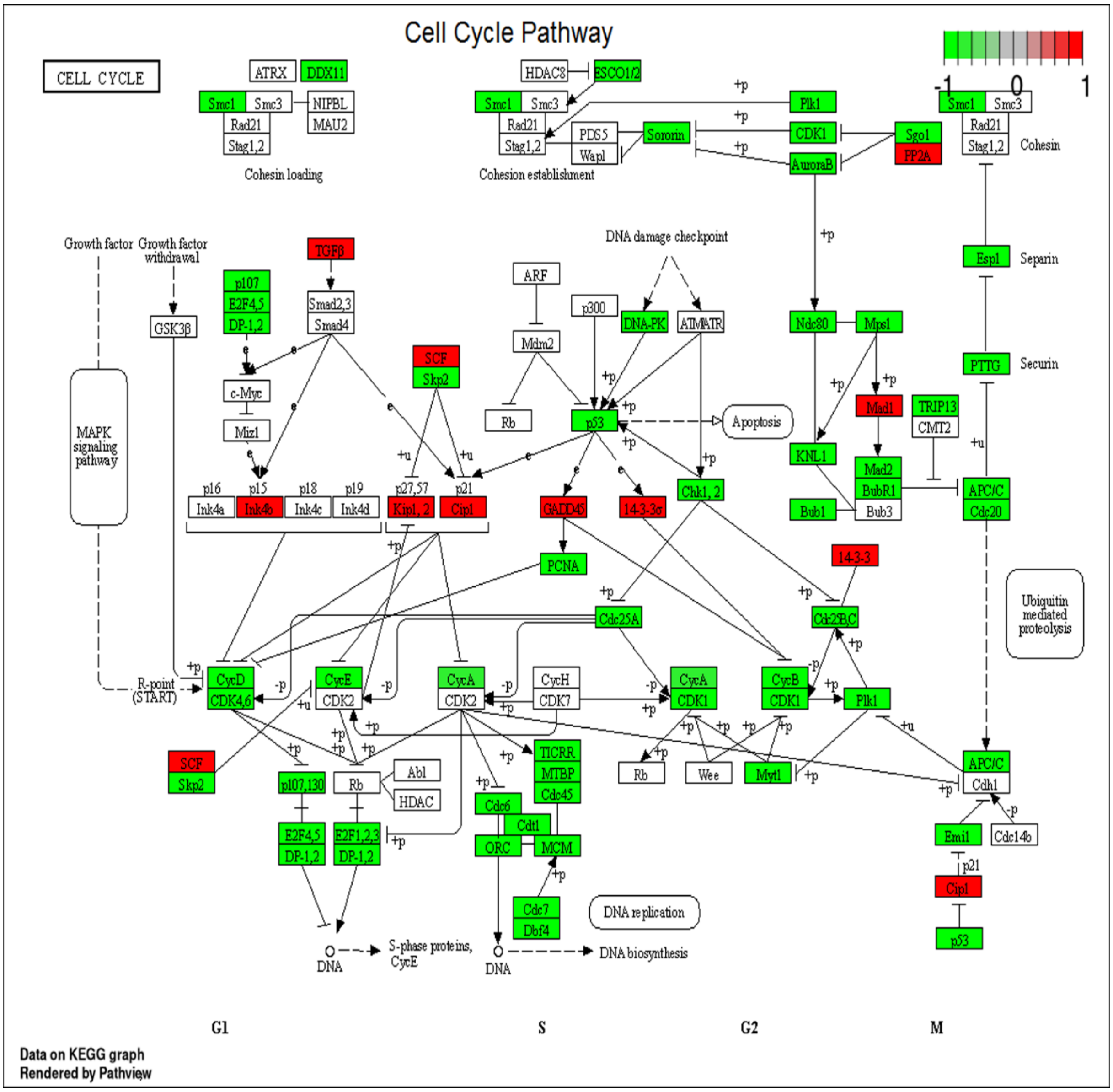


**Figure S12.** The identified differentially and statistically significant Cell cycle KEGG pathway (p-value<0.05) for IIOC019 Oro (N=6, including P8 and P24) as compared to IIOC019 Adh (N=6, including P8 and P24) cohort using GAGE v2.40.2 and visualized in Pathview v1.30.1 R package.

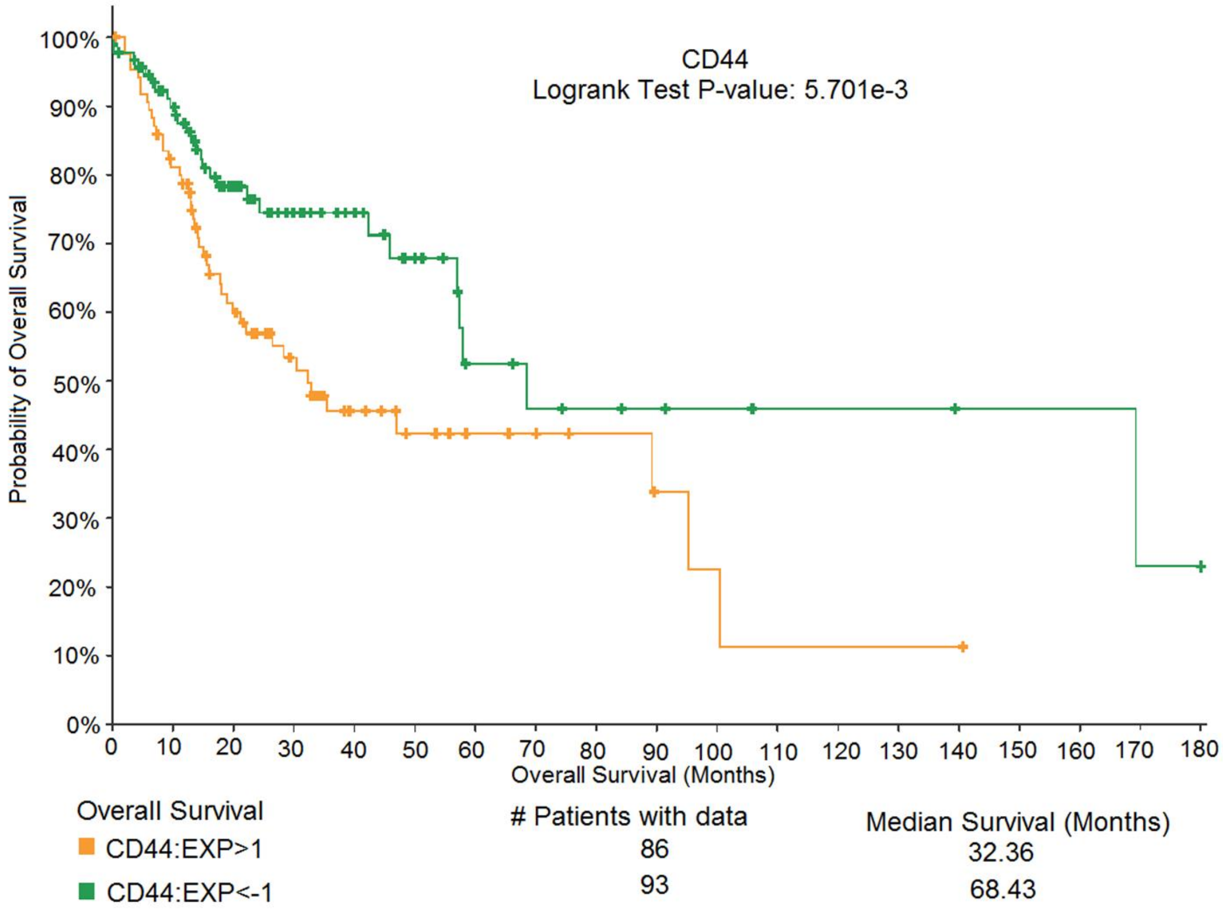


**Figure S13.** The plot showing Kaplan–Meier survival analysis correlating mRNA gene expression (expression Z-score or EXP indicating expression Z-score relative to diploid tumor samples) with overall survival outcome for CD44 with statistically significant Logrank test p-value (5.701e-3).

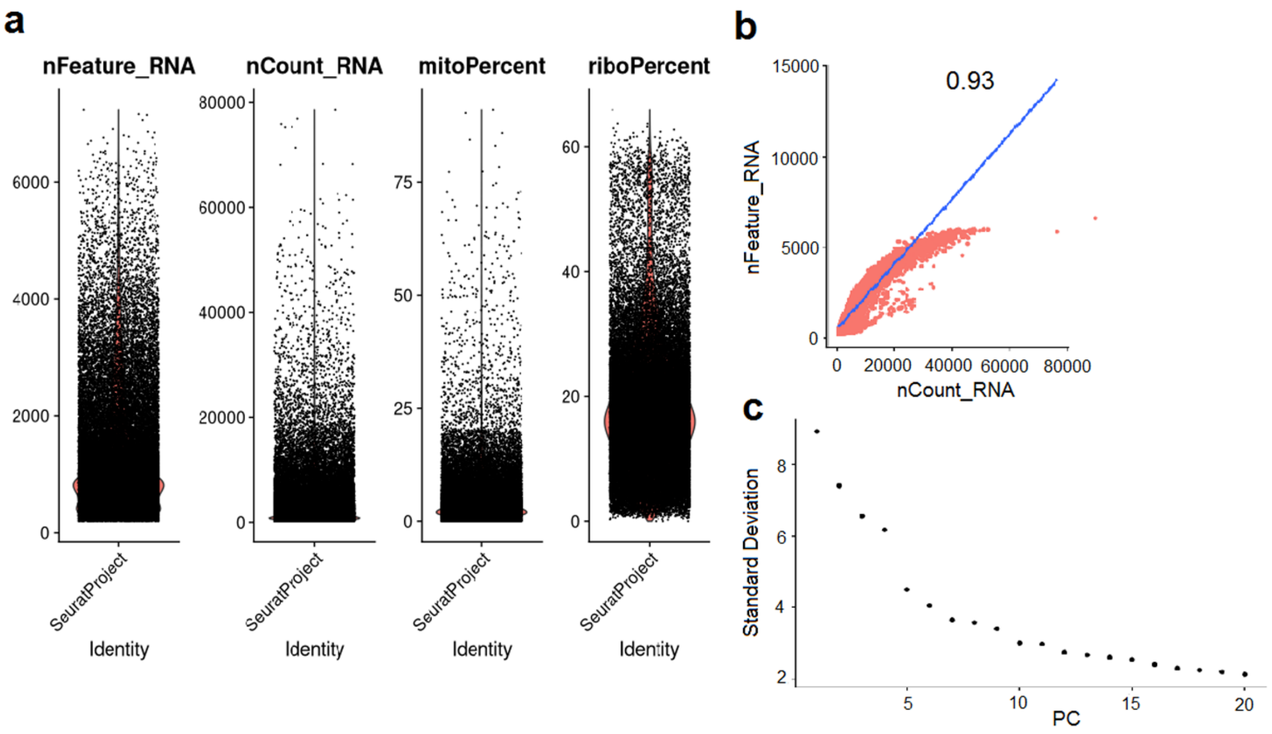


**Figure S14**. scRNA-seq analysis was performed to understand SON-mediated cancer stemness in HNSCC (N=8) using Seurat v4.3.0.1 R package, **a** Quality control showing the features, transcripts, mitochondrial and ribosomal content. **b** FeatureScatter plot showing genes and counts after quality control **c** Elbow plot used to identify the principal component (dims = 1:15) for harmony corrections.

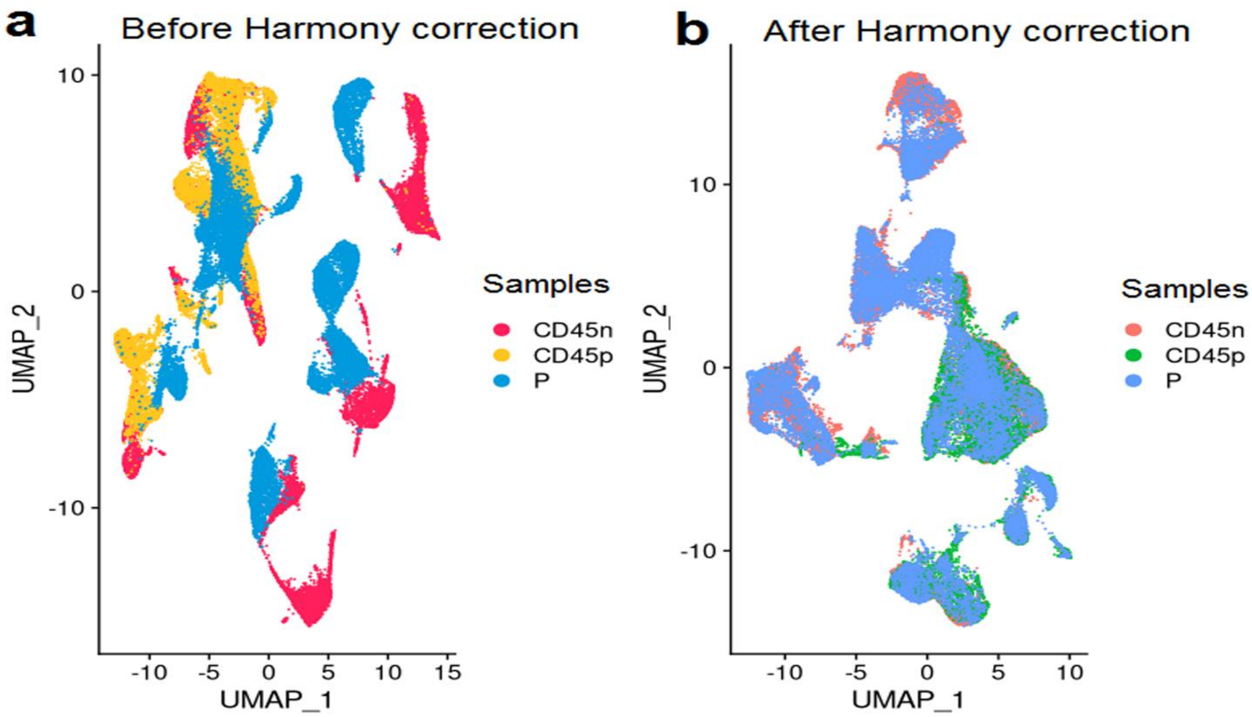

**Figure S15.** scRNA-seq analysis was performed to understand SON-mediated cancer stemness in HNSCC (N=8) using the Seurat v4.3.0.1 and Harmony v0.1.1 R package, UMAP plot showing cell cluster **a** before **b** after Harmony corrected cell clusters (N=39313). CD45n for non-immune cells, CD45p for immune cells (public data source from [2]), and P for (CD45n:CD45p::1:1) (public data source from [3]) in the plot.

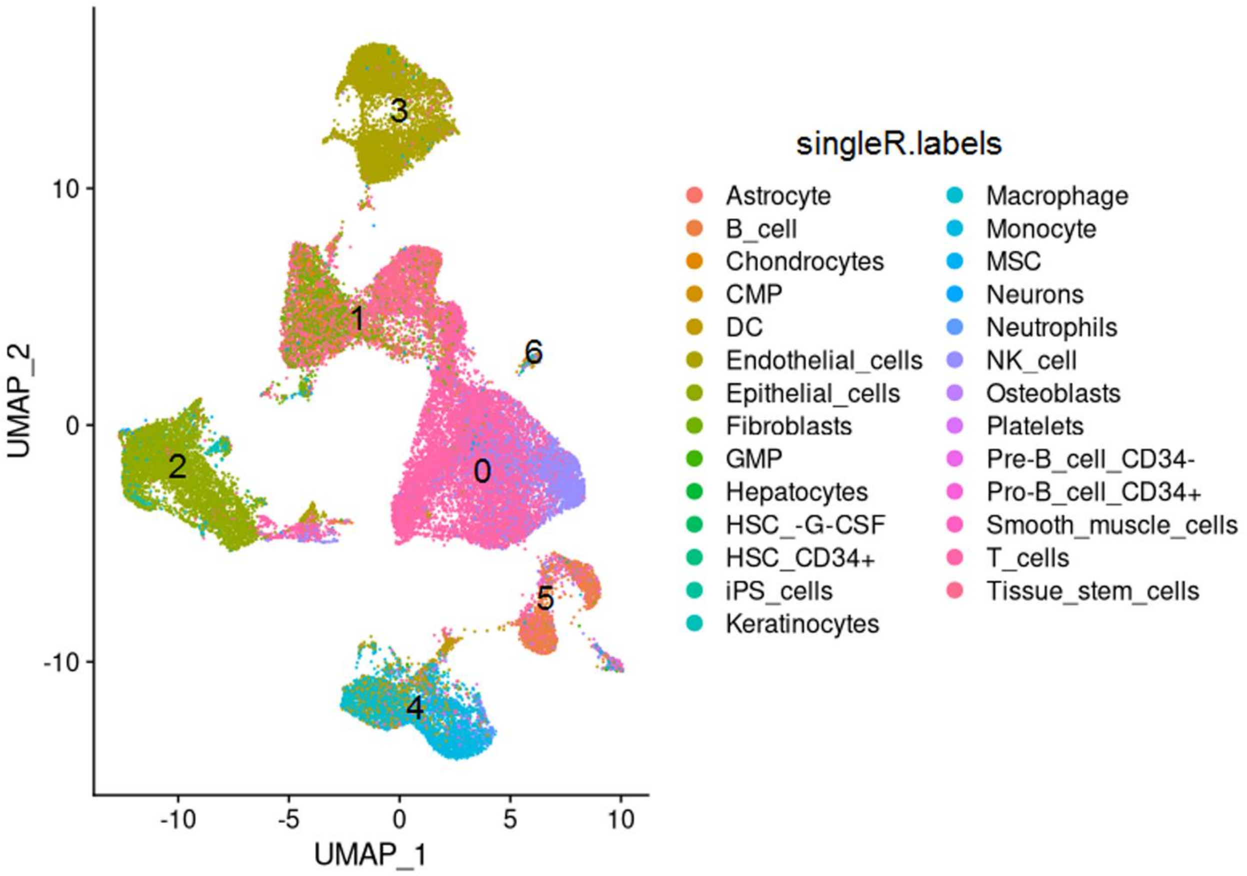


**Figure S16.** scRNA-seq analysis was performed to understand SON-mediated cancer stemness in HNSCC (N=8), a UMAP plot showing annotated cell types using singleR v1.4.1 R package with HumanPrimaryCellAtlasData as a reference from the celldex v1.0.0 R package. Here, the clusters are numbered as 0,1,2,3,4,5 and 6.

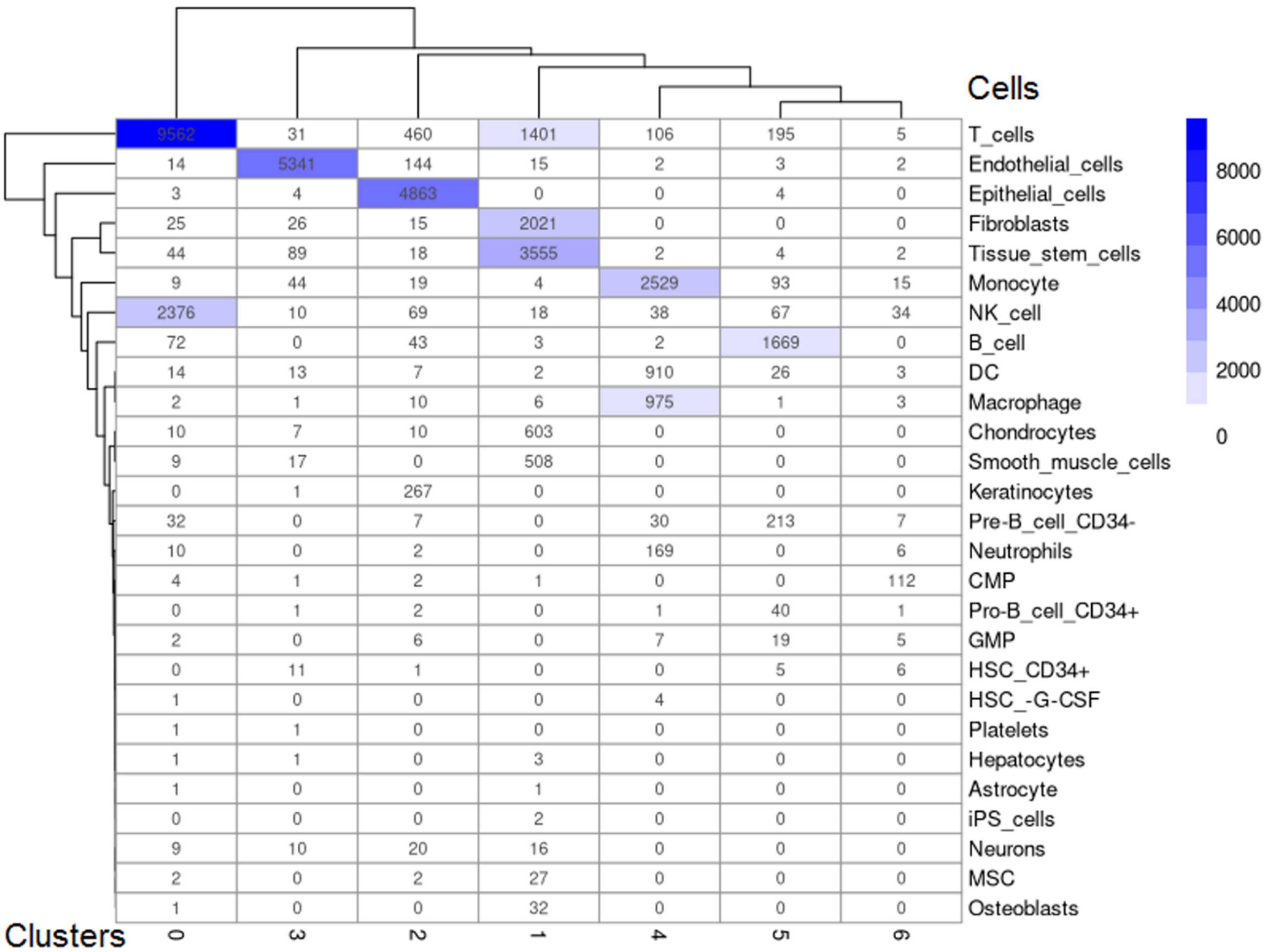


**Figure S17**. scRNA-seq analysis was performed to understand SON-mediated cancer stemness in HNSCC (N=8), a heatmap showing annotated cell types and associated counts using singleR v1.4.1 R package with HumanPrimaryCellAtlasData as a reference from the celldex v1.0.0 R package.

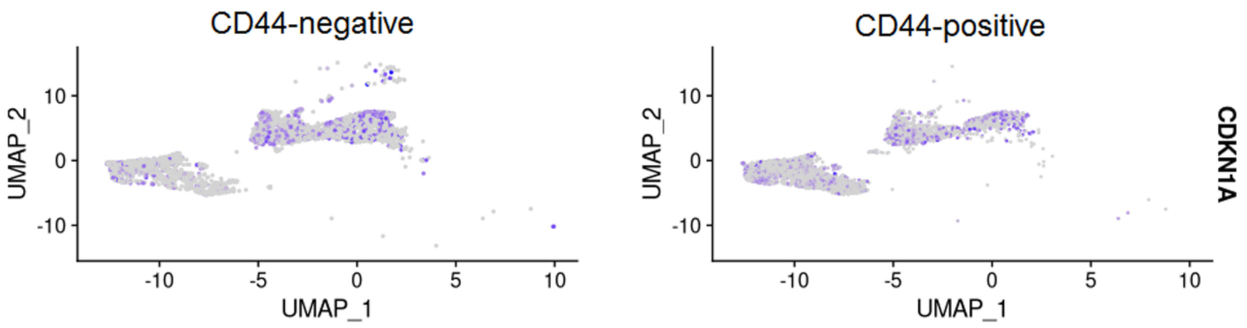


**Figure S18.** scRNA-seq analysis was performed to understand SON-mediated cancer stemness in HNSCC (N=8) using Seurat v4.3.0.1 R package, FeaturePlot showing TP53 known downstream target CDKN1A/p21 lower expression in CD44-positive as compared to CD44-negative cells in the Tissue stem cells and Epithelial cells cell types.

**Supplementary information**

#### S1 Data sources

The novel OSCC-GB cell line was derived from the gingivobuccal cavity of a 56-year-old female patient [4] with tumor tissue collected from KMIO hospital, Bangalore. The Indian patient had a history of smokeless tobacco consumption for over 20 years, with no records of cigarette smoking or alcohol use. The clinicopathological details of the patients include cT4aN2bM0, stage IV, neoadjuvant chemotherapy followed by surgery and subsequent adjuvant radiotherapy + chemotherapy, post-treatment rapid progression and death after 6 months. The cell line is named IIOC019, reflecting its ethnic origin, institute, cancer site, and sample number —Indian Institute of Science Oral Cancer 019 (IIOC019).

The resected tissue was processed following the method described earlier by our group [5] with slight modifications. The resected tumour tissue was rinsed in phosphate buffer saline supplemented with antibiotics, then subjected to mechanical and enzymatic digestion and cultured in standard tissue culture plastic dishes following relevant guidelines and regulations. Differential centrifugation isolated initial oral organoids, which were then trypsinized to obtain a single-cell suspension. Cells were routinely subcultured at a 1:3 split ratio and grown in DMEM media supplemented with 10% FBS and antibiotics. Additionally, these single cells were cultured under ultra-low attachment conditions in serum-free media supplemented with specific growth factors for 7 days to generate orospheres, following the established method for the orosphere assay [6] with minor modifications. An orosphere is defined as a non-adherent colony consisting of at least 25 cells or a sphere measuring over 50 μm in size [6]. This study used IIOC019 passage numbers P8 as early passage and P24 as late passage cells for each condition (Adherent/Adh and Orosphere/Oro) to perform RNA-seq experiments (Figure S1).

#### S2 Quality control, sequence alignments, and pre-processing of BAM files

A detailed version with commands for each step is mentioned in our previously published paper [1]. Briefly, ICGC acquired BAM files and converted them into FASTQ files using the SamToFastq tool from the Genome Analysis Toolkit (GATK) version 4.1.6.0 [7]. Only reads with itself and mate mapped from BAM files are used for further processing. Both the IIOC019 and FaDu cell lines, as well as the ICGC normal control patient OSCC-GB FASTQ files containing paired-end reads, were rigorously checked for base quality, adapter contamination, and other quality parameters using FastQC version 0.11.8 (http://www.bioinformatics.babraham.ac.uk/projects/fastqc). We applied Trimmomatic v0.39 to filter out low-quality reads and trim adapter sequences (sliding window of 4 with Q > 20 and minimum length 36) (**Table S3**) [8]. The quality-checked FASTQ files were aligned to the human genome, utilizing Gencode GRCh38.p12 and its corresponding gene annotation file (GENCODE release 31) [9]. This genome alignment was carried out using the transcript-aware STAR (Spliced Transcripts Alignment to a Reference) version 2.7.1a aligner, involving genome indexing and subsequent read alignment (chr 1-22, XYM). The resulting BAM files were inspected for the number of uniquely mapped reads in the respective STAR log files [10]. Before somatic variant calling, the BAM files underwent preprocessing following the GATK best practices. The preprocessing steps included adding reading groups and sorting by coordinate  (AddOrReplaceReadGroups), marking duplicate reads (MarkDuplicates), splitting into intron and exon segments (SplitNCigarReads), and recalibrating base scores using dbSNP b151 build on the GRCh38 reference (BaseRecalibrator, ApplyBQSR) [7, 11].

#### S3 Creation of the panel of normals (PoN)

The Panel of Normals (PoN) was created using ICGC RNA-seq data from normal controls adjacent to the tumor of the Indian OSCC-GB patient samples (N=30; age 32-59; tumor stage III-IV). GATK preprocessed BAM files underwent analysis through the Mutect2 pipeline to identify the normal variants in at least 3/30 samples (CreateSomaticPanelOfNormals) [7, 12]. This RNA-seq-derived PoN filter accounts for variants originating from ‘normal contamination’ in the tumor samples and addresses specific RNA sequencing artifacts.

#### S4  Somatic variants identification

Somatic variants were identified using GATK-preprocessed BAM files with the Mutect2 pipeline (GATK v4.1.6.0) in single-sample/tumor-only mode, focusing on genome intervals (chr 1-22, XY) following GATK best practices (https://gatk.broadinstitute.org/hc/en-us/articles/360035531132). Selecting potential somatic variants involved filtering steps, including checking against PoN variants, gnomAD germline variants [13], addressing soft-clipped bases, and mitigating strand-orientation bias artifacts. Further refinement included stringent criteria, such as identifying SNP clusters within a 35-base pair window, ensuring a minimum of ten supporting reads for variants with at least two for alternate reads and with at least 1 read for each strand, a minimum allele frequency of 0.01, and a Phred-scaled quality score (GERMQ) of 40 or higher (FilterMutectCalls, VariantFiltration, SelectVariants). Further, the known RNA editing sites from the RADAR database were also filtered out [14]. Throughout this process, strict adherence to read quality parameters was maintained, ensuring a thorough, high-quality screening through the GATK pipeline [12].

#### S5 Variant annotation

Somatic variants were annotated using Ensembl Variant Effect Predictor (VEP v97) [15] with the human GRCh38.p12 gene assembly, considering a single Ensembl transcript per variant. Variants information was incorporated from the Single Nucleotide Polymorphism Database (dbSNP) build 151 [11] and the Catalogue of Somatic Mutations in Cancer variants (COSMIC v88) [16]. Predictions of missense mutation's functional impact, involving amino acid substitutions, were generated through Polyphen2 v2.2.2 [17]. Following annotation, the VCF files underwent a rigorous filtering procedure, excluding variants with minor allele frequencies below 0.05 (gnomAD r2.1 AF<0.05), removing dbSNP variants using BCFtools v1.9, and filtering out non-coding variants (./filter_vep). For coding indels, the SIFT Indel algorithm (https://sift.bii.a-star.edu.sg/www/SIFT_indels2.html) was utilized to predict their functional impact, with a focus on prioritizing frameshift variants. This prioritization was based on factors such as amino acid conservation, DNA sequence conservation, indel location within the transcript, and indel proximity to the exon boundary [18].

#### S6 Mutation annotation analysis

The Variant Call Format (VCFv4.2) files containing coding sequence variants were converted to Mutation Annotation Format (MAF) using the vcf2maf v1.6.18 tool from https://github.com/mskcc/vcf2maf. The MAF files identified genes with mutations, including novel and COSMIC variants, in at least three or more IIOC019 Adh and IIOC019 Oro cohorts (N=6) were used for downstream analysis leveraging the Maftools (v2.0.16) Bioconductor v3.6 R package. Differential mutational analysis compared IIOC019 Oro and IIOC019 Adh cohorts (N=6) and at early passage (P8) and late passage (P24) for the subgroups of IIOC019, applying Fisher's exact test to all genes with Maftools-mafCompare.

**Table S1.** Details of reads generated in RNA-sequencing of FaDu (P7) and IIOC019 (Indian origin, stage IV, cT4aN2bM0, 56 years, female) with sample name/ID corresponding to Table S1 for each condition's Adh and Oro, IIOC019 were grown in early (P8) and late passage (P24) in three technical replicates each (N=3).

#### Supplementary information

| S.NO | Type of Library | Lane No | Sample Name | Million reads (Paired end-2X125bp) | | Total Number of paired end reads (Million) | % >= Q30 | Mean Quality Score |
|---|---|---|---|---|---|---|---|---|
| | | | | Read 1 | Read 2 | | | |
| 1 | mRNA | 1 | JS1 | 31.33 | 31.33 | 62.66 | 92.73 | 35.39 |
| 2 | | | JS2 | 30.53 | 30.53 | 61.06 | 92.30 | 35.30 |
| 3 | | | JS3 | 28.79 | 28.79 | 57.58 | 92.65 | 35.37 |
| 4 | | | JS4 | 34.34 | 34.34 | 68.67 | 93.20 | 35.48 |
| 5 | | | JS5 | 32.72 | 32.72 | 65.44 | 92.87 | 35.41 |
| 6 | | | JS6 | 29.52 | 29.52 | 59.05 | 92.43 | 35.34 |
| 7 | | | JS7 | 29.34 | 29.34 | 58.69 | 92.28 | 35.31 |
| 8 | | | JS8 | 27.90 | 27.90 | 55.81 | 92.18 | 35.30 |
| 9 | | | JS9 | 27.18 | 27.18 | 54.35 | 91.76 | 35.21 |
| 10 | mRNA | 2 | JS10 | 30.49 | 30.49 | 60.98 | 91.45 | 35.16 |
| 11 | | | JS11 | 31.00 | 31.00 | 61.99 | 91.46 | 35.16 |
| 12 | | | JS12 | 31.43 | 31.43 | 62.86 | 91.66 | 35.20 |
| 13 | | | JS13 | 30.05 | 30.05 | 60.10 | 91.82 | 35.21 |
| 14 | | | JS14 | 30.74 | 30.74 | 61.47 | 91.52 | 35.15 |
| 15 | | | JS15 | 30.26 | 30.26 | 60.52 | 91.82 | 35.21 |
| 16 | | | JS16 | 31.45 | 31.45 | 62.90 | 92.14 | 35.27 |
| 17 | | | JS17 | 31.85 | 31.85 | 63.70 | 91.92 | 35.23 |
| 18 | | | JS18 | 31.80 | 31.80 | 63.60 | 91.97 | 35.23 |

**Table S2.** The parameters used to trim the adapter sequences from FASTQ files using the Trimmomatic v0.39 package.

| | |
|---|---|
| trimmomatic PE | -threads 8<br>-baseout<br>ILLUMINACLIP:TruSeq3-PE-2.fa:2:30:10:2:keepBothReads<br>LEADING:3<br>TRAILING:3<br>SLIDINGWINDOW:4:20<br>MINLEN:36 |

**Table S3.** The parameters used to extract reads counts from BAM files using Subread v2.0.1 package.

| | |
|---|---|
| featureCounts | -T 8<br>-a gencode.v31.annotation.gtf<br>-g gene_id<br>-o readsCounts.txt |

**Table S4.** Sample characteristics of HNSCC patients used for the scRNA-seq analysis.

| Patients | Gender | Age | Origin | Tumour Grade |
|---|---|---|---|---|

**Supplementary information**

| | | | | |
|---|---|---|---|---|
| Kurten et al. [2] | | | | |
| HN01_CD45n<br>HN01_CD45p | M | 70-79 | Oral cavity | pT4N0M0 |
| HN05_CD45n<br>HN05_CD45p | F | 50-59 | Oral cavity | T3N3M0 |
| HN06_CD45n<br>HN06_CD45p | M | 30-39 | Oral cavity | T3N1M0 |
| HN09_CD45n<br>HN09_CD45p | F | 70-79 | Oral cavity | T3N2bM0 |
| Quah et al. [3] | | | | |
| HN237<br>(CD45+:CD45-:: 1:1) | NA* | L Mandibular alveolous; SCC involving bone | | pT4apN1 |
| HN257<br>(CD45+:CD45-:: 1:1) | | Rt Mandibular alveolus; SCC involving bone | | pT4aN3b |
| HN272<br>(CD45+:CD45-:: 1:1) | | L Buccal mucosa; SCC with Mandibular bony invasion | | pT4aN2b |
| HN251<br>(CD45+:CD45-:: 1:1) | | R Lateral tongue; SCC extends into the floor of the mouth, infiltrates into the right submandibular sublingual gland | | pT3N3b |

*males and females, aged 21–85 were mentioned in the source study

**Table S5.** The list of identified distinctly mutated somatic variants for IIOC019 Oro (N=6, including P8 and P24) as compared to IIOC019 Adh (N=6, including P8 and P24) cohort applying Fisher's exact test (p-value<0.05, highlighted in bold) on all genes using Maftools-mafCompare v2.6.05 R package.

| Hugo Gene Symbol (Gene name) | Chromosome | Change in amino acids (number of patients with recurrent mutations) | Oro | Adh | p-value |
|---|---|---|---|---|---|
| **ERBB receptor feedback inhibitor 1 (ERRFI1)** | **13** | **p.N135Kfs*13 (6)** | **6** | **0** | **0.002164502** |
| **Heterogeneous nuclear ribonucleoprotein H2 (HNRNPH2)** | **X** | **p.L381Ffs*14 (6)** | **6** | **0** | **0.002164502** |
| **Breast cancer anti-estrogen resistance 3 (BCAR3)** | **1** | **p.G57Rfs*9 (5)** | **5** | **0** | **0.015151515** |
| **Heterogeneous nuclear ribonucleoprotein A3 (HNRNPA3)** | **2** | **p.R167Efs*9 (5)** | **5** | **0** | **0.015151515** |
| **M-phase phosphoprotein 8 (MPHOSPH8)** | **13** | **p.A173Sfs*24 (5); p.L157Ifs*11 (1)** | **5** | **0** | **0.015151515** |

| | | | | | |
|---|---|---|---|---|---|
| **NK-tumor recognition protein (NKTR)** | **3** | **p.N670Kfs*5 (3); p.N841Kfs*4 (2); p.V993Sfs*14 (1)** | **5** | **0** | **0.015151515** |
| **RE1-silencing transcription factor (REST)** | **4** | **p.S578Kfs*10 (5)** | **5** | **0** | **0.015151515** |
| **SON DNA and RNA binding protein (SON)** | **21** | **p.E2187Rfs*4 (5); p.S799= (1)** | **5** | **0** | **0.015151515** |
| **Transcription elongation factor A like 9 (TCEAL9)** | **X** | **p.M7Nfs*8 (5)** | **5** | **0** | **0.015151515** |
| **Thyroid hormone receptor interactor 4 (TRIP4)** | **15** | **p.G61Rfs*15 (5)** | **5** | **0** | **0.015151515** |
| **YTH domain-containing family protein 3 (YTHDF3)** | **8** | **p.P131Sfs*3 (5)** | **5** | **0** | **0.015151515** |
| AC242842.3 | 1 | p.T3825Yfs*25 (4) | 4 | 0 | 0.060606061 |
| Aldehyde dehydrogenase 1 family member A3 (ALDH1A3) | 15 | p.I132Hfs*8 (2); p.N447Kfs*35 (2) | 4 | 0 | 0.060606061 |
| CDKN2A interacting protein (CDKN2AIP) | 4 | p.N441Qfs*2 (4) | 4 | 0 | 0.060606061 |
| Eukaryotic translation termination factor 1 (ETF1) | 5 | p.D433* (4) | 4 | 0 | 0.060606061 |
| Nucleoporin 160 (NUP160) | 11 | p.Q981* (4) | 4 | 0 | 0.060606061 |

| | | | | | |
|---|---|---|---|---|---|
| Replication factor C subunit 1 (RFC1) | 4 | p.E545Gfs*17 (4) | 4 | 0 | 0.060606061 |
| SFI1 centrin binding protein (SFI1) | 22 | p.A642V (1); p.E170D (1); p.H398Y (1); p.M614L (1); p.S765T (1) | 4 | 0 | 0.060606061 |
| **Cell division cycle 23 (CDC23)** | **5** | **p.L234Sfs*23 (6)** | **0** | **6** | **0.002164502** |
| **Centromere protein F (CENPF)** | **1** | **p.Q110Tfs*10 (5); p.T2854Nfs*6 (3)** | **0** | **6** | **0.002164502** |
| **Rac GTPase activating protein 1 (RACGAP1)** | **12** | **p.A627Cfs*18 (6)** | **0** | **6** | **0.002164502** |
| **Topoisomerase (DNA) II binding protein 1 (TOPBP1)** | **3** | **p.N1514Kfs*2 (6); p.Y156Ifs*31 (1)** | **0** | **6** | **0.002164502** |
| **Basonuclin 1 (BNC1)** | **15** | **p.E315Rfs*8 (5)** | **0** | **5** | **0.015151515** |
| **Deltex 3 like, E3 ubiquitin ligase (DTX3L)** | **3** | **p.S233Kfs*5 (5); p.K607Rfs*21 (1)** | **0** | **5** | **0.015151515** |
| **Lamin B1 (LMNB1)** | **5** | **p.Q183Tfs*5 (5); p.S158Kfs*30 (1)** | **0** | **5** | **0.015151515** |

| | | | | | |
|---|---|---|---|---|---|
| AC022966.1 | 17 | p.G922A (1); p.Q1062* (1); p.Q221* (1); p.T1065I (1) | 0 | 4 | 0.060606061 |
| AC026470.1 | 16 | p.P522Sfs*2 (3); p.E185Rfs*9 (1) | 0 | 4 | 0.060606061 |
| AC106886.5 | 16 | p.Y1959Lfs*2 (4) | 0 | 4 | 0.060606061 |
| Anillin (ANLN) | 7 | p.H477Tfs*27 (3); p.S67* (1) | 0 | 4 | 0.060606061 |
| Rho GTPase activating protein 29 (ARHGAP29) | 1 | p.E680Gfs*17 (2); p.R13Tfs*11 (1); p.T247I (1) | 0 | 4 | 0.060606061 |
| ATPase 13A3 (ATP13A3) | 3 | p.I1079Yfs*52 (4) | 0 | 4 | 0.060606061 |
| Cell division cycle associated 7 (CDCA7) | 2 | p.A168Sfs*13 (4) | 0 | 4 | 0.060606061 |
| CCAAT enhancer binding protein zeta (CEBPZ) | 2 | p.R470Sfs*10 (3); p.E171* (1) | 0 | 4 | 0.060606061 |

| | | | | | |
|---|---|---|---|---|---|
| Chromodomain helicase DNA binding protein 7 (CHD7) | 8 | p.E664Rfs*12 (3); p.G1982= (1); p.K643Efs*33 (1); p.S1524P (1); p.Y759= (1) | 0 | 4 | 0.060606061 |
| DEAD-box helicase 46 (DDX46) | 5 | p.F242Ifs*2 (4) | 0 | 4 | 0.060606061 |
| GTPase activating protein and VPS9 domains 1 (GAPVD1) | 9 | p.S186Ffs*2 (4); p.I327T (1) | 0 | 4 | 0.060606061 |
| Jumonji domain containing 1C (JMJD1C) | 10 | p.D560Rfs*3 (4); p.N1371Kfs*23 (1); p.S1963= (1) | 0 | 4 | 0.060606061 |
| Kinesin family member 20B (KIF20B) | 10 | p.N1132Kfs*5 (3); p.E1727Rfs*27 (2) | 0 | 4 | 0.060606061 |
| Lysine methyltransferase 2A (KMT2A) | 11 | p.L3237Tfs*5 (4); | 0 | 4 | 0.060606061 |
| Kinetochore scaffold 1 (KNL1) | 15 | p.R1788Kfs*12 (4); p.N1222Kfs*8 (1) | 0 | 4 | 0.060606061 |

| | | | | | |
|---|---|---|---|---|---|
| Maestro heat like repeat family member 1 (MROH1) | 8 | p.E860D (1); p.G555A (1); p.Q794* (1); p.V796L (1) | 0 | 4 | 0.060606061 |
| Non-SMC condensin I complex subunit G (NCAPG) | 4 | p.I662Nfs*7 (4); p.S1002Kfs*2 (1) | 0 | 4 | 0.060606061 |
| Phosphoribosyl pyrophosphate amidotransferase (PPAT) | 4 | p.F373Ifs*9 (4) | 0 | 4 | 0.060606061 |
| Ribosomal protein L26 like 1 (RPL26L1) | 5 | p.I118Nfs*3 (3); p.V55A (1) | 0 | 4 | 0.060606061 |
| Spliceosome associated factor 1 (SART1) | 11 | p.H38Tfs*11 (4) | 0 | 4 | 0.060606061 |
| Solute carrier family 30 member 5 (SLC30A5) | 5 | p.V281Cfs*32 (4) | 0 | 4 | 0.060606061 |
| Structural maintenance of chromosomes 3 (SMC3) | 10 | p.I189Nfs*3 (4) | 0 | 4 | 0.060606061 |
| Spindle apparatus coiled-coil protein 1 (SPDL1) | 5 | p.M89Nfs*20 (4) | 0 | 4 | 0.060606061 |
| Trimethylguanosine synthase 1 (TGS1) | 8 | p.L755Ffs*4 (4); p.K724N (1) | 0 | 4 | 0.060606061 |
| THAP domain containing 12 (THAP12) | 11 | p.P675Sfs*2 (4) | 0 | 4 | 0.060606061 |

| | | | | | |
|---|---|---|---|---|---|
| Tripartite motif containing 5 (TRIM5) | 11 | p.N390Kfs*2 (4) | 0 | 4 | 0.060606061 |
| Ubinuclein 1 (UBN1) | 16 | p.R262Kfs*6 (4) | 0 | 4 | 0.060606061 |
| URB2 ribosome biogenesis homolog (URB2) | 1 | p.A448Sfs*24 (4); p.S1391= (1) | 0 | 4 | 0.060606061 |
| Ubiquitin specific peptidase 11 (USP11) | X | p.C784= (1); p.D650N (1); p.F831L (1); p.K279I (1); p.S391F (1) | 0 | 4 | 0.060606061 |
| Zinc finger DHHC-type palmitoyltransferase 17 (ZDHHC17) | 12 | p.G157A (1); p.Q267L (1); p.S57T (1); p.V233= (1); p.W315* (1) | 0 | 4 | 0.060606061 |
| Zinc finger protein 318 (ZNF318) | 6 | p.A1217Gfs*11 (3); p.N708Kfs*9 (1) | 0 | 4 | 0.060606061 |

Note the genes mutated in IIOC019 Oro and IIOC019 Adh cohorts were highlighted in red and blue, respectively.

The different scientific notation for representing types in mutation; for example, a frameshift insertion (p.E2187Rfs*4 or p.E2187RfsTer4), a missense mutation (p.A642V),a silent mutation (p.S799= or p.S799S), and a nonsense mutation (p.D433* or p.D433Ter).

**Table S6.** The functional impact prediction for the frameshift insertion somatic variants/mutations in the identified distinctly mutated DNA damage-related response genes for IIOC019 Oro (N=6, including P8 and P24) as compared to IIOC019 Adh (N=6, including P8 and P24) cohort using SIFT Indel algorithm (https://sift.bii.a-star.edu.sg/www/SIFT_indels2.html).

| HGNC | Chromosome | Position | Strand | Allele | Mutation | Effect | Confidence Score | Indel Location | Causes NMD |
|---|---|---|---|---|---|---|---|---|---|
| MPHOSPH8 | 13 | 19646583 | 1 | A | p.A173Sfs*24 (5) | damaging | 0.858 | 20% | YES |
| SON | 21 | 33559670 | 1 | A | p.E2187Rfs*4 (5) | damaging | 0.858 | 95% | YES |
| REST | 4 | 56930583 | 1 | A | p.S578Kfs*10 (5) | damaging | 0.858 | 52% | NO |
| HNRNPH2 | X | 101413123 | 1 | T | p.L381Ffs*14 (6) | damaging | 0.858 | 53% | N/A |
| ERRFI1 | 1 | 8014194 | 1 | T | p.N135Kfs*13 (6) | damaging | 0.858 | 29% | NO |
| BCAR3 | 1 | 93674762 | 1 | T | p.G57Rfs*9 (5) | damaging | 0.579 | 7% | YES |
| HNRNPA3 | 2 | 177216126 | 1 | A | p.R167Efs*9 (5) | damaging | 0.858 | 43% | YES |
| YTHDF3 | 8 | 63186386 | 1 | T | p.P131Sfs*3 (5) | damaging | 0.858 | 21% | YES |
| TRIP4 | 15 | 64394018 | 1 | A | p.G61Rfs*15 (5) | damaging | 0.858 | 10% | YES |
| TCEAL9 | X | 103357697 | 1 | A | p.M7Nfs*8 (5) | damaging | 0.579 | 2% | N/A |

| | | | | | | | | | |
|---|---|---|---|---|---|---|---|---|---|
| BNC1 | 15 | 83264308 | 1 | T | p.E315Rfs*8 (5) | damaging | 0.858 | 32% | YES |
| TOPBP1 | 3 | 133601277 | 1 | T | p.N1514Kfs*2 (6) | neutral | 0.961 | 99% | NO |
| DTX3L | 3 | 122568780 | 1 | A | p.S233Kfs*5 (5) | damaging | 0.858 | 31% | YES |
| LMNB1 | 5 | 126805594 | 1 | A | p.Q183Tfs*5 (5) | damaging | 0.858 | 31% | YES |
| CENPF | 1 | 214614990 | 1 | A | p.Q110Tfs*10 (5) | damaging | 0.579 | 3% | YES |
| CDC23 | 5 | 138198737 | 1 | A | p.L234Sfs*23 (6) | damaging | 0.858 | 39% | YES |
| RACGAP1 | 12 | 49990288 | 1 | A | p.A627Cfs*18 (6) | neutral | 0.961 | 99% | NO |

Note the genes mutated in IIOC019 Oro and IIOC019 Adh cohorts were highlighted in red and blue, respectively.

**Table S7.** The list of identified distinctly mutated somatic variants for early passage (P8) IIOC019 Oro as compared to IIOC019 Adh cohorts (N=3) by applying Fisher's exact test on all genes using Maftools-mafCompare v2.6.05 R package.

| Hugo Gene Symbol (Gene name) | Chromosome | Change in amino acids (number of patients with recurrent mutations) | Oro (P8) | Adh (P8) | p-value |
|---|---|---|---|---|---|
| BBX high mobility group box domain containing (BBX) | 3 | p.G62C (1); p.K453Efs*13 ( 1 ); p.V531Sfs*27 (1) | 3 | 0 | 0.1 |
| BCAR3 adaptor protein (BCAR3) | 1 | p.G57Rfs*9 (3) | 3 | 0 | 0.1 |
| Basic leucine zipper and W2 domains 1 (BZW1) | 2 | p.E342Rfs*29 (3) | 3 | 0 | 0.1 |
| Coiled-coil domain containing 14 (CCDC14) | 3 | p.F528Ifs*2 (3); p.E346K (1) | 3 | 0 | 0.1 |
| Chromodomain helicase DNA binding protein 1 (CHD1) | 5 | p.L534Ffs*31 (2); p.S1569Kfs*2 (1); p.N310Kfs*12 (1); p.A76P ( 1 ) | 3 | 0 | 0.1 |
| Chromosome segregation 1 like (CSE1L) | 20 | p.E398* (3) | 3 | 0 | 0.1 |

| | | | | | |
|---|---|---|---|---|---|
| DLG associated protein 5 (DLGAP5) | 14 | p.E743Rfs*25 (3) | 3 | 0 | 0.1 |
| Deoxynucleotidyltransferase terminal interacting protein 2 (DNTTIP2) | 1 | p.T160Nfs*8 (3) | 3 | 0 | 0.1 |
| ERBB receptor feedback inhibitor 1 (ERRFI1) | 1 | p.N135Kfs*13 (3) | 3 | 0 | 0.1 |
| Eukaryotic translation termination factor 1 (ETF1) | 5 | p.D433* (3) | 3 | 0 | 0.1 |
| General transcription factor IIIC subunit 2 (GTF3C2) | 2 | p.L250Sfs*4 (3); p.A149= (1) | 3 | 0 | 0.1 |
| Heterogeneous nuclear ribonucleoprotein H2 (HNRNPH2) | X | p.L381Ffs*14 (3) | 3 | 0 | 0.1 |
| Heat shock protein 90 alpha family class A member 1 (HSP90AA1) | 3 | p.C542Mfs*7 (3) | 3 | 0 | 0.1 |
| M-phase phosphoprotein 8 (MPHOSPH8) | 13 | p.A173Sfs*24 (3); p.L157Ifs*11 (1) | 3 | 0 | 0.1 |

| | | | | | |
|---|---|---|---|---|---|
| Natural killer cell triggering receptor (NKTR) | 3 | p.N670Kfs*5 (2); p.N841Kfs*4 (1) | 3 | 0 | 0.1 |
| NAD(P)H quinone dehydrogenase 1 (NQO1) | 16 | p.E242Rfs*5 (3) | 3 | 0 | 0.1 |
| Prolyl 4-hydroxylase subunit alpha 2 (P4HA2) | 5 | p.T261Nfs*9 (3) | 3 | 0 | 0.1 |
| Pleckstrin homology domain interacting protein (PHIP) | 6 | p.L590Ffs*3 (2); p.K177R (1); p.T936I (1); p.L125= (1) | 3 | 0 | 0.1 |
| Protein phosphatase 4 regulatory subunit 1 (PPP4R1) | 18 | p.P427Tfs*3 (3) | 3 | 0 | 0.1 |
| Protein tyrosine phosphatase non-receptor type 13 (PTPN13) | 4 | p.S1620R (3); p.L1096= (1) | 3 | 0 | 0.1 |
| REST corepressor 1 (RCOR1) | 14 | p.V425Cfs*11 (3) | 3 | 0 | 0.1 |
| Retroelement silencing factor 1 (RESF1) | 12 | p.S1005Kfs*4 (3); p.L1341Ffs*4 (2) | 3 | 0 | 0.1 |

| | | | | | |
|---|---|---|---|---|---|
| RE1 silencing transcription factor (REST) | 4 | p.S578Kfs*10 (3) | 3 | 0 | 0.1 |
| SH3 domain containing kinase binding protein 1 (SH3KBP1) | X | p.P403Afs*5 (3) | 3 | 0 | 0.1 |
| SON DNA and RNA binding protein (SON) | 21 | p.E2187Rfs*4 (3) | 3 | 0 | 0.1 |
| THO complex subunit 2 (THOC2) | X | p.T1385Nfs*20 (2); p.E959G (1); p.G1206= (1) | 3 | 0 | 0.1 |
| Thyroid hormone receptor interactor 4 (TRIP4) | 15 | p.G61Rfs*15 (3) | 3 | 0 | 0.1 |
| WW domain binding protein 4 (WBP4) | 13 | p.S271Kfs*12 (3) | 3 | 0 | 0.1 |
| Zinc finger protein 354A (ZNF354A) | 5 | p.I319Nfs*4 (3); p.T222Nfs*4 (1) | 3 | 0 | 0.1 |
| AC022966.1 | 17 | p.Q1062* (1); p.Q221* (1); p.T1065I (1) | 0 | 3 | 0.1 |
| AC026470.1 | 16 | p.P522Sfs*2 (3) | 0 | 3 | 0.1 |

| | | | | | |
|---|---|---|---|---|---|
| ATPase 13A3 (ATP13A3) | 3 | p.I1079Yfs*52 (3) | 0 | 3 | 0.1 |
| Bromodomain adjacent to zinc finger domain 1B (BAZ1B) | 7 | p.E823Rfs*6 (3); p.E739* (1) | 0 | 3 | 0.1 |
| Baculoviral IAP repeat containing 2 (BIRC2) | 11 | p.L199Ffs*19 (3); p.M39Nfs*4 (2) | 0 | 3 | 0.1 |
| Biorientation of chromosomes in cell division 1 like 1 (BOD1L1) | 4 | p.T836Nfs*2 (3); p.S352Efs*2 (1) | 0 | 3 | 0.1 |
| BRCA1 interacting helicase 1 (BRIP1) | 17 | p.T733Nfs*4 (2); p.Q582* (1); p.E472V (1) | 0 | 3 | 0.1 |
| Caspase recruitment domain family member 11 (CARD11) | 7 | p.E54K (3) | 0 | 3 | 0.1 |
| Cell division cycle and apoptosis regulator 1 (CCAR1) | 10 | p.D396* (3) | 0 | 3 | 0.1 |
| Cell division cycle 23 (CDC23) | 5 | p.L234Sfs*23 (3) | 0 | 3 | 0.1 |
| Cell division cycle associated 2 (CDCA2) | 8 | p.N696Kfs*2 (3) | 0 | 3 | 0.1 |

| | | | | | |
|---|---|---|---|---|---|
| Centromere protein F (CENPF) | 1 | p.Q110Tfs*10 (3) | 0 | 3 | 0.1 |
| Centrosomal protein 55 (CEP55) | 10 | p.D247Rfs*3 (3); p.P223Afs*2 (1) | 0 | 3 | 0.1 |
| DEAD box protein 58 (DDX58) | 9 | p.A297Cfs*8 (2); p.I97Nfs*2 (2); p.M760Nfs*4 (2) | 0 | 3 | 0.1 |
| Eukaryotic translation initiation factor 2 subunit gamma (EIF2S3) | X | p.I441Nfs*4 (3) | 0 | 3 | 0.1 |
| Eukaryotic translation initiation factor 3 subunit I (EIF3I) | 1 | p.D126* (3) | 0 | 3 | 0.1 |
| Follistatin (FST) | 5 | p.C256Mfs*16 (3) | 0 | 3 | 0.1 |
| Jumonji domain containing 1C (JMJD1C) | 10 | p.D560Rfs*3 (3); p.N1371Kfs*23 (1) | 0 | 3 | 0.1 |
| Laminin subunit alpha 3 (LAMA3) | 18 | p.L3055Ifs*11 (3); p.T2580Nfs*8 (1) | 0 | 3 | 0.1 |
| La ribonucleoprotein 7 (LARP7) | 4 | p.K34Efs*6 (3) | 0 | 3 | 0.1 |
| Lamin B1 (LMNB1) | 5 | p.Q183Tfs*5 (3) | 0 | 3 | 0.1 |

| | | | | | |
|---|---|---|---|---|---|
| Minichromosome maintenance complex component 4 (MCM4) | 8 | p.S442Ffs*9 (3) | 0 | 3 | 0.1 |
| Rac GTPase activating protein 1 (RACGAP1) | 12 | p.A627Cfs*18 (3) | 0 | 3 | 0.1 |
| Splicing regulatory glutamic acid and lysine rich protein 1 (SREK1) | 5 | p.R471Kfs*5 (3); p.S477Ifs*16 (2); p.S40Ffs*29 (1) | 0 | 3 | 0.1 |
| DNA topoisomerase II binding protein 1 (TOPBP1) | 3 | p.N1514Kfs*2 (3); p.Y156Ifs*31 (1) | 0 | 3 | 0.1 |
| TTK protein kinase (TTK) | 6 | p.*858Mfs*7 (2)<br>p.T806S (1) | 0 | 3 | 0.1 |
| Vascular endothelial zinc finger 1 (VEZF1) | 17 | p.R184* (3) | 0 | 3 | 0.1 |
| VPS29 retromer complex component (VPS29) | 12 | p.L57Tfs*23 (3) | 0 | 3 | 0.1 |
| Zinc finger CCHC-type containing 9 (ZCCHC9) | 5 | p.E66Rfs*5 (3) | 0 | 3 | 0.1 |

267

Note the genes mutated in IIOC019 Oro and IIOC019 Adh cohorts were highlighted in red and blue, respectively.

**Table S8.** The list of identified distinctly mutated somatic variants for late passage (P24) IIOC019 Oro as compared to IIOC019 Adh cohorts (N=3) by applying Fisher's exact test on all genes using Maftools-mafCompare v2.6.05 R package.

| Hugo Gene Symbol (Gene name) | Chromosome | Change in amino acids (number of patients with recurrent mutations) | Oro (P24) | Adh (P24) | p-value |
|---|---|---|---|---|---|
| Aldehyde dehydrogenase 1 family member A3 (ALDH1A3) | 15 | p.N447Kfs*35 (2); p.I132Hfs*8 (1) | 3 | 0 | 0.1 |
| AT-rich interaction domain 4B (ARID4B) | 1 | p.T1001Nfs*6 (3) | 3 | 0 | 0.1 |
| Catenin alpha 1 (CTNNA1) | 5 | p.K488* (3) | 3 | 0 | 0.1 |
| ERBB receptor feedback inhibitor 1 (ERRFI1) | 1 | p.N135Kfs*13 (3) | 3 | 0 | 0.1 |
| Heterogeneous nuclear ribonucleoprotein A3 (HNRNPA3) | 2 | p.R167Efs*9 (3) | 3 | 0 | 0.1 |
| Heterogeneous nuclear ribonucleoprotein H2 (HNRNPH2) | X | p.L381Ffs*14 (3) | 3 | 0 | 0.1 |

| | | | | | |
|---|---|---|---|---|---|
| La ribonucleoprotein 7 (LARP7) | 4 | p.K34Efs*6 (3); p.E29Rfs*3 (1) | 3 | 0 | 0.1 |
| OCRL inositol polyphosphate-5-phosphatase (OCRL) | X | p.E900K (3) | 3 | 0 | 0.1 |
| O-GlcNAcase (OGA) | 10 | p.V851Sfs*3 (3) | 3 | 0 | 0.1 |
| PNN interacting serine and arginine rich protein (PNISR) | 6 | p.E498Rfs*3 (3); p.H751Tfs*5 (1); p.A270Gfs*6 (1) | 3 | 0 | 0.1 |
| Protein tyrosine phosphatase domain containing 1 (PTPDC1) | 9 | p.R797Kfs*9 (3) | 3 | 0 | 0.1 |
| Protein tyrosine phosphatase non-receptor type 2 (PTPN2) | 18 | p.Q411Sfs*9 (3) | 3 | 0 | 0.1 |
| REST corepressor 1 (RCOR1) | 14 | p.V425Cfs*11 (3) | 3 | 0 | 0.1 |
| Retroelement silencing factor 1 (RESF1) | 12 | p.S1005Kfs*4 (3); p.L1341Ffs*4 (3) | 3 | 0 | 0.1 |
| SFI1 centrin binding protein (SFI1) | 22 | p.M614L (1); p.A642V (1); p.S765T (1); p.E170D (1) | 3 | 0 | 0.1 |

| | | | | | |
|---|---|---|---|---|---|
| SH3 domain containing kinase binding protein 1 (SH3KBP1) | X | p.P403Afs*5 (3) | 3 | 0 | 0.1 |
| Transcription elongation factor A like 9 (TCEAL9) | X | p.M7Nfs*8 (3) | 3 | 0 | 0.1 |
| VPS29 retromer complex component (VPS29) | 12 | p.L57Tfs*23 (3) | 3 | 0 | 0.1 |
| YTH N6-methyladenosine RNA binding protein F3 (YTHDF3) | 8 | p.P131Sfs*3 (3) | 3 | 0 | 0.1 |
| A-kinase anchoring protein 12 (AKAP12) | 6 | p.N1461Kfs*2 (3); p.F299Ifs*22 (1) | 0 | 3 | 0.1 |
| Ankyrin repeat domain containing 11 (ANKRD11) | 16 | p.E805Rfs*5 (3); p.H632Tfs*2 (2); p.S1279Kfs*4 (1) | 0 | 3 | 0.1 |
| Bromodomain adjacent to zinc finger domain 1A (BAZ1A) | 14 | p.F477Lfs*11 (2); p.E341Rfs*5 (1) | 0 | 3 | 0.1 |
| BMS1 ribosome biogenesis factor (BMS1) | 10 | p.H68Afs*7 (3) | 0 | 3 | 0.1 |

| | | | | | |
|---|---|---|---|---|---|
| Basonuclin zinc finger protein 1 (BNC1) | 15 | p.E315Rfs*8 (3) | 0 | 3 | 0.1 |
| Biorientation of chromosomes in cell division 1 like 1 (BOD1L1) | 4 | p.S352Efs*2 (2); p.E828Rfs*7 (1); p.S1108Efs*3 (1); p.E769= (1) | 0 | 3 | 0.1 |
| Caspase recruitment domain family member 11 (CARD11) | 7 | p.E54K (3) | 0 | 3 | 0.1 |
| Cell division cycle and apoptosis regulator 1 (CCAR1) | 10 | p.D396* (3) | 0 | 3 | 0.1 |
| Cell division cycle 23 (CDC23) | 5 | p.L234Sfs*23 (3) | 0 | 3 | 0.1 |
| Cell division cycle associated 2 (CDCA2) | 8 | p.N696Kfs*2 (3) | 0 | 3 | 0.1 |
| Centromere protein F (CENPF) | 1 | p.T2854Nfs*6 (3); p.Q110Tfs*10 (2) | 0 | 3 | 0.1 |
| Chromodomain helicase DNA binding protein 1 (CHD1) | 5 | p.L534Ffs*31 (2); p.N310Kfs*12 (1) | 0 | 3 | 0.1 |
| Chromosome segregation 1 like (CSE1L) | 20 | p.E398* (3); p.I825Nfs*5 (1) | 0 | 3 | 0.1 |

| | | | | | |
|---|---|---|---|---|---|
| DLG associated protein 5 (DLGAP5) | 14 | p.L337Ffs*15 (3); p.E743Rfs*25 (1) | 0 | 3 | 0.1 |
| Deltex E3 ubiquitin ligase 3L (DTX3L) | 3 | p.S233Kfs*5 (3) | 0 | 3 | 0.1 |
| Eukaryotic translation initiation factor 2 subunit gamma (EIF2S3) | X | p.I441Nfs*4 (3) | 0 | 3 | 0.1 |
| Eukaryotic translation initiation factor 3 subunit I (EIF3I) | 1 | p.D126* (3) | 0 | 3 | 0.1 |
| FAM111 trypsin like peptidase A (FAM111A) | 11 | p.Q481Afs*3 (3); p.N340Kfs*6 (2); p.V427Cfs*2 (1) | 0 | 3 | 0.1 |
| Follistatin (FST) | 5 | p.C256Mfs*16 (3) | 0 | 3 | 0.1 |
| Heterogeneous nuclear ribonucleoprotein D (HNRNPD) | 4 | p.I184Nfs*12 (3) | 0 | 3 | 0.1 |
| Lysine methyltransferase 2A (KMT2A) | 11 | p.L3237Tfs*5 (3) | 0 | 3 | 0.1 |

| | | | | | |
|---|---|---|---|---|---|
| Minichromosome maintenance complex component 4 (MCM4) | 8 | p.S442Ffs*9 (3) | 0 | 3 | 0.1 |
| Methionyl aminopeptidase 2 (METAP2) | 12 | p.R39Kfs*19 (3) | 0 | 3 | 0.1 |
| Marker of proliferation Ki-67 (MKI67) | 10 | p.T1732Nfs*34 (3) | 0 | 3 | 0.1 |
| Maestro heat like repeat family member 1 (MROH1) | 8 | p.Q794* (1); p.G555A (1); p.E860D (1) | 0 | 3 | 0.1 |
| NIPBL cohesin loading factor (NIPBL) | 5 | p.P567Afs*2 (2); p.T36S (1); p.H2292Sfs*48 (1) | 0 | 3 | 0.1 |
| Nucleolar protein 8 (NOL8) | 9 | p.R430Kfs*17 (3) | 0 | 3 | 0.1 |
| DNA polymerase theta (POLQ) | 3 | p.I1168Nfs*3 (3); p.F361L (1) | 0 | 3 | 0.1 |
| Protein phosphatase 4 regulatory subunit 3A (PPP4R3A) | 14 | p.L566Ffs*3 (3) | 0 | 3 | 0.1 |
| Rac GTPase activating protein 1 (RACGAP1) | 12 | p.A627Cfs*18 (3) | 0 | 3 | 0.1 |

| | | | | | |
|---|---|---|---|---|---|
| Replication timing regulatory factor 1 (RIF1) | 2 | p.N1654Kfs*2 (2); p.A1765Gfs*16 (1); p.L963M (1) | 0 | 3 | 0.1 |
| Remodeling and spacing factor 1 (RSF1) | 11 | p.I307Dfs*7 (2); p.R387Tfs*4 (1) | 0 | 3 | 0.1 |
| Semaphorin 3A (SEMA3A) | 7 | p.K320* (1); p.Y369* (1); p.W469Y470delinsCD (1); p.P465= (1) | 0 | 3 | 0.1 |
| Spindle apparatus coiled-coil protein 1 (SPDL1) | 5 | p.M89Nfs*20 (3) | 0 | 3 | 0.1 |
| SURP and G-patch domain containing 2 (SUGP2) | 19 | p.E434* (3); p.S340Ffs*19 (3) | 0 | 3 | 0.1 |
| Transcription activation suppressor family member 2 (TASOR2) | 10 | p.D1860Rfs*8 (1); p.V27I (1); p.N1059Kfs*15 (1); | 0 | 3 | 0.1 |
| Terminal nucleotidyltransferase 4A (TENT4A) | 5 | p.E188* (3) | 0 | 3 | 0.1 |
| THAP domain containing 12 (THAP12) | 11 | p.P675Sfs*2 (3) | 0 | 3 | 0.1 |

| | | | | | |
|---|---|---|---|---|---|
| THO complex subunit 2 (THOC2) | X | p.T1385Nfs*20 (3); p.E927Rfs*14 (1); p.N1531Kfs*2 (1) | 0 | 3 | 0.1 |
| DNA topoisomerase II binding protein 1 (TOPBP1) | 3 | p.N1514Kfs*2 (3) | 0 | 3 | 0.1 |
| Tripartite motif containing 5 (TRIM5) | 11 | p.N390Kfs*2 (3) | 0 | 3 | 0.1 |
| UPF2 regulator of nonsense mediated mRNA decay (UPF2) | 10 | p.K96Efs*51 (2); p.E90Rfs*57 (1); p.D76Rfs*71 (1) | 0 | 3 | 0.1 |
| Vascular endothelial zinc finger 1 (VEZF1) | 17 | p.R184* (3) | 0 | 3 | 0.1 |
| Zinc finger CCHC-type containing 9 (ZCCHC9) | 5 | p.E66Rfs*5 (3) | 0 | 3 | 0.1 |

Note the genes mutated in IIOC019 Oro and IIOC019 Adh cohorts were highlighted in red and blue, respectively.

**Table S9.** The list of identified distinctly mutated somatic variants for passage FaDu Oro as compared to FaDu Adh cohorts (N=3) by applying Fisher's exact test on all genes using Maftools-mafCompare v2.6.05 R package.

| Hugo Gene Symbol (Gene name) | Chromosome | Change in amino acids (number of patients with recurrent mutations) | Oro | Adh | p-value |
|---|---|---|---|---|---|
| AT-hook containing transcription factor 1 (AHCTF1) | 1 | p.L1290Pfs*5 (2); p.L2057Tfs*6 (1); p.V1243= (1) | 3 | 0 | 0.1 |
| ALG10 alpha-1,2-glucosyltransferase (ALG10) | 12 | p.S296Ffs*15 (3) | 3 | 0 | 0.1 |
| Ankyrin repeat domain 12 (ANKRD12) | 18 | p.T396Nfs*14 (2); p.T1126Nfs*7 (1); p.I767Nfs*15 (1); p.K1844= (1) | 3 | 0 | 0.1 |
| AT-rich interaction domain 1B (ARID1B) | 6 | p.A363V (3); p.M1004I (1); p.T1167= (1) | 3 | 0 | 0.1 |
| UDP-GlcNAc:betaGal beta-1,3-N- | 3 | p.W32Lfs*6 (3) | 3 | 0 | 0.1 |

| | | | | | |
|---|---|---|---|---|---|
| acetylglucosaminyltransferase 5 (B3GNT5) | | | | | |
| Bromodomain adjacent to zinc finger domain 1B (BAZ1B) | 7 | p.E823Rfs*6 (3) | 3 | 0 | 0.1 |
| Baculoviral IAP repeat containing 2 (BIRC2) | 11 | p.M39Nfs*4 (3); p.L199Ffs*19 (2) | 3 | 0 | 0.1 |
| Calpain 7 (CAPN7) | 3 | p.D800Gfs*3 (3) | 3 | 0 | 0.1 |
| Coiled-coil domain containing 14 (CCDC14) | 3 | p.F528Ifs*2 (3) | 3 | 0 | 0.1 |
| Cyclin dependent kinase 1 (CDK1) | 10 | p.I35Nfs*6 (3) | 3 | 0 | 0.1 |
| Centrosomal protein 192 (CEP192) | 18 | p.F15L (2); p.P13S (1); p.V1937Sfs*6 (1); p.V1504L (1); p.E1623* (1); p.S1914Ffs*13 (1); p.E277= (1) | 3 | 0 | 0.1 |

| | | | | | |
|---|---|---|---|---|---|
| Cleavage stimulation factor subunit 2 tau variant (CSTF2T) | 10 | p.D454Y (3) | 3 | 0 | 0.1 |
| Kinesin family member 21A (KIF21A) | 12 | p.E1232K (1); p.A310V (1); p.D691H (1); p.T787I (1) | 3 | 0 | 0.1 |
| La ribonucleoprotein 7, transcriptional regulator (LARP7) | 4 | p.E29Rfs*3 (3) ; p.K34Efs*6 (2) | 3 | 0 | 0.1 |
| LPS responsive beige-like anchor protein (LRBA) | 4 | p.N851S (1); p.V2066E (1); p.C358S (1); p.F357I (1); p.A205= (1) | 3 | 0 | 0.1 |
| Mitogen-activated protein kinase kinase kinase 9 (MAP3K9) | 14 | p.I479Mfs*17 (3) | 3 | 0 | 0.1 |
| MutS homolog 6 (MSH6) | 2 | p.R248Tfs*8 (2); p.D135* (1) | 3 | 0 | 0.1 |
| Methylenetetrahydrofolate dehydrogenase (NADP+ dependent) 1 like (MTHFD1L) | 6 | p.G923Rfs*12 (3) | 3 | 0 | 0.1 |

| | | | | | |
|---|---|---|---|---|---|
| Peptidylprolyl isomerase G (PPIG) | 2 | p.N420Kfs*2 (3) | 3 | 0 | 0.1 |
| Protein tyrosine phosphatase non-receptor type 2 (PTPN2) | 18 | p.Q411Sfs*9 (3) | 3 | 0 | 0.1 |
| REX4 homolog, 3'-5' exonuclease (REXO4) | 9 | p.E109Rfs*48 (3) | 3 | 0 | 0.1 |
| SH3 domain containing kinase binding protein 1 (SH3KBP1) | X | p.P403Afs*5 (3) | 3 | 0 | 0.1 |
| Solute carrier family 25 member 36 (SLC25A36) | 3 | p.Q266Sfs*38 (3) | 3 | 0 | 0.1 |
| Solute carrier organic anion transporter family member 1B3 (SLCO1B3) | 12 | p.F651Ifs*12 (3) | 3 | 0 | 0.1 |
| Structural maintenance of chromosomes 3 (SMC3) | 10 | p.I189Nfs*3 (3) | 3 | 0 | 0.1 |

| | | | | | |
|---|---|---|---|---|---|
| Spectrin repeat containing nuclear envelope protein 2 (SYNE2) | 14 | p.T6593Nfs*2 (3); p.L4533Ifs*3 (1) | 3 | 0 | 0.1 |
| Ubiquitin specific peptidase 8 (USP8) | 15 | p.S452Ffs*9 (2); p.N476Kfs*35 (1) | 3 | 0 | 0.1 |
| Vacuolar protein sorting 13 homolog C (VPS13C) | 15 | p.V3437L (1); p.G987A (1); p.F3356V (1); p.N3101K (1); p.R3198T (1); p.A2187V (1) | 3 | 0 | 0.1 |
| Xanthine dehydrogenase (XDH) | 2 | p.D652H (3) | 3 | 0 | 0.1 |
| YEATS domain containing 4 (YEATS4) | 12 | p.T190Nfs*4 (3) | 3 | 0 | 0.1 |
| Zinc finger and BTB domain containing 1 (ZBTB1) | 14 | p.Q525Afs*4 (3) | 3 | 0 | 0.1 |
| Zinc finger CCHC-type containing 9 (ZCCHC9) | 5 | p.E66Rfs*5 (3) | 3 | 0 | 0.1 |

| | | | | | |
|---|---|---|---|---|---|
| Zinc finger protein 292 (ZNF292) | 6 | p.E2088Rfs*16 (3) | 3 | 0 | 0.1 |
| Zinc finger protein 318 (ZNF318) | 6 | p.A1217Gfs*11 (3); p.N708Kfs*9 (1) | 3 | 0 | 0.1 |
| AT-rich interaction domain 4B (ARID4B) | 1 | p.T1001Nfs*6 (3) | 0 | 3 | 0.1 |
| ATRX chromatin remodeler (ATRX) | X | p.M1? (1); p.T1358Dfs*4 (1); p.V469L (1) | 0 | 3 | 0.1 |
| BLM RecQ like helicase (BLM) | 15 | p.K195* (3); p.Q1201Tfs*16 (1); p.K1147* (1) | 0 | 3 | 0.1 |
| Biorientation of chromosomes in cell division 1 like 1 (BOD1L1) | 4 | p.I1681F (1); p.S352Efs*2 (1); p.S1108Efs*3 (1); p.T1158Nfs*9 (1); p.E828Rfs*7 (1) | 0 | 3 | 0.1 |
| Cell division cycle associated 7 (CDCA7) | 2 | p.A168Sfs*13 (3) | 0 | 3 | 0.1 |

| | | | | | |
|---|---|---|---|---|---|
| Carbohydrate sulfotransferase 11 (CHST11) | 12 | p.Q291L (3) | 0 | 3 | 0.1 |
| ELAV like RNA binding protein 2 (ELAVL2) | 9 | p.T239Nfs*29 (3) | 0 | 3 | 0.1 |
| FAM111 trypsin like peptidase A (FAM111A) | 11 | p.Q481Afs*3 (3); p.N340Kfs*6 (1); p.G229= (1) | 0 | 3 | 0.1 |
| FA complementation group D2 (FANCD2) | 3 | p.R926Pfs*6 (3) | 0 | 3 | 0.1 |
| F-box protein 38 (FBXO38) | 5 | p.F168C (2); p.A478T (1) | 0 | 3 | 0.1 |
| G elongation factor mitochondrial 1 (GFM1) | 3 | p.I46Nfs*11 (2); p.I292Nfs*6 (1) | 0 | 3 | 0.1 |
| HAUS augmin like complex subunit 5 (HAUS5) | 19 | p.A486S (3) | 0 | 3 | 0.1 |
| HECT and RLD domain containing E3 ubiquitin protein ligase 2 (HERC2) | 15 | p.E1670D (3); p.H2741Tfs*12 (1) | 0 | 3 | 0.1 |

| | | | | | |
|---|---|---|---|---|---|
| HEXIM P-TEFb complex subunit 1 (HEXIM1) | 17 | p.H153Tfs*2 (3) | 0 | 3 | 0.1 |
| Heat shock protein 90 beta family member 1 (HSP90B1) | 12 | p.W485Lfs*14 (3) | 0 | 3 | 0.1 |
| Uncharacterized protein KIAA2026 (KIAA2026) | 9 | p.I1876Nfs*18 (3); p.I1099Nfs*10 (2) | 0 | 3 | 0.1 |
| LDL receptor related protein 1 (LRP1) | 12 | p.Q344E (1); p.Y2030F (1); p.C4210F (1); p.Q4203L (1); | 0 | 3 | 0.1 |
| Methylenetetrahydrofolate dehydrogenase, cyclohydrolase and formyltetrahydrofolate synthetase 1 (MTHFD1) | 14 | p.N682* (3); p.P247Tfs*9 (1) | 0 | 3 | 0.1 |
| Non-SMC condensin I complex subunit G (NCAPG) | 4 | p.I662Nfs*7 (3); p.S1002Kfs*2 (1) | 0 | 3 | 0.1 |
| Neuropilin and tolloid like 2 (NETO2) | 16 | p.A337Sfs*5 (2); p.T29Nfs*36 (1); p.V377Gfs*9 (1) | 0 | 3 | 0.1 |

| | | | | | |
|---|---|---|---|---|---|
| Nucleolar protein 8 (NOL8) | 9 | p.R257Kfs*4 (2); p.D1050* (1) | 0 | 3 | 0.1 |
| NRDE-2, necessary for RNA interference, domain containing (NRDE2) | 14 | p.L766Tfs*7 (3)<br>p.D1022* (2); p.N681Qfs*8 (1); p.R90Kfs*7 (1); p.D1164= (1) | 0 | 3 | 0.1 |
| Protein kinase, DNA-activated, catalytic subunit (PRKDC) | 8 | p.N2830Kfs*16 (3); p.N4037Kfs*12 (2); p.R2704Kfs*11 (1) | 0 | 3 | 0.1 |
| Retroelement silencing factor 1 (RESF1) | 12 | p.L1341Ffs*4 (2) ; p.S1005Kfs*4 (1) | 0 | 3 | 0.1 |
| Senataxin (SETX) | 9 | p.L459Sfs*9 (1) ; p.C292F (1) ; p.G1252Efs*4 (1) ; p.H291= (1) | 0 | 3 | 0.1 |
| SURP and G-patch domain containing 2 (SUGP2) | 19 | p.E434* (2); p.C431= (2); p.S340Ffs*19 (1) | 0 | 3 | 0.1 |

| | | | | | |
|---|---|---|---|---|---|
| SZT2 subunit of KICSTOR complex (SZT2) | 1 | p.G2459R (1); p.N3055Y (1); p.A33V (1); p.R25= (1) | 0 | 3 | 0.1 |
| Trio Rho guanine nucleotide exchange factor (TRIO) | 5 | p.D2646Rfs*2 (3) | 0 | 3 | 0.1 |
| Ubiquitin specific peptidase 34 (USP34) | 2 | p.L3265F (2); p.E3340* (1) | 0 | 3 | 0.1 |
| YTH N6-methyladenosine RNA binding protein F3 (YTHDF3) | 8 | p.P131Sfs*3 (3) | 0 | 3 | 0.1 |
| Zinc finger protein 609 (ZNF609) | 15 | p.K940N (3); p.P640Tfs*3 (2); p.D929N (2) | 0 | 3 | 0.1 |
| Zinc finger protein 622 (ZNF622) | 5 | p.Y4H (3) | 0 | 3 | 0.1 |

Note the genes mutated in FaDu Oro and FaDu Adh cohorts were highlighted in red and blue, respectively. Here, ATRX: p.M1? represents a Translation_Start_Site mutation.

**Table S10.** The list of identified statistically significant DDR KEGG pathways (p-value<0.05) for IIOC019 Oro (N=6, including P8 and P24) as compared to IIOC019 Adh (N=6, including P8 and P24) cohort using GAGE v2.40.2 R package.

| **Pathways** | **IIOC019 (P8+P24)** | **IIOC019 (P8)** | **IIOC019 (P24)** |
|---|---|---|---|
| hsa04110 Cell cycle | + | + | + |
| hsa03440 Homologous recombination | + | | + |
| hsa03410 Base excision repair | + | | + |
| hsa03430 Mismatch repair | + | | + |
| hsa03420 Nucleotide excision repair | + | | + |
| hsa04115 p53 signaling pathway | + | | + |

Note the down-regulated pathways were highlighted in green.

**Table S11.** The list of top 10 identified differentially and statistically significant Biological Process (BP) gene ontology (p-value<0.05) for IIOC019 Oro (N=6, including P8 and P24) as compared to IIOC019 Adh (N=6, including P8 and P24) cohort using GO.db v3.12.1 and GOstats v2.56.0 R package.

| GOBPID | Term | GOBPID | Term |
|---|---|---|---|
| GO:0006810 | transport | GO:0007049 | cell cycle |
| GO:0006629 | lipid metabolic process | GO:0000278 | mitotic cell cycle |
| GO:0002275 | myeloid cell activation involved in immune response | GO:0022402 | cell cycle process |

| | | | |
|---|---|---|---|
| GO:0031424 | keratinization | GO:1903047 | mitotic cell cycle process |
| GO:0045055 | regulated exocytosis | GO:0046483 | heterocycle metabolic process |
| GO:0043299 | leukocyte degranulation | GO:0006139 | nucleobase-containing compound metabolic process |
| GO:0006887 | exocytosis | GO:0006725 | cellular aromatic compound metabolic process |
| GO:0002444 | myeloid leukocyte mediated immunity | GO:0090304 | nucleic acid metabolic process |
| GO:0140352 | export from cell | GO:0006259 | DNA metabolic process |
| GO:0016192 | vesicle-mediated transport | GO:0051276 | chromosome organization |

Note the up-regulated GO and down-regulated GO were highlighted in red and green, respectively.

**Table S12.** The list of top 10 identified differentially and statistically significant Cellular Component (CC) gene ontology (p-value<0.05) for IIOC019 Oro (N=6, including P8 and P24) as compared to IIOC019 Adh (N=6, including P8 and P24) cohort using GO.db v3.12.1 and GOstats v2.56.0 R package.

| GOCCID | Term | GOCCID | Term |
|---|---|---|---|
| GO:0005576 | extracellular region | GO:0031981 | nuclear lumen |
| GO:0005615 | extracellular space | GO:0005654 | nucleoplasm |

| | | | |
|---|---|---|---|
| GO:0031982 | vesicle | GO:0005634 | nucleus |
| GO:0071944 | cell periphery | GO:0005694 | chromosome |
| GO:0005886 | plasma membrane | GO:0031974 | membrane-enclosed lumen |
| GO:0031410 | cytoplasmic vesicle | GO:0043233 | organelle lumen |
| GO:0097708 | intracellular vesicle | GO:0070013 | intracellular organelle lumen |
| GO:0031224 | intrinsic component of membrane | GO:0043228 | non-membrane-bounded organelle |
| GO:0099503 | secretory vesicle | GO:0043232 | intracellular non-membrane-bounded organelle |
| GO:0030141 | secretory granule | GO:0043231 | intracellular membrane-bounded organelle |

Note the up-regulated GO and down-regulated GO were highlighted in red and green, respectively.

**Table S13.** The list of top 10 identified differentially and statistically significant Molecular Function (MF) gene ontology (p-value<0.05) for IIOC019 Oro (N=6, including P8 and P24) as compared to IIOC019 Adh (N=6, including P8 and P24) cohort using GO.db v3.12.1 and GOstats v2.56.0 R package.

| GOMFID | Term | GOMFID | Term |
|---|---|---|---|
| GO:0005216 | ion channel activity | GO:0003676 | nucleic acid binding |

| | | | |
|---|---|---|---|
| GO:0030414 | peptidase inhibitor activity | GO:1901363 | heterocyclic compound binding |
| GO:0015267 | channel activity | GO:0097159 | organic cyclic compound binding |
| GO:0022803 | passive transmembrane transporter activity | GO:0003677 | DNA binding |
| GO:0004866 | endopeptidase inhibitor activity | GO:0003723 | RNA binding |
| GO:0004857 | enzyme inhibitor activity | GO:0005524 | ATP binding |
| GO:0061135 | endopeptidase regulator activity | GO:0030554 | adenyl nucleotide binding |
| GO:0022890 | inorganic cation transmembrane transporter activity | GO:0032559 | adenyl ribonucleotide binding |
| GO:0004867 | serine-type endopeptidase inhibitor activity | GO:0140097 | catalytic activity, acting on DNA |
| GO:0061134 | peptidase regulator activity | GO:0035639 | purine ribonucleoside triphosphate binding |

Note the up-regulated GO and down-regulated GO were highlighted in red and green, respectively.
